# EZH2 deletion does not impact acinar cell regeneration but restricts progression to pancreatic cancer in mice

**DOI:** 10.1101/2023.09.25.559339

**Authors:** Emilie Jaune-Pons, Xiaoyi Wang, Fatemeh Mousavi, Samad Elkaoutari, Kurt Berger, Charis Johnson, Mickenzie M. Martin, Saloni Aggarwal, Sukhman Brar, Khalid Muhammad, Joanna Ryan, Parisa Shooshtari, Angela J. Mathison, Nelson Dusetti, Raul Urrutia, Gwen Lomberk, Christopher L. Pin

## Abstract

Enhancer of Zeste Homologue 2 (EZH2) is part of the Polycomb Repressor Complex 2, which induces trimethylation of lysine 27 on histone 3 (H3K27me3) and promotes genes repression. EZH2 is overexpressed in many cancers including pancreatic ductal adenocarcinoma (PDAC). Previous studies in mice attributed both pro-oncogenic and tumor suppressive functions to EZH2. Deletion of the EZH2 enhances *de novo* KRAS-driven neoplasia following pancreatic injury by preventing acinar cell regeneration, while increased EZH2 expression in PDAC is correlated to poor prognosis, suggesting a context-dependant effect for EZH2 in PDAC progression. In this study, we examined EZH2 function in pre-and early neoplastic stages of PDAC. Using an inducible model to generate deletion of EZH2 only in adult acinar cells (EZH2^ΔSET^), we showed loss of EZH2 activity did not prevent acinar cell regeneration in the absence of oncogenic KRAS (KRAS^G12D^), nor lead to increased PanIN formation in the presence of KRAS^G12D^ in adult mice. However, loss of EZH2 did reduce recruitment of inflammatory cells and, when combined with a PDAC model, promoted widespread PDAC progression. Loss of EZH2 function also correlated to remodeling of the tumor microenvironment, which favors cancer cell progression. This study suggests expression of EZH2 in adult acinar cells restricts PDAC initiation and progression by affecting both the tumour microenvironment and acinar cell differentiation.

## Introduction

Pancreatic ductal adenocarcinoma (PDAC) is the most common form of pancreatic cancer with the worst five-year survival, ∼10%, of any of the major cancers (Pancreatic Cancer Facts, PANCAN). The principle driver mutation in PDAC is activating *KRAS* mutations, which occurs in >90 % of PDAC patients (1). Oncogenic KRAS (KRAS^G12D^) mutations appear at early stages of the disease but are not enough to induce PDAC on their own (2, 3). A multitude of studies indicate environmental stressors, in addition to somatic mutations in *KRAS,* are required for PDAC progression. Chronic inflammation is associated with increased sensitivity to KRAS^G12D^, indicating environmental factors contribute to progression (4). Based on these findings, there is increasing interest in the epigenomes of PDAC patients and the role for epigenetic mediators in initiation and progression of this cancer. Mutations in several genes encoding epigenetic modifiers, including *ARID1A* and *KMT2D* (5), are found in PDAC patients, and activation of KRAS^G12D^ is associated with extensive changes in the epigenetic profile of cells (6). In addition, Enhancer of Zeste Homolog 2 (EZH2) is highly expressed in a subset of PDAC tumours and is correlated to poor prognosis (7).

EZH2 is a histone-lysine N-methyltransferase enzyme and part of the Polycomb Repressive Complex 2 (PRC2), which plays a critical role in cell fate specification during embryonic development (8, 9). EZH2 induces trimethylation of H3K27 (H3K27me3), a histone modification linked to chromatin remodeling and gene repression (10). EZH2 is overexpressed in many cancers (7, 11), but both pro-oncogenic and tumor suppressive roles have be reported in the context of PDAC (12, 13). In the developing pancreas, EZH2 plays a repressive role for neoplastic lesion formation, as deletion of the SET domain, which is responsible for methyltransferase activity, reduces acinar cell regeneration after injury and increases pancreatic intraepithelial neoplasia (PanIN) initiation and tumor progression (12). Using a similar model, embryonic loss of EZH2 methyltransferase activity in mice expressing KRAS^G12D^ initially favoured PanIN progression, but reduced PanIN maintenance in aged mice compare to KRAS^G12D^ alone (14). This study proposed a role for EZH2 in NFATc1 regulation and PDAC progression, suggesting the effects of EZH2 extend to altering the tumour microenvironment (14). More recent studies showed *EZH2* deletion in pancreatic cancer cells increased GATA6 expression, a marker of classical PDAC subtype, indicating EZH2 promotes a more aggressive, basal-like PDAC subtype (13). Coupled with the findings that increased EZH2 expression correlated to more advanced disease and increased therapeutic resistance (15, 16), it appears EZH2’s role differs between early stages of PDAC initiation and later progression and resistance.

In this study, we examined EZH2 function in pre-neoplastic stages of PDAC, focusing on EZH2’s impact on acinar cell regeneration and PanIN initiation in adult mice. Since PDAC patients typically present later in life (>60 years of age), we developed a pre-clinical model which allows KRAS^G12D^ induction in adult acinar cells of the pancreas, instead of using embryonic induction of KRAS^G12D^ (12, 14). While we employed a similar approach to alter EZH2 function, in which the SET domain of EZH2 (EZH2^ΔSET^) is deleted, we used an inducible cre recombinase that promoted deletion only in acinar cells of the adult pancreas. Unlike previous studies, our results indicate loss of EZH2 activity has minimal impact on acinar cell regeneration and does not increase PanIN initiation, but initially favours more advanced PanIN lesion development in the context of KRAS^G12D^. Interestingly, loss of EZH2 SET activity in combination with KRAS^G12D^, induces reprogramming of the genome based on H3K27me3 enrichment and reduces immune cell recruitment in response to injury. Conversely, deleting EZH2^ΔSET^ in a susceptible mouse model for PDAC (*Mist1^creERT/-^KRAS^G12D^*) greatly enhanced PanIN progression and PDAC formation.

This study highlights several, context dependent roles for EZH2 in PDAC initiation and progression. EZH2 helps mediate *KRAS^G12D^*-induced reprogramming of the acinar cell genome, primes immune and inflammatory genes in these cells which allows for a differential immune response, and is required for long term expansion of pre-neoplastic lesions.

## Results

### KRAS^G12D^ promotes widespread epigenetic remodeling with no effect on acinar cell morphology

To understand the molecular and epigenetic responses to oncogenic KRAS, KRAS^G12D^ was induced specifically in acinar cells of 2-4-month-old *Mist1^creERT/+^KRAS^G12D^* mice by tamoxifen (TX) gavage (**Figure S1A**). Twenty-two days after initial TX treatment, mice were sacrificed, and the pancreata examined by histological and molecular means. H&E staining showed no differences in acinar cell morphology following KRAS^G12D^ activation (**Figure S1B**). RNA-seq analysis identified 380 differentially expressed genes (DEGs; 211 increased and 169 decreased; FDR ≤ 0.05) in response to KRAS^G12D^ compared to WT pancreata (**Figure S1C; Appendix 1**). Gene Ontology (GO) analysis using these 380 DEGs revealed 27 enriched pathways (p_adj_ ≤ 0.05) (**Figure S1D, Table S1**) including several relating to cell development and secretion. In general, the effects of KRAS^G12D^ on acinar cell morphology and gene expression were limited.

Conversely, ChIP-seq for histone modifications in the same tissue samples revealed widespread effects on H3K27me3 and H3K4me3 enrichment across the genome. ChIP-seq identified 2,527 H3K4me3 and 7,343 H3K27me3 unique enrichment sites (p_adj_ ≤ 0.05) in either WT or *Mist1^creERT/+^KRAS^G12D^* tissue (**Figure 1A**). The total number of H3K4me3-enriched regions was relatively the same between WT and *Mist1^creERT/+^KRAS^G12D^* tissue (21,334 *vs*. 21,137, respectively; ∼0.9% decrease). However, the number of sites enriched for H3K27me3 was ∼33.2% higher in *Mist1^creERT/+^KRAS^G12D^* pancreata (16,141 *vs*. 12,114 in WT tissue; **Figure 1A**). The distribution of H3K4me3 and H3K27me3 enrichment sites within the genome was similar between genotypes (**Figure 1B**) and heatmaps confirmed little change in H3K4me3 enrichment within 2 kb of transcriptional start sites (TSSs) (**Figure 1C**). Similar analysis for H3K27me3 identified uniquely enriched TSSs in both *Mist1^creERT/+^KRAS^G12D^*and wild type pancreata (boxed areas, **Figure 1C**). Comparing TSSs between genotypes supported a general increase in K27me3 enrichment in *Mist1^creERT/+^KRAS^G12D^*pancreata (**Figure 1D**) while H3K4me3-enriched TSSs, which were more numerous, were uniformly distributed between the two genotypes (**Figure 1D**). These results suggest KRAS^G12D^ increases widespread H3K27me3 enrichment in acinar cells.

**Figure 1:**
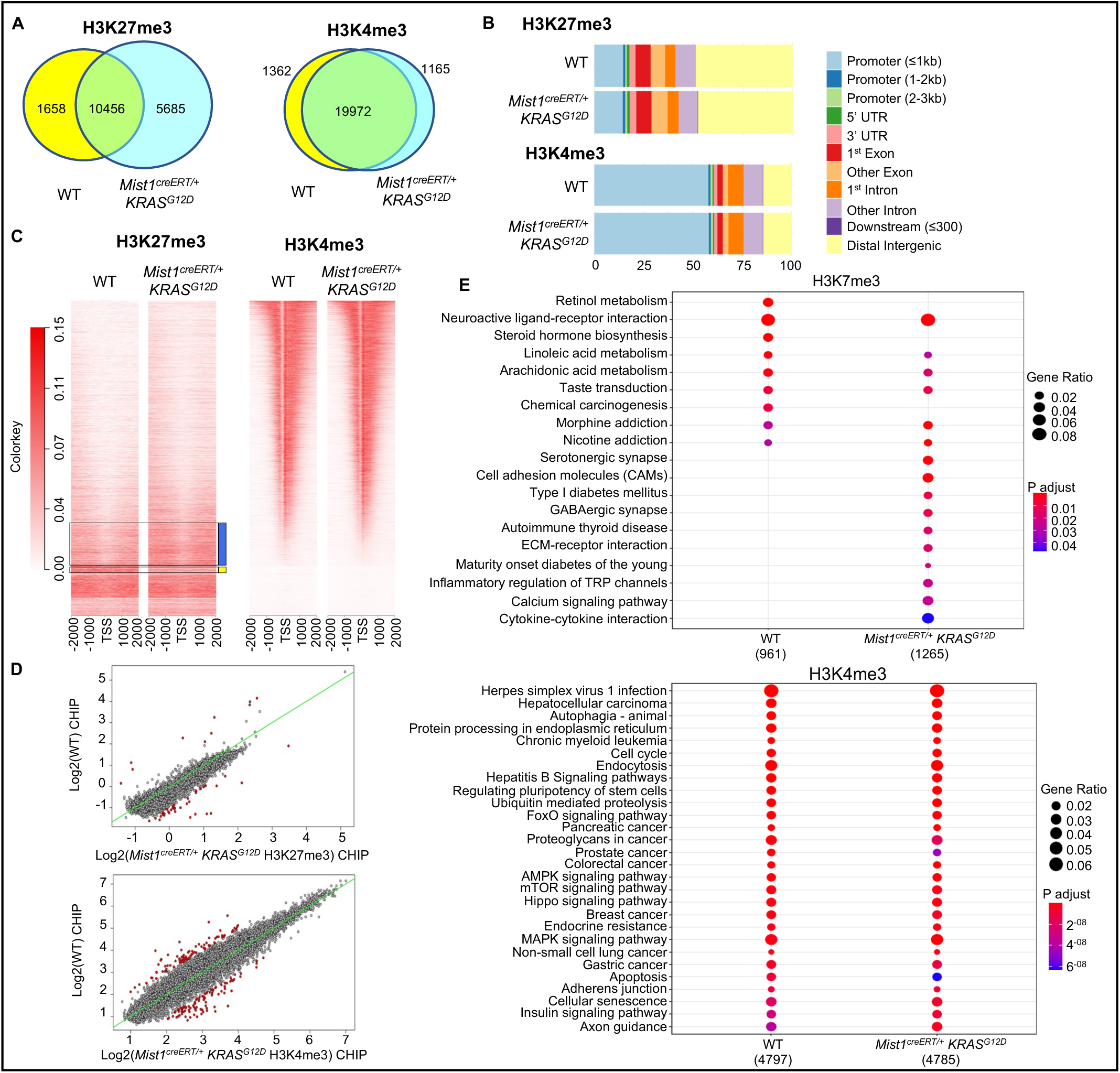
*KRAS^G12D^* promotes increased H3K27me3 enrichment in pancreatic acinar cells. **(A)** Venn diagram shows overlap of H3K4me3 and H3K27me3 enrichment sites based on ChIP-seq of pancreatic tissue from WT and *Mist1^creERT/+^KRAS^G12D^*mice 22 days after KRAS^G12D^ induction. **(B)** Genome distribution of H3K4me3 and H3K27me3 enrichment in WT and *Mist1^creERT/+^KRAS^G12D^*pancreatic tissue. **(C)** Heatmaps show H3K4me3 and H3K27me3 enrichment within 2 kb of transcriptional start sites (TSS) of all genes. **(D)** Comparison of called H3K4me3 and H3K27me3 enrichment at TSSs between WT and *Mist1^creERT/+^KRAS^G12D^*acinar cells. Red dots represent significantly dysregulated genes based on enrichment. Green line indicates expectation for equal enrichment between genotypes. **(E)** KEGG analysis identifying the pathways enriched for H3K4me3 (top 28 pathways) or H3K27me3 (all the pathways) in WT and *Mist1^creERT/+^KRAS^G12D^* pancreatic tissue (p adjusted ≤ 0.05).

To determine the molecular pathways associated with altered epigenetic enrichment patterns, genes were annotated in which the TSS occurred within +/-2 kB of a called enrichment peak. 1,991 genes were identified based on H3K27me3 enrichment in WT tissue *vs*. 2,684 in *Mist1^creERT/+^KRAS^G12D^*tissue, a 35% increase in H3K27me3-enriched genes (**Figure S1E**). 11,698 and 11,618 genes were identified based on H3K4me3 for WT and *Mist1^creERT/+^KRAS^G12D^* tissue, respectively. KEGG analysis of H3K4me3-annotated genes identified similar, enriched pathways between *Mist1^creERT/+^KRAS^G12D^* and WT data sets (**Table S2**; top 28 pathways are shown in **Figure 1E**). However, H3K27me3 enrichment identified more uniquely enriched pathways in *Mist1^creERT/+^KRAS^G12D^*compared to WT tissue, including pathways associated with ECM interaction, inflammation, and metabolism (**Table S2** and **Figure 1E**). This analysis suggests KRAS^G12D^ significantly increases H3K27me3 enrichment in acinar cells, which was not reflected at the level of gene transcription. This alteration in epigenetic marks without similar changes in expression underlies gene priming in which the activation of a gene by external cues is dictated by it’s the epigenetic program. Such priming had been identified in pancreatic development but has also been identified in adult tissue (17–19).

### EZH2 methyltransferase restricts KRAS^G12D^-mediated PanIN progression following injury

Since H3K27 trimethylation involves EZH2 (20, 21), we sought to determine the effects of deleting EZH2 methyltransferase activity in the context of pancreatic pathology. We used a similar model as previous studies, which targets the *Ezh2* gene by flanking exons 16 to 19, which encompass the SET domain, with loxP sites (12, 22). Unlike previous studies, we employed the *Mist1* cre-driver to promote inducible and acinar-specific *Ezh2* deletion and KRAS^G12D^ activation (**Figure S2A**).

To induce PanIN formation, we coupled activation of KRAS^G12D^ expression with acute injury using a two-day cerulein regimen 15 and 17 days after initial TX treatment (**Figure S2B;** (23)). PanIN progression was compared 35 days after initial cerulein treatment in *Mist1^creERT/+^KRAS^G12D^*and *Mist1^creERT/+^KRAS^LSL-G12D^Ezh2^ΔSET/ΔSET^* (referred to as *KRAS^G12D^Ezh2^ΔSET^*) mice (**Figure S2B**). C57Bl6 mice, or mice carrying only the *Mist1^creERT^* allele, were used as wild type controls since loss of a single *Mist1* allele has no effects on gene expression (both indicated as WT). We also included mice carrying the *Mist1^creERT^*allele and homozygous for the EZH2 ΔSET allele (*Mist1^creERT/+^Ezh2^ΔSET^*, called *EZH2^ΔSET^*). No group showed any overt differences based on final weights regardless of whether mice were treated with cerulein or saline (**Figure S2C**). Similarly, pancreas weight as a percentage of body weight showed no differences at the time of dissection (**Figure S2D**).

Histological analysis of WT and *EZH2^ΔSET^* pancreatic tissue showed no lesions or alterations in acinar cell regeneration (**Figure S2E**). This differed from previous studies showing an absence of EZH2 restricted acinar cell regeneration (12). Since the previous study used a longer, recurrent model of CIP, we next examined the response of *Ezh2^ΔSET^* mice to twice daily injections of 250 μg/kg cerulein over two weeks (**Figure S3A**; (24)). As previously reported, increased EZH2 accumulation was observed in response to injury in tissue from WT animals, with EZH2 completely ablated in *Ezh2^ΔSET^* tissue (**Figure S3B**). However, recurrent injury still showed no differences in body (**Figure S3C**) or pancreas to body weight (**Figure S3D**), pancreatic morphology (**Figure S3E**), or amylase accumulation (**Figure S3F**) between genotypes. Both WT and *EZH2^ΔSET^* pancreatic tissue showed increased CK19 accumulation following cerulein treatment based on IHC analysis (**Figure S3G**) but this was not different between genotypes. This suggests loss of EZH2^ΔSET^ in mature acinar cells does not restrict acinar cell regeneration as previously reported.

We next performed H&E histology to determine the impact of EZH2^ΔSET^ deletion on KRAS^G12D^-induced PanIN progression following acute injury. Saline-treated *Mist1^creERT/+^KRAS^G12D^* and *KRAS^G12D^Ezh2^ΔSET^* pancreatic tissue showed sporadic lesions (<1% of the entire tissue area). Cerulein treatment resulted in significantly more damaged regions (ADM and PanINs with surrounding stroma) in *Mist1^creERT/+^KRAS^G12D^* (29 ± 11.54%) and *KRAS^G12D^Ezh2^ΔSET^* (13.67 ± 3.175%) mice (**Figures 2A**, **S4A**), while there was no significant difference between *Mist1^creERT/+^KRAS^G12D^*and *KRAS^G12D^Ezh2^ΔSET^* tissue (**Figure 2A**). *Mist1^creERT/+^KRAS^G12D^*mice had significantly more pancreatic damage compared to WT and *Ezh2^ΔSET^* mice, while *KRAS^G12D^Ezh2^ΔSET^* showed a trend towards increased damage. To specifically identify ADM and PanIN lesions, we compared the ratio of CK19 to amylase accumulation by IHC, which identify duct/ADM/PanINs and acinar cells, respectively (**Figure 2B**). This analysis showed a trend towards increased CK19 accumulation in *KRAS^G12D^Ezh2^ΔSET^* mice, but the difference was not significant (p=0.219). To better characterize PanIN progression, we performed Alcian Blue (**Figures 2C, S4B**) and PAS (Periodic Acid Schiff) histology (**Figure 2D, S4C**), which identifies acidic mucins and mucin-polysaccharides, respectively. For both stains, *KRAS^G12D^Ezh2^ΔSET^* mice showed more extensive staining of PanINs compared to *Mist1^creERT/+^KRAS^G12D^*mice, suggesting EZH2 does not restrict initiation of KRAS^G12D^-mediated PanIN formation, but may limit progression to more advanced PanIN lesions.

**Figure 2:**
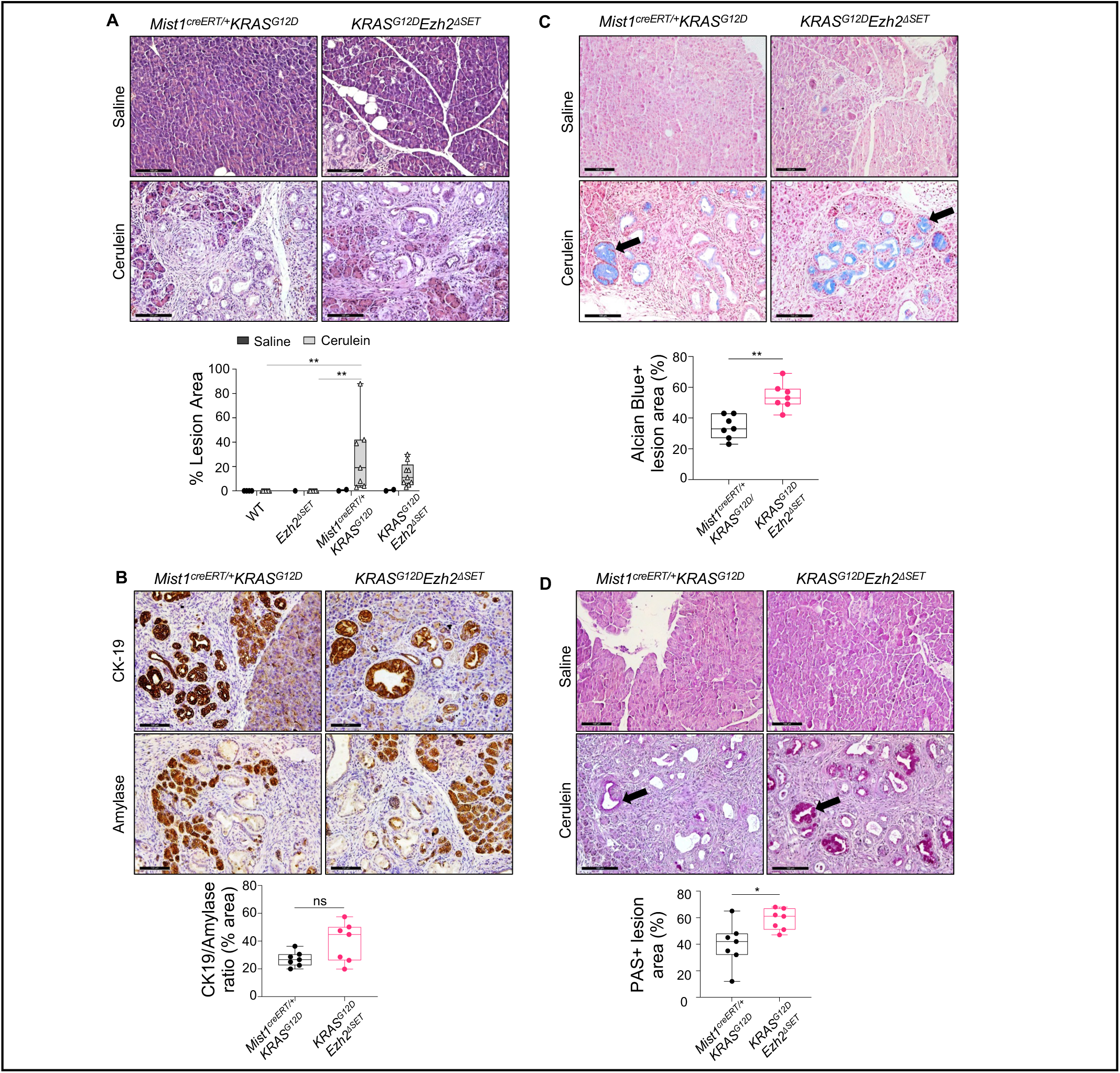
Loss of EZH2 methyltransferase activity increases *KRAS^G12D^*-mediated PanIN progression. Histological and quantitative analysis of *Mist1^creERT/+^KRAS^G12D^*and *KRAS^G12D^Ezh2^ΔSET^*mice 51 days after initial tamoxifen gavage to induce KRAS^G12D^ expression and 35 days after two days of treatment with vehicle (saline) or cerulein. **(A)** Representative H&E images of pancreatic tissue. Boxplots indicate the amount of lesion area determined as a percentage of the entire pancreatic tissue. Significance is measured by one-way ANOVA follow by Tukey’s post hoc tests. **(B)** Representative IHC for CK19 or amylase followed by counterstaining with hematoxylin. Boxplots compare the ratio of CK19^+^/amylase^+^ tissue. Significance is measured by two-tailed unpaired Mann-Whitney test. Representative images of **(C)** alcian blue histology or **(D**) Periodic acid-Schiff (PSA) histology showing advanced lesions (arrows) saline or cerulein-treated *Mist1^creERT/+^KRAS^G12D^*and *KRAS^G12D^Ezh2^ΔSET^*mice. Boxplots compare the amount of positively stained area as a percentage of ADM/PanIN lesions. Significance is measured by two-tailed unpaired Mann-Whitney test. In all cases, scale bar = 100 µm. For graphs, individual mice are shown, and data represents the mean ± min to max. p value*≤0.05, **≤0.01. ns=not-significant.

To identify the molecular mechanisms EZH2 regulates in the presence of KRAS^G12D^, H3K27me3 enrichment patterns were compared between *KRAS^G12D^Ezh2^ΔSET^*, *Mist1^creERT/+^KRAS^G12D^*and WT pancreatic tissue 22 days after KRAS^G12D^ activation. At this time point, the pancreatic tissue still retained normal histology in all genotypes (**Figures S2B, S5A**). Unlike *Mist1^creERT/+^KRAS^G12D^*tissue, the number of H3K27me3 enrichment sites was similar between WT (12,166) and *KRAS^G12D^Ezh2^ΔSET^* (12,647; ∼4% increase) tissues suggesting the absence of EZH2 reduces the ability of KRAS^G12D^ to reprogram the genome (**Figure 3A**). Analysis of H3K4me3 identified 21,463 (WT) *vs*. 21,938 (*KRAS^G12D^Ezh2^ΔSET^*) enrichment sites, a 2.3% increase, similar to that observed for *Mist1^creERT/+^KRAS^G12D^* tissue, while *Mist1^creERT/+^KRAS^G12D^* had 21.8% fewer H3K27me3-enriched sites than *KRAS^G12D^Ezh2^ΔSET^* tissue (16,458 vs 12,872, respectively; **Figure 3B**). Assessment of TSSs within 3 kB of H3K27me3 enrichment identified ∼19.5% fewer called genes in *KRAS^G12D^Ezh2^ΔSET^* tissue compared to *Mist1^creERT/+^KRAS^G12D^* (2,160 vs. 2,684 genes, respectively) (**Figure 3B**), but similar to the number of annotated genes in WT tissue (1,991) (**Figure 3A**). Comparing WT, *KRAS^G12D^Ezh2^ΔSET^* and *Mist1^creERT/+^KRAS^G12D^* annotated genes identified 611 uniquely enriched genes in *Mist1^creERT/+^KRAS^G12D^* tissue (**Figure 3C**, **Appendix 2)**. These genes, which are enriched only in the presence of EZH2, represent potential direct targets of EZH2 reprogramming in the context of KRAS^G12D^.

**Figure 3:**
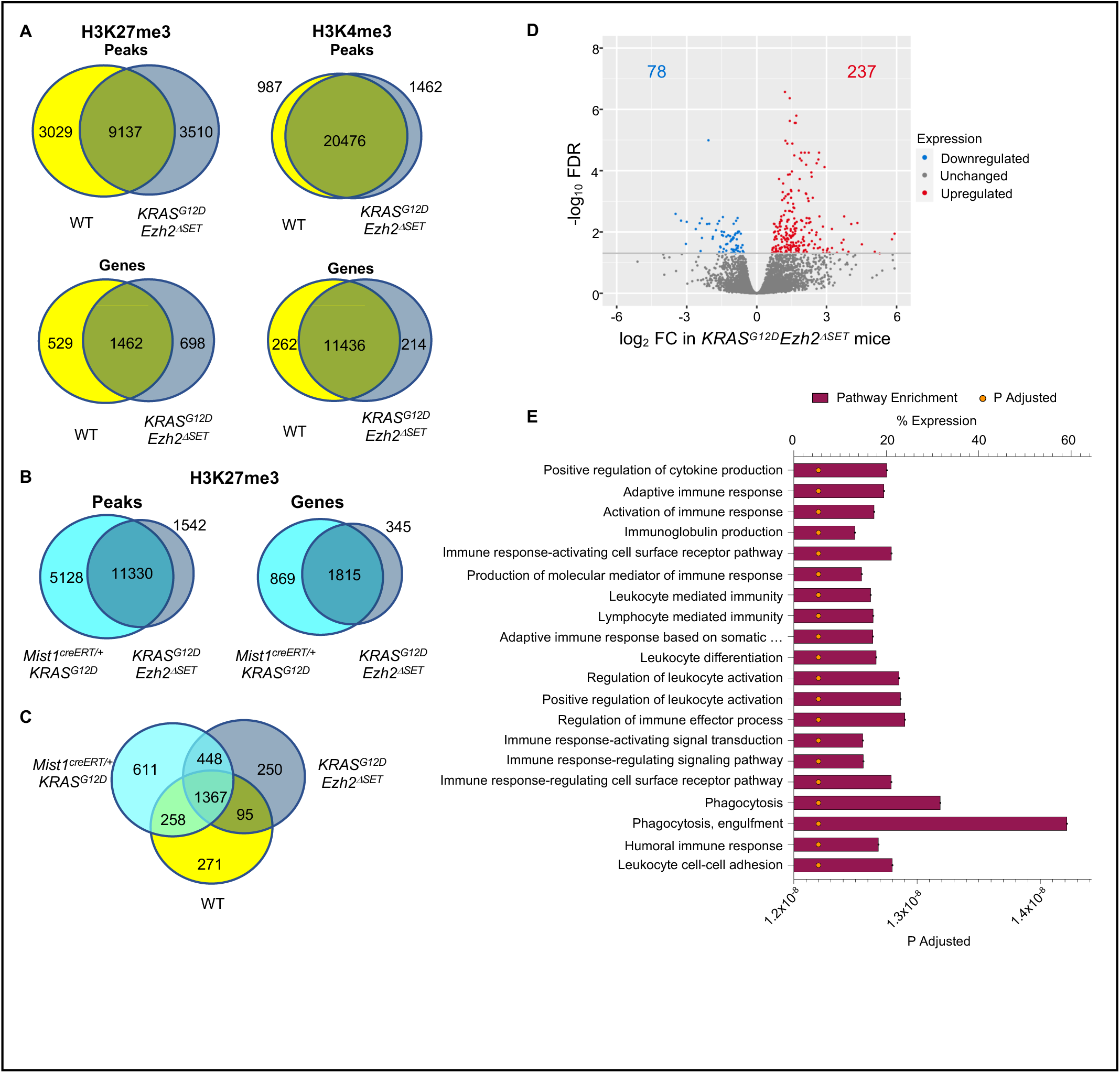
Loss of EZH2 methyltransferase activity alters the effects of *KRAS^G12^* on expression of genes linked to the tissue microenvironment. Venn diagrams comparing the number of enrichment sites or annotated genes within 3 kb of an enrichment site based on ChIP-Seq for H3K4me3 and H3K27me3 on pancreatic tissue chromatin from **(A)** wild type (WT) and *KRAS^G12D^Ezh2^ΔSET^*, (B) *Mist1^creERT/+^KRAS^G12D^*and *KRAS^G12D^Ezh2^ΔSET^*mice or **(C)** WT, *Mist1^creERT/+^KRAS^G12D^*, and *KRAS^G12D^Ezh2^ΔSET^*mice, 22 days after tamoxifen gavage. **(D)** Volcano plot showing genes with a log2 fold change (FC) between *KRAS^G12D^Ezh2^ΔSET^* and *Mist1^creERT/+^KRAS^G12D^*pancreatic tissue 22 days after activating KRAS^G12D^. Significantly lower expressed genes (78 genes) are shown in blue while significantly higher expressed genes (237 genes) are shown in red. Significance is determined as FDR ≤ 0.05. **(E)** Gene Ontology pathway analysis was performed using significantly different expressed genes between *KRAS^G12D^EZH2^ΔSET^* and *Mist1^creERT/+^KRAS^G12D^* tissue. Top 20 pathways are shown. Bars represent the percentage of genes enriched in each pathway while yellow dots indicate adjusted P values.

To determine if these differences were mirrored by changes in gene expression, we compared ChIP-seq results to RNA-Seq analysis performed at the same time. Fewer DEGs were identified between *Mist1^creERT/+^KRAS^G12D^*and *KRAS^G12D^Ezh2^ΔSET^* tissue (315 in total; 237 up and 78 down; **Figure 3D**) compared to the changes in H3K27me3 enrichment. GseGO analysis of these DEGs identified dysregulated immune-related pathways (all top 20 pathways; p_adj_ ≤ 0.05; **Table S3** and **Figure 3E**) and pathways linked to ECM organization. Comparisons of DEGs between WT, *KRAS^G12D^Ezh2^ΔSET^* and *Mist1^creERT/+^KRAS^G12D^*tissue identified 81 DEGs with decreased expression in *Mist1^creERT/+^KRAS^G12D^*tissue relative to both *WT* and *KRAS^G12D^Ezh2^ΔSET^* tissue. Since these genes require EZH2 to be downregulated (**Figure S5B, Appendix 3**), they may represent direct targets of EZH2 during *KRAS^G12D^*-mediated reprogramming. Conversely, 80 DEGs were significantly higher in *Mist1^creERT/+^KRAS^G12D^*compared to WT and *KRAS^G12D^Ezh2^ΔSET^* tissue, which suggests upregulation requires EZH2 (**Appendix 3**).

Based on pathway analysis, EZH2 appears to regulate immune pathways triggered by *KRAS^G12D^*, consistent with previous studies suggesting KRAS works through NFATc1 to affect the inflammatory response (25). However, RNA-seq data showed *Nfatc1* was not differentially expressed between genotypes (**Figure S5C**). To identify the impact of EZH2 SET domain deletion on immune cell infiltration, the accumulation of immune cells into *Mist1^creERT/+^KRAS^G12D^*and *KRAS^G12D^Ezh2^ΔSET^* tissue was compared two weeks after injury by flow cytometry (**Figure 4**). In *Mist1^creERT/+^KRAS^G12D^*mice, 9.24% of the total cells were CD45^+^ (immune cells), while in *KRAS^G12D^Ezh2^ΔSET^* tissue, infiltrating CD45^+^ immune cells made up less than 2% (1.79%) of the total cell number (**Figure 4A**). This is consistent with the RNA-seq data indicating immune-related genes were increased only in *Mist1^creERT/+^KRAS^G12D^*even before overt morphological changes were observed. CD3^+^ T-Cells, CD3^+^CD4^+^ T-cells (**Figure 4B**), and CD163^+^ M2 macrophages (**Figure 4C**) all appeared to be reduced in *KRAS^G12D^Ezh2^ΔSET^* tissue suggesting reduced inflammation.

**Figure 4:**
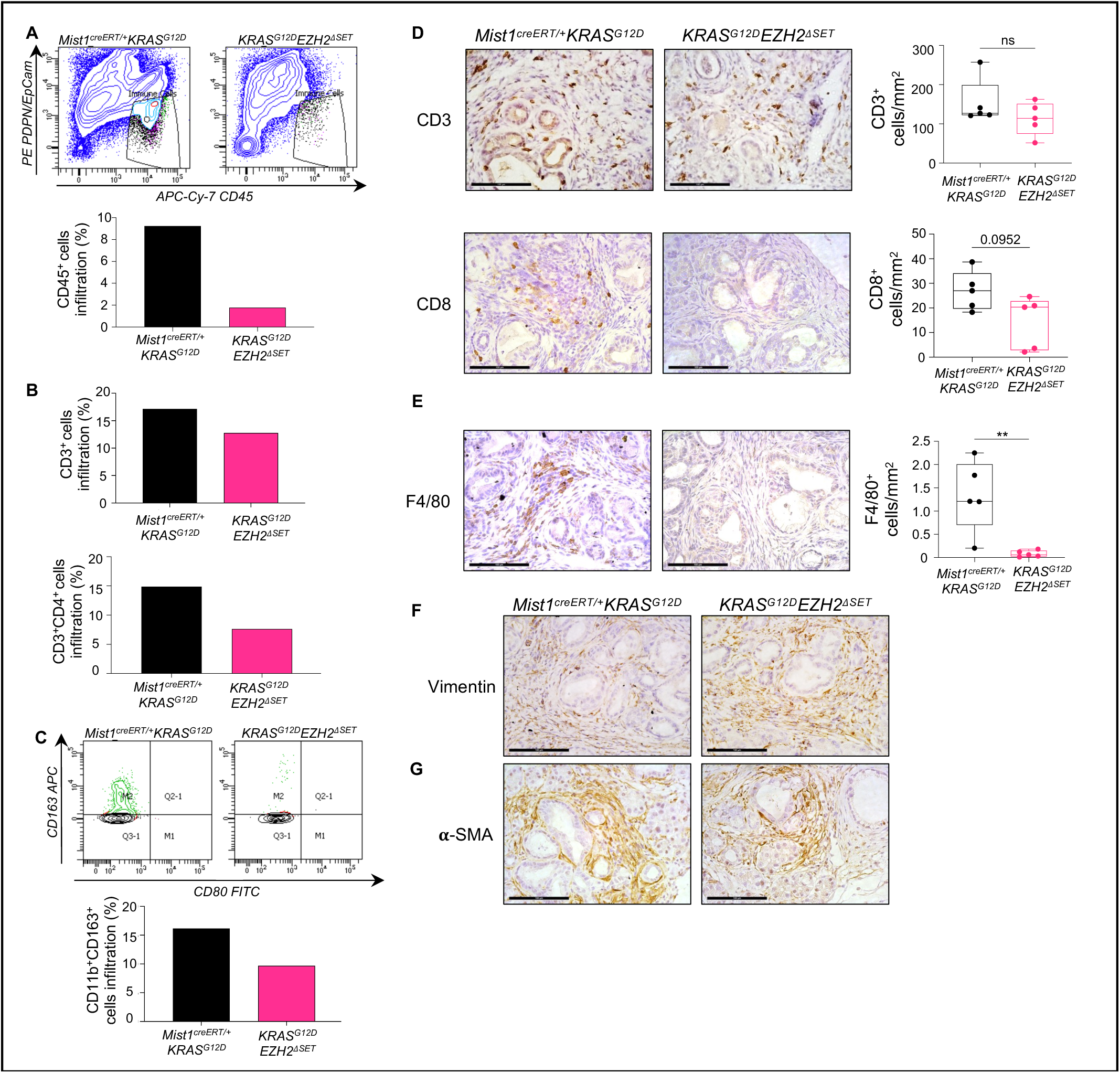
EZH2 deletion induces an immunosuppressive environment but don’t impact stroma organization after acute cerulein treatment. Flow cytometry analysis of CD45^+^ immune cells infiltration **(A)**, CD45^+^CD3^+^CD4^+^ T-cell infiltration **(B)** and CD45^+^CD11b^+^CD163^+^ M2 macrophage infiltration **(C)** in *Mist1^creERT/+^KRAS^G12D^*and *KRAS^G12D^Ezh2^ΔSET^*following cerulein treatment (n=1). Representative images of IHC for **(D)** CD3 **(E)** CD8, and **(F)** F4/80 positive cells in pancreas tissue from *Mist1^creERT/+^KRAS^G12D^* and *KRAS^G12D^Ezh2^ΔSET^* mice 51 days after tamoxifen gavage and 35 days following saline or cerulein treatment. Scale bar = 100 µm. Boxplots compare the mean number of positive cells and individual values are included. Data are shown as mean ± min to max (n=3). Significance was measured using a two-tailed unpaired Mann-Whitney test. **p≤0.01. **(G)** Representative images of IHC for vimentin or α-SMA staining on pancreatic tissue. Scale bar = 100 µm.

To confirm these findings, we performed IHC for the various immune cells on tissue 22 days after TX gavage, or five-weeks after cerulein treatment. IHC on pancreatic tissue 22 days after TX gavage showed negligible accumulation of CD3^+^, CD8^+^ and F4/80^+^ cells, regardless of genotype (**Figure S5D**). Five weeks after acute cerulein induction, *Mist1^creERT/+^KRAS^G12D^*tissue showed accumulation of CD3^+^ (T lymphocytes), CD8^+^ (cytotoxic T cells) and F4/80^+^ (macrophages) cells surrounding PanIN lesions (**Figures 4D-E**). Consistent with the flow cytometry data, *KRAS^G12D^Ezh2^ΔSET^* tissue showed reduced numbers of F4/80^+^ immune cells compared to *Mist1^creERT/+^KRAS^G12D^*mice but no significant changes in CD3^+^ T-cells. While a trend was observed for reduced accumulation of CD8^+^ T-cells, the difference between genotypes was not significantly different (**Figure 4D-E**). Similar analysis for vimentin and α-SMA, markers of cancer-associated fibroblasts, revealed no difference between *KRAS^G12D^Ezh2^ΔSET^* and *Mist1^creERT/+^KRAS^G12D^*tissue (**Figure 4F-G**). Combined, this data suggests an EZH2-dependent mechanism by which KRAS^G12D^ reprograms the acinar cell epigenome to promote differential infiltration of immune cells upon injury.

### Loss of EZH2 activity promotes rapid progression of PDAC in Mist1^creERT/-^KRAS^G12D^ model

Our findings regarding the role of EZH2 in early PanIN progression were different from previous studies, suggesting EZH2’s ability to restrict PanIN progression may differ depending on the extent of acinar cell differentiation or susceptibility of the model used. Previous studies using the *Mist1^creERT^* model showed a dramatic increase in KRAS^G12D^-mediated PDAC progression when MIST1 was completely absent, indicating loss of MIST1 increased sensitivity to KRAS^G12D^ (26). Therefore, we generated *Mist1^creERT/creERT^KRAS^G12D^* mice (indicated as *Mist1^creERT/-^KRAS^G12D^*) with and without the EZH2 ΔSET domain (*Mist1^creERT/-^KRAS^G12D^Ezh2^ΔSET^*; indicated as *MKE*; **Figure S6A**). 2-4 month-old mice were treated with tamoxifen and were followed for 2 months (**Figure S6B**).

Gross morphological analysis following TX gavage revealed no significant differences in weight between WT, *Mist1^creERT/+^KRAS^G12D^*, *KRAS^G12D^Ezh2^ΔSET^*, *Mist1^creERT/-^KRAS^G12D^* and *MKE* cohorts (**Figure S6C**). However, three *MKE* mice needed to be sacrificed within two months of TX treatment due to weight loss >15% of their initial body weight. Several mice expressing KRAS^G12D^ developed tumours within their oral mucosa (data not shown), likely due to *Mist1^creERT^* activity in this tissue, forcing us to cease the experiment at 60 days after initial TX treatment. Upon dissection, the pancreata of most genotypes appeared normal. However, fibrotic pancreatic masses were observed within *Mist1^creERT/-^KRAS^G12D^*(**Figure S6D**; blue arrows) that were not observed in *Mist1^creERT/+^KRAS^G12D^*and *KRAS^G12D^Ezh2^ΔSET^*, consistent with these mice developing preneoplastic nodules. Surprising, the absence of EZH2 in *Mist1^creERT/-^KRAS^G12D^* mice (*MKE*) dramatically increased nodule formation within the pancreas. In at least three *MKE* mice, multiple lesions appeared within the livers suggesting metastasis (pink arrows; **Figure S6D**), that were never observed in any other line. To confirm the presence of liver metastasis, we performed H&E and staining in *MKE* liver and identified presence of lesions (**Figure S6E**).

Following dissection, we examined EZH2 accumulation and observed increased mRNA (**Figure 5A**) and protein levels (**Figure 5B**) for EZH2 in *Mist1^creERT/-^KRAS^G12D^*pancreatic tissue that was decreased in *MKE* mice. Western blot analysis also showed a significant reduction in amylase expression in *MKE* tissue consistent with negligible acinar tissue in these mice (**Figure 5B**). H&E staining confirmed an almost complete loss of acinar tissue (**Figure 5C**) and development of high grade PanINs and PDAC in *MKE* mice (**Figure 5C** and **D**), while *Mist1^creERT/-^KRAS^G12D^* mice showed progression to PanINs (**Figure 5C**). Consistent with earlier experiments, *Mist1^creERT/+^KRAS^G12D^*and *KRAS^G12D^Ezh2^ΔSET^* mice showed few lesions, which consisted of ADM (acinar-to-duct cell metaplasia) and low grade PanINs. *MKE* pancreatic tissue also exhibited widespread fibrosis and invasive PDAC throughout the pancreas, different from all other genotypes.

**Figure 5.**
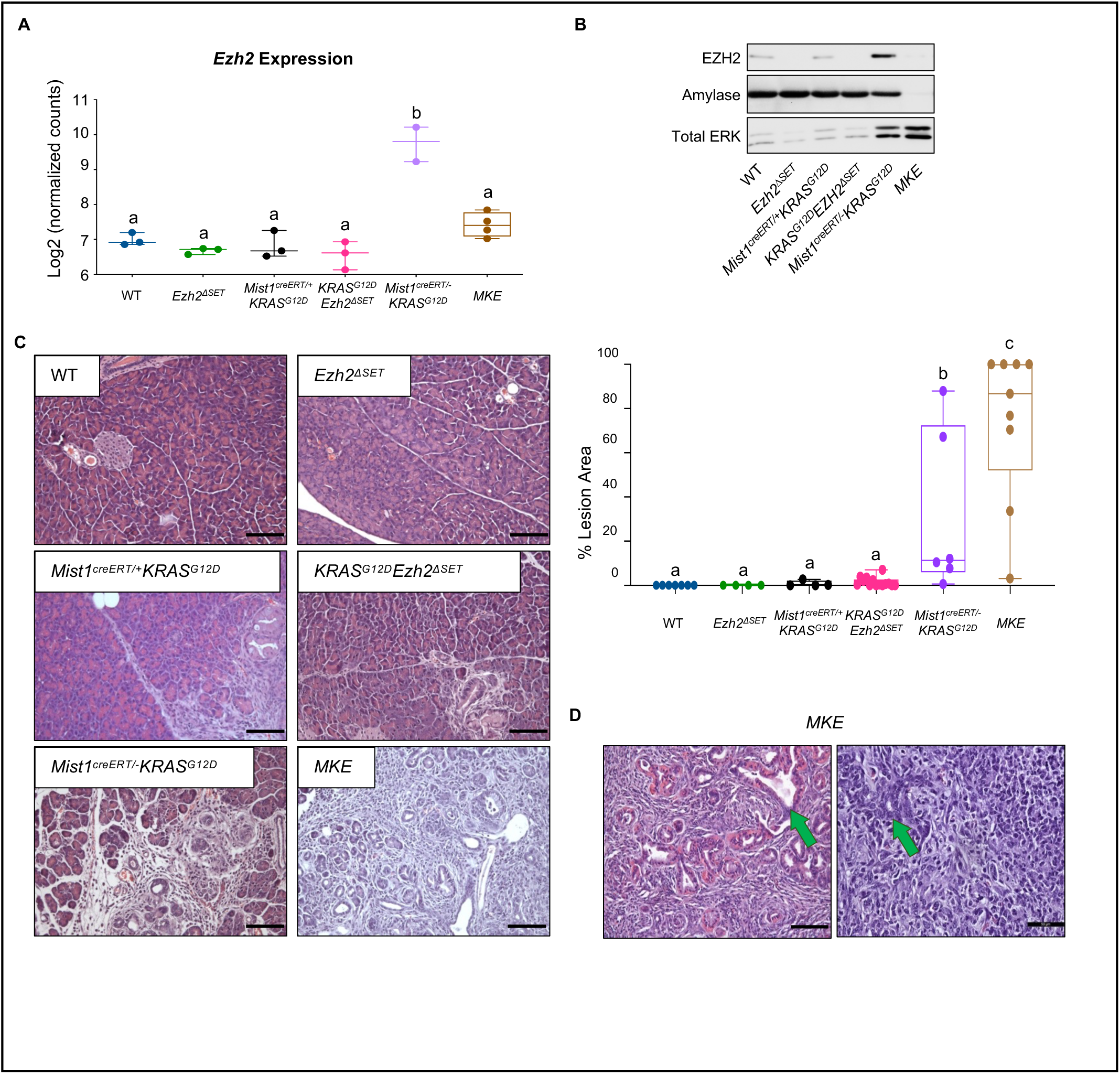
Combined loss of MIST1 and EZH2^ΔSET^ promotes rapid loss of acinar tissue in the presence of *KRAS^G12D^*. **(A)** RNA-seq analysis revealed significantly increased *Ezh2* mRNA in *Mist1^creERT/-^KRAS^G12D^*pancreatic tissue 22 days after KRAS^G12D^ induction relative to all other genotypes p≤0.001. **(B)** Representative western blots for EZH2, amylase or total ERK, 60 days after KRAS^G12D^ induction. **(C)** Representative H&E-stained pancreatic sections from wild type (WT), *Mist1^creERT/+^Ezh2^ΔSET^* (*Ezh2^ΔSET^*), *Mist1^creERT/+^KRAS^G12D^*, *KRAS^G12D^Ezh2^ΔSET^*, *Mist1^creERT/-^ KRAS^G12D^*, and *MKE* mice 60 days after KRAS^G12D^ induction. Scale bar = 100 µm. Right panel show a boxplot which quantifies the % lesion area in all genotypes based on H&E staining. **(D)** Higher magnification images of H&E-stained pancreatic tissue from *MKE* mice. Green arrows indicate high grade PanIN lesions and putative PDAC that is only found in these animals. Scale bar= 50 µm. Data are shown as mean ± min to max. Significance is measured by one-way ANOVA follow by a Tukey’s post-hoc test. Different letter indicate statistically different p values.

Biochemical analysis supported the histological analysis. Significantly reduced or negligible amylase expression was observed in all *MKE* mice (**Figures 5B, 6A, 6B**), indicating an absence of acinar tissue in these mice. Conversely, examination of ERK, a pathway activated by KRAS and linked to tumor development, showed higher levels of accumulation in *Mist1^creERT/-^KRAS^G12D^* and MKE extracts (**Figure 5B**), supporting progression from normal to diseased tissue. The increased development of extensive PanIN lesions in *MKE* mice was confirmed by IHC for CK19 (**Figure 6C**) and IF for SOX9 (**Figures 6D, 6E)**. Combined, this data indicated *EZH2^ΔSET^* deletion significantly increased KRAS^G12D^-mediated PanIN progression to PDAC in a susceptible model (*Mist1^creERT/-^KRAS^G12D^*).

**Figure 6:**
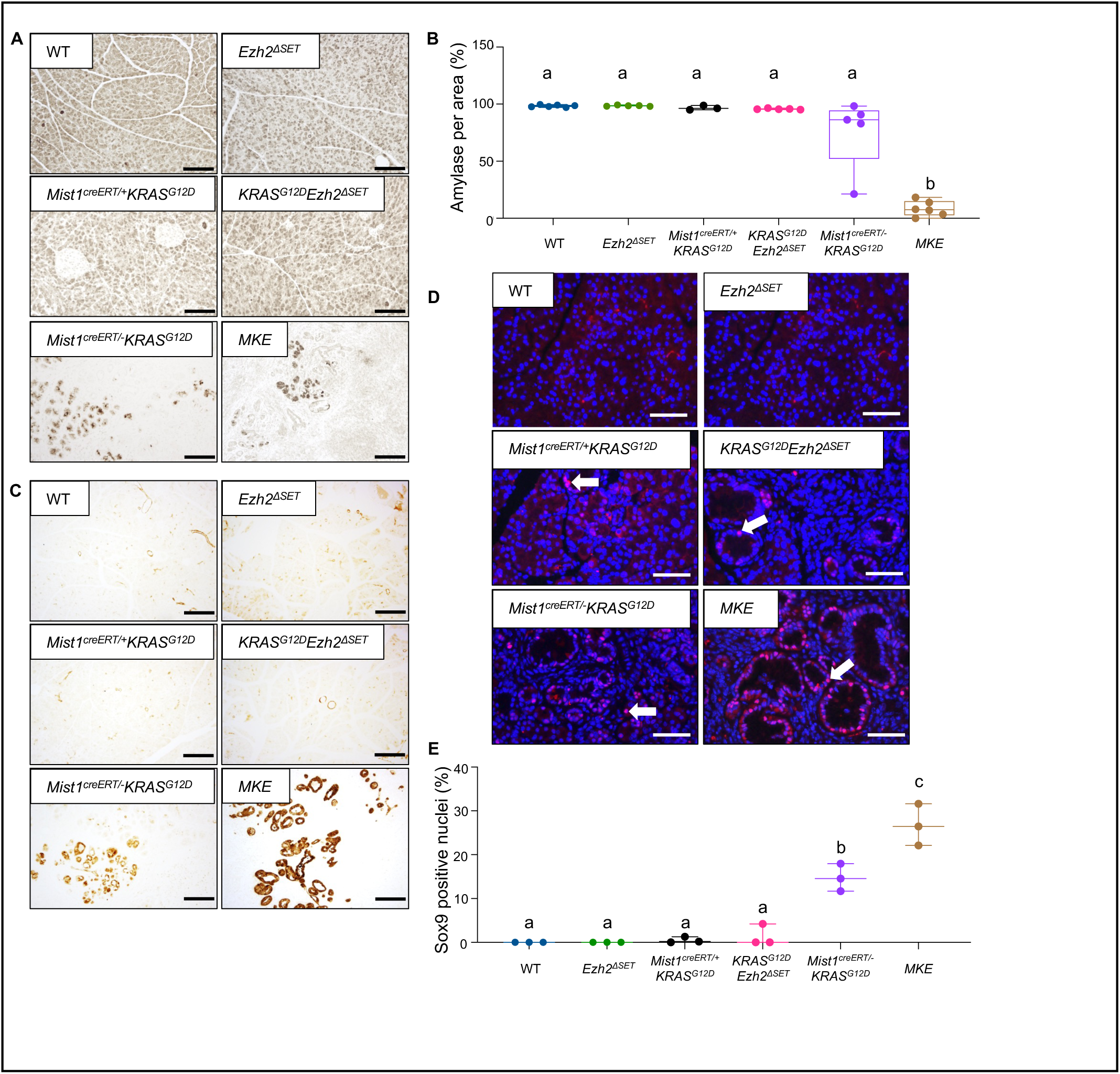
*MKE* mice exhibit extensive ductal and PanIN lesion progression. Representative IHC for **(A)** amylase or **(C)** CK-19 on pancreatic tissue from wild type (WT), *_Mist1_creERT/+_Ezh2_ΔSET* _(*Ezh2*_*ΔSET*_), *Mist1*_*creERT/+_KRAS_G12D*_, *KRAS*_*G12D_Ezh2_ΔSET*_, *Mist1*_*creERT/-KRAS^G12D^*, and *MKE* mice 60 days after KRAS^G12D^ induction. Scale bar = 100 µm. **(B**) Quantification of amylase staining in the various genotypes based on IHC staining. **(D)** Representative immunofluorescence for SOX9 on pancreatic section from *Mist1^creERT/+^KRAS^G12D^*, *KRAS^G12D^Ezh2^ΔSET^*, *Mist1^creERT/-^KRAS^G12D^*, and *MKE* mice 60 days after KRAS^G12D^ induction. Nuclei are counterstained with DAPI. White arrow represents positive SOX9 staining. Scale bar = 50 µm. **(E)** Quantification of SOX9 staining in the different mouse lines based on IF staining. In all cases, data are shown as mean ± min to max. Significance is measured by one-way ANOVA followed by a Tukey’s post hoc test. Different letters indicate statistically different p values.

To understand how *EZH2^ΔSET^* deletion enhanced PDAC progression, we aligned RNA-seq data from *Mist1^creERT/-^KRAS^G12D^*and *MKE* samples with the earlier 22-day RNA-Seq data (**Figure 3**). At this time point, no pre-neoplastic lesions were observed in most of the genotypes, while histological analysis of *MKE* tissue readily identified focal ADM lesions throughout the tissue (**Figure 7A**). Therefore, we expected the transcriptome to be vastly different in these mice, and principal component analysis confirmed *MKE* mice clustered away from all other genotypes (**Figure 7B**). Global RNA expression analysis between *Mist1^creERT/-^KRAS^G12D^*and *MKE* tissue identified 7,636 DEGs (3,843 up and 3,793 down) (**Appendix 4**) between these two genotypes (**Figure 7C**), and Gene Set Enrichment Analysis (GSEA) identified >150 significantly altered pathways (**Table S4**) including many within the top 20 enriched pathways related to TME remodeling and inflammation (**Figure 7D**). These results suggest a profound effect of deleting EZH2 on the TME of *MKE* mice that exists prior to overt morphological changes. In confirmation that EZH2 affects pro-inflammatory pathways, we observed an increase in the inflammation response based on GSEA specifically in *MKE* mice (**Figure 7E**), and *Ptgs2* expression, which encodes the pro-inflammatory protein COX2 (**Figure 7F**) is markedly increased only in *MKE* tissue. GSEA analysis also identified an increase in extracellular matrix organization specifically in *MKE* tissue (**Figure 7G**), and trichrome blue histology confirmed increased fibrosis in *MKE* pancreata (**Figure 7H**). Importantly, these pathways were enriched before the tissue showed significant transformation and fibrosis which reinforces the hypothesis that EZH2 play a role in priming TME genes at an early time-point in PDAC development. As expected, loss of EZH2 also correlated to downregulation of pathways related to the nucleosome and chromatin remodelling suggesting significant effects on the acinar cell genome as confirmed by the GSEA plot analysis which show a decrease in nucleosome assembly in *MKE* tissue (**Figure 7G**).

**Figure 7:**
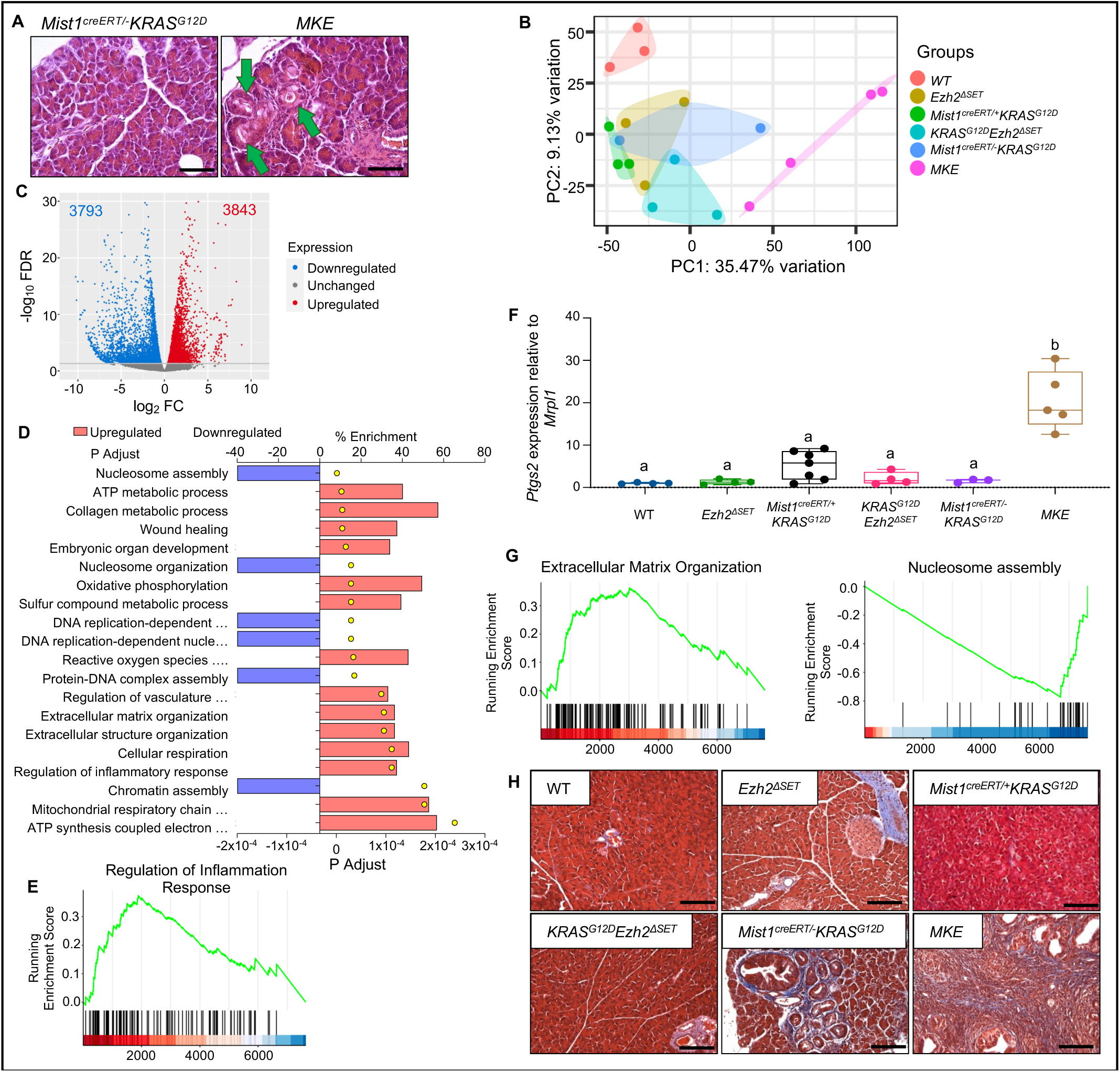
Acinar-specific deletion of *Ezh2^ΔSET^* in *KRAS^G12D^*-mediated PDAC alters the tumour microenvironment. **(A)** Representative H&E staining of pancreatic tissue from *Mist1^creERT/-^KRAS^G12D^* and *MKE* mice 22 days after KRAS^G12D^ induction. Green arrows indicate ADM in *MKE* mice. Scale bar = 50 µm. **(B)** Principal component clustering of mouse samples based on RNA-Seq data 22 days after KRAS^G12D^ induction. **(C)** Volcano plot showing differentially expressed genes between *Mist1^creERT/-^KRAS^G12D^*and *MKE* mice 22 days after KRAS^G12D^ induction and based on RNA-Seq analysis. Genes with significantly lower (3793 genes) or higher (3843 genes) expression in *MKE* mice are indicated in blue and red, respectively. Significance was determined with a FDR ≤ 0.05. **(D)** Top 20 pathways identified by gseGO analysis based on RNA-Seq analysis. (p adjusted ≤ 0.05). **(E)** GSEA analysis representing the KEGG term “Regulation of inflammation response” is increased in *MKE* tissue compared to *Mist1^creERT/-^KRAS^G12D^.* **(F)** qRT-PCR for *Ptgs2* expression in wild type (WT), *_Mist1_creERT/+_Ezh2_ΔSET* _(*Ezh2*_*ΔSET*_), *Mist1*_*creERT/+_KRAS_G12D*_, *KRAS*_*G12D_Ezhz2_ΔSET*_, *Mist1*_*creERT/-KRAS^G12D^*, and *MKE* mice pancreatic tissue 22 days after KRAS^G12D^ induction. Data are shown as mean ± min to max. Significance was determined with a one-way ANOVA follow by Tukey’s correction post hoc analysis. *MKE* is significantly different from all the other genotypes p≤0.0001. **(G)** GSEA analysis indicated increased enrichment of genes involved in “Extracellular matrix organization” or decreased enrichment of genes involved in “Nucleosome assembly” in *MKE* tissue relative to *Mist1^creERT/-^KRAS^G12D^*. **(H)** Representative trichrome blue staining of pancreas section from wild type, *Mist1^creERT/+^EZH2^ΔSET^*, *Mist1^creERT/+^KRAS^G12D^*, *KRAS^G12D^EZH2^ΔSET^*, *Mist1^creERT/-^KRAS^G12D^*, and *MKE* mice 60 days after KRAS^G12D^ induction. Scale bar = 100 µm.

Given the timing of these changes (i.e. before significant pathology is observed in *MKE* mice), these findings suggest loss of EZH2 has a direct, cell autonomous effect on acinar cell differentiation. To circumnavigate the potential contribution of external influences on PDAC progression in *MKE* mice, we cultured acinar cells from all genotypes 22 days after TX gavage in a 3D collagen matrix (**Figure 8A**) and assessed ADM formation over 9 days. We observed a significant difference in ADM size and morphology specific to *MKE* cultures (**Figure 8B**). Consistent with previous studies, a portion of acini developed ADM structures regardless of genotype three days into culture. No difference in the size of these clusters was observed over the course of the experiment for all genotypes except *MKE* cultures. *MKE* acini more rapidly developed into ADM structures with almost 100% of the clusters forming ADM by day 3. These acini continued to increase in size throughout the culture period and showed no signs of undergoing apoptosis or necrosis. Acinar cells derived from all other genotypes showed significantly increased ADM relative to WT cultures; however, except for *Mist1^creERT/-^KRAS^G12D^*cultures, the number of ADM structures decreased beginning at day 5 with most clusters dying by day 9 (**Figure 8C**). Staining for Ki67 identified proliferating cells only in *MKE* and *KRAS^G12D^EZH2*^Δ*SET*^ ADM, consistent with previous reports of EZH2-mediated regulation of cell cycle genes (**Figure S7**; (12, 27, 28)).

**Figure 8:**
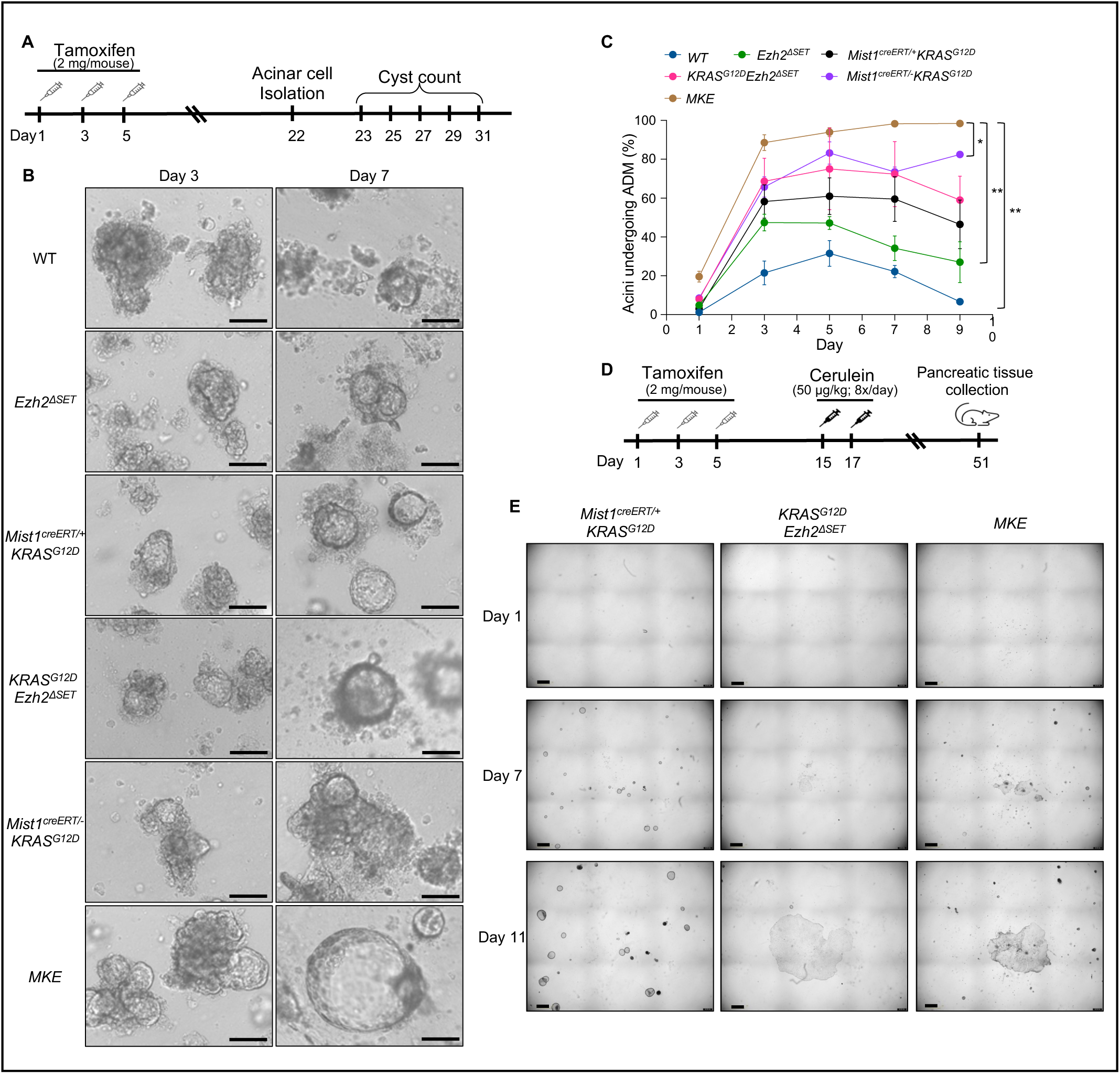
Acinar cells *EZH2^ΔSET^* deletion has different cell autonomous roles depending on the context in which *KRAS^G12D^* is expressed. **(A)** Experimental design for acinar cell isolation and embedding into collagen 22 days after KRAS^G12D^ induction. **(B)** Representative images of cell clusters three and seven days after acinar cell isolation. Genotypes included wild type (WT), *_Mist1_creERT/+_Ezh2_ΔSET* _(*Ezh2*_*ΔSET*_), *Mist1*_*creERT/+_KRAS_G12D*_, *KRAS*_*G12D_Ezh2_ΔSET*_, *Mist1*_*creERT/-KRAS^G12D^*, and *MKE.* Scale bar = 100 µm. **(C)** Quantification of the percentage of cell clusters with visible ADM, one to nine days after acinar cell isolation. 50 or more clusters were counted for each condition. Data are shown as mean ± SEM. Significance is measured by two-way Anova follow by Dunnett’s correction. *p≤0.05, **p≤0.01, ***p≤0.001. **(D)** Experimental timeline of TX gavage and acute cerulein-induced pancreatitis in *Mist1^creERT/+^KRAS^G12D^*, *KRAS^G12D^Ezh2^ΔSET^* and MKE mice. **(E)** Representative images of organoid development in matrigel 1, 7 and 11 days after cell isolation. Genotypes included *Mist1^creERT/+^KRAS^G12D^*, *KRAS^G12D^Ezh2^ΔSET^* and *MKE.* Scale bar = 800 µm.

Based on these 3D collagen cultures, it appears loss of acinar-specific EZH2 deletion favours ADM progression in a susceptible model. To further investigate the role of EZH2 in PanINs and PDAC development, we developed 3D organoid cultures from *Mist1^creERT/+^KRAS^G12D^, KRAS^G12D^Ezh2^ΔSET^* and *MKE* pancreatic tissue two weeks following acute cerulein induction (**Figure 8D**). After tissue isolation, organoid formation and morphology was monitored. Organoids were readily observed in *Mist1^creERT/+^KRAS^G12D^* cultures, but surprisingly, the *KRAS^G12D^Ezh2^ΔSET^* cultures showed few, smaller organoids, a difference that was maintained over several passages (**Figure S8A-B**). Interestingly, for the establishment of organoids, loss of EZH2 seems to slow down development of large cyst organoids in *MKE* cultures compare to *Mist1^creERT/+^KRAS^G12D^*(**Figure 8E**). But only one and two passages after establishment, *MKE* organoids showed rapid growth and larger cyst-like organoids compared to the other genotypes (**Figure S8A-B**). These 3D cultures highlight that cell autonomous events are, at least partially, responsible for the *MKE* phenotype and increased progression to PDAC observed in *MKE* mice and support a contextual role for EZH2 in early PDAC progression.

## Discussion

In this study, we examined the role of EZH2 in initiating pre-neoplastic lesions and progression to PDAC and its impact on epigenetic remodeling in the presence of KRAS^G12D^. Using a model allowing inducible loss of the EZH2 SET domain in acinar cells of adult mice, we showed EZH2 is required for reprogramming the acinar cell genome in response to constitutive activation of KRAS^G12D^. EZH2 is dispensable for acinar cell regeneration following pancreatic injury but restricts PanIN progression following acute injury combined with *KRAS^G12D^*. While this difference did not result in high-grade PanIN lesions, loss of EZH2^ΔSET^ activity greatly enhanced PDAC progression in mice susceptible to KRAS^G12D^, leading to spontaneous loss of acinar tissue, significant fibrosis, and PDAC within 60 days. This is the first study that examines changes in acinar cell H3K27me3 enrichment profiles directly related to KRAS^G12D^ expression, how these changes are affected by EZH2 function, and shows context-specific roles for EZH2 that promote or restrict early PanIN progression. This study also highlights the importance of epigenetic reprogramming in the context of PDAC and suggests EZH2 restricts early PanIN progression to PDAC through priming of immune and inflammatory genes.

### KRAS^G12D^ promotes epigenetic repression of the acinar cell genome

Our findings support a model in which KRAS^G12D^ promotes general epigenetic repression within the pancreas prior to overt morphological changes. Global enrichment of H3K27me3 was increased in *Mist1^creERT/+^KRAS^G12D^* compared to wild type tissue, while global H3K4me3 enrichment was similar between KRAS^G12D^ +/-tissue. This is consistent with studies showing increased expression and activity of DNA methyltransferases, Histone Deacetylases, and PRC1 and 2 in PDAC (29–32), all of which promote epigenetic repression. Importantly, epigenetic reprogramming does not accompany widespread transcriptomic dysregulation, suggesting changes in the epigenome predate transcriptional differences and may be masked until additional environmental stresses are present. We previously characterized similar epigenetic reprogramming of acinar cells in response to chronic stress which suggested reprogramming alters the molecular response to subsequent acute stimuli (33). One mechanism that underlies reprogramming involved changes to “poised” genes, which have bivalent epigenetic enrichment for active and repressive epigenetic marks. This epigenetic bivalency allows repressed genes to be rapidly activated and involves H3K27me3 enrichment. The widespread enrichment of H3K27me3 following KRAS^G12D^ activation suggests EZH2, in part, regulates reprogramming. In support of these findings, deletion of EZH2 in the presence of KRAS^G12D^ resulted in H3K27me3 enrichment levels comparable to WT tissue.

### Loss of EZH2 leads to epigenetic reprogramming of pathways involved inflammation

While EZH2 has been targeted in several other studies examining its role in PDAC (15, 34, 35), this is the first study that examines global H3K27me3 enrichment in the context of EZH2 loss of function. H3K27me3 ChIP-Seq revealed increased enrichment of immune-related pathways 22 days after KRAS^G12D^ induction that appears to prime the genome for a differential inflammatory response since no immune cell infiltration was observed in the pancreas until after induction of injury. Two and five weeks after pancreatic injury, *Mist1^creERT/+^KRAS^G12D^*mice showed an increase in CD45^+^ immune cell infiltration such as CD3^+^, CD8^+^ lymphocytes and CD163^+^ M2 macrophages cells. The accumulation of each of these cell types was reduced in the absence of EZH2^ΔSET^ activity, suggesting loss of EZH2 activity in *KRAS^G12D^Ezh2^ΔSET^* mice drives an immune cold environment. A global reduction of CD45^+^ immune cell infiltration and decreased accumulation of CD4^+^ and CD8^+^ cells would promote PDAC progression as their presence is associated with improved prognosis of PDAC patients (36–38). These findings support recent studies that propose a direct role for EZH2 in immune cell recruitment and activation in cancer (39, 40) and suggest EZH2 plays a protective role in early PanIN development by increasing specific CD45^+^ immune cell infiltration such as CD4^+^ and CD8^+^ cell recruitment. However, the absence of EZH2 activity also results in decreased accumulation of F4/80^+^ and CD163^+^ M2 macrophages cells, which are generally associated with enhanced PDAC progression since these favor immunosuppressive environment (41, 42). The contradiction supports a more complex involvement of EZH2 that is likely stage dependent. This is supported by analysis of organoids developed from *KRAS^G12D^Ezh2^ΔSET^* and *Mist1^creERT/+^KRAS^G12D^*pancreatic tissue, which showed no difference *in vivo*, but exhibited significantly different abilities *ex vivo*. Organoids developed from *KRAS^G12D^Ezh2^ΔSET^* tissue had reduced size and number suggesting Ezh2-negative lesions have a reduced ability for long term progression. This phenomenon is consistent with observations in Chen et al (14), which showed the increased PanIN progression initially observed in the absence of EZH2 was not maintained at later stages.

### Loss of EZH2^ΔSET^ activity enhances a susceptible environment for KRAS^G12D^-mediated PDAC

As mentioned, while our findings suggest a protective role for EZH2 in limiting early PanIN progression, previous studies on EZH2 show a more critical role in early stages of PDAC. Using the same floxed *EZH2^ΔSET^* allele, Mallen-St Clair et al (12) showed acinar cell regeneration was restricted following cerulein-induced injury and KRAS^G12D^ increased PanIN initiation progression, consistent with a more restrictive role for EZH2 methyltransferase function (12, 14). As mentioned above, Chen et al (14) supported these findings but suggested EZH2 was necessary for maintaining pre-neoplastic lesions, with fewer PanIN lesions apparent in older mice. Our results revealed negligible effects on acinar cell regeneration and *Mist1^creERT/+^KRAS^G12D^* and *KRAS^G12D^Ezh2^ΔSET^* mice developed similar numbers of PanINs lesions following injury. We suggest the discrepancy in our results arises, in part, from the *cre* driver used in the two studies having different effects on susceptibility to KRAS^G12D^. Previous studies achieved *Ezh2^ΔSET^* deletion by targeting a non-inducible *cre* recombinase to the *Ptf1a* or *Pdx1* genes resulting in *KRAS^G12D^*activation in early pancreatic development, prior to differentiation of mature pancreatic cell types. EZH2 is important for early development and specification (8) of acinar and liver cells from a common endodermal origin. In the absence of EZH2, epigenetic programs that fix in the differentiation status of mature cell types are absent. *Mist1^creERT^*mice allow *Ezh2^ΔSET^* deletion and *KRAS^G12D^* activation only in mature acinar cells when mature epigenetic programs are already in place. Therefore, epigenetic programs that establish an adult phenotype are not affected. In addition, haploinsufficiency for *Ptf1A* likely affects the response to KRAS^G12D^. Loss of a single *Ptf1a* allele alters the cell fate of acinar cells (43) which increases the potential for undergoing ADM. Conversely, loss of a single *Mist1* allele shows no differences in acinar cell function, response to injury or gene expression when compared to wild type litter mates. Only when MIST1 is completely absent do acinar cells show incomplete differentiation and increased sensitivity to injury and KRAS^G12D^ (26, 44). In support of the importance of the cre driver for studying PanIN progression, comparison of *Ptf1a^creERT/+^KRAS^G12D^*mice to *Mist1^creERT/+^KRAS^G12D^*or *Elastase^creERT/+^KRAS^G12D^* mice showed marked differences in sensitivity to cerulein-induced injury (Mousavi et al, in preparation).

### Loss of EZH2 methyltransferase activity leads to both cell autonomous and non-cell autonomous effects on PDAC progression

Despite the differences, both the current study and Chen et al (14)vconfirm a protective role for EZH2 in restricting early PanIN progression and PDAC development. We suggest this effect of EZH2 is through both cell autonomous and non-cell autonomous effects within the pancreas. RNA sequencing at 22 days revealed loss of EZH2 activity in *Mist1^creERT/-^KRAS^G12D^*mice (i.e. *MKE*) leads to activation of pathways involved in TME remodeling that favour aggressive PDAC progression (45–47) and the rapid progression to ADM and PanINs appears to be independent of the TME. RNA-seq analysis also revealed loss of EZH2 had a significant impact on pathways affecting chromatin stability in *MKE* acini suggesting a cell autonomous role for EZH2 in acinar cell reprogramming. This role was confirmed by culturing acinar cells of all genotypes in collagen 22 days after tamoxifen-induced recombination or culturing organoids from *Mist1^creERT/+^KRAS^G12D^* and MKE genotypes following acute cerulein treatment. In collagen cultures, *MKE* acini showed rapid ADM compared to other genotypes, with increased proliferation, and maintained survival over the length of culture. In matrigel cultures, *MKE* organoids show rapid growth and formed and maintained larger cyst structures compared to the *Mist1^creERT/+^KRAS^G12D^* cultures.

Targeting EZH2 function has been suggested as a possible therapy based on *in vitro* and xenograft data showing EZH2 inhibitors can enhance sensitivity to traditional chemotherapy (48). While our results support and extend findings of the importance of EZH2 in restricting progression to PDAC, they do not agree with studies on PDAC cell lines or tissue obtained from patients. Increased EZH2 expression in PDAC is correlated to worse prognosis and resistance to therapy (13, 15, 34). Crucially, these previous studies suggest the effects of EZH2 are independent of its methyltransferase activity. Therefore, it is likely EZH2 has additional, non-PRC2 functions relevant to late stage PDAC, which our study does not addressed. However, our findings suggest targeting EZH2 in PDAC with pharmacological inhibitors must be approached with caution.

To conclude, our study shows EZH2 limits progression from acinar cells to late stage PDAC through reprogramming of inflammatory and extracellular matrix genes. These effects are likely through both non-cell autonomous and cell autonomous mechanisms. Loss of EZH2 alters pathways that promote inflammation and fibrosis, thereby affecting the TME, but also enhances ADM in the absence of the TME. This work highlights a complex role for EZH2 in initiating and progression of pancreatic cancer. While our findings support a tumour suppressive role in restricting PanIN and PDAC formation, future studies are needed to determine if these effects are simply due to PRC2-related functions or additional modes of EZH2 activity.

## Materials and Methods

### Mouse models

In all experiments, both male and female mice were used to reach significance. Mice were given normal chow and water ad libitum throughout the experiment. C57/Bl6 mice containing *loxP* sites flanking exons 16 to 19 of the *Ezh2* gene (encompass the SET domain; *Ezh2^ΔSET/ΔSET^*), an oncogenic KRAS^G12D^ within the *Kras* locus and downstream of a *loxP-stop-loxP* (*LSL*) cassette (*Kras^LSL-G12D^*), or an inducible cre recombinase (creERT) targeted to the *Mist1* coding region (*Mist1^creERT^*), have been used and described previously (12, 17, 22, 49, 50). Mating of these transgenic lines lead to eight distinct genotypes which were confirmed before and after experimentation using the primers indicated in **Table S5**. To induce *loxP* recombination, two to four-month-old mice were gavaged 3 times over 5 days with 2 mg tamoxifen (TX; Sigma #T5648) in corn oil (Sigma #C8267). This regime has been used previously to induce >95% recombination in acinar cells of the *Mist1^creERT^* line (26, 51). Mice were sacrificed either 22 days or 60 days after the initial TX gavage or treated with cerulein to induce acute or recurrent pancreatic injury (see below). Pancreatic tissue was weighed and processed for paraffin sectioning, RNA, chromatin, or protein isolation.

### Cerulein-induced pancreatitis

To induce acute pancreatic injury, 2-4-month-old mice received eight hourly intraperitoneal injections of cerulein (50 mg/kg, MedChemExpress, #FI-6934) 15 and 17 days after the first dose of TX. Control mice received 0.9% saline solution. Mice were weighed every day to monitor weight changes and health, then sacrificed 14 or 35 days after initiating acute CIP.

To induce recurrent pancreatic injury, mice received intraperitoneal injections of cerulein (250 μg/kg body weight) or 0.9% saline solution (control) twice daily (9:00 h and 15:00 h) for 14 days. Mice were weighed daily to determine changes in body weight. Mice were sacrificed 7 days after the last cerulein injections.

### RNA isolation, RNA-seq and data analysis

RNA was isolated from whole pancreatic tissue of mice 22 days after TX induction using Trizol (Invitrogen, #15596018) followed by the Pure link kit following manufacturer’s instructions (Invitrogen, #12183018A). RNA was prepared for RNA-seq as previously described (33). Two (for *Mist1^creERT/-^KRAS^G12D^*), three (*WT*, *EZH2^ΔSET^*, *Mist1^creERT/+^KRAS^G12D^* and *KRAS^G12D^EZH2^ΔSET^*), or four (*MKE*) biological replicates per group were sequenced using the Illumina NextSeq High Output 150 cycle (paired-end sequencing) sequencing kits. The complete RNA-seq data can be found at GEO accession GSE (currently in process). RNA-seq reads were aligned to mouse genome mm10 and sorted by coordinate using STAR v2.7.9a (52). Gene counts were generated using the featureCounts function of the Subread v2.0.3 aligner (53) and the subsequent differential expression analysis performed using the edgeR v3.321 package (54, 55). Functional and enrichment analysis, including KEGG and Gene Ontology (GO) pathway analyses and Gene Set Enrichment, were performed using clusterProfiler v3.18.1 R package (56). A threshold of p-adjusted ≤ 0.05 cut off was used for all differential expression and pathway analyses. PCA plots (PCAtools: Everything principal components analysis; Blighe and Lun, 2020 R package version 2.2.0), Venn diagrams (VennDiagram: Generate high-resolution Venn and Euler plots; Chen, 2022 R package version 1.7.3), and dotplots (enrichplot: Visualization of functional enrichment results; Guangchuang, 2021 R package version 1.10.2) were generated using the corresponding R package.

### ChIP-Sequencing and data analysis

Chromatin was isolated from pancreatic tissue of mice 22 days after TX gavage. The ChIP-seq protocol was followed as previously described (33). Antibodies against trimethylated H3K27 (H3K27me3; Millipore Sigma #07-449) or trimethylated H3K4 (H3K4me3; Millipore Sigma #04-745) were used for immunoprecipitation and subsequent next-generation sequencing performed using Illumina NextSeq High Output 150 cycle sequencing kit. The complete ChIP-seq data can be found at GEO accession GSE (currently in process). Raw data were first checked for read quality using FastQC and aligner against the mouse genome (mm10) using bowtie2 tool (57). Identification of the peaks for each sample was performed using Homer FindPeaks tool with the “histone” mode, which searches for broad regions of enrichment of variable width by comparing both local background and corresponding input samples. Genomic annotation and visualization of the peaks was performed using ChIPSeeker R package and *TxDb.Mmusculus.UCSC.mm10.knownGene* library. To define the target genes with significant ChIP enrichment, we defined the promoter region of +/-3 kb from the TSS (transcription start site). Genes overlapping at least one identified peak were considered target genes for a given sample. KEGG enrichment analysis was performed based on the resulting lists of target genes using ClusterProfiler R package. Heatmap visualization of the ChIP enrichment was performed using ngs.plot tool (58) with decreasing ranking of genes based on the ChIP enrichment level among the gene body.

### Flow Cytometry

Pancreatic tissue was cut into small pieces, placed in GentleMACS C tubes, and lysed with Tumor Dissociation Kit according to supplier’s protocol (Miltenyi, #130-095-929). After dissociation, cell pellets were filtered using 100 µM cell strainers. Cells suspensions were incubated 30 min at 4°C in the dark with the following antibodies (PE-PDPN, Thermofisher #12538180; PE-EpCam, Miltenyi #130117864; APC-CD163, Thermofisher #17163180; FITC-CD80, Miltenyi #130102532; BV450-CD11b, BD Biosciences #560456; BV605-CD8, BD Biosciences #563152; APC-Cy7-CD45, BD Biosciences #561037; PerCP-Cy5.5-CD3, BD Biosciences #560527; PE-Cy5-CD4, BD Biosciences #561836). Following labelling, cells were rinsed with PBS, fixed with 2% PFA for 10 minutes, resuspended in FACS buffer (PBS, BSA 5%, EDTA 5 mM) and then analysed by flow cytometry using the corresponding lasers (LSR-II cytometer, BD Biosciences).

### Real-time qRT-PCR analysis

Real time qRT-PCR was performed on cDNA samples prepared as described (33). Expression of *Ptgs2* were normalized to mitochondrial ribosomal protein L1 (*Mrpl1*). ViiA 7 RUO software (Applied Biosystems) was used to calculate the amount of RNA relative to wild type animals for the equivalent time points. Primer sequences are shown in **Table S6**.

### Tissue fixation and histology

For histological analysis, pancreatic tissue was isolated from the head and tail of the pancreas and processed as described (33). To assess overall histology and identify differences in pancreatic tissue architecture, sections were stained with H&E. Lesions area were quantified using ImageJ as percent of total tissue area. Mucin accumulation was visualized using an Alcian Blue stain kit (Abcam, #ab150662) and staining quantified as a percentage of the whole tissue area. Periodic-acid Schiff staining was also performed (Sigma-Aldrich Kit; #3951 and 3952) and quantified by scoring PanIN lesions as PAS+ (>50%), partially PAS+ (<50%), or PAS negative. To assess fibrosis, paraffin sections were stained using Trichrome Blue (Abcam, #ab150686). Lesions and other staining were scored over at least three sections from both the duodenal and splenic regions of the pancreas.

### Immunohistochemistry and immunofluorescence

IHC was performed on paraffin sections as described (33). Following antigen retrieval, sections were permeabilized with 0.2% Triton-X (BDH, #R06433) in PBS, rinsed, then blocked in 5% sheep serum in PBS for 1 hour at room temperature. Primary antibodies were diluted in 5% sheep serum in PBS and incubated overnight at 4°C. Primary antibodies included rabbit amylase (Cell signaling Technology, #4017, 1:400), rabbit CK19 (Abcam, #15463, 1:200), rabbit CD3 (BD Biosciences, #560591, 1:200), rabbit CD8 (Thermofisher, #98941, 1:200), rabbit F4/80 (Abcam, #ab111101, 1:100), rabbit α-SMA (Cell signaling technology, #19245, 1:200), rabbit Vimentin (Cell signaling technology, #5741, 1:400). Sections were washed, then incubated in biotinylated mouse α-rabbit IgG secondary antibody (in 5% sheep serum, Vector, #PK-4001, 1:1000) for 30 min at room temperature. Finally, sections were incubated in AB reagent for 30 min at room temperature and visualized using ImmPACT DAB Peroxidase (HRP) substrate (Vector, #PK-4001/SK-4105). Slides were counterstained with hematoxylin (Biocare Medical, #CATHE-M) and imaged using Leica Microscope DM5500B (Leica Microsystems) and LAS V4.4 software.

IF analysis was performed on paraffin-embedded tissue sections for SOX9 and for KI67, acinar cells were fixed in PFA 3% and then embedded in paraffin. Slides were prepared as for IHC except for quenching with hydrogen peroxidase (Fisher Scientific, #H325) for SOX9. Primary antibody is rabbit SOX9 (Millipore Sigma, #AB5535, 1:250) and mouse KI67 (BD Biosciences, #550609, 1:250). After washing, slides were incubated in α-rabbit or α-mouse IgG conjugated to TRITC (Jackson ImmunoResearch, #711-025-152 and #715-025-150, 1:300) diluted in 5% sheep serum in PBS. Prior to mounting in Vectashield Permafluor mountant (Thermo Fisher Scientific, #SP15), sections were incubated in DAPI (Thermo Fisher Scientific, #62248). Staining was visualized using Leica DFC365 FX camera on the Leica DM5500B microscope. Images were taken on Leica LASV4.4 software.

### Protein isolation and Western blotting

Pancreatic protein was isolated as described (59) and quantified using a Bradford protein assay (Bio-Rad, #5000006). Isolated protein was resolved by SDS-PAGE and transferred to polyvinylidene fluoride membrane (Bio-Rad, #162-0177). Western blot analysis was carried out as described (60) using antibodies specific for rabbit EZH2 (Cell signaling technology, #5246, 1:1000), rabbit Amylase (Abcam, #ab21156, 1:8000), rabbit Total ERK (Cell signaling technology, #9102, 1:1000). After washing, blots were incubated in α-rabbit HRP antibody (Cell signaling technology, #7074, 1:3000). Blots were visualized using the VersaDoc Imaging System with Quantity One 1-D Analysis software (Bio-Rad).

### Acinar cell isolation and 3D-Collagen culture

Acinar cells were isolated and embedded in collagen as previously described (61). Cyst formation was assessed every day until day 9 in culture. At day 7, some cultures were processed for paraffin sectioning and IF analysis for Ki67. Representative images were taken with an upright Leica microscope.

### Organoid isolation and 3D-Matrigel culture

The middle section of the pancreas was isolated and digested based on previously published protocols with some modifications (62). Pancreata was digested by incubation in 1 mg/ml of collagenase/dispase for 20 minutes at 37°C in a rotating incubator. Digested tissue was washed with DMEM/F12 containing with 10 mM HEPES, 1% glutamax, 1% PenStrep and 100 µg/mL primocin and centrifuged at 300 g for 5 minutes. Supernatant was aspirated and tissue resuspended in StemPro Accutase (Gibco, cat #A11105-01) and incubated for 45 minutes at 37°C in a rotating incubator. The resulting slurry was filtered through a 70 µm nylon mesh filter and cells resuspended in feeding media (63) with 5% Matrigel. 30 000 cells were seeded on a layer of 100% Matrigel (Corning, cat #356230). After first passage, organoids were reseeded into 100% Matrigel domes for experimental analysis according to (64). For passaging, organoids were incubated in 1 mg/ml of collagenase/dispase for 2 hours at 37°C, then rinsed with wash media and centrifuged at 300 g for 5 minutes. Supernatants were aspirated and cells resuspended in StemPro Accutase and incubated for 45 minutes at 37°C in a rotating incubator. Cells were centrifuged at 300 g for 5 minutes and supernatant aspirated. 5 000 cells were reseeded at equal densities in 100% Matrigel and supplemented with feeding media.

### Statistical analysis

For ADM 3D-culture quantification, we used two-way ANOVA follow by Dunnett’s correction. For *in vivo* experiment, when two conditions were compared, a two-tailed unpaired Mann Whitney test was used. For more than two conditions comparison, one-way ANOVAs follow by Tukey’s correction were performed.

## Study Approval

All experiments on mice were approved by the Animal Care Committee at the University of Western Ontario (Protocol #2020-057).

## Author contributions

## Acknowledgments

CIHR, CRS, Baker Centre

See if we can get Gabe to read this

