## Supplementary Figures for "EZH2 deletion does not impact acinar cell regeneration but restricts progression to pancreatic cancer in mice"

### Slide 1
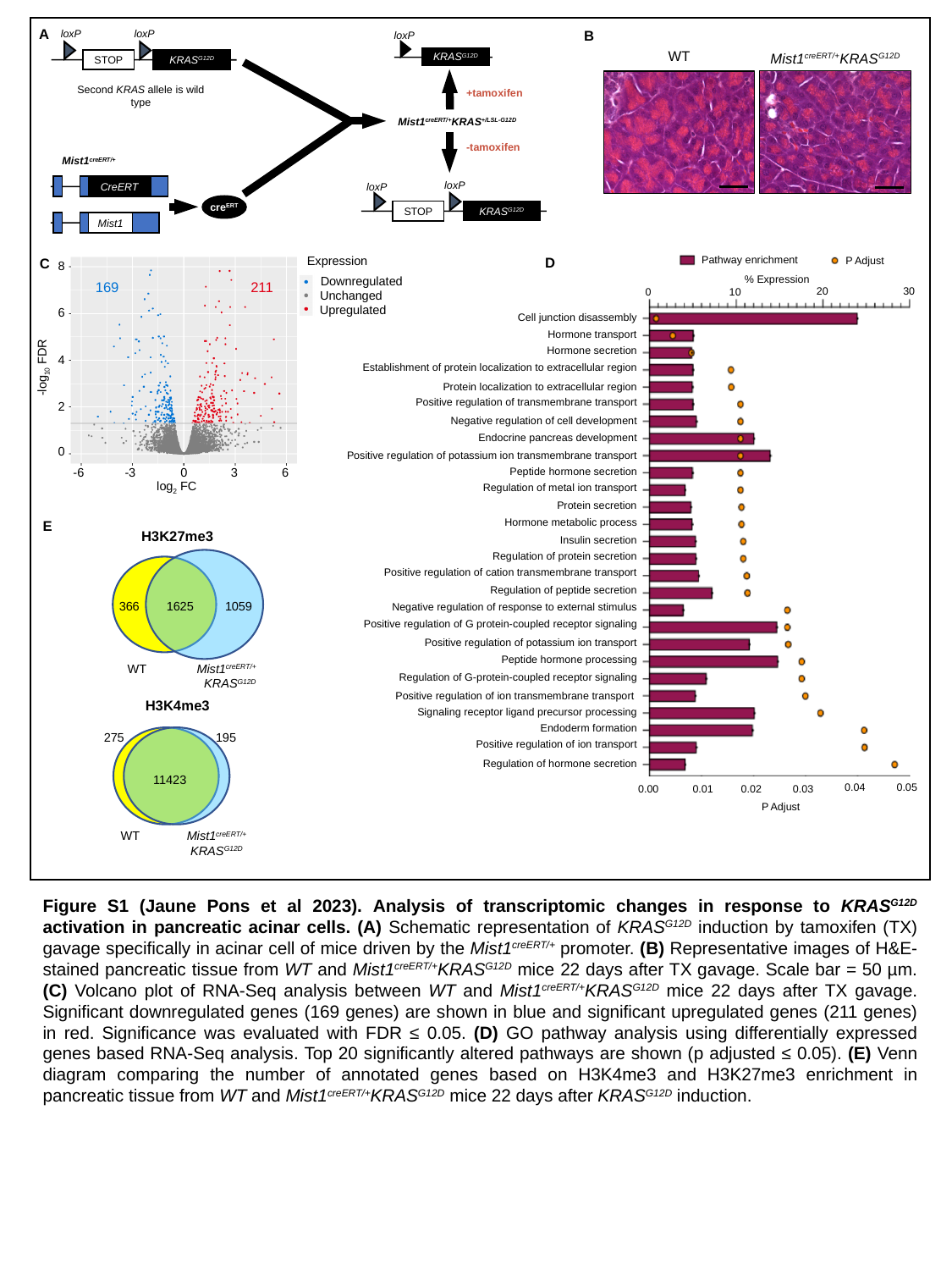

A
B
loxP
loxP
loxP
KRAS^G12D^
KRAS^G12D^
STOP
Second KRAS allele is wild type
+tamoxifen
Mist1^creERT/+^KRAS^+/LSL-G12D^
-tamoxifen
Mist1^creERT/+^
loxP
loxP
CreERT
KRAS^G12D^
STOP
cre^ERT^
Mist1
WT
Mist1^creERT/+^KRAS^G12D^
Pathway enrichment
D
P Adjust
% Expression
30
20
10
0
Cell junction disassembly
Hormone transport
Hormone secretion
Establishment of protein localization to extracellular region
Protein localization to extracellular region
Positive regulation of transmembrane transport
Negative regulation of cell development
Endocrine pancreas development
Positive regulation of potassium ion transmembrane transport
Peptide hormone secretion
Regulation of metal ion transport
Protein secretion
Hormone metabolic process
Insulin secretion
Regulation of protein secretion
Positive regulation of cation transmembrane transport
Regulation of peptide secretion
Negative regulation of response to external stimulus
Positive regulation of G protein-coupled receptor signaling
Positive regulation of potassium ion transport
Peptide hormone processing
Regulation of G-protein-coupled receptor signaling
Positive regulation of ion transmembrane transport
Signaling receptor ligand precursor processing
Endoderm formation
Positive regulation of ion transport
Regulation of hormone secretion
0.04
0.05
0.00
0.01
0.02
0.03
P Adjust
Expression
8
Downregulated
169
211
Unchanged
Upregulated
6
4
-log_10_ FDR
2
0
-6
-3
0
3
6
log_2_ FC
C
E
H3K27me3
366
1625
1059
Mist1^creERT/+^ KRAS^G12D^
WT
H3K4me3
275
195
11423
WT
Mist1^creERT/+^ KRAS^G12D^
Figure S1 (Jaune Pons et al 2023). Analysis of transcriptomic changes in response to KRAS^G12D^ activation in pancreatic acinar cells. (A) Schematic representation of KRAS^G12D^ induction by tamoxifen (TX) gavage specifically in acinar cell of mice driven by the Mist1^creERT/+^ promoter. (B) Representative images of H&E-stained pancreatic tissue from WT and Mist1^creERT/+^KRAS^G12D^ mice 22 days after TX gavage. Scale bar = 50 µm. (C) Volcano plot of RNA-Seq analysis between WT and Mist1^creERT/+^KRAS^G12D^ mice 22 days after TX gavage. Significant downregulated genes (169 genes) are shown in blue and significant upregulated genes (211 genes) in red. Significance was evaluated with FDR ≤ 0.05. (D) GO pathway analysis using differentially expressed genes based RNA-Seq analysis. Top 20 significantly altered pathways are shown (p adjusted ≤ 0.05). (E) Venn diagram comparing the number of annotated genes based on H3K4me3 and H3K27me3 enrichment in pancreatic tissue from WT and Mist1^creERT/+^KRAS^G12D^ mice 22 days after KRAS^G12D^ induction.

### Slide 2
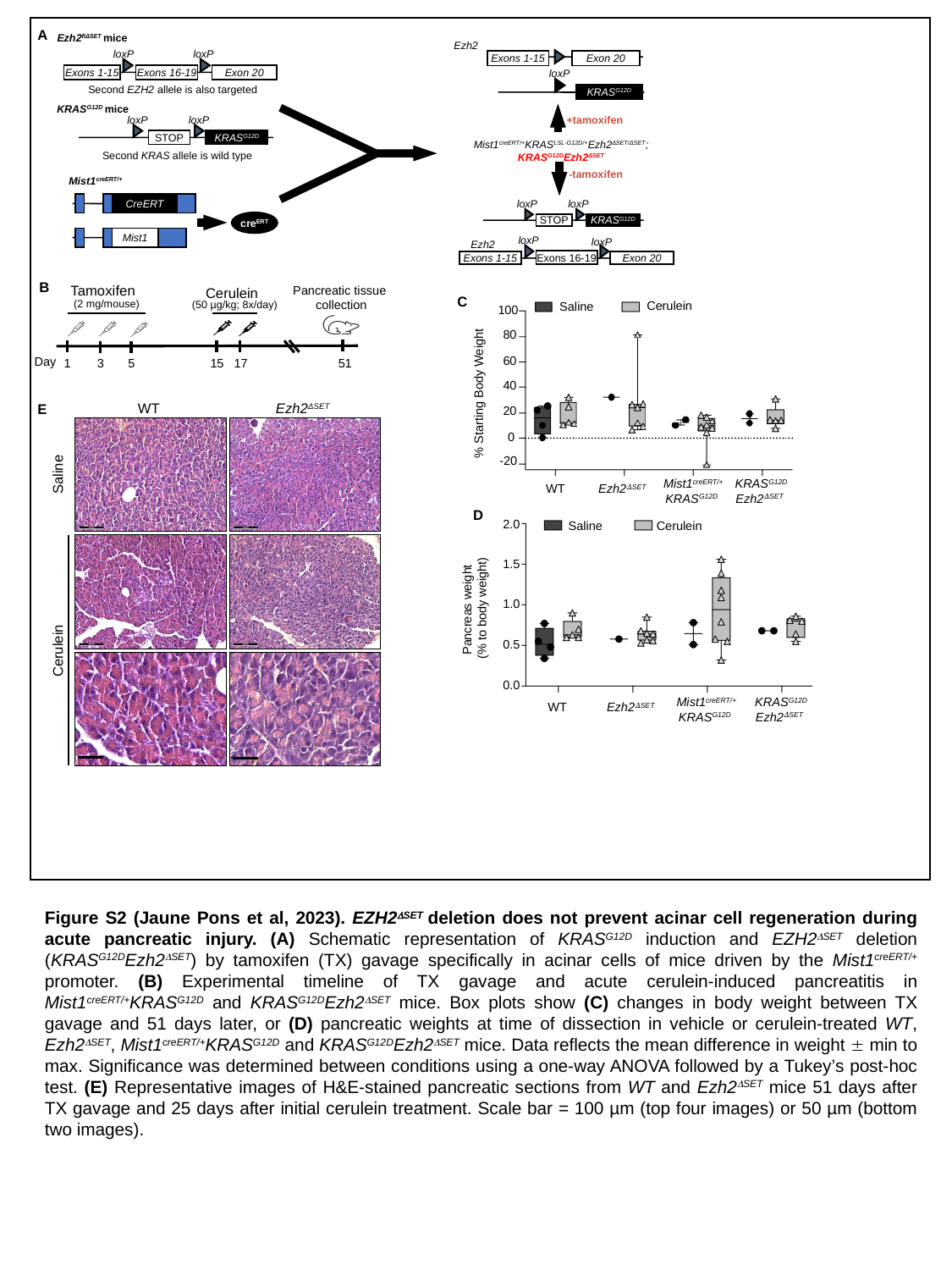

A
Ezh2^flΔSET^ mice
Ezh2
loxP
loxP
Exons 1-15
Exon 20
Exons 1-15
Exons 16-19
Exon 20
loxP
Second EZH2 allele is also targeted
KRAS^G12D^
KRAS^G12D^ mice
loxP
loxP
+tamoxifen
KRAS^G12D^
STOP
Mist1^creERT/+^KRAS^LSL-G12D/+^Ezh2^ΔSET/ΔSET^; KRAS^G12D^Ezh2^ΔSET^
Second KRAS allele is wild type
-tamoxifen
Mist1^creERT/+^
loxP
loxP
CreERT
KRAS^G12D^
STOP
cre^ERT^
loxP
Mist1
loxP
Ezh2
Exons 16-19
Exons 1-15
Exon 20
B
Tamoxifen
Cerulein
(2 mg/mouse)
(50 µg/kg; 8x/day)
Day
1
3
5
15
17
51
Pancreatic tissue
collection
C
Cerulein
Saline
100
80
60
40
% Starting Body Weight
20
0
-20
Mist1^creERT/+^
KRAS^G12D^
KRAS^G12D^
Ezh2^𝛥SET^
WT
Ezh2^𝛥SET^
WT
Ezh2^ΔSET^
E
Saline
D
2.0
Saline
Cerulein
1.5
Pancreas weight
(% to body weight)
1.0
0.5
0.0
Mist1^creERT/+^
KRAS^G12D^
KRAS^G12D^
Ezh2^𝛥SET^
WT
Ezh2^𝛥SET^
Cerulein
Figure S2 (Jaune Pons et al, 2023). EZH2^ΔSET^ deletion does not prevent acinar cell regeneration during acute pancreatic injury. (A) Schematic representation of KRAS^G12D^ induction and EZH2^ΔSET^ deletion (KRAS^G12D^Ezh2^ΔSET^) by tamoxifen (TX) gavage specifically in acinar cells of mice driven by the Mist1^creERT/+^ promoter. (B) Experimental timeline of TX gavage and acute cerulein-induced pancreatitis in Mist1^creERT/+^KRAS^G12D^ and KRAS^G12D^Ezh2^ΔSET^ mice. Box plots show (C) changes in body weight between TX gavage and 51 days later, or (D) pancreatic weights at time of dissection in vehicle or cerulein-treated WT, Ezh2^ΔSET^, Mist1^creERT/+^KRAS^G12D^ and KRAS^G12D^Ezh2^ΔSET^ mice. Data reflects the mean difference in weight ± min to max. Significance was determined between conditions using a one-way ANOVA followed by a Tukey’s post-hoc test. (E) Representative images of H&E-stained pancreatic sections from WT and Ezh2^ΔSET^ mice 51 days after TX gavage and 25 days after initial cerulein treatment. Scale bar = 100 µm (top four images) or 50 µm (bottom two images).

### Slide 3
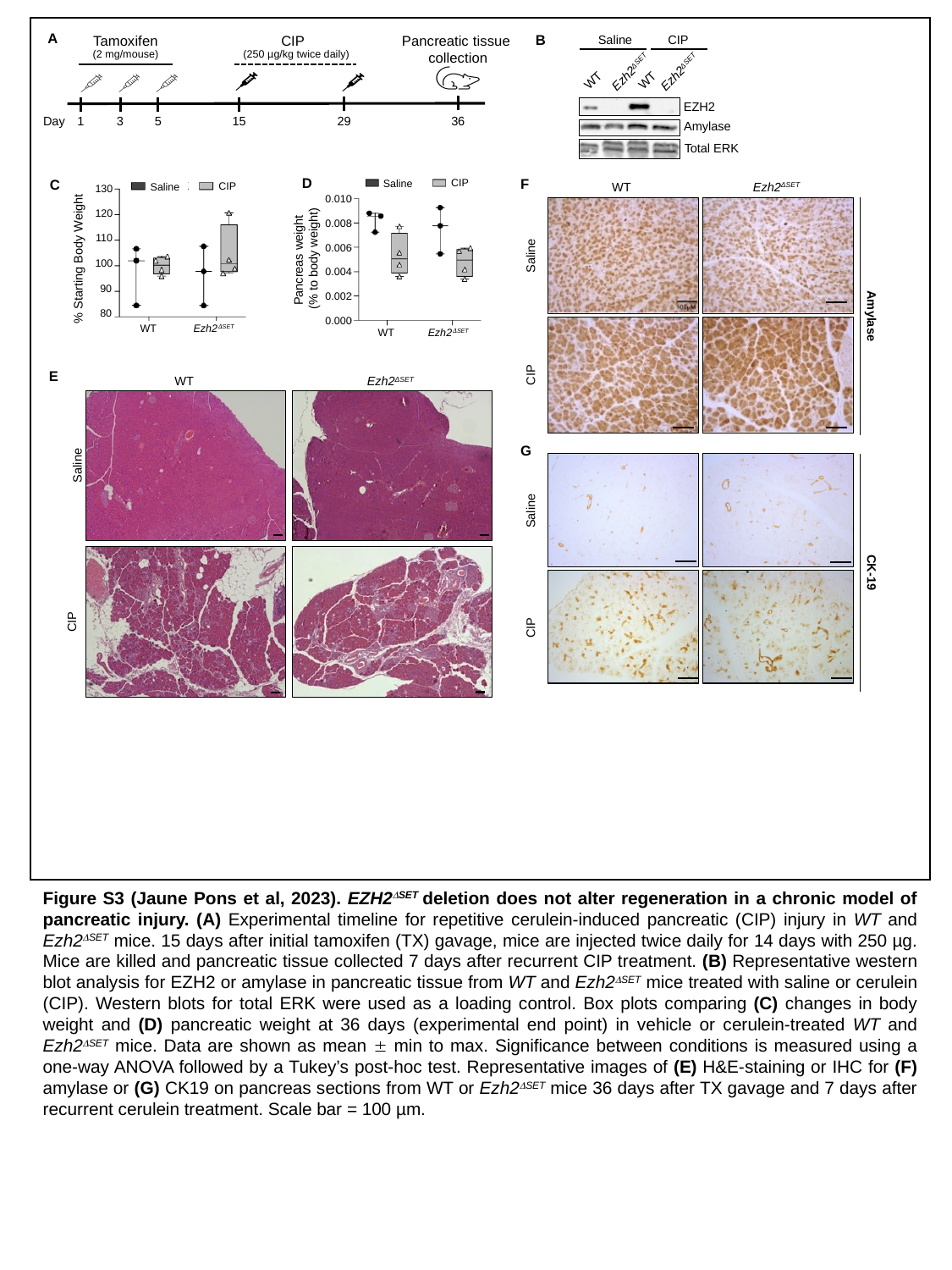

A
B
Pancreatic tissue
collection
Tamoxifen
CIP
(2 mg/mouse)
(250 µg/kg twice daily)
Day
1
3
5
15
29
36
Saline
CIP
Ezh2^ΔSET^
Ezh2^ΔSET^
WT
WT
EZH2
Amylase
Total ERK
D
F
CIP
Saline
0.010
0.008
0.006
Pancreas weight
(% to body weight)
0.004
0.002
0.000
Ezh2^𝛥SET^
WT
C
WT
Ezh2^ΔSET^
CIP
Saline
130
120
110
% Starting Body Weight
100
90
80
WT
Ezh2^𝛥SET^
Saline
Amylase
CIP
E
WT
Ezh2^ΔSET^
G
Saline
Saline
CK-19
CIP
CIP
Figure S3 (Jaune Pons et al, 2023). EZH2^ΔSET^ deletion does not alter regeneration in a chronic model of pancreatic injury. (A) Experimental timeline for repetitive cerulein-induced pancreatic (CIP) injury in WT and Ezh2^ΔSET^ mice. 15 days after initial tamoxifen (TX) gavage, mice are injected twice daily for 14 days with 250 µg. Mice are killed and pancreatic tissue collected 7 days after recurrent CIP treatment. (B) Representative western blot analysis for EZH2 or amylase in pancreatic tissue from WT and Ezh2^ΔSET^ mice treated with saline or cerulein (CIP). Western blots for total ERK were used as a loading control. Box plots comparing (C) changes in body weight and (D) pancreatic weight at 36 days (experimental end point) in vehicle or cerulein-treated WT and Ezh2^ΔSET^ mice. Data are shown as mean ± min to max. Significance between conditions is measured using a one-way ANOVA followed by a Tukey’s post-hoc test. Representative images of (E) H&E-staining or IHC for (F) amylase or (G) CK19 on pancreas sections from WT or Ezh2^ΔSET^ mice 36 days after TX gavage and 7 days after recurrent cerulein treatment. Scale bar = 100 µm.

### Slide 4
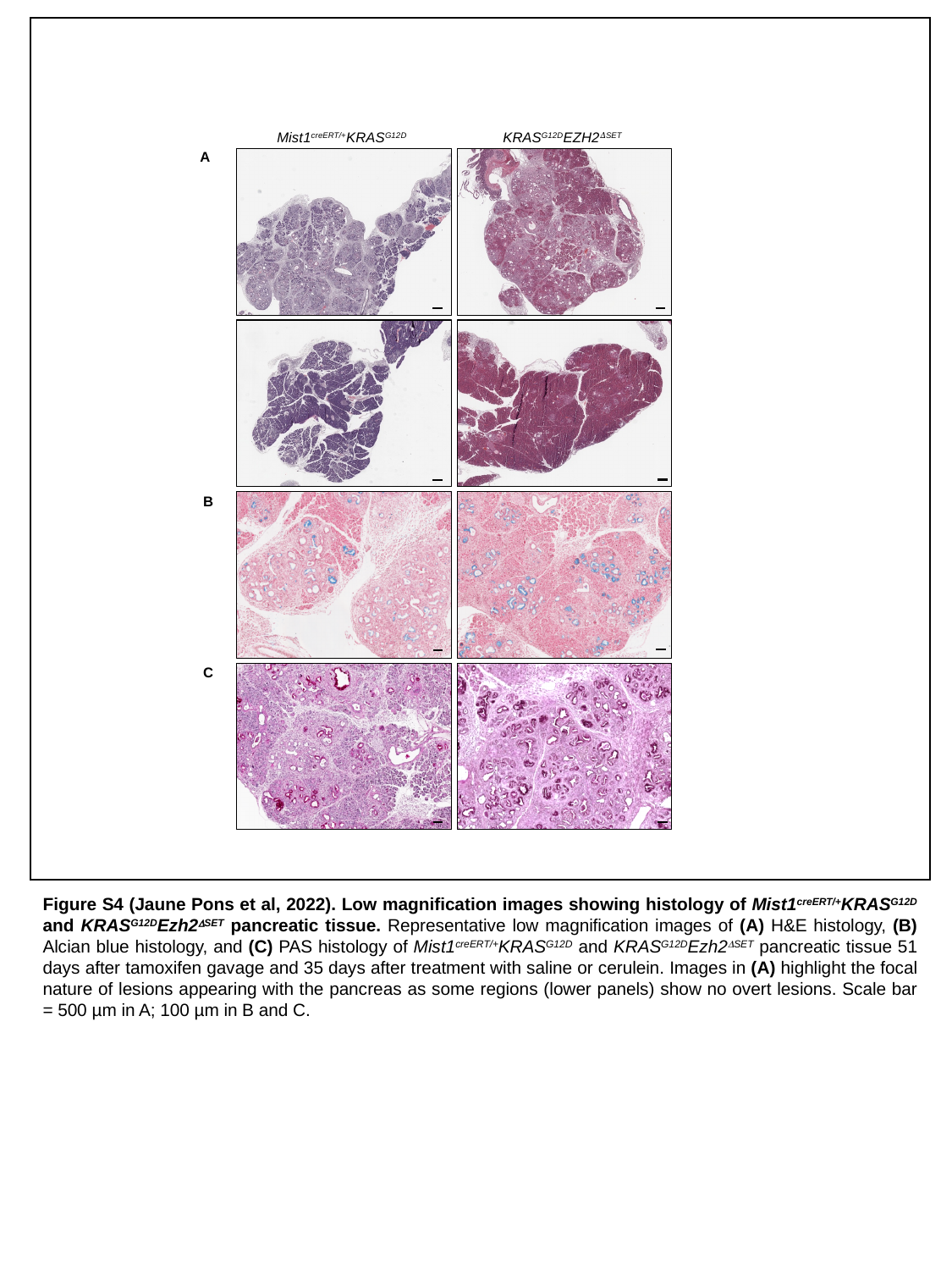

Mist1^creERT/+^KRAS^G12D^
KRAS^G12D^EZH2^𝛥SET^
A
B
C
Figure S4 (Jaune Pons et al, 2022). Low magnification images showing histology of Mist1^creERT/+^KRAS^G12D^ and KRAS^G12D^Ezh2^ΔSET^ pancreatic tissue. Representative low magnification images of (A) H&E histology, (B) Alcian blue histology, and (C) PAS histology of Mist1^creERT/+^KRAS^G12D^ and KRAS^G12D^Ezh2^ΔSET^ pancreatic tissue 51 days after tamoxifen gavage and 35 days after treatment with saline or cerulein. Images in (A) highlight the focal nature of lesions appearing with the pancreas as some regions (lower panels) show no overt lesions. Scale bar = 500 µm in A; 100 µm in B and C.

### Slide 5
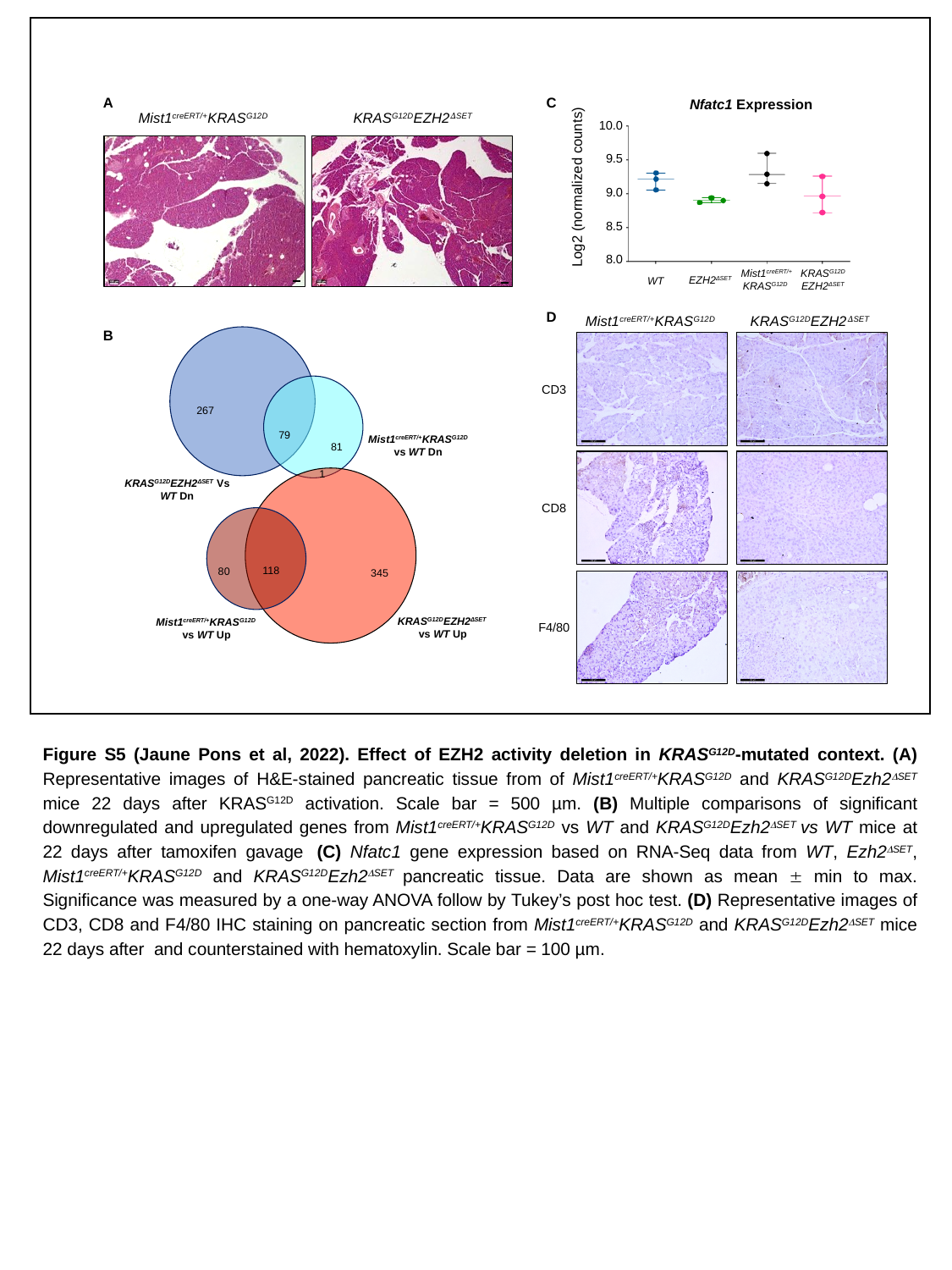

A
C
Nfatc1 Expression
10.0
9.5
Log2 (normalized counts)
9.0
8.5
8.0
Mist1^creERT/+^ KRAS^G12D^
KRAS^G12D^
EZH2^ΔSET^
EZH2^ΔSET^
WT
Mist1^creERT/+^KRAS^G12D^
KRAS^G12D^EZH2^𝛥SET^
D
Mist1^creERT/+^KRAS^G12D^
KRAS^G12D^EZH2^𝛥SET^
B
267
79
Mist1^creERT/+^KRAS^G12D^ vs WT Dn
81
1
KRAS^G12D^EZH2^ΔSET^ Vs WT Dn
118
80
345
KRAS^G12D^EZH2^ΔSET^ vs WT Up
Mist1^creERT/+^KRAS^G12D^ vs WT Up
CD3
CD8
F4/80
Figure S5 (Jaune Pons et al, 2022). Effect of EZH2 activity deletion in KRAS^G12D^-mutated context. (A) Representative images of H&E-stained pancreatic tissue from of Mist1^creERT/+^KRAS^G12D^ and KRAS^G12D^Ezh2^ΔSET^ mice 22 days after KRAS^G12D^ activation. Scale bar = 500 µm. (B) Multiple comparisons of significant downregulated and upregulated genes from Mist1^creERT/+^KRAS^G12D^ vs WT and KRAS^G12D^Ezh2^ΔSET^ vs WT mice at 22 days after tamoxifen gavage  (C) Nfatc1 gene expression based on RNA-Seq data from WT, Ezh2^ΔSET^, Mist1^creERT/+^KRAS^G12D^ and KRAS^G12D^Ezh2^ΔSET^ pancreatic tissue. Data are shown as mean ± min to max. Significance was measured by a one-way ANOVA follow by Tukey’s post hoc test. (D) Representative images of CD3, CD8 and F4/80 IHC staining on pancreatic section from Mist1^creERT/+^KRAS^G12D^ and KRAS^G12D^Ezh2^ΔSET^ mice 22 days after and counterstained with hematoxylin. Scale bar = 100 µm.

### Slide 6
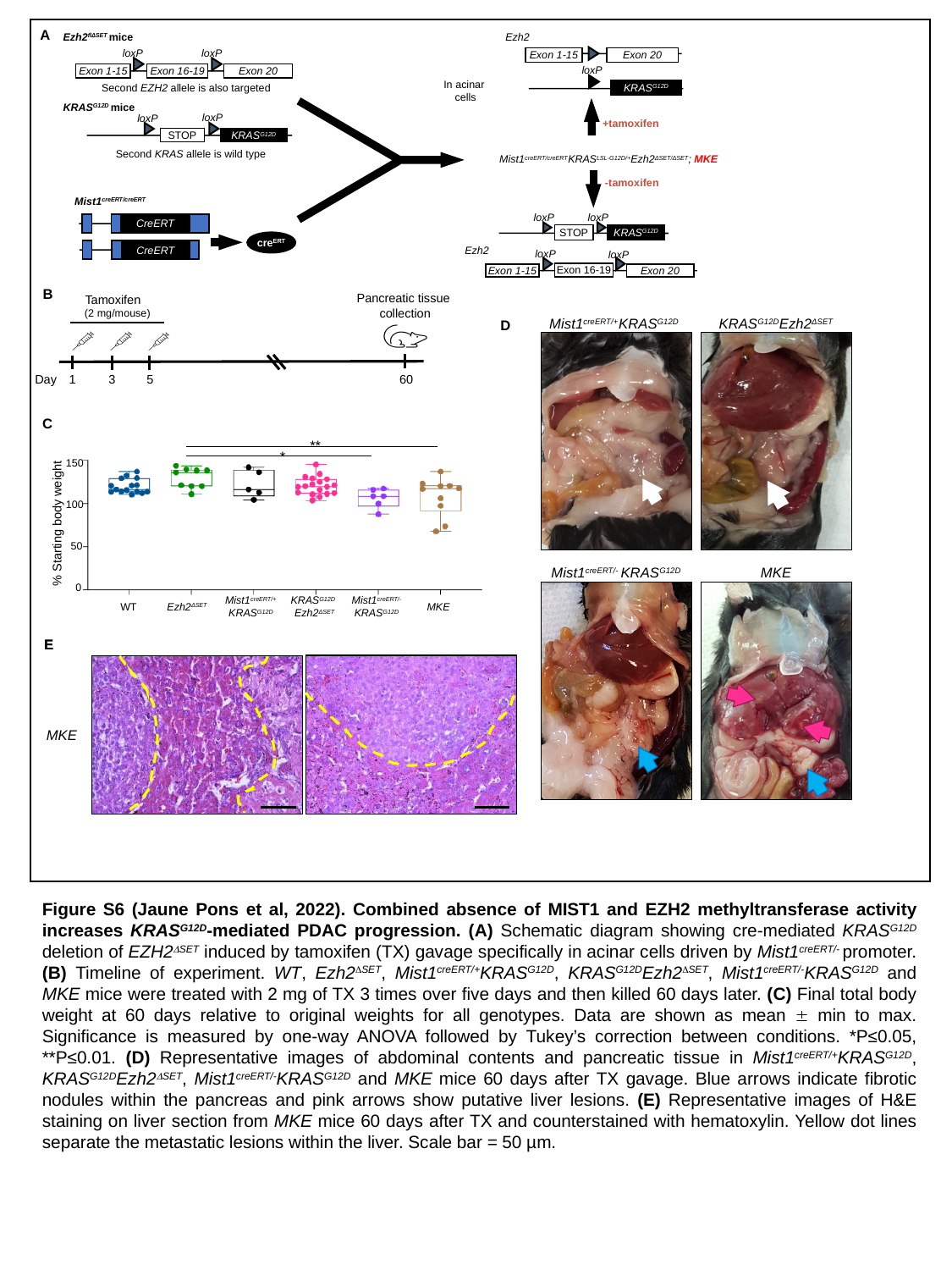

A
Ezh2^flΔSET^ mice
Ezh2
loxP
loxP
Exon 1-15
Exon 20
loxP
Exon 1-15
Exon 16-19
Exon 20
Second EZH2 allele is also targeted
KRAS^G12D^
In acinar
cells
KRAS^G12D^ mice
loxP
loxP
+tamoxifen
KRAS^G12D^
STOP
Second KRAS allele is wild type
Mist1^creERT/creERT^KRAS^LSL-G12D/+^Ezh2^ΔSET/ΔSET^; MKE
-tamoxifen
Mist1^creERT/creERT^
loxP
loxP
CreERT
STOP
KRAS^G12D^
cre^ERT^
Ezh2
CreERT
loxP
loxP
Exon 16-19
Exon 1-15
Exon 20
B
Pancreatic tissue
collection
Tamoxifen
(2 mg/mouse)
Day
1
3
5
60
Mist1^creERT/+^KRAS^G12D^
KRAS^G12D^Ezh2^ΔSET^
D
C
**
*
150
100
% Starting body weight
50
0
Mist1^creERT/-^ KRAS^G12D^
Mist1^creERT/+^
KRAS^G12D^
KRAS^G12D^
Ezh2^ΔSET^
WT
Ezh2^ΔSET^
MKE
Mist1^creERT/-^ KRAS^G12D^
MKE
E
E
MKE
Figure S6 (Jaune Pons et al, 2022). Combined absence of MIST1 and EZH2 methyltransferase activity increases KRAS^G12D^-mediated PDAC progression. (A) Schematic diagram showing cre-mediated KRAS^G12D^ deletion of EZH2^ΔSET^ induced by tamoxifen (TX) gavage specifically in acinar cells driven by Mist1^creERT/-^ promoter. (B) Timeline of experiment. WT, Ezh2^ΔSET^, Mist1^creERT/+^KRAS^G12D^, KRAS^G12D^Ezh2^ΔSET^, Mist1^creERT/-^KRAS^G12D^ and MKE mice were treated with 2 mg of TX 3 times over five days and then killed 60 days later. (C) Final total body weight at 60 days relative to original weights for all genotypes. Data are shown as mean ± min to max. Significance is measured by one-way ANOVA followed by Tukey’s correction between conditions. *P≤0.05, **P≤0.01. (D) Representative images of abdominal contents and pancreatic tissue in Mist1^creERT/+^KRAS^G12D^, KRAS^G12D^Ezh2^ΔSET^, Mist1^creERT/-^KRAS^G12D^ and MKE mice 60 days after TX gavage. Blue arrows indicate fibrotic nodules within the pancreas and pink arrows show putative liver lesions. (E) Representative images of H&E staining on liver section from MKE mice 60 days after TX and counterstained with hematoxylin. Yellow dot lines separate the metastatic lesions within the liver. Scale bar = 50 µm.

### Slide 7
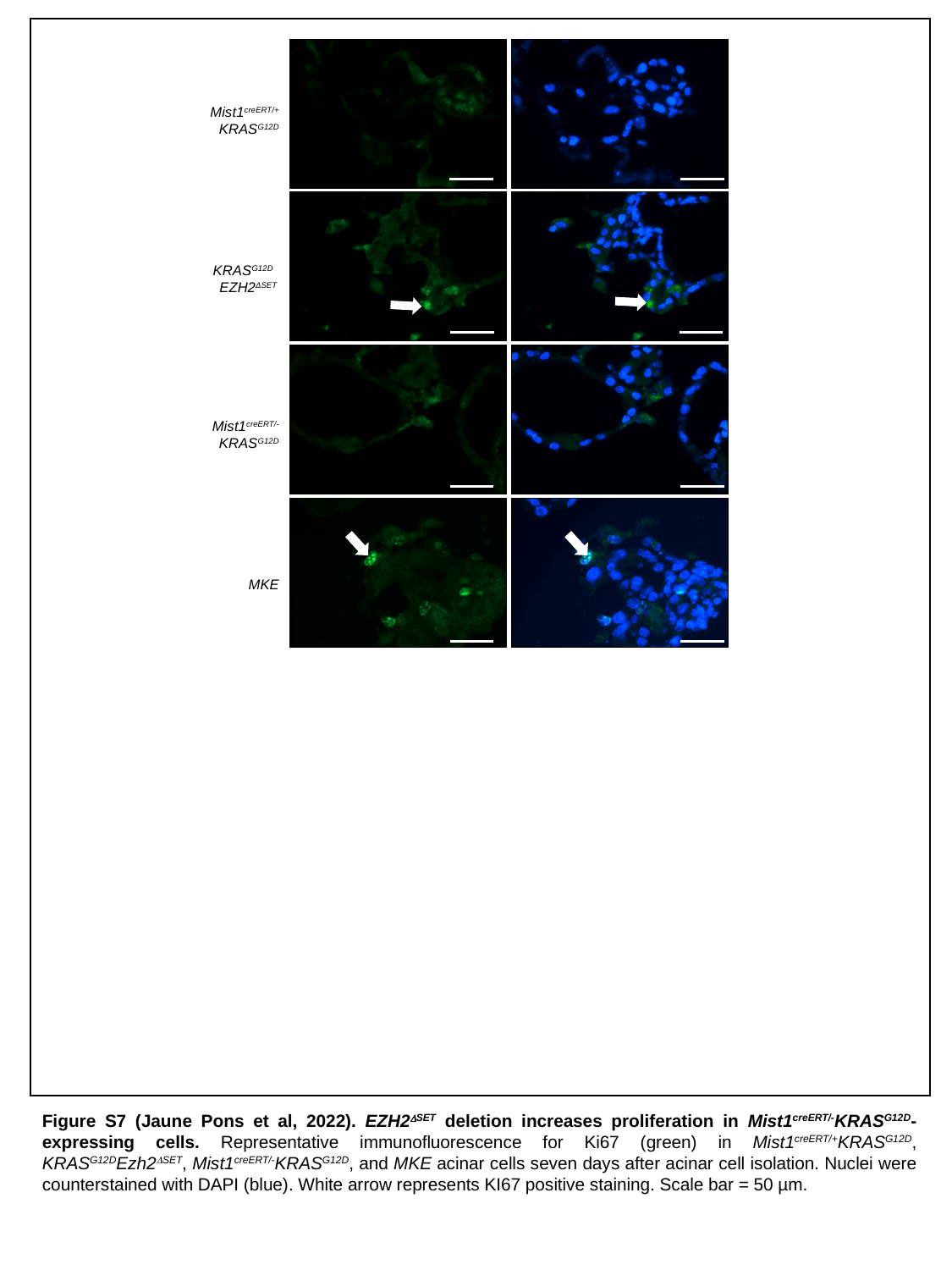

Mist1^creERT/+^
KRAS^G12D^
KRAS^G12D^
EZH2^ΔSET^
Mist1^creERT/-^KRAS^G12D^
x
MKE
x
Figure S7 (Jaune Pons et al, 2022). EZH2^ΔSET^ deletion increases proliferation in Mist1^creERT/-^KRAS^G12D^-expressing cells. Representative immunofluorescence for Ki67 (green) in Mist1^creERT/+^KRAS^G12D^, KRAS^G12D^Ezh2^ΔSET^, Mist1^creERT/-^KRAS^G12D^, and MKE acinar cells seven days after acinar cell isolation. Nuclei were counterstained with DAPI (blue). White arrow represents KI67 positive staining. Scale bar = 50 µm.

### Slide 8
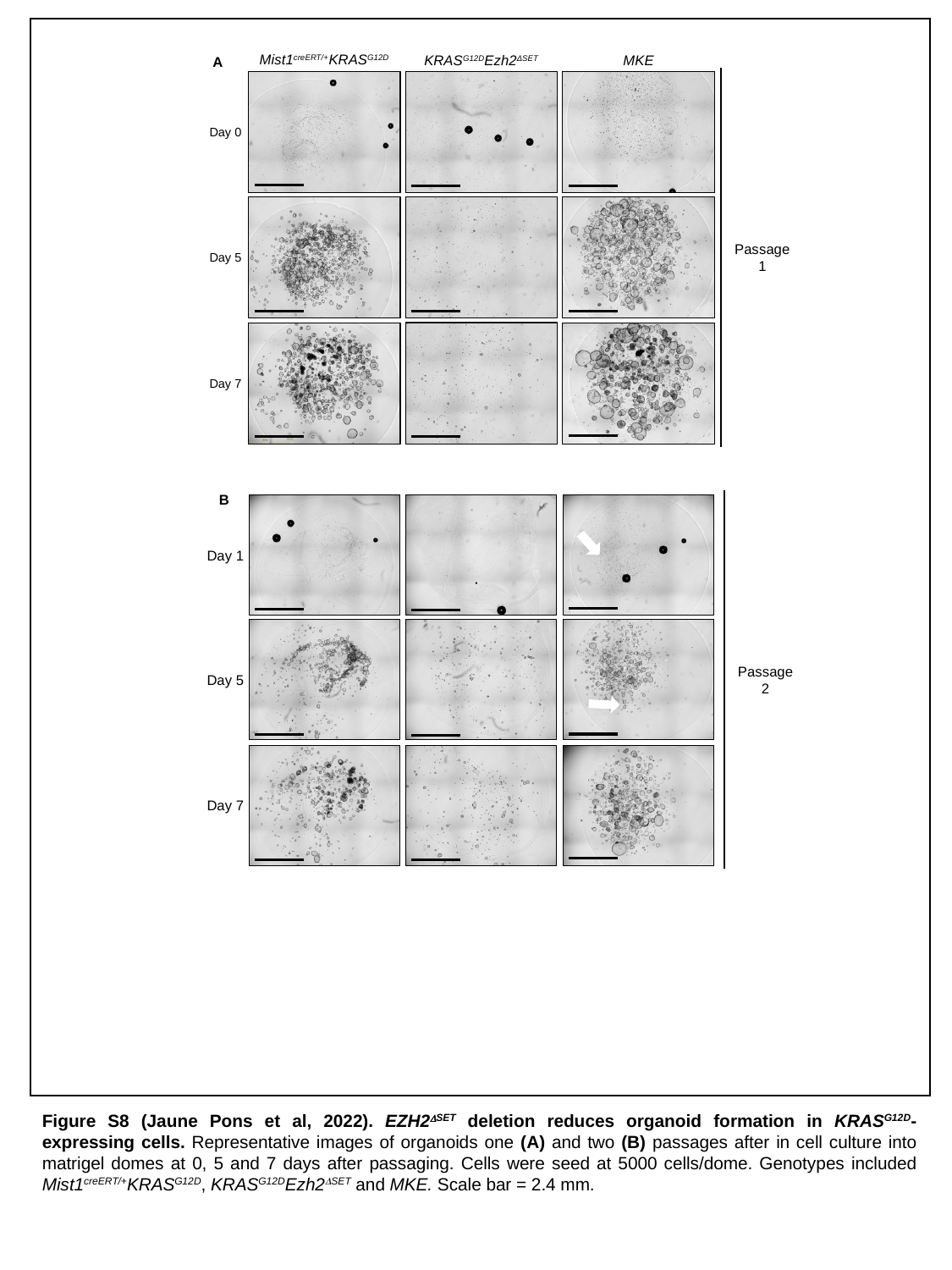

KRAS^G12D^Ezh2^ΔSET^
MKE
Mist1^creERT/+^KRAS^G12D^
A
Day 0
Passage 1
Day 5
Day 7
B
x
Day 1
Passage 2
Day 5
Day 7
Figure S8 (Jaune Pons et al, 2022). EZH2^ΔSET^ deletion reduces organoid formation in KRAS^G12D^-expressing cells. Representative images of organoids one (A) and two (B) passages after in cell culture into matrigel domes at 0, 5 and 7 days after passaging. Cells were seed at 5000 cells/dome. Genotypes included Mist1^creERT/+^KRAS^G12D^, KRAS^G12D^Ezh2^ΔSET^ and MKE. Scale bar = 2.4 mm.
