## Supplementary Table S5 for "EZH2 deletion does not impact acinar cell regeneration but restricts progression to pancreatic cancer in mice"

**Supplemental Table S2. RT-qPCR primers**

| **Gene** | **Forward** | **Reverse** |
| --- | --- | --- |
| *Mrpl1* | 5'-TTGGATATGCCAAGTGACCA2-3’ | 5'-GCTTCTGCCGTTTGAGTTTC-3’ |
| *Ptgs2* | 5'-AGGACTCTGCTCACGAAGGA-3’ | 5'-TCATACATTCCCCACGGTTT-3’ |
