## Supplementary Table S4 for "EZH2 deletion does not impact acinar cell regeneration but restricts progression to pancreatic cancer in mice"

**Supplemental Table S4. Significant gene set enrichment GO pathways enriched in *MKE* *vs* *Mist1^creERT/-^KRAS^G12D^* in RNA-Sequencing**

| **ID** | **Description** | **Enrichment Score** | **NES** | **P adjust** | **Core enrichment** |
| --- | --- | --- | --- | --- | --- |
| GO:0006334 | nucleosome assembly | -0,773677383 | -2.372314668 | 1.32E-06 | H4c14/H4c12/H4c8/H1f4/H3c4/H4f16/H3c6/H1f3/H1f5/H2bc3/H4c3/H3c1/H3c7/H4c1/H2bc7/H3c3/H3c2/H4c4/H4c18/H4c6/H1f1 |
| GO:0046034 | ATP metabolic process | 0,402367449 | 2.34745849 | 1.05E-05 | Atp5k/ATP8/Cox6a1/Atp5j2/Ddit4/Atp5e/Trem2/Cox5b/COX2/Cox4i2/Ier3/Cox7a1/Antkmt/Atp5g3/Chchd10/Gck/ND4L/Tspo/Ins2/ATP6/Ndufb6/Uqcrq/Pgam1/Cyc1/Mif/Eif6/Galk1/COX3/Ndufc2/Gpd1/Uqcr10/Eno3/Ndufv3/Uqcc2/Atp5d/Nupr1/Ldhd/Sdhc/Ndufa8/Khk/Ndufb9/Ndufb8/Ndufa12/Slc25a33/Pink1/Ak1/CYTB/Atp1a2/Aldoc/Eno2/Cox5a/Guk1/Uqcrfs1/Gadd45gip1/Sdhd/Coq7/Atp5g1/Atp5j/Uqcc3/Ndufa7/Ndufs8/Coq9/Park7/Uqcrh/Uqcrc1/Cox4i1/Cox7a2/Flcn/Bad/Slc25a23/Pgk1/Ak2/Aldoa/Ndufv1/Bcl2l1/Taz/Iscu |
| GO:0032963 | collagen metabolic process | 0,573375375 | 2.792596572 | 1.20E-05 | Retn/Mmp23/Scx/Mmp2/Adam15/Ccn2/Rgcc/Mmp9/Mfap4/Serpinh1/Rcn3/Id1/Eng/Prdx5/Cygb/P3h4/Mmp28/Mmp11/Col1a2/Mrc2/Col1a1/Adamts2/P3h3/Ctsk/Vim/Wnt4/Pdgfb/Mmp19/Rapgef3/Mmp14/Col5a1/Tnxb/Bmp4/Tgfb3/Smpd3/Mmp15/Ppard |
| GO:0042060 | wound healing | 0,374682357 | 2.221390754 | 1.20E-05 | Treml1/Vkorc1/Timp1/Insl3/F2rl3/Anxa8/Cldn4/Bnc1/Wfdc1/Alox15/Gpx1/Aqp1/Tmeff2/Clec10a/Ccm2l/Tspan8/Pf4/Serping1/Fgf1/Entpd2/Ins2/Thbd/Anxa2/BC024139/Tnfrsf12a/Cd34/Cadm4/Apoe/Cav1/Smoc2/Ajuba/Eng/Bloc1s4/Selp/Fbln1/C1qtnf1/Cdkn1a/Anxa1/Proc/Cd151/Fkbp10/Cldn3/Col1a1/Cela2a/F3/Adrb1/Bloc1s3/Plau/Gata2/Wnt4/Pdgfb/Col3a1/Gas6/Erbb2/Acvrl1/Col5a1/Srf/Hmox1/Gnas/Axl/Evpl/Wnt5a/Serpine2/Vegfb/Actg1/Plet1/Ppard |
| GO:0048568 | embryonic organ development | 0,340207194 | 2.027610144 | 1.92E-05 | Xist/Tbx4/Rnf112/Tcf21/Gjb5/Snai1/Nkx3-2/Ccdc103/Cebpb/Id3/Twist1/Sox18/Crb2/Gli1/Hand2/Mdfi/Wnt7b/Wnt11/Fzd2/Mfap2/Wnt2/Pbx4/Lif/Gas1/Adm/Col13a1/Vash1/Rarres2/Junb/Pkdcc/Wnt9a/Hoxb6/Cebpa/Erf/Mfap5/Eng/Rbp4/Osr1/Tbx2/Hoxb7/Cdkn1c/Epha2/Epn1/Slc39a3/Cdk20/Smo/Naglu/Plcd3/Sox17/Gja5/Gata2/Edn1/Socs3/Pdgfb/Aldh1a2/Mmp14/Krt8/Sod1/Chst11/Srf/Gnas/Bmp4 |
| GO:0034728 | nucleosome organization | -0,66781337 | -2.178966457 | 2.98E-05 | H4c14/H4c12/H4c8/H1f4/H3c4/H4f16/H3c6/H1f3/H1f5/H2bc3/H4c3/H3c1/H3c7/H4c1/H2bc7/H3c3/H3c2/H4c4/H4c18/H4c6/H1f1 |
| GO:0006119 | oxidative phosphorylation | 0,495136371 | 2.586992818 | 2.98E-05 | Cox6a1/Atp5j2/Cox5b/COX2/Cox4i2/Cox7a1/Antkmt/Chchd10/ND4L/ATP6/Ndufb6/Uqcrq/Cyc1/COX3/Ndufc2/Uqcr10/Ndufv3/Uqcc2/Atp5d/Nupr1/Sdhc/Ndufa8/Ndufb9/Ndufb8/Ndufa12/Slc25a33/Pink1/CYTB/Cox5a/Uqcrfs1/Gadd45gip1/Sdhd/Coq7/Atp5j/Uqcc3/Ndufa7/Ndufs8/Coq9/Park7/Uqcrh/Uqcrc1/Cox4i1/Cox7a2/Slc25a23/Pgk1/Ndufv1/Taz/Iscu/Ppif/Ndufs6/Shmt2/Atp5pb/Dnajc15 |
| GO:0006790 | sulfur compound metabolic process | 0,394039611 | 2.210970482 | 2.98E-05 | Gstp2/Pdk4/Gstm3/Gpx1/Dgat2/Bola2/Slc27a3/Ggt6/Chpf/Chst14/Chac1/Ciapin1/Chst7/Nat8/Gpx3/Gpx4/Cdo1/Mpst/Gamt/Tst/Dpep1/Hmgcl/Gstm2/Iba57/Dcn/Sod1/Chst11/Gstm6/Bgn/Fasn/Gnmt/Acaa2/Mvd/Ppcs/Pemt/Spock2/Dgat1/Ciao2b/Mlycd/Ggt5/Sulf1/Phgdh/Hscb/Mpc2/Gsta4/Apip/Comt/Gstt1/Pmvk/Mat1a/Acot7/Chst12/Gstk1/Glrx3/Park7/Isca2/Gstp1/Gstm1/Mgst1/Idua/Oplah/Hmgn5/Csad/Tpst2/Mthfr/Iscu/Acot8/Xylt2/Suox/Tpst1/Nubp2/Pcyox1l |
| GO:0006335 | DNA replication-dependent nucleosome assembly | -0,847802694 | -2.1638095 | 2.98E-05 | H4c14/H4c12/H4c8/H3c4/H4f16/H3c6/H4c3/H3c1/H4c1/H3c2/H4c4/H4c18/H4c6 |
| GO:0034723 | DNA replication-dependent nucleosome organization | -0,847802694 | -2.1638095 | 2.98E-05 | H4c14/H4c12/H4c8/H3c4/H4f16/H3c6/H4c3/H3c1/H4c1/H3c2/H4c4/H4c18/H4c6 |
| GO:0072593 | reactive oxygen species metabolic process | 0,429251086 | 2.406419873 | 3.44E-05 | Hbb-bt/Cryab/Ptgis/Pdk4/Hba-a1/Ddit4/Sod3/COX2/Gpx1/Ndufaf2/Spr/Ier3/Ngfr/Romo1/Hbb-b1/Tspo/Nos3/Ccn2/Ins2/Ndufa13/Ddah2/Cd34/Klf2/Cav1/Sesn1/Fbln5/Cyba/Prdx5/Gpx3/Agtr1a/Eif6/Cdkn1a |
| GO:0065004 | protein-DNA complex assembly | -0,638596948 | -2.12094844 | 3.64E-05 | H4c14/Mcmdc2/H4c12/H4c8/H1f4/H3c4/H4f16/H3c6/H1f3/H1f5/H2bc3/H4c3/H3c1/H3c7/H4c1/H2bc7/H3c3/H3c2/H4c4/H4c18/H4c6/H1f1 |
| GO:1901342 | regulation of vasculature development | 0,330764702 | 2.027266523 | 9.09E-05 | Sfrp2/Ptgis/Pgf/Rhob/Ccl24/Aqp1/Glul/Ngfr/Angpt4/Ecm1/Lif/Cma1/Hspb1/Adm/Wt1/Aplnr/Pf4/Vash1/Tmem100/Fgf1/Emc10/Nos3/Tnfrsf12a/Cd34/Rgcc/Serpinf1/Klf2/Mmp9/Smoc2/Sphk1/Adm2/Id1/Eng/Fbln5/Rnh1/Ramp2/Agtr1a/Anxa1/Sparc/Tie1/Anxa3/Ptn/Epha2/Epn1/F3/Card10/Bmper/Thbs2/Cela1/Grn/Ccl11/Gata2/Wnt4/Hdac7/Lrg1/Pdgfb/Rapgef3/Acvrl1/Cib1/Dcn/Reck/Hmox1/Mapk7/Bmp4/Naxe/Srpx2/Wnt5a/Klf4/Vegfb |
| GO:0030198 | extracellular matrix organization | 0,363061284 | 2.189499433 | 9.62E-05 | Eln/Sfrp2/Loxl1/Adamtsl5/Col16a1/Mmp23/Cma1/Vit/Papln/Wt1/Scx/Col13a1/Fbln2/Mmp2/Ccn2/Anxa2/Dpt/Rgcc/Mmp9/Pmp22/Col18a1/Mfap4/Aebp1/Serpinh1/Cav1/Smoc2/Eng/Fbln5/Ramp2/Fbln1/P3h4/Mmp28/Mmp11/Col5a3/Kazald1/Fkbp10/Col1a2/Col1a1/Adamts2/Olfml2b/Loxl2/Adamtsl4/Apbb1/Col3a1/Mmp19/Mmp14/Ccdc80/Col5a1/Reck/Vtn/Efemp2/Col5a2/Tnxb/Fgfr4/Phldb1/Spock2/Hpn/Smpd3/Lamb2/Mmp15/Loxl3/Sulf1/Cav2/Emilin1/Spint2/Crtap/Ddr1/Ltbp3/Adamtsl1/Lum/Adamts10/Nid1/Scara3/Col4a2/Lamb3/Lamb1/Idua/Antxr1/Ercc2/Col4a1/Qsox1/Flot1/Washc1/Crispld2/Prdx4/Aplp1/Gfod2/Col14a1/Colgalt1/Col15a1/Tnfrsf1a |
| GO:0043062 | extracellular structure organization | 0,363061284 | 2.189499433 | 9.62E-05 | Eln/Sfrp2/Loxl1/Adamtsl5/Col16a1/Mmp23/Cma1/Vit/Papln/Wt1/Scx/Col13a1/Fbln2/Mmp2/Ccn2/Anxa2/Dpt/Rgcc/Mmp9/Pmp22/Col18a1/Mfap4/Aebp1/Serpinh1/Cav1/Smoc2/Eng/Fbln5/Ramp2/Fbln1/P3h4/Mmp28/Mmp11/Col5a3/Kazald1/Fkbp10/Col1a2/Col1a1/Adamts2/Olfml2b/Loxl2/Adamtsl4/Apbb1/Col3a1/Mmp19/Mmp14/Ccdc80/Col5a1/Reck/Vtn/Efemp2/Col5a2/Tnxb/Fgfr4/Phldb1/Spock2/Hpn/Smpd3/Lamb2/Mmp15/Loxl3/Sulf1/Cav2/Emilin1/Spint2/Crtap/Ddr1/Ltbp3/Adamtsl1/Lum/Adamts10/Nid1/Scara3/Col4a2/Lamb3/Lamb1/Idua/Antxr1/Ercc2/Col4a1/Qsox1/Flot1/Washc1/Crispld2/Prdx4/Aplp1/Gfod2/Col14a1/Colgalt1/Col15a1/Tnfrsf1a |
| GO:0045333 | cellular respiration | 0,432139618 | 2.280978875 | 0.000111799 | Cox6a1/Cox5b/COX2/Cox4i2/Ndufaf2/Sdhaf4/Bax/ND4L/Nfatc4/Ndufb6/Uqcrq/Cyc1/Ndufs7/Cisd1/COX3/Bloc1s1/Ndufc2/Ndufa5/Gpd1/Uqcr10/Ndufv3/Atp5d/Chchd4/Sdhc/Ndufa8/Ndufb9/Slc25a22/Ndufb8/Ndufa12/Pink1/ND1/CYTB/Nop53/Sirt3/Cox5a/Uqcrfs1/Sdhd/Coq7/Uqcc3/Ndufa7/Ndufs8/Sdhb/Bnip3/Coq9/Park7/Uqcrh/Uqcrc1/Coq10a/Cox4i1/Flcn/Slc25a23/Ndufv1/Taz/Iscu/Ndufs6/Oxa1l/Mdh2/Shmt2/Dnajc15/Mdh1/Ndufs4/Dguok/ND2 |
| GO:0050727 | regulation of inflammatory response | 0,373844815 | 2.160902072 | 0.000111799 | Hamp/Ptgis/Cebpb/C2cd4b/Trem2/Wfdc1/Ccn3/Gpx2/Ccl24/Alox15/Gpx1/Ier3/Trpv4/Il17d/Per1/Cma1/Ins2/Ctla2a/Serpinf1/Pmp22/Cebpa/Metrnl/Apoe/Sphk1/Mif/Anxa1/Proc/Gpx4/Rps19/Rhbdd3/Gprc5b/Ndufc2/Tff2/Aoc3/Fabp4/C1qtnf12/Grn/Ddt/Gps2/Zp3/Il17rc/Socs3/Alox5ap/Zfp36/Nupr1/Tgm2/Sod1/Psmb4/Ace/Fndc4/Tradd/Pycard/Wnt5a |
| GO:0031497 | chromatin assembly | -0,631636068 | -2.086959752 | 0.000177823 | H4c14/H4c12/H4c8/H1f4/H3c4/H4f16/H3c6/H1f3/H1f5/H2bc3/H4c3/H3c1/H3c7/H4c1/H2bc7/H3c3/H3c2/H4c4/H4c18/H4c6/H1f1 |
| GO:0033108 | mitochondrial respiratory chain complex assembly | 0,528836125 | 2.548299874 | 0.000177823 | Ndufa2/Sdhaf4/Ndufa11/Ndufa6/Cox14/Ndufaf8/Ndufb10/Ndufb7/Ndufa13/Ndufb11/Ndufb6/Ndufa1/Ndufa3/Ndufs7/Sdhaf1/Ndufc1/Ndufb3/Ndufc2/1190007I07Rik/Ndufa5/Uqcr10/Uqcc2/Ndufaf3/Chchd4/Ndufa8/Ndufb9/Ndufb8/Slc25a33 |
| GO:0042773 | ATP synthesis coupled electron transport | 0,567386154 | 2.504598357 | 0.000239037 | Cox6a1/Cox5b/COX2/Cox4i2/ND4L/Ndufb6/Uqcrq/Cyc1/COX3/Ndufc2/Uqcr10/Ndufv3/Sdhc/Ndufa8/Ndufb9/Ndufb8/Ndufa12/Pink1/CYTB/Cox5a/Uqcrfs1/Sdhd/Coq7/Uqcc3/Ndufa7/Ndufs8/Coq9/Park7/Uqcrh/Uqcrc1/Cox4i1/Ndufv1/Taz/Iscu/Ndufs6/Dnajc15/Dguok/ND2 |
| GO:0022900 | electron transport chain | 0,497550081 | 2.470326329 | 0.000258545 | Cox6a1/Cox5b/COX2/Cox4i2/Ndufaf2/ND4L/Ndufb6/Uqcrq/Cyc1/COX3/Ndufb3/Ndufc2/Ndufa5/Gpd1/Uqcr10/Ndufv3/Sdhc/Ndufa8/Ndufb9/Cyb561/Slc25a22/Ndufb8/Ndufa12/Pink1/CYTB/Cox5a/Uqcrfs1/Sdhd/Coq7/Uqcc3/Ndufa7/Ndufs8/Sdhb/Coq9/Park7/Uqcrh/Uqcrc1/Cox4i1/Ndufv1/Taz/Iscu |
| GO:0045765 | regulation of angiogenesis | 0,330628191 | 1.933643654 | 0.000291132 | Sfrp2/Ptgis/Pgf/Rhob/Ccl24/Aqp1/Glul/Ngfr/Angpt4/Ecm1/Lif/Cma1/Hspb1/Adm/Aplnr/Pf4/Vash1/Fgf1/Emc10/Nos3/Tnfrsf12a/Cd34/Rgcc/Serpinf1/Klf2/Mmp9/Smoc2/Sphk1/Adm2/Id1/Eng/Fbln5/Rnh1/Ramp2/Agtr1a/Anxa1/Sparc/Tie1/Anxa3/Ptn/Epha2/Epn1/F3/Card10/Bmper/Thbs2/Cela1/Grn/Ccl11/Gata2/Hdac7/Lrg1/Rapgef3/Acvrl1/Cib1/Dcn/Reck/Hmox1/Mapk7/Naxe/Srpx2/Wnt5a/Klf4/Vegfb |
| GO:0050678 | regulation of epithelial cell proliferation | 0,321770272 | 1.880074974 | 0.00032839 | Wnt10b/Sfrp2/Twist1/Pgf/Nrarp/Bnc1/Gli1/Wfdc1/Ccl24/Fmc1/Gpx1/Glul/Ngfr/Wnt2/Ecm1/Gas1/Aplnr/Bax/Lims2/Vash1/Nme2/Cdkn2b/Fgf1/Reg1/Emc10/Nr4a1/Dlk1/Rgcc/Serpinf1/Sfn/Apoe/Cav1/Id1/Eng/Cyba/Osr1/Agtr1a/Sparc/Ptn/Htra1/F3/Pygo2/Smo/Pold4/Jun/Grn/Plau/Ccl11/Gata2/Lrg1/Pdgfb/Aldh1a2/Erbb2/Zfp36/Acvrl1/Atoh8/Zfas1/Nupr1/Hmox1/Aimp1/Bmp4/Wnt5a/Hpn/Vegfb/Ppard/Sfrp1/Sulf1/Egfl7/Cav2/Zfp703/Pdcd6/Hras/Vegfa |
| GO:0006333 | chromatin assembly or disassembly | -0,591403951 | -1.98823778 | 0.000385397 | H4c14/H4c12/H4c8/H1f4/H3c4/H4f16/H3c6/H1f3/H1f5/H2bc3/H4c3/H3c1/H3c7/H4c1/H2bc7/H3c3/H3c2/H4c4/H4c18/H4c6/H1f1 |
| GO:0042775 | mitochondrial ATP synthesis coupled electron transport | 0,567236842 | 2.488159378 | 0.000533691 | Cox6a1/Cox5b/Cox4i2/Ndufb6/Uqcrq/Cyc1/COX3/Ndufc2/Uqcr10/Ndufv3/Sdhc/Ndufa8/Ndufb9/Ndufb8/Ndufa12/Pink1/CYTB/Cox5a/Uqcrfs1/Sdhd/Coq7/Uqcc3/Ndufa7/Ndufs8/Coq9/Park7/Uqcrh/Uqcrc1/Cox4i1/Ndufv1/Taz/Iscu/Ndufs6/Dnajc15/Dguok/ND2 |
| GO:0010257 | NADH dehydrogenase complex assembly | 0,602541603 | 2.464705511 | 0.000533691 | Ndufa2/Ndufa11/Ndufa6/Ndufaf8/Ndufb10/Ndufb7/Ndufa13/Ndufb11/Ndufb6/Ndufa1/Ndufa3/Ndufs7/Ndufc1/Ndufb3/Ndufc2/Ndufa5/Ndufaf3/Ndufa8/Ndufb9/Ndufb8 |
| GO:0032981 | mitochondrial respiratory chain complex I assembly | 0,602541603 | 2.464705511 | 0.000533691 | Ndufa2/Ndufa11/Ndufa6/Ndufaf8/Ndufb10/Ndufb7/Ndufa13/Ndufb11/Ndufb6/Ndufa1/Ndufa3/Ndufs7/Ndufc1/Ndufb3/Ndufc2/Ndufa5/Ndufaf3/Ndufa8/Ndufb9/Ndufb8 |
| GO:0071824 | protein-DNA complex subunit organization | -0,559624028 | -1.906324535 | 0.000615425 | H4c14/Mcmdc2/H4c12/H4c8/H1f4/H3c4/H4f16/H3c6/H1f3/H1f5/H2bc3/H4c3/H3c1/H3c7/H4c1/H2bc7/H3c3/H3c2/H4c4/H4c18/H4c6/H1f1 |
| GO:0022904 | respiratory electron transport chain | 0,491753299 | 2.408789663 | 0.000872262 | Cox6a1/Cox5b/COX2/Cox4i2/Ndufaf2/ND4L/Ndufb6/Uqcrq/Cyc1/COX3/Ndufc2/Ndufa5/Gpd1/Uqcr10/Ndufv3/Sdhc/Ndufa8/Ndufb9/Slc25a22/Ndufb8/Ndufa12/Pink1/CYTB/Cox5a/Uqcrfs1/Sdhd/Coq7/Uqcc3/Ndufa7/Ndufs8/Sdhb/Coq9/Park7/Uqcrh/Uqcrc1/Cox4i1/Ndufv1/Taz/Iscu |
| GO:0015980 | energy derivation by oxidation of organic compounds | 0,336666424 | 1.924706136 | 0.001198847 | Cox6a1/Gcgr/Cavin3/Cox5b/COX2/Cox4i2/Ndufaf2/Sdhaf4/Gck/Bax/ND4L/C1qtnf2/Nfatc4/Ins2/Ndufb6/Uqcrq/Cyc1/Ndufs7/Cisd1/COX3/Bloc1s1/Adrb1/Ndufc2/Ndufa5/Slc37a4/Gpd1/Uqcr10/Ndufv3/Atp5d/Chchd4/Sdhc/Ndufa8/Khk/Gnas/Ndufb9/Gnmt/Slc25a22/Ndufb8/Ndufa12/Pink1/ND1/CYTB/Nop53/Sirt3/Cox5a/Uqcrfs1/Sdhd/Coq7/Uqcc3/Ndufa7/Ndufs8/Sdhb/Bnip3/Coq9/Park7/Uqcrh/Uqcrc1/Coq10a/Cox4i1/Stbd1/Flcn/Ppp1ca/Slc25a23/Ndufv1/Taz/Iscu/Ppp1cc/Gaa/Ndufs6/Oxa1l/Mdh2/Akt2/Shmt2 |
| GO:0006631 | fatty acid metabolic process | 0,343881113 | 1.988483676 | 0.001514143 | Ptgis/Pdk4/Twist1/Pla2g1b/Alox15/Gpx1/Cyp2s1/Dgat2/Ltc4s/Cpt1c/C1qtnf2/Ins2/Prxl2b/Slc27a3/Cav1/Sphk1/Dbi/Mid1ip1/Cygb/Slc27a1/Mif/Eif6/Anxa1/Alkbh7/Gpx4/Acads/Acaa1a/Ptges/Slc45a3/Pck2/Lypla2/Ptgs1/Fabp4/Cyp2d22/Ech1/Edn1/Alox5ap/Gstm2/Abcd4/Tnxb/Fasn/Acaa2/Nr1h2/Pla2g15/Dgat1/Etfa/Abhd16a/Echs1/Mlycd/Ppard/Ggt5/Tysnd1/Aacs/Mcat/Acadvl/Cryl1/Ilvbl/Elovl1/Acot7/Prkag1/Etfb |
| GO:0009150 | purine ribonucleotide metabolic process | 0,31855913 | 1.885619748 | 0.001514143 | Atp5k/ATP8/Pdk4/Atp5j2/Ddit4/Atp5e/Trem2/Cox5b/COX2/Ier3/Antkmt/Dgat2/Atp5g3/Gck/Nme2/Ins2/Npr1/Hint1/Adcy1/Slc27a3/Nme4/ATP6/Aprt/Pgam1/Cyc1/Mif/Eif6/Galk1/Epha2/Ndufc2/Gpd1/Npr2/Eno3/Hmgcl/Nudt18/Atp5d/Nupr1/Ldhd/Khk/Fasn/Acaa2/Mvd/Ppcs/Dgat1/Ak1/Mlycd/Aldoc/Eno2/Mpc2/Guk1 |
| GO:0006323 | DNA packaging | -0,560448725 | -1.898848122 | 0.001514143 | H4c14/H4c12/H4c8/H1f4/H3c4/H4f16/H3c6/H1f3/H1f5/H2bc3/H4c3/H3c1/H3c7/H4c1/H2bc7/H3c3/H3c2/H4c4/H4c18/H4c6/H1f1 |
| GO:0009152 | purine ribonucleotide biosynthetic process | 0,434064845 | 2.274309058 | 0.001514143 | Atp5k/ATP8/Pdk4/Atp5j2/Atp5e/Trem2/Cox5b/COX2/Antkmt/Atp5g3/Nme2/Npr1/Adcy1/Nme4/ATP6/Aprt/Cyc1/Ndufc2/Npr2/Atp5d/Ldhd/Ppcs/Ak1/Mlycd/Mpc2/Guk1/Atp5g1/Atp5j/Acot7/Uqcc3/Flcn/Ak2/Aldoa/Bcl2l1/Taz/Coasy/Map2k1/Sphk2/Atp5pb/Adssl1/Ppcdc/Nme6/Dguok/Pdk2/Atp5h/Nme1/Dcakd/Pkm/Papss1/Gcdh/Atp5c1/Atp5b/Impdh1/Pdhb/Hprt |
| GO:0001935 | endothelial cell proliferation | 0,406153796 | 2.124037952 | 0.001514143 | Pgf/Nrarp/Ccl24/Ngfr/Wnt2/Ecm1/Aplnr/Vash1/Emc10/Nr4a1/Dlk1/Cd34/Rgcc/Apoe/Cav1/Eng/Cyba/Agtr1a/Sparc/Epha2/F3/Bmper/Pold4/Loxl2/Jun/Ccl11/Gata2/Lrg1/Pdgfb/Aldh1a2/Acvrl1/Atoh8/Hmox1/Aimp1/Bmp4/Wnt5a/Vegfb/Scarb1/Sulf1/Egfl7/Cav2/Pdcd6/Vegfa/Thap1 |
| GO:0042254 | ribosome biogenesis | 0,31876629 | 1.924011315 | 0.001801829 | Rpp25/Rps16/Rps10/Rplp0/Rpl3/Rps21/Mrpl20/Glul/Rpl35a/Rps17/Imp3/Eif6/Ddx28/Rpl14/Rps19/Npm3/Rpl26/Tsr3/Rrs1/Gtf3a/Rrp9/Pop5/Rps8/Rpl10a/Pih1d1/Mrpl36/Nob1/Exosc5/Nle1/Pop7/Mpv17l2/Nop53/Gar1/Exosc7/Wdr18/Rpl12/Ppan/Rps5/Riox1/Fam207a/Rps7/Emg1/Exosc4/Abt1/Sirt7/Mrpl11/Nsun5/Ercc2/Isg20/Ddx54/Nop10/Znhit3/Traf7/Rpusd4/Nop16/Rbfa/Zfp622/Noc4l/Bud23/Wdr74/C1qbp/Bop1/Mrps7/Tbl3/Malsu1/Pak1ip1/Fbl/Mrto4/Mrps11/Lyar/Rps14/Rpl35/Utp3/Noc2l/Rps28/Eral1/Rpl7l1/Rsl24d1/Rrp36/Rrp1/Ebna1bp2/Gtf2h5/Ran/Surf6/Sbds/Nop9/Rps15/Pes1/Ngdn/Utp11/Nhp2/Sart1/C1d/Ddx49/Rps27l |
| GO:0052547 | regulation of peptidase activity | 0,303374414 | 1.840034629 | 0.002657148 | Cryab/Spink1/Timp1/Sfrp2/Prr7/Pcsk1n/Anxa8/Cldn4/Crb2/Slpi/Wfdc1/Pi16/Col7a1/Nol3/Gpx1/Aqp1/Ngfr/Ecm1/Papln/Bax/Serping1/Wfdc2/Nr4a1/Bbc3/Ccn2/Timp2/Gsn/Ndufa13/Serpinf1/Mmp9/Sfn/Wnt9a/Serpinh1/Cav1/Rcn3/Cstb/Cst3/Cldn3/Svbp/Bok/F3/Serpinb6a/Dpep1/Pih1d1/Grn/Tbc1d10a/Herpud1/Gas6/Renbp/Cdkn2d/Pebp1/Reck/Vtn/Nle1/Ubxn1/Atp13a2/Mtch1/Pycard/Serpine2/Klf4/Htra2/Mgmt |
| GO:0072521 | purine-containing compound metabolic process | 0,301970193 | 1.766040415 | 0.002837455 | Atp5k/ATP8/Pdk4/Atp5j2/Ddit4/Atp5e/Trem2/Cox5b/COX2/Ier3/Antkmt/Dgat2/Atp5g3/Gck/Nme2/Oasl2/Nos3/Ins2/Npr1/Urah/Hint1/Adcy1/Slc27a3/Nme4/ATP6/Aprt/Pgam1/Cyc1/Mif/Eif6/Galk1/Epha2/Gamt/Ndufc2/Gpd1/Npr2/Eno3/Hmgcl/Nudt18/Atp5d/Nupr1/Ldhd/Khk/Fasn/Gnmt/Acaa2/Mvd/Ppcs/Pemt/Dgat1/Ak1/Mlycd/Aldoc/Eno2/Mpc2/Guk1/Macrod1/Itpa/Ak7/Atp5g1/Atp5j/Pmvk/Acot7/Uqcc3 |
| GO:0046390 | ribose phosphate biosynthetic process | 0,413411803 | 2.159997583 | 0.002895116 | Atp5k/ATP8/Pdk4/Atp5j2/Atp5e/Trem2/Cox5b/COX2/Antkmt/Atp5g3/Nme2/Npr1/Adcy1/Nme4/ATP6/Aprt/Cyc1/Ndufc2/Npr2/Atp5d/Ldhd/Ppcs/Ak1/Mlycd/Mpc2/Guk1/Atp5g1/Atp5j/Acot7/Uqcc3/Flcn/Ak2/Tkt/Aldoa/Bcl2l1/Taz/Coasy/Map2k1/Sphk2/Atp5pb/Adssl1/Ppcdc/Nme6/Dguok/Pdk2/Atp5h/Nme1/Dcakd/Pkm/Papss1/Gcdh/Atp5c1/Atp5b/Impdh1/Pdhb/Hprt |
| GO:0019693 | ribose phosphate metabolic process | 0,31112276 | 1.877876459 | 0.003335589 | Atp5k/ATP8/Pdk4/Atp5j2/Ddit4/Atp5e/Trem2/Cox5b/COX2/Ier3/Antkmt/Dgat2/Atp5g3/Gck/Nme2/Ins2/Npr1/Hint1/Adcy1/Slc27a3/Nme4/ATP6/Aprt/Pgam1/Cyc1/Mif/Eif6/Galk1/Epha2/Ndufc2/Gpd1/Npr2/Eno3/Hmgcl/Nudt18/Atp5d/Nupr1/Ldhd/Khk/Fasn/Acaa2/Mvd/Ppcs/Dgat1/Nudt14/Ak1/Mlycd/Aldoc/Eno2/Mpc2/Guk1 |
| GO:0006753 | nucleoside phosphate metabolic process | 0,302528643 | 1.804250097 | 0.003425595 | Atp5k/ATP8/Pdk4/Atp5j2/Ddit4/Atp5e/Trem2/Cox5b/COX2/Ier3/Antkmt/Dgat2/Atp5g3/Gck/Nme2/Oasl2/Entpd2/Nos3/Ins2/Npr1/Naprt/Dnph1/Hint1/Adcy1/Slc27a3/Nme4/ATP6/Aprt/Pgam1/Cyc1/Mif/Eif6/Galk1/Epha2/Ndufc2/Upp1/Dctpp1/Gpd1/Npr2/Eno3/Hmgcl/Nudt18/Atp5d/Nupr1/Ldhd/Khk/Fasn/Acaa2/Mvd/Ppcs/Naxe/Dgat1/Nt5c3b/Nudt14/Ak1/Mlycd/Aldoc/Eno2/Mpc2/Guk1/Itpa/Ak7/Atp5g1/Atp5j/Pmvk/Acot7/Uqcc3/Nt5m |
| GO:0015985 | energy coupled proton transport, down electrochemical gradient | 0,680886962 | 2.325599183 | 0.003543999 | Atp5k/ATP8/Atp5j2/Atp5e/Cox5b/Antkmt/Atp5g3/ATP6/Cyc1 |
| GO:0015986 | ATP synthesis coupled proton transport | 0,680886962 | 2.325599183 | 0.003543999 | Atp5k/ATP8/Atp5j2/Atp5e/Cox5b/Antkmt/Atp5g3/ATP6/Cyc1 |
| GO:0006163 | purine nucleotide metabolic process | 0,301937146 | 1.828328864 | 0.003905893 | Atp5k/ATP8/Pdk4/Atp5j2/Ddit4/Atp5e/Trem2/Cox5b/COX2/Ier3/Antkmt/Dgat2/Atp5g3/Gck/Nme2/Oasl2/Nos3/Ins2/Npr1/Hint1/Adcy1/Slc27a3/Nme4/ATP6/Aprt/Pgam1/Cyc1/Mif/Eif6/Galk1/Epha2/Ndufc2/Gpd1/Npr2/Eno3/Hmgcl/Nudt18/Atp5d/Nupr1/Ldhd/Khk/Fasn/Acaa2/Mvd/Ppcs/Dgat1/Ak1/Mlycd/Aldoc/Eno2/Mpc2/Guk1 |
| GO:0006336 | DNA replication-independent nucleosome assembly | -0,790861086 | -1.961553763 | 0.003907426 | H4c11/H4c14/H4c12/H4c8/H4f16/H4c3/H4c1/H4c4/H4c18/H4c6 |
| GO:0009260 | ribonucleotide biosynthetic process | 0,412345834 | 2.162190856 | 0.004208275 | Atp5k/ATP8/Pdk4/Atp5j2/Atp5e/Trem2/Cox5b/COX2/Antkmt/Atp5g3/Nme2/Npr1/Adcy1/Nme4/ATP6/Aprt/Cyc1/Ndufc2/Npr2/Atp5d/Ldhd/Ppcs/Ak1/Mlycd/Mpc2/Guk1/Atp5g1/Atp5j/Acot7/Uqcc3/Flcn/Ak2/Aldoa/Bcl2l1/Taz/Coasy/Map2k1/Sphk2/Atp5pb/Adssl1/Ppcdc/Nme6/Dguok/Pdk2/Atp5h/Nme1/Dcakd/Pkm/Papss1/Gcdh/Atp5c1/Atp5b/Impdh1/Pdhb/Hprt |
| GO:0009117 | nucleotide metabolic process | 0,302980369 | 1.76823437 | 0.004442401 | Atp5k/ATP8/Pdk4/Atp5j2/Ddit4/Atp5e/Trem2/Cox5b/COX2/Ier3/Antkmt/Dgat2/Atp5g3/Gck/Nme2/Oasl2/Nos3/Ins2/Npr1/Naprt/Dnph1/Hint1/Adcy1/Slc27a3/Nme4/ATP6/Aprt/Pgam1/Cyc1/Mif/Eif6/Galk1/Epha2/Ndufc2/Upp1/Dctpp1/Gpd1/Npr2/Eno3/Hmgcl/Nudt18/Atp5d/Nupr1/Ldhd/Khk/Fasn/Acaa2/Mvd/Ppcs/Naxe/Dgat1/Nt5c3b/Ak1/Mlycd/Aldoc/Eno2/Mpc2/Guk1/Itpa/Ak7/Atp5g1/Atp5j/Pmvk/Acot7/Uqcc3/Nt5m |
| GO:0006575 | cellular modified amino acid metabolic process | 0,401309868 | 2.096767292 | 0.004552902 | Gstp2/Ckm/Gstm3/Dio3/Gpx1/Cpt1c/Ckb/Ggt6/Chac1/Nat8/Gpx3/Slc27a1/Serinc2/Gpx4/Gamt/Dpep1/Ctsk/Plod2/Gstm2/Sod1/Gstm6/Gnmt/Pemt/Hpn/Abhd16a/Ggt5/Gsta4/Ckmt1/Gstt1/Mthfd2/Gstk1/Park7/Gstp1/Gstm1/Mgst1/Ctsb/Oplah/Hmgn5/Kyat1/Mthfr/Fpgs/Shmt2/Pcyox1l/Ptdss2/Icmt/Gch1/Crat/Gstz1/Hagh/Plod3/Aldh1l1/Ggh/Nfe2l1/Abhd12/Ctsl |
| GO:0006749 | glutathione metabolic process | 0,561297485 | 2.235028567 | 0.004872959 | Gstp2/Gstm3/Gpx1/Ggt6/Chac1/Nat8/Gpx3/Gpx4/Dpep1/Gstm2/Sod1/Gstm6/Ggt5/Gsta4/Gstt1/Gstk1/Park7/Gstp1/Gstm1/Mgst1/Oplah/Hmgn5 |
| GO:1904018 | positive regulation of vasculature development | 0,358572572 | 1.954068428 | 0.005318985 | Sfrp2/Ptgis/Pgf/Rhob/Ccl24/Aqp1/Angpt4/Ecm1/Cma1/Hspb1/Adm/Aplnr/Tmem100/Fgf1/Emc10/Nos3/Cd34/Mmp9/Smoc2/Sphk1/Adm2/Eng/Ramp2/Agtr1a/Anxa1/Tie1/Anxa3/F3/Bmper/Cela1/Grn/Ccl11/Gata2/Hdac7/Lrg1/Pdgfb/Rapgef3/Acvrl1/Cib1/Hmox1/Srpx2/Wnt5a/Klf4/Vegfb |
| GO:0045653 | negative regulation of megakaryocyte differentiation | -0,780148516 | -1.934983633 | 0.006627423 | H4c11/H4c14/H4c12/H4c8/H4f16/H4c3/H4c1/H4c4/H4c18/H4c6 |
| GO:0032964 | collagen biosynthetic process | 0,569936533 | 2.229143599 | 0.006627423 | Scx/Ccn2/Rgcc/Serpinh1/Rcn3/Eng/Prdx5/Cygb/P3h4/Col1a1/P3h3/Vim/Wnt4/Pdgfb/Rapgef3/Col5a1/Bmp4/Tgfb3/Ppard/Emilin1 |
| GO:0048645 | animal organ formation | 0,554148335 | 2.206561391 | 0.006627423 | Nkx3-2/Gata5/Hand2/Wnt11/Wnt2/Wt1/Lrp2/Fgf1/Ntf5/Gng5/Lemd2/Map2k2/Fgfr4/Bmp4/Wnt5a/Rdh10/Sulf1 |
| GO:0008630 | intrinsic apoptotic signaling pathway in response to DNA damage | 0,407189482 | 2.071703914 | 0.006640166 | Tpt1/Snai1/Phlda3/Ddit4/Crip1/Ier3/Bax/Bbc3/Nfatc4/Sfn/Ackr3/Zfp385a/Steap3/Hic1/Mif/Cdkn1a/Epha2/Bok/Rpl26/E2f1/Bcl2l2/Tmem109/Ifi204/Cdkn2d/Nupr1/Hmox1/Pycard/Htra2/Tmem161a |
| GO:0009259 | ribonucleotide metabolic process | 0,311249169 | 1.851601002 | 0.006640166 | Atp5k/ATP8/Pdk4/Atp5j2/Ddit4/Atp5e/Trem2/Cox5b/COX2/Ier3/Antkmt/Dgat2/Atp5g3/Gck/Nme2/Ins2/Npr1/Hint1/Adcy1/Slc27a3/Nme4/ATP6/Aprt/Pgam1/Cyc1/Mif/Eif6/Galk1/Epha2/Ndufc2/Gpd1/Npr2/Eno3/Hmgcl/Nudt18/Atp5d/Nupr1/Ldhd/Khk/Fasn/Acaa2/Mvd/Ppcs/Dgat1/Ak1/Mlycd/Aldoc/Eno2/Mpc2/Guk1 |
| GO:0006342 | chromatin silencing | -0,636251306 | -1.939245258 | 0.006836341 | H1f4/H2ac20/H2al1m/H1f3/H1f5/H2ac10/H2ap/H2ac15/H2ac4/H2al3/H2ac6/H1f1 |
| GO:0001936 | regulation of endothelial cell proliferation | 0,396611327 | 2.031274943 | 0.006946842 | Pgf/Nrarp/Ccl24/Ngfr/Wnt2/Ecm1/Aplnr/Vash1/Emc10/Nr4a1/Dlk1/Rgcc/Apoe/Cav1/Eng/Cyba/Agtr1a/Sparc/F3/Pold4/Jun/Ccl11/Gata2/Lrg1/Pdgfb/Aldh1a2/Acvrl1/Atoh8/Hmox1/Aimp1/Bmp4/Wnt5a/Vegfb/Sulf1/Egfl7/Cav2/Pdcd6/Vegfa |
| GO:0016053 | organic acid biosynthetic process | 0,346937034 | 1.944959952 | 0.006946842 | Ptgis/Pdk4/Pla2g1b/Alox15/Glul/Ltc4s/Nags/Rbp1/Prxl2b/Sphk1/Sdsl/Chst14/Mid1ip1/Mif/Eif6/Anxa1/Tha1/Ptges/Gamt/Slc45a3/Ptgs1/Plod2/Edn1/Alox5ap/Pdgfb/Aldh1a2/Gstm2/Pycrl/Pycr2/Mgst3/Fgfr4/Fasn/Nr1h2/Smpd3/Rdh10/Mlycd/Ggt5/Ldhb/Phgdh/Mcat/Acadvl/Apip/Ilvbl/Elovl1/Acot7/Prkag1/Chst12/Park7/Scap/Csad/Ptges2/Got1/Mthfr/Nr1h3/Ap2a1/Tecr/Abat/Fpgs/Srebf1/Shmt2 |
| GO:0072330 | monocarboxylic acid biosynthetic process | 0,400201444 | 2.050764747 | 0.00722674 | Ptgis/Pdk4/Pla2g1b/Alox15/Ltc4s/Rbp1/Prxl2b/Sphk1/Chst14/Mid1ip1/Mif/Eif6/Anxa1/Ptges/Gamt/Slc45a3/Ptgs1/Edn1/Aldh1a2/Gstm2/Fgfr4/Fasn/Nr1h2/Rdh10/Mlycd/Ldhb/Mcat/Acadvl/Elovl1/Acot7/Prkag1/Chst12/Park7 |
| GO:0050679 | positive regulation of epithelial cell proliferation | 0,34739614 | 1.909974402 | 0.008217093 | Twist1/Pgf/Nrarp/Bnc1/Ccl24/Glul/Wnt2/Ecm1/Gas1/Aplnr/Nme2/Fgf1/Reg1/Emc10/Nr4a1/Cav1/Id1/Cyba/Osr1/Agtr1a/Ptn/Htra1/F3/Smo/Pold4/Jun/Grn/Ccl11/Gata2/Lrg1/Pdgfb/Erbb2/Acvrl1/Hmox1/Bmp4/Wnt5a/Hpn/Vegfb/Sfrp1/Egfl7/Cav2/Zfp703/Pdcd6/Hras/Vegfa |
| GO:0010817 | regulation of hormone levels | 0,277968221 | 1.682273173 | 0.008217093 | Ren1/Retn/Pcsk1n/Dio3/Pim3/Ccn3/Ltbp4/Cyp46a1/Ndufaf2/Aqp1/Gal/Glul/Cyp2s1/Trpv4/Lif/Inhbb/Adm/Lrp2/Gck/Dkk3/Selenom/Igfbp3/Rbp1/Chga/Inha/Rbp4/Agtr1a/C1qtnf1/Anxa1/Fdx1/Slc7a5/Cela2a/Cry2/Pck2/Ctsk/C1qtnf12/Cyp2d22/Pcsk4/Capn10/Wnt4/Edn1/Aldh1a2/Spp1/Uqcc2/Ildr2/Clcn2/Fgfr4/Aimp1/Gnas/Adh1/Ace/Ece2/Slc8b1/Dgat1/Hpn/Smpd3/Stard3/Slc7a8/Rdh10/Niban2/Ppard/Scarb1/Sfrp1/Sirt3/Nlgn2/Mpc2/Aacs |
| GO:0009145 | purine nucleoside triphosphate biosynthetic process | 0,483135757 | 2.116996669 | 0.008217093 | Atp5k/ATP8/Atp5j2/Atp5e/Trem2/Cox5b/COX2/Antkmt/Atp5g3/Nme2/Nme4/ATP6/Cyc1 |
| GO:0009201 | ribonucleoside triphosphate biosynthetic process | 0,483135757 | 2.116996669 | 0.008217093 | Atp5k/ATP8/Atp5j2/Atp5e/Trem2/Cox5b/COX2/Antkmt/Atp5g3/Nme2/Nme4/ATP6/Cyc1 |
| GO:0009206 | purine ribonucleoside triphosphate biosynthetic process | 0,483135757 | 2.116996669 | 0.008217093 | Atp5k/ATP8/Atp5j2/Atp5e/Trem2/Cox5b/COX2/Antkmt/Atp5g3/Nme2/Nme4/ATP6/Cyc1 |
| GO:0045669 | positive regulation of osteoblast differentiation | 0,516362711 | 2.194836478 | 0.008341717 | Tmem119/Wnt10b/Sfrp2/Cebpb/Cebpd/Jund/Wnt7b/Cebpa/Pdlim7/Gdf10/Wnt4/Clic1/Ifi204/Lrp3/Atraid/Gnas/Bmp4 |
| GO:0034724 | DNA replication-independent nucleosome organization | -0,770757996 | -1.932176848 | 0.008620185 | H4c11/H4c14/H4c12/H4c8/H4f16/H4c3/H4c1/H4c4/H4c18/H4c6 |
| GO:0042445 | hormone metabolic process | 0,441892332 | 2.129344272 | 0.008620185 | Ren1/Pcsk1n/Dio3/Cyp46a1/Gal/Cyp2s1/Adm/Dkk3/Selenom/Rbp1/Rbp4/Fdx1/Ctsk/Cyp2d22/Pcsk4/Wnt4/Aldh1a2/Spp1/Clcn2/Adh1/Ace/Ece2/Hpn/Stard3/Rdh10/Scarb1 |
| GO:0050728 | negative regulation of inflammatory response | 0,461088868 | 2.143946461 | 0.008620185 | Hamp/Ptgis/Wfdc1/Ccn3/Gpx2/Gpx1/Ier3/Ins2/Ctla2a/Serpinf1/Metrnl/Apoe/Proc/Rps19/Tff2/C1qtnf12/Grn/Gps2/Socs3/Zfp36/Sod1/Psmb4/Fndc4/Pycard/Ppard |
| GO:0046394 | carboxylic acid biosynthetic process | 0,344895013 | 1.896223274 | 0.008947457 | Ptgis/Pdk4/Pla2g1b/Alox15/Glul/Ltc4s/Nags/Rbp1/Prxl2b/Sphk1/Sdsl/Chst14/Mid1ip1/Mif/Eif6/Anxa1/Tha1/Ptges/Gamt/Slc45a3/Ptgs1/Plod2/Edn1/Alox5ap/Pdgfb/Aldh1a2/Gstm2/Pycrl/Pycr2/Mgst3/Fgfr4/Fasn/Nr1h2/Smpd3/Rdh10/Mlycd/Ggt5/Ldhb/Phgdh/Mcat/Acadvl/Apip/Ilvbl/Elovl1/Acot7/Prkag1/Chst12/Park7/Scap/Ptges2/Got1/Mthfr/Nr1h3/Ap2a1/Tecr/Abat/Fpgs/Srebf1/Shmt2 |
| GO:0006164 | purine nucleotide biosynthetic process | 0,378341774 | 1.950364343 | 0.008987781 | Atp5k/ATP8/Pdk4/Atp5j2/Atp5e/Trem2/Cox5b/COX2/Antkmt/Atp5g3/Nme2/Oasl2/Nos3/Npr1/Adcy1/Nme4/ATP6/Aprt/Cyc1 |
| GO:0071103 | DNA conformation change | -0,515871981 | -1.777850375 | 0.009337825 | H4c14/Mcmdc2/H4c12/H4c8/H1f4/H3c4/H4f16/H3c6/H1f3/H1f5/H2bc3/H4c3/H3c1/H3c7/H4c1/H2bc7/H3c3/H3c2/H4c4/H4c18/H4c6/H1f1 |
| GO:0044282 | small molecule catabolic process | 0,312932659 | 1.791900005 | 0.009830346 | Twist1/Tdh/Cyp46a1/Cbr3/Nos3/Ddah2/Apoe/Sdsl/Akr1b8/Dbi/Galk1/Cdo1/Acads/Hsd3b7/Acaa1a/Tha1/Pck2/Lypla2/Upp1/Prodh/Hmgcl/Bckdha/Pycrl/Nudt18/Renbp/Ldhd/Abcd4/Khk/Qdpr/Adh1/Acaa2/Etfa/Nagk/Abhd16a/Amdhd2/Echs1/Mlycd/Ppard/Scarb1/Tysnd1/Pnkd/Acadvl/Gcat/Hyal2/Ilvbl/Mat1a/Acot7/Etfb/Park7/Idua/Echdc2/Mgat1/Haghl/Bad/Csad/Acad8/Got1/Acot8/Aldh4a1/Abat/Gnpda1/Akt2/Shmt2/Pcyox1l/Thnsl2/Pfkl/Slc27a4/Gcsh/Lrp5/Crat/Tpi1/Bckdk/Gstz1/Eci1/Pfkm/Srd5a3/Hagh/Aldh1l1/Gcdh/Akr1a1/Abcd1/Akt1/Pex2/Pex5/Cpt2/Hprt/Adh5/Decr1 |
| GO:0045444 | fat cell differentiation | 0,32759562 | 1.825437723 | 0.009830346 | Wnt10b/Ccdc85b/Sfrp2/Retn/Cebpb/Cebpd/Gpx1/Trpv4/Inhbb/Fndc5/Clip3/Rarres2/Nr4a1/Dlk1/Cebpa/Metrnl/Zfp385a/Atf5/Gdf10/Jdp2/Tmem120a/Mmp11/Pex11a/Medag/Selenbp1/Adrb1/Fabp4/E2f1/Tlcd3b/Gps2/Gata2/Angptl8/Lrg1/Tgfb1i1/Lrp3/Zfp36 |
| GO:0045766 | positive regulation of angiogenesis | 0,362410861 | 1.929120686 | 0.010084785 | Sfrp2/Ptgis/Pgf/Rhob/Ccl24/Aqp1/Angpt4/Ecm1/Cma1/Hspb1/Adm/Aplnr/Fgf1/Emc10/Nos3/Cd34/Mmp9/Smoc2/Sphk1/Adm2/Eng/Ramp2/Agtr1a/Anxa1/Tie1/Anxa3/F3/Bmper/Cela1/Grn/Ccl11/Gata2/Hdac7/Lrg1/Rapgef3/Acvrl1/Cib1/Hmox1/Srpx2/Wnt5a/Klf4/Vegfb |
| GO:0045814 | negative regulation of gene expression, epigenetic | -0,620571646 | -1.915488051 | 0.010256544 | H1f4/H2ac20/H2al1m/H1f3/H1f5/H2ac10/H2ap/H2ac15/H2ac4/H2al3/H2ac6/H1f1 |
| GO:0048562 | embryonic organ morphogenesis | 0,338994196 | 1.855850943 | 0.010513193 | Tcf21/Nkx3-2/Ccdc103/Twist1/Crb2/Gli1/Hand2/Mdfi/Wnt11/Fzd2/Mfap2/Gas1/Wnt9a/Hoxb6/Mfap5/Eng/Rbp4/Osr1/Tbx2/Hoxb7/Epha2/Slc39a3/Smo/Naglu/Sox17/Gata2/Edn1/Aldh1a2/Mmp14/Sod1/Chst11/Srf/Gnas/Bmp4/Wnt5a/Tgfb3/Hpn/Rdh10/Mafb |
| GO:0016485 | protein processing | 0,383565961 | 1.964462115 | 0.010575238 | Ren1/Clec3b/Pcsk1n/Klk1b26/Nol3/Klk1b9/Gas1/Gsn/Plat/Ctla2a/Klk1b8/Aebp1/Klk1b11/Chac1/C1ra/Tmem98/Adamts2/F3/Pcsk4/Plau/Angptl8/Tmem208/Tbc1d10a/Mmp14/Cpxm1/Rce1/Ece2/Pycard/Serpine2/Hpn/Tysnd1/Hgfac |
| GO:0051604 | protein maturation | 0,319965389 | 1.768752495 | 0.010575238 | Ren1/Clec3b/Pcsk1n/Tspan17/Klk1b26/Nol3/Klk1b9/Gas1/Gsn/Plat/Ctla2a/Klk1b8/Aebp1/Serpinh1/Klk1b11/Chac1/C1ra/Tmem98/Adamts2/F3/Cdk20/Dohh/Glrx5/Pcsk4/Plau/Angptl8/Tmem208/Tbc1d10a/Mmp14/Cpxm1/Naa80/Chchd4/Rce1/Ece2/Pycard/Serpine2/Hpn/Ciao2b/Hscb/Tysnd1/Hgfac/Dhps/Glrx3/Notch4/Isca2/Spcs1/Stub1/Enpep/Spg7/Lmf2/Bad/Pgk1 |
| GO:0090092 | regulation of transmembrane receptor protein serine/threonine kinase signaling pathway | 0,333887753 | 1.857220221 | 0.010784819 | Sfrp4/Sfrp2/Crb2/Ccn3/Dact2/Inhbb/Lrp2/Chrd/Cdkn2b/Nbl1/Tgfbr3l/Inha/Cav1/Kcp/Gdf10/Eng/Pmepa1/Htra3/Fbxl15/Htra1/Bmper/Fstl1/Bambi/Fzd1/Lemd2/Fstl3/Lrg1/Rgma/Tgfb1i1/Eid2/Fkbp8/Acvrl1/Chst11/Bmp4/Wnt5a/Tgfb3/Chrdl2/Cilp/Sfrp1/Sulf1/Cav2/Zfp703/Smad6/Emilin1 |
| GO:0010466 | negative regulation of peptidase activity | 0,352661724 | 1.861467705 | 0.011776539 | Cryab/Spink1/Timp1/Sfrp2/Pcsk1n/Anxa8/Crb2/Slpi/Wfdc1/Pi16/Col7a1/Nol3/Gpx1/Aqp1/Ecm1/Papln/Serping1/Wfdc2/Nr4a1/Timp2/Serpinf1/Mmp9/Sfn/Wnt9a/Serpinh1/Cstb/Cst3/Serpinb6a/Dpep1/Pih1d1/Herpud1/Gas6/Renbp/Cdkn2d/Pebp1/Reck/Vtn |
| GO:0060485 | mesenchyme development | 0,30829796 | 1.774168434 | 0.011776539 | Tcf21/Nrtn/Snai1/Sfrp2/Gata5/Twist1/Crb2/Hand2/Wnt7b/Wnt11/Wnt2/Mcrip1/Wt1/Scx/Tmem100/Adam15/Nos3/Sema6b/Rgcc/Tgfbr3l/Eng/Dact3/Sema6c/Osr1/Tbx2/Col1a1/Sema3f/Bambi/Smo/Loxl2/Sema3g/Wnt4/Edn1/Aldh1a2/Tgfb1i1/Acvrl1/Phldb1/Bmp4/Wnt5a/Tgfb3/Hpn/Mad2l2/Rdh10/Meox1/Sfrp1/Loxl3/Pef1/Zfp703/Sema3b/Pdcd6/Vegfa |
| GO:0090287 | regulation of cellular response to growth factor stimulus | 0,307732413 | 1.770913872 | 0.011813101 | Sfrp4/Sfrp2/Crb2/Ngfr/Lrp2/Chrd/Cdkn2b/Fgf1/Nbl1/Cadm4/Tgfbr3l/Cav1/Smoc2/Kcp/Eng/Pmepa1/Htra3/Fbxl15/Htra1/Bmper/Cd63/Fstl1/Bambi/Fzd1/Lemd2/Fstl3/Wnt4/Lrg1/Rgma/Pdgfb/Tgfb1i1/Eid2/Fkbp8/Acvrl1/Dcn/Tmem204/Chst11/Bmp4/Wnt5a/Tgfb3/Vegfb/Chrdl2/Sfrp1/Sulf1/Gpc1/Cav2/Zfp703/Pdcd6/Smad6/Emilin1/Adgra2/Bcl9l |
| GO:0010594 | regulation of endothelial cell migration | 0,359277666 | 1.878892686 | 0.011813101 | Rhob/Glul/Angpt4/Hspb1/Vash1/Fgf1/Emc10/Nos3/Rgcc/Serpinf1/Apoe/Smoc2/Anxa1/Sparc/Anxa3/Epha2/Svbp/Card10/Bmper/Grn/Bsg/Gata2/Hdac7/Pdgfb/Acvrl1/Cib1/Dcn/Atoh8/Hmox1/Bmp4/Srpx2/Wnt5a/Klf4/Map2k3/Hdac5/Pdcd6/Vegfa/Adgra2/Dll4/Rras |
| GO:1901293 | nucleoside phosphate biosynthetic process | 0,349974153 | 1.826204745 | 0.011813101 | Atp5k/ATP8/Pdk4/Atp5j2/Atp5e/Trem2/Cox5b/COX2/Antkmt/Atp5g3/Nme2/Oasl2/Nos3/Npr1/Naprt/Adcy1/Nme4/ATP6/Aprt/Cyc1 |
| GO:0050900 | leukocyte migration | 0,322690523 | 1.798105402 | 0.013639769 | S100a14/Pgf/Trem2/Ccn3/Gp2/Ccl24/Trpv4/Ecm1/Pf4/Rarres2/Dusp1/Rpl13a/Cd34/Mmp9/Nbl1/Chga/Il34/Selp/Mif/Anxa1/Mmp28/Plvap/Ptn/Thy1/Rps19/Hsd3b7/Aoc3/Dpep1/Slc37a4/Nkx2-3/Podxl2/Ddt/Ccl11/Zp3/Il17rc/Edn1/Pdgfb/Spp1/Gas6/Mmp14/Aimp1/Pycard/Wnt5a/Slc8b1/Smpd3/Vegfb/Cmklr1/Ppib/Cxcl14/Vegfa/Emilin1/Rhog |
| GO:1901615 | organic hydroxy compound metabolic process | 0,265043554 | 1.624459686 | 0.013879653 | Saa1/Snai1/Dio3/Hand2/Pgp/Alox15/Cyp46a1/Spr/Pltp/Dgat2/Dkk3/Akr7a5/Fgf1/Npr1/Apoc1/Rbp1/Pmp22/Cebpa/Apoe/Sphk1/Akr1b8/Rbp4/Itpka/Agtr1a/Galk1/Pth1r/Fdx1/Hsd3b7/Acaa1a/Pcbd2/Plpp2/Pck2/P2ry6/Ctsk/Slc37a4/Cyp2d22/Gpd1/Ddt/Isyna1/Cln6/Wnt4/Ldhd/Sod1/Clcn2/Fgfr4/Npc2/Qdpr/Adh1/Mvd/Ip6k1/Wnt5a/Hpn/Rdh10/Scarb1/Plpp1/Ldhb/Pnkd/Fdxr/Tm7sf2/Acadvl/Coq2/Comt/Gfi1/Apobr/Ly6e/Dpm2 |
| GO:0050886 | endocrine process | 0,476075692 | 2.0227918 | 0.015649633 | Ren1/Retn/Klk1b26/Aqp1/Gal/Lif/Inhbb/Nos3/Selenom/Inha/Cyba/Agtr1a/Mif/C1qtnf1/Cry2/Adrb1/Gja5/Edn1/Fgfr4/Gnas/Ace |
| GO:0009142 | nucleoside triphosphate biosynthetic process | 0,459640375 | 2.069382053 | 0.015720146 | Atp5k/ATP8/Atp5j2/Atp5e/Trem2/Cox5b/COX2/Antkmt/Atp5g3/Nme2/Nme4/ATP6/Cyc1 |
| GO:0043542 | endothelial cell migration | 0,32012579 | 1.787560757 | 0.015720146 | Ccn3/Rhob/Gpx1/Glul/Angpt4/Hspb1/Vash1/Fgf1/Emc10/Nr4a1/Nos3/Rgcc/Serpinf1/Apoe/Smoc2/Anxa1/Sparc/Anxa3/Epha2/Svbp/Card10/Bmper/Loxl2/Grn/Bsg/Gata2/Rab13/Hdac7/Pdgfb/Acvrl1/Cib1/Dcn/Atoh8/Srf/Hmox1/Bmp4/Srpx2/Wnt5a/Klf4/Scarb1/Map2k3/Hdac5/Pdcd6/Vegfa/Adgra2/Dll4/Rras/Ptp4a3/Plekhg5 |
| GO:0030574 | collagen catabolic process | 0,620763409 | 2.12024456 | 0.015950949 | Mmp23/Mmp2/Adam15/Mmp9/Mmp28/Mmp11/Mrc2/Adamts2/Ctsk/Mmp19/Mmp14 |
| GO:0007006 | mitochondrial membrane organization | 0,381534625 | 1.955109786 | 0.015950949 | Timm13/Nol3/Ier3/Siva1/Romo1/Chchd10/Bax/Bbc3/Ndufa13/Micos13/Slc25a4/Alkbh7/Chchd6/Bok/Timm50/Bcl2l2/Acaa2/Eya2/Pink1/Rcc1l/Pdcd5/Uqcc3/Bnip3/Tomm22/Spg7/Timm9/Bad/Bcl2l1/Taz/Bloc1s2/Plekhf1/Mul1/Ppif/Oxa1l/Rhot2/Timm22/Afg3l1/Slc25a5/Samm50/Gsk3a/Tmem11/Adck1/Slc35f6 |
| GO:0051271 | negative regulation of cellular component movement | 0,28161147 | 1.684589953 | 0.015950949 | Timp1/Sfrp2/Ccn3/Rhob/Wnt11/Tmeff2/Angpt4/Dbn1/Igfbp5/Chrd/Vash1/Adam15/Dusp1/Sema6b/Igfbp3/Podn/Rgcc/Serpinf1/Nbl1/Apoe/Eng/Sema6c/Cygb/Fbln1/Mif/Mmp28/Tie1/Ptn/Thy1/Cldn3/Apex1/Svbp/Sema3f/Card10/Dpep1/Cd63/Ddt/Sema3g/Wnt4/Pdgfb/Col3a1/Acvrl1/Dcn/Mapk15/Reck/Srf/Wnt5a/Klf4/Arap3/Ppard/Robo4/Sfrp1/Sulf1/Egfl7/Sema3b/Hdac5/Emilin1/Miip/Spint2/Hyal2/Cd9/Dll4/Rras |
| GO:0072522 | purine-containing compound biosynthetic process | 0,374130319 | 1.916136759 | 0.015950949 | Atp5k/ATP8/Pdk4/Atp5j2/Atp5e/Trem2/Cox5b/COX2/Antkmt/Atp5g3/Nme2/Oasl2/Nos3/Npr1/Adcy1/Nme4/ATP6/Aprt/Cyc1 |
| GO:0009165 | nucleotide biosynthetic process | 0,354996581 | 1.828307677 | 0.015950949 | Atp5k/ATP8/Pdk4/Atp5j2/Atp5e/Trem2/Cox5b/COX2/Antkmt/Atp5g3/Nme2/Oasl2/Nos3/Npr1/Naprt/Adcy1/Nme4/ATP6/Aprt/Cyc1 |
| GO:2000027 | regulation of animal organ morphogenesis | 0,359798232 | 1.887872756 | 0.015950949 | Sfrp2/Gata5/Twist1/Gli1/Hand2/Wnt7b/Wnt11/Fzd2/Ngfr/Wnt2/Lif/Wt1/Bax/Lims2/Fgf1/Apcdd1/Cd34/Plekha4/Eng/Tbx2/Agtr1a/Hoxb7/Nkd1/Gng5/Ptn/Smo/Fzd1/Plau/Wnt4/Edn1 |
| GO:0051346 | negative regulation of hydrolase activity | 0,264381898 | 1.552946031 | 0.016287777 | Cryab/Spink1/Timp1/Apcs/Sfrp2/Pcsk1n/Anxa8/Crb2/Ppp1r35/Slpi/Wfdc1/Pi16/Col7a1/Nol3/Gpx1/Aqp1/Ecm1/Ppp1r1b/Papln/Dbn1/Serping1/Wfdc2/Nr4a1/Nos3/Timp2/Apoc1/Serpinf1/Mmp9/Sfn/Wnt9a/Serpinh1/Cstb/Ccdc8/Anxa1/Cst3/Cry2/Serpinb6a/Gpsm1/Dpep1/Bod1/Pih1d1/Arl2/Angptl8/Herpud1/Gas6/Renbp/Cdkn2d/Pebp1/Reck/Vtn/Gnai2/Nle1/Ubxn1/Serpine2/Tnni3/Hpn/Klf4 |
| GO:0009144 | purine nucleoside triphosphate metabolic process | 0,406005402 | 1.940535954 | 0.01636303 | Atp5k/ATP8/Atp5j2/Atp5e/Trem2/Cox5b/COX2/Antkmt/Atp5g3/Nme2/Nme4/ATP6/Cyc1 |
| GO:0097529 | myeloid leukocyte migration | 0,383063477 | 1.964622151 | 0.01636303 | S100a14/Pgf/Trem2/Ccn3/Gp2/Ccl24/Trpv4/Pf4/Rarres2/Dusp1/Rpl13a/Nbl1/Chga/Il34/Mif/Anxa1/Mmp28/Rps19/Dpep1/Slc37a4/Ddt/Ccl11/Il17rc/Edn1/Pdgfb/Spp1/Mmp14 |
| GO:0001570 | vasculogenesis | 0,446560224 | 2.04099305 | 0.01636303 | Sox18/Wnt7b/Adm/Wt1/Aplnr/Tmem100/Fgf1/Junb/Cd34/Ackr3/Tgfbr3l/Cav1/Eng/Ramp2/Tie1/Epha2/Smo/Sox17/Hdac7 |
| GO:0044060 | regulation of endocrine process | 0,585761928 | 2.111232734 | 0.016898068 | Ren1/Retn/Gal/Lif/Inhbb/Inha/Agtr1a/C1qtnf1/Cry2/Adrb1/Gja5/Fgfr4/Gnas |
| GO:0050878 | regulation of body fluid levels | 0,309058503 | 1.759313878 | 0.016898068 | Treml1/Vkorc1/F2rl3/Anxa8/Cldn4/Aqp1/Aqp5/Trpv4/Adm/Aplnr/Tspan8/Pf4/Serping1/Entpd2/Thbd/Anxa2/Cd34/Sfn/Apoe/Zfp385a/Cav1/Aprt/Bloc1s4/Cyba/Selp/Fbln1/C1qtnf1/Proc/Oas2/Cdo1/Cela2a/F3/Adrb1/Bloc1s3/Plau/Gja5/Edn1/Pdgfb/Gas6/Gnai2/Srf/Gnas/Axl/Nr1h2 |
| GO:0030510 | regulation of BMP signaling pathway | 0,433345303 | 2.014945907 | 0.020621966 | Sfrp4/Sfrp2/Crb2/Lrp2/Chrd/Nbl1/Cav1/Kcp/Eng/Htra3/Fbxl15/Htra1/Bmper/Fstl1/Fzd1/Lemd2/Fstl3/Rgma/Fkbp8/Acvrl1/Bmp4/Wnt5a/Chrdl2/Sfrp1/Sulf1 |
| GO:0002687 | positive regulation of leukocyte migration | 0,429592103 | 1.997494479 | 0.021358099 | S100a14/Pgf/Trem2/Ccl24/Trpv4/Rarres2/Mmp9/Il34/Selp/Plvap/Ptn/Thy1/Aoc3/Zp3/Edn1/Gas6/Mmp14/Pycard/Wnt5a/Vegfb/Cmklr1/Cxcl14/Vegfa |
| GO:0008217 | regulation of blood pressure | 0,374064599 | 1.959934131 | 0.021507352 | Ren1/Asic2/Klk1b26/Klk1b9/Ier3/Adm/Rarres2/Nos3/Klk1b8/Cd34/Klk1b11/Adm2/Eng/Cyba/Ramp2/Agtr1a/Mif/Col1a2/Adrb1/Aoc3/Ptgs1/Gja5/Edn1/Pdgfb/Gas6/Acvrl1/Sod1/Hmox1/Gnas/Ace/Smtn/Tnni3/Atp1a2 |
| GO:0009199 | ribonucleoside triphosphate metabolic process | 0,41944005 | 1.944657561 | 0.021852729 | Atp5k/ATP8/Atp5j2/Atp5e/Trem2/Cox5b/COX2/Antkmt/Atp5g3/Nme2/Nme4/ATP6/Cyc1 |
| GO:0009141 | nucleoside triphosphate metabolic process | 0,386836398 | 1.920635077 | 0.021852729 | Atp5k/ATP8/Atp5j2/Atp5e/Trem2/Cox5b/COX2/Antkmt/Atp5g3/Nme2/Nme4/ATP6/Cyc1/Ndufc2/Dctpp1/Atp5d/Ldhd/Ak1/Guk1/Itpa/Ak7/Atp5g1/Atp5j/Uqcc3/Flcn/Nudt16/Ak2/Aldoa/Bcl2l1/Taz/Map2k1/Sphk2/Atp5pb/Nme6/Dguok/Atp5h/Nme1/Pkm/Ran |
| GO:0040013 | negative regulation of locomotion | 0,279004419 | 1.682581 | 0.022478944 | Timp1/Sfrp2/Ccn3/Rhob/Wnt11/Tmeff2/Angpt4/Dbn1/Igfbp5/Chrd/Vash1/Adam15/Dusp1/Sema6b/Igfbp3/Podn/Rgcc/Serpinf1/Nbl1/Apoe/Eng/Sema6c/Cygb/Fbln1/Mif/Mmp28/Tie1/Ptn/Thy1/Cldn3/Apex1/Svbp/Sema3f/Card10/Dpep1/Cd63/Ddt/Sema3g/Wnt4/Pdgfb/Col3a1/Acvrl1/Dcn/Mapk15/Reck/Srf/Wnt5a/Klf4/Arap3/Ppard/Robo4/Sfrp1/Sulf1/Egfl7/Sema3b/Hdac5/Emilin1/Miip/Spint2/Hyal2 |
| GO:0006754 | ATP biosynthetic process | 0,475684517 | 2.021930914 | 0.02476853 | Atp5k/ATP8/Atp5j2/Atp5e/Trem2/Cox5b/COX2/Antkmt/Atp5g3/ATP6/Cyc1 |
| GO:0044342 | type B pancreatic cell proliferation | 0,644738587 | 2.069627419 | 0.025520916 | Ccn3/Fmc1/Igfbp4/Igfbp5/Reg1/Nr4a1/Igfbp3 |
| GO:1901568 | fatty acid derivative metabolic process | 0,439164358 | 1.977195413 | 0.027517539 | Ptgis/Alox15/Gpx1/Cyp2s1/Dgat2/Ltc4s/Prxl2b/Sphk1/Mif/Anxa1/Gpx4/Ptges/Lypla2/Ptgs1/Dpep1/Cyp2d22/Edn1/Alox5ap/Hmgcl/Gstm2/Mgst3/Dgat1/Abhd16a/Ggt5 |
| GO:0006839 | mitochondrial transport | 0,308872941 | 1.721110673 | 0.027693182 | Atp5j2/Cox5b/Timm13/Nol3/Ier3/Antkmt/Siva1/Romo1/Chchd10/Bax/Bbc3/Tomm7/Ndufa13/Nptx1/ATP6/Cyc1/Slc25a4/Alkbh7/Bok/Tmem14c/Timm50/Smdt1/Tst/Tomm5/Bcl2l2/Timm17b/Atp5d/Tomm40l/Chchd4/Slc25a28/Slc25a22/Acaa2/Eya2/Dnlz/Slc25a33/Pink1/Slc8b1/Mpc2/Slc25a1/Slc25a38/Sfxn3/Pdcd5/Atp5j/Bnip3/Tomm22/Spg7/Slc25a29/Timm9/Bad/Slc25a23/Slc25a10/Bcl2l1/Bloc1s2/Aip/Plekhf1/Timm44/Mul1/Ppif/Srebf1/Oxa1l/Rhot2/Timm22/Atp5pb/Grpel1/Dnajc15 |
| GO:0003073 | regulation of systemic arterial blood pressure | 0,457676364 | 1.966417648 | 0.027693182 | Ren1/Asic2/Klk1b26/Klk1b9/Ier3/Adm/Rarres2/Nos3/Klk1b8/Klk1b11/Eng/Cyba/Agtr1a/Mif/Adrb1/Gja5/Edn1/Pdgfb/Gas6/Ace/Smtn/Tnni3 |
| GO:0002685 | regulation of leukocyte migration | 0,36207607 | 1.898594588 | 0.027911758 | S100a14/Pgf/Trem2/Ccn3/Ccl24/Trpv4/Ecm1/Rarres2/Dusp1/Mmp9/Nbl1/Il34/Selp/Mif/Anxa1/Mmp28/Plvap/Ptn/Thy1/Aoc3/Ddt/Zp3/Edn1/Gas6/Mmp14/Pycard/Wnt5a/Slc8b1/Smpd3/Vegfb/Cmklr1/Cxcl14/Vegfa/Emilin1/Rhog/Cd9 |
| GO:0045861 | negative regulation of proteolysis | 0,262667123 | 1.592564131 | 0.028217374 | Cryab/Spink1/Timp1/Sfrp2/Pcsk1n/Anxa8/Crb2/Slpi/Wfdc1/Pi16/Col7a1/Nol3/Gpx1/Aqp1/Ecm1/Gas1/Papln/Serping1/Wfdc2/Nr4a1/Timp2/Ins2/Plat/Ctla2a/Serpinf1/Mmp9/Sfn/Wnt9a/Serpinh1/Chac1/Cstb/Cst3/Tmem98/Serpinb6a/Ube2g2/Dpep1/Pih1d1/Derl3/Herpud1/Gas6/Renbp/Cdkn2d/Pebp1/Reck/Vtn/Nle1/Nr1h2/Ubxn1 |
| GO:0032102 | negative regulation of response to external stimulus | 0,279007533 | 1.659797608 | 0.0292687 | Hamp/Ptgis/Wfdc1/Ccn3/Nenf/Gpx2/Gpx1/Ier3/Dbn1/Tspan8/Serping1/Ins2/Ctla2a/Dusp1/Thbd/Sema6b/Anxa2/Cd34/Serpinf1/Nbl1/Metrnl/Apoe/Sema6c/Mif/C1qtnf1/Proc/Mmp28/Nfkbil1/Rps19/Cldn3/Sema3f/Htra1/Tspan6/Tff2/C1qtnf12/Grn/Ddt/Sema3g/Plau/Gps2/Wnt4/Socs3/Rgma/Pdgfb/Zfp36/Sod1/Psmb4/Fndc4/Pycard/Wnt5a/Serpine2 |
| GO:0042273 | ribosomal large subunit biogenesis | 0,458217761 | 2.009951987 | 0.0292687 | Rplp0/Rpl3/Mrpl20/Rpl35a/Eif6/Ddx28/Rpl14/Rpl26/Rrs1/Gtf3a/Rpl10a/Nle1/Nop53/Rpl12/Ppan/Mrpl11/Znhit3/Traf7/Nop16/Zfp622/Wdr74/Bop1/Malsu1/Pak1ip1/Mrto4/Rpl35/Noc2l/Rpl7l1/Rsl24d1/Ebna1bp2/Surf6/Pes1 |
| GO:0062012 | regulation of small molecule metabolic process | 0,268555873 | 1.592195421 | 0.030346476 | Snai1/Pdk4/Twist1/Gcgr/Ddit4/Trem2/COX2/Pgp/Ier3/Antkmt/Dgat2/Igfbp4/Gck/Dkk3/Fgf1/C1qtnf2/Nos3/Ins2/Igfbp3/Apoe/Cav1/Dbi/Mid1ip1/Pgam1/Mif/Cox11/Eif6/C1qtnf1/Anxa1/Pth1r/Lcmt1/Gpt2/Slc45a3/P2ry6/Ndufc2/C1qtnf12/Gpd1/Wnt4/Pdgfb/Cd320/Nupr1/Sod1/Clcn2/Fgfr4/Khk/Gnmt/Nr1h2/Dgat1/Smpd3/Rdh10/Mlycd/Ppard |
| GO:0007178 | transmembrane receptor protein serine/threonine kinase signaling pathway | 0,272034921 | 1.587632558 | 0.03078672 | Amhr2/Sfrp4/Sfrp2/Crb2/Ccn3/Dact2/Ltbp4/Inhbb/Scx/Lrp2/Chrd/Tmem100/Cdkn2b/Nbl1/Tgfbr3l/Inha/Cav1/Id1/Kcp/Gdf10/Eng/Pmepa1/Htra3/Fbxl15/Col1a2/Htra1/Bmper/Fstl1/Bambi/Fzd1/Jun/Lemd2/Fstl3/Slc39a5/Vim/Lrrc32/Lrg1/Rgma/Col3a1/Tgfb1i1/Eid2/Fkbp8/Acvrl1/Atoh8/Chst11/Bmp4/Wnt5a/Tgfb3/Smpd3/Chrdl2/Cilp/Sfrp1/Sulf1/Cav2/Zfp703/Meg3/Smad6/Emilin1/Itgb5/Bcl9l/Stoml1/Ltbp3/Megf8/Ovol2/Zyx |
| GO:0016458 | gene silencing | -0,489634399 | -1.691811532 | 0.030804002 | H1f4/H3c4/Mir200a/H2ac20/H2al1m/H3c6/H1f3/H1f5/H2ac10/H2ap/H3c1/Mirlet7c-1/Kcnq1ot1/H2ac15/H3c2/H2ac4/H2al3/H2ac6/H1f1 |
| GO:0048762 | mesenchymal cell differentiation | 0,305900002 | 1.689149764 | 0.031340739 | Tcf21/Nrtn/Snai1/Sfrp2/Gata5/Twist1/Crb2/Hand2/Wnt11/Wnt2/Mcrip1/Tmem100/Adam15/Sema6b/Rgcc/Tgfbr3l/Eng/Dact3/Sema6c/Osr1/Col1a1/Sema3f/Bambi/Smo/Loxl2/Sema3g/Wnt4/Edn1/Aldh1a2/Tgfb1i1/Phldb1/Bmp4/Wnt5a/Tgfb3/Hpn/Mad2l2/Rdh10/Sfrp1/Loxl3/Pef1/Zfp703/Sema3b/Pdcd6/Vegfa |
| GO:0009991 | response to extracellular stimulus | 0,269200761 | 1.649877026 | 0.031417566 | Hamp/Bglap3/Sfrp2/Fam107a/Pcsk1n/Pdk4/Gcgr/Nenf/Map1lc3a/Glul/Trpv4/Inhbb/Gck/Cdkn2b/Rbp1/Mmp9/Eif4ebp1/Apoe/Sesn1/Slc39a4/Higd1a/Cdkn1a/Gsdmd/Pck2/Adrb1/Fstl1/Dctpp1/Jun/Capn10/Gas2l1/Slc39a5/Kif26a/Wnt4/Arsa/Spp1/Gas6/Zfp36/Cdkn2d/Atn1/Sod1/Srf/Axl/Bhlha15/Pemt |
| GO:0019932 | second-messenger-mediated signaling | 0,265340213 | 1.566921398 | 0.032852525 | Mt1/Treml1/Mt2/Spink1/Insl3/Gcgr/Ramp3/Trem2/Aqp1/Gal/Adm/Aplnr/Fhl2/Gck/Pf4/Tmem100/Nme2/Nos3/Nfatc4/Ins2/Npr1/Lmcd1/Hint1/Ackr3/Adcy1/Ptgfr/Chga/Apoe/Sphk1/Adm2/Ramp2/Selp/Agtr1a/Pth1r/Adgrd1/Ramp1/Rasd1/P2ry6/Adrb1/Tff2/Npr2/Edn1/Homer3/Pdgfb/Cdc34/Cib1/Pebp1/Gnai2/Mapk7/Selenon/Tmem38a/Gnas/Bhlha15/Rundc3a |
| GO:0046456 | icosanoid biosynthetic process | 0,554394777 | 1.998177663 | 0.033013929 | Ptgis/Alox15/Ltc4s/Prxl2b/Sphk1/Mif/Anxa1/Ptges/Ptgs1/Edn1/Alox5ap/Mgst3 |
| GO:0048771 | tissue remodeling | 0,323375665 | 1.687410825 | 0.034834506 | Hamp/Fcgr4/Tmem119/Timp1/Pdk4/Hand2/Nol3/Lif/Clec10a/Igfbp5/Bax/Mmp2/Nos3/Dlk1/Mmp9/Cav1/P3h4/Pth1r/Ptn/Epha2/Gpr137/Ctsk/Cela1/Gja5/Spp1/Mmp14/Acvrl1/Tgm2 |
| GO:0001666 | response to hypoxia | 0,320457432 | 1.685039613 | 0.035280916 | Cryab/Ptgis/Twist1/Pgf/Ddit4/Sod3/Nol3/Aqp1/Fndc1/Mmp2/Plat/Cd34/Rgcc/Eif4ebp1/Cav1/Ajuba/Eng/Kcnk3/Higd1a/Chchd2/Plk3/Loxl2/Plau/Edn1/Slc2a8/Hmox1/Acaa2/Tgfb3/Egln2/Pink1/Vegfb/Ppard/Nop53/Prmt2/Vegfa/Sdhd/Usf1 |
| GO:0035924 | cellular response to vascular endothelial growth factor stimulus | 0,49993929 | 1.95536942 | 0.035280916 | Pgf/Hspb1/Nr4a1/Cadm4/Smoc2/Sphk1/Ramp2/Anxa1/Cd63/Pdgfb/Dcn/Vegfb/Map2k3/Vegfa/Adgra2/Dll4/Ptp4a3 |
| GO:0006633 | fatty acid biosynthetic process | 0,410693304 | 1.909619852 | 0.035368659 | Ptgis/Pdk4/Pla2g1b/Alox15/Ltc4s/Prxl2b/Sphk1/Mid1ip1/Mif/Eif6/Anxa1/Ptges/Slc45a3/Ptgs1/Edn1/Gstm2/Fasn/Nr1h2/Mlycd/Mcat/Acadvl/Elovl1/Acot7/Prkag1/Scap/Ptges2/Nr1h3/Tecr |
| GO:0071772 | response to BMP | 0,360165255 | 1.863635654 | 0.035368659 | Sfrp4/Sfrp2/Gata5/Crb2/Scx/Lrp2/Chrd/Tmem100/Nbl1/Cav1/Id1/Kcp/Eng/Htra3/Fbxl15/Htra1/Bmper/Fstl1/Fzd1/Lemd2/Fstl3/Slc39a5/Rgma/Fkbp8/Acvrl1/Bmp4/Wnt5a/Tgfb3/Smpd3/Chrdl2/Sfrp1/Sulf1 |
| GO:0071773 | cellular response to BMP stimulus | 0,360165255 | 1.863635654 | 0.035368659 | Sfrp4/Sfrp2/Gata5/Crb2/Scx/Lrp2/Chrd/Tmem100/Nbl1/Cav1/Id1/Kcp/Eng/Htra3/Fbxl15/Htra1/Bmper/Fstl1/Fzd1/Lemd2/Fstl3/Slc39a5/Rgma/Fkbp8/Acvrl1/Bmp4/Wnt5a/Tgfb3/Smpd3/Chrdl2/Sfrp1/Sulf1 |
| GO:0009205 | purine ribonucleoside triphosphate metabolic process | 0,431485289 | 2.03272119 | 0.036341296 | Atp5k/ATP8/Atp5j2/Atp5e/Trem2/Cox5b/COX2/Antkmt/Atp5g3/Nme2/Nme4/ATP6/Cyc1 |
| GO:0045598 | regulation of fat cell differentiation | 0,35632334 | 1.825918811 | 0.037637116 | Wnt10b/Ccdc85b/Sfrp2/Cebpb/Trpv4/Fndc5/Rarres2/Dlk1/Cebpa/Metrnl/Zfp385a/Jdp2/Mmp11/Medag/E2f1/Tlcd3b/Gps2/Gata2/Tgfb1i1/Lrp3/Zfp36 |
| GO:0036293 | response to decreased oxygen levels | 0,310832499 | 1.674038154 | 0.038226113 | Cryab/Ptgis/Twist1/Pgf/Ddit4/Sod3/Nol3/Aqp1/Fndc1/Mmp2/Plat/Cd34/Rgcc/Eif4ebp1/Cav1/Ajuba/Eng/Kcnk3/Higd1a/Chchd2/Plk3/Loxl2/Plau/Edn1/Slc2a8/Hmox1/Acaa2/Tgfb3/Egln2/Pink1/Vegfb/Ppard/Nop53/Prmt2/Vegfa/Sdhd/Usf1 |
| GO:0010595 | positive regulation of endothelial cell migration | 0,412329517 | 1.926642405 | 0.038594866 | Rhob/Angpt4/Hspb1/Fgf1/Emc10/Nos3/Smoc2/Anxa1/Sparc/Anxa3/Grn/Bsg/Gata2/Hdac7/Pdgfb/Cib1/Atoh8/Hmox1/Bmp4/Srpx2/Wnt5a/Map2k3/Pdcd6/Vegfa/Adgra2/Rras |
| GO:0032368 | regulation of lipid transport | 0,378528068 | 1.854170576 | 0.039388403 | Ren1/Retn/Gal/Pltp/Anxa2/Igfbp3/Apoc1/Apoe/Cav1/Dbi/Agtr1a/Mif/C1qtnf1/Ptges/Cry2/Zdhhc8/Abcg8/Gps2/Edn1/Spp1/Nr1h2/Lamtor1/Naxe/Dgat1 |
| GO:1901570 | fatty acid derivative biosynthetic process | 0,522496174 | 1.99266197 | 0.039665129 | Ptgis/Alox15/Ltc4s/Prxl2b/Sphk1/Mif/Anxa1/Ptges/Ptgs1/Edn1/Alox5ap/Hmgcl/Mgst3 |
| GO:0006690 | icosanoid metabolic process | 0,456031125 | 1.938392771 | 0.041061377 | Ptgis/Alox15/Gpx1/Cyp2s1/Ltc4s/Prxl2b/Sphk1/Mif/Anxa1/Gpx4/Ptges/Lypla2/Ptgs1/Dpep1/Cyp2d22/Edn1/Alox5ap/Gstm2/Mgst3/Abhd16a/Ggt5 |
| GO:0097193 | intrinsic apoptotic signaling pathway | 0,26303802 | 1.612106831 | 0.041061377 | Tpt1/Snai1/Phlda3/Cebpb/Ddit4/Nol3/Gpx1/Crip1/Ier3/Hspb1/Bax/Bbc3/Nfatc4/Ins2/Ndufa13/Mmp9/Sfn/Hint1/Ackr3/Zfp385a/Cav1/Chac1/Steap3/Hic1/Mif/Cdkn1a/Epha2/Bok/Rpl26/E2f1/Bcl2l2/Tmem109/Herpud1/Ifi204/Grina/Cdkn2d/Nupr1/Sod1/Hmox1/Mapk7/Pycard/Pink1/Htra2/Tmem161a/Triap1/Hras/Rps7/Rack1/Rps3/Bnip3/Park7/Syvn1/Casp9/Tmbim6/Flcn/Pias4/Bad/Mapk8ip1/Ybx3/Bcl2l1/Atf4/Ppm1f/Ubb/Clu/Plekhf1/Bcap31/Zfp622/Map2k1/Tnfrsf1a/Stk11/Ppif/Ndufs3 |
| GO:0001523 | retinoid metabolic process | 0,627062474 | 2.012886641 | 0.041410755 | Cyp2s1/Rarres2/Rbp1/Akr1b8/Rbp4/Aldh1a2/Adh1/Dgat1/Rdh10/Ppard |
| GO:0016101 | diterpenoid metabolic process | 0,627062474 | 2.012886641 | 0.041410755 | Cyp2s1/Rarres2/Rbp1/Akr1b8/Rbp4/Aldh1a2/Adh1/Dgat1/Rdh10/Ppard |
| GO:0030308 | negative regulation of cell growth | 0,328059297 | 1.712128437 | 0.041694189 | Cryab/Ccdc85b/Sfrp2/Ccn3/Pi16/Wnt11/Gas1/Wt1/Bbc3/Adam15/Ndufa13/Sema6b/Sesn1/Dact3/Sema6c/Sertad1/Cdkn1a/Sema3f/Sema3g/Gas2l1/Sox17/Apbb1/Rgma/Acvrl1/Cdkn2d/Aatk/Wnt5a/Serpine2/Ppard/Sfrp1/Cirbp/Sema3b/Cdkn2c/Hyal2/Rack1/Rgs2/Gja1/Ip6k2/Cdk5/Flcn/Cfl1/Dnajb2/Osgin1/Ulk1/Fhl1/Stk11/Mul1/Sphk2/P3h1/Ndufs3 |
| GO:0010712 | regulation of collagen metabolic process | 0,516698614 | 1.973042298 | 0.041694189 | Retn/Scx/Ccn2/Rgcc/Mfap4/Eng/Prdx5/Cygb/Vim/Wnt4/Pdgfb/Rapgef3/Bmp4/Tgfb3/Ppard/Emilin1/Arrb2/Idua/Got1/Fgfr3 |
| GO:0030324 | lung development | 0,308253997 | 1.679853536 | 0.041747066 | Tbx4/Tcf21/Mgp/Gli1/Wnt7b/Wnt2/Lif/Igfbp5/Fgf1/Nos3/Ccn2/Pkdcc/Rpl13a/Klf2/Cebpa/Rcn3/Id1/Rbp4/Sparc/Adamts2/Aldh1a2/Mmp14/Map2k2/Srf/Fgfr4/Selenon/Bmp4/Ace/Wnt5a/Tgfb3/Smpd3/Rdh10/Loxl3 |
| GO:0061041 | regulation of wound healing | 0,347283426 | 1.766913106 | 0.044106219 | Vkorc1/Insl3/Cldn4/Wfdc1/Tspan8/Serping1/Thbd/Anxa2/Tnfrsf12a/Cd34/Cadm4/Apoe/Cav1/Smoc2/Selp/C1qtnf1/Anxa1/Proc/Cldn3/Cela2a/Plau/Wnt4/Pdgfb |
| GO:0031348 | negative regulation of defense response | 0,353292298 | 1.851096404 | 0.044463234 | Hamp/Ptgis/Wfdc1/Ccn3/Gpx2/Gpx1/Ier3/Serping1/Ins2/Ctla2a/Serpinf1/Metrnl/Apoe/Proc/Rps19/Htra1/Tspan6/Tff2/C1qtnf12/Grn/Gps2/Socs3/Zfp36/Sod1/Psmb4/Fndc4/Pycard |
| GO:0045652 | regulation of megakaryocyte differentiation | -0,653038132 | -1.814498033 | 0.044932691 | H4c14/H4c12/H4c8/H4f16/H4c3/H4c1/H4c4/H4c18/H4c6 |
| GO:0030323 | respiratory tube development | 0,304068329 | 1.664646481 | 0.046242075 | Tbx4/Tcf21/Mgp/Gli1/Wnt7b/Wnt2/Lif/Igfbp5/Fgf1/Nos3/Ccn2/Pkdcc/Rpl13a/Klf2/Cebpa/Rcn3/Id1/Rbp4/Sparc/Adamts2/Aldh1a2/Mmp14/Map2k2/Srf/Fgfr4/Selenon/Bmp4/Ace/Wnt5a/Tgfb3/Smpd3/Rdh10/Loxl3 |
| GO:0006979 | response to oxidative stress | 0,243344788 | 1.450317248 | 0.046916504 | Cryab/Rnf112/Endog/Rhob/Sod3/Gpx2/Nol3/Gpx1/Reg3b/Atox1/Aqp1/Ngfr/Ngb/Ndufa6/Romo1/Hspb1/Mmp2/Nos3/Naprt/Mmp9/Psmb5/Apoe/Sphk1/Sesn1/Fbln5/Prdx5/Gpx3/Cygb/Anxa1/Gpx4/Cst3/Apex1/Plk3/Gpx8/Ptgs1/Fzd1/Jun/Ccs/Mapk13/Pycr2/Chchd4/Sod1/Hmox1/Mapk7/Selenon/Axl/Bmp4/Atp13a2/Ndufa12/Pink1/ND1/Smpd3/Htra2/Sirt3/Tmem161a/Prnp/Hyal2/Coq7/Mgat3/Rack1/Rps3/Stx2/Lonp1/Ndufs8/Bnip3/Park7/Xrcc1/Gstp1/Stx4a/Mgst1/Bcar1/Adprhl2/Ercc2/Rcan1/Cfl1/Tor1a/Atf4/Prdx4/Rela/Zfp622/Ogg1/Map2k1/Keap1/Ppif/Agap3/Arl6ip5/Txnip/Hnrnpm/Pcgf2/Txn1/Mapk3/Aifm2/Aldh2/Sigmar1 |
| GO:0007599 | hemostasis | 0,352618075 | 1.812624801 | 0.047008031 | Treml1/Vkorc1/F2rl3/Anxa8/Tspan8/Pf4/Serping1/Entpd2/Thbd/Anxa2/Cd34/Apoe/Zfp385a/Cav1/Bloc1s4/Selp/Fbln1/C1qtnf1/Proc/Cela2a/F3/Bloc1s3/Plau/Pdgfb/Gas6/Srf/Gnas/Axl |
| GO:0003156 | regulation of animal organ formation | 0,614618615 | 2.042695408 | 0.047339508 | Gata5/Hand2/Wnt11/Wnt2/Wt1/Fgf1/Gng5/Fgfr4/Bmp4/Wnt5a/Sulf1 |
| GO:0034440 | lipid oxidation | 0,428370092 | 1.877025339 | 0.047744995 | Pdk4/Twist1/Alox15/Dgat2/C1qtnf2/Dbi/Cygb/Acads/Acaa1a/Abcd4/Acaa2/Dgat1/Etfa/Echs1/Mlycd/Ppard/Tysnd1/Acadvl/Ilvbl/Etfb/Echdc2/Akt2/Crat/Eci1/Gcdh/Abcd1/Akt1/Pex2/Pex5/Cpt2/Adh5/Decr1/Lonp2 |
| GO:0030509 | BMP signaling pathway | 0,352034095 | 1.809622869 | 0.048052182 | Sfrp4/Sfrp2/Crb2/Scx/Lrp2/Chrd/Tmem100/Nbl1/Cav1/Id1/Kcp/Eng/Htra3/Fbxl15/Htra1/Bmper/Fstl1/Fzd1/Lemd2/Fstl3/Slc39a5/Rgma/Fkbp8/Acvrl1/Bmp4/Wnt5a/Tgfb3/Smpd3/Chrdl2/Sfrp1/Sulf1 |
| GO:0006636 | unsaturated fatty acid biosynthetic process | 0,569573357 | 1.945402701 | 0.048172284 | Ptgis/Alox15/Prxl2b/Sphk1/Mif/Anxa1/Ptges/Ptgs1/Edn1/Gstm2 |
| GO:0007596 | blood coagulation | 0,346969464 | 1.77798645 | 0.048409599 | Treml1/Vkorc1/F2rl3/Anxa8/Tspan8/Pf4/Serping1/Entpd2/Thbd/Anxa2/Cd34/Apoe/Cav1/Bloc1s4/Selp/Fbln1/C1qtnf1/Proc/Cela2a/F3/Bloc1s3/Plau/Pdgfb/Gas6/Srf/Gnas/Axl |
| GO:0050817 | coagulation | 0,346969464 | 1.77798645 | 0.048409599 | Treml1/Vkorc1/F2rl3/Anxa8/Tspan8/Pf4/Serping1/Entpd2/Thbd/Anxa2/Cd34/Apoe/Cav1/Bloc1s4/Selp/Fbln1/C1qtnf1/Proc/Cela2a/F3/Bloc1s3/Plau/Pdgfb/Gas6/Srf/Gnas/Axl |
| GO:0010951 | negative regulation of endopeptidase activity | 0,340492928 | 1.785420476 | 0.048851146 | Cryab/Spink1/Timp1/Sfrp2/Pcsk1n/Anxa8/Crb2/Nol3/Gpx1/Aqp1/Serping1/Nr4a1/Timp2/Serpinf1/Mmp9/Sfn/Wnt9a/Serpinh1/Serpinb6a/Dpep1/Pih1d1/Herpud1/Gas6/Renbp/Cdkn2d/Reck/Vtn/Nle1 |
| GO:0097549 | chromatin organization involved in negative regulation of transcription | -0,528843303 | -1.710323696 | 0.049779537 | H1f4/H2ac20/H2al1m/H1f3/H1f5/H2ac10/H2ap/H2ac15/H2ac4/H2al3/H2ac6/H1f1 |
