## Supplementary Table S3 for "EZH2 deletion does not impact acinar cell regeneration but restricts progression to pancreatic cancer in mice"

**Supplemental Table S3. Significant GseGO pathways enriched in *Mist1^creERT/+^KRAS^G12D^* *vs KRAS^G12D^EZH2^ΔSET^* by RNA-Sequencing**

| ID | Description | P adjust | Gene ID |
| --- | --- | --- | --- |
| GO:0001819 | positive regulation of cytokine production | 1,22E-08 | Pou2af1/Scimp/Ptpn22/Cd2/Tlr1/Adipoq/Il1rl1/Mcoln2/Sash3/Cd40/Il33/Card11/H2-Q7/Rasgrp1/Cd300c2/Ltb/Casp4/Nfam1/Cd74/Ly9/Ptprc/Cd3e/Syk/Ifi204/Ifi205/Vegfd/C3/Irf4/Anxa1/Wnt11/C5ar1/Il27ra/Hpse/Il18/Cd209d/Ccr7/Egr1/Pou2f2/Sulf1/Aim2/Lep/Lum/Il16/Flt3/Panx1/Rab7b/Cd84/Slc11a1/Cd28/Tlr9/Vtcn1/Wnt5a/Carmil2/Tlr7/Irf8/Sulf2/Gsdmd/Havcr2/Nr4a3/Tnfsf9/Gprc5b/Cd34/Cebpb/Cd1d1/Cd83/Cd274/Casp1/Ccdc88b/Sphk1/Ccr2/Mndal/Tspan6/Gpsm3/Serpine1/Il1rl2/Ifi211/Rgcc/Tlr5/Clec9a/B2m/Tlr4/Ffar3/Tlr8/Pf4/H2-T23/Tyrobp/Runx1/Ccl5/Cd6/Card9/Spon2/Panx2/Icosl/Postn/Unc93b1/Ccbe1/Trem2/Irf7/Ccr5/Isl1/Stat1/Thbs1/Abcc8/Dlk1/Cadm1/Cyp1b1/Fcer1g/H2-M3/C3ar1/P2rx7 |
| GO:0002250 | adaptive immune response | 1,22E-08 | Ighg2b/Ighv1-53/Ighv11-2/Ighv1-55/Ighg3/Ighg2c/Lax1/Igkc/Fcer2a/Jchain/Ighd/Siglecg/Cr2/Igkv5-48/Cd19/Alox15/Cd79b/Klhl6/Cd4/Gadd45g/Il7r/Slamf7/Ighm/Cd79a/Il1rl1/Itk/Mcoln2/Btla/Iglc2/Masp2/Sash3/H2-DMb1/Cd40/Il33/Exo1/Igha/Btk/Myo1g/H2-Q7/Fcgr3/Samsn1/H2-Q6/Spn/Cd74/Ly9/H2-Eb1/Ptprc/Cd3e/Syk/Camk4/Fcgr2b/Trbc2/C3/H2-Aa/Irf4/Anxa1/H2-DMa/C4b/Cd48/Il27ra/H2-Ab1/Dpp4/Il18/Stx11/Igkv4-55/Ccr7/Pirb/Pou2f2/Serpina3g/Fcgr1/Clec4a2/C1qc/Pla2g4a/Cd84/Slc11a1/Cd28/C1qb/Cd86/Vtcn1/Vsir/Cd55/Adgre1/Serping1/Havcr2/Cd1d1/Fgl1/Cd274/Ccr2/Arg2/Lat2/C1s1/C1ra/Nfkbid/Parp3/Tnfrsf13b/Tnfrsf21/Pik3cd/Prkcb/C2/B2m/Hspa8/Cd44/Ephb6/Icam1/Adcy7/H2-T23/Apcs/Gpr183/Azgp1/Raet1e/Skap1/Tec/Hfe/Rorc/Tnfsf13b/Icosl/Dclre1c/Inpp5d/H2-T10/Unc93b1/Tnfrsf1b/Bach2/Irf7/Emp2 |
| GO:0002253 | activation of immune response | 1,22E-08 | Stap1/Ighg2b/Ighv1-53/Ighv11-2/Ighv1-55/Ighg3/Ighg2c/Cfd/Lax1/Igkc/Ptpn22/Ighd/Cr2/Ms4a1/Cd19/Cd79b/Zbp1/Klhl6/Cd22/Vsig4/Cd5l/Ighm/Cd79a/Itk/Ifi209/Blk/Iglc2/Masp2/Igha/Fyb2/Card11/Myo1g/Lck/Nfam1/Spn/Ptprc/Cd3e/Cfp/Syk/Fcgr2b/Ifi204/Ifi205/Trbc2/C3/C5ar1/C4b/Ccr7/Aim2/Nckap1l/Lgals3/C1qc/Cd28/C1qb/Tlr9/Fcna/Vtcn1/Carmil2/Cd55/Serping1/Thy1/Nr4a3/Cd300a/Mill2/Mndal/Lat2/Themis2/Ifi211/C1s1/C1ra/Rgcc/Lcp2/Nfkbid/Tnfrsf21/Cfh/Prkcb/C2/Tlr4/Fyb/Cacnb3/Apcs/Cd38/Sh2b2/Skap1/Lpxn/Tec/Vav3/Icosl |
| GO:0002377 | immunoglobulin production | 1,22E-08 | Igkv8-16/Igkv8-30/Igkv14-111/Igkv5-39/Igkv3-5/Iglv1/Lax1/Igkv6-23/Ighd/Siglecg/Igkv10-96/Igkv5-48/Mzb1/Cd22/Igkv4-91/Il7r/Ighm/Sash3/Cd40/Il33/Exo1/Igkv9-120/Btk/Igkv3-4/Card11/Igkv14-126/Ptprc/Cd37/Fcgr2b/Il27ra/H2-Ab1/Igkv4-55/Pou2f2/Igkv12-46/Cd28/Tlr9/Cd86 |
| GO:0002429 | immune response-activating cell surface receptor signaling pathway | 1,22E-08 | Stap1/Ighg2b/Ighv1-53/Ighv11-2/Ighv1-55/Ighg3/Ighg2c/Lax1/Igkc/Ptpn22/Ighd/Ms4a1/Cd19/Cd79b/Klhl6/Cd22/Ighm/Cd79a/Itk/Blk/Iglc2/Igha/Fyb2/Card11/Myo1g/Lck/Nfam1/Spn/Ptprc/Cd3e/Syk/Fcgr2b/Trbc2/C5ar1/Ccr7/Nckap1l/Lgals3/Cd28/Vtcn1/Carmil2/Thy1/Nr4a3/Cd300a/Mill2/Lat2/Themis2/Lcp2/Nfkbid/Tnfrsf21/Prkcb/Fyb/Cacnb3/Cd38/Sh2b2/Skap1/Lpxn/Tec/Vav3/Icosl |
| GO:0002440 | production of molecular mediator of immune response | 1,22E-08 | Igkv8-16/Igkv8-30/Igkv14-111/Igkv5-39/Igkv3-5/Iglv1/Scimp/Lax1/Ptpn22/Igkv6-23/Ighd/Siglecg/Igkv10-96/Igkv5-48/Mzb1/Cd22/Igkv4-91/Il7r/Ighm/Sash3/Cd40/Il33/Exo1/Igkv9-120/Btk/Igkv3-4/Card11/Cd74/Igkv14-126/Ptprc/Cd37/Fcgr2b/Il27ra/H2-Ab1/Irak3/Il18/Igkv4-55/Pou2f2/Lacc1/Slc11a1/Igkv12-46/Cd28/Tlr9/Cd86/Vsir/Wnt5a/Nr4a3/Gprc5b/Ccr2/Parp3/Itm2a/B2m/Tlr4/Ffar3/H2-T23/Igkv3-2/Hmox1/Spon2/Hfe/Tnfsf13b/Icosl/Igkv6-32/Tnfrsf1b |
| GO:0002443 | leukocyte mediated immunity | 1,22E-08 | Stap1/Ighg2b/Ighv1-53/Ighv11-2/Ighv1-55/Ighg3/Ighg2c/Scimp/Igkc/Fcer2a/Ighd/Cr2/Cd19/Il7r/Ighm/Rac2/Blk/Iglc2/Masp2/Sash3/Cd40/Exo1/Igha/Btk/Myo1g/H2-Q7/Fcgr3/Dnase1l3/Rasgrp1/H2-Q6/Spn/Cd74/Coro1a/Ptprc/Syk/Fcgr2b/Trbc2/C3/H2-DMa/C4b/Itgam/Il27ra/H2-Ab1/Dpp4/Il18/Myo1f/Stx11/Pirb/Pou2f2/Fcgr1/Lep/C1qc/Cd84/Slc11a1/Cd28/C1qb/Tlr9/Vsir/Cd55/Serping1/Havcr2/Nr4a3/Cd1d1/Cd300a/Ncf1/Ccr2/Lat2/C1s1/C1ra/Parp3/C2/B2m/Hspa8/Ephb6/Tlr4/Icam1/H2-T23/Tyrobp/Apcs/Hmox1/Azgp1/Raet1e/Spon2/Hfe/Icosl/Inpp5d/H2-T10/Tnfrsf1b/Irf7/Emp2/Anxa3/Batf/Itgb2/Paxip1/Cadm1/Fcer1g/Fas/Tap1/H2-M3/P2rx7/Vav1 |
| GO:0002449 | lymphocyte mediated immunity | 1,22E-08 | Ighg2b/Ighv1-53/Ighv11-2/Ighv1-55/Ighg3/Ighg2c/Igkc/Fcer2a/Ighd/Cr2/Cd19/Il7r/Ighm/Iglc2/Masp2/Sash3/Cd40/Exo1/Igha/Btk/Myo1g/H2-Q7/Fcgr3/Rasgrp1/H2-Q6/Spn/Cd74/Coro1a/Ptprc/Fcgr2b/Trbc2/C3/H2-DMa/C4b/Il27ra/H2-Ab1/Dpp4/Il18/Stx11/Pirb/Pou2f2/Fcgr1/Lep/C1qc/Slc11a1/Cd28/C1qb/Vsir/Cd55/Serping1/Havcr2/Cd1d1/Ccr2/C1s1/C1ra/Parp3/C2/B2m/Hspa8/Ephb6/Icam1/H2-T23/Apcs/Azgp1/Raet1e/Hfe/Icosl/Inpp5d/H2-T10/Tnfrsf1b/Irf7/Emp2/Batf/Paxip1/Cadm1/Fcer1g/Fas/Tap1/H2-M3/P2rx7/Vav1/Muc4/Was/Arg1 |
| GO:0002460 | adaptive immune response based on somatic recombination of immune receptors built from immunoglobulin superfamily domains | 1,22E-08 | Ighg2b/Ighv1-53/Ighv11-2/Ighv1-55/Ighg3/Ighg2c/Igkc/Fcer2a/Ighd/Cr2/Cd19/Klhl6/Cd4/Gadd45g/Il7r/Ighm/Il1rl1/Iglc2/Masp2/Sash3/Cd40/Il33/Exo1/Igha/Btk/Myo1g/H2-Q7/Fcgr3/H2-Q6/Spn/Cd74/Ly9/Ptprc/Fcgr2b/Trbc2/C3/Irf4/Anxa1/H2-DMa/C4b/Il27ra/H2-Ab1/Dpp4/Il18/Stx11/Ccr7/Pirb/Pou2f2/Fcgr1/C1qc/Pla2g4a/Slc11a1/Cd28/C1qb/Vsir/Cd55/Serping1/Havcr2/Cd1d1/Cd274/Ccr2/C1s1/C1ra/Nfkbid/Parp3/C2/B2m/Hspa8/Ephb6/Icam1/H2-T23/Apcs/Azgp1/Raet1e/Hfe/Rorc/Tnfsf13b/Icosl/Inpp5d/H2-T10/Tnfrsf1b/Bach2/Irf7/Emp2 |
| GO:0002521 | leukocyte differentiation | 1,22E-08 | Pou2af1/Igkc/Ptpn22/Cr2/Ms4a1/Cd19/Pck1/Spib/Cd79b/Cd4/Gadd45g/Rhoh/Il7r/Batf3/Ighm/Cd79a/Adipoq/Itk/Dock2/Blk/Sash3/Ccr1/Fos/Ikzf1/Btk/Card11/Lck/Rasgrp1/Nfam1/Spn/Cd74/Ly9/Ptprc/Cd3e/Syk/Camk4/Spi1/H2-Aa/Irf4/Anxa1/Runx3/Lrrc17/H2-DMa/Myb/Itgam/H2-Ab1/Il2rg/Il2ra/Il18/Ikzf3/Ccr7/Pirb/Egr1/Pou2f2/Trf/Lep/Nckap1l/C1qc/Flt3/Car2/Axl/Cd28/Tlr9/Dock10/Vsir/Hcls1/Carmil2/Runx2/Satb1/Tnfsf9/Cebpb/Cd1d1/Bmp4/Cd83/Ccr2/Ntrk1/Itgb3/Rassf2/Pparg/Ubash3b/Trpm2/Myc/Il1rl2/Lif/Snx10/Nfkbid/Mfng/Lyl1/Nrros/Itm2a/B2m/Cd44/Il3ra/Kitl/Pf4/Fzd9/Tyrobp/Runx1/Apcs/Ccl5/Trib1/Mpzl2/Gpr183/Prex1/Gpc3/Anxa2/Csf1r/Erbb2/Rorc/Dclre1c/Inpp5d/Pir/Zbtb16/Trem2/Tgfbr2/Zeb1/Batf/Jun/Lepr/Dlk1/Csf1/Fcer1g/Fas/H2-M3/Pla2g2d |
| GO:0002694 | regulation of leukocyte activation | 1,22E-08 | Stap1/Ighg2b/Ighv1-53/Ighv11-2/Ighv1-55/Ighg3/Ighg2c/Lax1/Igkc/Ptpn22/Slfn1/Ighd/Siglecg/Cd19/Pck1/Cd4/Mzb1/Cd22/Vsig4/Rhoh/Il7r/Slamf7/Ighm/Bank1/Il1rl1/Rac2/Btla/Blk/Iglc2/Sash3/Cd40/Il33/Ikzf1/Igha/Btk/Card11/Tspan32/Lck/Samsn1/Rasgrp1/Nfam1/Lgals7/Spn/Cd74/Coro1a/Ptprc/Cd3e/Syk/Cd37/Fcgr2b/Trbc2/Tbc1d10c/H2-Aa/Anxa1/Runx3/Rasal3/H2-DMa/Myb/Itgam/Il27ra/H2-Ab1/Dpp4/Il2rg/Il2ra/Il18/Ikzf3/Vcam1/Ccr7/Slc4a1/Lep/Ptpre/Nckap1l/Lgals3/Pla2g4a/Il16/Ripor2/Flt3/Axl/Cd84/Cd28/Tlr9/Cd86/Vtcn1/Mfhas1/Vsir/Sh3kbp1/Wnt5a/Carmil2/Dock8/Thy1/Havcr2/Nr4a3/Tnfsf9/Cebpb/Cd1d1/Cd300a/Bmp4/Fgl1/Cd83/Cd274/Ccdc88b/Sphk1/Ccr2/Adora2a/Lrrk2/Arg2/Pparg/Il1rl2/Nfkbid/Parp3/Tnfrsf13b/Efnb3/Tnfrsf21/Syt11/Cd44/Ephb6/Tlr4/H2-T23/Tyrobp/Runx1/Cd38/Ccl5/Gpr183/Cd6/Hmox1/Gsn/Erbb2/Hfe/Rorc/Tnfsf13b/Vav3/Icosl/Inpp5d/Pla2g2f/Tnfrsf1b/Zbtb16/Trem2/Tgfbr2/Plscr1/Zeb1/Thbs1/Cav1/Itgal/Itgb2/Irs2/Paxip1/Bst1/Fcer1g/Fas/Sirpa/H2-M3/Pla2g2d/Gal |
| GO:0002696 | positive regulation of leukocyte activation | 1,22E-08 | Stap1/Ighg2b/Ighv1-53/Ighv11-2/Ighv1-55/Ighg3/Ighg2c/Igkc/Ptpn22/Ighd/Pck1/Cd4/Rhoh/Il7r/Ighm/Il1rl1/Iglc2/Sash3/Cd40/Il33/Ikzf1/Igha/Card11/Lck/Rasgrp1/Spn/Cd74/Coro1a/Ptprc/Cd3e/Syk/Trbc2/H2-Aa/Anxa1/Runx3/Rasal3/H2-DMa/Myb/Itgam/Il27ra/H2-Ab1/Dpp4/Il2rg/Il2ra/Il18/Vcam1/Ccr7/Slc4a1/Lep/Nckap1l/Pla2g4a/Il16/Axl/Cd28/Tlr9/Cd86/Vtcn1/Vsir/Sh3kbp1/Wnt5a/Carmil2/Dock8/Thy1/Havcr2/Nr4a3/Tnfsf9/Cd1d1/Cd83/Cd274/Ccdc88b/Ccr2/Lrrk2/Il1rl2/Nfkbid/Efnb3/Ephb6/Tlr4/H2-T23/Tyrobp/Runx1/Cd38/Ccl5/Gpr183/Cd6/Tnfsf13b/Vav3/Icosl/Inpp5d/Zbtb16/Trem2/Tgfbr2/Thbs1/Cav1/Itgal/Itgb2/Irs2/Paxip1/Bst1/Fcer1g/Sirpa/H2-M3 |
| GO:0002697 | regulation of immune effector process | 1,22E-08 | Stap1/Ighg2b/Scimp/Ptpn22/Fcer2a/Siglecg/Pck1/Mzb1/Cd22/Vsig4/Cd5l/Il7r/Ighm/Rac2/Blk/Sash3/Cd40/Il33/Btk/Tspan32/H2-Q7/Fcgr3/Dnase1l3/Rasgrp1/H2-Q6/Spn/Cd74/Ptprc/Syk/Cd37/Fcgr2b/C3/Anxa1/Myb/Itgam/Il27ra/Dpp4/Irak3/Il2ra/Il18/Ccr7/Fcgr1/Aim2/Lep/Apobec3/Lgals3/Cd84/Cd28/Tlr9/Vsir/Wnt5a/Cd55/Serping1/Havcr2/Nr4a3/Gprc5b/Cd1d1/Cd300a/Mill2/Ccr2/Tspan6/Nfkbid/Parp3/Cfh/B2m/Hspa8/Tlr4/Ffar3/H2-T23/Tyrobp/Htra1/Ccl5/Hmox1/Azgp1/Spon2/Hfe/H2-T10/Tnfrsf1b/Stat1/Itgb2/Paxip1/Cadm1/Tgfb3/Fcer1g/Vpreb3/Tap1/H2-M3/P2rx7/Vav1/Muc4/Was/Arg1/H2-T24/Il12rb1/Dtx3l/Hmces/Hk1/Tnfsf13/Gpatch3/Il18r1/Lag3/Jak3/Tgfb2/Ffar2/Tek/Arrb2/Clec2d/C1qbp/Gfer/Dusp10/Sting1/Stx7/Foxf1/Unc13d/Lyn |
| GO:0002757 | immune response-activating signal transduction | 1,22E-08 | Stap1/Ighg2b/Ighv1-53/Ighv11-2/Ighv1-55/Ighg3/Ighg2c/Lax1/Igkc/Ptpn22/Ighd/Ms4a1/Cd19/Cd79b/Klhl6/Cd22/Ighm/Cd79a/Itk/Blk/Iglc2/Igha/Fyb2/Card11/Myo1g/Lck/Nfam1/Spn/Ptprc/Cd3e/Syk/Fcgr2b/Trbc2/C5ar1/Ccr7/Nckap1l/Lgals3/Cd28/Vtcn1/Carmil2/Thy1/Nr4a3/Cd300a/Mill2/Lat2/Themis2/Lcp2/Nfkbid/Tnfrsf21/Prkcb/Tlr4/Fyb/Cacnb3/Cd38/Sh2b2/Skap1/Lpxn/Tec/Vav3/Icosl |
| GO:0002764 | immune response-regulating signaling pathway | 1,22E-08 | Stap1/Ighg2b/Ighv1-53/Ighv11-2/Ighv1-55/Ighg3/Ighg2c/Lax1/Igkc/Ptpn22/Ighd/Ms4a1/Cd19/Cd79b/Klhl6/Cd22/Ighm/Cd79a/Itk/Btla/Blk/Iglc2/Cd40/Igha/Fyb2/Card11/Myo1g/Lck/Nfam1/Spn/Ptprc/Cd3e/Syk/Fcgr2b/Trbc2/C5ar1/Ccr7/Nckap1l/Lgals3/Cd28/Vtcn1/Carmil2/Thy1/Nr4a3/Cd300a/Mill2/Lat2/Themis2/Lcp2/Nfkbid/Tnfrsf21/Prkcb/Tlr4/Fyb/Cacnb3/Cd38/Sh2b2/Skap1/Lpxn/Tec/Vav3/Icosl |
| GO:0002768 | immune response-regulating cell surface receptor signaling pathway | 1,22E-08 | Stap1/Ighg2b/Ighv1-53/Ighv11-2/Ighv1-55/Ighg3/Ighg2c/Lax1/Igkc/Ptpn22/Ighd/Ms4a1/Cd19/Cd79b/Klhl6/Cd22/Ighm/Cd79a/Itk/Btla/Blk/Iglc2/Cd40/Igha/Fyb2/Card11/Myo1g/Lck/Nfam1/Spn/Ptprc/Cd3e/Syk/Fcgr2b/Trbc2/C5ar1/Ccr7/Nckap1l/Lgals3/Cd28/Vtcn1/Carmil2/Thy1/Nr4a3/Cd300a/Mill2/Lat2/Themis2/Lcp2/Nfkbid/Tnfrsf21/Prkcb/Fyb/Cacnb3/Cd38/Sh2b2/Skap1/Lpxn/Tec/Vav3/Icosl |
| GO:0006909 | phagocytosis | 1,22E-08 | Stap1/Ighg2b/Ighv1-53/Ighv11-2/Ighv1-55/Ighg3/Ighg2c/Igkc/Ighd/Alox15/Marco/Rhoh/Ighm/Adipoq/Rac2/Dock2/Iglc2/Il2rb/Igha/Cd209b/Msr1/Myo1g/Fcgr3/Ncf2/Coro1a/Ptprc/Syk/Fcgr2b/Trbc2/C3/Anxa1/Itgam/Il2rg/Ccr7/Fcgr1/Pld4/Ncf4/Lep/Nckap1l/Axl/Rab7b/Slc11a1/Arhgap25/Irf8/Cd300a/Colec12/Sphk1/Ccr2/Itgb3/Pparg/Mst1r/Syt11/Hck/C2/Bin2/Hspa8/Tlr4/Tyrobp/Elmo1/Rab31/Gsn/Siglece/Spon2/Rab20/Trem2/Anxa3/Xkr4/Thbs1/Itgal/Itgb2/Lepr/Pros1/Gulp1/Cnn2/Fcer1g/Sirpa/P2rx7/Vav1 |
| GO:0006911 | phagocytosis, engulfment | 1,22E-08 | Stap1/Ighg2b/Ighv1-53/Ighv11-2/Ighv1-55/Ighg3/Ighg2c/Igkc/Ighd/Alox15/Marco/Rhoh/Ighm/Rac2/Iglc2/Igha/Msr1/Fcgr3/Fcgr2b/Trbc2/C3/Itgam/Fcgr1/Nckap1l/Arhgap25/Cd300a/Pparg/Bin2/Elmo1/Rab31/Gsn/Siglece/Trem2/Xkr4/Thbs1/Itgb2/Gulp1/Fcer1g/Sirpa |
| GO:0006959 | humoral immune response | 1,22E-08 | Ighg2b/Ighv1-53/Ighv11-2/Ighv1-55/Ighg3/Ighg2c/Cfd/Igkc/Fcer2a/Jchain/Ighd/Cr2/Vsig4/Cd5l/Ighm/Reg2/Iglc2/Masp2/Exo1/Cxcl13/Wfdc17/Igha/Slpi/Ptprc/Cfp/Cd37/Fcgr2b/Trbc2/C3/Reg3a/H2-DMa/C4b/H2-Ab1/Ccr7/Trf/Npy/Reg3b/Lgals3/C1qc/Ccl11/Rarres2/C1qb/Vip/Fcna/Cd55/Serping1/Ccr2/Ppbp/C1s1/C1ra/Rgcc/Tnfrsf21/Cfh/C2/B2m/Pf4/H2-T23/Apcs/Gpr183/Spon2/Reg1 |
| GO:0007159 | leukocyte cell-cell adhesion | 1,22E-08 | Sell/Lax1/Ptpn22/Slfn1/Chst4/Pck1/Cd4/Vsig4/Rhoh/Il7r/Rac2/Btla/Sash3/Ikzf1/Card11/Lck/Rasgrp1/Lgals7/Spn/Cd74/Coro1a/Ptprc/Cd3e/Syk/Itgb7/H2-Aa/Anxa1/Runx3/Rasal3/H2-DMa/Myb/Itgam/Il27ra/H2-Ab1/Dpp4/Il2rg/Il2ra/Il18/Vcam1/Ccr7/Slc4a1/Selplg/Lep/Nckap1l/Lgals3/Ripor2/Cd28/Cd86/Vtcn1/Vsir/Carmil2/Dock8/Thy1/Havcr2/Nr4a3/Tnfsf9/Cebpb/Cd1d1/Cd300a/Bmp4/Fgl1/Cd83/Cd274/Ccdc88b/Ccr2/Adora2a/Arg2/Il1rl2/Nfkbid/Efnb3/Tnfrsf21/Cd44/Ephb6/Icam1/H2-T23/Runx1/Stk10/Ccl5/Cd6/Skap1/Erbb2/Hfe/Tnfsf13b/Icosl/Pla2g2f/Zbtb16/Tgfbr2/Itga4/Fut4/Cav1/Itgal/Itgb2/Fermt3/Sirpa/H2-M3/Pla2g2d |
| GO:0009617 | response to bacterium | 1,22E-08 | Cyp2b10/Stap1/Coch/Ighg2b/Ighv1-53/Ighv11-2/Ighv1-55/Saa3/Ighg3/Ighg2c/Cyp2e1/Cfd/Scimp/Igkc/Ptpn22/Jchain/Car3/Ighd/Ms4a1/Pck1/Cd79b/Tlr1/Klhl6/Cd4/Il7r/Ighm/Lyz1/Adipoq/Bank1/Cd52/Iglc2/Plac8/Cxcl13/Wfdc17/Igha/Gpm6a/Scd1/Slpi/Cd209b/Upk1b/Pygl/Ncf2/Spn/Mt2/Gbp8/Syk/Fcgr2b/Ifi204/Ifi205/Trbc2/Irf5/Vegfd/Thrsp/C3/Lyz2/C5ar1/Fkbp5/Tmem255a/Il27ra/Irak3/Ly86/Il18/Cd209d/Myo1f/Ikzf3/Ccr7/Cd180/Fcgr1/Trf/Npy/Reg3b/Hp/Axl/Rarres2/Gas5/Cd84/Gbp2/Slc11a1/Nlrc5/Vim/Vip/Tlr9/Cd86/Wnt5a/Sp110/Fer1l6/Gstm3/Cd68/Irf8/Lilrb4a/Gsdmd/Havcr2/Zfp36/Cebpb/Ptgfr/Colec12/Ccdc80/Ncf1/Cd274/Casp1/Bnip3/Psmb9/Iigp1/Mndal/Usp18/Arg2/Ppbp/Serpine1/Ifi211/Alpk1/Oas2/Ifit1/Syt11/Tlr5/Bmp6/Hck/Abcd2/Mrc1/B2m/Tlr4/Pf4/Bmp2/Scn7a/B3galt5/Lypd8/H2-T23/Slfn2/Lcn2/Mpeg1/Prkd1/Ccl5/Ggt5/Ptgir/Trib1/Cd6/Card9/Gpc3/Spon2/Pmp22/Tnfrsf1b/Nexn/Herc6/Plscr4/Trem2/Noct/Anxa3/Plscr1/Trim30a/Stat1/Defb1/Cav1/Adamts5/H2bc6/Resf1/Rpl13a/Serpina3n/Fcer1g/Sirpa/H2-M3 |
| GO:0010324 | membrane invagination | 1,22E-08 | Stap1/Ighg2b/Ighv1-53/Ighv11-2/Ighv1-55/Ighg3/Ighg2c/Igkc/Ighd/Alox15/Marco/Rhoh/Ighm/Rac2/Iglc2/Igha/Msr1/Aurkb/Fcgr3/Fcgr2b/Trbc2/C3/Itgam/Fcgr1/Nckap1l/Arhgap25/Cd300a/Pparg/Syt11/Bin2/Elmo1/Rab31/Gsn/Siglece/Trem2/Spire2/Spire1/Xkr4/Thbs1/Itgb2/Gulp1/Fcer1g/Sirpa |
| GO:0016064 | immunoglobulin mediated immune response | 1,22E-08 | Ighg2b/Ighv1-53/Ighv11-2/Ighv1-55/Ighg3/Ighg2c/Igkc/Fcer2a/Ighd/Cr2/Cd19/Ighm/Iglc2/Masp2/Cd40/Exo1/Igha/Btk/Fcgr3/Cd74/Ptprc/Fcgr2b/Trbc2/C3/H2-DMa/C4b/Il27ra/H2-Ab1/Pou2f2/Fcgr1/C1qc/Cd28/C1qb/Cd55/Serping1/C1s1/C1ra/Parp3/C2/H2-T23/Apcs |
| GO:0019724 | B cell mediated immunity | 1,22E-08 | Ighg2b/Ighv1-53/Ighv11-2/Ighv1-55/Ighg3/Ighg2c/Igkc/Fcer2a/Ighd/Cr2/Cd19/Ighm/Iglc2/Masp2/Cd40/Exo1/Igha/Btk/Fcgr3/Cd74/Ptprc/Fcgr2b/Trbc2/C3/H2-DMa/C4b/Il27ra/H2-Ab1/Pirb/Pou2f2/Fcgr1/C1qc/Cd28/C1qb/Cd55/Serping1/C1s1/C1ra/Parp3/C2/H2-T23/Apcs |
| GO:0022407 | regulation of cell-cell adhesion | 1,22E-08 | Lax1/Ptpn22/Slfn1/Alox15/Pck1/Cd4/Vsig4/Rhoh/Il7r/Adipoq/Btla/Blk/Sash3/Cxcl13/Ikzf1/Card11/Lck/Rasgrp1/Lgals7/Spn/Cd74/Coro1a/Ptprc/Cd3e/Syk/H2-Aa/Anxa1/Runx3/Rasal3/H2-DMa/Myb/Il27ra/H2-Ab1/Dpp4/Il2rg/Il2ra/Il18/Vcam1/Ccr7/Efna5/Slc4a1/Lep/Nckap1l/Lgals3/Akna/Ripor2/Pdpn/Cd28/Ephb3/Cd86/Vtcn1/Vsir/Wnt5a/Carmil2/Dock8/Zdhhc2/Thy1/Havcr2/Tenm3/Nr4a3/Tnfsf9/Cebpb/Cd1d1/Cd300a/Bmp4/Fgl1/Cd83/Cd274/Ccdc88b/Ccr2/Adora2a/Arg2/Ubash3b/C1qtnf1/Il1rl2/Rgcc/Nfkbid/Efnb3/Tnfrsf21/Bmp6/Tnr/Magi2/Cd44/Ephb6/Icam1/Bmp2/H2-T23/Runx1/Ccl5/Cd6/Skap1/Erbb2/Hfe/Tnfsf13b/Icosl/Pla2g2f/Adrb2/Zbtb16/Ccr5/Tgfbr2/Itga4/Fut4/Cav1/Itgal/Itgb2/Fermt3/Sirpa/H2-M3/Pla2g2d/Plpp3/Muc4/Arg1/Adk/Spta1/Il12rb1 |
| GO:0030098 | lymphocyte differentiation | 1,22E-08 | Pou2af1/Igkc/Ptpn22/Cr2/Ms4a1/Cd19/Pck1/Cd79b/Cd4/Gadd45g/Rhoh/Il7r/Ighm/Cd79a/Itk/Dock2/Sash3/Ikzf1/Btk/Card11/Lck/Rasgrp1/Nfam1/Spn/Cd74/Ly9/Ptprc/Cd3e/Syk/Spi1/H2-Aa/Irf4/Anxa1/Runx3/H2-DMa/Myb/H2-Ab1/Il2rg/Il2ra/Il18/Ikzf3/Ccr7/Egr1/Pou2f2/Lep/Nckap1l/Flt3/Axl/Cd28/Tlr9/Dock10/Vsir/Carmil2/Runx2/Satb1/Tnfsf9/Cd1d1/Bmp4/Cd83/Ccr2/Ntrk1/Il1rl2/Nfkbid/Mfng/Lyl1/Itm2a/B2m/Cd44/Fzd9/Runx1/Mpzl2/Gpr183/Prex1/Erbb2/Rorc/Dclre1c/Inpp5d |
| GO:0032943 | mononuclear cell proliferation | 1,22E-08 | Ptpn22/Slfn1/Ighd/Siglecg/Cr2/Cd19/Cd4/Mzb1/Cd22/Vsig4/Il7r/Ighm/Cd79a/Rac2/Btla/Dock2/Blk/Sash3/Cd40/Btk/Card11/Rasgrp1/Lgals7/Spn/Cd74/Coro1a/Ptprc/Cd3e/Syk/Fcgr2b/H2-Aa/Anxa1/Rasal3/Itgam/Il27ra/H2-Ab1/Il2ra/Il18/Ikzf3/Vcam1/Ccr7/Cd180/Slc4a1/Lep/Nckap1l/Lgals3/Ripor2/Flt3/Slc11a1/Cd28/Tlr9/Cd86/Vtcn1/Vsir/Carmil2/Dock8/Lilrb4a/Satb1/Havcr2/Tnfsf9/Cebpb/Cd1d1/Cd300a/Elf4/Bmp4/Cd274/Ccdc88b/Ccr2/Arg2/Tnfrsf13b/Tnfrsf21/Cd44/Ephb6/Tlr4/H2-T23/Tyrobp/Cd38/Ccl5/Gpr183/Cd6/Erbb2/Tnfsf13b/Vav3/Icosl/Inpp5d/Pla2g2f/Tnfrsf1b/Cxcr4/Tgfbr2/Itgal/Itgb2/Irs2/Bst1/Csf1/H2-M3/Pla2g2d/Gal/P2rx7 |
| GO:0032944 | regulation of mononuclear cell proliferation | 1,22E-08 | Ptpn22/Slfn1/Ighd/Siglecg/Cd4/Mzb1/Cd22/Vsig4/Ighm/Rac2/Btla/Blk/Sash3/Cd40/Btk/Card11/Lgals7/Spn/Cd74/Coro1a/Ptprc/Cd3e/Syk/Fcgr2b/H2-Aa/Anxa1/Rasal3/Il27ra/H2-Ab1/Il2ra/Il18/Ikzf3/Vcam1/Ccr7/Slc4a1/Lep/Nckap1l/Lgals3/Ripor2/Cd28/Tlr9/Cd86/Vtcn1/Vsir/Carmil2/Havcr2/Tnfsf9/Cebpb/Cd1d1/Cd300a/Bmp4/Cd274/Ccdc88b/Ccr2/Arg2/Tnfrsf13b/Tnfrsf21/Cd44/Tlr4/H2-T23/Tyrobp/Cd38/Ccl5/Gpr183/Cd6/Erbb2/Tnfsf13b/Vav3/Icosl/Inpp5d/Pla2g2f/Tnfrsf1b/Tgfbr2/Itgal/Irs2/Bst1/Csf1/H2-M3/Pla2g2d/Gal/Arg1/Adk/Spta1/Il12rb1 |
| GO:0042110 | T cell activation | 1,22E-08 | Lax1/Ptpn22/Slfn1/Cd2/Pck1/Treml2/Cd4/Gadd45g/Vsig4/Rhoh/Il7r/Itk/Rac2/Btla/Dock2/Sash3/Ikzf1/Card11/Lck/Rasgrp1/Lgals7/Spn/Cd74/Ly9/Coro1a/Ptprc/Jaml/Cd3e/Syk/H2-Aa/Irf4/Anxa1/Wdfy4/Runx3/Rasal3/H2-DMa/Myb/Cd48/Itgam/Il27ra/H2-Ab1/Dpp4/Il2rg/Il2ra/Il18/Stx11/Vcam1/Ccr7/Egr1/Slc4a1/Lep/Clec4a2/Nckap1l/Lgals3/Apbb1ip/Ripor2/Flt3/Slc11a1/Cd28/Cd86/Vtcn1/Vsir/Carmil2/Dock8/Runx2/Lilrb4a/Satb1/Thy1/Havcr2/Tnfsf9/Cebpb/Cd1d1/Cd300a/Elf4/Bmp4/Fgl1/Cd83/Mill2/Cd274/Ccdc88b/Ccr2/Adora2a/Arg2/Il1rl2/Nfkbid/Efnb3/Tnfrsf21/B2m/Cd44/Ephb6/Icam1/H2-T23/Runx1/Ccl5/Mpzl2/Gpr183/Cd6/Prex1/Gsn/Erbb2/Hfe/Rorc/Tnfsf13b/Icosl/Pla2g2f/Tnfrsf1b/Zbtb16/Cxcr4/Tgfbr2/Zeb1/Batf/Cav1/Itgal/Itgb2/Lepr/Fcer1g/Fas/Sirpa/H2-M3/Pla2g2d/P2rx7/Vav1/Lcp1/Tcf7/Was/Arg1/Adk/Spta1/Il12rb1 |
| GO:0042113 | B cell activation | 1,22E-08 | Ighg2b/Ighv1-53/Ighv11-2/Ighv1-55/Ighg3/Ighg2c/Pou2af1/Lax1/Igkc/Ighd/Siglecg/Cr2/Ms4a1/Cd19/Cd79b/Mzb1/Cd22/Il7r/Ighm/Cd79a/Bank1/Btla/Blk/Iglc2/Sash3/Cd40/Exo1/Ikzf1/Igha/Btk/Card11/Samsn1/Rasgrp1/Nfam1/Cd74/Ptprc/Syk/Fcgr2b/Trbc2/Tbc1d10c/Myb/Il27ra/H2-Ab1/Il2rg/Ikzf3/Pou2f2/Cd180/Nckap1l/Cxcr5/Flt3/Cd28/Tlr9/Cd86/Dock10/Sh3kbp1/Cd300a/Ntrk1/Lat2/Parp3/Mfng/Tnfrsf13b/Lyl1/Tnfrsf21/Pik3cd/Prkcb/Itm2a/Tlr4/Fzd9/Tyrobp/Cd38/Gpr183/Tnfsf13b/Vav3/Icosl/Dclre1c/Inpp5d |
| GO:0042742 | defense response to bacterium | 1,22E-08 | Coch/Ighg2b/Ighv1-53/Ighv11-2/Ighv1-55/Ighg3/Ighg2c/Igkc/Jchain/Ighd/Cd4/Il7r/Ighm/Lyz1/Iglc2/Plac8/Cxcl13/Wfdc17/Igha/Scd1/Slpi/Spn/Gbp8/Syk/Trbc2/Lyz2/C5ar1/Il27ra/Cd209d/Myo1f/Fcgr1/Trf/Npy/Reg3b/Hp/Rarres2/Gbp2/Slc11a1/Vip/Tlr9/Irf8/Gsdmd/Havcr2/Cebpb/Colec12/Ncf1/Iigp1/Arg2/Serpine1/Syt11/Tlr5/Hck/B2m/Tlr4/Lypd8/H2-T23/Lcn2/Mpeg1/Prkd1/Card9 |
| GO:0045087 | innate immune response | 1,22E-08 | Coch/Ighg2b/Ighv1-53/Ighv11-2/Ighv1-55/Ighg3/Ighg2c/Cfd/Igkc/Jchain/Ighd/Siglecg/Cr2/Sla/Ear2/Treml2/Zbp1/Tlr1/Marco/Vsig4/Ccl8/Slamf7/Ighm/Ccl6/Mcoln2/Ifi209/Blk/Iglc2/Masp2/Ccr1/Cd40/Ccl9/Wfdc17/Igha/Btk/Cybb/Slpi/Padi4/Lck/H2-Q7/Rasgrp1/Casp4/Cd74/Ly9/Coro1a/Gbp8/H2-Eb1/Trim5/Cfp/Syk/Ifi204/Ifi205/Trbc2/Irf5/C3/H2-Aa/Tlr13/Anxa1/H2-Ab1/Irak3/Ly86/Il18/Myo1f/Stx11/Cd180/Fcgr1/Aim2/Pld4/Lacc1/Npy/Lep/Clec4a2/Apobec3/Lgals3/C1qc/Ccl11/Trim59/Axl/Rarres2/Rab7b/Smpdl3b/Cd84/Trim14/Gbp2/Oasl2/Slc11a1/Nlrc5/C1qb/Vim/Nmi/Vip/Tlr9/Fcna/Mfhas1/Wnt5a/Ccl24/Sp110/Tlr7/Cd55/Serping1/Irf8/Trim30d/Gsdmd/Havcr2/Cd1d1/Elf4/Ciita/Casp1/Iigp1/Mndal/Arg2/Pparg/Mst1r/Il1rl2/Ifi211/Alpk1/C1s1/C1ra/Oas2/Ifit1/Actg1/Pik3cd/Tlr5/Hck/Cfh/Mrc1/C2/B2m/Trim34a/Tlr4/Tlr8/Mmp2/Aqp4/H2-T23/Ifit2/Apcs/Trim12c/Lcn2/Mpeg1/Prkd1/Ccl5/Evl/Card9/Gsn/Spon2/Rab20/Csf1r/Tifa/Pstpip1/Ifitm1/Pla2g2f/Unc93b1/Trim35/Herc6/Trem2/Irf7/Plscr1/Trim30a/Stat1/Ifi203/Defb1/H2bc6/Trim12a/Rpl13a/Cadm1/Csf1/Parp14/Fcer1g/Tap1/Sirpa/H2-M3/Polr3h/Vav1/Hmgb2/Ptk2b/Was/Arg1/Gbp2b/Capg/Lrp8 |
| GO:0046651 | lymphocyte proliferation | 1,22E-08 | Ptpn22/Slfn1/Ighd/Siglecg/Cr2/Cd19/Cd4/Mzb1/Cd22/Vsig4/Il7r/Ighm/Cd79a/Rac2/Btla/Dock2/Blk/Sash3/Cd40/Btk/Card11/Rasgrp1/Lgals7/Spn/Cd74/Coro1a/Ptprc/Cd3e/Syk/Fcgr2b/H2-Aa/Anxa1/Rasal3/Itgam/Il27ra/H2-Ab1/Il2ra/Il18/Ikzf3/Vcam1/Ccr7/Cd180/Slc4a1/Lep/Nckap1l/Lgals3/Ripor2/Flt3/Slc11a1/Cd28/Tlr9/Cd86/Vtcn1/Vsir/Carmil2/Dock8/Lilrb4a/Satb1/Havcr2/Tnfsf9/Cebpb/Cd1d1/Cd300a/Elf4/Bmp4/Cd274/Ccdc88b/Ccr2/Arg2/Tnfrsf13b/Tnfrsf21/Cd44/Ephb6/Tlr4/H2-T23/Tyrobp/Cd38/Ccl5/Gpr183/Cd6/Erbb2/Tnfsf13b/Vav3/Icosl/Inpp5d/Pla2g2f/Tnfrsf1b/Cxcr4/Tgfbr2/Itgal/Itgb2/Irs2/Bst1/H2-M3/Pla2g2d/Gal/P2rx7 |
| GO:0050670 | regulation of lymphocyte proliferation | 1,22E-08 | Ptpn22/Slfn1/Ighd/Siglecg/Cd4/Mzb1/Cd22/Vsig4/Ighm/Rac2/Btla/Blk/Sash3/Cd40/Btk/Card11/Lgals7/Spn/Cd74/Coro1a/Ptprc/Cd3e/Syk/Fcgr2b/H2-Aa/Anxa1/Rasal3/Il27ra/H2-Ab1/Il2ra/Il18/Ikzf3/Vcam1/Ccr7/Slc4a1/Lep/Nckap1l/Lgals3/Ripor2/Cd28/Tlr9/Cd86/Vtcn1/Vsir/Carmil2/Havcr2/Tnfsf9/Cebpb/Cd1d1/Cd300a/Bmp4/Cd274/Ccdc88b/Ccr2/Arg2/Tnfrsf13b/Tnfrsf21/Cd44/Tlr4/H2-T23/Tyrobp/Cd38/Ccl5/Gpr183/Cd6/Erbb2/Tnfsf13b/Vav3/Icosl/Inpp5d/Pla2g2f/Tnfrsf1b/Tgfbr2/Itgal/Irs2/Bst1/H2-M3/Pla2g2d/Gal/Arg1/Adk/Spta1/Il12rb1 |
| GO:0050778 | positive regulation of immune response | 1,22E-08 | Stap1/Coch/Ighg2b/Ighv1-53/Ighv11-2/Ighv1-55/Ighg3/Ighg2c/Cfd/Scimp/Lax1/Igkc/Ptpn22/Fcer2a/Ighd/Cr2/Ms4a1/Cd19/Pck1/Cd79b/Zbp1/Klhl6/Cd4/Cd22/Vsig4/Cd5l/Ighm/Cd79a/Itk/Ifi209/Blk/Iglc2/Masp2/Sash3/Cd40/Il33/Igha/Fyb2/Btk/Card11/Myo1g/Lck/H2-Q7/Fcgr3/Rasgrp1/H2-Q6/Nfam1/Spn/Cd74/Ptprc/Cd3e/Cfp/Syk/Fcgr2b/Ifi204/Ifi205/Trbc2/C3/Anxa1/C5ar1/H2-DMa/C4b/Myb/Itgam/Il27ra/H2-Ab1/Il18/Ccr7/Fcgr1/Aim2/Nckap1l/Lgals3/C1qc/Pla2g4a/Slc11a1/Nlrc5/Cd28/C1qb/Tlr9/Fcna/Vtcn1/Wnt5a/Carmil2/Cd55/Serping1/Thy1/Havcr2/Nr4a3/Gprc5b/Cd1d1/Cd300a/Mill2/Cd274/Ccr2/Mndal/Lat2/Themis2/Ifi211/C1s1/C1ra/Rgcc/Lcp2/Nfkbid/Tnfrsf21/Cfh/Prkcb/C2/B2m/Hspa8/Cd44/Tlr4/Ffar3/Tlr8/Fyb/Cacnb3/Mmp2/H2-T23/Apcs/Cd38/Sh2b2/Card9/Azgp1/Spon2/Skap1/Lpxn/Tec/Tnfsf13b/Vav3/Icosl |
| GO:0050851 | antigen receptor-mediated signaling pathway | 1,22E-08 | Stap1/Ighg2b/Ighv1-53/Ighv11-2/Ighv1-55/Ighg3/Ighg2c/Lax1/Igkc/Ptpn22/Ighd/Ms4a1/Cd19/Cd79b/Klhl6/Cd22/Ighm/Cd79a/Itk/Blk/Iglc2/Igha/Fyb2/Card11/Lck/Nfam1/Spn/Ptprc/Cd3e/Syk/Fcgr2b/Trbc2/Ccr7/Nckap1l/Lgals3/Cd28/Vtcn1/Carmil2/Thy1/Cd300a/Lat2/Themis2/Lcp2/Nfkbid/Tnfrsf21/Prkcb/Fyb/Cacnb3/Cd38/Sh2b2/Skap1/Lpxn/Tec/Vav3/Icosl |
| GO:0050853 | B cell receptor signaling pathway | 1,22E-08 | Stap1/Ighg2b/Ighv1-53/Ighv11-2/Ighv1-55/Ighg3/Ighg2c/Igkc/Ighd/Ms4a1/Cd19/Cd79b/Klhl6/Cd22/Ighm/Cd79a/Itk/Blk/Iglc2/Igha/Lck/Nfam1/Ptprc/Syk/Fcgr2b/Trbc2 |
| GO:0050863 | regulation of T cell activation | 1,22E-08 | Lax1/Ptpn22/Slfn1/Pck1/Cd4/Vsig4/Rhoh/Il7r/Rac2/Btla/Sash3/Ikzf1/Card11/Lck/Rasgrp1/Lgals7/Spn/Cd74/Coro1a/Ptprc/Cd3e/Syk/H2-Aa/Anxa1/Runx3/Rasal3/H2-DMa/Myb/Il27ra/H2-Ab1/Dpp4/Il2rg/Il2ra/Il18/Vcam1/Ccr7/Slc4a1/Lep/Nckap1l/Lgals3/Ripor2/Cd28/Cd86/Vtcn1/Vsir/Carmil2/Dock8/Thy1/Havcr2/Tnfsf9/Cebpb/Cd1d1/Cd300a/Bmp4/Fgl1/Cd83/Cd274/Ccdc88b/Ccr2/Adora2a/Arg2/Il1rl2/Nfkbid/Efnb3/Tnfrsf21/Cd44/Ephb6/H2-T23/Runx1/Ccl5/Cd6/Gsn/Erbb2/Hfe/Rorc/Tnfsf13b/Icosl/Pla2g2f/Tnfrsf1b/Zbtb16/Tgfbr2/Zeb1/Cav1/Itgal/Sirpa/H2-M3/Pla2g2d/Tcf7/Arg1/Adk/Spta1/Il12rb1 |
| GO:0050864 | regulation of B cell activation | 1,22E-08 | Ighg2b/Ighv1-53/Ighv11-2/Ighv1-55/Ighg3/Ighg2c/Igkc/Ighd/Siglecg/Cd19/Mzb1/Cd22/Ighm/Bank1/Btla/Blk/Iglc2/Sash3/Cd40/Igha/Btk/Card11/Samsn1/Nfam1/Cd74/Ptprc/Syk/Fcgr2b/Trbc2/Tbc1d10c/Il27ra/Il2rg/Ikzf3/Nckap1l/Flt3/Cd28/Tlr9/Sh3kbp1/Cd300a/Parp3/Tnfrsf13b/Tnfrsf21/Tlr4/Tyrobp/Cd38/Gpr183/Tnfsf13b/Vav3/Inpp5d |
| GO:0050865 | regulation of cell activation | 1,22E-08 | Stap1/Ighg2b/Ighv1-53/Ighv11-2/Ighv1-55/Ighg3/Ighg2c/Lax1/Igkc/Ptpn22/Slfn1/Ighd/Siglecg/Cd19/Pck1/Cd4/Mzb1/Cd22/Vsig4/Rhoh/Il7r/Slamf7/Ighm/Bank1/Il1rl1/Rac2/Btla/Blk/Iglc2/Sash3/Cd40/Il33/Ikzf1/Igha/Btk/Card11/Tspan32/Lck/Samsn1/Rasgrp1/Nfam1/Lgals7/Spn/Cd74/Coro1a/Ptprc/Cd3e/Syk/Cd37/Fcgr2b/Trbc2/Tbc1d10c/H2-Aa/Anxa1/Runx3/Rasal3/H2-DMa/Myb/Itgam/Il27ra/H2-Ab1/Dpp4/Il2rg/Il2ra/Il18/Ikzf3/Vcam1/Ccr7/Slc4a1/Lep/Ptpre/Nckap1l/Lgals3/Pla2g4a/Il16/Ripor2/Flt3/Pdpn/Axl/Cd84/Cd28/Tlr9/Plek/Cd86/Vtcn1/Mfhas1/Vsir/Sh3kbp1/Wnt5a/Carmil2/Dock8/Thy1/Havcr2/Nr4a3/Tnfsf9/Cebpb/Cd1d1/Cd300a/Bmp4/Fgl1/Cd83/Cd274/Ccdc88b/Sphk1/Ccr2/Adora2a/Lrrk2/Arg2/Ccn2/Pparg/Ubash3b/C1qtnf1/Il1rl2/Nfkbid/Parp3/Tnfrsf13b/Efnb3/Tnfrsf21/Syt11/Cd44/Ephb6/Tlr4/H2-T23/Tyrobp/Runx1/Cd38/Ccl5/Gpr183/Cd6/Hmox1/Gsn/Tec/Erbb2/Hfe/Rorc/Tnfsf13b/Vav3/Icosl/Inpp5d/Pla2g2f/Tnfrsf1b/Adrb2/Zbtb16/Trem2/Tgfbr2/Plscr1/Zeb1/Thbs1/Cav1/Itgal/Itgb2/Irs2/Paxip1/Bst1/Fcer1g/Fas/Sirpa/H2-M3/Pla2g2d/Gal |
| GO:0050867 | positive regulation of cell activation | 1,22E-08 | Stap1/Ighg2b/Ighv1-53/Ighv11-2/Ighv1-55/Ighg3/Ighg2c/Igkc/Ptpn22/Ighd/Pck1/Cd4/Rhoh/Il7r/Ighm/Il1rl1/Iglc2/Sash3/Cd40/Il33/Ikzf1/Igha/Card11/Lck/Rasgrp1/Spn/Cd74/Coro1a/Ptprc/Cd3e/Syk/Trbc2/H2-Aa/Anxa1/Runx3/Rasal3/H2-DMa/Myb/Itgam/Il27ra/H2-Ab1/Dpp4/Il2rg/Il2ra/Il18/Vcam1/Ccr7/Slc4a1/Lep/Nckap1l/Pla2g4a/Il16/Pdpn/Axl/Cd28/Tlr9/Plek/Cd86/Vtcn1/Vsir/Sh3kbp1/Wnt5a/Carmil2/Dock8/Thy1/Havcr2/Nr4a3/Tnfsf9/Cd1d1/Cd83/Cd274/Ccdc88b/Ccr2/Lrrk2/Ccn2/Il1rl2/Nfkbid/Efnb3/Ephb6/Tlr4/H2-T23/Tyrobp/Runx1/Cd38/Ccl5/Gpr183/Cd6/Tnfsf13b/Vav3/Icosl/Inpp5d/Zbtb16/Trem2/Tgfbr2/Thbs1/Cav1/Itgal/Itgb2/Irs2/Paxip1/Bst1/Fcer1g/Sirpa/H2-M3 |
| GO:0051249 | regulation of lymphocyte activation | 1,22E-08 | Ighg2b/Ighv1-53/Ighv11-2/Ighv1-55/Ighg3/Ighg2c/Lax1/Igkc/Ptpn22/Slfn1/Ighd/Siglecg/Cd19/Pck1/Cd4/Mzb1/Cd22/Vsig4/Rhoh/Il7r/Slamf7/Ighm/Bank1/Rac2/Btla/Blk/Iglc2/Sash3/Cd40/Ikzf1/Igha/Btk/Card11/Lck/Samsn1/Rasgrp1/Nfam1/Lgals7/Spn/Cd74/Coro1a/Ptprc/Cd3e/Syk/Fcgr2b/Trbc2/Tbc1d10c/H2-Aa/Anxa1/Runx3/Rasal3/H2-DMa/Myb/Il27ra/H2-Ab1/Dpp4/Il2rg/Il2ra/Il18/Ikzf3/Vcam1/Ccr7/Slc4a1/Lep/Nckap1l/Lgals3/Ripor2/Flt3/Axl/Cd28/Tlr9/Cd86/Vtcn1/Vsir/Sh3kbp1/Carmil2/Dock8/Thy1/Havcr2/Tnfsf9/Cebpb/Cd1d1/Cd300a/Bmp4/Fgl1/Cd83/Cd274/Ccdc88b/Ccr2/Adora2a/Arg2/Il1rl2/Nfkbid/Parp3/Tnfrsf13b/Efnb3/Tnfrsf21/Cd44/Ephb6/Tlr4/H2-T23/Tyrobp/Runx1/Cd38/Ccl5/Gpr183/Cd6/Gsn/Erbb2/Hfe/Rorc/Tnfsf13b/Vav3/Icosl/Inpp5d/Pla2g2f/Tnfrsf1b/Zbtb16/Tgfbr2/Zeb1/Cav1/Itgal/Irs2/Paxip1/Bst1/Fas/Sirpa/H2-M3/Pla2g2d/Gal |
| GO:0051251 | positive regulation of lymphocyte activation | 1,22E-08 | Ighg2b/Ighv1-53/Ighv11-2/Ighv1-55/Ighg3/Ighg2c/Igkc/Ptpn22/Ighd/Pck1/Cd4/Rhoh/Il7r/Ighm/Iglc2/Sash3/Cd40/Ikzf1/Igha/Card11/Lck/Rasgrp1/Spn/Cd74/Coro1a/Ptprc/Cd3e/Syk/Trbc2/H2-Aa/Anxa1/Runx3/Rasal3/H2-DMa/Myb/Il27ra/H2-Ab1/Dpp4/Il2rg/Il2ra/Il18/Vcam1/Ccr7/Slc4a1/Lep/Nckap1l/Axl/Cd28/Tlr9/Cd86/Vtcn1/Vsir/Sh3kbp1/Carmil2/Dock8/Thy1/Havcr2/Tnfsf9/Cd1d1/Cd83/Cd274/Ccdc88b/Ccr2/Il1rl2/Nfkbid/Efnb3/Ephb6/Tlr4/H2-T23/Tyrobp/Runx1/Cd38/Ccl5/Gpr183/Cd6/Tnfsf13b/Vav3/Icosl/Inpp5d/Zbtb16/Tgfbr2/Cav1/Itgal/Irs2/Paxip1/Bst1 |
| GO:0070661 | leukocyte proliferation | 1,22E-08 | Ptpn22/Slfn1/Ighd/Siglecg/Cr2/Cd19/Cd4/Mzb1/Cd22/Vsig4/Il7r/Ighm/Cd79a/Rac2/Btla/Dock2/Blk/Sash3/Cd40/Il33/Btk/Card11/Rasgrp1/Lgals7/Spn/Cd74/Coro1a/Ptprc/Cd3e/Syk/Fcgr2b/H2-Aa/Anxa1/Rasal3/Itgam/Il27ra/H2-Ab1/Il2ra/Il18/Ikzf3/Vcam1/Ccr7/Cd180/Slc4a1/Lep/Nckap1l/Lgals3/Ripor2/Flt3/Slc11a1/Cd28/Tlr9/Cd86/Vtcn1/Vsir/Carmil2/Dock8/Lilrb4a/Satb1/Havcr2/Tnfsf9/Cebpb/Cd1d1/Cd300a/Elf4/Bmp4/Cd274/Ccdc88b/Ccr2/Arg2/Tnfrsf13b/Tnfrsf21/Cd44/Kitl/Ephb6/Tlr4/H2-T23/Tyrobp/Cd38/Ccl5/Gpr183/Cd6/Csf1r/Erbb2/Tnfsf13b/Vav3/Icosl/Inpp5d/Pla2g2f/Tnfrsf1b/Trem2/Cxcr4/Tgfbr2/Itgal/Itgb2/Irs2/Bst1/Csf1/H2-M3/Pla2g2d/Gal/P2rx7 |
| GO:0070663 | regulation of leukocyte proliferation | 1,22E-08 | Ptpn22/Slfn1/Ighd/Siglecg/Cd4/Mzb1/Cd22/Vsig4/Ighm/Rac2/Btla/Blk/Sash3/Cd40/Il33/Btk/Card11/Lgals7/Spn/Cd74/Coro1a/Ptprc/Cd3e/Syk/Fcgr2b/H2-Aa/Anxa1/Rasal3/Il27ra/H2-Ab1/Il2ra/Il18/Ikzf3/Vcam1/Ccr7/Slc4a1/Lep/Nckap1l/Lgals3/Ripor2/Cd28/Tlr9/Cd86/Vtcn1/Vsir/Carmil2/Havcr2/Tnfsf9/Cebpb/Cd1d1/Cd300a/Bmp4/Cd274/Ccdc88b/Ccr2/Arg2/Tnfrsf13b/Tnfrsf21/Cd44/Kitl/Tlr4/H2-T23/Tyrobp/Cd38/Ccl5/Gpr183/Cd6/Csf1r/Erbb2/Tnfsf13b/Vav3/Icosl/Inpp5d/Pla2g2f/Tnfrsf1b/Tgfbr2/Itgal/Irs2/Bst1/Csf1/H2-M3/Pla2g2d/Gal/Arg1/Adk/Spta1/Il12rb1 |
| GO:0099024 | plasma membrane invagination | 1,22E-08 | Stap1/Ighg2b/Ighv1-53/Ighv11-2/Ighv1-55/Ighg3/Ighg2c/Igkc/Ighd/Alox15/Marco/Rhoh/Ighm/Rac2/Iglc2/Igha/Msr1/Aurkb/Fcgr3/Fcgr2b/Trbc2/C3/Itgam/Fcgr1/Nckap1l/Arhgap25/Cd300a/Pparg/Bin2/Elmo1/Rab31/Gsn/Siglece/Trem2/Spire2/Spire1/Xkr4/Thbs1/Itgb2/Gulp1/Fcer1g/Sirpa |
| GO:1903037 | regulation of leukocyte cell-cell adhesion | 1,22E-08 | Lax1/Ptpn22/Slfn1/Pck1/Cd4/Vsig4/Rhoh/Il7r/Btla/Sash3/Ikzf1/Card11/Lck/Rasgrp1/Lgals7/Spn/Cd74/Coro1a/Ptprc/Cd3e/Syk/H2-Aa/Anxa1/Runx3/Rasal3/H2-DMa/Myb/Il27ra/H2-Ab1/Dpp4/Il2rg/Il2ra/Il18/Vcam1/Ccr7/Slc4a1/Lep/Nckap1l/Lgals3/Ripor2/Cd28/Cd86/Vtcn1/Vsir/Carmil2/Dock8/Thy1/Havcr2/Nr4a3/Tnfsf9/Cebpb/Cd1d1/Cd300a/Bmp4/Fgl1/Cd83/Cd274/Ccdc88b/Ccr2/Adora2a/Arg2/Il1rl2/Nfkbid/Efnb3/Tnfrsf21/Cd44/Ephb6/Icam1/H2-T23/Runx1/Ccl5/Cd6/Skap1/Erbb2/Hfe/Tnfsf13b/Icosl/Pla2g2f/Zbtb16/Tgfbr2/Itga4/Fut4/Cav1/Itgal/Itgb2/Sirpa/H2-M3/Pla2g2d |
| GO:0050900 | leukocyte migration | 2,57E-08 | Stap1/Sell/Ptpn22/Chst4/Rhoh/Ccl8/Ccl6/Mcoln2/Rac2/Ccr1/Il33/Ccl9/Cxcl13/Ptpro/Myo1g/Fcgr3/Spn/Cd74/Coro1a/Jaml/Syk/Pgf/Itgb7/Vegfd/Anxa1/C5ar1/Ccn3/Itgam/Il27ra/Dpp4/Vcam1/Ccr7/P2ry12/Selplg/Lep/Nckap1l/Lgals3/Cxcr5/Ccl11/Il16/Ripor2/Rarres2/Pla2g7/Nbl1/Thbs4/Wnt5a/Ccl24/Dock8/Thy1/Cd34/Cd300a/Dusp1/Ccr2/Itgb3/Cd200r1/Gpsm3/Ppbp/Trpm2/Serpine1/Dpep1/Kitl/Icam1/Pf4/Stk10/Ccl5/Gpr183/Prex1/B4galt1/Ecm1/Smpd3/Csf1r/Vav3/Trem2/Vegfc/Ccl27a/Itga4/Fut4/Mmp9/Thbs1/Itgal/Itgb2/Bst1/Csf3r/Rpl13a/Cadm1/Csf1/Fcer1g/Sirpa/C3ar1/Vav1 |
| GO:0002683 | negative regulation of immune system process | 2,71E-08 | Stap1/Lax1/Ptpn22/Slfn1/Siglecg/Hoxa5/Alox15/Cd22/Vsig4/Il7r/H2-Ob/Adipoq/Bank1/Il1rl1/Btla/Blk/Ccr1/Il33/Btk/Tspan32/Samsn1/Lgals7/Spn/Cd74/Ptprc/Cd37/Fcgr2b/Tbc1d10c/H2-Aa/Anxa1/Runx3/Lrrc17/Ccn3/Il27ra/H2-Ab1/Dpp4/Irak3/Il2ra/Il18/Lgals3/C1qc/Ripor2/Flt3/Axl/Smpdl3b/Cd84/Nlrc5/Nbl1/Nmi/Cd86/Vtcn1/Vsir/Serping1/Cd68/Thy1/Havcr2/Zfp36/Cebpb/Cd300a/Bmp4/Fgl1/Mill2/Cd274/Dusp1/Ccr2/Adora2a/Cd200r1/Arg2/Tspan6/Pparg/Ubash3b/Myc/Nfkbid/Parp3/Tnfrsf13b/Tnfrsf21/Syt11/Cd44/Kitl/Pf4/H2-T23/Tyrobp/Htra1/Runx1/Apcs/Trib1/Hmox1/Tsc22d3/Klf13/Lpxn/Erbb2/Hfe/Inpp5d/Pla2g2f/Zbtb16/Lmo2/Thbs1/Tgfb3/Parp14/Fcer1g/Fas/Vpreb3/Tap1/H2-M3/Pla2g2d/Gal/Gpx1/Tob2/Ptk2b/Muc4/Arg1 |
| GO:0031347 | regulation of defense response | 3,08E-08 | Stap1/Coch/Cma1/Ighg2b/Pbk/Ptpn22/Siglecg/Alox15/Treml2/Zbp1/Vsig4/Adipoq/Il1rl1/Ifi209/Ccr1/Il33/Btk/Tspan32/Fcgr3/Dnase1l3/Rasgrp1/Casp4/Spn/Cd74/Cd37/Fcgr2b/Ifi204/Ifi205/C3/Anxa1/Ccn3/Per1/Irak3/Il2ra/Myo1f/Ccr7/Fcgr1/Aim2/Pld4/Ptgis/Lacc1/Ctss/Lep/Cckbr/Apobec3/Akna/Pla2g4a/Il16/Smpdl3b/Nlrc5/Cd28/Nmi/Tlr9/Mfhas1/Wnt5a/Ccl24/Serping1/Wfdc1/Havcr2/Zfp36/Gprc5b/Cebpb/Mill2/Casp1/Alox5ap/Sphk1/Ccr2/Mndal/Adora2a/Pik3cg/Lrrk2/Usp18/Cd200r1/Arg2/Tspan6/Pparg/Gpsm3/Serpine1/Il1rl2/Ifi211/Aoah/Syt11/Abcd2/Cd44/Tlr4/Ffar3/Tlr8/Adcy7/Mmp2/H2-T23/Htra1/Ccl5/Cd6/Card9/Siglece/Pmp22/Tnfrsf1b/Adrb2/Nr1d2/Trem2/Esr1/Irf7/Ccr5/Plscr1/Isl1/Stat1/Ifi203/Casp12/Bst1/Cadm1/Parp14/Fcer1g/Tap1/Sirpa/H2-M3/Pla2g2d |
| GO:1903039 | positive regulation of leukocyte cell-cell adhesion | 4,69E-08 | Ptpn22/Pck1/Cd4/Rhoh/Il7r/Sash3/Ikzf1/Card11/Lck/Rasgrp1/Spn/Cd74/Coro1a/Ptprc/Cd3e/Syk/H2-Aa/Anxa1/Runx3/Rasal3/H2-DMa/Myb/Il27ra/H2-Ab1/Dpp4/Il2rg/Il2ra/Il18/Vcam1/Ccr7/Slc4a1/Lep/Nckap1l/Cd28/Cd86/Vtcn1/Vsir/Carmil2/Dock8/Thy1/Havcr2/Nr4a3/Tnfsf9/Cd1d1/Cd83/Cd274/Ccdc88b/Ccr2/Il1rl2/Nfkbid/Efnb3/Cd44/Ephb6/Icam1/H2-T23/Runx1/Ccl5/Cd6/Skap1/Tnfsf13b/Icosl/Zbtb16/Tgfbr2/Itga4/Fut4/Cav1/Itgal/Itgb2 |
| GO:0002699 | positive regulation of immune effector process | 4,83E-08 | Stap1/Ighg2b/Scimp/Ptpn22/Fcer2a/Pck1/Mzb1/Rac2/Blk/Sash3/Cd40/Il33/Btk/H2-Q7/Fcgr3/Rasgrp1/H2-Q6/Cd74/Ptprc/Syk/Cd37/C3/Anxa1/Myb/Itgam/Il18/Ccr7/Fcgr1/Cd84/Cd28/Tlr9/Wnt5a/Nr4a3/Gprc5b/Cd1d1/Cd300a/Ccr2/Nfkbid/B2m/Hspa8/Tlr4/Ffar3/H2-T23/Tyrobp/Hmox1/Azgp1/Spon2/H2-T10/Itgb2/Paxip1/Cadm1/Fcer1g/H2-M3/P2rx7/Vav1 |
| GO:0002455 | humoral immune response mediated by circulating immunoglobulin | 5,06E-08 | Ighg2b/Ighv1-53/Ighv11-2/Ighv1-55/Ighg3/Ighg2c/Igkc/Fcer2a/Ighd/Cr2/Ighm/Iglc2/Masp2/Exo1/Igha/Ptprc/Fcgr2b/Trbc2/C3/H2-Ab1/C1qc/C1qb/Cd55/Serping1/C1s1/C1ra/C2 |
| GO:0019221 | cytokine-mediated signaling pathway | 6,17E-08 | Stap1/Zbp1/Cd4/Ccl8/Adipoq/Ccl6/Il2rb/Ccr1/Il33/Ccl9/Cxcl13/Casp4/Cd74/Ptprc/Syk/Irf5/Spi1/Il27ra/Il2rg/Irak3/Il18/Pirb/Egr1/Aim2/Lep/Rps6ka5/Ccl11/Il16/Flt3/Csf2rb/Axl/Nlrc5/Cntfr/Nmi/Wnt5a/Ccl24/Csf2ra/Traip/Casp1/Sphk1/Ccr2/Iigp1/Ackr3/Pparg/Ppbp/Duox1/Traf1/Oas2/Grem2/Thpo/Il3ra/Pf4/Ccl5/St18/Sh2b2/Card14/Ecm1/Csf1r/Ifitm1/Klf6/Trem2/Cxcr4/Irf7/Stat1/Slc27a1/Cav1/Lepr/Csf3r/Csf1/Parp14/Fcer1g/Il10ra/Ptk2b/Arg1/Il12rb1/Acsl1/Numbl/Robo1/Foxo3/Irgm2/Bbs4/Il12rb2/Il18r1/Jak3/Stat3/Nfkbia/Il15ra/Mt3/Myd88/Foxc1 |
| GO:0002819 | regulation of adaptive immune response | 7,04E-08 | Ighg2b/Fcer2a/Alox15/Cd4/Il7r/Il1rl1/Sash3/Cd40/Il33/Btk/H2-Q7/Fcgr3/Samsn1/H2-Q6/Spn/Cd74/Ptprc/Fcgr2b/C3/Anxa1/H2-DMa/Cd48/Il27ra/H2-Ab1/Dpp4/Il18/Ccr7/Fcgr1/Pla2g4a/Slc11a1/Cd28/Vsir/Cd55/Havcr2/Cd1d1/Cd274/Ccr2/Nfkbid/Parp3/B2m/Hspa8/Cd44/Adcy7/H2-T23/Azgp1/Skap1/Hfe/Tnfsf13b/H2-T10/Tnfrsf1b/Irf7/Paxip1/Fcer1g/H2-M3/P2rx7/Muc4/Was/Arg1/H2-T24/Il12rb1 |
| GO:0042098 | T cell proliferation | 9,46E-08 | Ptpn22/Slfn1/Cd4/Vsig4/Rac2/Btla/Dock2/Sash3/Card11/Rasgrp1/Lgals7/Spn/Coro1a/Ptprc/Cd3e/Syk/H2-Aa/Anxa1/Rasal3/Itgam/Il27ra/H2-Ab1/Il2ra/Il18/Vcam1/Ccr7/Slc4a1/Lep/Nckap1l/Lgals3/Ripor2/Slc11a1/Cd28/Cd86/Vtcn1/Vsir/Carmil2/Dock8/Lilrb4a/Satb1/Havcr2/Tnfsf9/Cebpb/Cd1d1/Elf4/Bmp4/Cd274/Ccdc88b/Ccr2/Arg2/Tnfrsf21/Cd44/Ephb6/H2-T23/Ccl5/Cd6/Erbb2/Tnfsf13b/Icosl/Pla2g2f/Tnfrsf1b/Cxcr4/Tgfbr2/Itgal/Itgb2/H2-M3/Pla2g2d/P2rx7/Arg1/Adk/Spta1/Il12rb1 |
| GO:0002274 | myeloid leukocyte activation | 9,63E-08 | Stap1/Tlr1/Rhoh/Batf3/Il1rl1/Rac2/Dock2/Blk/Il33/Btk/Tspan32/Fcgr3/Dnase1l3/Rasgrp1/Syk/Cd37/Camk4/Spi1/Irf4/C5ar1/Cd48/Itgam/Il18/Myo1f/Stx11/Pirb/Ptpre/Pla2g4a/Il16/Cd84/Slc11a1/Tlr9/Mfhas1/Wnt5a/Tlr7/Havcr2/Nr4a3/Tnfsf9/Cd1d1/Cd300a/Casp1/Sphk1/Ccr2/Lrrk2/Lat2/Pparg/Lcp2/Syt11/Tlr4/Tlr8/Tyrobp/Ccl5/Hmox1/Trem2/Anxa3/Tgfbr2/Plscr1/Batf/Thbs1/Itgb2/Jun |
| GO:0050870 | positive regulation of T cell activation | 1,12E-07 | Ptpn22/Pck1/Cd4/Rhoh/Il7r/Sash3/Ikzf1/Card11/Lck/Rasgrp1/Spn/Cd74/Coro1a/Ptprc/Cd3e/Syk/H2-Aa/Anxa1/Runx3/Rasal3/H2-DMa/Myb/Il27ra/H2-Ab1/Dpp4/Il2rg/Il2ra/Il18/Vcam1/Ccr7/Slc4a1/Lep/Nckap1l/Cd28/Cd86/Vtcn1/Vsir/Carmil2/Dock8/Thy1/Havcr2/Tnfsf9/Cd1d1/Cd83/Cd274/Ccdc88b/Ccr2/Il1rl2/Nfkbid/Efnb3/Ephb6/H2-T23/Runx1/Ccl5/Cd6/Tnfsf13b/Icosl/Zbtb16/Tgfbr2/Cav1/Itgal |
| GO:0050871 | positive regulation of B cell activation | 1,94E-07 | Ighg2b/Ighv1-53/Ighv11-2/Ighv1-55/Ighg3/Ighg2c/Igkc/Ighd/Ighm/Iglc2/Sash3/Cd40/Igha/Card11/Cd74/Ptprc/Syk/Trbc2/Il2rg/Nckap1l/Cd28/Tlr9/Sh3kbp1 |
| GO:0006956 | complement activation | 2,23E-07 | Ighg2b/Ighv1-53/Ighv11-2/Ighv1-55/Ighg3/Ighg2c/Cfd/Igkc/Ighd/Cr2/Vsig4/Cd5l/Ighm/Iglc2/Masp2/Igha/Cfp/Trbc2/C3/C4b/C1qc/C1qb/Fcna/Cd55/Serping1/C1s1/C1ra/Rgcc/Cfh/C2 |
| GO:0006910 | phagocytosis, recognition | 2,29E-07 | Ighg2b/Ighv1-53/Ighv11-2/Ighv1-55/Ighg3/Ighg2c/Igkc/Ighd/Ighm/Iglc2/Igha/Cd209b/Fcgr3/Trbc2/Fcgr1 |
| GO:0022409 | positive regulation of cell-cell adhesion | 2,47E-07 | Ptpn22/Alox15/Pck1/Cd4/Rhoh/Il7r/Sash3/Cxcl13/Ikzf1/Card11/Lck/Rasgrp1/Spn/Cd74/Coro1a/Ptprc/Cd3e/Syk/H2-Aa/Anxa1/Runx3/Rasal3/H2-DMa/Myb/Il27ra/H2-Ab1/Dpp4/Il2rg/Il2ra/Il18/Vcam1/Ccr7/Slc4a1/Lep/Nckap1l/Pdpn/Cd28/Cd86/Vtcn1/Vsir/Wnt5a/Carmil2/Dock8/Thy1/Havcr2/Nr4a3/Tnfsf9/Cd1d1/Cd83/Cd274/Ccdc88b/Ccr2/Il1rl2/Nfkbid/Efnb3/Cd44/Ephb6/Icam1/H2-T23/Runx1/Ccl5/Cd6/Skap1/Tnfsf13b/Icosl/Zbtb16/Ccr5/Tgfbr2/Itga4/Fut4/Cav1/Itgal/Itgb2 |
| GO:0006935 | chemotaxis | 4,12E-07 | Stap1/Saa3/Sell/Egr2/Ear2/Rhoh/Ccl8/Ccl6/Sema3d/Rac2/Dock2/Ccr1/Ccl9/Cxcl13/Prtg/Lrp2/Ptpro/Fcgr3/Reln/Cd74/Ackr4/Coro1a/Plxnc1/Jaml/Syk/Pgf/Enpp2/Fgf1/Vegfd/Anxa1/C5ar1/Runx3/Ccn3/Cxcr6/Itgam/Lsp1/Dpp4/Vcam1/Fn1/Sema3e/Ccr7/Efna5/P2ry12/Epha5/Ackr2/Nckap1l/Lgals3/Cxcr5/Ccl11/Il16/Ripor2/Vstm2l/Rarres2/Pla2g7/Sema3a/Nbl1/Ephb3/Thbs4/Wnt5a/Ccl24/Wnt3/Nr4a3/Cdk5r1/Gli2/Dusp1/B3gnt2/Sema6c/Ccn1/Ccr2/Ntrk1/Pik3cg/Itgb3/Ackr3/Gpsm3/Ppbp/Trpm2/Serpine1/Efnb3/Pik3cd/Bin2/Dpep1/Xcr1/Ephb6/Pf4/Epha8/Dpysl2/Prkd1/Ccl5/Kif5a/Tubb2b/Gpr183/Evl/Prex1/Rtn4r/Dpysl5/Nrp2/Plxna4/Csf1r/Erbb2/Chn1/Vav3/Ncam1/Gas1/Flrt3/Nexn/Trem2/Cxcr4/Vegfc/Ccr5/Vangl2/Ccl27a/Isl1/Rpl24/Cmtm3/Pax6/Thbs1/Apbb1/Itgb2/Cysltr1/Bst1/Csf3r/Rpl13a/Csf1/Fcer1g/Emb/Nptn/C3ar1/Vav1/Myh10/Sema3c/Lrp1/Hmgb2/Ptk2b |
| GO:0002822 | regulation of adaptive immune response based on somatic recombination of immune receptors built from immunoglobulin superfamily domains | 5,46E-07 | Ighg2b/Fcer2a/Cd4/Il7r/Il1rl1/Sash3/Cd40/Il33/Btk/H2-Q7/Fcgr3/H2-Q6/Spn/Ptprc/Fcgr2b/C3/Anxa1/H2-DMa/Il27ra/H2-Ab1/Dpp4/Il18/Ccr7/Fcgr1/Pla2g4a/Slc11a1/Cd28/Vsir/Cd55/Havcr2/Cd1d1/Cd274/Ccr2/Nfkbid/Parp3/B2m/Hspa8/H2-T23/Azgp1/Hfe/Tnfsf13b/H2-T10/Tnfrsf1b/Paxip1/Fcer1g/H2-M3/P2rx7/Muc4/Was/Arg1/H2-T24/Il12rb1 |
| GO:0042100 | B cell proliferation | 6,99E-07 | Ighd/Siglecg/Cr2/Cd19/Mzb1/Cd22/Il7r/Ighm/Cd79a/Btla/Blk/Sash3/Cd40/Btk/Card11/Rasgrp1/Cd74/Ptprc/Fcgr2b/Ikzf3/Cd180/Nckap1l/Tlr9/Cd300a/Tnfrsf13b/Tnfrsf21/Tlr4/Tyrobp/Cd38/Gpr183/Tnfsf13b/Vav3/Inpp5d |
| GO:0042330 | taxis | 7,10E-07 | Stap1/Saa3/Sell/Egr2/Ear2/Rhoh/Ccl8/Ccl6/Sema3d/Rac2/Dock2/Ccr1/Ccl9/Cxcl13/Prtg/Lrp2/Ptpro/Fcgr3/Reln/Cd74/Ackr4/Coro1a/Plxnc1/Jaml/Syk/Pgf/Enpp2/Fgf1/Vegfd/Anxa1/C5ar1/Runx3/Ccn3/Cxcr6/Itgam/Lsp1/Dpp4/Vcam1/Fn1/Sema3e/Ccr7/Efna5/P2ry12/Epha5/Ackr2/Nckap1l/Lgals3/Cxcr5/Ccl11/Il16/Ripor2/Vstm2l/Rarres2/Pla2g7/Sema3a/Nbl1/Ephb3/Thbs4/Wnt5a/Ccl24/Wnt3/Nr4a3/Cdk5r1/Gli2/Dusp1/B3gnt2/Sema6c/Ccn1/Ccr2/Ntrk1/Pik3cg/Itgb3/Ackr3/Gpsm3/Ppbp/Trpm2/Serpine1/Efnb3/Pik3cd/Bin2/Dpep1/Xcr1/Ephb6/Pf4/Epha8/Dpysl2/Prkd1/Ccl5/Kif5a/Tubb2b/Gpr183/Evl/Prex1/Rtn4r/Dpysl5/Nrp2/Plxna4/Csf1r/Erbb2/Chn1/Vav3/Ncam1/Gas1/Flrt3/Nexn/Trem2/Cxcr4/Vegfc/Ccr5/Vangl2/Ccl27a/Isl1/Rpl24/Cmtm3/Pax6/Thbs1/Apbb1/Itgb2/Cysltr1/Bst1/Csf3r/Rpl13a/Csf1/Fcer1g/Emb/Nptn/C3ar1/Vav1/Myh10/Sema3c/Lrp1/Hmgb2/Ptk2b |
| GO:0002703 | regulation of leukocyte mediated immunity | 7,92E-07 | Stap1/Ighg2b/Scimp/Fcer2a/Il7r/Rac2/Blk/Sash3/Cd40/Btk/H2-Q7/Fcgr3/Dnase1l3/Rasgrp1/H2-Q6/Spn/Ptprc/Syk/Fcgr2b/C3/Itgam/Il27ra/Dpp4/Il18/Fcgr1/Lep/Cd84/Cd28/Tlr9/Vsir/Cd55/Havcr2/Cd1d1/Cd300a/Ccr2/Parp3/B2m/Hspa8/Tlr4/H2-T23/Tyrobp/Hmox1/Azgp1/Hfe/H2-T10/Tnfrsf1b/Itgb2/Paxip1/Cadm1/Fcer1g/Tap1/H2-M3/P2rx7/Vav1/Muc4/Was/Arg1/H2-T24/Hmces/Tnfsf13/Il18r1/Lag3/Jak3/Arrb2/Clec2d/Gfer/Stx7/Foxf1/Unc13d/Lyn |
| GO:0046631 | alpha-beta T cell activation | 1,01E-06 | Ptpn22/Gadd45g/Itk/Btla/Dock2/Sash3/Ikzf1/Lgals7/Spn/Ly9/Ptprc/Cd3e/Syk/Irf4/Anxa1/Wdfy4/Runx3/Rasal3/Myb/H2-Ab1/Il2rg/Il18/Ccr7/Clec4a2/Nckap1l/Cd28/Vsir/Lilrb4a/Satb1/Cebpb/Cd1d1/Cd300a/Elf4/Cd83/Cd274/Ccr2/Adora2a/Arg2/Nfkbid/Cd44/H2-T23/Runx1/Gpr183/Hfe/Rorc/Zbtb16/Tgfbr2/Batf |
| GO:0042129 | regulation of T cell proliferation | 1,04E-06 | Ptpn22/Slfn1/Cd4/Vsig4/Rac2/Btla/Sash3/Card11/Lgals7/Spn/Coro1a/Ptprc/Cd3e/Syk/H2-Aa/Anxa1/Rasal3/Il27ra/H2-Ab1/Il2ra/Il18/Vcam1/Ccr7/Slc4a1/Lep/Nckap1l/Lgals3/Ripor2/Cd28/Cd86/Vtcn1/Vsir/Carmil2/Havcr2/Tnfsf9/Cebpb/Cd1d1/Bmp4/Cd274/Ccdc88b/Ccr2/Arg2/Tnfrsf21/Cd44/H2-T23/Ccl5/Cd6/Erbb2/Tnfsf13b/Icosl/Pla2g2f/Tnfrsf1b/Tgfbr2/Itgal/H2-M3/Pla2g2d/Arg1/Adk/Spta1/Il12rb1 |
| GO:0006897 | endocytosis | 1,45E-06 | Stap1/Ighg2b/Ighv1-53/Ighv11-2/Ighv1-55/Ighg3/Ighg2c/Igkc/Ighd/Alox15/Cd22/Marco/Rhoh/Ighm/Rac2/Dock2/Iglc2/Lrp2/Igha/Cd209b/Msr1/Fcgr3/Fcho1/Syk/Rspo1/Fcgr2b/Trbc2/C3/Pacsin1/Itgam/Cd209d/Ccr7/Fcgr1/Trf/Nckap1l/Lgals3/Syt2/Scara5/Axl/Mx2/Arhgap25/Sh3kbp1/Wnt5a/Dnm1/Cd300a/Colec12/Akap5/Sphk1/Siglec1/Clip3/Pik3cg/Bmp2k/Lrrk2/Itgb3/Ackr3/Pparg/Sh3gl2/Serpine1/Snx10/Actg1/Syt11/Epn3/Mrc1/Bin2/Clec9a/B2m/Magi2/Snph/Sfrp4/Dpysl2/Elmo1/Fchsd2/Mx1/Pla2r1/Rab31/Gsn/Siglece/Gpc3/Snap25/Anxa2/Hfe/Pstpip1/Dab2/Ppp3cc/Slc2a4/Adrb2/Trem2/Tgfbr2/Itga4/Xkr4/Thbs1/Cav1/Itgb2/Arf6/Ehd3/Gulp1/Arc/Fcer1g/Fnbp1/Il10ra/Sirpa/Rin2/Ston1/Adm |
| GO:0050727 | regulation of inflammatory response | 2,15E-06 | Stap1/Cma1/Ighg2b/Pbk/Siglecg/Alox15/Zbp1/Adipoq/Il1rl1/Il33/Btk/Fcgr3/Dnase1l3/Casp4/Spn/Fcgr2b/C3/Anxa1/Ccn3/Per1/Il2ra/Ccr7/Fcgr1/Pld4/Ptgis/Lacc1/Ctss/Lep/Akna/Pla2g4a/Il16/Smpdl3b/Cd28/Tlr9/Mfhas1/Wnt5a/Ccl24/Wfdc1/Zfp36/Gprc5b/Cebpb/Casp1/Alox5ap/Sphk1/Ccr2/Adora2a/Pik3cg/Lrrk2/Usp18/Cd200r1/Pparg/Gpsm3/Serpine1/Il1rl2/Aoah/Syt11/Abcd2/Cd44/Tlr4/Ffar3/Adcy7/H2-T23/Ccl5/Cd6/Siglece/Pmp22/Tnfrsf1b/Adrb2/Nr1d2/Trem2/Esr1/Ccr5/Isl1/Casp12/Bst1/Fcer1g/Sirpa/Pla2g2d |
| GO:0006958 | complement activation, classical pathway | 2,19E-06 | Ighg2b/Ighv1-53/Ighv11-2/Ighv1-55/Ighg3/Ighg2c/Igkc/Ighd/Cr2/Ighm/Iglc2/Masp2/Igha/Trbc2/C3/C1qc/C1qb/Cd55/Serping1/C1s1/C1ra/C2/Apcs |
| GO:1902107 | positive regulation of leukocyte differentiation | 2,48E-06 | Pck1/Cd4/Rhoh/Il7r/Sash3/Ccr1/Fos/Ikzf1/Lck/Rasgrp1/Cd74/Ptprc/Syk/H2-Aa/Anxa1/Runx3/H2-DMa/Myb/Itgam/Il2rg/Il2ra/Il18/Ccr7/Nckap1l/Car2/Axl/Vsir/Hcls1/Carmil2/Tnfsf9/Cd1d1/Cd83/Ccr2/Itgb3/Il1rl2/Lif/Nfkbid/Kitl/Pf4/Tyrobp/Runx1/Ccl5/Trib1/Csf1r/Inpp5d/Zbtb16/Trem2/Tgfbr2/Jun/Dlk1/Csf1/H2-M3 |
| GO:0048002 | antigen processing and presentation of peptide antigen | 3,10E-06 | H2-Ob/H2-DMb2/H2-DMb1/H2-Q7/Fcgr3/H2-Q6/Cd74/H2-Eb1/Marchf1/Fcgr2b/H2-Aa/H2-DMa/H2-Ab1/Ctse/Fcgr1/Ctss/Clec4a2/Slc11a1/Ifi30/B2m/H2-T23/Azgp1/Hfe/H2-T10/Unc93b1/Trem2/Fcer1g/Tap1/H2-M3 |
| GO:1903708 | positive regulation of hemopoiesis | 3,10E-06 | Hoxa5/Pck1/Cd4/Rhoh/Il7r/Sash3/Ccr1/Fos/Ikzf1/Lck/Rasgrp1/Cd74/Ptprc/Syk/H2-Aa/Anxa1/Runx3/H2-DMa/Myb/Itgam/Il2rg/Il2ra/Il18/Ccr7/Lep/Nckap1l/Car2/Axl/Rab7b/Vsir/Hcls1/Carmil2/Tnfsf9/Cd1d1/Cd83/Ccr2/Itgb3/Il1rl2/Lif/Nfkbid/Thpo/Kitl/Pf4/Tyrobp/Runx1/Ccl5/Trib1/Csf1r/Inpp5d/Zbtb16/Trem2/Tgfbr2/Stat1/Jun/Dlk1/Csf1/Mturn/H2-M3 |
| GO:0002695 | negative regulation of leukocyte activation | 3,10E-06 | Lax1/Ptpn22/Slfn1/Vsig4/Bank1/Btla/Blk/Btk/Tspan32/Samsn1/Lgals7/Spn/Cd74/Cd37/Fcgr2b/Tbc1d10c/H2-Aa/Anxa1/Runx3/H2-Ab1/Il2ra/Lgals3/Ripor2/Flt3/Axl/Cd84/Cd86/Vtcn1/Vsir/Havcr2/Cebpb/Cd300a/Bmp4/Fgl1/Cd274/Ccr2/Adora2a/Arg2/Pparg/Nfkbid/Parp3/Tnfrsf13b/Tnfrsf21/Syt11/Cd44/Tyrobp/Runx1/Hmox1/Erbb2/Hfe/Inpp5d/Pla2g2f |
| GO:0097529 | myeloid leukocyte migration | 3,11E-06 | Stap1/Sell/Rhoh/Ccl8/Ccl6/Mcoln2/Rac2/Ccr1/Ccl9/Cxcl13/Ptpro/Fcgr3/Cd74/Jaml/Syk/Pgf/Vegfd/Anxa1/C5ar1/Ccn3/Itgam/Dpp4/Ccr7/P2ry12/Nckap1l/Lgals3/Ccl11/Ripor2/Rarres2/Pla2g7/Nbl1/Thbs4/Ccl24/Cd300a/Dusp1/Ccr2/Cd200r1/Ppbp/Serpine1/Dpep1/Pf4/Ccl5/Prex1/Csf1r/Vav3/Trem2/Vegfc/Thbs1/Itgb2/Bst1/Csf3r/Rpl13a/Csf1/Fcer1g/Sirpa/C3ar1/Vav1 |
| GO:0051250 | negative regulation of lymphocyte activation | 3,77E-06 | Lax1/Ptpn22/Slfn1/Vsig4/Bank1/Btla/Blk/Btk/Samsn1/Lgals7/Spn/Cd74/Fcgr2b/Tbc1d10c/H2-Aa/Anxa1/Runx3/H2-Ab1/Il2ra/Lgals3/Ripor2/Flt3/Axl/Cd86/Vtcn1/Vsir/Havcr2/Cebpb/Cd300a/Bmp4/Fgl1/Cd274/Adora2a/Arg2/Nfkbid/Parp3/Tnfrsf13b/Tnfrsf21/Cd44/Tyrobp/Runx1/Erbb2/Hfe/Inpp5d/Pla2g2f |
| GO:0032609 | interferon-gamma production | 4,45E-06 | Ptpn22/Cd2/Gadd45g/Il1rl1/Itk/Sash3/Il33/H2-Q7/Rasgrp1/Lgals7/Spn/Cd3e/Runx3/Il27ra/Il18/Ccr7/Flt3/Axl/Slc11a1/Tlr9/Vtcn1/Vsir/Wnt5a/Carmil2/Tlr7/Irf8/Havcr2/Tnfsf9/Cd1d1/Cd274/Ccr2/Tlr4/Tlr8/Runx1 |
| GO:0060326 | cell chemotaxis | 5,56E-06 | Stap1/Saa3/Sell/Ccl8/Ccl6/Rac2/Ccr1/Ccl9/Cxcl13/Ptpro/Fcgr3/Cd74/Ackr4/Coro1a/Jaml/Syk/Pgf/Enpp2/Fgf1/Vegfd/Anxa1/C5ar1/Ccn3/Cxcr6/Itgam/Dpp4/Vcam1/Ccr7/Ackr2/Nckap1l/Lgals3/Cxcr5/Ccl11/Il16/Ripor2/Rarres2/Pla2g7/Nbl1/Thbs4/Wnt5a/Ccl24/Dusp1/Ccr2/Ackr3/Gpsm3/Ppbp/Trpm2/Serpine1/Bin2/Dpep1/Xcr1/Pf4/Prkd1/Ccl5/Gpr183/Prex1/Csf1r/Vav3/Cxcr4/Vegfc/Ccr5/Ccl27a/Thbs1/Itgb2/Bst1/Csf3r/Rpl13a/Csf1/Fcer1g/C3ar1/Vav1/Hmgb2/Ptk2b |
| GO:0032103 | positive regulation of response to external stimulus | 7,63E-06 | Stap1/Coch/Ighg2b/Sell/Scimp/Zbp1/Il1rl1/Ifi209/Rac2/Ccr1/Il33/Cxcl13/Btk/Fcgr3/Rasgrp1/Cd74/Pgf/Ifi204/Ifi205/Vegfd/C3/C5ar1/Ly86/Fn1/Ccr7/P2ry12/Cd180/Fcgr1/Aim2/Ctss/Nckap1l/Pla2g4a/Il16/Ripor2/Rarres2/Nlrc5/Pla2g7/Cd28/Tlr9/Thbs4/Wnt5a/Ccl24/Havcr2/Gprc5b/Cebpb/Alox5ap/Ccr2/Mndal/Pik3cg/Lrrk2/Gpsm3/Serpine1/Ifi211/Bmp6/Tlr4/Ffar3/Tlr8/Mmp2/H2-T23/Prkd1/Ccl5/Tubb2b/Cd6/Card9/Csf1r/Trem2/Cxcr4/Vegfc/Irf7/Ccr5/Ccl27a/Plscr1/Ifi203/Thbs1/Cadm1/Csf1/Fcer1g/H2-M3/C3ar1/Vav1/Igf1r/Lrp1/Hmgb2/Ptk2b/Arg1 |
| GO:0030217 | T cell differentiation | 8,05E-06 | Ptpn22/Pck1/Cd4/Gadd45g/Rhoh/Il7r/Itk/Dock2/Sash3/Ikzf1/Card11/Lck/Rasgrp1/Spn/Cd74/Ly9/Ptprc/Cd3e/Syk/H2-Aa/Irf4/Anxa1/Runx3/H2-DMa/Myb/Il2rg/Il2ra/Il18/Ccr7/Egr1/Lep/Nckap1l/Flt3/Cd28/Vsir/Carmil2/Runx2/Satb1/Tnfsf9/Cd1d1/Bmp4/Cd83/Ccr2/Il1rl2/Nfkbid/B2m/Cd44/Runx1/Mpzl2/Gpr183/Prex1/Erbb2/Rorc/Zbtb16/Tgfbr2/Zeb1/Batf/Lepr/Fcer1g/Fas/H2-M3/Pla2g2d/Vav1/Tcf7 |
| GO:0002263 | cell activation involved in immune response | 8,64E-06 | Pou2af1/Cd19/Pck1/Gadd45g/Rac2/Dock2/Blk/Cd40/Il33/Exo1/Btk/Dnase1l3/Rasgrp1/Spn/Cd74/Ly9/Coro1a/Ptprc/Syk/Irf4/Anxa1/Myb/Itgam/Il27ra/Il18/Myo1f/Stx11/Ccr7/Cd180/Lgals3/Apbb1ip/Cd84/Slc11a1/Cd28/Dock10/Havcr2/Nr4a3/Cd300a/Ccr2/Lat2/Nfkbid/Parp3/Mfng/Itm2a/Tlr4/Icam1/Tyrobp/Gpr183/Hmox1/Rorc/Icosl/Trem2/Anxa3/Batf/Itgal/Itgb2/Paxip1/Fcer1g/H2-M3/Lcp1/Lrp1/Ptk2b |
| GO:0002366 | leukocyte activation involved in immune response | 8,80E-06 | Pou2af1/Cd19/Pck1/Gadd45g/Rac2/Dock2/Blk/Cd40/Il33/Exo1/Btk/Dnase1l3/Rasgrp1/Spn/Cd74/Ly9/Coro1a/Ptprc/Syk/Irf4/Anxa1/Myb/Itgam/Il27ra/Il18/Myo1f/Stx11/Ccr7/Cd180/Lgals3/Apbb1ip/Cd84/Slc11a1/Cd28/Dock10/Havcr2/Nr4a3/Cd300a/Ccr2/Lat2/Nfkbid/Parp3/Mfng/Itm2a/Tlr4/Icam1/Tyrobp/Gpr183/Hmox1/Rorc/Icosl/Trem2/Anxa3/Batf/Itgal/Itgb2/Paxip1/Fcer1g/H2-M3/Lcp1 |
| GO:1902105 | regulation of leukocyte differentiation | 8,80E-06 | Pck1/Cd4/Rhoh/Il7r/Adipoq/Sash3/Ccr1/Fos/Ikzf1/Card11/Lck/Rasgrp1/Nfam1/Cd74/Ptprc/Syk/H2-Aa/Anxa1/Runx3/Lrrc17/H2-DMa/Myb/Itgam/Il2rg/Il2ra/Il18/Ikzf3/Ccr7/Nckap1l/C1qc/Flt3/Car2/Axl/Cd28/Tlr9/Vsir/Hcls1/Carmil2/Tnfsf9/Cebpb/Cd1d1/Bmp4/Cd83/Ccr2/Itgb3/Rassf2/Ubash3b/Myc/Il1rl2/Lif/Nfkbid/Cd44/Kitl/Pf4/Tyrobp/Runx1/Apcs/Ccl5/Trib1/Csf1r/Erbb2/Rorc/Inpp5d/Zbtb16/Trem2/Tgfbr2/Zeb1/Jun/Dlk1/Csf1/Fas/H2-M3 |
| GO:0030595 | leukocyte chemotaxis | 9,08E-06 | Stap1/Sell/Ccl8/Ccl6/Rac2/Ccr1/Ccl9/Cxcl13/Ptpro/Fcgr3/Cd74/Coro1a/Jaml/Syk/Pgf/Vegfd/Anxa1/C5ar1/Ccn3/Itgam/Dpp4/Ccr7/Nckap1l/Lgals3/Cxcr5/Ccl11/Il16/Ripor2/Rarres2/Pla2g7/Nbl1/Thbs4/Wnt5a/Ccl24/Dusp1/Ccr2/Gpsm3/Ppbp/Trpm2/Serpine1/Dpep1/Pf4/Ccl5/Gpr183/Prex1/Csf1r/Vav3/Vegfc/Ccl27a/Thbs1/Itgb2/Bst1/Csf3r/Rpl13a/Csf1/Fcer1g/C3ar1/Vav1 |
| GO:0032635 | interleukin-6 production | 9,48E-06 | Pou2af1/Scimp/Ptpn22/Tlr1/Prg4/Bank1/Il33/Cd74/Syk/Vegfd/Il27ra/Irak3/Il18/Pou2f2/Lep/Nckap1l/Il16/Flt3/Rab7b/Cd84/Cd300ld/Tlr9/Wnt5a/Tlr7/Havcr2/Tnfsf9/Cebpb/Klf2/Cd200r1/Il1rl2/Syt11/Tlr4/Tlr8/Aqp4/Tyrobp/Card9/Spon2/Inpp5d/Unc93b1/Trem2/Ccr5/Isl1/Trim30a/Fcer1g/Sirpa/P2rx7 |
| GO:0045582 | positive regulation of T cell differentiation | 1,60E-05 | Pck1/Rhoh/Il7r/Sash3/Ikzf1/Lck/Rasgrp1/Cd74/Ptprc/Syk/H2-Aa/Anxa1/Runx3/H2-DMa/Myb/Il2rg/Il2ra/Il18/Ccr7/Nckap1l/Vsir/Carmil2/Tnfsf9/Cd1d1/Cd83/Ccr2/Il1rl2/Nfkbid |
| GO:0008037 | cell recognition | 2,04E-05 | Ighg2b/Ighv1-53/Ighv11-2/Ighv1-55/Ighg3/Ighg2c/Igkc/Ighd/Ighm/Dock2/Iglc2/Prtg/Igha/Cd209b/Fcgr3/Trbc2/Ccr7/Fcgr1/Lgals3/Vstm2l/Spa17/Sema3a/Ephb3/Wnt5a/Dock8/Havcr2/Cdk5r1/Colec12/Efnb3/Tnfrsf21/Cd6/B4galt1/Folr2/Ncam1/Nexn/Cadm1/Emb/Sirpa/Nptn/Ptprz1 |
| GO:1903706 | regulation of hemopoiesis | 2,35E-05 | Hoxa5/Pck1/Cd4/Rhoh/Il7r/Adipoq/Sash3/Ccr1/Fos/Ikzf1/Card11/Lck/Rasgrp1/Nfam1/Cd74/Ptprc/Syk/Spi1/H2-Aa/Anxa1/Runx3/Lrrc17/H2-DMa/Myb/Itgam/Il2rg/Il2ra/Il18/Ikzf3/Ccr7/Lep/Nckap1l/C1qc/Flt3/Car2/Axl/Rab7b/Cd28/Tlr9/Vsir/Hcls1/Carmil2/Zfp36/Tnfsf9/Cebpb/Cd1d1/Bmp4/Cd83/P4htm/Ccr2/Itgb3/Rassf2/Ubash3b/Myc/Il1rl2/Lif/Nfkbid/Thpo/B2m/Cd44/Kitl/Pf4/Tyrobp/Runx1/Apcs/Ccl5/Trib1/Klf13/Csf1r/Erbb2/Rorc/Inpp5d/Zbtb16/Trem2/Lmo2/Tgfbr2/Zeb1/Stat1/Jun/Csf3r/Dlk1/Csf1/Mturn/Fas/H2-M3/Tcf7/Tob2/Hmgb2/Ptk2b |
| GO:0050866 | negative regulation of cell activation | 2,42E-05 | Lax1/Ptpn22/Slfn1/Vsig4/Bank1/Btla/Blk/Btk/Tspan32/Samsn1/Lgals7/Spn/Cd74/Cd37/Fcgr2b/Tbc1d10c/H2-Aa/Anxa1/Runx3/H2-Ab1/Il2ra/Lgals3/Ripor2/Flt3/Axl/Cd84/Cd86/Vtcn1/Vsir/Havcr2/Cebpb/Cd300a/Bmp4/Fgl1/Cd274/Ccr2/Adora2a/Arg2/Pparg/Ubash3b/C1qtnf1/Nfkbid/Parp3/Tnfrsf13b/Tnfrsf21/Syt11/Cd44/Tyrobp/Runx1/Hmox1/Erbb2/Hfe/Inpp5d/Pla2g2f/Adrb2 |
| GO:0045785 | positive regulation of cell adhesion | 3,07E-05 | Ptpn22/Alox15/Pck1/Cd4/Rhoh/Il7r/Sash3/Cxcl13/Ikzf1/Card11/Lck/Rasgrp1/Spn/Cd74/Coro1a/Ptprc/Cd3e/Syk/Enpp2/H2-Aa/Ndnf/Anxa1/Runx3/Rasal3/H2-DMa/Myb/Il27ra/H2-Ab1/Dpp4/Il2rg/Il2ra/Il18/Vcam1/Fn1/Ccr7/P2ry12/Fbln1/Slc4a1/Dbn1/Lep/Nckap1l/Apbb1ip/Vit/Pdpn/Chrd/Cd28/Cd86/Vtcn1/Vsir/Wnt5a/Carmil2/Dock8/Spock2/Thy1/Havcr2/Nr4a3/Tnfsf9/Cd1d1/Ccdc80/Cd83/S100a10/Cd274/Ccdc88b/Ccn1/Ccr2/Itgb3/Il1rl2/Lif/Nfkbid/Efnb3/Cd44/Ephb6/Icam1/H2-T23/Runx1/Ccl5/Cd6/Prex1/Skap1/Erbb2/Tnfsf13b/Vav3/Icosl/Dab2/Nid1/Zbtb16/Vegfc/Ccr5/Emp2/Tgfbr2/Itga4/Fut4/Thbs1/Cav1/Itgal/Abi3bp/Itgb2/Lims2/Dusp26/Csf1/Sirpa/H2-M3/Vav1/Rin2/Cyth3/Plpp3/Ptk2b/Adk/Spta1/Il12rb1 |
| GO:0045621 | positive regulation of lymphocyte differentiation | 3,17E-05 | Pck1/Rhoh/Il7r/Sash3/Ikzf1/Lck/Rasgrp1/Cd74/Ptprc/Syk/H2-Aa/Anxa1/Runx3/H2-DMa/Myb/Il2rg/Il2ra/Il18/Ccr7/Nckap1l/Axl/Vsir/Carmil2/Tnfsf9/Cd1d1/Cd83/Ccr2/Il1rl2/Nfkbid |
| GO:0002821 | positive regulation of adaptive immune response | 3,59E-05 | Ighg2b/Fcer2a/Cd4/Sash3/Cd40/Btk/H2-Q7/Fcgr3/H2-Q6/Cd74/Ptprc/C3/Anxa1/H2-DMa/Il27ra/H2-Ab1/Il18/Ccr7/Fcgr1/Pla2g4a/Slc11a1/Cd28/Cd1d1/Cd274/Ccr2/Nfkbid/B2m/Hspa8/Cd44/H2-T23/Azgp1/Skap1/Tnfsf13b/H2-T10/Paxip1/Fcer1g/H2-M3/P2rx7/H2-T24/Il12rb1 |
| GO:0030183 | B cell differentiation | 3,59E-05 | Pou2af1/Igkc/Cr2/Ms4a1/Cd19/Cd79b/Ighm/Cd79a/Ikzf1/Btk/Card11/Nfam1/Ptprc/Syk/Myb/H2-Ab1/Il2rg/Ikzf3/Pou2f2/Nckap1l/Flt3/Tlr9/Dock10/Ntrk1/Mfng/Lyl1/Itm2a/Fzd9/Gpr183/Dclre1c/Inpp5d |
| GO:0050671 | positive regulation of lymphocyte proliferation | 3,62E-05 | Ptpn22/Ighd/Cd4/Ighm/Sash3/Cd40/Card11/Spn/Cd74/Coro1a/Ptprc/Cd3e/Syk/Anxa1/Rasal3/Il27ra/Il2ra/Il18/Vcam1/Ccr7/Slc4a1/Lep/Nckap1l/Cd28/Tlr9/Cd86/Vtcn1/Carmil2/Havcr2/Tnfsf9/Cd1d1/Cd274/Ccdc88b/Ccr2/Tlr4/H2-T23/Cd38/Ccl5/Gpr183/Cd6/Tnfsf13b/Vav3/Icosl/Tgfbr2/Itgal/Irs2/Bst1 |
| GO:0032946 | positive regulation of mononuclear cell proliferation | 4,26E-05 | Ptpn22/Ighd/Cd4/Ighm/Sash3/Cd40/Card11/Spn/Cd74/Coro1a/Ptprc/Cd3e/Syk/Anxa1/Rasal3/Il27ra/Il2ra/Il18/Vcam1/Ccr7/Slc4a1/Lep/Nckap1l/Cd28/Tlr9/Cd86/Vtcn1/Carmil2/Havcr2/Tnfsf9/Cd1d1/Cd274/Ccdc88b/Ccr2/Tlr4/H2-T23/Cd38/Ccl5/Gpr183/Cd6/Tnfsf13b/Vav3/Icosl/Tgfbr2/Itgal/Irs2/Bst1/Csf1/Adk/Spta1/Il12rb1 |
| GO:0032675 | regulation of interleukin-6 production | 4,68E-05 | Pou2af1/Scimp/Ptpn22/Tlr1/Prg4/Bank1/Il33/Cd74/Syk/Vegfd/Il27ra/Irak3/Pou2f2/Nckap1l/Il16/Flt3/Rab7b/Cd84/Cd300ld/Tlr9/Wnt5a/Tlr7/Havcr2/Tnfsf9/Cebpb/Klf2/Cd200r1/Il1rl2/Syt11/Tlr4/Tlr8/Aqp4/Tyrobp/Card9/Spon2/Inpp5d/Unc93b1/Trem2/Ccr5/Isl1/Trim30a/Fcer1g/Sirpa/P2rx7 |
| GO:0030888 | regulation of B cell proliferation | 5,14E-05 | Ighd/Siglecg/Mzb1/Cd22/Ighm/Btla/Blk/Sash3/Cd40/Btk/Card11/Cd74/Ptprc/Fcgr2b/Ikzf3/Nckap1l/Tlr9/Cd300a/Tnfrsf13b/Tnfrsf21/Tlr4/Tyrobp/Cd38/Gpr183/Tnfsf13b/Vav3/Inpp5d |
| GO:0070665 | positive regulation of leukocyte proliferation | 5,16E-05 | Ptpn22/Ighd/Cd4/Ighm/Sash3/Cd40/Card11/Spn/Cd74/Coro1a/Ptprc/Cd3e/Syk/Anxa1/Rasal3/Il27ra/Il2ra/Il18/Vcam1/Ccr7/Slc4a1/Lep/Nckap1l/Cd28/Tlr9/Cd86/Vtcn1/Carmil2/Havcr2/Tnfsf9/Cd1d1/Cd274/Ccdc88b/Ccr2/Kitl/Tlr4/H2-T23/Cd38/Ccl5/Gpr183/Cd6/Csf1r/Tnfsf13b/Vav3/Icosl/Tgfbr2/Itgal/Irs2/Bst1/Csf1/Adk/Spta1/Il12rb1 |
| GO:0045619 | regulation of lymphocyte differentiation | 6,27E-05 | Pck1/Rhoh/Il7r/Sash3/Ikzf1/Card11/Lck/Rasgrp1/Nfam1/Cd74/Ptprc/Syk/H2-Aa/Anxa1/Runx3/H2-DMa/Myb/Il2rg/Il2ra/Il18/Ikzf3/Ccr7/Nckap1l/Flt3/Axl/Cd28/Tlr9/Vsir/Carmil2/Tnfsf9/Cd1d1/Bmp4/Cd83/Ccr2/Il1rl2/Nfkbid/Cd44/Runx1/Erbb2/Rorc/Inpp5d/Zbtb16/Tgfbr2/Zeb1 |
| GO:0071219 | cellular response to molecule of bacterial origin | 6,31E-05 | Stap1/Scimp/Ptpn22/Tlr1/Cxcl13/Fcgr2b/Ly86/Il18/Cd180/Axl/Cd84/Gbp2/Vim/Cd86/Wnt5a/Cd68/Irf8/Lilrb4a/Havcr2/Zfp36/Cebpb/Cd274/Casp1/Ppbp/Serpine1/Bmp6/Mrc1/B2m/Tlr4/Pf4/Ccl5/Trib1/Cd6/Spon2/Tnfrsf1b/Plscr4/Trem2/Plscr1/Stat1 |
| GO:0002706 | regulation of lymphocyte mediated immunity | 6,83E-05 | Ighg2b/Fcer2a/Il7r/Sash3/Cd40/Btk/H2-Q7/Fcgr3/Rasgrp1/H2-Q6/Spn/Ptprc/Fcgr2b/C3/Il27ra/Dpp4/Il18/Fcgr1/Lep/Cd28/Vsir/Cd55/Havcr2/Cd1d1/Ccr2/Parp3/B2m/Hspa8/H2-T23/Azgp1/Hfe/H2-T10/Tnfrsf1b/Paxip1/Cadm1/Fcer1g/Tap1/H2-M3/P2rx7/Vav1/Muc4/Was/Arg1/H2-T24/Hmces/Tnfsf13/Il18r1/Lag3/Arrb2/Clec2d/Gfer/Stx7 |
| GO:0097530 | granulocyte migration | 7,76E-05 | Sell/Rhoh/Ccl8/Ccl6/Mcoln2/Rac2/Ccl9/Cxcl13/Fcgr3/Cd74/Jaml/Syk/Anxa1/C5ar1/Itgam/Dpp4/Ccr7/Nckap1l/Lgals3/Ccl11/Ripor2/Rarres2/Thbs4/Ccl24/Cd300a/Ppbp/Dpep1/Pf4/Ccl5/Prex1/Csf1r/Vav3/Thbs1/Itgb2/Bst1/Csf3r/Csf1/Fcer1g/Sirpa/C3ar1/Vav1 |
| GO:0043410 | positive regulation of MAPK cascade | 8,04E-05 | Scimp/Ptpn22/Alox15/Cd4/Marco/Gadd45g/Ccl8/Ighm/Bank1/Ccl6/Gadd45b/Ccr1/Cd40/Adrb3/Ccl9/Rasgrp1/Adra1b/Cd74/Ptprc/Syk/Fcgr2b/Fgf1/C3/Igfbp6/Nox4/C5ar1/Tbx1/Ccr7/Fgf9/Fgd2/Trf/Lep/Ccl11/Nek10/Gpr39/Cd84/Tlr9/Mfhas1/Wnt5a/Ccl24/Mdfi/Timp2/Xdh/Havcr2/Sorbs3/Peli2/Bmp4/Ncf1/Gadd45a/Glipr2/Igfbp4/Sphk1/Ntrk1/Map3k6/Pik3cg/Lrrk2/Itgb3/Rassf2/Ccn2/Ackr3/C1qtnf1/Traf1/Mst1r/Lif/Thpo/Cd44/Kitl/Tlr4/Icam1/Epha8/Bmp2/Ar/Pik3r5/Igfbp3/Ccl5/Tnik/Gpr183/Card9/Nrxn1/Cdon/Map4k1/Iqgap3/Csf1r/Erbb2/Dab2/Cavin3/Tgfa/Adrb2/Trem2/Esr1/Vangl2/Dnajc27/Thbs1/Mapk8ip2/Jun/Zeb2/Lepr/Ankrd6/Tgfb3/Ndrg4/Hand2/Cdh2/P2rx7/Igf1r |
| GO:0046634 | regulation of alpha-beta T cell activation | 8,28E-05 | Ptpn22/Btla/Sash3/Ikzf1/Lgals7/Ptprc/Cd3e/Syk/Anxa1/Runx3/Rasal3/Myb/H2-Ab1/Il2rg/Il18/Ccr7/Nckap1l/Cd28/Vsir/Cd1d1/Cd300a/Cd83/Cd274/Ccr2/Adora2a/Arg2/Nfkbid/Cd44/H2-T23/Runx1/Hfe/Zbtb16/Tgfbr2 |
| GO:0002478 | antigen processing and presentation of exogenous peptide antigen | 8,69E-05 | H2-DMb2/H2-DMb1/Fcgr3/Cd74/H2-Eb1/Fcgr2b/H2-Aa/H2-DMa/H2-Ab1/Ctse/Fcgr1/Ctss/Clec4a2/Ifi30/B2m/H2-T23/Unc93b1/Fcer1g/Tap1/H2-M3 |
| GO:0070664 | negative regulation of leukocyte proliferation | 1,02E-04 | Slfn1/Vsig4/Btla/Blk/Il33/Btk/Lgals7/Spn/Fcgr2b/H2-Aa/H2-Ab1/Il2ra/Ripor2/Cd86/Vtcn1/Vsir/Havcr2/Cebpb/Cd300a/Bmp4/Cd274/Arg2/Tnfrsf13b/Tnfrsf21/Cd44/Tyrobp/Erbb2/Inpp5d/Pla2g2f/H2-M3/Pla2g2d/Gal/Arg1 |
| GO:0034341 | response to interferon-gamma | 1,03E-04 | Ccl8/Ccl6/Cd40/Ccl9/H2-Q7/Gbp8/H2-Eb1/H2-Aa/H2-Ab1/Stx11/Ccl11/Rab7b/Gbp2/Slc11a1/Nlrc5/Vim/Nmi/Ccl24/Irf8/Ciita/Casp1/Pparg/Actg1/Mrc1/Tlr4/Aqp4/Ccl5/Evl/Gsn/Rab20/Ifitm1/Stat1/Rpl13a/Parp14/Sirpa/Was/Arg1/Gbp2b/Capg/Il12rb1 |
| GO:0050764 | regulation of phagocytosis | 1,04E-04 | Stap1/Ighg2b/Alox15/Adipoq/Dock2/Il2rb/Cd209b/Fcgr3/Ptprc/Syk/Fcgr2b/C3/Il2rg/Ccr7/Fcgr1/Nckap1l/Slc11a1/Cd300a/Sphk1/Pparg/Syt11/Hck/C2/Hspa8/Rab31/Siglece/Trem2/Pros1/Cnn2/Fcer1g/Sirpa |
| GO:0019884 | antigen processing and presentation of exogenous antigen | 1,14E-04 | H2-DMb2/H2-DMb1/Fcgr3/Cd74/H2-Eb1/Fcgr2b/H2-Aa/H2-DMa/H2-Ab1/Ctse/Fcgr1/Ctss/Clec4a2/Cd1d1/Ifi30/B2m/H2-T23/Unc93b1/Fcer1g/Tap1/H2-M3 |
| GO:1903555 | regulation of tumor necrosis factor superfamily cytokine production | 1,19E-04 | Ptpn22/Cd2/Adipoq/Sash3/Rasgrp1/Ptprc/Irak3/Il18/Ccr7/Lep/Clec4a2/Flt3/Axl/Cd300ld/Tlr9/Vsir/Havcr2/Zfp36/Cd34/Cd274/Arg2/Tlr4/Pf4/H2-T23/Card9/Spon2/Trem2/Ccr5/Isl1/Trim30a/Abcc8/Fcer1g/Sirpa |
| GO:0002824 | positive regulation of adaptive immune response based on somatic recombination of immune receptors built from immunoglobulin superfamily domains | 1,22E-04 | Ighg2b/Fcer2a/Cd4/Sash3/Cd40/Btk/H2-Q7/Fcgr3/H2-Q6/Ptprc/C3/Anxa1/H2-DMa/Il27ra/H2-Ab1/Il18/Ccr7/Fcgr1/Pla2g4a/Slc11a1/Cd28/Cd1d1/Cd274/Ccr2/Nfkbid/B2m/Hspa8/H2-T23/Azgp1/Tnfsf13b/H2-T10/Paxip1/Fcer1g/H2-M3/P2rx7/H2-T24/Il12rb1 |
| GO:0050729 | positive regulation of inflammatory response | 1,69E-04 | Stap1/Ighg2b/Zbp1/Il1rl1/Il33/Btk/Fcgr3/C3/Ccr7/Fcgr1/Ctss/Pla2g4a/Il16/Cd28/Tlr9/Wnt5a/Ccl24/Gprc5b/Cebpb/Alox5ap/Ccr2/Pik3cg/Lrrk2/Gpsm3/Serpine1/Tlr4/Ffar3/H2-T23/Ccl5/Cd6 |
| GO:0032945 | negative regulation of mononuclear cell proliferation | 1,78E-04 | Slfn1/Vsig4/Btla/Blk/Btk/Lgals7/Spn/Fcgr2b/H2-Aa/H2-Ab1/Il2ra/Ripor2/Cd86/Vtcn1/Vsir/Havcr2/Cebpb/Cd300a/Bmp4/Cd274/Arg2/Tnfrsf13b/Tnfrsf21/Cd44/Tyrobp/Erbb2/Inpp5d/Pla2g2f/H2-M3/Pla2g2d/Gal/Arg1 |
| GO:0002831 | regulation of response to biotic stimulus | 1,82E-04 | Coch/Scimp/Ptpn22/Treml2/Zbp1/Vsig4/Ifi209/Ccr1/Tspan32/Rasgrp1/Spn/Cd74/Cd37/Ifi204/Ifi205/Irak3/Il2ra/Ly86/Myo1f/Cd180/Aim2/Lep/Apobec3/Smpdl3b/Cd84/Nlrc5/Nmi/Tlr9/Wnt5a/Serping1/Havcr2/Mill2/Cd274/Mndal/Arg2/Tspan6/Pparg/Ifi211/Syt11/Bmp6/Tlr4/Tlr8/Mmp2/H2-T23/Htra1/Ccl5/Trib1/Card9/Trem2/Irf7/Plscr1/Stat1/Ifi203/Cadm1/Parp14/Tap1/H2-M3/Vav1/Hmgb2/Muc4/Arg1/Lrp8/Il12rb1/Dtx3l/Irgm2/Gpatch3/Lag3/Ffar2/Arrb2/Clec2d/C1qbp/Gfer/Apoe/Dusp10/Sting1/Txk |
| GO:0046633 | alpha-beta T cell proliferation | 1,82E-04 | Ptpn22/Btla/Dock2/Lgals7/Ptprc/Cd3e/Syk/Rasal3/Il18/Cd28/Vsir/Lilrb4a/Elf4/Cd274/Ccr2/Arg2/Cd44/H2-T23 |
| GO:0007162 | negative regulation of cell adhesion | 1,96E-04 | Lax1/Ptpn22/Slfn1/Vsig4/Adipoq/Btla/Fam107a/Lgals7/Spn/Cd74/Tnc/Plxnc1/Ptprc/Enpp2/H2-Aa/Anxa1/Runx3/Dab1/H2-Ab1/Nat8f2/Il2ra/Myo1f/Sema3e/Efna5/Epha5/Fbln1/Lgals3/Akna/Ripor2/Nat8f5/Cd86/Vtcn1/Vsir/Nat8/Havcr2/Cebpb/Cd300a/Bmp4/Fgl1/Cd274/Dusp1/Adora2a/Arg2/Ubash3b/C1qtnf1/Serpine1/Rgcc/Nfkbid/Tnfrsf21/Bmp6/Tnr/Cd44/Bmp2/Mmp2/Runx1/Coro2b/Plxna4/Lpxn/Erbb2/Hfe/Postn/Pla2g2f/Adrb2 |
| GO:0002705 | positive regulation of leukocyte mediated immunity | 1,99E-04 | Stap1/Ighg2b/Scimp/Fcer2a/Sash3/Cd40/Btk/H2-Q7/Fcgr3/Rasgrp1/H2-Q6/Ptprc/Syk/C3/Itgam/Il18/Fcgr1/Cd28/Cd1d1/B2m/Hspa8/H2-T23/Tyrobp/Azgp1/H2-T10/Itgb2/Paxip1/Cadm1/Fcer1g/H2-M3/P2rx7/Vav1/Arg1/H2-T24 |
| GO:0031349 | positive regulation of defense response | 2,12E-04 | Stap1/Coch/Ighg2b/Zbp1/Il1rl1/Ifi209/Il33/Btk/Fcgr3/Rasgrp1/Cd74/Ifi204/Ifi205/C3/Ccr7/Fcgr1/Aim2/Ctss/Cckbr/Pla2g4a/Il16/Nlrc5/Cd28/Tlr9/Wnt5a/Ccl24/Havcr2/Gprc5b/Cebpb/Alox5ap/Ccr2/Mndal/Pik3cg/Lrrk2/Gpsm3/Serpine1/Ifi211/Tlr4/Ffar3/Tlr8/Mmp2/H2-T23/Ccl5/Cd6/Card9/Trem2/Irf7/Ccr5/Plscr1/Ifi203/Cadm1/Fcer1g/H2-M3/Vav1/Hmgb2/Arg1 |
| GO:0050672 | negative regulation of lymphocyte proliferation | 2,25E-04 | Slfn1/Vsig4/Btla/Blk/Btk/Lgals7/Spn/Fcgr2b/H2-Aa/H2-Ab1/Il2ra/Ripor2/Cd86/Vtcn1/Vsir/Havcr2/Cebpb/Cd300a/Bmp4/Cd274/Arg2/Tnfrsf13b/Tnfrsf21/Cd44/Tyrobp/Erbb2/Inpp5d/Pla2g2f/H2-M3/Pla2g2d/Gal/Arg1 |
| GO:0032649 | regulation of interferon-gamma production | 2,44E-04 | Ptpn22/Cd2/Il1rl1/Sash3/Il33/H2-Q7/Rasgrp1/Lgals7/Cd3e/Il27ra/Il18/Ccr7/Flt3/Axl/Slc11a1/Tlr9/Vsir/Wnt5a/Carmil2/Tlr7/Irf8/Havcr2/Tnfsf9/Cd1d1/Cd274/Ccr2/Tlr4/Tlr8/Runx1 |
| GO:0071216 | cellular response to biotic stimulus | 2,56E-04 | Stap1/Scimp/Ptpn22/Tlr1/Cxcl13/Btk/Syk/Fcgr2b/Ly86/Il18/Cd180/Txnip/Axl/Cd84/Gbp2/Vim/Cd86/Wnt5a/Cd68/Irf8/Lilrb4a/Havcr2/Zfp36/Cebpb/Cd274/Casp1/Ppbp/Serpine1/Bmp6/Mrc1/B2m/Tlr4/Pf4/Ccl5/Trib1/Cd6/Spon2/Tnfrsf1b/Plscr4/Trem2/Plscr1/Stat1 |
| GO:0007059 | chromosome segregation | 2,56E-04 | Sycp1/Esco2/Tex15/Ccnb1/Ndc80/Cenpe/Mki67/Knl1/Nek2/Cenpf/Aurkb/Nuf2/Dlgap5/Septin1/Knstrn/Bub1b/Ube2c/Ncaph/Rgs14/Haspin/Top2a/Gem/Cdc6/Cdc20/Nusap1/Cep85/Kif4/Ccne1/Cdca2/Kif22/Birc5/Ect2/Fmn2/Racgap1/Smc4/Kif23/Rad18/Psrc1/Mad2l1bp/Ncapg/Spag5/Mis18a/Dsn1/Cdca8/Espl1/A730008H23Rik/Axin2 |
| GO:0002685 | regulation of leukocyte migration | 2,74E-04 | Stap1/Sell/Ptpn22/Rhoh/Rac2/Ccr1/Il33/Cxcl13/Spn/Cd74/Pgf/Vegfd/Anxa1/C5ar1/Ccn3/Il27ra/Dpp4/Ccr7/P2ry12/Nckap1l/Lgals3/Ripor2/Rarres2/Pla2g7/Nbl1/Thbs4/Wnt5a/Ccl24/Dock8/Thy1/Cd300a/Dusp1/Ccr2/Itgb3/Cd200r1/Gpsm3/Serpine1/Kitl/Icam1/Stk10/Ccl5/Ecm1/Smpd3/Csf1r/Trem2/Vegfc/Ccl27a/Itga4/Fut4/Mmp9/Thbs1/Bst1/Csf1/C3ar1 |
| GO:0071346 | cellular response to interferon-gamma | 2,83E-04 | Ccl8/Ccl6/Ccl9/H2-Q7/Gbp8/H2-Ab1/Stx11/Ccl11/Rab7b/Gbp2/Nlrc5/Vim/Nmi/Ccl24/Irf8/Casp1/Pparg/Actg1/Mrc1/Tlr4/Aqp4/Ccl5/Evl/Gsn/Rab20/Stat1/Rpl13a/Parp14/Sirpa/Was/Arg1/Gbp2b/Capg/Il12rb1 |
| GO:0045580 | regulation of T cell differentiation | 3,12E-04 | Pck1/Rhoh/Il7r/Sash3/Ikzf1/Card11/Lck/Rasgrp1/Cd74/Ptprc/Syk/H2-Aa/Anxa1/Runx3/H2-DMa/Myb/Il2rg/Il2ra/Il18/Ccr7/Nckap1l/Cd28/Vsir/Carmil2/Tnfsf9/Cd1d1/Bmp4/Cd83/Ccr2/Il1rl2/Nfkbid/Cd44/Runx1/Erbb2/Rorc/Zbtb16/Tgfbr2/Zeb1 |
| GO:0042102 | positive regulation of T cell proliferation | 3,18E-04 | Ptpn22/Cd4/Sash3/Card11/Spn/Coro1a/Ptprc/Cd3e/Syk/Anxa1/Rasal3/Il27ra/Il2ra/Il18/Vcam1/Ccr7/Slc4a1/Lep/Nckap1l/Cd28/Cd86/Vtcn1/Carmil2/Havcr2/Tnfsf9/Cd1d1/Cd274/Ccdc88b/Ccr2/H2-T23/Ccl5/Cd6/Tnfsf13b/Icosl/Tgfbr2/Itgal |
| GO:0002237 | response to molecule of bacterial origin | 3,92E-04 | Stap1/Scimp/Ptpn22/Tlr1/Cxcl13/Slpi/Ncf2/Fcgr2b/Irf5/C5ar1/Irak3/Ly86/Il18/Ccr7/Cd180/Axl/Cd84/Gbp2/Slc11a1/Vim/Tlr9/Cd86/Wnt5a/Cd68/Irf8/Lilrb4a/Havcr2/Zfp36/Cebpb/Ptgfr/Cd274/Casp1/Ppbp/Serpine1/Bmp6/Mrc1/B2m/Tlr4/Pf4/Ccl5/Ggt5/Ptgir/Trib1/Cd6/Card9/Spon2/Tnfrsf1b/Plscr4/Trem2/Noct/Plscr1/Stat1/Rpl13a/Sirpa/H2-M3/P2rx7/Adm |
| GO:1990266 | neutrophil migration | 4,30E-04 | Sell/Rhoh/Ccl8/Ccl6/Mcoln2/Rac2/Ccl9/Cxcl13/Fcgr3/Cd74/Jaml/Syk/C5ar1/Itgam/Dpp4/Ccr7/Nckap1l/Lgals3/Ccl11/Ripor2/Thbs4/Ccl24/Ppbp/Dpep1/Pf4/Ccl5/Prex1/Vav3/Itgb2/Bst1/Csf3r/Fcer1g/C3ar1/Vav1 |
| GO:0045088 | regulation of innate immune response | 4,35E-04 | Coch/Treml2/Zbp1/Vsig4/Ifi209/Ccr1/Rasgrp1/Cd74/Ifi204/Ifi205/Irak3/Myo1f/Aim2/Lep/Smpdl3b/Nlrc5/Nmi/Tlr9/Wnt5a/Serping1/Havcr2/Mndal/Pparg/Ifi211/Tlr4/Tlr8/Mmp2/H2-T23/Card9/Trem2/Irf7/Plscr1/Ifi203/Cadm1/Parp14/Tap1/H2-M3/Vav1/Hmgb2/Arg1/Lrp8/Irgm2/Lag3/Ffar2/Arrb2/Clec2d/Gfer/Apoe/Dusp10/Sting1/Txk |
| GO:0050868 | negative regulation of T cell activation | 4,72E-04 | Lax1/Ptpn22/Slfn1/Vsig4/Btla/Lgals7/Spn/Cd74/H2-Aa/Anxa1/Runx3/H2-Ab1/Il2ra/Lgals3/Ripor2/Cd86/Vtcn1/Vsir/Havcr2/Cebpb/Cd300a/Bmp4/Fgl1/Cd274/Adora2a/Arg2/Nfkbid/Tnfrsf21/Cd44/Runx1/Erbb2/Hfe/Pla2g2f |
| GO:0032680 | regulation of tumor necrosis factor production | 4,79E-04 | Ptpn22/Cd2/Adipoq/Sash3/Rasgrp1/Ptprc/Irak3/Il18/Ccr7/Lep/Clec4a2/Flt3/Axl/Cd300ld/Tlr9/Vsir/Havcr2/Zfp36/Cd34/Arg2/Tlr4/Pf4/H2-T23/Card9/Spon2/Trem2/Ccr5/Isl1/Trim30a/Abcc8/Fcer1g/Sirpa |
| GO:0071706 | tumor necrosis factor superfamily cytokine production | 4,81E-04 | Ptpn22/Cd2/Adipoq/Sash3/Rasgrp1/Ptprc/Irak3/Il18/Ccr7/Lep/Clec4a2/Flt3/Axl/Cd300ld/Tlr9/Vsir/Havcr2/Zfp36/Cd34/Cd274/Arg2/Tlr4/Pf4/H2-T23/Card9/Spon2/Trem2/Ccr5/Isl1/Trim30a/Abcc8/Fcer1g/Sirpa |
| GO:0046635 | positive regulation of alpha-beta T cell activation | 4,95E-04 | Ptpn22/Sash3/Ikzf1/Ptprc/Cd3e/Syk/Anxa1/Runx3/Rasal3/Myb/H2-Ab1/Il2rg/Il18/Ccr7/Nckap1l/Cd28/Cd1d1/Cd83/Ccr2/Nfkbid/H2-T23/Runx1/Zbtb16/Tgfbr2 |
| GO:0098813 | nuclear chromosome segregation | 5,02E-04 | Sycp1/Esco2/Tex15/Ccnb1/Ndc80/Cenpe/Knl1/Nek2/Cenpf/Aurkb/Nuf2/Septin1/Knstrn/Bub1b/Ube2c/Ncaph/Haspin/Top2a/Gem/Cdc6/Cdc20/Nusap1/Kif4/Ccne1/Kif22/Birc5/Ect2/Fmn2/Racgap1/Smc4/Kif23/Psrc1/Mad2l1bp/Ncapg/Spag5/Dsn1/Cdca8/Espl1/Axin2 |
| GO:0002444 | myeloid leukocyte mediated immunity | 5,14E-04 | Stap1/Ighg2b/Rac2/Blk/Btk/Fcgr3/Dnase1l3/Rasgrp1/Syk/C3/Itgam/Myo1f/Stx11/Fcgr1/Cd84/Nr4a3/Cd300a/Ncf1/Ccr2/Lat2/H2-T23/Tyrobp/Hmox1/Spon2 |
| GO:0032640 | tumor necrosis factor production | 5,57E-04 | Ptpn22/Cd2/Adipoq/Sash3/Rasgrp1/Ptprc/Irak3/Il18/Ccr7/Lep/Clec4a2/Flt3/Axl/Cd300ld/Tlr9/Vsir/Havcr2/Zfp36/Cd34/Arg2/Tlr4/Pf4/H2-T23/Card9/Spon2/Trem2/Ccr5/Isl1/Trim30a/Abcc8/Fcer1g/Sirpa |
| GO:0071222 | cellular response to lipopolysaccharide | 5,90E-04 | Stap1/Scimp/Ptpn22/Cxcl13/Ly86/Il18/Cd180/Axl/Cd84/Gbp2/Vim/Cd86/Cd68/Irf8/Lilrb4a/Havcr2/Zfp36/Cebpb/Cd274/Casp1/Ppbp/Serpine1/Bmp6/Mrc1/B2m/Tlr4/Pf4/Ccl5/Trib1/Cd6/Spon2/Tnfrsf1b/Plscr4/Trem2/Plscr1/Stat1 |
| GO:0002700 | regulation of production of molecular mediator of immune response | 5,91E-04 | Scimp/Ptpn22/Siglecg/Mzb1/Cd22/Ighm/Sash3/Cd40/Il33/Cd74/Ptprc/Cd37/Fcgr2b/Il27ra/Irak3/Il18/Cd28/Tlr9/Vsir/Wnt5a/Nr4a3/Gprc5b/Ccr2/Parp3/B2m/Tlr4/Ffar3/H2-T23/Hmox1/Spon2/Hfe/Tnfrsf1b/Paxip1/Tgfb3/Fcer1g/Vpreb3/H2-M3/Arg1/Hmces/Hk1/Tnfsf13/Il18r1/Jak3/Tgfb2/Ffar2/Tek |
| GO:0050854 | regulation of antigen receptor-mediated signaling pathway | 5,91E-04 | Stap1/Ptpn22/Cd19/Cd22/Blk/Card11/Lck/Nfam1/Ptprc/Fcgr2b/Ccr7/Lgals3/Thy1/Cd300a |
| GO:0001818 | negative regulation of cytokine production | 6,07E-04 | Ptpn22/Prg4/Vsig4/Adipoq/Bank1/Il1rl1/Il33/Btk/Lgals7/Ptprc/Fcgr2b/Anxa1/Wnt11/Il27ra/Errfi1/Irak3/Ccr7/Ffar1/Clec4a2/Nckap1l/Flt3/Axl/Cd84/Slc11a1/Nmi/Tlr9/Vsir/Ptprs/Havcr2/Zfp36/Cd34/Traip/Cd83/Cd274/Klf2/Nav3/Cd200r1/Arg2/Pparg/Rgcc/Tnfrsf21/Syt11/Abcd2/Tlr4/Tlr8/Adcy7/Aqp4/Tyrobp/Hmox1/Inhbb/Hfe/Inpp5d/Trem2/Trim30a/Thbs1/Srgn/Tgfb3/Sirpa/Nptn |
| GO:0002504 | antigen processing and presentation of peptide or polysaccharide antigen via MHC class II | 6,41E-04 | H2-Ob/H2-DMb2/H2-DMb1/Cd74/H2-Eb1/Marchf1/Fcgr2b/H2-Aa/H2-DMa/H2-Ab1/Ctse/Ctss/Ifi30/Unc93b1/Trem2/Thbs1/Fcer1g |
| GO:0046640 | regulation of alpha-beta T cell proliferation | 7,32E-04 | Ptpn22/Btla/Lgals7/Ptprc/Cd3e/Syk/Rasal3/Il18/Cd28/Vsir/Cd274/Ccr2/Arg2/Cd44/H2-T23 |
| GO:0006482 | protein demethylation | 8,18E-04 | Kdm2b/Uty/Kdm5d |
| GO:0008214 | protein dealkylation | 8,18E-04 | Kdm2b/Uty/Kdm5d |
| GO:0071621 | granulocyte chemotaxis | 8,50E-04 | Sell/Ccl8/Ccl6/Rac2/Ccl9/Cxcl13/Fcgr3/Cd74/Jaml/Syk/Anxa1/C5ar1/Itgam/Dpp4/Ccr7/Nckap1l/Lgals3/Ccl11/Ripor2/Rarres2/Thbs4/Ccl24/Ppbp/Dpep1/Pf4/Ccl5/Prex1/Csf1r/Vav3/Thbs1/Itgb2/Bst1/Csf3r/Csf1/Fcer1g/C3ar1/Vav1 |
| GO:0032743 | positive regulation of interleukin-2 production | 1,01E-03 | Sash3/Card11/Ptprc/Cd3e/Irf4/Anxa1/Cd28/Vtcn1/Carmil2/Cd1d1/Cd83/Ccr2/Runx1 |
| GO:0002495 | antigen processing and presentation of peptide antigen via MHC class II | 1,06E-03 | H2-Ob/H2-DMb2/H2-DMb1/Cd74/H2-Eb1/Marchf1/Fcgr2b/H2-Aa/H2-DMa/H2-Ab1/Ctse/Ctss/Ifi30/Unc93b1/Trem2/Fcer1g |
| GO:0030593 | neutrophil chemotaxis | 1,07E-03 | Sell/Ccl8/Ccl6/Rac2/Ccl9/Cxcl13/Fcgr3/Cd74/Jaml/Syk/C5ar1/Itgam/Dpp4/Ccr7/Nckap1l/Lgals3/Ccl11/Ripor2/Thbs4/Ccl24/Ppbp/Dpep1/Pf4/Ccl5/Prex1/Vav3/Itgb2/Bst1/Csf3r/Fcer1g/C3ar1/Vav1 |
| GO:0018108 | peptidyl-tyrosine phosphorylation | 1,10E-03 | Stap1/Ptpn22/Sla/Cd4/Sfrp2/Ighm/Adipoq/Bank1/Itk/Blk/Cd40/Btk/Lck/Samsn1/Reln/Cd74/Ptprc/Cd3e/Syk/Enpp2/Nox4/Errfi1/Il18/Efna5/Fcgr1/Lep/Efemp1/Thbs4/Hcls1/Thy1/Gprc5b/Cd300a/Ncf1/Ntrk1/Itgb3/Lif/Ddr2/Bmp6/Hck/Cd44/Kitl/Tlr4/Icam1/Ccl5/Csf1r/Tec/Erbb2/Tgfa/Trem2/Vegfc/Wee1/Isl1/Abi2/Bmx/Cav1/Itgb2/Bst1/Parp14/Igf1r/Ptprz1/Plpp3/Ptk2b/Lrp8/Il12rb1/Pdgfra/Fgf7/Hk1/Dmtn/Il12rb2/Arl2bp/Adra2a/Jak3/Pdgfrb/Epha7/Prnp/Tek/Arrb2/Cblb/Lyn/Txk/Acvr1/Adora1/Dyrk3/Prmt2/Vtn |
| GO:0007204 | positive regulation of cytosolic calcium ion concentration | 1,27E-03 | Gpr174/P2ry10/Ms4a1/Cd19/Cd4/Cd52/Ccr1/Cxcl13/Lck/Adra1b/Ackr4/Coro1a/Ptprc/Nmb/C5ar1/Cxcr6/Ccr7/Gna15/Grin2c/Ffar1/Ackr2/Cckbr/Cxcr5/Adrb1/Gpr39/Fzd2/Trpv2/Gpr65/Thy1/Avpr1a/Fam155a/Ptgfr/Bmp4/Akap5/Ccr2/Pik3cg/Itgb3/Slc8a1/Ackr3/Ubash3b/Cacna1g/C1qtnf1/Trpm2/Myo5a/Tmem28/Pth1r/Jph3/Xcr1/Cacnb3/Prkd1/Cd38/Ptgir/Gsto1/Grin1/Trpc1/Cacna1i/Adrb2/Cxcr4/Esr1/Ccr5/Galr2/Cav1/Cacna2d1/C3ar1/P2rx7/Adm/Lrp1/Ptk2b/Oxtr |
| GO:0031343 | positive regulation of cell killing | 1,27E-03 | Stap1/Fcer2a/Cd5l/H2-Q7/Rasgrp1/H2-Q6/Ptprc/Syk/Itgam/Cd1d1/B2m/Hspa8/H2-T23/Tyrobp/Azgp1/H2-T10/Ccr5/Cadm1/H2-M3/P2rx7/Vav1/Arg1/H2-T24 |
| GO:0071887 | leukocyte apoptotic process | 1,29E-03 | Il7r/Blk/Fcmr/Aurkb/Cd74/Fcgr2b/Anxa1/Itgam/Il2ra/Il18/Ccr7/Lgals3/Axl/Plekho2/Hcls1/Wnt5a/Dock8/Nr4a3/Bmp4/Cd274/Siglec1/Arg2/Myc/Nfkbid/Tnfrsf21/Pik3cd/Cd44/Kitl/Ccl5/Tsc22d3/Rorc/Ccr5/Irs2/Fcer1g/Fas/P2rx7 |
| GO:0016577 | histone demethylation | 1,29E-03 | Kdm2b/Uty/Kdm5d |
| GO:0070076 | histone lysine demethylation | 1,29E-03 | Kdm2b/Uty/Kdm5d |
| GO:0051480 | regulation of cytosolic calcium ion concentration | 1,29E-03 | Gpr174/P2ry10/Ms4a1/Cd19/Cd4/Cd52/Ccr1/Cxcl13/Lck/Adra1b/Ackr4/Coro1a/Ptprc/Nmb/C5ar1/Cxcr6/Calb1/Ccr7/Gna15/Grin2c/Ffar1/Ackr2/Atp1a2/Cckbr/Kcnk3/Cxcr5/Adrb1/Gpr39/Fzd2/Trpv2/Gpr65/Wnt5a/Thy1/Avpr1a/Fam155a/Ptgfr/Bmp4/Akap5/Ccr2/Pik3cg/Itgb3/Slc8a1/Ackr3/Ubash3b/Cacna1g/C1qtnf1/Trpm2/Myo5a/Tmem28/Pth1r/Jph3/Xcr1/Cacnb3/Fzd9/Prkd1/Cd38/Ptgir/Gsto1/Grin1/Smpd3/Trpc1/Npy1r/Cacna1i/Adrb2/Cxcr4/Esr1/Ccr5/Galr2/Cav1/Cacna2d1/Scgn/C3ar1/P2rx7/Adm/Lrp1/Ptk2b/Oxtr |
| GO:0009615 | response to virus | 1,31E-03 | Pou2af1/Ptpn22/Zbp1/Slfn8/Batf3/Cd40/Il33/Tspan32/H2-Q7/Oas1a/Spn/Ptprc/Cd37/Irf5/Ifi27l2a/Tlr13/Wdfy4/Irak3/Il2ra/Pou2f2/Aim2/Apobec3/Flt3/Oasl2/Tlr9/Cd86/Tlr7/Rnasel/Gli2/Mill2/Bnip3/Tspan6/Ddit4/Mst1r/Itgax/Oas2/Ifit1/Slfn9/Trim34a/Tlr8/Ifit2/Htra1/Lcn2/Ccl5/Card9/Ifit3b/Spon2/Ifitm1/Unc93b1/Irf7/Plscr1/Trim30a/Stat1 |
| GO:0002285 | lymphocyte activation involved in immune response | 1,32E-03 | Pou2af1/Cd19/Pck1/Gadd45g/Cd40/Exo1/Spn/Cd74/Ly9/Coro1a/Ptprc/Irf4/Anxa1/Myb/Il27ra/Il18/Stx11/Ccr7/Cd180/Lgals3/Apbb1ip/Slc11a1/Cd28/Dock10/Havcr2/Ccr2/Nfkbid/Parp3/Mfng/Itm2a/Tlr4/Icam1/Gpr183/Rorc/Icosl/Batf/Itgal/Paxip1/Fcer1g/H2-M3/Lcp1/Ptk2b |
| GO:0050766 | positive regulation of phagocytosis | 1,44E-03 | Stap1/Ighg2b/Dock2/Il2rb/Cd209b/Fcgr3/Ptprc/Fcgr2b/C3/Il2rg/Ccr7/Fcgr1/Nckap1l/Slc11a1/Pparg/C2/Hspa8/Rab31/Trem2/Pros1/Fcer1g/Sirpa/Lrp1 |
| GO:0022408 | negative regulation of cell-cell adhesion | 1,48E-03 | Lax1/Ptpn22/Slfn1/Vsig4/Adipoq/Btla/Lgals7/Spn/Cd74/H2-Aa/Anxa1/Runx3/H2-Ab1/Il2ra/Lgals3/Akna/Ripor2/Cd86/Vtcn1/Vsir/Havcr2/Cebpb/Cd300a/Bmp4/Fgl1/Cd274/Adora2a/Arg2/Ubash3b/C1qtnf1/Rgcc/Nfkbid/Tnfrsf21/Bmp6/Tnr/Cd44/Bmp2/Runx1/Erbb2/Hfe/Pla2g2f/Adrb2 |
| GO:1903038 | negative regulation of leukocyte cell-cell adhesion | 1,56E-03 | Lax1/Ptpn22/Slfn1/Vsig4/Btla/Lgals7/Spn/Cd74/H2-Aa/Anxa1/Runx3/H2-Ab1/Il2ra/Lgals3/Ripor2/Cd86/Vtcn1/Vsir/Havcr2/Cebpb/Cd300a/Bmp4/Fgl1/Cd274/Adora2a/Arg2/Nfkbid/Tnfrsf21/Cd44/Runx1/Erbb2/Hfe/Pla2g2f |
| GO:2000106 | regulation of leukocyte apoptotic process | 1,63E-03 | Il7r/Blk/Fcmr/Aurkb/Cd74/Fcgr2b/Anxa1/Il18/Ccr7/Lgals3/Axl/Hcls1/Wnt5a/Dock8/Nr4a3/Bmp4/Cd274/Siglec1/Arg2/Myc/Nfkbid/Pik3cd/Cd44/Kitl/Ccl5/Tsc22d3/Rorc/Ccr5/Irs2/Fcer1g/P2rx7 |
| GO:0048872 | homeostasis of number of cells | 1,86E-03 | Gpr174/Tex15/Hoxa5/Il7r/Sash3/Ikzf1/Card11/Ccnb2/Chst3/Cd74/Coro1a/Fcgr2b/Spi1/Anxa1/Ccn3/Myb/Itgam/Il2ra/Il18/Ccr7/Pirb/Adgrf4/Slc4a1/Ildr2/Nckap1l/Flt3/Axl/Klf1/Col14a1/Dock10/Mfhas1/Hcls1/Carmil2/Lilrb4a/Zfp36/Bmp4/Klf2/P4htm/Ccr2/Rassf2/Tnfrsf13b/Pik3cd/Cfh/B2m/Cd44/Kitl/Heph/Gpr183/Hmox1/Sh2b2/Tsc22d3/Klf13/Akt3/Tnfsf13b/Inpp5d/Lmo2/Stat1/Csf1/Fcer1g/Fas/Cdh2/P2rx7/Hmgb2/Gcnt4/Spta1/Tgfbr3/L3mbtl3/Gpam/Foxo3/Smo/Dmtn/Bcl2l11/Lpcat3/Slc37a4/Jak3/Stat3 |
| GO:0018212 | peptidyl-tyrosine modification | 2,01E-03 | Stap1/Ptpn22/Sla/Cd4/Sfrp2/Ighm/Adipoq/Bank1/Itk/Blk/Cd40/Btk/Lck/Samsn1/Reln/Cd74/Ptprc/Cd3e/Syk/Enpp2/Nox4/Errfi1/Il18/Efna5/Fcgr1/Lep/Efemp1/Thbs4/Hcls1/Thy1/Gprc5b/Cd300a/Ncf1/Ntrk1/Itgb3/Lif/Ddr2/Bmp6/Hck/Cd44/Kitl/Tlr4/Icam1/Ccl5/Csf1r/Tec/Erbb2/Tgfa/Trem2/Vegfc/Wee1/Isl1/Abi2/Bmx/Cav1/Itgb2/Bst1/Parp14/Igf1r/Ptprz1/Plpp3/Ptk2b/Lrp8/Il12rb1/Pdgfra/Fgf7/Hk1/Dmtn/Il12rb2/Arl2bp/Adra2a/Jak3/Pdgfrb/Epha7/Prnp/Tek/Arrb2/Cblb/Lyn/Txk/Acvr1/Adora1/Dyrk3/Prmt2/Vtn |
| GO:0019886 | antigen processing and presentation of exogenous peptide antigen via MHC class II | 2,07E-03 | H2-DMb2/H2-DMb1/Cd74/H2-Eb1/Fcgr2b/H2-Aa/H2-DMa/H2-Ab1/Ctse/Ctss/Ifi30 |
| GO:0002886 | regulation of myeloid leukocyte mediated immunity | 2,13E-03 | Stap1/Ighg2b/Rac2/Blk/Btk/Fcgr3/Dnase1l3/Syk/C3/Itgam/Fcgr1/Cd84/Cd300a/Ccr2/H2-T23/Tyrobp/Hmox1 |
| GO:0032663 | regulation of interleukin-2 production | 2,14E-03 | Vsig4/Sash3/Card11/Ptprc/Cd3e/Irf4/Anxa1/Cd28/Vtcn1/Carmil2/Havcr2/Zfp36/Cd34/Cd1d1/Cd83/Nav3/Ccr2/Runx1/Card9 |
| GO:0050830 | defense response to Gram-positive bacterium | 2,21E-03 | Il7r/Lyz1/Scd1/Gbp8/Lyz2/C5ar1/Il27ra/Myo1f/Npy/Reg3b/Rarres2/Gbp2/Vip/Gsdmd/Havcr2/Ncf1/Hck/B2m/H2-T23/Mpeg1/Card9/Defb1/H2bc6/P2rx7/Adm/Hmgb2/Gbp2b/Rpl39/Ang |
| GO:0050000 | chromosome localization | 2,22E-03 | Ccnb1/Ndc80/Cenpe/Cenpf/Aurkb/Nuf2/Septin1/Gem/Kif22/Birc5/Fmn2/Psrc1/Spag5/Cdca8 |
| GO:0051303 | establishment of chromosome localization | 2,22E-03 | Ccnb1/Ndc80/Cenpe/Cenpf/Aurkb/Nuf2/Septin1/Gem/Kif22/Birc5/Fmn2/Psrc1/Spag5/Cdca8 |
| GO:0019882 | antigen processing and presentation | 2,43E-03 | H2-Ob/Ighm/H2-DMb2/H2-DMb1/H2-Q7/Fcgr3/H2-Q6/Cd74/H2-Eb1/Marchf1/Fcgr2b/H2-Aa/Wdfy4/H2-DMa/H2-Ab1/Ctse/Ccr7/Fcgr1/Ctss/Clec4a2/Flt3/Slc11a1/Psmb8/Cd68/Cd1d1/Psmb9/Ifi30/B2m/Icam1/H2-T23/Azgp1/Hfe/H2-T10/Unc93b1/Trem2/Thbs1/Rab32/Fcer1g/Tap1/H2-M3/Was/H2-T24 |
| GO:0002573 | myeloid leukocyte differentiation | 2,48E-03 | Spib/Cd4/Batf3/Adipoq/Ccr1/Fos/Ikzf1/Cd74/Camk4/Spi1/Irf4/Lrrc17/Itgam/Ccr7/Pirb/Trf/C1qc/Car2/Hcls1/Tnfsf9/Cebpb/Bmp4/Itgb3/Rassf2/Pparg/Ubash3b/Myc/Lif/Snx10/Nrros/Il3ra/Kitl/Pf4/Tyrobp/Runx1/Apcs/Ccl5/Trib1/Gpr183/Gpc3/Anxa2/Csf1r/Inpp5d/Pir/Trem2/Tgfbr2/Batf/Jun/Dlk1/Csf1/Fcer1g |
| GO:0030099 | myeloid cell differentiation | 2,53E-03 | Stap1/Hoxa5/Spib/Cd4/Batf3/Adipoq/Ccr1/Fos/Ikzf1/Cd74/Camk4/Spi1/Irf4/Lrrc17/Myb/Itgam/Ccr7/Pirb/Adgrf4/Slc4a1/Trf/Lep/Nckap1l/C1qc/Car2/Rab7b/Klf1/Mfhas1/Hcls1/Irf8/Zfp36/Tnfsf9/Cebpb/Bmp4/Klf2/P4htm/Itgb3/Rassf2/Pparg/Ubash3b/Pip4k2a/Zfp385a/Myc/Lif/Snx10/Thpo/Nrros/B2m/Il3ra/Kitl/Pf4/Heph/Tyrobp/Runx1/Apcs/Ccl5/Trib1/Gpr183/Gpc3/Anxa2/Klf13/Csf1r/Inpp5d/Dab2/Pir/Zbtb16/Trem2/Lmo2/Tgfbr2/Plscr1/Batf/Stat1/Jun/Csf3r/Dlk1/Csf1/Mturn/Fcer1g/Fas |
| GO:0032102 | negative regulation of response to external stimulus | 2,98E-03 | Stap1/Pbk/Siglecg/Vsig4/Adipoq/Sema3d/Ccr1/Cxcl13/Spn/Fcgr2b/Ccn3/Dpp4/Irak3/Il2ra/Sema3e/Dbn1/Ptgis/Lep/Smpdl3b/Nlrc5/Sema3a/Nbl1/Nmi/Mfhas1/Wnt5a/Serping1/Ptprs/Wnt3/Wfdc1/Havcr2/Zfp36/Cd34/Mill2/Dusp1/Sema6c/Adora2a/Cd200r1/Arg2/Tspan6/Tspan8/Pparg/Ubash3b/C1qtnf1/Serpine1/Aoah/Syt11/Tnr/Abcd2/Cd44/Cacnb3/Adcy7/H2-T23/Htra1/Trib1/Rtn4r/Nrxn1/Grin1/Siglece/Anxa2/Tfpi/Tnfrsf1b/Adrb2/Nr1d2/Isl1/Thbs1/Abcc8/Pros1/Parp14/Tap1/Sirpa/H2-M3/Gpx1/Sema3c/Arg1/Sema3b/Cd109/Wnt4/Robo1/Nt5e/Pdgfra/Thbd/Bbs4/Fndc4/Gpatch3/Lpcat3/Cd200/Arrb2/Clec2d/C1qbp/Gfer/Apoe/Dusp10/Foxf1 |
| GO:0008608 | attachment of spindle microtubules to kinetochore | 3,10E-03 | Ccnb1/Ndc80/Cenpe/Knl1/Nek2/Aurkb/Nuf2/Knstrn/Ect2/Racgap1/Spag5 |
| GO:0000819 | sister chromatid segregation | 3,23E-03 | Esco2/Ccnb1/Ndc80/Cenpe/Nek2/Aurkb/Nuf2/Knstrn/Bub1b/Ube2c/Ncaph/Haspin/Top2a/Cdc6/Cdc20/Nusap1/Kif4/Kif22/Birc5/Racgap1/Smc4/Kif23/Psrc1/Mad2l1bp/Ncapg/Spag5/Dsn1/Cdca8/Espl1/Axin2/Ncapd2/Slf1/Mapk15/Prc1/Hdac8/Tacc3/Rrs1/Stag2/Cdt1 |
| GO:0002712 | regulation of B cell mediated immunity | 3,23E-03 | Ighg2b/Fcer2a/Cd40/Btk/Fcgr3/Ptprc/Fcgr2b/C3/Il27ra/Fcgr1/Cd28/Cd55/Parp3/H2-T23 |
| GO:0002889 | regulation of immunoglobulin mediated immune response | 3,23E-03 | Ighg2b/Fcer2a/Cd40/Btk/Fcgr3/Ptprc/Fcgr2b/C3/Il27ra/Fcgr1/Cd28/Cd55/Parp3/H2-T23 |
| GO:0050852 | T cell receptor signaling pathway | 3,31E-03 | Ptpn22/Itk/Fyb2/Card11/Lck/Spn/Ptprc/Cd3e/Ccr7/Lgals3/Cd28/Vtcn1/Carmil2/Thy1/Cd300a/Themis2/Lcp2/Nfkbid/Tnfrsf21/Fyb/Cacnb3/Skap1/Tec/Icosl |
| GO:0042130 | negative regulation of T cell proliferation | 3,47E-03 | Slfn1/Vsig4/Btla/Lgals7/Spn/H2-Aa/H2-Ab1/Il2ra/Ripor2/Cd86/Vtcn1/Vsir/Havcr2/Cebpb/Bmp4/Cd274/Arg2/Tnfrsf21/Cd44/Erbb2/Pla2g2f/H2-M3/Pla2g2d/Arg1 |
| GO:0045639 | positive regulation of myeloid cell differentiation | 3,48E-03 | Hoxa5/Cd4/Ccr1/Fos/Ikzf1/Cd74/Itgam/Lep/Nckap1l/Car2/Rab7b/Hcls1/Itgb3/Lif/Thpo/Kitl/Pf4/Tyrobp/Runx1/Ccl5/Trib1/Csf1r/Inpp5d/Trem2/Stat1/Jun/Dlk1/Csf1/Mturn/Hmgb2/Ror2/Foxo3 |
| GO:0002708 | positive regulation of lymphocyte mediated immunity | 3,66E-03 | Ighg2b/Fcer2a/Sash3/Cd40/Btk/H2-Q7/Fcgr3/Rasgrp1/H2-Q6/Ptprc/C3/Il18/Fcgr1/Cd28/Cd1d1/B2m/Hspa8/H2-T23/Azgp1/H2-T10/Paxip1/Cadm1/Fcer1g/H2-M3/P2rx7/Vav1/H2-T24/Hmces/Tnfsf13/Il18r1/Lag3 |
| GO:1902850 | microtubule cytoskeleton organization involved in mitosis | 4,19E-03 | Ccnb1/Ndc80/Cenpe/Kifc1/Nek2/Kif11/Aurkb/Sapcd2/Nuf2/Tpx2/Dlgap5/Hspa1a/Cenpa/Cdc20/Efhc1/Ripor2/Nusap1/Stmn1/Kif4/Birc5/Parp3/Racgap1/Gpsm2/Kif23/Psrc1/Dynlt1b/Map9/Pax6 |
| GO:0032729 | positive regulation of interferon-gamma production | 4,21E-03 | Ptpn22/Cd2/Sash3/H2-Q7/Rasgrp1/Cd3e/Il27ra/Il18/Flt3/Slc11a1/Tlr9/Wnt5a/Carmil2/Tlr7/Irf8/Havcr2/Tnfsf9/Cd1d1/Ccr2/Tlr4/Tlr8/Runx1 |
| GO:0032623 | interleukin-2 production | 4,23E-03 | Vsig4/Sash3/Card11/Ptprc/Cd3e/Irf4/Anxa1/Slc11a1/Cd28/Vtcn1/Carmil2/Havcr2/Zfp36/Cd34/Cd1d1/Cd83/Nav3/Ccr2/Runx1/Card9 |
| GO:0070302 | regulation of stress-activated protein kinase signaling cascade | 4,23E-03 | Scimp/Pbk/Ptpn22/Gadd45g/Sfrp2/Gadd45b/Rasgrp1/Foxm1/Syk/Fcgr2b/Per1/Ccr7/Fgd2/Trf/Lep/Tlr9/Mfhas1/Wnt5a/Mdfi/Xdh/Ncf1/Dusp1/Gadd45a/Sphk1/Rassf2/Ccn2/Traf1/Tlr4/Bmp2/Sfrp4/Tnxb/Tnik/Card9/Map4k1/Dab2/Vangl2/Mapk8ip2/Zeb2/Ankrd6/Fas/Hand2/Sirpa/Igf1r/Ptk2b/Dact1/Ror2/Cdc42se1/Stk3/Pak1/Tiam1/Map3k20/Eda2r/Tgfb2/Myd88/Edn1/Dusp10/Lyn |
| GO:0000070 | mitotic sister chromatid segregation | 4,23E-03 | Ccnb1/Ndc80/Cenpe/Nek2/Aurkb/Nuf2/Knstrn/Bub1b/Ube2c/Ncaph/Haspin/Cdc6/Cdc20/Nusap1/Kif4/Kif22/Birc5/Racgap1/Smc4/Kif23/Psrc1/Mad2l1bp/Ncapg/Spag5/Dsn1/Cdca8/Espl1 |
| GO:0002833 | positive regulation of response to biotic stimulus | 4,63E-03 | Coch/Scimp/Zbp1/Ifi209/Rasgrp1/Cd74/Ifi204/Ifi205/Ly86/Cd180/Aim2/Nlrc5/Tlr9/Wnt5a/Havcr2/Cd274/Mndal/Ifi211/Bmp6/Tlr4/Tlr8/Mmp2/H2-T23/Card9/Irf7/Plscr1/Ifi203/Cadm1/H2-M3/Vav1/Hmgb2/Arg1 |
| GO:0007052 | mitotic spindle organization | 4,68E-03 | Ccnb1/Ndc80/Cenpe/Kifc1/Nek2/Kif11/Aurkb/Nuf2/Tpx2/Dlgap5/Hspa1a/Cdc20/Efhc1/Ripor2/Stmn1/Kif4/Birc5/Parp3/Racgap1/Gpsm2/Kif23/Psrc1 |
| GO:0051310 | metaphase plate congression | 5,12E-03 | Ccnb1/Ndc80/Cenpe/Cenpf/Aurkb/Nuf2/Septin1/Gem/Kif22/Psrc1/Spag5/Cdca8 |
| GO:0032872 | regulation of stress-activated MAPK cascade | 5,14E-03 | Scimp/Pbk/Ptpn22/Gadd45g/Sfrp2/Gadd45b/Rasgrp1/Foxm1/Syk/Fcgr2b/Per1/Ccr7/Fgd2/Trf/Lep/Tlr9/Mfhas1/Wnt5a/Mdfi/Xdh/Ncf1/Dusp1/Gadd45a/Sphk1/Rassf2/Ccn2/Traf1/Tlr4/Bmp2/Sfrp4/Tnxb/Tnik/Card9/Map4k1/Dab2/Vangl2/Mapk8ip2/Zeb2/Ankrd6/Fas/Hand2/Sirpa/Igf1r/Ptk2b/Dact1/Ror2/Cdc42se1/Stk3/Pak1/Tiam1/Map3k20/Eda2r/Tgfb2/Myd88/Edn1/Dusp10 |
| GO:0002688 | regulation of leukocyte chemotaxis | 5,23E-03 | Stap1/Sell/Rac2/Ccr1/Cxcl13/Cd74/Pgf/Vegfd/C5ar1/Ccn3/Dpp4/Ccr7/Nckap1l/Ripor2/Rarres2/Pla2g7/Nbl1/Thbs4/Wnt5a/Dusp1/Ccr2/Gpsm3/Serpine1/Ccl5/Csf1r/Vegfc/Ccl27a/Thbs1/Bst1/Csf1/C3ar1 |
| GO:0001909 | leukocyte mediated cytotoxicity | 5,47E-03 | Stap1/Il7r/H2-Q7/Fcgr3/Dnase1l3/Rasgrp1/H2-Q6/Coro1a/Ptprc/Itgam/Il18/Stx11/Fcgr1/Lep/Havcr2/Cd1d1/Ncf1/B2m/Hspa8/H2-T23/Tyrobp/Azgp1/Raet1e/H2-T10/Emp2/Cadm1/Tap1/H2-M3/P2rx7/Vav1/Muc4/Arg1/H2-T24/Lag3/Arrb2/Clec2d/Gfer/Stx7/Unc13d |
| GO:0001959 | regulation of cytokine-mediated signaling pathway | 5,65E-03 | Stap1/Zbp1/Adipoq/Casp4/Cd74/Ptprc/Syk/Irak3/Axl/Nlrc5/Wnt5a/Traip/Casp1/Sphk1/Pparg/Ecm1/Trem2/Cxcr4/Irf7/Cav1/Csf1/Parp14 |
| GO:0001912 | positive regulation of leukocyte mediated cytotoxicity | 5,68E-03 | Stap1/H2-Q7/Rasgrp1/H2-Q6/Ptprc/Itgam/Cd1d1/B2m/Hspa8/H2-T23/Tyrobp/Azgp1/H2-T10/Cadm1/H2-M3/P2rx7/Vav1/Arg1/H2-T24 |
| GO:0002548 | monocyte chemotaxis | 5,95E-03 | Ccl8/Ccl6/Ccr1/Ccl9/Ptpro/Anxa1/Ccn3/Lgals3/Ccl11/Pla2g7/Nbl1/Ccl24/Dusp1/Ccr2/Serpine1/Ccl5 |
| GO:0051315 | attachment of mitotic spindle microtubules to kinetochore | 6,07E-03 | Ndc80/Cenpe/Aurkb/Nuf2 |
| GO:0035456 | response to interferon-beta | 6,45E-03 | Ifi209/Ifi206/Ifi207/Ifi204/Ifi205/Igtp/Ifi47/Ifi213/Aim2/Gbp2/Iigp1/Mndal/Ifi211/Ifit1/Xaf1/9930111J21Rik1/Ifitm1/Tgtp2/Plscr1/Stat1/Ifi203 |
| GO:0002637 | regulation of immunoglobulin production | 6,48E-03 | Siglecg/Mzb1/Cd22/Ighm/Sash3/Cd40/Il33/Ptprc/Cd37/Fcgr2b/Il27ra/Cd28/Tlr9/Parp3/H2-T23 |
| GO:0071674 | mononuclear cell migration | 6,48E-03 | Ccl8/Ccl6/Ccr1/Ccl9/Ptpro/Jaml/Anxa1/C5ar1/Ccn3/Ccr7/Lgals3/Ccl11/Rarres2/Pla2g7/Nbl1/Ccl24/Dusp1/Ccr2/Serpine1/Ccl5/Csf1r/Ccl27a/Thbs1/Csf1/Sirpa/C3ar1 |
| GO:0030198 | extracellular matrix organization | 6,48E-03 | Cma1/Adamts19/Sfrp2/Wt1/Col28a1/Angptl7/Adamtsl2/Col8a2/Loxl4/Adamts3/Ndnf/Dpp4/Fn1/Antxr1/Sulf1/Fbln1/Dpt/Tgfbi/Ctss/Lum/Lgals3/Vit/Pdpn/Aebp1/Loxl1/Adamtsl1/Col14a1/Tnfrsf11b/Reck/Carmil2/Mmp23/Spock2/Sulf2/Ccdc80/Col4a6/Olfml2b/Ccn1/Itgb3/Ccn2/Adamts14/Mmp19/Rgcc/Mmp17/Adamtsl5/Ntng2/Ddr2/Tnr/Mmp2/Tnxb/Lmx1b/B4galt1/Anxa2/Mmp3/Smpd3/Postn/Pmp22/Nid1/Tnfrsf1b/Adamtsl4/Sh3pxd2b/Mmp9/Cav1/Apbb1/Abi3bp/Adamts5/Kif9/Adamts6/Cyp1b1 |
| GO:0043062 | extracellular structure organization | 6,48E-03 | Cma1/Adamts19/Sfrp2/Wt1/Col28a1/Angptl7/Adamtsl2/Col8a2/Loxl4/Adamts3/Ndnf/Dpp4/Fn1/Antxr1/Sulf1/Fbln1/Dpt/Tgfbi/Ctss/Lum/Lgals3/Vit/Pdpn/Aebp1/Loxl1/Adamtsl1/Col14a1/Tnfrsf11b/Reck/Carmil2/Mmp23/Spock2/Sulf2/Ccdc80/Col4a6/Olfml2b/Ccn1/Itgb3/Ccn2/Adamts14/Mmp19/Rgcc/Mmp17/Adamtsl5/Ntng2/Ddr2/Tnr/Mmp2/Tnxb/Lmx1b/B4galt1/Anxa2/Mmp3/Smpd3/Postn/Pmp22/Nid1/Tnfrsf1b/Adamtsl4/Sh3pxd2b/Mmp9/Cav1/Apbb1/Abi3bp/Adamts5/Kif9/Adamts6/Cyp1b1 |
| GO:0045058 | T cell selection | 6,68E-03 | Cd4/Dock2/Card11/Spn/Cd74/Ly9/Ptprc/Cd3e/Syk/Irf4/H2-DMa/Ccr7/Cd28/Cd1d1 |
| GO:0035710 | CD4-positive, alpha-beta T cell activation | 6,69E-03 | Gadd45g/Sash3/Lgals7/Spn/Ly9/Irf4/Anxa1/Runx3/Myb/Il2rg/Il18/Ccr7/Nckap1l/Vsir/Lilrb4a/Satb1/Cebpb/Cd83/Cd274/Ccr2/Arg2/Nfkbid/Cd44/Runx1/Gpr183/Rorc/Tgfbr2/Batf |
| GO:0002888 | positive regulation of myeloid leukocyte mediated immunity | 6,75E-03 | Stap1/Ighg2b/Btk/Fcgr3/Syk/C3/Itgam/Fcgr1 |
| GO:0009611 | response to wounding | 6,77E-03 | Siglecg/Alox15/Scnn1g/Tmeff2/Blk/Bnc1/Slc1a2/Tspan32/F5/Tnc/Mmrn1/Jaml/Syk/Fgf1/C3/Ndnf/Reg3a/Anxa1/Cldn1/Hpse/Fn1/P2ry12/Grin2c/Fbln1/Slc4a1/Mustn1/Pla2g4a/Adrb1/Pdpn/Bex1/Axl/Slc11a1/Plek/Dcbld2/Prr33/Wnt5a/Carmil2/Serping1/Ptprs/Lilrb4a/Sulf2/Wfdc1/Zfp36/Cd34/Ccn1/Ccr2/Itgb3/Tspan8/Ubash3b/C1qtnf1/Duox1/Serpine1/F13a1/Actg1/Clec10a/Syt11/Tnr/Cfh/Anxa8/Cd44/Tlr4/Pf4/Timp1/Mmp2/Tyrobp/Evl/Hmox1/B4galt1/Rtn4r/Myof/Ppl/Anxa2/Tec/Erbb2/Tfpi/Dpysl3/Flrt3/Adrb2/Cxcr4/Vangl2/Thbs1/Cav1/Jun/Abcc8/Pros1/Gli1/Cnn2/Fermt3/Fcer1g/5430416N02Rik/Adm/Igf1r/Lcp1/Myh10/Gpx1/Plpp3/2810403D21Rik/Lrp1/Scnn1b |
| GO:1900744 | regulation of p38MAPK cascade | 6,77E-03 | Ptpn22/Gadd45g/Gadd45b/Per1/Lep/Mfhas1/Xdh/Ncf1/Dusp1/Gadd45a/Sphk1/Bmp2 |
| GO:0032732 | positive regulation of interleukin-1 production | 7,02E-03 | Casp4/Ifi204/Ifi205/Ccr7/Egr1/Aim2/Il16/Panx1/Wnt5a/Gsdmd/Havcr2/Casp1/Mndal/Ifi211/Tlr4/Tlr8/Tyrobp/Panx2/Ccr5/Isl1/P2rx7/Hk1/Trim16/Stat3 |
| GO:0031341 | regulation of cell killing | 7,49E-03 | Stap1/Fcer2a/Cd5l/Il7r/H2-Q7/Dnase1l3/Rasgrp1/H2-Q6/Ptprc/Syk/Itgam/Lep/Cd55/Havcr2/Cd1d1/Cfh/B2m/Hspa8/H2-T23/Tyrobp/Azgp1/H2-T10/Ccr5/Cadm1/Tap1/H2-M3/P2rx7/Vav1/Muc4/Arg1/H2-T24/Bcl2l11/Lag3/Arrb2/Clec2d/Gfer/Stx7 |
| GO:0001776 | leukocyte homeostasis | 7,49E-03 | Gpr174/Ccnb2/Chst3/Cd74/Coro1a/Fcgr2b/Anxa1/Itgam/Il2ra/Il18/Pirb/Nckap1l/Flt3/Axl/Dock10/Lilrb4a/Ccr2/Tnfrsf13b/Pik3cd/Cfh/Cd44/Kitl/Gpr183/Sh2b2/Tsc22d3/Tnfsf13b/Fcer1g/Fas/P2rx7/Spta1/Gpam/Bcl2l11/Slc37a4/Jak3/Tgfb2 |
| GO:0001910 | regulation of leukocyte mediated cytotoxicity | 7,75E-03 | Stap1/Il7r/H2-Q7/Dnase1l3/Rasgrp1/H2-Q6/Ptprc/Itgam/Lep/Havcr2/Cd1d1/B2m/Hspa8/H2-T23/Tyrobp/Azgp1/H2-T10/Cadm1/Tap1/H2-M3/P2rx7/Vav1/Muc4/Arg1/H2-T24/Lag3/Arrb2/Clec2d/Gfer/Stx7 |
| GO:0035458 | cellular response to interferon-beta | 7,88E-03 | Ifi209/Ifi206/Ifi207/Ifi204/Ifi205/Igtp/Ifi47/Ifi213/Aim2/Gbp2/Iigp1/Mndal/Ifi211/Ifit1/9930111J21Rik1/Tgtp2/Stat1/Ifi203 |
| GO:0045730 | respiratory burst | 8,13E-03 | Jchain/Rac2/Cybb/Ncf2/Ncf4/Slc11a1/Ncf1 |
| GO:0043032 | positive regulation of macrophage activation | 8,23E-03 | Stap1/Il1rl1/Il33/Pla2g4a/Wnt5a/Havcr2/Cd1d1/Lrrk2/Tlr4/Trem2/Thbs1 |
| GO:0032496 | response to lipopolysaccharide | 8,34E-03 | Stap1/Scimp/Ptpn22/Cxcl13/Slpi/Ncf2/Irak3/Ly86/Il18/Ccr7/Cd180/Axl/Cd84/Gbp2/Slc11a1/Vim/Cd86/Cd68/Irf8/Lilrb4a/Havcr2/Zfp36/Cebpb/Ptgfr/Cd274/Casp1/Ppbp/Serpine1/Bmp6/Mrc1/B2m/Tlr4/Pf4/Ccl5/Ggt5/Ptgir/Trib1/Cd6/Spon2/Tnfrsf1b/Plscr4/Trem2/Noct/Plscr1/Stat1/Rpl13a/Sirpa/P2rx7/Adm/Hmgb2/Abca1/Cnr1/Irgm2/Il12rb2/Plcg2/Nfkbia/Ncl/Myd88/Dusp10/Lyn/Cxcl16 |
| GO:0030335 | positive regulation of cell migration | 8,43E-03 | Retn/Sell/Sema3d/Rac2/Ccr1/Cd40/Cxcl13/Fam107a/Reln/Spn/Cd74/Coro1a/Ptprc/Pgf/Enpp2/Fgf1/Vegfd/Anxa1/Wnt11/Nox4/C5ar1/Cldn1/Il18/Myo1f/Fn1/Sema3e/Ccr7/Fgf9/P2ry12/Egr1/Igfbp5/Fbln1/Nckap1l/Lgals3/Ccl11/Ripor2/Pdpn/Rarres2/Pla2g7/Sema3a/Vsir/Thbs4/Wnt5a/Ccl24/Carmil2/Dock8/Thy1/Nr4a3/Bmp4/Cd274/Glipr2/Sema6c/Ccn1/Sphk1/Ccr2/Itgb3/Slc8a1/Ackr3/Gpsm3/Serpine1/Myc/Myo5a/Itgax/Actg1/Ddr2/Pik3cd/Kitl/Icam1/Bmp2/Mmp2/Prkd1/Ccl5/Hmox1/Prex1/Grin1/Akt3/Kif20b/Mmp3/Csf1r/Fbxo31/Postn/Dab2/Ccbe1/Mgat5/Trem2/Cxcr4/Vegfc/Anxa3/Tgfbr2/Ccl27a/Itga4/Fut4/Pax6/Mmp9/Thbs1/Cav1/Jun/Irs2/Rhob/Arf6/Gli1/Fermt3/Cyp1b1/Csf1/C3ar1/Rin2/Igf1r/Rapgef3/Ptprz1/Plpp3/Sema3c/Lrp1/Ptk2b/Sema3b/Adgra2/Ror2/Pdgfra/Carmil1/Fgf7/Sun2/Smo/Mmp14/Dmtn/Pak1/Atp8a1/Tiam1/Plcg2/Adra2a/Hdac4/Stat3/Syde1/Pdgfrb/Tgfb2/Tek/C1qbp/Edn1/Foxf1/Lyn/Gpnmb/Cxcl16/Acvr1/Myadm/Tmsb4x/Vtn/Sema4b/Mcu/Cpeb1/Lamb1/Stat5a/Lgmn/Pcsk5/Ptp4a1/Hspb1/Lpar1/Kit/Fgfr1/Camk2d |
| GO:0033674 | positive regulation of kinase activity | 8,43E-03 | Stap1/Cd19/Tlr1/Cd4/Gadd45g/Ighm/Adipoq/Cenpe/Gadd45b/Cd40/Clspn/Rasgrp1/Reln/Tpx2/Cd74/Ptprc/Syk/Fgf1/Igfbp6/Nox4/Dab1/Il18/Ccr7/Egr1/Fgd2/Epha5/Fcgr1/Trf/Lep/Cdc6/Flt3/Nek10/Gpr39/Axl/Slc11a1/Tlr9/Ephb3/Cd86/Wnt5a/Mdfi/Gprc5b/Mmd/Cdk5r1/Cd300a/Bmp4/Ncf1/Gadd45a/Ccn1/Ntrk1/Map3k6/Pik3cg/Lrrk2/Itgb3/Rassf2/Mst1r/Ect2/Rgcc/Gckr/Ddr2/Kitl/Ephb6/Tlr4/Epha8/Bmp2/Pik3r5/Cdc25b/Tnik/Pfkfb1/Psrc1/Ccdc88a/Nrxn1/Map4k1/Iqgap3/Csf1r/Erbb2/Vav3/Dab2/Tgfa/Adrb2/Als2/Axin2/Vegfc/Emp2/Vangl2/Tgfbr2/Thbs1/Slc27a1/Zeb2/Agap2/Tgfb3/Csf1/P2rx7/Igf1r/Ptk2b/Lrp8/Npm1/Dusp12/Acsl1/Ang/Ror2/Lpar2/Robo1/Pdgfra/Irgm2/Stk3/Pak1/Tiam1/Map3k20/Adra2a/Pdgfrb/Tgfb2/Epha7/Prnp/Tek/Arrb2 |
| GO:0048525 | negative regulation of viral process | 8,50E-03 | Slpi/Oas1a/Fbln1/Fam111a/Apobec3/Trim59/Trim14/Oasl2/Zfp36/Rnasel/Oas2/Apcs/Ccl5/Gsn/Ifitm1/Trim35/Plscr1/Stat1/Jun/Resf1 |
| GO:0045860 | positive regulation of protein kinase activity | 9,42E-03 | Stap1/Tlr1/Cd4/Gadd45g/Ighm/Adipoq/Cenpe/Gadd45b/Cd40/Clspn/Rasgrp1/Reln/Tpx2/Cd74/Ptprc/Syk/Fgf1/Igfbp6/Nox4/Dab1/Il18/Ccr7/Egr1/Fgd2/Fcgr1/Trf/Lep/Cdc6/Nek10/Gpr39/Slc11a1/Tlr9/Cd86/Wnt5a/Mdfi/Gprc5b/Mmd/Cdk5r1/Cd300a/Bmp4/Ncf1/Gadd45a/Ccn1/Map3k6/Pik3cg/Lrrk2/Itgb3/Rassf2/Mst1r/Ect2/Rgcc/Ddr2/Kitl/Tlr4/Bmp2/Pik3r5/Cdc25b/Tnik/Psrc1/Ccdc88a/Nrxn1/Map4k1/Iqgap3/Csf1r/Erbb2/Dab2/Tgfa/Adrb2/Als2/Axin2/Vegfc/Emp2/Vangl2/Tgfbr2/Thbs1/Slc27a1/Zeb2/Agap2/Tgfb3/Csf1 |
| GO:0048285 | organelle fission | 9,61E-03 | Sycp1/Tex15/Ccnb1/Ndc80/Cenpe/Kifc1/Mki67/Nek2/Gimap3/Kif11/Aurkb/Fignl1/Nuf2/Tpx2/Septin1/Rspo1/Knstrn/Bub1b/Ube2c/Ncaph/Haspin/Top2a/Hspa1a/Cdc6/Cdc20/Mybl1/Ripor2/Mx2/Cd28/Nusap1/Wnt5a/Cep85/Dnm1/Kif4/Ccne1/Bmp4/Ppp2r2b/Bnip3/Kif22/Sphk1/Dcn/Birc5/Fmn2/Rgcc/Lif/Racgap1/Mnd1/Smc4/Kif23/Cdc25b/Mx1/Psrc1/Mad2l1bp/Inf2/Kif20b/Smpd3/Ppargc1a/Ncapg/Spag5/Dsn1/Cdca8/Tgfa/Espl1/Esr1/Map9/Spire2/Rpl24/Spire1/Rad54l/Pex11a/Ncapd2/Suv39h2/Rad51/Ccne2/Slf1/Igf1r/Lpin1/Acox1/Mapk15/Cntd1/Ccdc8/Prc1/Wnt4/Ube2s/Tacc3/Rrs1/Stag2/Mybl2/Bcl2l11/Cdt1/Ndc1/Syde1/Pdgfrb |
| GO:0002183 | cytoplasmic translational initiation | 9,67E-03 | Mcts2/Eif2s3y |
| GO:0046638 | positive regulation of alpha-beta T cell differentiation | 9,85E-03 | Sash3/Ikzf1/Syk/Anxa1/Runx3/Myb/Il2rg/Il18/Ccr7/Nckap1l/Cd1d1/Cd83/Ccr2/Nfkbid/Runx1/Zbtb16/Tgfbr2 |
| GO:0045807 | positive regulation of endocytosis | 9,96E-03 | Stap1/Ighm/Lrp2/Syk/C3/Ccr7/Fcgr1/Trf/Nckap1l/Axl/Wnt5a/Dnm1/Clip3/Lrrk2/Pparg/Serpine1/B2m/Magi2/Sfrp4/Rab31/Gpc3/Anxa2/Hfe/Dab2/Ppp3cc/Trem2/Cav1 |
| GO:0032715 | negative regulation of interleukin-6 production | 9,96E-03 | Ptpn22/Prg4/Bank1/Il27ra/Irak3/Nckap1l/Flt3/Cd84/Tlr9/Havcr2/Klf2/Cd200r1/Syt11/Tlr4/Aqp4/Inpp5d/Trim30a/Sirpa |
| GO:0002526 | acute inflammatory response | 1,09E-02 | Ighg2b/Saa3/Cd163/Btk/Fcgr3/Dnase1l3/Spn/Fcgr2b/C3/Mylk3/Reg3a/Fn1/Ccr7/Fcgr1/Reg3b/Hp/Alox5ap/Pik3cg/Pparg/Ephb6/Ffar3/Icam1/H2-T23/Ccl5/Cd6/B4galt1/Ccr5/Plscr1/Serpina3n/Fcer1g/Pla2g2d |
| GO:0071425 | hematopoietic stem cell proliferation | 1,13E-02 | Prg4/Sfrp2/Wnt10b/Wnt2b/Wnt5a/Cd34/Thpo/Kitl |
| GO:0038066 | p38MAPK cascade | 1,17E-02 | Ptpn22/Gadd45g/Gadd45b/Per1/Lep/Mfhas1/Xdh/Zfp36/Ncf1/Dusp1/Gadd45a/Sphk1/Bmp2 |
| GO:0060099 | regulation of phagocytosis, engulfment | 1,20E-02 | Stap1/Alox15/C3/Fcgr1/Nckap1l/Cd300a/Pparg/Rab31/Siglece/Trem2 |
| GO:0060759 | regulation of response to cytokine stimulus | 1,20E-02 | Stap1/Zbp1/Adipoq/Casp4/Cd74/Ptprc/Syk/Irak3/Axl/Nlrc5/Wnt5a/Traip/Casp1/Sphk1/Pparg/Tlr4/Ecm1/Trem2/Cxcr4/Irf7/Cav1/Csf1/Parp14 |
| GO:0002702 | positive regulation of production of molecular mediator of immune response | 1,20E-02 | Scimp/Ptpn22/Mzb1/Sash3/Cd40/Il33/Cd74/Ptprc/Cd37/Il18/Cd28/Tlr9/Wnt5a/Nr4a3/Gprc5b/B2m/Tlr4/Ffar3/H2-T23/Spon2/Paxip1/Fcer1g/H2-M3/Hmces/Hk1/Tnfsf13/Il18r1/Ffar2/Tek |
| GO:0042060 | wound healing | 1,20E-02 | Alox15/Scnn1g/Tmeff2/Blk/Bnc1/Tspan32/F5/Mmrn1/Jaml/Syk/Fgf1/C3/Ndnf/Reg3a/Anxa1/Cldn1/Hpse/Fn1/P2ry12/Fbln1/Slc4a1/Mustn1/Pla2g4a/Adrb1/Pdpn/Axl/Slc11a1/Plek/Dcbld2/Wnt5a/Carmil2/Serping1/Lilrb4a/Wfdc1/Cd34/Ccn1/Ccr2/Itgb3/Tspan8/Ubash3b/C1qtnf1/Duox1/Serpine1/F13a1/Actg1/Clec10a/Syt11/Cfh/Anxa8/Cd44/Tlr4/Pf4/Timp1/Evl/Hmox1/B4galt1/Myof/Ppl/Anxa2/Tec/Erbb2/Tfpi/Adrb2/Cxcr4/Vangl2/Thbs1/Cav1/Abcc8/Pros1/Cnn2/Fermt3/Fcer1g/Myh10/Gpx1/Plpp3/Scnn1b |
| GO:1990868 | response to chemokine | 1,20E-02 | Ccl8/Ccl6/Ccr1/Ccl9/Cxcl13/Ccl11/Ripor2/Ccl24/Dock8/Dusp1/Ackr3/Ppbp/Thpo/Pf4/Ccl5/Trem2/Cxcr4 |
| GO:1990869 | cellular response to chemokine | 1,20E-02 | Ccl8/Ccl6/Ccr1/Ccl9/Cxcl13/Ccl11/Ripor2/Ccl24/Dock8/Dusp1/Ackr3/Ppbp/Thpo/Pf4/Ccl5/Trem2/Cxcr4 |
| GO:1905517 | macrophage migration | 1,30E-02 | Stap1/Mcoln2/C5ar1/Ccr7/P2ry12/Lgals3/Rarres2/Ccr2/Cd200r1/Ccl5/Csf1r/Trem2/Thbs1/Rpl13a/Csf1/C3ar1/Ptk2b/Ror2/Mmp14/Cd200 |
| GO:0050792 | regulation of viral process | 1,30E-02 | Cd4/Slpi/Oas1a/Cd74/Trim5/Top2a/Fbln1/Fam111a/Apobec3/Trim59/Trim14/Oasl2/Trim30d/Zfp36/Rnasel/Oas2/Hspa8/Trim34a/Apcs/Trim12c/Ccl5/Gsn/Csf1r/Ifitm1/Trim35/Cxcr4/Plscr1/Trim30a/Stat1/Jun/Trim12a/Resf1/Kpna2 |
| GO:0010038 | response to metal ion | 1,30E-02 | Alox15/Fosb/Fibin/Fos/Lck/Cybrd1/Mt2/Mt1/Aldob/Kcnip3/Nptx1/Fn1/Trf/Pla2g4a/Gpr39/Syt2/Xdh/Pde1c/Ncf1/Bnip3/Crip1/Alox5ap/Lrrk2/Pparg/Cacna1g/Trpm2/Ect2/Hvcn1/Sod3/Syt11/Bmp6/B2m/Adcy7/Hmox1/Gsn/Nrxn1/Smpd3/Trpc1/Hfe/Pcdh15/Ncam1/Mmp9/Thbs1/Cav1/Syt9/Slc30a1/Jun/Kcnma1/Abcc8/Fas/P2rx7/Slc25a24 |
| GO:0046632 | alpha-beta T cell differentiation | 1,40E-02 | Gadd45g/Itk/Sash3/Ikzf1/Spn/Ly9/Syk/Irf4/Anxa1/Runx3/Myb/Il2rg/Il18/Ccr7/Nckap1l/Satb1/Cd1d1/Cd83/Ccr2/Nfkbid/Runx1/Gpr183/Rorc/Zbtb16/Tgfbr2/Batf/Pla2g2d/Tcf7 |
| GO:0071347 | cellular response to interleukin-1 | 1,41E-02 | Saa3/Zbp1/Ccl8/Ccl6/Cd40/Ccl9/Mylk3/Irak3/Fn1/Egr1/Rps6ka5/Ccl11/Ccl24/Cebpb/Serpine1/Ccl5/St18 |
| GO:0050730 | regulation of peptidyl-tyrosine phosphorylation | 1,45E-02 | Stap1/Ptpn22/Cd4/Sfrp2/Ighm/Adipoq/Bank1/Cd40/Lck/Samsn1/Reln/Cd74/Ptprc/Cd3e/Syk/Enpp2/Nox4/Errfi1/Il18/Efna5/Fcgr1/Lep/Thbs4/Hcls1/Thy1/Gprc5b/Cd300a/Ncf1/Itgb3/Lif/Bmp6/Cd44/Kitl/Tlr4/Icam1/Ccl5/Csf1r/Tec/Tgfa/Trem2/Vegfc/Isl1/Cav1/Itgb2/Bst1/Parp14/Ptprz1/Plpp3/Ptk2b/Lrp8 |
| GO:0050869 | negative regulation of B cell activation | 1,47E-02 | Bank1/Btla/Blk/Btk/Samsn1/Fcgr2b/Tbc1d10c/Flt3/Cd300a/Parp3/Tnfrsf13b/Tnfrsf21/Tyrobp/Inpp5d |
| GO:0002286 | T cell activation involved in immune response | 1,49E-02 | Pck1/Gadd45g/Spn/Cd74/Ly9/Irf4/Anxa1/Myb/Il18/Stx11/Ccr7/Lgals3/Apbb1ip/Slc11a1/Havcr2/Ccr2/Nfkbid/Icam1/Gpr183/Rorc/Batf/Itgal/Fcer1g/H2-M3/Lcp1 |
| GO:0001906 | cell killing | 1,53E-02 | Stap1/Fcer2a/Cd5l/Il7r/Cxcl13/H2-Q7/Fcgr3/Dnase1l3/Rasgrp1/H2-Q6/Coro1a/Ptprc/Syk/C3/Itgam/Il18/Stx11/Fcgr1/Lep/Lgals3/Ccl11/Cd55/Havcr2/Cd1d1/Ncf1/Cfh/B2m/Hspa8/Pf4/H2-T23/Tyrobp/Azgp1/Raet1e/H2-T10/Ccr5/Emp2/Ccl27a/Cadm1/Tap1/H2-M3/P2rx7/Vav1/Muc4/Arg1/H2-T24 |
| GO:0051607 | defense response to virus | 1,53E-02 | Ptpn22/Zbp1/Slfn8/Cd40/Il33/Tspan32/H2-Q7/Oas1a/Spn/Ptprc/Cd37/Irf5/Il2ra/Aim2/Apobec3/Oasl2/Tlr9/Cd86/Tlr7/Rnasel/Mill2/Bnip3/Tspan6/Ddit4/Itgax/Oas2/Ifit1/Slfn9/Trim34a/Tlr8/Ifit2/Htra1/Ccl5/Card9/Ifit3b/Spon2/Ifitm1/Unc93b1/Irf7/Plscr1/Trim30a/Stat1/Trim12a/Polr3h/Gbp2b/Il12rb1/Dtx3l/Serinc5/Bst2/Gpam |
| GO:0048535 | lymph node development | 1,53E-02 | Il7r/Cxcl13/Ikzf1/Ltb/Ccr7/Cxcr5/Cd248/Flt3/Pdpn |
| GO:0071459 | protein localization to chromosome, centromeric region | 1,56E-02 | Ndc80/Knl1/Aurkb/Bub1b/Haspin/Cenpa/Cdk1 |
| GO:0042116 | macrophage activation | 1,60E-02 | Stap1/Tlr1/Il1rl1/Il33/Syk/C5ar1/Itgam/Pla2g4a/Cd84/Slc11a1/Tlr9/Mfhas1/Wnt5a/Tlr7/Havcr2/Cd1d1/Casp1/Sphk1/Lrrk2/Pparg/Syt11/Tlr4/Tlr8/Tyrobp |
| GO:0032637 | interleukin-8 production | 1,61E-02 | Ptpn22/Cd2/Tlr1/Adipoq/Cd74/Ptprc/Syk/Anxa1/Il18/Lep/Tlr9/Wnt5a/Tlr7/Serpine1/Tlr5/Tlr4/Tlr8 |
| GO:0007080 | mitotic metaphase plate congression | 1,61E-02 | Ccnb1/Ndc80/Cenpe/Aurkb/Nuf2/Kif22/Psrc1/Cdca8 |
| GO:2000107 | negative regulation of leukocyte apoptotic process | 1,68E-02 | Il7r/Fcmr/Aurkb/Cd74/Fcgr2b/Il18/Ccr7/Axl/Hcls1/Dock8/Bmp4/Arg2/Cd44/Kitl/Ccl5/Tsc22d3/Rorc/Ccr5/Irs2/Fcer1g |
| GO:0019722 | calcium-mediated signaling | 1,79E-02 | Siglecg/Cd4/Cd22/Ccr1/Ackr4/Ptprc/Cd3e/Syk/Tbc1d10c/Cxcr6/Vcam1/Ccr7/P2ry12/Ackr2/Tmem100/Cxcr5/Calml4/Avpr1a/Ptgfr/Akap5/Sphk1/Ccr2/Lrrk2/Slc8a1/Lat2/Ackr3/Trpm2/Myo5a/Jph3/Xcr1/Gsto1/Fhl2/Ncam1/Ppp3cc/Trem2/Cxcr4/Ccr5/Itgal/Bst1/Adgrb2/Ptk2b/Cmya5/Irgm2/Dmtn/Ncald/Hdac4/Mapk7/Prnp/Itpr1/Edn1/L1cam |
| GO:0002763 | positive regulation of myeloid leukocyte differentiation | 1,82E-02 | Cd4/Ccr1/Fos/Ikzf1/Cd74/Itgam/Car2/Hcls1/Itgb3/Lif/Kitl/Pf4/Tyrobp/Runx1/Ccl5/Trib1/Csf1r/Trem2/Jun/Dlk1/Csf1 |
| GO:0051403 | stress-activated MAPK cascade | 1,82E-02 | Scimp/Pbk/Ptpn22/Gadd45g/Sfrp2/Gadd45b/Rasgrp1/Foxm1/Syk/Fcgr2b/Per1/Ccr7/Fgd2/Trf/Lep/Tlr9/Mfhas1/Wnt5a/Mdfi/Xdh/Zfp36/Ncf1/Dusp1/Gadd45a/Sphk1/Rassf2/Ccn2/Traf1/Tlr4/Bmp2/Sfrp4/Cryab/Tnxb/Tnik/Trib1/Card9/Map4k1/Fgf12/Dab2/Vangl2/Mapk8ip2/Zeb2/Ankrd6/Fas/Hand2/Sirpa/Igf1r/Ptk2b/Dact1/Ror2/Cdc42se1/Stk3/Pak1/Tiam1/Map3k20/Eda2r/Tgfb2/Myd88/Edn1/Dusp10 |
| GO:0043383 | negative T cell selection | 1,83E-02 | Dock2/Spn/Cd74/Ptprc/Cd3e/Ccr7/Cd28 |
| GO:0045060 | negative thymic T cell selection | 1,83E-02 | Dock2/Spn/Cd74/Ptprc/Cd3e/Ccr7/Cd28 |
| GO:0046641 | positive regulation of alpha-beta T cell proliferation | 1,83E-02 | Ptpn22/Ptprc/Cd3e/Syk/Rasal3/Il18/Cd28/Ccr2/H2-T23/Tgfbr2 |
| GO:0002714 | positive regulation of B cell mediated immunity | 1,86E-02 | Ighg2b/Fcer2a/Cd40/Btk/Fcgr3/Ptprc/C3/Fcgr1/Cd28 |
| GO:0002891 | positive regulation of immunoglobulin mediated immune response | 1,86E-02 | Ighg2b/Fcer2a/Cd40/Btk/Fcgr3/Ptprc/C3/Fcgr1/Cd28 |
| GO:0000280 | nuclear division | 1,94E-02 | Sycp1/Tex15/Ccnb1/Ndc80/Cenpe/Kifc1/Mki67/Nek2/Kif11/Aurkb/Fignl1/Nuf2/Tpx2/Septin1/Rspo1/Knstrn/Bub1b/Ube2c/Ncaph/Haspin/Top2a/Hspa1a/Cdc6/Cdc20/Mybl1/Ripor2/Cd28/Nusap1/Wnt5a/Cep85/Kif4/Ccne1/Bmp4/Kif22/Sphk1/Birc5/Fmn2/Rgcc/Lif/Racgap1/Mnd1/Smc4/Kif23/Cdc25b/Psrc1/Mad2l1bp/Kif20b/Smpd3/Ncapg/Spag5/Dsn1/Cdca8/Tgfa/Espl1/Esr1/Map9/Spire2/Rpl24/Spire1/Rad54l/Ncapd2/Suv39h2/Rad51/Ccne2/Slf1/Igf1r/Mapk15/Cntd1/Ccdc8/Prc1/Wnt4/Ube2s/Tacc3/Rrs1/Stag2/Mybl2/Bcl2l11/Cdt1/Ndc1/Syde1/Pdgfrb |
| GO:0032755 | positive regulation of interleukin-6 production | 1,94E-02 | Pou2af1/Scimp/Tlr1/Il33/Cd74/Syk/Vegfd/Pou2f2/Il16/Rab7b/Tlr9/Wnt5a/Tlr7/Tnfsf9/Il1rl2/Tlr4/Tlr8/Tyrobp/Card9/Spon2/Unc93b1/Ccr5/Isl1/Fcer1g/P2rx7 |
| GO:0007186 | G protein-coupled receptor signaling pathway | 1,97E-02 | Gpr174/P2ry10/Marco/Gpr1/Ccl8/Ppy/Ccl6/Rac2/Adgre4/Ccr1/Adrb3/Ccl9/Gpr132/Gabra3/Plcb2/Adra1b/Ackr4/Syk/Adgrd1/Mrap/Gpsm1/C3/Anxa1/Nmb/C5ar1/Cxcr6/Rgs14/Rgs2/Gna15/P2ry12/Adgrf4/Ffar1/Ackr2/Npy/Cckbr/Cxcr5/Ccl11/Adrb1/Gabrq/Gpr39/Car2/Sstr3/Fzd2/Vip/Adgrg2/Plek/Gpr65/Ccl24/Adgre1/Stmn1/Dnm1/Gabbr2/Avpr1a/Gprc5b/Cdk5r1/Ptgfr/S1pr4/Akap5/Ccr2/Adora2a/Pik3cg/Itgb3/Gpr34/Ackr3/Palm/Pth1r/Xcr1/Ffar3/Pf4/Gng2/Fzd9/Adcy7/Pik3r5/Gpr160/Kcnk2/Ccl5/Ptgir/Gpr183/Prex1/Gabrb3/Gng4/Npy1r/Rgs17/P2ry14/Dynlt1b/Adrb2/Gnb4/Cxcr4/Esr1/Ccr5/Acpp/Alox8/Galr2/Cav1/Cysltr1/Cish/Adgre5/Rgs10/C3ar1/Rgs11/Gal/Adgrb2/Vav1/Adm/Gpr162/Lrp1/Ptk2b/Tas1r1/Oxtr/Cacnb4/Abca1/Gabbr1/Cnr1/Glrb/Adgra2/Lpar2/Gpr161/Fzd1/Rgs16/Ucn3/Gkap1/Smo/Baiap3/Adra2a/Ramp1/Sctr/Bicd1/Ffar2/Arrb2/Arrdc3/Edn1/Gcgr/Apoe |
| GO:0045089 | positive regulation of innate immune response | 2,05E-02 | Coch/Zbp1/Ifi209/Rasgrp1/Cd74/Ifi204/Ifi205/Aim2/Nlrc5/Tlr9/Wnt5a/Havcr2/Mndal/Ifi211/Tlr4/Tlr8/Mmp2/H2-T23/Card9/Irf7/Plscr1/Ifi203/Cadm1/H2-M3/Vav1/Hmgb2 |
| GO:0010035 | response to inorganic substance | 2,05E-02 | Alox15/Pon1/Fosb/Fibin/Fos/Lck/Cybrd1/Mt2/Mt1/Aldob/Anxa1/Kcnip3/Nptx1/Fn1/Ccr7/Slc4a1/Trf/Reg3b/Hp/Pla2g4a/Rnf112/Gpr39/Syt2/Axl/Xdh/Pde1c/Avpr1a/Nr4a3/Ncf1/Bnip3/Crip1/Alox5ap/Sphk1/Lrrk2/Pparg/Cacna1g/Trpm2/Ect2/Hk3/Hvcn1/Sod3/Syt11/Bmp6/B2m/Adcy7/Ucp2/Mmp2/Cryab/Selenow/Lcn2/Hmox1/Gsn/Nrxn1/Smpd3/Trpc1/Hfe/Pcdh15/Ncam1/Mmp9/Thbs1/Cav1/Syt9/Slc30a1/Jun/Kcnma1/Rhob/Abcc8/Cyp1b1/Fas/Sirpa/Rad51/Lig1/P2rx7/Slc25a24/Gpx1 |
| GO:0019932 | second-messenger-mediated signaling | 2,05E-02 | Siglecg/Cd4/Cd22/Adgre4/Ccr1/Adrb3/Adra1b/Mt2/Mt1/Ackr4/Ptprc/Cd3e/Syk/Adgrd1/Mrap/Tbc1d10c/Ndnf/Cxcr6/Vcam1/Rgs2/Ccr7/P2ry12/Epha5/Ackr2/Tmem100/Prkar2b/Cxcr5/Adrb1/Vip/Adgrg2/Gpr65/Calml4/Adgre1/Avpr1a/Ptgfr/S1pr4/Akap5/Sphk1/Ccr2/Adora2a/Lrrk2/Slc8a1/Lat2/Ackr3/Rasd1/Trpm2/Myo5a/Pth1r/Jph3/Xcr1/Pf4/Adcy7/Ptgir/Gsto1/Fhl2/Ncam1/Npr1/Ppp3cc/Adrb2/Tcp11/Trem2/Cxcr4/Ccr5/Thbs1/Pde9a/Itgal/Bst1/Adgre5/Rasd2/Gal/Adgrb2/Adm/Pde7b/Ptk2b/Abca1/Prkar1b/Cnr1/Cmya5/Lpar2/Gpr161/Ucn3/Irgm2/Dmtn/Mafa/Ncald/Adra2a/Hdac4/Ramp1/Mapk7/Sctr/Prnp/Itpr1/Gucy1a2/Arrdc3/Edn1/Gcgr/Apoe/Ube2b/L1cam/Chga |
| GO:0072676 | lymphocyte migration | 2,15E-02 | Ccl8/Ccl6/Ccl9/Cxcl13/Myo1g/Spn/Itgb7/Il27ra/Ccr7/Ccl11/Ripor2/Wnt5a/Ccl24/Dock8/Ccr2/Itgb3/Cd200r1/Icam1/Stk10/Ccl5/Gpr183/Ecm1/Ccl27a/Itga4/Itgal/Cadm1 |
| GO:0006873 | cellular ion homeostasis | 2,26E-02 | Gpr174/P2ry10/Ms4a1/Cd19/Cd4/Cd52/Atp1a3/Ccr1/Cd40/Cxcl13/Lrp2/Atp6v0a4/Lck/Adra1b/Mt2/Mt1/Ackr4/Coro1a/Ptprc/Nmb/C5ar1/Cxcr6/Calb1/Ccr7/Gna15/Grin2c/Ffar1/Ackr2/Slc4a1/Lacc1/Trf/Atp1a2/Cckbr/Efhc1/Kcnk3/Cxcr5/Slc4a10/Adrb1/Gpr39/Scara5/Car2/Slc11a1/Fzd2/Trpv2/Gpr65/Tbxas1/Wnt5a/Kel/Mcub/Thy1/Avpr1a/Fam155a/Ptgfr/Bmp4/Bnip3/Akap5/Sgk1/Ccr2/Pik3cg/Lrrk2/Itgb3/Slc8a1/Ackr3/Ubash3b/Cacna1g/C1qtnf1/Trpm2/Myc/Myo5a/Hvcn1/Tmem28/Bmp6/Pth1r/Prkcb/Jph3/Xcr1/Cacnb3/Fzd9/Heph/Atp2c2/Lcn2/Slc4a4/Prkd1/Cd38/Ccl5/Ptgir/Gsto1/Hmox1/Grin1/Car7/Smpd3/Trpc1/Rab20/Npy1r/Hfe/Cacna1i/Adrb2/Cxcr4/Esr1/Ccr5/Galr2/Cav1/Slc30a1/Kcnma1/Cacna2d1/Scgn/Nptn/C3ar1/P2rx7/Adm/Lrp1/Ptk2b/Maip1 |
| GO:0002275 | myeloid cell activation involved in immune response | 2,26E-02 | Rac2/Dock2/Blk/Il33/Btk/Dnase1l3/Rasgrp1/Syk/Itgam/Myo1f/Stx11/Cd84/Havcr2/Nr4a3/Cd300a/Ccr2/Lat2/Tyrobp/Hmox1/Trem2/Anxa3/Itgb2/Fcer1g |
| GO:1905153 | regulation of membrane invagination | 2,32E-02 | Stap1/Alox15/C3/Fcgr1/Nckap1l/Cd300a/Pparg/Syt11/Rab31/Siglece/Trem2 |
| GO:0030003 | cellular cation homeostasis | 2,36E-02 | Gpr174/P2ry10/Ms4a1/Cd19/Cd4/Cd52/Atp1a3/Ccr1/Cd40/Cxcl13/Lrp2/Atp6v0a4/Lck/Adra1b/Mt2/Mt1/Ackr4/Coro1a/Ptprc/Nmb/C5ar1/Cxcr6/Calb1/Ccr7/Gna15/Grin2c/Ffar1/Ackr2/Slc4a1/Lacc1/Trf/Atp1a2/Cckbr/Efhc1/Kcnk3/Cxcr5/Slc4a10/Adrb1/Gpr39/Scara5/Car2/Slc11a1/Fzd2/Trpv2/Gpr65/Wnt5a/Kel/Mcub/Thy1/Avpr1a/Fam155a/Ptgfr/Bmp4/Bnip3/Akap5/Sgk1/Ccr2/Pik3cg/Lrrk2/Itgb3/Slc8a1/Ackr3/Ubash3b/Cacna1g/C1qtnf1/Trpm2/Myc/Myo5a/Hvcn1/Tmem28/Bmp6/Pth1r/Prkcb/Jph3/Xcr1/Cacnb3/Fzd9/Heph/Atp2c2/Lcn2/Slc4a4/Prkd1/Cd38/Ccl5/Ptgir/Gsto1/Hmox1/Grin1/Car7/Smpd3/Trpc1/Rab20/Npy1r/Hfe/Cacna1i/Adrb2/Cxcr4/Esr1/Ccr5/Galr2/Cav1/Slc30a1/Kcnma1/Cacna2d1/Scgn/Nptn/C3ar1/P2rx7/Adm/Lrp1/Ptk2b/Maip1 |
| GO:0002224 | toll-like receptor signaling pathway | 2,36E-02 | Scimp/Ptpn22/Tlr1/Cd40/Irf4/Tlr13/Irak3/Rab7b/Smpdl3b/Tlr9/Cd86/Mfhas1/Tlr7/Ptprs/Havcr2/Cd300a/Colec12/Tlr5/Tlr4/Tlr8/Unc93b1/Esr1/Irf7/Trim30a/Cav1/Arf6 |
| GO:0050777 | negative regulation of immune response | 2,44E-02 | Alox15/Vsig4/Il7r/Il1rl1/Ccr1/Il33/Samsn1/Spn/Ptprc/Fcgr2b/Anxa1/Il27ra/Irak3/Il2ra/Lgals3/Smpdl3b/Cd84/Nlrc5/Nmi/Vsir/Serping1/Havcr2/Cd300a/Ccr2/Arg2/Pparg/Parp3/H2-T23/Hmox1/Hfe/Inpp5d/Tgfb3/Parp14/Tap1/H2-M3/Gpx1/Muc4/Arg1 |
| GO:0030101 | natural killer cell activation | 2,44E-02 | Ptpn22/Slamf7/Il2rb/Ikzf1/Rasgrp1/Coro1a/Ptprc/Il18/Stx11/Lep/Axl/Havcr2/Elf4/H2-T23/Tyrobp |
| GO:0050920 | regulation of chemotaxis | 2,52E-02 | Stap1/Sell/Sema3d/Rac2/Ccr1/Cxcl13/Cd74/Pgf/Fgf1/Vegfd/C5ar1/Ccn3/Dpp4/Fn1/Sema3e/Ccr7/P2ry12/Nckap1l/Il16/Ripor2/Rarres2/Pla2g7/Sema3a/Nbl1/Thbs4/Wnt5a/Wnt3/Dusp1/Sema6c/Ccr2/Gpsm3/Serpine1/Prkd1/Ccl5/Tubb2b/Gpr183/Plxna4/Csf1r/Trem2/Cxcr4/Vegfc/Ccl27a/Thbs1/Bst1/Csf1/C3ar1/Sema3c/Ptk2b/Sema3b/Adgra2/Robo1/Scg2/Pdgfra |
| GO:1903900 | regulation of viral life cycle | 2,52E-02 | Cd4/Slpi/Oas1a/Cd74/Trim5/Top2a/Fam111a/Apobec3/Trim59/Trim14/Oasl2/Trim30d/Rnasel/Oas2/Hspa8/Trim34a/Apcs/Trim12c/Ccl5/Gsn/Ifitm1/Trim35/Plscr1/Trim30a/Trim12a/Resf1/Kpna2 |
| GO:0036037 | CD8-positive, alpha-beta T cell activation | 2,52E-02 | Ptpn22/Wdfy4/Runx3/Clec4a2/Nckap1l/Vsir/Lilrb4a/Satb1/Cd274/H2-T23/Runx1/Hfe |
| GO:0050855 | regulation of B cell receptor signaling pathway | 2,52E-02 | Stap1/Cd19/Cd22/Blk/Nfam1/Fcgr2b/Cd300a/Prkcb/Lpxn/Cmtm3 |
| GO:0002709 | regulation of T cell mediated immunity | 2,53E-02 | Il7r/Sash3/H2-Q7/H2-Q6/Spn/Ptprc/Dpp4/Il18/Vsir/Cd1d1/Ccr2/B2m/Hspa8/H2-T23/Azgp1/Hfe/H2-T10/Tnfrsf1b/H2-M3/P2rx7/Muc4/Was/Arg1/H2-T24 |
| GO:0050731 | positive regulation of peptidyl-tyrosine phosphorylation | 2,53E-02 | Stap1/Cd4/Ighm/Adipoq/Bank1/Cd40/Lck/Reln/Cd74/Ptprc/Cd3e/Syk/Enpp2/Nox4/Il18/Efna5/Fcgr1/Lep/Thbs4/Hcls1/Gprc5b/Ncf1/Itgb3/Lif/Bmp6/Cd44/Kitl/Tlr4/Icam1/Ccl5/Csf1r/Tec/Tgfa/Trem2/Vegfc/Isl1/Parp14/Ptprz1/Plpp3/Ptk2b/Lrp8 |
| GO:0043903 | regulation of symbiotic process | 2,63E-02 | Cd4/Slpi/Oas1a/Cd74/Trim5/Top2a/Fbln1/Fam111a/Apobec3/Trim59/Trim14/Oasl2/Trim30d/Zfp36/Rnasel/Oas2/Hspa8/Trim34a/Apcs/Trim12c/Ccl5/Gsn/Csf1r/Ifitm1/Trim35/Cxcr4/Ccr5/Plscr1/Trim30a/Stat1/Cav1/Jun/Trim12a/Resf1/Kpna2 |
| GO:0043372 | positive regulation of CD4-positive, alpha-beta T cell differentiation | 2,64E-02 | Sash3/Anxa1/Myb/Il2rg/Il18/Ccr7/Nckap1l/Cd83/Ccr2/Nfkbid |
| GO:0007051 | spindle organization | 2,64E-02 | Ccnb1/Ndc80/Cenpe/Kifc1/Nek2/Kif11/Aurkb/Nuf2/Tpx2/Dlgap5/Septin1/Knstrn/Rgs14/Hspa1a/Cdc20/Efhc1/Ripor2/Trim36/Stmn1/Kif4/Birc5/Parp3/Racgap1/Gpsm2/Kif23/Psrc1/Spag5/Espl1/Map9 |
| GO:0045059 | positive thymic T cell selection | 2,66E-02 | Dock2/Cd74/Ptprc/Cd3e/H2-DMa/Cd1d1 |
| GO:0070227 | lymphocyte apoptotic process | 2,66E-02 | Il7r/Blk/Fcmr/Aurkb/Cd74/Il2ra/Lgals3/Wnt5a/Dock8/Bmp4/Cd274/Siglec1/Arg2/Myc/Nfkbid/Tnfrsf21/Cd44/Ccl5/Tsc22d3/Rorc/Irs2/Fas/P2rx7/Gpam/Bcl2l11/Jak3/Tgfb2/Dffa/Lyn |
| GO:0002687 | positive regulation of leukocyte migration | 2,66E-02 | Sell/Rac2/Ccr1/Cxcl13/Spn/Cd74/Pgf/Vegfd/C5ar1/Ccr7/P2ry12/Nckap1l/Lgals3/Ripor2/Rarres2/Pla2g7/Thbs4/Wnt5a/Ccl24/Dock8/Thy1/Ccr2/Itgb3/Gpsm3/Serpine1/Kitl/Icam1/Ccl5/Csf1r/Trem2/Vegfc/Ccl27a/Itga4/Fut4/Mmp9/Thbs1/Csf1/C3ar1 |
| GO:0046718 | viral entry into host cell | 2,67E-02 | Cd4/Cd74/Trim5/Trim59/Axl/Trim14/Trim30d/Siglec1/Itgb3/Trim34a/Apcs/Trim12c/Gsn/Ifitm1/Trim30a/Cav1/Trim12a |
| GO:0072503 | cellular divalent inorganic cation homeostasis | 2,74E-02 | Gpr174/P2ry10/Ms4a1/Cd19/Cd4/Cd52/Ccr1/Cd40/Cxcl13/Lck/Adra1b/Mt2/Mt1/Ackr4/Coro1a/Ptprc/Nmb/C5ar1/Cxcr6/Calb1/Ccr7/Gna15/Grin2c/Ffar1/Ackr2/Atp1a2/Cckbr/Efhc1/Kcnk3/Cxcr5/Adrb1/Gpr39/Slc11a1/Fzd2/Trpv2/Gpr65/Wnt5a/Kel/Mcub/Thy1/Avpr1a/Fam155a/Ptgfr/Bmp4/Bnip3/Akap5/Ccr2/Pik3cg/Itgb3/Slc8a1/Ackr3/Ubash3b/Cacna1g/C1qtnf1/Trpm2/Myo5a/Tmem28/Pth1r/Prkcb/Jph3/Xcr1/Cacnb3/Fzd9/Atp2c2/Prkd1/Cd38/Ccl5/Ptgir/Gsto1/Grin1/Smpd3/Trpc1/Npy1r/Cacna1i/Adrb2/Cxcr4/Esr1/Ccr5/Galr2/Cav1/Slc30a1/Cacna2d1/Scgn/Nptn/C3ar1/P2rx7/Adm/Lrp1/Ptk2b/Maip1 |
| GO:0042088 | T-helper 1 type immune response | 2,76E-02 | Gadd45g/Il1rl1/Il33/Spn/Anxa1/Il27ra/H2-Ab1/Il18/Ccr7/Pla2g4a/Slc11a1/Havcr2/Ccr2 |
| GO:0051091 | positive regulation of DNA-binding transcription factor activity | 2,76E-02 | Cd40/Card11/Reln/Nfam1/Wnt10b/Trim5/Fcgr2b/Irak3/Il18/Epha5/Aim2/Rps6ka5/Rab7b/Cd84/Trim14/Fzd2/Tlr9/Hcls1/Wnt5a/Trim30d/Sphk1/Ntrk1/Rhebl1/Pparg/Traf1/Rgcc/Ddr2/Prkcb/Trim34a/Tlr4/Icam1/Ar/Trim12c/Prkd1/Cytl1/Card14/Ripk4/Ppargc1a/Arid5b/Esr1/Anxa3/Trim30a/Cav1/Itgb2/Trim12a/Plpp3/Lrp8/Npm1/Trim37/Tgfbr3/Lrrfip1/Fzd1/Smo/Stk3/Cd200/Il18r1/Eda2r/Hdac4/Stat3/Myd88/Edn1/Sting1 |
| GO:0140014 | mitotic nuclear division | 2,77E-02 | Ccnb1/Ndc80/Cenpe/Kifc1/Mki67/Nek2/Kif11/Aurkb/Nuf2/Tpx2/Knstrn/Bub1b/Ube2c/Ncaph/Haspin/Hspa1a/Cdc6/Cdc20/Ripor2/Cd28/Nusap1/Cep85/Kif4/Bmp4/Kif22/Sphk1/Birc5/Rgcc/Racgap1/Smc4/Kif23/Psrc1/Mad2l1bp/Kif20b/Smpd3/Ncapg/Spag5/Dsn1/Cdca8/Tgfa/Espl1/Esr1/Map9/Rpl24/Ncapd2/Slf1/Igf1r/Ccdc8/Prc1/Ube2s/Tacc3/Rrs1/Stag2/Mybl2/Cdt1 |
| GO:0061640 | cytoskeleton-dependent cytokinesis | 2,81E-02 | Cep55/Ckap2/Aurkb/Septin1/Cenpa/Efhc1/Septin3/Trim36/Nusap1/Kif20a/Stmn1/Kif4/Anln/Birc5/Ect2/Fmn2/Racgap1/Kif23/Kif20b/Map9/Spire2/Spire1/Rhob/Myh10/Prc1/Bbs4/Efhc2/Septin6/Rab35 |
| GO:0032731 | positive regulation of interleukin-1 beta production | 2,81E-02 | Casp4/Ifi204/Ifi205/Ccr7/Egr1/Aim2/Panx1/Wnt5a/Gsdmd/Casp1/Mndal/Ifi211/Tlr4/Tlr8/Tyrobp/Ccr5/Isl1/P2rx7/Hk1/Trim16/Stat3 |
| GO:0097012 | response to granulocyte macrophage colony-stimulating factor | 2,84E-02 | Cd4/Csf2ra/Pde1b/Zfp36 |
| GO:0050873 | brown fat cell differentiation | 2,98E-02 | Adipoq/Plac8/Adrb3/Scd1/Mrap/Rgs2/Lep/Adrb1/Fndc5/Rarres2/Ebf2/Cebpb/Bnip3/Pparg/Sh2b2/Ppargc1a/Slc2a4/Adrb2/Pex11a |
| GO:0031098 | stress-activated protein kinase signaling cascade | 3,02E-02 | Scimp/Pbk/Ptpn22/Gadd45g/Sfrp2/Gadd45b/Rasgrp1/Foxm1/Syk/Fcgr2b/Per1/Errfi1/Ccr7/Fgd2/Trf/Lep/Tlr9/Mfhas1/Wnt5a/Mdfi/Xdh/Zfp36/Ncf1/Dusp1/Gadd45a/Sphk1/Rassf2/Ccn2/Traf1/Tlr4/Bmp2/Sfrp4/Cryab/Tnxb/Tnik/Trib1/Card9/Map4k1/Fgf12/Dab2/Vangl2/Mapk8ip2/Zeb2/Ankrd6/Fas/Hand2/Sirpa/Igf1r/Ptk2b/Dact1/Ror2/Cdc42se1/Stk3/Pak1/Tiam1/Map3k20/Eda2r/Tgfb2/Myd88/Edn1/Dusp10/Lyn |
| GO:1903975 | regulation of glial cell migration | 3,02E-02 | Stap1/P2ry12/Efemp1/Vim/Ccr2/Gpr183/Grin1/Trem2/Csf1/Fas/Ptprz1/Lrp1 |
| GO:0043405 | regulation of MAP kinase activity | 3,15E-02 | Lax1/Ptpn22/Gadd45g/Sfrp2/Ighm/Adipoq/Gadd45b/Cd40/Rasgrp1/Cd74/Ptprc/Syk/Fgf1/Igfbp6/Nox4/Irak3/Rgs14/Rgs2/Dusp2/Ccr7/Fgd2/Trf/Nek10/Gpr39/Tlr9/Wnt5a/Mdfi/Cd300a/Bmp4/Dusp1/Gadd45a/Map3k6/Pik3cg/Lrrk2/Mst1r/Kitl/Uchl1/Tlr4/Bmp2/Pik3r5/Tnxb/Tnik/Trib1/Map4k1/Iqgap3/Erbb2/Dab2/Tgfa/Vangl2/Spred3/Thbs1/Cav1/Zeb2/Tgfb3/P2rx7/Igf1r |
| GO:0060627 | regulation of vesicle-mediated transport | 3,32E-02 | Stap1/Ighg2b/Alox15/Cd22/Ighm/Adipoq/Rac2/Dock2/Blk/Il2rb/Lrp2/Cd209b/Fcgr3/Septin1/Coro1a/Ptprc/Syk/Rspo1/Fcgr2b/C3/Anxa1/Pacsin1/Itgam/Il2rg/Ccr7/Fcgr1/Trf/Nckap1l/Lgals3/Rims3/Pla2g4a/Syt2/Axl/Cd84/Stxbp5l/Slc11a1/Wnt5a/Zdhhc2/Dnm1/Cdk5r1/Cd300a/Akap5/Sphk1/Ccr2/Clip3/Adora2a/Bmp2k/Lrrk2/Itgb3/Pparg/Sh3gl2/Cacna1g/Serpine1/Myo5a/Syt11/Hck/Prkcb/C2/B2m/Magi2/Hspa8/Snph/Sfrp4/Rab31/Hmox1/Nrxn1/Siglece/Gpc3/Anxa2/Smpd3/Doc2b/Hfe/Cacna1i/Ncam1/Dab2/Ppp3cc/Gas1/Slc2a4/Adrb2/Tcp11/Trem2/Cadps/Cav1/Syt9/Itgb2/Pros1/Cnn2/Arc/Ndrg4/Fcer1g/Sirpa/Cdh2/Ston1/Bsn/Stx1b/Lrp1 |
| GO:0036230 | granulocyte activation | 3,50E-02 | Dnase1l3/Syk/Itgam/Il18/Myo1f/Stx11/Il16/Cd300a/Ccr2/Tyrobp/Ccl5/Anxa3/Itgb2/Fcer1g |
| GO:0001960 | negative regulation of cytokine-mediated signaling pathway | 3,53E-02 | Stap1/Adipoq/Ptprc/Irak3/Nlrc5/Traip/Pparg/Ecm1/Cav1/Parp14/Arg1/Robo1 |
| GO:0072678 | T cell migration | 3,54E-02 | Cxcl13/Myo1g/Spn/Itgb7/Il27ra/Ccr7/Ripor2/Wnt5a/Dock8/Ccr2/Itgb3/Cd200r1/Icam1/Ccl5/Gpr183/Ecm1/Ccl27a/Itga4/Itgal |
| GO:0032956 | regulation of actin cytoskeleton organization | 3,59E-02 | Stap1/Alox15/Rhoh/Tmeff2/Rac2/Arhgdib/Fam107a/Coro1a/Mylk3/Wnt11/Nox4/Cotl1/Myo1f/Sema3e/Efna5/Epha5/Dbn1/Nckap1l/Ccl11/Plek/Gpr65/Hcls1/Rnd3/Ccl24/Carmil2/Stmn1/Sorbs3/Cdk5r1/S100a10/Itgb3/Plekhh2/Ccn2/Trpm2/Ect2/Fmn2/Rgcc/Actg1/Hck/Fhod3/Icam1/Fchsd2/Coro2b/Evl/Ccdc88a/Prex1/Gsn/Iqgap3/Csf1r/Tmsb10/Vangl2/Ccl27a/Spire2/Sh3pxd2b/Abi2/Spire1/Rhob/Arf6/Bst1/Tgfb3/Ppm1e/Rapgef3/Lrp1/Ptk2b/Was/Capg/Kank4/Spta1/Pdlim4/Wasf3/Wnt4/Add3/Bst2/Carmil1/Bbs4/Dmtn/Pak1 |
| GO:0000910 | cytokinesis | 3,59E-02 | Cep55/Ckap2/Aurkb/Septin1/Cdc6/Cenpa/Efhc1/Cxcr5/Septin3/Trim36/Nusap1/E2f7/Kif20a/Stmn1/Kif4/Anln/Birc5/Ect2/Fmn2/Racgap1/Plk3/Kif23/Cdc25b/Klhl13/Kif20b/Pstpip1/Map9/Spire2/Spire1/Rhob/Igf1r/Myh10/Prc1/Plk4/Bbs4/Efhc2/Septin6/Rab35/Plk5/Cdc14a/Stx2 |
| GO:0070555 | response to interleukin-1 | 3,69E-02 | Saa3/Zbp1/Ccl8/Ccl6/Cd40/Ccl9/Mylk3/Anxa1/Irak3/Fn1/Egr1/Rps6ka5/Ccl11/Ccl24/Cebpb/Serpine1/Ccl5/St18 |
| GO:0045637 | regulation of myeloid cell differentiation | 3,74E-02 | Hoxa5/Cd4/Adipoq/Ccr1/Fos/Ikzf1/Cd74/Spi1/Lrrc17/Itgam/Lep/Nckap1l/C1qc/Car2/Rab7b/Hcls1/Zfp36/Cebpb/P4htm/Itgb3/Rassf2/Ubash3b/Myc/Lif/Thpo/B2m/Kitl/Pf4/Tyrobp/Runx1/Apcs/Ccl5/Trib1/Klf13/Csf1r/Inpp5d/Zbtb16/Trem2/Lmo2/Stat1/Jun/Csf3r/Dlk1/Csf1/Mturn/Fas/Tob2/Hmgb2/Ptk2b |
| GO:0045061 | thymic T cell selection | 3,75E-02 | Dock2/Card11/Spn/Cd74/Ptprc/Cd3e/H2-DMa/Ccr7/Cd28/Cd1d1 |
| GO:0048645 | animal organ formation | 3,79E-02 | Gata5/Wt1/Lrp2/Wnt2b/Fgf1/Wnt11/Tbx1/Sulf1/Wnt5a/Gli2/Bmp4/Hoxa3/Bmp2/Ar/Axin2/Tgfbr2/Isl1/Hand2/Eya1 |
| GO:0046328 | regulation of JNK cascade | 3,85E-02 | Ptpn22/Gadd45g/Sfrp2/Gadd45b/Rasgrp1/Syk/Fcgr2b/Per1/Ccr7/Fgd2/Trf/Tlr9/Mfhas1/Wnt5a/Mdfi/Ncf1/Gadd45a/Rassf2/Ccn2/Traf1/Tlr4/Sfrp4/Tnxb/Tnik/Card9/Map4k1/Dab2/Vangl2/Mapk8ip2/Zeb2/Ankrd6/Sirpa/Igf1r/Ptk2b/Dact1/Ror2/Cdc42se1/Stk3/Pak1/Tiam1/Map3k20/Eda2r/Myd88/Edn1/Dusp10 |
| GO:1902622 | regulation of neutrophil migration | 3,86E-02 | Sell/Rhoh/Rac2/Cd74/C5ar1/Dpp4/Ccr7/Nckap1l/Ripor2/Thbs4 |
| GO:0002437 | inflammatory response to antigenic stimulus | 3,93E-02 | Ighg2b/Btk/Fcgr3/Rasgrp1/Spn/Fcgr2b/C3/Il2ra/Ccr7/Fcgr1/Cd28/Cd68/Lilrb4a/Ephb6/Icam1/H2-T23/Cd6/Cysltr1/Fcer1g/Pla2g2d/Gpx1/Hmgb2/Ahcy/Cnr1 |
| GO:0002920 | regulation of humoral immune response | 3,95E-02 | Fcer2a/Vsig4/Cd5l/Ptprc/Cd37/Fcgr2b/C3/H2-DMa/Ccr7/Cd55/Serping1/Cfh |
| GO:0032633 | interleukin-4 production | 4,04E-02 | Itk/Sash3/Il33/Cd3e/Syk/Irf4/Cd28/Vtcn1/Havcr2/Cebpb/Cd1d1/Cd83/H2-T23/Icosl |
| GO:0033077 | T cell differentiation in thymus | 4,09E-02 | Il7r/Dock2/Card11/Rasgrp1/Spn/Cd74/Ptprc/Cd3e/H2-DMa/Il2rg/Ccr7/Cd28/Cd1d1/Bmp4/Nfkbid/B2m/Mpzl2/Erbb2/Rorc/Zeb1 |
| GO:0043277 | apoptotic cell clearance | 4,09E-02 | Alox15/Marco/Rhoh/Rac2/C3/Anxa1/Axl/Ccr2/Itgb3/C2/Tyrobp/Trem2/Xkr4/Thbs1 |
| GO:0006874 | cellular calcium ion homeostasis | 4,11E-02 | Gpr174/P2ry10/Ms4a1/Cd19/Cd4/Cd52/Ccr1/Cd40/Cxcl13/Lck/Adra1b/Ackr4/Coro1a/Ptprc/Nmb/C5ar1/Cxcr6/Calb1/Ccr7/Gna15/Grin2c/Ffar1/Ackr2/Atp1a2/Cckbr/Efhc1/Kcnk3/Cxcr5/Adrb1/Gpr39/Fzd2/Trpv2/Gpr65/Wnt5a/Kel/Mcub/Thy1/Avpr1a/Fam155a/Ptgfr/Bmp4/Bnip3/Akap5/Ccr2/Pik3cg/Itgb3/Slc8a1/Ackr3/Ubash3b/Cacna1g/C1qtnf1/Trpm2/Myo5a/Tmem28/Pth1r/Prkcb/Jph3/Xcr1/Cacnb3/Fzd9/Atp2c2/Prkd1/Cd38/Ccl5/Ptgir/Gsto1/Grin1/Smpd3/Trpc1/Npy1r/Cacna1i/Adrb2/Cxcr4/Esr1/Ccr5/Galr2/Cav1/Slc30a1/Cacna2d1/Scgn/Nptn/C3ar1/P2rx7/Adm/Lrp1/Ptk2b/Maip1 |
| GO:0032677 | regulation of interleukin-8 production | 4,11E-02 | Ptpn22/Cd2/Tlr1/Adipoq/Cd74/Ptprc/Syk/Anxa1/Il18/Tlr9/Wnt5a/Tlr7/Serpine1/Tlr5/Tlr4/Tlr8 |
| GO:0070228 | regulation of lymphocyte apoptotic process | 4,14E-02 | Il7r/Blk/Fcmr/Aurkb/Cd74/Lgals3/Wnt5a/Dock8/Bmp4/Cd274/Siglec1/Arg2/Myc/Nfkbid/Cd44/Ccl5/Tsc22d3/Rorc/Irs2/P2rx7/Gpam/Jak3/Tgfb2/Lyn |
| GO:0070374 | positive regulation of ERK1 and ERK2 cascade | 4,23E-02 | Scimp/Ptpn22/Alox15/Cd4/Marco/Ccl8/Ccl6/Ccr1/Ccl9/Rasgrp1/Cd74/Ptprc/C3/Nox4/C5ar1/Ccl11/Mfhas1/Ccl24/Havcr2/Bmp4/Glipr2/Itgb3/Ccn2/Ackr3/Thpo/Cd44/Tlr4/Icam1/Bmp2/Ccl5/Gpr183/Nrxn1/Csf1r/Cavin3/Trem2/Esr1/Dnajc27/Jun/Ndrg4/Hand2/Lrp1/Ptk2b |
| GO:0032652 | regulation of interleukin-1 production | 4,23E-02 | Casp4/Ifi204/Ifi205/Anxa1/Errfi1/Ccr7/Egr1/Ffar1/Aim2/Il16/Panx1/Wnt5a/Gsdmd/Havcr2/Casp1/Sphk1/Mndal/Arg2/Ifi211/Tlr4/Tlr8/Aqp4/Tyrobp/Panx2/Trem2/Ccr5/Isl1/Sirpa/P2rx7 |
| GO:0070098 | chemokine-mediated signaling pathway | 4,26E-02 | Ccl8/Ccl6/Ccr1/Ccl9/Cxcl13/Ccl11/Ccl24/Ackr3/Ppbp/Thpo/Pf4/Ccl5/Trem2/Cxcr4 |
| GO:0071385 | cellular response to glucocorticoid stimulus | 4,34E-02 | Pck1/Fam107a/Anxa1/Zfp36/Bmp4/Sgk1/Ddit4 |
| GO:0002689 | negative regulation of leukocyte chemotaxis | 4,37E-02 | Stap1/Ccn3/Dpp4/Nbl1/Dusp1 |
| GO:0002474 | antigen processing and presentation of peptide antigen via MHC class I | 4,37E-02 | H2-Q7/Fcgr3/H2-Q6/Fcgr1/Clec4a2/Ifi30/B2m/H2-T23/Azgp1/Hfe/H2-T10/Fcer1g/Tap1/H2-M3/H2-T24 |
| GO:0045123 | cellular extravasation | 4,37E-02 | Sell/Chst4/Spn/Jaml/Itgb7/Itgam/Il27ra/Vcam1/Selplg/Lep/Ripor2/Thy1/Ccr2/Icam1/Ccl5/Itga4/Fut4/Itgal/Itgb2/Sirpa |
| GO:0048193 | Golgi vesicle transport | 4,39E-02 | Rab10/Sar1b/Sec31b/Ap1g2/Lamp1/Vti1b/Scyl1/Tbc1d20/Blzf1/Vps51/Rab1b/Copa/Scfd1/Trappc4/Ergic3/Rab8a/Gga2/Mia3/Wipi1/Pkdcc/Mia2/Trappc1/Dnm2/Arl1/Slc30a6/Rnf215/Tmed3/Vamp7/Rab6a/Trappc11/Trappc10/Kif1c/Arfgef2/Sec24c/Sec23a/Rab33b/Trappc2/Myo18a/Whamm/Copb1/Vps35l/Golga4/Pgap1/Prepl/Vps41/Arcn1/Sar1a/Rer1/Cux1/Cog8/Exoc1/Cog3/Snx1/Trappc6a/Tmed9/Plcb3/Snx12/Cnst/Yif1b/Atp9a/Dop1a/Rabif/Sec13/Kif16b/Copz2/Myo1b/Ykt6/Sort1/Vti1a/Krt18/Nsf/Lman2/Cog1/Arf4/Nbea/Optn/Llgl1/Ergic2/Arf1/Arfgap2/Rab13/Sec31a/Vcp/Cope/Uso1/Tmed6/Ergic1/Dnajc28/Cog7/Golph3l/Ier3ip1/Stx6/Arfgap3/Ap3d1/Copg1/Gosr2/Rab1a/Kdelr2/Htt/Bet1/Gbf1/Copz1/Stx5a/Tmed4/Gga3/Tmed10/Rab26/Sec22b/Kdelr3/Sec16a/Cog6/Sec16b/Preb/Creb3l2/Golga7/Lman1/Yipf5/Rint1/Hyou1/Lyplal1/Stxbp6/Sec23b/Nbas/Bnip1/Nkd2/Sec24d/Slc10a7/Rangrf/Rab6b |
| GO:0030100 | regulation of endocytosis | 4,46E-02 | Stap1/Alox15/Cd22/Ighm/Lrp2/Syk/Rspo1/C3/Pacsin1/Ccr7/Fcgr1/Trf/Nckap1l/Lgals3/Axl/Wnt5a/Dnm1/Cd300a/Akap5/Sphk1/Clip3/Bmp2k/Lrrk2/Itgb3/Pparg/Sh3gl2/Serpine1/Syt11/B2m/Magi2/Snph/Sfrp4/Rab31/Siglece/Gpc3/Anxa2/Hfe/Dab2/Ppp3cc/Slc2a4/Trem2 |
| GO:0002675 | positive regulation of acute inflammatory response | 4,47E-02 | Ighg2b/Btk/Fcgr3/C3/Ccr7/Fcgr1/Alox5ap/Pik3cg/Ffar3/H2-T23/Ccl5/Ccr5/Fcer1g |
| GO:0001773 | myeloid dendritic cell activation | 4,50E-02 | Batf3/Dock2/Tspan32/Cd37/Camk4/Spi1/Irf4/Pirb/Havcr2/Tnfsf9 |
| GO:0007632 | visual behavior | 4,54E-02 | Cacna1e/Atp1a3/Slc1a2/Lrrn4/Adra1b/Rgs14/Atp1a2/Pde1b/Ppp1r1b/Sgk1/Grin1/Ap1s2/Apbb1/Abcc8/Ndrg4 |
| GO:0032615 | interleukin-12 production | 4,54E-02 | Scimp/Cd40/Ltb/Syk/Irak3/Ccr7/Lep/Il16/Flt3/Tlr9/Irf8/Tnfsf9/Tlr4/Tlr8/Unc93b1/Isl1/Thbs1/H2-M3 |
| GO:0043368 | positive T cell selection | 4,55E-02 | Dock2/Spn/Cd74/Ly9/Ptprc/Cd3e/Irf4/H2-DMa/Cd1d1 |
| GO:0001914 | regulation of T cell mediated cytotoxicity | 4,59E-02 | Il7r/H2-Q7/H2-Q6/Ptprc/Cd1d1/B2m/Hspa8/H2-T23/Azgp1/H2-T10/H2-M3/P2rx7/Muc4/H2-T24 |
| GO:0031935 | regulation of chromatin silencing | 4,63E-02 | Cdc45/Uty |
| GO:0002791 | regulation of peptide secretion | 4,65E-02 | Siglecg/Cacna1e/Tlr1/Cd22/Glul/Blk/Cd40/Il33/Cd300c2/Cd74/Nnat/Syk/Anxa1/Ccn3/Cd209d/Efna5/Epha5/Ffar1/Lep/Cckbr/Per2/Adrb1/Gpr39/Panx1/Stxbp5l/Tlr9/Pde1c/Cd34/Casp1/Adora2a/Pparg/Trpm2/Birc5/Rgcc/Lif/Syt11/Tlr5/Krt20/Clec9a/Tlr4/Ffar3/Tlr8/Cpt1a/Slc16a1/Ucp2/H2-T23/Cd38/Ccl5/Pim3/Capn10/Nrxn1/Inhbb/Doc2b/Hfe/Bmp8a/Trem2/Vegfc/Isl1/Abcg1/Pax6/Apbb1/Syt9/Irs2/Arf6/Lepr/Abcc8/Kcnj11/Rfx6/Srgn/Cadm1/Tgfb3/Vpreb3/P2rx7/Myh10/Oxct1/Lrp1/Gabbr1/Cnr1/Ang/Gpam/Ucn3/Nlgn2/Dynll1/Camk2n1/Baiap3/Tiam1/Cd200/Adra2a/Tgfb2/Ffar2/Apoe/Cltrn/Lyn/Chga/Itsn1/Syt4/Adora1/Syt7/Slc30a8/Mcu/Rbp4/Ndufaf2/Hmgn3/Gpr68/Slc9b2/Foxp3 |
| GO:0055074 | calcium ion homeostasis | 4,71E-02 | Gpr174/P2ry10/Ms4a1/Cd19/Cd4/Cd52/Ccr1/Cd40/Cxcl13/Lck/Adra1b/Ackr4/Coro1a/Ptprc/Nmb/C5ar1/Cxcr6/Calb1/Ccr7/Gna15/Grin2c/Ffar1/Ackr2/Atp1a2/Cckbr/Efhc1/Kcnk3/Cxcr5/Adrb1/Gpr39/Fzd2/Trpv2/Gpr65/Wnt5a/Kel/Mcub/Thy1/Avpr1a/Fam155a/Ptgfr/Bmp4/Bnip3/Akap5/Ccr2/Pik3cg/Itgb3/Slc8a1/Ackr3/Ubash3b/Cacna1g/C1qtnf1/Trpm2/Myo5a/Snx10/Tmem28/Pth1r/Prkcb/Jph3/Xcr1/Cacnb3/Fzd9/Atp2c2/Prkd1/Cd38/Ccl5/Ptgir/Gsto1/Grin1/Smpd3/Reg1/Trpc1/Npy1r/Cacna1i/Adrb2/Cxcr4/Esr1/Ccr5/Galr2/Cav1/Slc30a1/Cacna2d1/Scgn/Nptn/C3ar1/P2rx7/Adm/Lrp1/Ptk2b/Maip1 |
| GO:0002698 | negative regulation of immune effector process | 4,81E-02 | Siglecg/Cd22/Vsig4/Il7r/Il33/Spn/Ptprc/Fcgr2b/Anxa1/Irak3/Il2ra/Lgals3/Cd84/Vsir/Serping1/Havcr2/Cd300a/Mill2/Ccr2/Tspan6/Parp3/H2-T23/Htra1/Hmox1/Hfe/Tgfb3/Vpreb3/Tap1/H2-M3/Muc4/Arg1/Gpatch3/Jak3/Tgfb2/Arrb2/Clec2d/C1qbp/Gfer/Dusp10/Foxf1 |
| GO:0045576 | mast cell activation | 4,81E-02 | Rhoh/Rac2/Blk/Btk/Fcgr3/Rasgrp1/Syk/Cd48/Ptpre/Cd84/Nr4a3/Cd300a/Lat2/Lcp2/Tlr4/Hmox1 |
| GO:0043508 | negative regulation of JUN kinase activity | 4,81E-02 | Ptpn22/Sfrp2 |
| GO:0032607 | interferon-alpha production | 4,81E-02 | Ptpn22/Flt3/Nmi/Tlr9/Tlr7/Ptprs/Havcr2/Tlr4/Tlr8/Irf7/Stat1 |
| GO:0002883 | regulation of hypersensitivity | 4,81E-02 | Ighg2b/Btk/Fcgr3/Spn/Fcgr2b/C3/Ccr7/Fcgr1 |
| GO:0051988 | regulation of attachment of spindle microtubules to kinetochore | 4,81E-02 | Ccnb1/Cenpe/Nek2/Aurkb/Knstrn/Ect2/Racgap1/Spag5 |
| GO:0007249 | I-kappaB kinase/NF-kappaB signaling | 4,85E-02 | Saa3/Cd4/Rhoh/Adipoq/Il1rl1/Cd40/Btk/Card11/Cd74/Trim5/Per1/Trim59/Trim14/Tlr9/Wnt5a/Tlr7/Trim30d/Gprc5b/Casp1/Traf1/Alpk1/Tifab/Nfkbid/Prkcb/Trim34a/Tlr4/Tlr8/Trim12c/Prkd1/Card9/Tifa/Irak1bp1/Esr1/S100a4/Trim30a/Stat1/Trim12a |
| GO:0016032 | viral process | 4,88E-02 | Zbp1/Cd4/Slpi/Oas1a/Cd74/Trim5/Vcam1/Top2a/Fbln1/Fam111a/Apobec3/Trim59/Axl/Trim14/Oasl2/Trim30d/Zfp36/Rnasel/Siglec1/Itgb3/Myc/Mst1r/Oas2/Hspa8/Trim34a/Icam1/Apcs/Trim12c/Ccl5/Gsn/Csf1r/Ifitm1/Trim35/Cxcr4/Ccr5/Plscr1/Trim30a/Stat1/Mmp9/Cav1/Jun/Trim12a/Resf1/Kpna2 |
| GO:0097011 | cellular response to granulocyte macrophage colony-stimulating factor stimulus | 4,89E-02 | Cd4/Csf2ra/Pde1b/Zfp36 |
