## Supplementary Table S2 for "EZH2 deletion does not impact acinar cell regeneration but restricts progression to pancreatic cancer in mice"

**Supplemental Table S2. Significant KEGG pathways enriched in *Mist1^creERT/+^KRAS^G12D^* *vs* WT for H3K4me3 and H3K27me3 ChIP-Sequencing**

| **H3K4me3** | | | |
| --- | --- | --- | --- |
| **ID** | **Description** | **P adjust** | **Gene ID** |
| mmu05168 | Herpes simplex virus 1 infection | 6,3426E-18 | Gm2026/Gm3055/Zfp984/Gm14308/Zfp850/Gm10778/Zfp729b/Zfp282/Zfp956/E430018J23Rik/AW146154/Zfp119a/Ticam1/Srsf9/Eif2b3/H2-Q6/H2-Q9/Srsf1/Zfp607b/Akt1/Akt2/Apaf1/Birc3/Birc2/B2m/Bad/Bak1/Bax/Bcl2/Bcl2l1/Bid/Calr/Casp3/Casp8/Casp9/Chuk/Socs3/Cycs/Daxx/Eif2s1/Eif2ak3/Eif2b4/Eif4ebp1/Fadd/Fas/Tlr3/Pdia3/H2-Ab1/H2-Bl/H2-D1/H2-Eb1/H2-K1/H2-M3/H2-DMa/H2-Q1/H2-Q10/H2-Q2/H2-Q4/H2-T22/H2-T23/H2-T24/Eif2ak1/Ifnar1/Ifnar2/Ifngr1/Ifngr2/Ikbkb/Irf9/Itga5/Jak1/Jak2/Zfp87/Zfp617/Nfkb1/Nfkbia/Pik3ca/Pik3cd/Pik3r1/Pik3r2/Pik3r3/Pml/Pou2f1/Pou2f2/Pou2f3/Ppp1cb/Ppp1cc/Eif2ak2/Ptpn11/Zfp286/Zfp184/Rela/Rheb/Ccl5/Srsf2/Srsf3/Srsf5/Sp100/Src/Srpk1/Zfp871/Stat1/Stat2/Eif2b1/Syk/Zfp658/Zfp719/Zfp180/Zfp677/Zfp947/Zfp748/Zfp273/Zfp583/Tap2/Tapbp/Zfp354a/Cgas/Zfp879/AU041133/Alyref/Eif2b2/Zfp455/Zfp595/Tnfrsf1a/Traf2/Traf3/Traf5/Traf6/Zfp7/Eif2b5/Zfp160/Zfp472/Zfp81/Zfp959/Srsf7/Zfp1/Zfp101/Zfp11/Zfp13/Zfp26/Zfp28/Zfp30/Zfp37/Zfp39/Zfp40/Zfp46/Zfp51/Zfp52/Zfp54/Zfp57/Zfp60/Zfp61/Zfp85/Zfp9/Zfp90/Zfp93/Zfp94/Zfp97/Zik1/Zim1/Mavs/Zfp334/Ddx58/Zfp189/Ifna13/Tnfrsf14/Zfp12/Zfp212/Zfp954/Zfp418/Zfp772/Zfp114/Zfp790/Zfp940/Zfp420/Zfp382/Zfp939/AI987944/Zfp768/Zfp764/Zfp958/Zfp930/Zfp868/Zfp961/Zfp612/Zfp426/Zfp809/Zfp599/Zfp810/Zfp65/Zfp825/Zfp709/Zfp938/Zfp454/Zfp867/Akt3/Zfp458/Zfp874a/Zfp58/Zfp647/Zfp641/Zfp760/Zfp994/Zfp799/Zfp870/Zfp952/Zfp563/Zfp119b/Rnasel/Tlr2/Zfp53/Zfp68/Zfp267/Ifna15/Zfp933/A430033K04Rik/Zfp128/Zfp324/Zfp568/Zfp14/Zfp473/Zfp791/Zfp317/Zfp937/Oas1a/Map3k7/Irak4/Zfp747/Eif2ak4/Zfp398/Zfp354b/Zfp354c/Zfp459/Zfp853/Zfp786/Zfp78/Zfp82/Zfp866/Card9/Zfp780b/Gm5141/Rsl1/Zfp948/Zfp229/Zfp69/Zfp667/Zfp456/Zfp874b/Zfp950/Zfp457/Zfp708/Zfp268/Zfp141/Zfp975/Zfp560/Zfp960/Nxf1/Irf7/Irf3/Zfp316/Zfp607a/Zfp108/Tyk2/Alyref2/Zfp386/Zfp113/Tbk1/Ikbke/Zfp235/Zfp111/Mtor/Zfp109/Srsf4/Zfp112/Nectin1/Zfp872/Zfp551/Zfp963/Zfp951/Gm14322/Gm14430/Zfp808/Zfp964/Tsc1/Gm14391/Tab1/Gm14326/Zfp991/H2-T-ps/Gm8909/Gm14434/Zfp869/Zfp442/Zfp605/Zfp788/Zfp169/Hcfc2/Srsf6/Tab2/Zfp707/Zfp746/Zfp688/Bst2/Zfp715/2810021J22Rik/Zfp619/Zfp597/Zfp689/Zfp626/Zfp935/Ifih1/Zfp251/Tradd/Zfp949/2610008E11Rik/Zfp157/Zfp661/Zfp558/Zfp777/Sting1/Zfp248/Zfp74/Zfp974/Zfp763/Zfp383/Zfp946/Zfp84/Pik3cb/Zfp773/Zfp266/Zfp712/Zfp623/9130019O22Rik |
| mmu05225 | Hepatocellular carcinoma | 3,8522E-14 | Gstt3/Ddb2/Braf/Raf1/Actb/Actg1/Akt1/Akt2/Apc/Axin1/Axin2/Bad/Bak1/Bax/Bcl2l1/Ctnnb1/Ccnd1/Cdk4/Cdk6/Cdkn1a/Cdkn2a/Gadd45a/Dvl1/Dvl2/Dvl3/E2f1/E2f3/Egfr/Frat1/Fzd1/Fzd3/Fzd4/Fzd5/Fzd6/Fzd7/Fzd8/Fzd9/Gab1/Grb2/Gsta3/Gsta4/Gstm1/Gstm2/Gstm3/Gstm4/Gstm5/Gstp2/Gstp1/Gstt1/Gstt2/Gsto1/Hmox1/Hras/Igf1r/Lef1/Lrp5/Lrp6/Smad2/Smad3/Smad4/Met/Myc/Gadd45b/Nfe2l2/Nqo1/Nras/Pik3ca/Pik3cd/Pik3r1/Pik3r2/Pik3r3/Prkcb/Prkcg/Plcg1/Pten/Rb1/Shc1/Shc3/Smarca4/Smarcb1/Smarcc1/Sos1/Sos2/Mgst2/Frat2/Tcf7/Tcf7l1/Tcf7l2/Wnt9a/Terc/Tert/Tgfa/Tgfb1/Tgfb2/Tgfb3/Tgfbr1/Tgfbr2/Wnt1/Wnt10b/Wnt11/Wnt9b/Wnt2/Wnt2b/Wnt4/Wnt5a/Wnt5b/Wnt6/Gstp3/Txnrd3/Plcg2/Akt3/Apc2/Gadd45g/Arid1b/E2f2/Map2k1/Map2k2/Mapk1/Mapk3/Brd7/Polk/Shc4/Peg12/Dpf1/Txnrd1/Keap1/Actl6a/Mgst1/Gsk3b/Mtor/Fzd2/Smarce1/Rps6kb2/Mgst3/Smarcd3/Smarca2/Gsto2/Gstm7/Dpf3/Phf10/Pik3cb/Smarcd2/Smarcd1/Csnk1a1/Arid1a |
| mmu04140 | Autophagy - animal | 8,809E-12 | Rubcn/Atg14/Prkaa1/Prkaa2/Raf1/Akt1/Akt2/Atg5/Bad/Bcl2/Bcl2l1/Bnip3/Rb1cc1/Cflar/Ctsb/Ctsd/Ctsl/Dapk2/Dapk3/Eif2s1/Eif2ak3/Hif1a/Hmgb1/Hras/Igf1r/Irs1/Irs3/Itpr1/Lamp1/Mtmr4/Rab8a/Mras/Nras/Sqstm1/Pdpk1/Pik3ca/Pik3cd/Pik3r1/Pik3r2/Pik3r3/Prkaca/Prkacb/Prkcd/Ppp2ca/Ppp2cb/Pten/Rab1a/Rab33b/Rab7/Rheb/Rras/Camkk2/Stk11/Tank/Atg9b/Zfyve1/Wdr41/Traf6/Ulk1/Vamp8/Pik3c3/Ambra1/Atg4d/Smcr8/Akt3/Atg4c/Atg9a/Map2k1/Map2k2/Map3k7/Mapk1/Mapk10/Mapk3/Mapk8/Mapk9/Eif2ak4/Ulk2/Atg2a/Irs2/Rragd/Wipi1/Rragc/Sh3glb1/Becn1/Tbk1/Gabarap/Mlst8/Mtor/Supt20/Rps6kb2/Nrbf2/Tsc1/Atg4b/Atg10/Rras2/Akt1s1/Stx17/Atg3/Atg101/Rraga/Trp53inp2/Dapk1/C9orf72/Atg16l2/Atg7/Mtmr3/Rptor/Ddit4/Pik3cb/Wipi2/Pik3r4/Atg16l1/Uvrag/Ern1/Gabarapl2/Mtmr14/Deptor |
| mmu04141 | Protein processing in endoplasmic reticulum | 8,809E-12 | Dnajc3/Rpn1/Txndc5/Ssr1/Edem2/Selenos/Sec13/Atxn3/Derl2/Atf4/Bag1/Bak1/Bax/Bcl2/Hyou1/Pdia4/Calr/Canx/Capn1/Capn2/Casp12/Atf6b/Cryaa/Cryab/Dnajc5/Ddost/Dnajc1/Eif2s1/Eif2ak3/Ube2j2/Sec63/Pdia3/Hspa5/Eif2ak1/Hspa1l/Dnaja1/Hsph1/Hspa1b/Hspa2/Hsp90ab1/Hsp90aa1/Stt3a/Man1a2/Ppp1r15a/Nfe2l2/Hspa4l/P4hb/Plaa/Prkcsh/Eif2ak2/Edem1/Rad23a/Rad23b/Rnf185/Rpn2/Sar1a/Sec23a/Sec61g/Sel1l/Bag2/Skp1/Ube2d1/Ckap4/Os9/Nploc4/Sec24c/Traf2/Ube2g2/Marchf6/Wfs1/Xbp1/Atf6/Man1b1/Man1c1/Amfr/Sec31b/Map2k7/Map3k5/Mapk10/Mapk8/Mapk9/Vcp/Cul1/Sec23b/Eif2ak4/Uggt1/Nsfl1c/Ero1a/Fbxo6/Prkn/Preb/Sec61a1/Rnf5/Ubqln1/Stub1/Rbx1/Dnaja2/Mbtps1/Ube2d2a/Dnajb12/Dnajb2/Mogs/Sec61a2/Ngly1/Ube4b/Herpud1/Hspbp1/Ssr2/Sar1b/Uggt2/Ubxn6/Dnajc10/Lman2/Ube2g1/Erp29/Ssr3/Ero1b/Derl1/Dnajb11/Stt3b/Sec31a/Sec62/Sec24d/Lman1/Derl3/Pdia6/Tram1/Syvn1/Svip/Sec24a/Ern1/Tusc3/Sil1/Rrbp1/Sec24b |
| mmu05220 | Chronic myeloid leukemia | 8,809E-12 | Ddb2/Braf/Raf1/Bcr/Abl1/Akt1/Akt2/Bad/Bak1/Bax/Bcl2l1/Runx1/Cbl/Ccnd1/Cdk4/Cdk6/Cdkn1a/Cdkn1b/Cdkn2a/Chuk/Crk/Crkl/Ctbp1/Ctbp2/Gadd45a/E2f1/E2f3/Mecom/Gab2/Grb2/Hdac2/Hras/Ikbkb/Smad3/Smad4/Mdm2/Myc/Gadd45b/Nfkb1/Nfkbia/Nras/Pik3ca/Pik3cd/Pik3r1/Pik3r2/Pik3r3/Ptpn11/Rb1/Rela/Shc1/Shc3/Sos1/Sos2/Stat5a/Stat5b/Tgfb1/Tgfb2/Tgfb3/Tgfbr1/Tgfbr2/Akt3/Gadd45g/E2f2/Map2k1/Map2k2/Mapk1/Mapk3/Polk/Shc4/Hdac1/Pik3cb |
| mmu04110 | Cell cycle | 3,4588E-11 | E2f4/Espl1/Cdc20/Abl1/Atm/Bub1/Bub1b/Bub3/Ccna2/Ccnb2/Ccnd1/Ccnd2/Ccnd3/Ccne1/Ccne2/Cdc25a/Cdc25b/Cdc25c/Cdk1/Cdc45/Cdc7/Cdk2/Cdk4/Cdk6/Cdk7/Cdkn1a/Cdkn1b/Cdkn2a/Cdkn2b/Cdkn2c/Cdkn2d/Chek1/Crebbp/Smc3/Gadd45a/E2f1/E2f3/E2f5/Hdac2/Mad1l1/Smad2/Smad3/Smad4/Mcm3/Mcm2/Mcm4/Mcm5/Mcm6/Anapc1/Mdm2/Myc/Gadd45b/Orc1/Orc2/Pcna/Plk1/Rad21/Rb1/Rbl1/Rbl2/Stag1/Tfdp2/Skp1/Tfdp1/Tgfb1/Tgfb2/Tgfb3/Cdc14b/Ttk/Wee1/Ywhae/Ywhag/Ywhah/Ywhaq/Ywhaz/Zbtb17/Cdc14a/Cdc6/Gadd45g/E2f2/Atr/Orc5/Ccnb1/Pkmyt1/Cul1/Dbf4/Skp2/Pttg1/Ep300/Hdac1/Anapc4/Cdc23/Ywhab/Sfn/Mad2l1/Anapc7/Fzr1/Rbx1/Orc6/Gsk3b/Anapc5/Cdc26/Ccnh/Cdc16/Mad2l2 |
| mmu04144 | Endocytosis | 3,6722E-11 | Ldlrap1/Dnm3/Chmp7/Rab31/Vps37c/Gbf1/Wwp1/Arrb1/Grk2/H2-Q6/H2-Q9/Vps4a/Ap2a1/Ap2a2/Ap2m1/Arf1/Arf3/Arf5/Arf6/Arpc1b/Capza1/Capza2/Capzb/Cav1/Cav2/Cbl/Cdc42/Clta/Cxcr4/Dab2/Asap1/Dnm1/Dnm2/Egfr/Ehd1/Epn1/Epn2/Eps15/Eps15l1/Acap3/Fgfr2/Fgfr3/Fgfr4/Grk4/Grk5/H2-Bl/H2-D1/H2-K1/H2-M3/H2-Q1/H2-Q10/H2-Q2/H2-Q4/H2-T22/H2-T23/H2-T24/Hgs/Hras/Hspa1l/Hspa1b/Hspa2/Igf1r/Igf2r/Itch/Kif5a/Kif5b/Kif5c/Ldlr/Smad2/Smad3/Mdm2/Rab8a/Nedd4/Pdcd6ip/Pdgfra/Pip5k1c/Pip5k1b/Pip5k1a/Prkci/Prkcz/Pld1/Pld2/Pml/Cyth1/Cyth2/Cyth3/Rab10/Rab11b/Rab4a/Rab5b/Rab5c/Rab7/Vps37d/Sh3gl2/Sh3gl1/Vps4b/Src/Chmp6/Stam/Cblb/Arfgef1/Asap2/Arap2/Agap3/Wipf1/Rab11fip3/Psd4/Eea1/Rufy1/Arrb2/Git1/Tgfbr1/Tgfbr2/Zfyve16/Traf6/Tfrc/Tsg101/Ubb/Washc5/Vps45/Snx32/Sh3glb2/Pip5kl1/Arfgap1/Spg20/Zfyve9/Asap3/Iqsec1/Ap2s1/Psd3/Chmp1a/Grk6/Git2/Rab11fip4/Rab5a/Snf8/Spg21/Washc2/Vps25/Vps26a/Bin1/Washc4/Zfyve27/Grk3/Vps37b/Agap1/Rab11fip5/Rab11a/Rabep1/Snx3/Sh3glb1/Stam2/Arpc3/Snx1/Ehd3/Pard6b/Arfgap3/Smurf2/Chmp4c/Chmp3/H2-T-ps/Gm8909/Vps28/Chmp1b/Washc3/Cltc/Rnf41/Snx2/Wipf2/Washc1/Chmp2b/Chmp2a/Vps26b/Snx4/Arap1/Smap2/Vps36/Stambp/Epn3/Ist1/Snx6/Mvb12b/Dnajc6/Wasl/Mvb12a/Psd2/Cltb/Rab11fip2/Chmp4b/Rab11fip1/Smurf1/Arpc2/Arfgap2/Rab35/Rbsn/Acap2/Cblc/Nedd4l/Usp8/Pard6g/Pard3/Smap1/Ehd4/Arfgef2 |
| mmu05161 | Hepatitis B | 4,2237E-11 | Ticam1/Ddb2/Braf/Raf1/Akt1/Akt2/Tirap/Apaf1/Birc5/Atf2/Atf4/Bad/Bax/Bcl2/Bid/Casp12/Casp3/Casp8/Casp9/Ccna2/Ccne1/Ccne2/Cdk2/Cdkn1a/Chuk/Creb1/Creb3/Crebbp/Atf6b/Cycs/Ddb1/E2f1/E2f3/Egr2/Egr3/Fadd/Fas/Fos/Tlr3/Grb2/Hras/Hspg2/Ifnar1/Ikbkb/Jak1/Jak2/Jak3/Jun/Smad3/Smad4/Mmp9/Myc/Nfatc1/Nfatc2/Nfatc3/Nfkb1/Nfkbia/Nras/Pcna/Pik3ca/Pik3cd/Pik3r1/Pik3r2/Pik3r3/Prkcb/Prkcg/Mapk11/Ptk2b/Rb1/Rela/Sos1/Sos2/Src/Stat1/Stat2/Stat3/Stat4/Stat5a/Stat5b/Stat6/Creb3l2/Tgfb1/Tgfb2/Tgfb3/Tgfbr1/Tgfbr2/Tlr4/Traf3/Traf6/Vdac3/Ticam2/Ywhaq/Ywhaz/Mavs/Ddx58/Ifna13/Creb5/Akt3/Tlr2/Ifna15/E2f2/Map2k1/Map2k2/Map2k3/Map2k4/Map2k6/Map2k7/Map3k1/Map3k7/Mapk1/Mapk10/Mapk13/Mapk14/Mapk3/Mapk8/Mapk9/Creb3l1/Irak4/Mapk12/Ep300/Irf7/Irf3/Ywhab/Tyk2/Tbk1/Ikbke/Tab1/Tab2/Ifih1/Nfatc4/Pik3cb/Creb3l4 |
| mmu04550 | Signaling pathways regulating pluripotency of stem cells | 5,7618E-11 | Raf1/Acvr1/Acvr1b/Acvr2a/Acvr2b/Akt1/Akt2/Apc/Zfhx3/Axin1/Axin2/Bmi1/Bmpr1a/Bmpr1b/Bmpr2/Ctnnb1/Dvl1/Dvl2/Dvl3/Lefty1/Smarcad1/Fgf2/Fgfr1/Fgfr2/Fgfr3/Fgfr4/Fzd1/Fzd3/Fzd4/Fzd5/Fzd6/Fzd7/Fzd8/Fzd9/Grb2/Hand1/Hesx1/Onecut1/Hras/Id1/Id2/Id3/Id4/Igf1/Igf1r/Il6st/Inhba/Inhbb/Isl1/Jak1/Jak2/Jak3/Jarid2/Klf4/Lhx5/Lif/Lifr/Smad1/Smad2/Smad3/Smad4/Smad5/Meis1/Myc/Nras/Otx1/Pax6/Pik3ca/Pik3cd/Pik3r1/Pik3r2/Pik3r3/Mapk11/Rest/Skil/Stat3/Tbx3/Tcf7/Tcf3/Wnt9a/Wnt1/Wnt10b/Wnt11/Wnt9b/Wnt2/Wnt2b/Wnt4/Wnt5a/Wnt5b/Wnt6/Pcgf2/Akt3/Apc2/Kat6a/Esrrb/Map2k1/Map2k2/Mapk1/Mapk13/Mapk14/Mapk3/Acvr1c/Mapk12/Lefty2/Rif1/Smad9/Gsk3b/Fzd2/Pcgf3/Pcgf1/Pcgf6/Nanog/Pik3cb/Pcgf5/Setdb1 |
| mmu04120 | Ubiquitin mediated proteolysis | 2,0894E-10 | Ube3c/Fbxw11/Nhlrc1/Fbxo4/Wwp1/Ddb2/Cdc20/Ube2q2/Aire/Ube3b/Birc3/Birc2/Brca1/Birc6/Btrc/Cbl/Socs3/Socs1/Ddb1/Ube2j2/Ubox5/Ube4a/Trip12/Herc2/Itch/Anapc1/Mgrn1/Mdm2/Pias2/Nedd4/Pml/Rnf7/Siah1a/Cblb/Skp1/Ube2d1/Cdc34/Ube2o/Ube2e2/Traf6/Ube2m/Ube2e3/Ube2e1/Ube2l3/Ube2i/Uba3/Ube2b/Ube2g2/Ube2h/Ube3a/Vhl/Pias3/Uba6/Fbxw8/Herc1/Klhl9/Rhobtb2/Cop1/Map3k1/Cul3/Ube2z/Cul1/Skp2/Prpf19/Fbxw7/Keap1/Prkn/Uba2/Anapc4/Cdc23/Ube2k/Anapc7/Fzr1/Stub1/Rbx1/Sae1/Pias1/Ube2d2a/Ube2l6/Pias4/Anapc5/Ube4b/Ppil2/Smurf2/Cdc26/Cul7/Ube2w/Wwp2/Fancl/Ube2g1/Herc4/Ube2r2/Elob/Ube2f/Ube2c/Trim37/Rhobtb1/Trim32/Cdc16/Ube2q1/Ubr5/Cul2/Ercc8/Herc3/Syvn1/Uba7/Cul5/Smurf1/Det1/Ube2ql1/Ube2s/Cblc/Nedd4l/Ube2n |
| mmu04068 | FoxO signaling pathway | 2,2153E-10 | Prkaa1/Prkaa2/Prkab2/Prkag2/Braf/Raf1/Akt1/Akt2/Atm/Bcl6/Bcl2l11/Bnip3/Cat/Ccnb2/Ccnd1/Ccnd2/Ccng2/Cdk2/Cdkn1a/Cdkn1b/Cdkn2b/Cdkn2d/Chuk/Plk3/Crebbp/Gadd45a/S1pr1/Egf/Egfr/G6pc/Grb2/Foxg1/Hras/Igf1/Igf1r/Ikbkb/Il10/Insr/Irs1/Irs3/Klf2/Sgk3/Smad3/Smad4/Mdm2/Gadd45b/Nlk/Nras/Pdpk1/Pik3ca/Pik3cd/Pik3r1/Pik3r2/Pik3r3/Plk1/Prkab1/Prkag1/Mapk11/Pten/Rbl2/Sgk1/Slc2a4/Plk2/Sos1/Sos2/Stat3/Stk11/Plk4/Tgfb1/Tgfb2/Tgfb3/Tgfbr1/Tgfbr2/Tnfsf10/Akt3/Gadd45g/Usp7/Map2k1/Map2k2/Mapk1/Mapk10/Mapk13/Mapk14/Mapk3/Mapk8/Mapk9/Homer1/Homer2/Homer3/Ccnb1/Sgk2/Csnk1e/Skp2/Mapk12/Ep300/Foxo6/Irs2/Foxo1/Foxo3/Gabarap/Stk4/Fbxo25/Fbxo32/G6pc3/Setd7/Pck2/Pik3cb/Gabarapl2/Sirt1 |
| mmu05212 | Pancreatic cancer | 3,0643E-10 | Ddb2/Braf/Raf1/Akt1/Akt2/Bad/Bak1/Bax/Bcl2l1/Brca2/Casp9/Ccnd1/Cdc42/Cdk4/Cdk6/Cdkn1a/Cdkn2a/Chuk/Gadd45a/E2f1/E2f3/Egf/Egfr/Erbb2/Ikbkb/Jak1/Rac3/Smad2/Smad3/Smad4/Gadd45b/Nfkb1/Pik3ca/Pik3cd/Pik3r1/Pik3r2/Pik3r3/Pld1/Pld2/Rac1/Rad51/Rb1/Rela/Ralgds/Ralbp1/Stat1/Stat3/Tgfa/Tgfb1/Tgfb2/Tgfb3/Tgfbr1/Tgfbr2/Vegfa/Akt3/Gadd45g/E2f2/Map2k1/Mapk1/Mapk10/Mapk3/Mapk8/Mapk9/Polk/Rala/Mtor/Rps6kb2/Ralb/Pik3cb |
| mmu05205 | Proteoglycans in cancer | 8,7871E-10 | Camk2d/Ank2/Braf/Raf1/Actb/Actg1/Akt1/Akt2/Ank3/Casp3/Ctnnb1/Cav1/Cav2/Cbl/Ccnd1/Cd44/Cd63/Cdc42/Cdkn1a/Col1a1/Ctsl/Cttn/Dcn/Ddx5/Egfr/Erbb2/Erbb3/Erbb4/Esr1/Drosha/Ptk2/Fas/Fgf2/Fgfr1/Fzd1/Fzd3/Fzd4/Fzd5/Fzd6/Fzd7/Fzd8/Fzd9/Gab1/Gpc1/Grb2/Ptpn6/Hbegf/Hif1a/Hras/Sdc2/Hspg2/Igf1/Igf1r/Ihh/Itga2/Itga5/Itgav/Itgb1/Itgb5/Itpr1/Itpr2/Itpr3/Kdr/Smad2/Mdm2/Met/Mmp9/Mras/Myc/Ppp1r12a/Nras/Pak1/Pdcd4/Pdpk1/Pik3ca/Pik3cd/Pik3r1/Pik3r2/Pik3r3/Prkaca/Prkacb/Prkcb/Prkcg/Plau/Plaur/Plcg1/Ppp1cb/Ppp1cc/Mapk11/Ptch1/Ptpn11/Pxn/Rac1/Rdx/Rock1/Rock2/Rps6/Rras/Shh/Slc9a1/Sos1/Sos2/Src/Stat3/Sdc1/Sdc4/Wnt9a/Tgfb1/Tgfb2/Thbs1/Tiam1/Timp3/Tlr4/Vav1/Vav2/Vegfa/Ezr/Vtn/Wnt1/Wnt10b/Wnt11/Wnt9b/Wnt2/Wnt2b/Wnt4/Wnt5a/Wnt5b/Wnt6/Ppp1r12c/Plcg2/Akt3/Tlr2/Map2k1/Map2k2/Mapk1/Mapk13/Mapk14/Mapk3/Flnb/Mapk12/Iqgap1/Smo/Frs2/Ppp1r12b/Mir21a/Mtor/Vav3/Fzd2/Rps6kb2/Rras2/Flnc/Arhgef12/Nanog/Mir10a/Plce1/Pik3cb/Eif4b/Tfap4 |
| mmu05215 | Prostate cancer | 1,2322E-09 | Etv5/Braf/Raf1/Akt1/Akt2/Atf4/Bad/Bcl2/Casp9/Ctnnb1/Ccnd1/Ccne1/Ccne2/Cdk2/Cdkn1a/Cdkn1b/Chuk/Creb1/Creb3/Crebbp/E2f1/E2f3/Egf/Egfr/Erbb2/Fgfr1/Fgfr2/Grb2/Gstp2/Gstp1/Hras/Hsp90ab1/Hsp90aa1/Igf1/Igf1r/Ikbkb/Lef1/Mdm2/Mmp9/Nfkb1/Nfkbia/Nras/Pdgfa/Pdgfb/Pdgfra/Pdgfrb/Pdpk1/Pik3ca/Pik3cd/Pik3r1/Pik3r2/Pik3r3/Plat/Plau/Pten/Rb1/Rela/Sos1/Sos2/Spint1/Creb3l2/Tcf7/Tcf7l1/Tcf7l2/Zeb1/Tgfa/Gstp3/Creb5/Akt3/Insrr/E2f2/Map2k1/Map2k2/Mapk1/Mapk3/Creb3l1/Ep300/Tmprss2/Pdgfc/Foxo1/Gsk3b/Mtor/Pdgfd/Pik3cb/Creb3l4 |
| mmu05210 | Colorectal cancer | 1,2322E-09 | Ddb2/Braf/Raf1/Akt1/Akt2/Apc/Birc5/Areg/Axin1/Axin2/Bad/Bak1/Bax/Bcl2/Bcl2l11/Casp3/Casp9/Ctnnb1/Ccnd1/Cdkn1a/Cycs/Gadd45a/Egf/Egfr/Fos/Grb2/Hras/Jun/Lef1/Rac3/Bbc3/Smad2/Smad3/Smad4/Mlh1/Msh2/Msh6/Myc/Gadd45b/Nras/Pik3ca/Pik3cd/Pik3r1/Pik3r2/Pik3r3/Rac1/Ralgds/Sos1/Sos2/Tcf7/Tcf7l1/Tcf7l2/Tgfa/Tgfb1/Tgfb2/Tgfb3/Tgfbr1/Tgfbr2/Akt3/Apc2/Gadd45g/Map2k1/Map2k2/Mapk1/Mapk10/Mapk3/Mapk8/Mapk9/Polk/Rala/Gsk3b/Mtor/Pmaip1/Rps6kb2/Ralb/Appl1/Pik3cb |
| mmu04152 | AMPK signaling pathway | 1,2322E-09 | Acacb/Prkaa1/Acaca/Prkaa2/Prkab2/Prkag2/Akt1/Akt2/Cab39/Ccna2/Ccnd1/Cd36/Cftr/Cpt1a/Creb1/Creb3/Eef2/Eef2k/Eif4ebp1/Fasn/Fbp2/Fbp1/G6pc/Gys1/Hmgcr/Hnf4a/Elavl1/Igf1/Igf1r/Insr/Irs1/Irs3/Lepr/Lipe/Pfkfb3/Rab8a/Pdpk1/Pfkfb2/Pfkl/Pfkm/Pik3ca/Pik3cd/Pik3r1/Pik3r2/Pik3r3/Pparg/Ppargc1a/Ppp2ca/Ppp2cb/Prkab1/Prkag1/Rab10/Rab11b/Rheb/Scd1/Scd2/Slc2a4/Camkk2/Srebf1/Creb3l2/Stk11/Ppp2r5d/Ulk1/Ppp2r5b/Ppp2r5a/Stradb/Creb5/Ppp2r3a/Akt3/Map3k7/Creb3l1/Ppp2r5c/Ppp2r5e/Pfkfb4/Scd3/Irs2/Ppp2r1a/Ppp2r2d/Pfkp/Foxo1/Foxo3/Mlycd/Mtor/Tbc1d1/Rps6kb2/Rab2a/Ppp2r3c/Tsc1/Akt1s1/Rab14/G6pc3/Adipor2/Cab39l/Ppp2r2a/Strada/Adipor1/Ppp2r1b/Crtc2/Rptor/Pck2/Pik3cb/Cpt1c/Creb3l4/Sirt1 |
| mmu04150 | mTOR signaling pathway | 1,5057E-09 | Prkaa1/Prkaa2/Atp6v1h/Prr5/Braf/Raf1/Sec13/Akt1/Akt2/Atp6v1a/Atp6v1e1/Cab39/Chuk/Dvl1/Dvl2/Dvl3/Eif4ebp1/Lpin1/Fzd1/Fzd3/Fzd4/Fzd5/Fzd6/Fzd7/Fzd8/Fzd9/Grb10/Grb2/Hras/Igf1/Igf1r/Ikbkb/Insr/Irs1/Lrp5/Lrp6/Nprl3/Slc3a2/Nras/Pdpk1/Pik3ca/Pik3cd/Pik3r1/Pik3r2/Pik3r3/Prkcb/Prkcg/Pten/Rheb/Rps6/Rps6ka1/Rps6ka2/Sgk1/Slc7a5/Sos1/Sos2/Stk11/Fnip1/Wnt9a/Tnfrsf1a/Ulk1/Wnt1/Wnt10b/Wnt11/Wnt9b/Wnt2/Wnt2b/Wnt4/Wnt5a/Wnt5b/Wnt6/Stradb/Mapkap1/Sesn2/Akt3/Mios/Map2k1/Map2k2/Mapk1/Mapk3/Slc38a9/Wdr24/Eif4e2/Skp2/Depdc5/Ulk2/Wdr59/Fnip2/Rragd/Rragc/Nprl2/Clip1/Gsk3b/Lamtor3/Mlst8/Mtor/Fzd2/Rps6kb2/Tsc1/Lamtor4/Atp6v1f/Atp6v1g1/Atp6v1c1/Lamtor1/Tbc1d7/Akt1s1/Rraga/Lamtor5/Atp6v1c2/Cab39l/Telo2/Castor1/Seh1l/Strada/Rptor/Ddit4/Pik3cb/Atp6v1e2/Eif4b/Rictor/Castor2/Lamtor2/Deptor |
| mmu04390 | Hippo signaling pathway | 1,5057E-09 | Fbxw11/Csnk1d/Scrib/Actb/Actg1/Amh/Apc/Birc2/Birc5/Areg/Axin1/Axin2/Bmp2/Bmp6/Bmp7/Bmp8a/Bmpr1a/Bmpr1b/Bmpr2/Btrc/Ctnna1/Ctnnb1/Ccnd1/Ccnd2/Ccnd3/Cdh1/Patj/Dlg1/Dlg4/Dvl1/Dvl2/Dvl3/Fgf1/Fzd1/Fzd3/Fzd4/Fzd5/Fzd6/Fzd7/Fzd8/Fzd9/Gli2/Id1/Id2/Ajuba/Lats1/Lef1/Llgl1/Bbc3/Crb1/Smad1/Smad2/Smad3/Smad4/Myc/Nf2/Prkci/Prkcz/Ppp1cb/Ppp1cc/Ppp2ca/Ppp2cb/Snai2/Trp53bp2/Wwc1/Tcf7/Tcf7l1/Tcf7l2/Ctnna3/Tead1/Tead2/Tead3/Tead4/Wnt9a/Llgl2/Tgfb1/Tgfb2/Tgfb3/Tgfbr1/Tgfbr2/Trp73/Wnt1/Wnt10b/Wnt11/Wnt9b/Wnt2/Wnt2b/Wnt4/Wnt5a/Wnt5b/Wnt6/Yap1/Ywhae/Ywhag/Ywhah/Ywhaq/Ywhaz/Apc2/Dlg2/Crb2/Gdf6/Csnk1e/Limd1/Frmd6/Lats2/Ppp2r1a/Ppp2r2d/Ywhab/Pals1/Stk3/Rassf1/Gsk3b/Fzd2/Pard6b/Sav1/Mob1b/Ppp2r2a/Rassf6/Ppp2r1b/Pard6g/Pard3/Nkd1/Wwtr1 |
| mmu05224 | Breast cancer | 1,8893E-09 | Ddb2/Braf/Raf1/Akt1/Akt2/Apc/Axin1/Axin2/Bak1/Bax/Brca1/Brca2/Ctnnb1/Ccnd1/Cdk4/Cdk6/Cdkn1a/Gadd45a/Dll1/Dll3/Dvl1/Dvl2/Dvl3/E2f1/E2f3/Egf/Egfr/Erbb2/Esr1/Esr2/Fgf1/Fgf15/Fgf18/Fgf2/Fgf5/Fgf8/Fgf9/Fgfr1/Flt4/Fos/Frat1/Fzd1/Fzd3/Fzd4/Fzd5/Fzd6/Fzd7/Fzd8/Fzd9/Grb2/Hes1/Hes5/Hey1/Hey2/Hras/Igf1/Igf1r/Jag1/Jun/Kit/Lef1/Lrp5/Lrp6/Myc/Gadd45b/Ncoa1/Ncoa3/Nfkb2/Notch1/Notch2/Notch3/Nras/Pik3ca/Pik3cd/Pik3r1/Pik3r2/Pik3r3/Pten/Rb1/Shc1/Shc3/Sos1/Sos2/Sp1/Frat2/Tcf7/Tcf7l1/Tcf7l2/Wnt9a/Wnt1/Wnt10b/Wnt11/Wnt9b/Wnt2/Wnt2b/Wnt4/Wnt5a/Wnt5b/Wnt6/Akt3/Apc2/Gadd45g/E2f2/Map2k1/Map2k2/Mapk1/Mapk3/Polk/Shc4/Peg12/Dll4/Heyl/Gsk3b/Mtor/Fzd2/Rps6kb2/Pik3cb/Csnk1a1 |
| mmu01522 | Endocrine resistance | 2,725E-09 | Adcy3/Braf/Raf1/Adcy6/Adcy8/Adcy9/Akt1/Akt2/Bad/Bax/Bcl2/Bik/Ccnd1/Cdk4/Cdkn1a/Cdkn1b/Cdkn2a/Cdkn2c/Dll1/Dll3/E2f1/E2f3/Egfr/Erbb2/Esr1/Esr2/Ptk2/Fos/Gnas/Grb2/Hbegf/Hras/Igf1/Igf1r/Jag1/Jun/Mdm2/Mmp9/Ncoa3/Notch1/Notch2/Notch3/Nras/Pik3ca/Pik3cd/Pik3r1/Pik3r2/Pik3r3/Prkaca/Prkacb/Med1/Mapk11/Rb1/Ncor1/Shc1/Shc3/Sos1/Sos2/Sp1/Src/Akt3/E2f2/Map2k1/Map2k2/Mapk1/Mapk10/Mapk13/Mapk14/Mapk3/Mapk8/Mapk9/Shc4/Abcb11/Mapk12/Dll4/Mtor/Rps6kb2/Carm1/Pik3cb/Gper1 |
| mmu04010 | MAPK signaling pathway | 2,9581E-09 | Mapkapk3/Arrb1/Braf/Rap1a/Raf1/Rasa2/Angpt1/Angpt2/Akt1/Akt2/Areg/Atf2/Atf4/Cacna1a/Cacna1d/Cacna1e/Cacna1g/Cacna2d1/Cacnb1/Cacnb2/Cacnb3/Cacnb4/Casp3/Cd14/Cdc25b/Cdc42/Chuk/Crk/Crkl/Csf1/Daxx/Gadd45a/Dusp2/Efna1/Efna2/Efna3/Efna4/Efna5/Egf/Egfr/Elk4/Epha2/Erbb2/Erbb3/Erbb4/Mecom/Fas/Fgf1/Fgf15/Fgf18/Fgf2/Fgf5/Fgf8/Fgf9/Fgfr1/Fgfr2/Fgfr3/Fgfr4/Flt3l/Flt4/Fos/Gna12/Gng12/Grb2/Nr4a1/Hras/Hspa1l/Hspb1/Hspa1b/Hspa2/Igf1/Igf1r/Ikbkb/Il1r1/Il1rap/Insr/Jun/Jund/Kdr/Kit/Stmn1/Rac3/Mapkapk2/Mapkapk5/Max/Mef2c/Met/Kitl/Mknk1/Mknk2/Mras/Mapt/Myc/Gadd45b/Nf1/Nfatc1/Nfatc3/Nfkb1/Nfkb2/Nlk/Nras/Dusp8/Pak1/Pdgfa/Pdgfb/Pdgfra/Pdgfrb/Pgf/Prkaca/Prkacb/Prkcb/Prkcg/Pla2g4a/Ppm1a/Ppm1b/Ppp3ca/Ppp3cb/Ppp3cc/Ppp3r1/Ppp5c/Mapk11/Mapk8ip1/Dusp1/Ptprr/Rac1/Rasgrp2/Rasgrf1/Rela/Relb/Rps6ka1/Rps6ka2/Rras/Sos1/Sos2/Rap1b/Arrb2/Taok1/Tgfa/Tgfb1/Tgfb2/Tgfb3/Tgfbr1/Tgfbr2/Rasa1/Tnfrsf1a/Traf2/Traf6/Vegfa/Vegfb/Vegfc/Pak2/Map4k3/Dusp7/Akt3/Gadd45g/Map2k5/Mapk7/Rasgrp3/Dusp5/Map2k1/Map2k2/Map2k3/Map2k4/Map2k6/Map2k7/Map3k1/Map3k12/Map3k2/Map3k3/Map3k5/Map3k7/Map3k8/Map4k1/Map4k2/Mapk1/Mapk10/Mapk13/Mapk14/Mapk3/Mapk8/Mapk9/Irak4/Map4k4/Ecsit/Flnb/Mapk12/Mapk8ip3/Dusp4/Pla2g4e/Taok3/Taok2/Map3k6/Map3k14/Pdgfc/Stk3/Rps6ka4/Lamtor3/Stk4/Dusp10/Map3k20/Tab1/Rras2/Dusp6/Tab2/Flnc/Dusp16/Tradd/Map3k13/Pdgfd/Dusp3/Rps6ka5/Rapgef2/Ntf5/Cacng7 |
| mmu05223 | Non-small cell lung cancer | 3,3985E-09 | Ddb2/Braf/Raf1/Akt1/Akt2/Bad/Bak1/Bax/Casp9/Ccnd1/Cdk4/Cdk6/Cdkn1a/Cdkn2a/Gadd45a/E2f1/E2f3/Egf/Egfr/Erbb2/Fhit/Grb2/Hras/Jak3/Gadd45b/Nras/Pdpk1/Pik3ca/Pik3cd/Pik3r1/Pik3r2/Pik3r3/Prkcb/Prkcg/Plcg1/Rb1/Rxra/Rxrg/Sos1/Sos2/Stat3/Stat5a/Stat5b/Tgfa/Rarb/Plcg2/Akt3/Gadd45g/E2f2/Map2k1/Map2k2/Mapk1/Mapk3/Polk/Rassf5/Rassf1/Foxo3/Stk4/Pik3cb/Eml4 |
| mmu05226 | Gastric cancer | 5,3617E-09 | Ddb2/Braf/Raf1/Akt1/Akt2/Apc/Axin1/Axin2/Bak1/Bax/Bcl2/Ctnna1/Ctnnb1/Ccnd1/Ccne1/Ccne2/Cdh1/Cdk2/Cdkn1a/Cdkn1b/Cdkn2b/Cdx2/Gadd45a/Dvl1/Dvl2/Dvl3/E2f1/E2f3/Egf/Egfr/Erbb2/Fgf1/Fgf15/Fgf18/Fgf2/Fgf5/Fgf8/Fgf9/Fgfr2/Frat1/Fzd1/Fzd3/Fzd4/Fzd5/Fzd6/Fzd7/Fzd8/Fzd9/Gab1/Grb2/Hras/Jup/Lef1/Lrp5/Lrp6/Smad2/Smad3/Smad4/Met/Mlh1/Myc/Gadd45b/Nras/Abcb1b/Abcb1a/Pik3ca/Pik3cd/Pik3r1/Pik3r2/Pik3r3/Rb1/Rxra/Rxrg/Shc1/Shc3/Shh/Sos1/Sos2/Frat2/Tcf7/Tcf7l1/Tcf7l2/Ctnna3/Wnt9a/Terc/Tert/Tgfb1/Tgfb2/Tgfb3/Tgfbr1/Tgfbr2/Rarb/Wnt1/Wnt10b/Wnt11/Wnt9b/Wnt2/Wnt2b/Wnt4/Wnt5a/Wnt5b/Wnt6/Akt3/Apc2/Gadd45g/E2f2/Map2k1/Map2k2/Mapk1/Mapk3/Polk/Shc4/Peg12/Gsk3b/Mtor/Fzd2/Rps6kb2/Pik3cb/Csnk1a1 |
| mmu04210 | Apoptosis | 9,2316E-09 | Raf1/Actb/Actg1/Parp1/Parp2/Akt1/Akt2/Apaf1/Birc3/Birc2/Birc5/Atf4/Atm/Bad/Bak1/Bax/Bcl2/Bcl2l1/Bid/Hrk/Bcl2l11/Capn1/Capn2/Casp12/Casp2/Casp3/Casp6/Casp7/Casp8/Casp9/Cflar/Chuk/Ctsb/Ctsc/Ctsd/Ctsh/Ctsl/Cycs/Daxx/Gadd45a/Dffa/Dffb/Eif2s1/Eif2ak3/Endog/Fadd/Fas/Fos/Hras/Ikbkb/Il3ra/Itpr1/Itpr2/Itpr3/Jun/Lmna/Lmnb1/Lmnb2/Bbc3/Mcl1/Gadd45b/Nfkb1/Nfkbia/Nras/Pdpk1/Pik3ca/Pik3cd/Pik3r1/Pik3r2/Pik3r3/Septin4/Ptpn13/Rela/Ripk1/Sptan1/Tnfrsf10b/Tnfrsf1a/Traf1/Traf2/Tnfsf10/Tuba1a/Tuba1b/Tuba3a/Tuba4a/Tuba1c/Ctso/Parp3/Akt3/Tubal3/Gadd45g/Map2k1/Map2k2/Map3k5/Mapk1/Mapk10/Mapk3/Mapk8/Mapk9/Tuba8/Map3k14/Ctsf/Pidd1/Pmaip1/Ctsz/Diablo/Dab2ip/Tradd/Pik3cb/Ern1 |
| mmu04520 | Adherens junction | 1,2227E-08 | Baiap2/Actn1/Acp1/Actb/Actg1/Ctnna1/Ctnnb1/Ctnnd1/Cdc42/Cdh1/Crebbp/Csnk2a1/Csnk2a2/Csnk2b/Egfr/Erbb2/Fer/Fgfr1/Fyn/Ptpn6/Igf1r/Insr/Lef1/Rac3/Smad3/Smad4/Met/Afdn/Nlk/Ptpn1/Ptprf/Ptprj/Ptprm/Nectin2/Rac1/Sorbs1/Snai2/Snai1/Src/Tcf7/Tcf7l1/Tcf7l2/Ctnna3/Tgfbr1/Tgfbr2/Tjp1/Vcl/Yes1/Farp2/Wasf2/Map3k7/Mapk1/Mapk3/Iqgap1/Ep300/Lmo7/Nectin1/Nectin3/Actn4/Nectin4/Wasl/Pard3/Ssx2ip |
| mmu04218 | Cellular senescence | 1,9964E-08 | Fbxw11/E2f4/Raf1/H2-Q6/H2-Q9/Akt1/Akt2/Slc25a4/Atm/Zfp36l1/Zfp36l2/Btrc/Cacna1d/Calm1/Calm2/Calm3/Capn1/Capn2/Ccna2/Ccnb2/Ccnd1/Ccnd2/Ccnd3/Ccne1/Ccne2/Cdc25a/Cdk1/Cdk2/Cdk4/Cdk6/Cdkn1a/Cdkn2a/Cdkn2b/Chek1/Gadd45a/E2f1/E2f3/E2f5/Eif4ebp1/Foxm1/Gata4/H2-Bl/H2-D1/H2-K1/H2-M3/H2-Q1/H2-Q10/H2-Q2/H2-Q4/H2-T22/H2-T23/H2-T24/Hipk1/Hipk2/Hipk3/Hras/Hus1/Itpr1/Itpr2/Itpr3/Smad2/Smad3/Mapkapk2/Mdm2/Mras/Mre11a/Mybl2/Myc/Gadd45b/Nfatc1/Nfatc2/Nfatc3/Nfkb1/Nras/Sqstm1/Pik3ca/Pik3cd/Pik3r1/Pik3r2/Pik3r3/Ppp1cb/Ppp1cc/Ppp3ca/Ppp3cb/Ppp3cc/Ppp3r1/Mapk11/Pten/Rad1/Rad50/Rad9a/Rb1/Rbbp4/Rbl1/Rbl2/Rela/Rheb/Rras/Mcu/Tgfb1/Tgfb2/Tgfb3/Tgfbr1/Tgfbr2/Vdac1/Vdac2/Vdac3/Lin54/Rad9b/Akt3/Ets1/Gadd45g/E2f2/Atr/Map2k1/Map2k2/Map2k3/Map2k6/Mapk1/Mapk13/Mapk14/Mapk3/Ccnb1/Nbn/Mapk12/Rassf5/Foxo1/Foxo3/Mtor/Trpm7/Trpv4/Tsc1/H2-T-ps/Gm8909/Rras2/Ppid/Lin9/Nfatc4/Pik3cb/Lin37/Sirt1 |
| mmu04910 | Insulin signaling pathway | 2,7781E-08 | Acacb/Phkb/Gck/Rhoq/Ppp1r3e/Prkaa1/Trip10/Acaca/Rapgef1/Prkaa2/Prkab2/Prkag2/Braf/Pygb/Pygl/Raf1/Akt1/Akt2/Bad/Calm1/Calm2/Calm3/Cbl/Socs3/Socs1/Crk/Crkl/Eif4ebp1/Fasn/Fbp2/Fbp1/Flot1/Flot2/G6pc/Grb2/Gys1/Hk1/Hk2/Hras/Ikbkb/Inppl1/Insr/Irs1/Irs3/Lipe/Mknk1/Mknk2/Nras/Pde3b/Pdpk1/Pik3ca/Pik3cd/Pik3r1/Pik3r2/Pik3r3/Prkaca/Prkacb/Prkci/Prkcz/Pklr/Ppargc1a/Ppp1cb/Ppp1cc/Inpp5k/Prkab1/Prkag1/Prkar1a/Prkar1b/Prkar2a/Prkar2b/Ptpn1/Ptprf/Rheb/Rps6/Sorbs1/Shc1/Shc3/Slc2a4/Sos1/Sos2/Srebf1/Cblb/Hkdc1/Socs2/Ppp1r3d/Akt3/Sh2b2/Ppp1r3b/Map2k1/Map2k2/Mapk1/Mapk10/Mapk3/Mapk8/Mapk9/Eif4e2/Shc4/Irs2/Ppp1r3c/Exoc7/Foxo1/Gsk3b/Mtor/Rps6kb2/Tsc1/G6pc3/Phkg2/Rptor/Pck2/Pik3cb |
| mmu04360 | Axon guidance | 3,7176E-08 | Unc5a/Unc5b/Camk2d/Sema3d/Raf1/Abl1/Srgap1/Rhod/Bmp7/Bmpr1b/Bmpr2/Cdc42/Cdk5/Cfl1/Cfl2/Cxcr4/Dpysl2/Efna1/Efna2/Efna3/Efna4/Efna5/Efnb2/Enah/Epha1/Epha2/Epha3/Epha4/Epha5/Epha7/Ephb3/Ephb4/Ephb6/Plxnb2/Ptk2/Fes/Fyn/Fzd3/Gnai1/Gnai2/Gnai3/Hras/Itgb1/Limk1/Limk2/Rac3/Ntng2/Met/Nck1/Nck2/Neo1/Nfatc2/Nfatc3/Nras/Ntn1/Ntn3/Pak1/Pik3ca/Pik3cd/Pik3r1/Pik3r2/Pik3r3/Prkcz/Plcg1/Plxna1/Plxna2/Ppp3ca/Ppp3cb/Ppp3cc/Ppp3r1/Ptch1/Ptpn11/Rac1/Robo3/Rock1/Rock2/Rras/Ryk/Cxcl12/Sema3a/Sema3c/Sema3f/Sema4a/Sema4b/Sema4c/Sema4d/Sema4f/Sema6a/Sema6b/Sema6c/Sema7a/Shh/Slit2/Slit3/Src/Unc5d/Pak6/Sema6d/Rasa1/Trpc1/Unc5c/Rnd1/Pak2/Wnt4/Wnt5a/Wnt5b/Ablim1/Pdk1/Ablim2/Ssh1/Plcg2/Plxnb1/Ssh2/Plxna4/Rgma/Ssh3/Srgap3/Mapk1/Mapk3/Sema4g/Ephb1/Ablim3/Smo/Rgs3/Plxnc1/Gsk3b/Ntn4/Pard6b/Dpysl5/Myl12a/Myl12b/Arhgef12/Nfatc4/Pik3cb/Ntng1/Pard6g/Pard3 |
| mmu01521 | EGFR tyrosine kinase inhibitor resistance | 4,3041E-08 | Nrg2/Braf/Raf1/Akt1/Akt2/Bad/Bax/Bcl2/Bcl2l1/Bcl2l11/Egf/Egfr/Eif4ebp1/Erbb2/Erbb3/Fgf2/Fgfr2/Fgfr3/Gab1/Gas6/Grb2/Hras/Igf1/Igf1r/Il6ra/Jak1/Jak2/Kdr/Met/Nf1/Nras/Pdgfa/Pdgfb/Pdgfra/Pdgfrb/Pik3ca/Pik3cd/Pik3r1/Pik3r2/Pik3r3/Prkcb/Prkcg/Plcg1/Pten/Rps6/Shc1/Shc3/Sos1/Sos2/Src/Stat3/Tgfa/Vegfa/Plcg2/Akt3/Map2k1/Map2k2/Mapk1/Mapk3/Eif4e2/Shc4/Pdgfc/Foxo3/Gsk3b/Mtor/Rps6kb2/Pdgfd/Pik3cb |
| mmu04668 | TNF signaling pathway | 1,2256E-07 | Akt1/Akt2/Birc3/Birc2/Atf2/Atf4/Bcl3/Casp3/Casp7/Casp8/Cebpb/Cflar/Chuk/Socs3/Creb1/Creb3/Atf6b/Csf1/Edn1/Fadd/Fas/Fos/Cxcl1/Icam1/Cxcl10/Ifi47/Ikbkb/Il15/Irf1/Itch/Jag1/Jun/Junb/Lif/Mmp14/Mmp9/Nfkb1/Nfkbia/Pik3ca/Pik3cd/Pik3r1/Pik3r2/Pik3r3/Mapk11/Ptgs2/Rela/Ripk1/Ccl5/Cx3cl1/Creb3l2/Tnfaip3/Tnfrsf1a/Tnfrsf1b/Traf1/Traf2/Traf3/Traf5/Vegfc/Creb5/Akt3/Nod2/Map2k1/Map2k3/Map2k4/Map2k6/Map2k7/Map3k5/Map3k7/Map3k8/Mapk1/Mapk10/Mapk13/Mapk14/Mapk3/Mapk8/Mapk9/Creb3l1/Mapk12/Map3k14/Ripk3/Rps6ka4/Tab1/Tab2/Dab2ip/Tradd/Pgam5/Rps6ka5/Dnm1l/Mlkl/Pik3cb/Creb3l4 |
| mmu04935 | Growth hormone synthesis, secretion and action | 1,2566E-07 | Adcy3/Raf1/Adcy6/Adcy8/Adcy9/Akt1/Akt2/Atf2/Atf4/Cacna1d/Socs3/Socs1/Creb1/Creb3/Crebbp/Atf6b/Bcar1/Crk/Crkl/Ptk2/Fos/Ghr/Gna11/Gnai1/Gnai2/Gnai3/Gnaq/Gnas/Grb2/Hras/Igf1/Igfals/Irs1/Irs3/Itpr1/Itpr2/Itpr3/Jak2/Junb/Nras/Pik3ca/Pik3cd/Pik3r1/Pik3r2/Pik3r3/Prkaca/Prkacb/Prkcb/Prkcg/Plcb1/Plcb2/Plcb3/Plcb4/Plcg1/Mapk11/Shc1/Shc3/Sst/Sstr1/Sstr2/Sos1/Sos2/Stat1/Stat3/Stat5a/Stat5b/Creb3l2/Socs2/Creb5/Plcg2/Akt3/Map2k1/Map2k2/Map2k3/Map2k4/Map2k6/Map3k1/Mapk1/Mapk10/Mapk13/Mapk14/Mapk3/Mapk8/Mapk9/Creb3l1/Shc4/Mapk12/Ep300/Irs2/Gsk3b/Mtor/Pik3cb/Creb3l4 |
| mmu05213 | Endometrial cancer | 1,2816E-07 | Ddb2/Braf/Raf1/Akt1/Akt2/Apc/Axin1/Axin2/Bad/Bak1/Bax/Casp9/Ctnna1/Ctnnb1/Ccnd1/Cdh1/Cdkn1a/Gadd45a/Egf/Egfr/Erbb2/Grb2/Hras/Lef1/Mlh1/Myc/Gadd45b/Nras/Pdpk1/Pik3ca/Pik3cd/Pik3r1/Pik3r2/Pik3r3/Pten/Sos1/Sos2/Tcf7/Tcf7l1/Tcf7l2/Ctnna3/Akt3/Apc2/Gadd45g/Map2k1/Map2k2/Mapk1/Mapk3/Polk/Foxo3/Gsk3b/Pik3cb |
| mmu05222 | Small cell lung cancer | 2,1874E-07 | Ddb2/Akt1/Akt2/Apaf1/Birc3/Birc2/Bak1/Bax/Bcl2/Bcl2l1/Casp3/Casp9/Ccnd1/Ccne1/Ccne2/Cdk2/Cdk4/Cdk6/Cdkn1a/Cdkn1b/Cdkn2b/Chuk/Col4a1/Cycs/Gadd45a/E2f1/E2f3/Ptk2/Fhit/Ikbkb/Itga2/Itga2b/Itga3/Itga6/Itgav/Itgb1/Lama1/Lama3/Lama4/Lama5/Lamb2/Lamb3/Lamc2/Max/Myc/Gadd45b/Nfkb1/Nfkbia/Nos2/Pik3ca/Pik3cd/Pik3r1/Pik3r2/Pik3r3/Pten/Ptgs2/Rb1/Rela/Rxra/Rxrg/Rarb/Traf1/Traf2/Traf3/Traf4/Traf5/Traf6/Zbtb17/Lamc1/Akt3/Gadd45g/E2f2/Polk/Skp2/Cks2/Pik3cb |
| mmu04070 | Phosphatidylinositol signaling system | 2,5938E-07 | Inpp5f/Synj1/Dgkz/Pi4kb/Dgkg/Dgkq/Cds2/Impa2/Pip4k2c/Calm1/Calm2/Calm3/Dgka/Inpp1/Inpp5b/Inppl1/Itpr1/Itpr2/Itpr3/Mtmr4/Inpp5j/Pik3c2a/Pik3c2g/Pik3ca/Pik3cd/Pik3r1/Pik3r2/Pik3r3/Pip5k1c/Pip4k2a/Pip5k1b/Pip5k1a/Prkcb/Prkcg/Plcb1/Plcb2/Plcb3/Plcb4/Plcd1/Plcg1/Inpp5k/Pten/Synj2/Inpp5a/Dgkb/Itpk1/Mtmr6/Pi4ka/Pik3c3/Dgkd/Ppip5k2/Itpka/Itpkc/Inpp4b/Plcg2/Pik3c2b/Bpnt2/Inpp4a/Ip6k1/Dgki/Itpkb/Ppip5k1/Dgkh/Mtmr7/Impa1/Dgke/Inpp5e/Pi4k2b/Ipmk/Pip4p2/Plce1/Mtmr3/Cds1/Pik3cb/Ippk/Ip6k2/Mtmr2/Sacm1l/Pi4k2a/Mtmr14 |
| mmu01524 | Platinum drug resistance | 2,5938E-07 | Gstt3/Akt1/Akt2/Apaf1/Birc3/Birc2/Birc5/Atm/Atp7b/Bad/Bak1/Bax/Bcl2/Bcl2l1/Bid/Brca1/Casp3/Casp8/Casp9/Cdkn1a/Cdkn2a/Cycs/Erbb2/Ercc1/Fadd/Fas/Gsta3/Gsta4/Gstm1/Gstm2/Gstm3/Gstm4/Gstm5/Gstp2/Gstp1/Gstt1/Gstt2/Gsto1/Bbc3/Mdm2/Mlh1/Msh2/Msh6/Pdpk1/Pik3ca/Pik3cd/Pik3r1/Pik3r2/Pik3r3/Rev3l/Slc31a1/Mgst2/Top2a/Top2b/Gstp3/Xpa/Akt3/Map3k5/Mapk1/Mapk3/Mgst1/Pmaip1/Mgst3/Gsto2/Gstm7/Pik3cb |
| mmu04115 | p53 signaling pathway | 2,5938E-07 | Ddb2/Apaf1/Atm/Bax/Bcl2/Bcl2l1/Bid/Casp3/Casp8/Casp9/Ccnb2/Ccnd1/Ccnd2/Ccnd3/Ccne1/Ccne2/Ccng1/Ccng2/Cd82/Cdk1/Cdk2/Cdk4/Cdk6/Cdkn1a/Cdkn2a/Chek1/Cycs/Gadd45a/Ei24/Sesn1/Fas/Igf1/Bbc3/Mdm2/Mdm4/Gadd45b/Pten/Rrm2/Siah1a/Thbs1/Trp73/Zmat3/Sesn2/Gadd45g/Atr/Cop1/Ccnb1/Gtse1/Siva1/Rrm2b/Ppm1d/Sfn/Pidd1/Pmaip1/Perp/Shisa5/Rprm/Steap3/Aifm2/Sesn3/Gorab |
| mmu05418 | Fluid shear stress and atherosclerosis | 2,5938E-07 | Gstt3/Prkaa1/Prkaa2/Actb/Actg1/Acvr1/Acvr2a/Acvr2b/Akt1/Akt2/Ass1/Bcl2/Bmpr1a/Bmpr1b/Bmpr2/Calm1/Calm2/Calm3/Ctnnb1/Cav1/Cav2/Chuk/Ctsl/Cyba/Edn1/Ptk2/Fos/Gpc1/Gsta3/Gsta4/Gstm1/Gstm2/Gstm3/Gstm4/Gstm5/Gstp2/Gstp1/Gstt1/Gstt2/Gsto1/Hmox1/Hsp90ab1/Hsp90aa1/Sdc2/Icam1/Ikbkb/Il1r1/Itga2b/Itgav/Jun/Kdr/Klf2/Arhgef2/Rac3/Sumo2/Mef2a/Mef2c/Mmp9/Nfe2l2/Nfkb1/Nqo1/Nppc/Sqstm1/Pdgfa/Pdgfb/Pik3ca/Pik3cd/Pik3r1/Pik3r2/Pik3r3/Prkcz/Plat/Mapk11/Dusp1/Rac1/Rela/Sumo3/Src/Sdc1/Sdc4/Mgst2/Thbd/Tnfrsf1a/Txn1/Sumo1/Vegfa/Gstp3/Akt3/Map2k5/Mapk7/Map2k4/Map2k6/Map2k7/Map3k5/Map3k7/Mapk10/Mapk13/Mapk14/Mapk8/Mapk9/Mapk12/Keap1/Txn2/Mgst1/Pias4/Trpv4/Mgst3/Gsto2/Gstm7/Mir10a/Pik3cb |
| mmu05165 | Human papillomavirus infection | 2,5938E-07 | Isg15/Tada3/Tubg1/Tubg2/Maml1/Itga9/Scrib/Ticam1/Atp6v1h/Raf1/H2-Q6/H2-Q9/Atp6v0b/Akt1/Akt2/Apc/Atm/Atp6v1a/Atp6v0d1/Atp6v1e1/Atp6v0e/Atp6v0a1/Atp6v0c/Axin1/Axin2/Bad/Bak1/Bax/Casp3/Casp8/Ctnnb1/Ccna2/Ccnd1/Ccnd2/Ccnd3/Ccne1/Ccne2/Cdc42/Cdk2/Cdk4/Cdk6/Cdkn1a/Cdkn1b/Chad/Chuk/Patj/Col4a1/Col1a1/Creb1/Creb3/Crebbp/Dlg1/Dvl1/Dvl2/Dvl3/E2f1/Egf/Egfr/Eif4ebp1/Fadd/Ptk2/Fas/Tlr3/Fzd1/Fzd3/Fzd4/Fzd5/Fzd6/Fzd7/Fzd8/Fzd9/Gnas/Grb2/Magi1/H2-Bl/H2-D1/H2-K1/H2-M3/H2-Q1/H2-Q10/H2-Q2/H2-Q4/H2-T22/H2-T23/H2-T24/Hdac2/Hes1/Hes2/Hes5/Hey1/Hey2/Hras/Ifnar1/Ifnar2/Ikbkb/Irf1/Irf9/Itga2/Itga2b/Itga3/Itga5/Itga6/Itga7/Itgav/Itgb1/Itgb5/Itgb6/Itgb7/Jag1/Jak1/Lama1/Lama3/Lama4/Lama5/Lamb2/Lamb3/Lamc2/Lfng/Llgl1/Mdm2/Mx2/Nfkb1/Notch1/Notch2/Notch3/Nras/Pdgfrb/Pik3ca/Pik3cd/Pik3r1/Pik3r2/Pik3r3/Pkm/Prkaca/Prkacb/Prkci/Prkcz/Ppp2ca/Ppp2cb/Eif2ak2/Psen1/Psmc1/Pten/Ptger4/Ptgs2/Itgb4/Pxn/Rb1/Rbl1/Rbl2/Rbpj/Rela/Rfng/Rheb/Sos1/Sos2/Spp1/Stat1/Stat2/Creb3l2/Tcf7/Tcf7l1/Tcf7l2/Wnt9a/Llgl2/Tert/Ppp2r5d/Thbs1/Atp6v0a2/Tnfrsf1a/Traf3/Ube3a/Vegfa/Vtn/Wnt1/Wnt10b/Wnt11/Wnt9b/Wnt2/Wnt2b/Wnt4/Wnt5a/Wnt5b/Wnt6/Crb3/Ppp2r5b/Lamc1/Ppp2r5a/Ifna13/Oasl1/Creb5/Ppp2r3a/Tbpl1/Akt3/Apc2/Dlg2/Oasl2/Itga8/Ifna15/Atr/Map2k1/Map2k2/Mapk1/Mapk3/Creb3l1/Ppp2r5c/Ppp2r5e/Slc9a3r1/Maml2/Tcirg1/Itgb8/Ep300/Maml3/Hdac1/Ppp2r1a/Ppp2r2d/Irf3/Tyk2/Hes6/Heyl/Pals1/Foxo1/Tbk1/Ikbke/Gsk3b/Mtor/Fzd2/Pard6b/Rps6kb2/Ppp2r3c/Tsc1/Atp6v1f/Atp6v1g1/Atp6v1c1/H2-T-ps/Gm8909/Atp6v1c2/Ubr4/Tradd/Ppp2r2a/Ppp2r1b/Nfx1/Pik3cb/Atp6v1e2/Creb3l4/Hes7/Csnk1a1/Pard6g/Pard3 |
| mmu04137 | Mitophagy - animal | 3,5321E-07 | Usp30/Atg5/Atf4/Bcl2l1/Bnip3/Bnip3l/Csnk2a1/Csnk2a2/Csnk2b/E2f1/Eif2ak3/Usp15/Hif1a/Hras/Jun/Mfn2/Mitf/Mras/Cited2/Nras/Sqstm1/Rab7/Rela/Rras/Sp1/Src/Atg9b/Tfeb/Rhot2/Ubb/Ulk1/Ambra1/Tbc1d17/Atg9a/Mapk10/Mapk8/Mapk9/Prkn/Tax1bp1/Becn1/Tbk1/Foxo3/Gabarap/Rhot1/Tomm7/Fis1/Tbc1d15/Rras2/Mfn1/Pink1/Optn/Pgam5/Usp8/Gabarapl2/Bcl2l13 |
| mmu04530 | Tight junction | 3,8672E-07 | Arhgef18/Synpo/Scrib/Prkaa1/Prkaa2/Prkab2/Prkag2/Actn1/Rap1a/Actb/Actg1/Cacna1d/Runx1/Ccnd1/Cd1d1/Cdc42/Cdk4/Cftr/Patj/Cldn1/Cldn3/Cttn/Dlg1/Erbb2/Gata4/Magi1/Hspa4/Itgb1/F11r/Jun/Arhgef2/Llgl1/Rab8a/Afdn/Mpdz/Myh11/Myh9/Myl6/Nedd4/Nf2/Ocln/Cldn11/Pcna/Prkaca/Prkacb/Prkce/Prkci/Prkcz/Ppp2ca/Ppp2cb/Prkab1/Prkag1/Rapgef6/Rac1/Rdx/Rock1/Rock2/Src/Stk11/Myl6b/Llgl2/Tiam1/Marveld2/Tjp1/Tjp2/Tuba1a/Tuba1b/Tuba3a/Tuba4a/Tuba1c/Vasp/Ezr/Crb3/Micall2/Rab8b/Tubal3/Dlg2/Actr3b/Map2k7/Map3k1/Map3k5/Mapk10/Mapk8/Mapk9/Slc9a3r1/Tjp3/Whamm/Ppp2r1a/Ppp2r2d/Cldn7/Tuba8/Epb41l4b/Cldn6/Cldn8/Myh13/Pals1/Amotl2/Ybx3/Cldn9/Cldn10/Pard6b/Actn4/Actr2/Myh7b/Myl12a/Jam2/Myl12b/Cgnl1/Rab13/Cgn/Cldn23/Myh14/Ppp2r2a/Igsf5/Marveld3/Ppp2r1b/Tjap1/Actr3/Amotl1/Rapgef2/Myh10/Nedd4l/Jam3/Pard6g/Pard3 |
| mmu04071 | Sphingolipid signaling pathway | 4,7772E-07 | Raf1/Adora1/Akt1/Akt2/Asah1/Bax/Bcl2/Bdkrb2/Bid/Ctsd/Degs1/S1pr1/S1pr3/Fyn/Gab2/Gna12/Gnai1/Gnai2/Gnai3/Gnaq/S1pr2/Hras/Rac3/Abcc1/Nfkb1/Nras/Nsmaf/Pdpk1/Pik3ca/Pik3cd/Pik3r1/Pik3r2/Pik3r3/Prkcb/Prkcg/Prkce/Prkcz/Plcb1/Plcb2/Plcb3/Plcb4/Pld1/Pld2/Ppp2ca/Ppp2cb/Mapk11/Pten/Rac1/Rela/Rock1/Rock2/Sgpl1/Smpd1/Smpd2/Sphk1/Sptlc2/Sgms1/Ppp2r5d/Tnfrsf1a/Traf2/Ppp2r5b/Ppp2r5a/Acer2/Ppp2r3a/Akt3/Cers6/Map2k1/Map2k2/Map3k5/Mapk1/Mapk10/Mapk13/Mapk14/Mapk3/Mapk8/Mapk9/Sptlc1/Ppp2r5c/Ppp2r5e/Mapk12/Kng2/Sgpp2/Ppp2r1a/Ppp2r2d/Asah2/Ppp2r3c/Cers4/Degs2/Tradd/Cers5/Ppp2r2a/Ppp2r1b/Sgms2/Pik3cb/Cers2/Sgpp1/S1pr5 |
| mmu05203 | Viral carcinogenesis | 7,2619E-07 | H4c17/Scrib/Cdc20/Actn1/H2-Q6/H2-Q9/Rasa2/Atf2/Atf4/Atp6v0d1/Bad/Bak1/Bax/Casp3/Casp8/Ccna2/Ccnd1/Ccnd2/Ccnd3/Ccne1/Ccne2/Cdk1/Cdc42/Cdk2/Cdk4/Cdk6/Cdkn1a/Cdkn1b/Cdkn2a/Cdkn2b/Chek1/Creb1/Creb3/Crebbp/Atf6b/Ddb1/Dlg1/Egr2/Egr3/Kat2a/Grb2/Gtf2h1/Gtf2h4/H2-Bl/H2-D1/H2-K1/H2-M3/H2-Q1/H2-Q10/H2-Q2/H2-Q4/H2-T22/H2-T23/H2-T24/Hdac2/Hdac3/Hdac5/Hnrnpk/Hpn/Hras/Il6st/Irf9/Jak1/Jak3/Jun/Ltbr/Hdac10/Lyn/Mad1l1/Mapkapk2/Mdm2/Nfkb1/Nfkb2/Nfkbia/Nras/Kat2b/Pik3ca/Pik3cd/Pik3r1/Pik3r2/Pik3r3/Pkm/Prkaca/Prkacb/Polb/Eif2ak2/Psmc1/Pxn/Rac1/Rb1/Rbl1/Rbl2/Rbpj/Rel/Rela/Scin/Sp100/Src/Stat3/Stat5a/Stat5b/Creb3l2/Hdac4/Syk/Traf1/Traf2/Traf3/Traf5/Ube3a/Vdac3/Ywhae/Ywhag/Ywhah/Ywhaq/Ywhaz/Gsn/Gtf2b/Creb5/Hdac11/Vac14/Gtf2a2/Tbpl1/Gtf2h2/Usp7/Mapk1/Mapk3/Creb3l1/Skp2/H4c3/H4c4/H4c6/H4c9/H4c11/H4c12/H4c18/H2bc1/H2bc6/H2bc11/H2bc13/H2bc14/H2bc18/H4f16/H4c1/H4c2/Ep300/Hdac1/Irf7/Irf3/Ywhab/Hdac7/Snd1/Pmaip1/Actn4/Snw1/H2-T-ps/Gm8909/Gtf2e2/Ubr4/H4c8/Tradd/Pik3cb/Creb3l4/H2bu2/Gtf2a1/Dnaja3/H4c14 |
| mmu04015 | Rap1 signaling pathway | 8,1991E-07 | Prkd2/Adcy3/Rapgef1/Braf/Rap1a/Raf1/Rap1gap/Actb/Actg1/Adcy6/Adcy8/Adcy9/Adora2a/Adora2b/Angpt1/Angpt2/Akt1/Akt2/Calm1/Calm2/Calm3/Ctnnb1/Ctnnd1/Cdc42/Cdh1/Cnr1/Bcar1/Crk/Crkl/Csf1/Efna1/Efna2/Efna3/Efna4/Efna5/Egf/Egfr/Enah/Epha2/F2r/Fgf1/Fgf15/Fgf18/Fgf2/Fgf5/Fgf8/Fgf9/Fgfr1/Fgfr2/Fgfr3/Fgfr4/Flt4/Gnai1/Gnai2/Gnai3/Gnao1/Gnaq/Gnas/Lpar1/Magi1/Hras/Id1/Igf1/Igf1r/Insr/Itga2b/Itgam/Itgb1/Kdr/Kit/Rac3/Met/Kitl/Afdn/Mras/Nras/P2ry1/Pdgfa/Pdgfb/Pdgfra/Pdgfrb/Pfn1/Pfn2/Pgf/Pik3ca/Pik3cd/Pik3r1/Pik3r2/Pik3r3/Prkcb/Prkcg/Prkci/Prkd1/Prkcz/Plcb1/Plcb2/Plcb3/Plcb4/Plcg1/Mapk11/Rapgef6/Rac1/Rasgrp2/Ralgds/Rras/Sipa1/Src/Rap1b/Sipa1l1/Rapgef5/Thbs1/Tiam1/Vasp/Vav1/Vav2/Vegfa/Vegfb/Vegfc/Rapgef3/Farp2/Akt3/Dock4/Fyb/Rasgrp3/Sipa1l2/Map2k1/Map2k2/Map2k3/Map2k6/Mapk1/Mapk13/Mapk14/Mapk3/Mapk12/Magi2/Lpar2/Rassf5/Apbb1ip/Pdgfc/Rala/Rapgef4/Vav3/Pard6b/Ralb/Lpar3/Tln2/Pdgfd/Plce1/Sipa1l3/Pik3cb/Prkd3/Rapgef2/Pard6g/Pard3/Magi3 |
| mmu05211 | Renal cell carcinoma | 8,2279E-07 | Rapgef1/Braf/Rap1a/Raf1/Egln1/Egln2/Egln3/Akt1/Akt2/Arnt/Bad/Cdc42/Cdkn1a/Crebbp/Crk/Crkl/Epas1/Fh1/Gab1/Grb2/Hif1a/Hras/Jun/Met/Nras/Pak1/Pdgfb/Pik3ca/Pik3cd/Pik3r1/Pik3r2/Pik3r3/Ptpn11/Rac1/Slc2a1/Sos1/Sos2/Pak6/Rap1b/Tgfa/Tgfb1/Tgfb2/Tgfb3/Vegfa/Vhl/Pak2/Akt3/Ets1/Map2k1/Map2k2/Mapk1/Mapk3/Ep300/Rbx1/Elob/Cul2/Pik3cb/Prcc |
| mmu04012 | ErbB signaling pathway | 1,1222E-06 | Nrg2/Camk2d/Braf/Raf1/Abl1/Abl2/Akt1/Akt2/Areg/Bad/Btc/Cbl/Cdkn1a/Cdkn1b/Crk/Crkl/Egf/Egfr/Eif4ebp1/Erbb2/Erbb3/Erbb4/Ptk2/Gab1/Grb2/Hbegf/Hras/Jun/Myc/Nck1/Nck2/Nras/Pak1/Pik3ca/Pik3cd/Pik3r1/Pik3r2/Pik3r3/Prkcb/Prkcg/Plcg1/Shc1/Shc3/Sos1/Sos2/Src/Stat5a/Stat5b/Cblb/Pak6/Tgfa/Pak2/Plcg2/Akt3/Map2k1/Map2k2/Map2k4/Map2k7/Mapk1/Mapk10/Mapk3/Mapk8/Mapk9/Shc4/Gsk3b/Mtor/Rps6kb2/Pik3cb/Nrg4 |
| mmu04310 | Wnt signaling pathway | 1,1935E-06 | Fbxw11/Prickle1/Lgr4/Camk2d/Semp2l2b/Apc/Axin1/Axin2/Btrc/Cacybp/Ctnnb1/Ccnd1/Ccnd2/Ccnd3/Crebbp/Csnk2a1/Csnk2a2/Csnk2b/Ctbp1/Ctbp2/Dvl1/Dvl2/Dvl3/Lgr5/Fosl1/Frat1/Fzd1/Fzd3/Fzd4/Fzd5/Fzd6/Fzd7/Fzd8/Fzd9/Invs/Jun/Lef1/Lrp5/Lrp6/Rac3/Smad3/Smad4/Myc/Nfatc1/Nfatc2/Nfatc3/Nlk/Prkaca/Prkacb/Prkcb/Prkcg/Plcb1/Plcb2/Plcb3/Plcb4/Ppard/Ppp3ca/Ppp3cb/Ppp3cc/Ppp3r1/Psen1/Rac1/Rock2/Ryk/Sfrp2/Sfrp1/Siah1a/Sox17/Rnf43/Daam1/Frat2/Skp1/Tcf7/Tcf7l1/Tcf7l2/Wnt9a/Wnt1/Wnt10b/Wnt11/Wnt9b/Wnt2/Wnt2b/Wnt4/Wnt5a/Wnt5b/Wnt6/Vangl1/Apc2/Wif1/Prickle2/Map3k7/Mapk10/Mapk8/Mapk9/Ror1/Ror2/Cul1/Csnk1e/Peg12/Cxxc4/Ep300/Znrf3/Sfrp5/Rbx1/Ruvbl1/Gsk3b/Fzd2/Ctnnbip1/Chd8/Bambi/Rspo3/Nfatc4/Cby1/Sost/Senp2/Daam2/Notum/Tbl1xr1/Csnk1a1/Vangl2/Nkd1 |
| mmu04919 | Thyroid hormone signaling pathway | 1,1935E-06 | Raf1/Actb/Actg1/Akt1/Akt2/Atp1a1/Atp1b1/Atp1b2/Atp1b3/Atp2a2/Bad/Casp9/Ctnnb1/Ccnd1/Crebbp/Dio1/Dio2/Esr1/Gata4/Kat2a/Hdac2/Hdac3/Hif1a/Hras/Itgav/Mdm2/Myc/Ncoa1/Ncoa2/Ncoa3/Notch1/Notch2/Notch3/Nras/Kat2b/Pdpk1/Pfkfb2/Pik3ca/Pik3cd/Pik3r1/Pik3r2/Pik3r3/Prkaca/Prkacb/Prkcb/Prkcg/Plcb1/Plcb2/Plcb3/Plcb4/Plcd1/Plcg1/Med1/Rheb/Rxra/Rxrg/Ncor1/Slc2a1/Slc9a1/Src/Stat1/Tbc1d4/Med16/Thra/Thrb/Wnt4/Plcg2/Med17/Akt3/Med24/Map2k1/Map2k2/Mapk1/Mapk3/Med13/Ep300/Med12l/Hdac1/Atp2a3/Rcan2/Rcan1/Pfkp/Foxo1/Gsk3b/Mtor/Med4/Med27/Med30/Slc16a10/Plce1/Pik3cb/Med13l |
| mmu05135 | Yersinia infection | 1,1935E-06 | Ticam1/Baiap2/Pkn2/Arhgef28/Actb/Actg1/Akt1/Akt2/Arf6/Cdc42/Chuk/Bcar1/Crk/Crkl/Ptk2/Fos/Ikbkb/Il10/Il18/Itga5/Itgb1/Jun/Lck/Limk1/Rac3/Nfatc1/Nfatc2/Nfatc3/Nfkb1/Nfkbia/Pik3ca/Pik3cd/Pik3r1/Pik3r2/Pik3r3/Pip5k1c/Pip5k1b/Pip5k1a/Plcg1/Mapk11/Ptk2b/Pxn/Rac1/Rela/Rock1/Rock2/Rps6ka1/Rps6ka2/Src/Wipf1/Tlr4/Traf2/Traf6/Vav1/Vav2/Zap70/Akt3/Fyb/Wasf2/Actr3b/Map2k1/Map2k2/Map2k3/Map2k4/Map2k6/Map2k7/Map3k7/Mapk1/Mapk10/Mapk13/Mapk14/Mapk3/Mapk8/Mapk9/Git2/Irak4/Mapk12/Pkn1/Dock1/Arhgef7/Irf3/Skap2/Tbk1/Gsk3b/Vav3/Tab1/Actr2/Pycard/Wipf2/Tab2/Arhgef12/Wasl/Actr3/Pik3cb |
| mmu00562 | Inositol phosphate metabolism | 1,6315E-06 | Inpp5f/Fig4/Synj1/Aldh6a1/Pi4kb/Impa2/Pip4k2c/Inpp1/Inpp5b/Inppl1/Mtmr4/Inpp5j/Minpp1/Pik3c2a/Pik3c2g/Pik3ca/Pik3cd/Pip5k1c/Pip4k2a/Pip5k1b/Pip5k1a/Plcb1/Plcb2/Plcb3/Plcb4/Plcd1/Plcg1/Inpp5k/Pten/Synj2/Inpp5a/Itpk1/Mtmr6/Tpi1/Pi4ka/Pik3c3/Pip5kl1/Itpka/Itpkc/Inpp4b/Plcg2/Pik3c2b/Bpnt2/Inpp4a/Plch1/Pik3cg/Itpkb/Mtmr7/Impa1/Inpp5e/Pi4k2b/Ipmk/Isyna1/Plce1/Mtmr3/Pik3cb/Ippk/Mtmr2/Sacm1l/Pi4k2a/Mtmr14 |
| mmu05231 | Choline metabolism in cancer | 2,1047E-06 | Slc44a1/Dgkz/Raf1/Dgkg/Dgkq/Akt1/Akt2/Chka/Pcyt1a/Dgka/Egf/Egfr/Eif4ebp1/Fos/Grb2/Hif1a/Hras/Jun/Rac3/Nras/Pdgfa/Pdgfb/Pdgfra/Pdgfrb/Pdpk1/Pik3ca/Pik3cd/Pik3r1/Pik3r2/Pik3r3/Pip5k1c/Pip5k1b/Pip5k1a/Prkcb/Prkcg/Lypla1/Pla2g4a/Plcg1/Pld1/Pld2/Plpp1/Rac1/Ralgds/Rheb/Slc22a5/Sos1/Sos2/Sp1/Chpt1/Slc44a3/Dgkb/Dgkd/Akt3/Wasf2/Map2k1/Map2k2/Mapk1/Mapk10/Mapk3/Mapk8/Mapk9/Slc22a4/Dgki/Pla2g4e/Dgkh/Plpp2/Pdgfc/Dgke/Slc22a21/Mtor/Rps6kb2/Tsc1/Plpp3/Slc44a2/Slc44a4/Pdgfd/Gpcpd1/Pik3cb |
| mmu04710 | Circadian rhythm | 2,7323E-06 | Fbxw11/Csnk1d/Prkaa1/Prkaa2/Prkab2/Prkag2/Arntl/Btrc/Clock/Creb1/Cry1/Cry2/Npas2/Per1/Per2/Per3/Prkab1/Prkag1/Rora/Rorc/Bhlhe40/Skp1/Nr1d1/Rorb/Cul1/Csnk1e/Fbxl3/Rbx1/Bhlhe41 |
| mmu05166 | Human T-cell leukemia virus 1 infection | 3,469E-06 | Trrap/Xpo1/Adcy3/Espl1/Cdc20/Fdps/H2-Q6/H2-Q9/Adcy6/Adcy8/Adcy9/Akt1/Akt2/Slc25a4/Atf2/Atf4/Atm/B2m/Bax/Bcl2l1/Bub1b/Bub3/Tspo/Calr/Canx/Ccna2/Ccnb2/Ccnd1/Ccnd2/Ccnd3/Ccne1/Ccne2/Cdk2/Cdk4/Cdkn1a/Cdkn2a/Cdkn2b/Cdkn2c/Chek1/Chuk/Creb1/Creb3/Crebbp/Atf6b/Dlg1/E2f1/E2f3/Egr1/Egr2/Elk4/Fos/Fosl1/Kat2a/H2-Ab1/H2-Bl/H2-D1/H2-Eb1/H2-K1/H2-M3/H2-DMa/H2-Q1/H2-Q10/H2-Q2/H2-Q4/H2-T22/H2-T23/H2-T24/Hras/Icam1/Ikbkb/Il15/Il15ra/Il1r1/Jak1/Jak3/Jun/Lck/Ltbr/Mad1l1/Smad2/Smad3/Smad4/Anapc1/Msx1/Msx2/Msx3/Myc/Nfatc1/Nfatc2/Nfatc3/Nfkb1/Nfkb2/Nfkbia/Nfyb/Nras/Kat2b/Pik3ca/Pik3cd/Pik3r1/Pik3r2/Pik3r3/Prkaca/Prkacb/Polb/Ppp3ca/Ppp3cb/Ppp3cc/Ppp3r1/Pten/Ran/Rasl2-9/Rb1/Rela/Relb/Spi1/Slc2a1/Stat5a/Stat5b/Creb3l2/Tcf3/Tert/Tgfb1/Tgfb2/Tgfb3/Tgfbr1/Tgfbr2/Tnfrsf1a/Cd40/Vdac1/Vdac2/Vdac3/Zfp36/Creb5/Vac14/Tbpl1/Akt3/Ets1/Ets2/E2f2/Atr/Map2k1/Map2k2/Map2k4/Map3k1/Map3k3/Mapk1/Mapk10/Mapk3/Mapk8/Mapk9/Creb3l1/Pttg1/Ep300/Crtc1/Anapc4/Cdc23/Map3k14/Mad2l1/Gps2/Anapc7/Anapc5/Cdc26/H2-T-ps/Gm8909/Cdc16/Crtc3/Tln2/Ranbp3/Nfatc4/Crtc2/Pik3cb/Creb3l4/Kat5 |
| mmu04146 | Peroxisome | 3,469E-06 | Pex12/Nudt19/Pecr/Acaa1a/Acox1/Cat/Ephx2/Acsl1/Gnpat/Hmgcl/Hsd17b4/Idh1/Phyh/Acot8/Amacr/Mpv17/Mvk/Nos2/Prdx1/Pex11a/Pex11b/Pex16/Pex7/Pipox/Pex19/Abcd3/Abcd4/Pex2/Pex5/Scp2/Slc25a17/Sod1/Hmgcll1/Paox/Acsl6/Xdh/Pex6/Agps/Mpv17l2/Acaa1b/Eci2/Decr2/Abcd2/Idh2/Dhrs4/Acsl5/Ech1/Prdx5/Hao2/Pex14/Pex3/Mlycd/Pxmp4/Pex10/Far1/Nudt7/Nudt12/Pmvk/Eci3/Pex11g/Ddo/Pex13/Pex26/Crot/Ehhadh/Gstk1/Acox3/Mpv17l |
| mmu05214 | Glioma | 3,8932E-06 | Ddb2/Camk2d/Braf/Raf1/Akt1/Akt2/Bak1/Bax/Calm1/Calm2/Calm3/Camk4/Ccnd1/Cdk4/Cdk6/Cdkn1a/Cdkn2a/Gadd45a/E2f1/E2f3/Egf/Egfr/Grb2/Hras/Igf1/Igf1r/Mdm2/Gadd45b/Nras/Pdgfa/Pdgfb/Pdgfra/Pdgfrb/Pik3ca/Pik3cd/Pik3r1/Pik3r2/Pik3r3/Prkcb/Prkcg/Plcg1/Pten/Rb1/Shc1/Shc3/Sos1/Sos2/Tgfa/Plcg2/Akt3/Gadd45g/E2f2/Map2k1/Map2k2/Mapk1/Mapk3/Polk/Shc4/Camk1/Mtor/Pik3cb |
| mmu05169 | Epstein-Barr virus infection | 4,1423E-06 | Isg15/Ddb2/H2-Q6/H2-Q9/Akt1/Akt2/Apaf1/B2m/Bak1/Bax/Bcl2/Bid/Bcl2l11/Calr/Casp3/Casp8/Casp9/Ccna2/Ccnd1/Ccnd2/Ccnd3/Ccne1/Ccne2/Entpd1/Cd247/Cd44/Cdk2/Cdk4/Cdk6/Cdkn1a/Cdkn1b/Chuk/Cycs/Gadd45a/E2f1/E2f3/Fadd/Fas/Pdia3/H2-Ab1/H2-Bl/H2-D1/H2-Eb1/H2-K1/H2-M3/H2-DMa/H2-Q1/H2-Q10/H2-Q2/H2-Q4/H2-T22/H2-T23/H2-T24/Hdac2/Hes1/Icam1/Cxcl10/Ifnar1/Ifnar2/Ikbkb/Irf9/Jak1/Jak3/Jun/Lyn/Mdm2/Psmd7/Myc/Gadd45b/Nedd4/Nfkb1/Nfkb2/Nfkbia/Nfkbib/Nfkbie/Pik3ca/Pik3cd/Pik3r1/Pik3r2/Pik3r3/Mapk11/Eif2ak2/Psmc1/Psmc2/Psmc3/Psmc5/Psmd4/Rac1/Rb1/Rbpj/Rela/Relb/Ripk1/Sem1/Ncor2/Stat1/Stat2/Stat3/Syk/Tap2/Tapbp/Entpd3/Psmd2/Tnfaip3/Cd40/Traf2/Traf3/Traf5/Traf6/Psmd3/Vim/Mavs/Ddx58/Ifna13/Plcg2/Akt3/Gadd45g/Psmc4/Psmd13/Tlr2/Ifna15/E2f2/Oas1a/Usp7/Map2k3/Map2k4/Map2k6/Map2k7/Map3k7/Mapk10/Mapk13/Mapk14/Mapk8/Mapk9/Irak4/Polk/Skp2/Mapk12/Hdac1/Map3k14/Irf7/Irf3/Tyk2/Adrm1/Tbk1/Ikbke/Psmd8/Psmd14/Sap30/Snw1/Psmd6/Tab1/H2-T-ps/Gm8909/Cir1/Psmd12/Psmc6/Tab2/Psmd11/Psmd1/Tradd/Pik3cb |
| mmu04931 | Insulin resistance | 5,9814E-06 | Acacb/Ppp1r3e/Prkaa1/Prkaa2/Prkab2/Prkag2/Pygb/Pygl/Ptpa/Akt1/Akt2/Cd36/Socs3/Cpt1a/Creb1/Creb3/G6pc/Gfpt1/Gfpt2/Gys1/Ikbkb/Insr/Irs1/Ppargc1b/Nfkb1/Nfkbia/Pdpk1/Pik3ca/Pik3cd/Pik3r1/Pik3r2/Pik3r3/Prkcb/Prkcd/Prkce/Prkcz/Ppara/Ppargc1a/Ppp1cb/Ppp1cc/Prkab1/Prkag1/Pten/Ptpn1/Ptpn11/Ptprf/Rela/Rps6ka1/Rps6ka2/Slc2a1/Slc2a2/Slc2a4/Srebf1/Mlxip/Stat3/Creb3l2/Tbc1d4/Mlx/Tnfrsf1a/Nr1h3/Nr1h2/Trib3/Ppp1r3d/Creb5/Akt3/Ppp1r3b/Mapk10/Mapk8/Mapk9/Creb3l1/Slc27a1/Slc27a4/Irs2/Ppp1r3c/Foxo1/Gsk3b/Mtor/Mlxipl/Rps6kb2/G6pc3/Crtc2/Pck2/Pik3cb/Oga/Creb3l4 |
| mmu05170 | Human immunodeficiency virus 1 infection | 6,2481E-06 | Fbxw11/Raf1/H2-Q6/H2-Q9/Akt1/Akt2/Ap1b1/Ap1g1/Ap1g2/Ap1m1/Ap1m2/Ap1s1/Atm/B2m/Bad/Bak1/Bax/Bcl2/Bcl2l1/Bid/Btrc/Calm1/Calm2/Calm3/Calr/Casp3/Casp8/Casp9/Ccnb2/Cd247/Cdc25c/Cdk1/Cfl1/Cfl2/Chek1/Chuk/Cxcr4/Crk/Crkl/Cycs/Ddb1/Fadd/Ptk2/Fas/Fos/Gna11/Gnai1/Gnai2/Gnai3/Gnao1/Gnaq/Gnb1/Gnb4/Gnb5/Gng10/Gng12/Gng2/Gng5/Gng7/Gng8/Gngt2/Pdia3/H2-Bl/H2-D1/H2-K1/H2-M3/H2-Q1/H2-Q10/H2-Q2/H2-Q4/H2-T22/H2-T23/H2-T24/Hras/Ikbkb/Itpr1/Itpr2/Itpr3/Jun/Limk1/Limk2/Rac3/Nfatc1/Nfatc2/Nfatc3/Nfkb1/Nfkbia/Nras/Pak1/Pik3ca/Pik3cd/Pik3r1/Pik3r2/Pik3r3/Prkcb/Prkcg/Plcg1/Ppp3ca/Ppp3cb/Ppp3cc/Ppp3r1/Mapk11/Ptk2b/Pxn/Rac1/Rela/Ripk1/Rnf7/Trim30d/Tap2/Tapbp/Skp1/Pak6/Cgas/Tlr4/Tnfrsf1a/Tnfrsf1b/Traf2/Traf5/Traf6/Wee1/Pak2/Ifna13/Plcg2/Akt3/Tlr2/Ifna15/Atr/Ap1s3/Map2k1/Map2k2/Map2k3/Map2k6/Map2k7/Map3k7/Mapk1/Mapk10/Mapk13/Mapk14/Mapk3/Mapk8/Mapk9/Irak4/Ccnb1/Cul1/Mapk12/Trim12c/Dcaf1/Irf3/Samhd1/Rbx1/Tbk1/Mtor/Rps6kb2/Gng11/Tab1/H2-T-ps/Gm8909/Elob/Tab2/Bst2/Tradd/Sting1/Nfatc4/Pik3cb/Cul5/Apobec3 |
| mmu04142 | Lysosome | 7,3481E-06 | Gga1/Ap4e1/Atp6v1h/Gusb/Manba/Abca2/Laptm4b/Atp6v0b/Acp2/Acp5/Aga/Ap1b1/Ap1g1/Ap1g2/Ap1m1/Ap1m2/Ap1s1/Ap3b1/Ap3b2/Ap3d1/Ap3s1/Ap3s2/Ap4m1/Ap4s1/Arsb/Asah1/Atp6v0d1/Atp6v0a1/Atp6v0c/Scarb2/Cd63/Tpp1/Cln3/Clta/Ctsb/Ctsc/Ctsd/Ctsh/Ctsl/Dnase2a/Gaa/Galc/Gba/Gm2a/Hexa/Hexb/Hyal1/Igf2r/Lamp1/Lipa/M6pr/Man2b1/Laptm4a/Naga/Neu1/Npc1/Slc11a1/Slc11a2/Ctsa/Ppt1/Lgmn/Psap/Pla2g15/Smpd1/Sort1/Cln5/Gnptg/Atp6v0a2/Ctso/Slc17a5/Ap1s3/Gga3/Tcirg1/Naglu/Nagpa/Gnptab/Galns/Hgsnat/Cd164/Ppt2/Abcb9/Ctsf/Litaf/Sumf1/Ctsz/Ap3m2/Cltc/Npc2/Fuca1/Arsg/Gga2/Cltb/Gns/Mcoln1 |
| mmu04722 | Neurotrophin signaling pathway | 7,4346E-06 | Rapgef1/Camk2d/Irak2/Braf/Rap1a/Raf1/Abl1/Akt1/Akt2/Atf4/Bad/Bax/Bcl2/Calm1/Calm2/Calm3/Camk4/Cdc42/Crk/Crkl/Gab1/Arhgdig/Grb2/Hras/Ikbkb/Irs1/Jun/Sh2b3/Zfp369/Mapkapk2/Nfkb1/Nfkbia/Nfkbib/Nfkbie/Nras/Pdpk1/Pik3ca/Pik3cd/Pik3r1/Pik3r2/Pik3r3/Prkcd/Plcg1/Mapk11/Psen1/Psen2/Ptpn11/Arhgdia/Rac1/Rela/Rps6ka1/Rps6ka2/Sh2b1/Shc1/Shc3/Sort1/Sos1/Sos2/Rap1b/Traf6/Trp73/Ywhae/Plcg2/Akt3/Sh2b2/Map2k5/Mapk7/Map2k1/Map2k2/Map2k7/Map3k1/Map3k3/Map3k5/Mapk1/Mapk10/Mapk13/Mapk14/Mapk3/Mapk8/Mapk9/Irak4/Shc4/Mapk12/Frs2/Foxo3/Gsk3b/Zfp110/Prdm4/Rps6ka5/Pik3cb/Kidins220/Ntf5 |
| mmu05160 | Hepatitis C | 9,5991E-06 | Ticam1/Braf/Raf1/Akt1/Akt2/Apaf1/Bad/Bak1/Bax/Bid/Casp3/Casp8/Casp9/Ctnnb1/Ccnd1/Cd81/Cdk2/Cdk4/Cdk6/Cdkn1a/Cflar/Chuk/Socs3/Cldn1/Cldn3/Cycs/E2f1/E2f3/Egf/Egfr/Eif2s1/Eif2ak3/Fadd/Fas/Tlr3/Grb2/Hras/Eif2ak1/Cxcl10/Ifnar1/Ifnar2/Ikbkb/Eif3e/Irf9/Jak1/Ldlr/Mx2/Myc/Nfkb1/Nfkbia/Nras/Ocln/Cldn11/Pik3ca/Pik3cd/Pik3r1/Pik3r2/Pik3r3/Ppara/Ppp2ca/Ppp2cb/Eif2ak2/Psme3/Rb1/Rela/Ripk1/Rxra/Sos1/Sos2/Scarb1/Stat1/Stat2/Stat3/Tnfrsf1a/Traf2/Traf3/Traf6/Nr1h3/Ywhae/Ywhag/Ywhah/Ywhaq/Ywhaz/Mavs/Ddx58/Ifna13/Akt3/Rnasel/Ifna15/E2f2/Oas1a/Map2k1/Map2k2/Mapk1/Mapk3/Eif2ak4/Ppp2r1a/Ppp2r2d/Cldn7/Irf7/Irf3/Ywhab/Cldn6/Cldn8/Tyk2/Pias1/Tbk1/Ikbke/Gsk3b/Cldn9/Rsad2/Cldn10/Tradd/Cldn23/Ppp2r2a/Ppp2r1b/Pik3cb |
| mmu04340 | Hedgehog signaling pathway | 1,3391E-05 | Csnk1g2/Fbxw11/Csnk1d/Arrb1/Grk2/Bcl2/Btrc/Ccnd1/Ccnd2/Gas1/Gli1/Gli2/Lrp2/Hhip/Ihh/Kif3a/Kif7/Mgrn1/Prkaca/Prkacb/Ptch1/Ptch2/Shh/Spop/Csnk1g1/Arrb2/Mosmo/Sufu/Cul3/Cul1/Megf8/Csnk1e/Smo/Grk3/Gsk3b/Cdon/Smurf2/Csnk1g3/Smurf1/Spopl/Csnk1a1 |
| mmu04330 | Notch signaling pathway | 1,3542E-05 | Maml1/Adam17/Crebbp/Ctbp1/Ctbp2/Dll1/Dll3/Dvl1/Dvl2/Dvl3/Dtx1/Kat2a/Hdac2/Hes1/Hes5/Hey1/Hey2/Jag1/Lfng/Notch1/Notch2/Notch3/Numb/Numbl/Kat2b/Psen1/Psen2/Rbpj/Rfng/Atxn1/Ncor2/Dtx4/Aph1b/Dtx3l/Aph1a/Maml2/Ep300/Maml3/Hdac1/Atxn1l/Dll4/Heyl/Snw1/Cir1/Aph1c/Dtx2 |
| mmu04932 | Non-alcoholic fatty liver disease (NAFLD) | 1,4779E-05 | Prkaa1/Prkaa2/Prkab2/Prkag2/Cox6b1/Akt1/Akt2/Atf4/Bax/Bid/Bcl2l11/Casp3/Casp7/Casp8/Cdc42/Cebpa/Socs3/Cox5a/Cox5b/Cox6a1/Cox6c/Cox7a1/Cox7a2/Cox7c/Cox8a/Cycs/Eif2s1/Eif2ak3/Fas/Ikbkb/Il6ra/Insr/Irs1/Itch/Jun/Lepr/Ndufa2/Ndufa4/Ndufs4/Ndufv1/Nfkb1/Pik3ca/Pik3cd/Pik3r1/Pik3r2/Pik3r3/Pklr/Ppara/Prkab1/Prkag1/Rac1/Rela/Rxra/Cox7a2l/Srebf1/Mlxip/Mlx/Tgfb1/Tnfrsf1a/Traf2/Nr1h3/Uqcrc1/Xbp1/Ndufs8/Ndufs2/Ndufs1/Ndufb6/Akt3/Map3k5/Mapk10/Mapk8/Mapk9/Irs2/Ndufs6/Ndufa4l2/Gsk3b/Mlxipl/Ndufs5/Gsk3a/Ndufb5/Ndufa3/Ndufa9/Uqcr10/Ndufb9/Ndufc1/Ndufa12/Ndufa7/Cyc1/Uqcr11/Uqcrfs1/Ndufb7/Sdhd/Sdha/Uqcrc2/Ndufa6/Ndufb8/Ndufa10/Sdhb/Ndufb4/Ndufc2/Ndufb2/Ndufa5/Ndufa8/Adipor2/Ndufab1/Adipor1/Ndufv2/Pik3cb/Ndufs7/Ern1 |
| mmu04350 | TGF-beta signaling pathway | 1,5245E-05 | E2f4/Amhr2/Acvr1/Acvr1b/Acvr2a/Acvr2b/Amh/Bmp2/Bmp6/Bmp7/Bmp8a/Bmpr1a/Bmpr1b/Bmpr2/Cdkn2b/Chrd/Crebbp/Dcn/E2f5/Lefty1/Fst/Id1/Id2/Id3/Id4/Inhba/Inhbb/Smad1/Smad2/Smad3/Smad4/Smad5/Smad6/Myc/Nbl1/Neo1/Pitx2/Ppp2ca/Ppp2cb/Rbl1/Rock1/Sp1/Thsd4/Skp1/Tfdp1/Tgfb1/Tgfb2/Tgfb3/Tgfbr1/Tgfbr2/Tgif1/Thbs1/Zfyve16/Tgif2/Zfyve9/Gdf6/Rgma/Mapk1/Mapk3/Ltbp1/Acvr1c/Cul1/Lefty2/Ep300/Ppp2r1a/Smad9/Rbx1/Rps6kb2/Smurf2/Hamp2/Bambi/Rgmb/Ppp2r1b/Smurf1 |
| mmu03050 | Proteasome | 2,0025E-05 | Psme4/Psmb9/Psmb8/Psmd7/Psma2/Psma3/Psmb1/Psmb4/Psmb5/Psmb6/Psmb7/Psmc1/Psmc2/Psmc3/Psmc5/Psmd4/Psme1/Psme2/Psme3/Sem1/Psmd2/Psmd3/Psmf1/Psmc4/Psmd13/Psma1/Psma5/Psma6/Psma7/Psmb2/Psmb3/Adrm1/Psmd8/Psmd14/Psmd6/Pomp/Psmd12/Psmc6/Psmd11/Psmd1 |
| mmu03018 | RNA degradation | 2,0032E-05 | Pan2/Cnot6/Skiv2l/Exosc4/Btg1/Btg2/Btg3/Ddx6/Eno1/Eno2/Hspd1/Hspa9/Pabpc1/Pfkl/Pfkm/Cnot7/Tent4a/Dis3l/Tent4b/Ttc37/Tob1/Patl1/Eno4/Exosc2/Pabpc4/Cnot6l/Cnot3/Cnot1/Edc4/Xrn1/Xrn2/Pabpc4l/Lsm2/Dcp1b/Zcchc7/Edc3/Lsm4/Exosc9/Exosc10/Cnot4/Pfkp/Tob2/C1d/Lsm7/Wdr61/Exosc3/Lsm5/Exosc7/Exosc1/Lsm1/Mphosph6/Cnot8/Dcps/Dcp2/Pnpt1/Cnot2/Dhx36/Exosc6/Pan3/Parn/Nudt16/Dcp1a/Lsm8/Lsm6/Cnot10 |
| mmu05235 | PD-L1 expression and PD-1 checkpoint pathway in cancer | 2,0568E-05 | Ticam1/Raf1/Akt1/Akt2/Tirap/Cd247/Chuk/Csnk2a1/Csnk2a2/Csnk2b/Egf/Egfr/Fos/Ptpn6/Hif1a/Hras/Ifngr1/Ifngr2/Ikbkb/Jak1/Jak2/Jun/Lck/Nfatc1/Nfatc2/Nfatc3/Nfkb1/Nfkbia/Nfkbib/Nfkbie/Nras/Pik3ca/Pik3cd/Pik3r1/Pik3r2/Pik3r3/Plcg1/Ppp3ca/Ppp3cb/Ppp3cc/Ppp3r1/Mapk11/Pten/Ptpn11/Rela/Stat1/Stat3/Tlr4/Traf6/Ticam2/Zap70/Akt3/Tlr2/Map2k1/Map2k2/Map2k3/Map2k6/Map3k3/Mapk1/Mapk13/Mapk14/Mapk3/Mapk12/Batf/Mtor/Rps6kb2/Cd274/Pik3cb/Eml4 |
| mmu05145 | Toxoplasmosis | 2,6575E-05 | Ppif/Akt1/Akt2/Alox5/Birc3/Birc2/Bad/Bcl2/Bcl2l1/Ciita/Casp3/Casp8/Casp9/Chuk/Socs1/Cycs/Gnai1/Gnai2/Gnai3/Gnao1/H2-Ab1/H2-Eb1/H2-DMa/Hspa1l/Hspa1b/Hspa2/Irgm1/Ifngr1/Ifngr2/Igtp/Ikbkb/Il10/Il10rb/Itga6/Itgb1/Jak1/Jak2/Lama1/Lama3/Lama4/Lama5/Lamb2/Lamb3/Lamc2/Ldlr/Ly96/Nfkb1/Nfkbia/Nfkbib/Nos2/Pdpk1/Mapk11/Rela/Stat1/Stat3/Tgfb1/Tgfb2/Tgfb3/Tlr4/Tnfrsf1a/Cd40/Traf6/Lamc1/Akt3/Tlr2/Map2k3/Map2k6/Map3k7/Mapk1/Mapk10/Mapk13/Mapk14/Mapk3/Mapk8/Mapk9/Irak4/Mapk12/Pik3cg/Pik3r5/Tyk2/Tab1/Tab2 |
| mmu04211 | Longevity regulating pathway | 2,8803E-05 | Adcy3/Prkaa1/Prkaa2/Prkab2/Prkag2/Ehmt2/Adcy6/Adcy8/Adcy9/Akt1/Akt2/Atg5/Atf2/Atf4/Bax/Camk4/Cat/Rb1cc1/Creb1/Creb3/Atf6b/Eif4ebp1/Sesn1/Hras/Igf1/Igf1r/Insr/Irs1/Irs3/Nfkb1/Nras/Pik3ca/Pik3cd/Pik3r1/Pik3r2/Pik3r3/Prkaca/Prkacb/Pparg/Ppargc1a/Prkab1/Prkag1/Rela/Rheb/Camkk2/Creb3l2/Stk11/Ulk1/Sesn2/Creb5/Akt3/Creb3l1/Eif4e2/Irs2/Foxo1/Foxo3/Mtor/Rps6kb2/Tsc1/Akt1s1/Atg101/Adipor2/Adipor1/Appl1/Rptor/Pik3cb/Sesn3/Ehmt1/Creb3l4/Sirt1 |
| mmu04540 | Gap junction | 4,0507E-05 | Adcy3/Csnk1d/Raf1/Adcy6/Adcy8/Adcy9/Adrb1/Cdk1/Egf/Egfr/Gja1/Gna11/Gnai1/Gnai2/Gnai3/Gnaq/Gnas/Lpar1/Grb2/Hras/Htr2a/Itpr1/Itpr2/Itpr3/Nras/Pdgfa/Pdgfb/Pdgfra/Pdgfrb/Prkaca/Prkacb/Prkcb/Prkcg/Plcb1/Plcb2/Plcb3/Plcb4/Prkg1/Sos1/Sos2/Src/Tjp1/Tuba1a/Tuba1b/Tuba3a/Tuba4a/Tuba1c/Tubb2a/Tubb3/Tubb5/Tubb4b/Gucy1a2/Tubal3/Map2k5/Mapk7/Map2k1/Map2k2/Map3k2/Mapk1/Mapk3/Tuba8/Gucy1b1/Pdgfc/Gucy1a1/Tubb6/Pdgfd/Tubb2b |
| mmu00310 | Lysine degradation | 4,2474E-05 | Nsd2/Ehmt2/Acat1/Acat2/Prdm2/Aldh7a1/Aldh2/Aldh3a2/Dld/Ezh1/Ezh2/Hadh/Nsd1/Plod1/Pipox/Ash1l/Setd1b/Dot1l/Dhtkd1/Prdm9/Kmt2a/Kmt5b/Smyd2/Kmt2c/Kmt5c/Setd1a/Nsd3/Colgalt1/Hykk/Setd2/Setdb2/Plod2/Plod3/Colgalt2/Gcdh/Aass/Kmt2d/Aldh9a1/Suv39h2/Kmt5a/Kmt2e/Smyd3/Aldh1b1/Phykpl/Setd7/Ehhadh/Setmar/Ehmt1/Dlst/Setdb1 |
| mmu04934 | Cushing syndrome | 5,1511E-05 | Adcy3/Camk2d/Orai1/Braf/Rap1a/Adcy6/Adcy8/Adcy9/Agtr1a/Ahr/Aip/Apc/Arnt/Atf2/Atf4/Axin1/Axin2/Cacna1d/Cacna1g/Ctnnb1/Ccnd1/Ccne1/Ccne2/Cdk2/Cdk4/Cdk6/Cdkn1a/Cdkn1b/Cdkn2a/Cdkn2b/Cdkn2c/Creb1/Creb3/Atf6b/Crhr2/Cyp11a1/Dvl1/Dvl2/Dvl3/E2f1/E2f3/Egfr/Wdr5/Fh1/Fzd1/Fzd3/Fzd4/Fzd5/Fzd6/Fzd7/Fzd8/Fzd9/Gna11/Gnai1/Gnai2/Gnai3/Gnaq/Gnas/Nr4a1/Itpr1/Itpr2/Itpr3/Ldlr/Lef1/Men1/Pbx1/Pde8a/Prkaca/Prkacb/Plcb1/Plcb2/Plcb3/Plcb4/Pomc/Rasd1/Rb1/Sp1/Scarb1/Creb3l2/Rbbp5/Tcf7/Tcf7l1/Tcf7l2/Kmt2a/Rap1b/Wnt9a/Pde8b/Wnt1/Wnt10b/Wnt11/Wnt9b/Wnt2/Wnt2b/Wnt4/Wnt5a/Wnt5b/Wnt6/Creb5/Apc2/Ash2l/E2f2/Map2k1/Map2k2/Mapk1/Mapk3/Nr5a1/Creb3l1/Nceh1/Kmt2d/Gsk3b/Fzd2/Wdr5b/Creb3l4/Usp8 |
| mmu05217 | Basal cell carcinoma | 6,6848E-05 | Ddb2/Apc/Axin1/Axin2/Bak1/Bax/Bmp2/Ctnnb1/Cdkn1a/Gadd45a/Dvl1/Dvl2/Dvl3/Fzd1/Fzd3/Fzd4/Fzd5/Fzd6/Fzd7/Fzd8/Fzd9/Gli1/Gli2/Hhip/Kif7/Lef1/Gadd45b/Ptch1/Ptch2/Shh/Tcf7/Tcf7l1/Tcf7l2/Wnt9a/Wnt1/Wnt10b/Wnt11/Wnt9b/Wnt2/Wnt2b/Wnt4/Wnt5a/Wnt5b/Wnt6/Apc2/Gadd45g/Sufu/Polk/Smo/Gsk3b/Fzd2 |
| mmu04114 | Oocyte meiosis | 7,0666E-05 | Fbxw11/Adcy3/Espl1/Cdc20/Camk2d/Adcy6/Adcy8/Adcy9/Btrc/Bub1/Calm1/Calm2/Calm3/Ccnb2/Ccne1/Ccne2/Cdc25c/Cdk1/Cdk2/Cpeb1/Smc3/Igf1/Igf1r/Itpr1/Itpr2/Itpr3/Mad1l1/Anapc1/Mos/Prkaca/Prkacb/Plk1/Ppp1cb/Ppp1cc/Ppp2ca/Ppp2cb/Ppp3ca/Ppp3cb/Ppp3cc/Ppp3r1/Mapk11/Rps6ka1/Rps6ka2/Slk/Aurka/Cpeb3/Skp1/Ppp2r5d/Ppp2r5b/Ywhae/Ywhag/Ywhah/Ywhaq/Ywhaz/Ppp2r5a/Cpeb2/Map2k1/Mapk1/Mapk13/Mapk14/Mapk3/Ccnb1/Pkmyt1/Ppp2r5c/Ppp2r5e/Cul1/Mapk12/Pttg1/Spdye4b/Stag3/Ppp2r1a/Anapc4/Cdc23/Ywhab/Mad2l1/Anapc7/Rbx1/Rec8/Anapc5/Cdc26/Fbxo5/Cpeb4/Cdc16/Spdya/Mad2l2/Sgo1/Ppp2r1b/Fbxo43 |
| mmu05017 | Spinocerebellar ataxia | 7,1348E-05 | Atg14/Atxn3/Afg3l1/Akt1/Akt2/Atp2a2/Cacna1a/Rb1cc1/Dab1/Fgf14/Gnaq/Grin2d/Itpr1/Itpr2/Itpr3/Kcnc3/Pik3ca/Pik3cd/Pik3r1/Pik3r2/Pik3r3/Prkcb/Prkcg/Plcb1/Plcb2/Plcb3/Plcb4/Rbpj/Rora/Atxn1/Atxn2/Slc1a6/Sp1/Sptbn2/Traf2/Ulk1/Vldlr/Xbp1/Pik3c3/Twnk/Ambra1/Gtf2b/Atxn2l/Tbpl1/Akt3/Grin3a/Map3k5/Mapk10/Mapk8/Mapk9/Ulk2/Atg2a/Atxn1l/Wipi1/Atp2a3/Atxn10/Becn1/Kcnd3/Mtor/Nrbf2/Bean1/Oma1/Atg101/Afg3l2/Cic/Opa1/Pik3cb/Wipi2/Pik3r4/Ern1/Pum1/Pum2/Kat5 |
| mmu04917 | Prolactin signaling pathway | 7,1931E-05 | Gck/Raf1/Akt1/Akt2/Ccnd1/Ccnd2/Cga/Cish/Socs3/Socs1/Esr1/Esr2/Fos/Grb2/Hras/Irf1/Jak2/Nfkb1/Nras/Pik3ca/Pik3cd/Pik3r1/Pik3r2/Pik3r3/Mapk11/Prlr/Socs7/Rela/Shc1/Shc3/Slc2a2/Sos1/Sos2/Src/Stat1/Stat3/Stat5a/Stat5b/Socs2/Tnfrsf11a/Akt3/Map2k1/Map2k2/Mapk1/Mapk10/Mapk13/Mapk14/Mapk3/Mapk8/Mapk9/Shc4/Mapk12/Socs6/Socs5/Foxo3/Gsk3b/Pik3cb |
| mmu05218 | Melanoma | 7,1931E-05 | Ddb2/Braf/Raf1/Akt1/Akt2/Bad/Bak1/Bax/Ccnd1/Cdh1/Cdk4/Cdk6/Cdkn1a/Cdkn2a/Gadd45a/E2f1/E2f3/Egf/Egfr/Fgf1/Fgf15/Fgf18/Fgf2/Fgf5/Fgf8/Fgf9/Fgfr1/Hras/Igf1/Igf1r/Mdm2/Met/Mitf/Gadd45b/Nras/Pdgfa/Pdgfb/Pdgfra/Pdgfrb/Pik3ca/Pik3cd/Pik3r1/Pik3r2/Pik3r3/Pten/Rb1/Akt3/Gadd45g/E2f2/Map2k1/Map2k2/Mapk1/Mapk3/Polk/Pdgfc/Pdgfd/Pik3cb |
| mmu04914 | Progesterone-mediated oocyte maturation | 7,1931E-05 | Adcy3/Braf/Kif22/Raf1/Adcy6/Adcy8/Adcy9/Akt1/Akt2/Bub1/Ccna2/Ccnb2/Cdc25a/Cdc25b/Cdc25c/Cdk1/Cdk2/Cpeb1/Gnai1/Gnai2/Gnai3/Hsp90ab1/Hsp90aa1/Igf1/Igf1r/Mad1l1/Anapc1/Mos/Pde3b/Pik3ca/Pik3cd/Pik3r1/Pik3r2/Pik3r3/Prkaca/Prkacb/Plk1/Mapk11/Rps6ka1/Rps6ka2/Stk10/Aurka/Cpeb3/Cpeb2/Akt3/Map2k1/Mapk1/Mapk10/Mapk13/Mapk14/Mapk3/Mapk8/Mapk9/Ccnb1/Pkmyt1/Mapk12/Spdye4b/Anapc4/Cdc23/Mad2l1/Anapc7/Fzr1/Anapc5/Cdc26/Cpeb4/Cdc16/Spdya/Mad2l2/Pik3cb |
| mmu04810 | Regulation of actin cytoskeleton | 7,4106E-05 | Brk1/Itga9/Mylk/Baiap2/Actn1/Braf/Raf1/Actb/Actg1/Pip4k2c/Apc/Arpc1b/Bdkrb2/Cdc42/Cfl1/Cfl2/Chrm1/Chrm3/Cxcr4/Bcar1/Crk/Crkl/Diaph1/Egf/Egfr/Enah/F2/F2r/Ptk2/Fgf1/Fgf15/Fgf18/Fgf2/Fgf5/Fgf8/Fgf9/Fgfr1/Fgfr2/Fgfr3/Fgfr4/Gna12/Gng12/Lpar1/Hras/Itga2/Itga2b/Itga3/Itga5/Itga6/Itga7/Itgam/Itgav/Itgb1/Itgb5/Itgb6/Itgb7/Limk1/Limk2/Rac3/Mos/Mras/Myh9/Mylpf/Ppp1r12a/Nras/Pak1/Pdgfa/Pdgfb/Pdgfra/Pdgfrb/Pfn1/Pfn2/Pik3ca/Pik3cd/Pik3r1/Pik3r2/Pik3r3/Pip5k1c/Pip4k2a/Pip5k1b/Pip5k1a/Ppp1cb/Ppp1cc/Itgb4/Pxn/Rac1/Rdx/Rock1/Rock2/Rras/Scin/Cxcl12/Cyfip1/Slc9a1/Sos1/Sos2/Src/Pak6/Git1/Tiam1/Spata13/Vav1/Vav2/Vcl/Ezr/Pak2/Arhgef4/Gsn/Ssh1/Ppp1r12c/Ssh2/Apc2/Insrr/Itga8/Wasf2/Chrm2/Ssh3/Map2k1/Map2k2/Mapk1/Mapk3/Iqgap1/Itgb8/Abi2/Ppp1r12b/Dock1/Itgad/Kng2/Iqgap3/Nckap1/Lpar2/Arhgef7/Iqgap2/Pdgfc/Arpc3/Diaph3/Vav3/Actn4/Rras2/Myl12a/Myl12b/Arhgef12/Pdgfd/Myh14/Wasl/Pik3cb/Arpc2/Cyfip2/Myh10 |
| mmu05167 | Kaposi sarcoma-associated herpesvirus infection | 7,4106E-05 | Atg14/Ticam1/Raf1/H2-Q6/H2-Q9/Angpt2/Akt1/Akt2/Bak1/Bax/Bid/Calm1/Calm2/Calm3/Casp3/Casp8/Casp9/Ctnnb1/Ccnd1/Cdk4/Cdk6/Cdkn1a/Chuk/Creb1/Crebbp/Cycs/E2f1/E2f3/Fadd/Fas/Fgf2/Fos/Tlr3/Gnb1/Gnb4/Gnb5/Gng10/Gng12/Gng2/Gng5/Gng7/Gng8/Gngt2/Cxcl1/H2-Bl/H2-D1/H2-K1/H2-M3/H2-Q1/H2-Q10/H2-Q2/H2-Q4/H2-T22/H2-T23/H2-T24/Hck/Hif1a/Hras/Icam1/Ifnar1/Ifnar2/Ifngr1/Ikbkb/Il6st/Irf9/Itpr1/Itpr2/Itpr3/Jak1/Jak2/Jun/Lyn/Mapkapk2/Myc/Nfatc1/Nfatc2/Nfatc3/Nfkb1/Nfkbia/Nras/Pdgfb/Pik3ca/Pik3cd/Pik3r1/Pik3r2/Pik3r3/Plcg1/Ppp3ca/Ppp3cb/Ppp3cc/Ppp3r1/Mapk11/Eif2ak2/Ptgs2/Rac1/Rb1/Rela/Src/Stat1/Stat2/Stat3/Syk/Tnfrsf1a/Traf2/Traf3/Ubb/Vegfa/Pik3c3/Zfp36/Ifna13/Plcg2/Akt3/Ifna15/E2f2/Map2k1/Map2k2/Map2k4/Map2k6/Map2k7/Mapk1/Mapk10/Mapk13/Mapk14/Mapk3/Mapk8/Mapk9/Prex1/Mapk12/Pik3cg/Pik3r5/Ep300/Irf7/Irf3/Rcan1/Tyk2/Becn1/Tbk1/Gabarap/Ikbke/Gsk3b/Mtor/Gng11/H2-T-ps/Gm8909/Atg3/Tradd/Nfatc4/Pik3cb/Gabarapl2 |
| mmu05163 | Human cytomegalovirus infection | 7,4842E-05 | Adcy3/Raf1/H2-Q6/H2-Q9/Adcy6/Adcy8/Adcy9/Akt1/Akt2/Atf2/Atf4/B2m/Bak1/Bax/Bid/Calm1/Calm2/Calm3/Calr/Casp3/Casp8/Casp9/Ctnnb1/Ccnd1/Cdk4/Cdk6/Cdkn1a/Cdkn2a/Chuk/Cxcr4/Creb1/Creb3/Atf6b/Bcar1/Crk/Crkl/Cycs/E2f1/E2f3/Egfr/Eif4ebp1/Fadd/Ptk2/Fas/Gna11/Gna12/Gnai1/Gnai2/Gnai3/Gnao1/Gnaq/Gnas/Gnb1/Gnb4/Gnb5/Gng10/Gng12/Gng2/Gng5/Gng7/Gng8/Gngt2/Grb2/Pdia3/H2-Bl/H2-D1/H2-K1/H2-M3/H2-Q1/H2-Q10/H2-Q2/H2-Q4/H2-T22/H2-T23/H2-T24/Hras/Ikbkb/Il10rb/Il1r1/Il6ra/Itgav/Itpr1/Itpr2/Itpr3/Jak1/Rac3/Mdm2/Myc/Nfatc1/Nfatc2/Nfatc3/Nfkb1/Nfkbia/Nras/Pdgfra/Pik3ca/Pik3cd/Pik3r1/Pik3r2/Pik3r3/Prkaca/Prkacb/Prkcb/Prkcg/Plcb1/Plcb2/Plcb3/Plcb4/Ppp3ca/Ppp3cb/Ppp3cc/Ppp3r1/Mapk11/Ptger2/Ptger3/Ptger4/Ptgs2/Ptk2b/Pxn/Rac1/Rb1/Rela/Rheb/Ripk1/Rock1/Rock2/Ccl5/Cx3cl1/Cxcl12/Sos1/Sos2/Sp1/Src/Stat3/Creb3l2/Arhgef11/Tap2/Tapbp/Cgas/Tnfrsf1a/Traf2/Traf5/Vegfa/Ifna13/Creb5/Akt3/Ifna15/E2f2/Map2k1/Map2k2/Map2k6/Mapk1/Mapk13/Mapk14/Mapk3/Creb3l1/Mapk12/Irf3/Tbk1/Gsk3b/Mtor/Rps6kb2/Tsc1/Gng11/H2-T-ps/Gm8909/Arhgef12/Tradd/Sting1/Nfatc4/Pik3cb/Akap13/Creb3l4 |
| mmu05202 | Transcriptional misregulation in cancer | 7,6488E-05 | Etv5/Nsd2/Ddb2/Jmjd1c/Supt3/Cebpe/Birc3/Atf1/Atm/Bak1/Bax/Bcl2l1/Bcl6/Bmi1/Runx2/Runx1/Runx1t1/Ccna2/Ccnd2/Cd14/Cdkn1a/Cdkn1b/Cdkn2c/Cebpa/Cebpb/Gadd45a/Ddx5/Elk4/Etv1/Etv6/Ewsr1/Eya1/Bmp2k/Ptk2/Fcgr1/Fli1/H3c14/H3f3a/H3f3b/Hdac2/Hhex/Hmga2/Hpgd/Id2/Igf1/Igf1r/Itgam/Itgb7/Jup/Klf3/Ldb1/Lmo2/Smad1/Maf/Max/Mdm2/Mef2c/Meis1/Men1/Met/Mitf/Mlf1/Aff1/Mmp9/Myc/Gadd45b/Nfkb1/Mycn/Nr4a3/Pax3/Pbx1/Pbx3/Pdgfa/Etv4/Per2/Cdk14/Plat/Plau/Pml/Pparg/Prom1/Rara/Rel/Rela/Rxra/Rxrg/Ncor1/Spi1/Six1/Six4/Sp1/Spint1/Dot1l/Slc45a3/Kmt2a/Zeb1/Tcf3/Tgfbr2/Tlx1/Cd40/Traf1/Wt1/Zbtb17/Fus/Zbtb16/Gadd45g/Fev/H3c7/Ss18/Polk/H3c3/H3c4/H3c2/H3c6/H3c10/H3c11/H3c13/Hdac1/Tmprss2/Fut8/Nupr1/Foxo1/Mllt1/Dusp6/Aspscr1/Mllt3/Taf15/Nfkbiz/Prcc |
| mmu04928 | Parathyroid hormone synthesis, secretion and action | 7,6941E-05 | Adcy3/Arrb1/Braf/Raf1/Pde4c/Adcy6/Adcy8/Adcy9/Atf2/Atf4/Bcl2/Bglap2/Runx2/Cdkn1a/Creb1/Creb3/Atf6b/Cyp27b1/Egfr/Egr1/Fgfr1/Slc34a3/Fos/Gata3/Gcm1/Gna11/Gna12/Gnai1/Gnai2/Gnai3/Gnaq/Gnas/Hbegf/Itpr1/Itpr2/Itpr3/Jund/Mafb/Lrp5/Lrp6/Mef2a/Mef2c/Mef2d/Mmp14/Mmp15/Mmp16/Naca/Nr4a2/Pde4a/Pde4b/Prkaca/Prkacb/Prkcb/Prkcg/Plcb1/Plcb2/Plcb3/Plcb4/Pld1/Pld2/Pthlh/Rxra/Rxrg/Sp1/Creb3l2/Arhgef11/Arrb2/Vdr/Creb5/Pde4d/Mmp17/Mmp25/Map2k1/Mapk1/Mapk3/Creb3l1/Slc9a3r1/Sost/Akap13/Creb3l4 |
| mmu04714 | Thermogenesis | 0,00010455 | Adcy3/Kdm3a/Prkaa1/Prkaa2/Prkab2/Prkag2/Cox6b1/Actb/Actg1/Adcy6/Adcy8/Adcy9/Atf2/Atp5a1/Atp5b/Atp5c1/Atp5pb/Atp5g1/Atp5j/Atp5k/Bmp8a/Cnr1/Cox17/Cox5a/Cox5b/Cox6a1/Cox6c/Cox7a1/Cox7a2/Cox7c/Cox8a/Cpt1a/Cpt2/Creb1/Creb3/Acsl1/Fgfr1/Gnas/Grb2/Hras/Lipe/Ndufa2/Ndufa4/Ndufs4/Ndufv1/Npr1/Nras/Prkaca/Prkacb/Pparg/Ppargc1a/Prkab1/Prkag1/Prkg1/Mapk11/Rheb/Rps6/Rps6ka1/Rps6ka2/Cox7a2l/Smarca4/Smarcb1/Smarcc1/Sos1/Sos2/Creb3l2/Acsl6/Uqcrc1/Ndufs8/Cox15/Ndufs2/Ndufs1/Atp5g3/Ndufb6/Cox18/Creb5/Mgll/Arid1b/Map2k3/Map3k5/Mapk13/Mapk14/Creb3l1/Atp5l/Kdm3b/Atp5o/Mapk12/Dpf1/Frs2/Zfp516/Ndufs6/Ndufa4l2/Acsl5/Sirt6/Coa3/Actl6a/Mlst8/Mtor/Slc25a20/Smarce1/Atp5j2/Rps6kb2/Ndufs5/Tsc1/Atp5d/Ndufb5/Ndufa3/Ndufa9/Uqcr10/Ndufb9/Cox16/Cox20/Ndufc1/Ndufa12/Ndufa7/Cyc1/Uqcr11/Uqcrfs1/Pnpla2/Ndufb7/Sdhd/Sdha/Smarcd3/Uqcrc2/Atp5e/Ndufa6/Smarca2/Ndufb8/Ndufa10/Akt1s1/Sdhb/Coa6/Atp5g2/Cox19/Coa4/Ndufb4/Ndufc2/Ndufb2/Ndufa5/Ndufa8/Ndufaf4/Ndufaf1/Coa7/Dpf3/Ndufab1/Prdm16/Ndufv2/Rptor/Ndufs7/Ndufaf6/Cpt1c/Creb3l4/Klb/Smarcd2/Smarcd1/Arid1a/Kdm1a |
| mmu00561 | Glycerolipid metabolism | 0,00012572 | Gpat4/Dgkz/Dgkg/Dgkq/Aldh7a1/Aldh2/Aldh3a2/Akr1b3/Akr1b7/Cel/Dgka/Dgat1/Lpin1/Gpam/Lipg/Lpl/Pnliprp1/Pnliprp2/Plpp1/Dgkb/Mboat1/Lclat1/Tkfc/Dgkd/Gpat3/Glyctk/Mgll/Agpat3/Dgki/Dgkh/Plpp2/Agpat5/Dgke/Aldh9a1/Akr1a1/Lpin2/Lpin3/Pnpla2/Mboat2/Agpat2/Dgat2/Akr1b10/Plpp3/Agpat4/Mogat1/Pnlip/Agk/Plpp5/Aldh1b1 |
| mmu05221 | Acute myeloid leukemia | 0,0001361 | Braf/Raf1/Cebpe/Akt1/Akt2/Bad/Runx1/Runx1t1/Ccna2/Ccnd1/Cd14/Cebpa/Chuk/Eif4ebp1/Fcgr1/Grb2/Hras/Ikbkb/Itgam/Jup/Kit/Lef1/Myc/Nfkb1/Nras/Per2/Pik3ca/Pik3cd/Pik3r1/Pik3r2/Pik3r3/Pim1/Pml/Ppard/Rara/Rela/Spi1/Sos1/Sos2/Stat3/Stat5a/Stat5b/Tcf7/Tcf7l1/Tcf7l2/Zbtb16/Akt3/Map2k1/Map2k2/Mapk1/Mapk3/Mtor/Rps6kb2/Dusp6/Pik3cb |
| mmu05010 | Alzheimer disease | 0,00015403 | Cox6b1/Adam10/Adam17/Apaf1/Apbb1/Apoe/App/Atp2a2/Atp5a1/Atp5b/Atp5c1/Atp5pb/Atp5g1/Atp5j/Bad/Bid/Cacna1d/Calm1/Calm2/Calm3/Capn1/Capn2/Casp12/Casp3/Casp7/Casp8/Casp9/Cdk5/Cdk5r1/Cox5a/Cox5b/Cox6a1/Cox6c/Cox7a1/Cox7a2/Cox7c/Cox8a/Cycs/Eif2ak3/Fadd/Fas/Gapdh/Gnaq/Grin2d/Itpr1/Itpr2/Itpr3/Lpl/Lrp1/Mme/Mapt/Ndufa2/Ndufa4/Ndufs4/Ndufv1/Plcb1/Plcb2/Plcb3/Plcb4/Ppp3ca/Ppp3cb/Ppp3cc/Ppp3r1/Psen1/Psen2/Rtn3/Cox7a2l/Snca/Aph1b/Tnfrsf1a/Uqcrc1/Ndufs8/Aph1a/Atf6/Ndufs2/Ndufs1/Atp5g3/Ndufb6/Bace1/Mapk1/Mapk3/Atp5o/Ndufs6/Ndufa4l2/Atp2a3/Bace2/Gsk3b/Ndufs5/Atp5d/Ndufb5/Ndufa3/Ndufa9/Uqcr10/Ndufb9/Ndufc1/Ndufa12/Ndufa7/Cyc1/Uqcr11/Uqcrfs1/Ndufb7/Sdhd/Sdha/Uqcrc2/Atp5e/Ndufa6/Ndufb8/Ndufa10/Sdhb/Atp5g2/Ndufb4/Ndufc2/Ndufb2/Ndufa5/Aph1c/Ndufa8/Rtn4/Ndufab1/Ndufv2/Ndufs7/Ern1 |
| mmu05016 | Huntington disease | 0,00016878 | Dnal1/Ppif/Dnah1/Cox6b1/Slc25a4/Ap2a1/Ap2a2/Ap2m1/Apaf1/Atp5a1/Atp5b/Atp5c1/Atp5pb/Atp5g1/Atp5j/Bax/Casp3/Casp8/Casp9/Clta/Cox5a/Cox5b/Cox6a1/Cox6c/Cox7a1/Cox7a2/Cox7c/Cox8a/Creb1/Creb3/Crebbp/Cycs/Dctn1/Dlg4/Dnah11/Gnaq/Gpx1/Gpx3/Hap1/Hdac2/Htt/Itpr1/Bbc3/Ndufa2/Ndufa4/Ndufs4/Ndufv1/Nrf1/Plcb1/Plcb2/Plcb3/Plcb4/Pparg/Ppargc1a/Rest/Polr2a/Polr2c/Polr2j/Cox7a2l/Sod1/Sp1/Creb3l2/Hip1/Tfam/Rcor1/Tgm2/Uqcrc1/Vdac1/Vdac2/Vdac3/Ndufs8/Ndufs2/Dnah7b/Ndufs1/Atp5g3/Taf4/Ndufb6/Polr2b/Creb5/Ap2s1/Tbpl1/Creb3l1/Atp5o/Dnah2/Ep300/Dnah6/Ndufs6/Ndufa4l2/Dnai2/Hdac1/Dnal4/Ndufs5/Atp5d/Ndufb5/Ndufa3/Ndufa9/Uqcr10/Ndufb9/Ndufc1/Ndufa12/Ndufa7/Polr2e/Cyc1/Uqcr11/Uqcrfs1/Ndufb7/Sdhd/Sdha/Uqcrc2/Atp5e/Ndufa6/Ndufb8/Ndufa10/Cltc/Gpx7/Dctn4/Sdhb/Polr2g/Atp5g2/Ndufb4/Ndufc2/Ndufb2/Ndufa5/Ndufa8/Dnai1/Polr2d/Gpx8/Dctn2/Ndufab1/Taf4b/Ndufv2/Ift57/Cltb/Ndufs7/Gpx6/Dnali1/Creb3l4 |
| mmu05216 | Thyroid cancer | 0,00018105 | Ddb2/Tpr/Braf/Bak1/Bax/Ctnnb1/Ccnd1/Cdh1/Cdkn1a/Gadd45a/Hras/Lef1/Myc/Gadd45b/Nras/Pparg/Ret/Rxra/Rxrg/Tcf7/Tcf7l1/Tcf7l2/Tfg/Gadd45g/Map2k1/Map2k2/Mapk1/Mapk3/Polk/Ncoa4/Tpm3/Ccdc6 |
| mmu04072 | Phospholipase D signaling pathway | 0,00018651 | Dnm3/Adcy3/Dgkz/Grm6/Raf1/Dgkg/Dgkq/Adcy6/Adcy8/Adcy9/Agtr1a/Akt1/Akt2/Arf1/Arf6/Dgka/Dnm1/Dnm2/Egf/Egfr/F2/F2r/Fyn/Gab1/Gab2/Gna12/Gnas/Lpar1/Grb2/Grm8/Hras/Insr/Kit/Kitl/Mras/Nras/Pdgfa/Pdgfb/Pdgfra/Pdgfrb/Pik3ca/Pik3cd/Pik3r1/Pik3r2/Pik3r3/Pip5k1c/Pip5k1b/Pip5k1a/Pla2g4a/Plcb1/Plcb2/Plcb3/Plcb4/Plcg1/Pld1/Pld2/Plpp1/Cyth1/Cyth2/Cyth3/Ptk2b/Ptpn11/Ralgds/Rheb/Rras/Shc1/Shc3/Sos1/Sos2/Sphk1/Syk/Dgkb/Rapgef3/Dgkd/Plcg2/Akt3/Map2k1/Map2k2/Mapk1/Mapk3/Shc4/Agpat3/Pik3cg/Dgki/Pik3r5/Pla2g4e/Dgkh/Plpp2/Agpat5/Lpar2/Pdgfc/Rala/Dgke/Rapgef4/Mtor/Ralb/Tsc1/Lpar3/Rras2/Lpar6/Agpat2/Plpp3/Agpat4/Pdgfd/Pik3cb |
| mmu01200 | Carbon metabolism | 0,00022437 | H6pd/Psph/Gck/Gldc/Aldh6a1/Me2/Psat1/Shmt2/Gpt2/Me3/Pgd/Acat1/Acat2/Pcca/Acads/Aco1/Aco2/Acox1/Adh5/Aldoa/Cat/Cs/Dld/Eno1/Eno2/Esd/Fbp2/Fbp1/Fh1/Gapdh/Glud1/Got1/Got2/Gpi1/Hk1/Hk2/Idh1/Idh3b/Mdh2/Mdh1/Mthfr/Mmut/Ogdh/Pcx/Pfkl/Pfkm/Pgam1/Pkm/Pklr/Rpia/Shmt1/Sucla2/Suclg2/Taldo1/Hkdc1/Tkt/Tpi1/Tkfc/Eno4/Hibch/Dlat/Glyctk/Phgdh/Ogdhl/Sdsl/Idh2/Prps1l3/Amt/Hao2/Pfkp/Suclg1/Pgls/Rpe/Pccb/Sdhd/Sdha/Sdhb/Idh3a/Gcsh/Pdhb/Acss1/Adpgk/Ehhadh/Idnk/Gpt/Dlst/Acox3 |
| mmu00230 | Purine metabolism | 0,00024981 | Ak6/Nt5m/Adcy3/Gmpr2/Nt5c3/Atic/Ampd2/Pde4c/Ada/Adcy6/Adcy8/Adcy9/Adk/Adsl/Adssl1/Adss/Ak1/Ak2/Ak4/Ampd3/Aprt/Entpd1/Entpd2/Entpd6/Entpd5/Dck/Fhit/Gart/Gda/Gucy2e/Guk1/Itpa/Nme7/Nme1/Nme2/Npr1/Pde1b/Pde1c/Pde3b/Pde4a/Pde4b/Pde6d/Pde7a/Pde8a/Pde9a/Enpp1/Pkm/Pklr/Pnp/Rrm1/Rrm2/Pde2a/Enpp3/Entpd3/Pde8b/Xdh/Enpp4/Gmps/Prune1/Ak5/Npr2/Nt5c1a/Ppat/Gucy1a2/Pfas/Pde4d/Impdh1/Impdh2/Nt5e/Papss1/Papss2/Pde10a/Pde5a/Dguok/Pde7b/Prps1l3/Rrm2b/Nt5c/Nudt5/Gucy1b1/Nme6/Ak3/Nme4/Gucy1a1/Gmpr/Adprm/Ntpcr/Pgm2/Pnp2/Nt5c3b/Hddc3/Ak8/Nudt16/Cant1/Nt5c2/Urah/Allc |
| mmu04962 | Vasopressin-regulated water reabsorption | 0,00025118 | Adcy3/Dync2h1/Adcy6/Adcy9/Aqp2/Aqp3/Creb1/Creb3/Dctn1/Dync1h1/Dync1i1/Dync1i2/Arhgdig/Gnas/Nsf/Prkaca/Prkacb/Arhgdia/Rab11b/Rab5b/Rab5c/Creb3l2/Stx4a/Dync2li1/Vamp2/Dctn6/Creb5/Dync1li2/Dync1li1/Creb3l1/Rab5a/Rab11a/Dctn4/Dynll2/Dctn2/Creb3l4 |
| mmu00480 | Glutathione metabolism | 0,00025374 | Gstt3/Ggct/Pgd/Ggt1/Gclc/Gclm/Gpx1/Gpx3/Gsr/Gss/Gsta3/Gsta4/Gstm1/Gstm2/Gstm3/Gstm4/Gstm5/Gstp2/Gstp1/Gstt1/Gstt2/Gsto1/Idh1/Anpep/Odc1/Rrm1/Rrm2/Ggt7/Srm/Mgst2/Gstp3/Ggt5/Idh2/Rrm2b/Mgst1/Gpx4/Txndc12/Nat8f1/Mgst3/Lap3/Gpx7/Chac2/Gsto2/Gstm7/Nat8/Chac1/Gpx8/Ggt6/Oplah/Gpx6/Gstk1 |
| mmu04920 | Adipocytokine signaling pathway | 0,00025374 | Acacb/Prkaa1/Prkaa2/Prkab2/Prkag2/Npy/Agrp/Akt1/Akt2/Cd36/Chuk/Socs3/Cpt1a/Acsl1/G6pc/Ikbkb/Irs1/Irs3/Jak2/Lepr/Nfkb1/Nfkbia/Nfkbib/Nfkbie/Pomc/Ppara/Ppargc1a/Prkab1/Prkag1/Ptpn11/Rela/Rxra/Rxrg/Slc2a1/Slc2a4/Camkk2/Stat3/Stk11/Acsl6/Tnfrsf1a/Tnfrsf1b/Traf2/Akt3/Mapk10/Mapk8/Mapk9/Irs2/Acsl5/Mtor/G6pc3/Adipor2/Tradd/Adipor1/Pck2/Cpt1c |
| mmu04370 | VEGF signaling pathway | 0,00034725 | Mapkapk3/Raf1/Akt1/Akt2/Bad/Casp9/Cdc42/Ptk2/Hras/Hspb1/Kdr/Rac3/Mapkapk2/Nfatc2/Nras/Pik3ca/Pik3cd/Pik3r1/Pik3r2/Pik3r3/Prkcb/Prkcg/Pla2g4a/Plcg1/Ppp3ca/Ppp3cb/Ppp3cc/Ppp3r1/Mapk11/Ptgs2/Pxn/Rac1/Sphk1/Src/Vegfa/Plcg2/Akt3/Map2k1/Map2k2/Mapk1/Mapk13/Mapk14/Mapk3/Mapk12/Pla2g4e/Pik3cb |
| mmu05100 | Bacterial invasion of epithelial cells | 0,00034725 | Dnm3/Actb/Actg1/Arpc1b/Ctnna1/Ctnnb1/Cav1/Cav2/Cbl/Cd2ap/Cdc42/Cdh1/Clta/Bcar1/Crk/Crkl/Cttn/Dnm1/Dnm2/Elmo2/Elmo1/Ptk2/Gab1/Itga5/Itgb1/Met/Pik3ca/Pik3cd/Pik3r1/Pik3r2/Pik3r3/Pxn/Rac1/Septin8/Shc1/Shc3/Src/Ctnna3/Vcl/Elmo3/Wasf2/Shc4/Dock1/Septin11/Septin9/Septin1/Rhog/Arpc3/Arhgef26/Cltc/Mad2l2/Wasl/Cltb/Pik3cb/Arpc2/Arhgap10 |
| mmu01212 | Fatty acid metabolism | 0,00034769 | Tecr/Acaca/Acat1/Acat2/Acadl/Acadm/Acadvl/Acaa1a/Acads/Acox1/Elovl3/Cpt1a/Cpt2/Acsl1/Fasn/Hadh/Hsd17b4/Elovl6/Ppt1/Scd1/Scd2/Scp2/Acsl6/Mcat/Hadhb/Cbr4/Acaa1b/Acsf3/Mecr/Scd3/Hacd1/Acsl5/Elovl1/Ppt2/Hsd17b12/Fads2/Hacd3/Hacd4/Acadsb/Elovl5/Hacd2/Oxsm/Ehhadh/Elovl7/Fads1/Cpt1c/Acox3/Elovl4 |
| mmu05230 | Central carbon metabolism in cancer | 0,00034769 | Sco2/Gck/Raf1/Akt1/Akt2/Egfr/Erbb2/Fgfr1/Fgfr2/Fgfr3/Gls/Hif1a/Hk1/Hk2/Hras/Kit/Ldha/Ldhb/Met/Myc/Nras/Pdgfra/Pdgfrb/Pfkl/Pfkm/Pgam1/Pik3ca/Pik3cd/Pik3r1/Pik3r2/Pik3r3/Pkm/Pten/Ret/Slc1a5/Slc2a1/Slc2a2/Slc7a5/Hkdc1/Gls2/Pdk1/Akt3/Map2k1/Map2k2/Mapk1/Mapk3/Tigar/Sirt6/Pfkp/Mtor/Pdhb/Pik3cb |
| mmu04151 | PI3K-Akt signaling pathway | 0,00034769 | Itga9/Prkaa1/Prkaa2/Pkn2/Raf1/Angpt1/Angpt2/Akt1/Akt2/Areg/Atf2/Atf4/Bad/Bcl2/Bcl2l1/Bcl2l11/Brca1/Casp9/Ccnd1/Ccnd2/Ccnd3/Ccne1/Ccne2/Cdc37/Cdk2/Cdk4/Cdk6/Cdkn1a/Cdkn1b/Chad/Chrm1/Chuk/Col4a1/Col1a1/Creb1/Creb3/Atf6b/Csf1/Efna1/Efna2/Efna3/Efna4/Efna5/Egf/Egfr/Eif4ebp1/Epha2/Epo/Epor/Erbb2/Erbb3/Erbb4/F2r/Ptk2/Fgf1/Fgf15/Fgf18/Fgf2/Fgf5/Fgf8/Fgf9/Fgfr1/Fgfr2/Fgfr3/Fgfr4/Flt3l/Flt4/G6pc/Ghr/Gnb1/Gnb4/Gnb5/Gng10/Gng12/Gng2/Gng5/Gng7/Gng8/Gngt2/Lpar1/Grb2/Magi1/Gys1/Nr4a1/Hras/Hsp90ab1/Hsp90aa1/Ifnar1/Ifnar2/Igf1/Igf1r/Ikbkb/Il3ra/Il4ra/Il6ra/Il7/Insr/Irs1/Itga2/Itga2b/Itga3/Itga5/Itga6/Itga7/Itgav/Itgb1/Itgb5/Itgb6/Itgb7/Jak1/Jak2/Jak3/Kdr/Kit/Lama1/Lama3/Lama4/Lama5/Lamb2/Lamb3/Lamc2/Sgk3/Mcl1/Mdm2/Met/Kitl/Myb/Myc/Nfkb1/Nras/Osm/Osmr/Pdgfa/Pdgfb/Pdgfra/Pdgfrb/Pdpk1/Pgf/Pik3ca/Pik3cd/Pik3r1/Pik3r2/Pik3r3/Ppp2ca/Ppp2cb/Prlr/Pten/Itgb4/Rac1/Rbl2/Rela/Rheb/Rps6/Rxra/Sgk1/Sos1/Sos2/Spp1/Creb3l2/Stk11/Syk/Ppp2r5d/Tgfa/Thbs1/Tlr4/Vegfa/Vegfb/Vegfc/Vtn/Ppp2r5b/Ywhae/Ywhag/Ywhah/Ywhaq/Ywhaz/Lamc1/Ppp2r5a/Ifna13/Creb5/Ppp2r3a/Akt3/Tlr2/Itga8/Ifna15/Chrm2/Phlpp2/Pkn3/Map2k1/Map2k2/Mapk1/Mapk3/Creb3l1/Ppp2r5c/Ppp2r5e/Eif4e2/Sgk2/Pik3cg/Pik3r5/Pkn1/Itgb8/Magi2/Ppp2r1a/Ppp2r2d/Lpar2/Ywhab/Pdgfc/Foxo3/Gsk3b/Mlst8/Mtor/Rps6kb2/Ppp2r3c/Tsc1/Lpar3/Gng11/Lpar6/G6pc3/Pdgfd/Ppp2r2a/Ppp2r1b/Crtc2/Rptor/Pck2/Ddit4/Pik3cb/Eif4b/Them4/Creb3l4/Ntf5/Pik3ap1 |
| mmu03030 | DNA replication | 0,00039729 | Rfc4/Fen1/Lig1/Mcm3/Mcm2/Mcm4/Mcm5/Mcm6/Pcna/Pola2/Pold1/Pold2/Pole/Pole2/Prim1/Prim2/Rfc1/Rfc2/Rnaseh1/Rpa2/Dna2/Ssbp1/Pole4/Rnaseh2b/Pold3/Rnaseh2c/Rfc3/Rnaseh2a/Pold4/Rfc5 |
| mmu03460 | Fanconi anemia pathway | 0,00045006 | Fancf/Faap24/Fancm/Rad51c/Blm/Brca1/Brca2/Ercc1/Fanca/Fancc/Hes1/Mlh1/Pms2/Rad51/Eme2/Rev3l/Rpa2/Fanci/Cenpx/Fancd2/Top3a/Top3b/Rmi2/Usp1/Palb2/Brip1/Atr/Poli/Eme1/Polk/Fan1/Ercc4/Slx4/Fancg/Fancl/Ube2t/Wdr48/Cenps/Telo2/Faap100/Fance |
| mmu04510 | Focal adhesion | 0,00047596 | Itga9/Mylk/Rapgef1/Actn1/Braf/Rap1a/Raf1/Actb/Actg1/Akt1/Akt2/Birc3/Birc2/Arhgap5/Bad/Bcl2/Capn2/Ctnnb1/Cav1/Cav2/Ccnd1/Ccnd2/Ccnd3/Cdc42/Chad/Col4a1/Col1a1/Bcar1/Crk/Crkl/Diaph1/Egf/Egfr/Erbb2/Ptk2/Flt4/Fyn/Grb2/Hras/Igf1/Igf1r/Itga2/Itga2b/Itga3/Itga5/Itga6/Itga7/Itgav/Itgb1/Itgb5/Itgb6/Itgb7/Jun/Kdr/Lama1/Lama3/Lama4/Lama5/Lamb2/Lamb3/Lamc2/Parvb/Rac3/Met/Mylpf/Ppp1r12a/Pak1/Pdgfa/Pdgfb/Pdgfra/Pdgfrb/Pdpk1/Pgf/Pik3ca/Pik3cd/Pik3r1/Pik3r2/Pik3r3/Pip5k1c/Prkcb/Prkcg/Ppp1cb/Ppp1cc/Pten/Itgb4/Pxn/Rac1/Rasgrf1/Rock1/Rock2/Shc1/Shc3/Sos1/Sos2/Spp1/Src/Pak6/Rap1b/Thbs1/Vasp/Vav1/Vav2/Vcl/Vegfa/Vegfb/Vegfc/Vtn/Pak2/Lamc1/Zyx/Ppp1r12c/Akt3/Itga8/Map2k1/Mapk1/Mapk10/Mapk3/Mapk8/Mapk9/Shc4/Flnb/Itgb8/Ppp1r12b/Dock1/Pdgfc/Gsk3b/Vav3/Parva/Actn4/Myl12a/Myl12b/Flnc/Tln2/Pdgfd/Pik3cb |
| mmu04014 | Ras signaling pathway | 0,00051382 | Rap1a/Raf1/Abl1/Abl2/Rasa2/Angpt1/Angpt2/Akt1/Akt2/Arf6/Bad/Bcl2l1/Calm1/Calm2/Calm3/Cdc42/Chuk/Csf1/Efna1/Efna2/Efna3/Efna4/Efna5/Egf/Egfr/Epha2/Fgf1/Fgf15/Fgf18/Fgf2/Fgf5/Fgf8/Fgf9/Fgfr1/Fgfr2/Fgfr3/Fgfr4/Flt3l/Flt4/Gab1/Gab2/Gnb1/Gnb4/Gnb5/Gng10/Gng12/Gng2/Gng5/Gng7/Gng8/Gngt2/Grb2/Hras/Igf1/Igf1r/Ikbkb/Insr/Kdr/Kit/Ksr1/Rac3/Met/Kitl/Afdn/Mras/Nf1/Nfkb1/Nras/Pak1/Pdgfa/Pdgfb/Pdgfra/Pdgfrb/Pgf/Pik3ca/Pik3cd/Pik3r1/Pik3r2/Pik3r3/Prkaca/Prkacb/Prkcb/Prkcg/Pla2g1b/Pla2g4a/Plcg1/Pld1/Pld2/Ptpn11/Rab5b/Rab5c/Rac1/Rasgrp2/Rasa3/Rasal1/Rasgrf1/Rel/Rela/Ralgds/Rgl1/Rgl2/Ralbp1/Rras/Shc1/Shc3/Sos1/Sos2/Pak6/Rap1b/Rapgef5/Tgfa/Rasa1/Tiam1/Vegfa/Vegfb/Vegfc/Pak2/Plaat3/Zap70/Rasal2/Plcg2/Akt3/Ets1/Ets2/Syngap1/Rasgrp3/Map2k1/Map2k2/Mapk1/Mapk10/Mapk3/Mapk8/Mapk9/Rab5a/Shc4/Rasal3/Pla2g4e/Ksr2/Pla2g6/Rasa4/Rassf5/Pdgfc/Rala/Rassf1/Tbk1/Stk4/Ralb/Gng11/Pla2g12a/Exoc2/Rras2/Pdgfd/Plce1/Pik3cb/Ntf5 |
| mmu03440 | Homologous recombination | 0,0005203 | Babam2/Rad51c/Atm/Bard1/Blm/Brca1/Brca2/Mre11a/Pold1/Pold2/Rad50/Rad51/Rad51b/Rad51d/Rad52/Rad54l/Rpa2/Uimc1/Sem1/Top3a/Top3b/Rbbp8/Palb2/Topbp1/Brip1/Eme1/Nbn/Ssbp1/Xrcc2/Rad54b/Pold3/Babam1/Pold4/Abraxas1 |
| mmu00670 | One carbon pool by folate | 0,00060357 | Mthfsl/Aldh1l1/Mthfs/Shmt2/Atic/Mthfd1/Dhfr/Gart/Mthfd2/Mthfr/Shmt1/Aldh1l2/Tyms/Mtr/Mthfd1l/Amt/Mthfd2l/Mtfmt |
| mmu00900 | Terpenoid backbone biosynthesis | 0,00062753 | Fdps/Acat1/Acat2/Fntb/Fnta/Ggps1/Hmgcr/Hmgcs2/Mvk/Mvd/Rce1/Hmgcs1/Zmpste24/Idi1/Nus1/Pdss1/Icmt/Pcyox1/Dhdds/Pmvk/Pdss2 |
| mmu04213 | Longevity regulating pathway - multiple species | 0,00065573 | Adcy3/Prkaa1/Prkaa2/Prkab2/Prkag2/Adcy6/Adcy8/Adcy9/Akt1/Akt2/Atg5/Cat/Cryab/Eif4ebp2/Hdac2/Foxa2/Hras/Hspa1l/Hspa1b/Hspa2/Igf1/Igf1r/Insr/Irs1/Irs3/Nras/Pik3ca/Pik3cd/Pik3r1/Pik3r2/Pik3r3/Prkaca/Prkacb/Prkab1/Prkag1/Clpb/Sod1/Akt3/Irs2/Hdac1/Foxo1/Foxo3/Mtor/Rps6kb2/Akt1s1/Rptor/Pik3cb/Sirt1 |
| mmu04926 | Relaxin signaling pathway | 0,00066317 | Adcy3/Arrb1/Raf1/Adcy6/Adcy8/Adcy9/Akt1/Akt2/Atf2/Atf4/Col4a1/Col1a1/Creb1/Creb3/Atf6b/Edn1/Ednrb/Egfr/Fos/Gna15/Gnai1/Gnai2/Gnai3/Gnao1/Gnas/Gnb1/Gnb4/Gnb5/Gng10/Gng12/Gng2/Gng5/Gng7/Gng8/Gngt2/Grb2/Hras/Jun/Smad2/Mmp9/Nfkb1/Nfkbia/Nos2/Nras/Pik3ca/Pik3cd/Pik3r1/Pik3r2/Pik3r3/Prkaca/Prkacb/Prkcz/Plcb1/Plcb2/Plcb3/Plcb4/Mapk11/Rela/Rln1/Shc1/Shc3/Sos1/Sos2/Src/Creb3l2/Arrb2/Tgfb1/Tgfbr1/Tgfbr2/Vegfa/Vegfb/Vegfc/Creb5/Akt3/Map2k1/Map2k2/Map2k4/Map2k7/Mapk1/Mapk10/Mapk13/Mapk14/Mapk3/Mapk8/Mapk9/Creb3l1/Shc4/Mapk12/Gng11/Pik3cb/Creb3l4 |
| mmu00513 | Various types of N-glycan biosynthesis | 0,00075452 | Alg9/Mgat4b/Rpn1/Ddost/Man2a2/B4galt1/Hexa/Hexb/Stt3a/Man1a2/Man2a1/Mgat1/Rpn2/St3gal3/Alg1/Mgat2/Alg12/Man1b1/Man1c1/Hexdc/Mgat4a/Glt28d2/B4galnt3/B4galnt4/B4galt2/Fut8/Alg2/B4galt3/Alg14/Stt3b/Chst8/Chst9/Tusc3 |
| mmu03420 | Nucleotide excision repair | 0,00081286 | Rfc4/Ddb2/Cdk7/Ddb1/Ercc1/Ercc2/Ercc3/Gtf2h1/Gtf2h4/Lig1/Mnat1/Pcna/Pold1/Pold2/Pole/Pole2/Rad23a/Rad23b/Rfc1/Rfc2/Rpa2/Xpa/Xpc/Ercc5/Gtf2h2/Ercc6/Ercc4/Rbx1/Ccnh/Pole4/Pold3/Rfc3/Pold4/Ercc8/Rfc5 |
| mmu00520 | Amino sugar and nucleotide sugar metabolism | 0,00088645 | Gck/Uap1/Cyb5r3/Pgm3/Mpi/Cmas/Gfpt1/Gfpt2/Galk1/Gpi1/Hexa/Hexb/Hk1/Hk2/Hkdc1/Ugp2/Gmds/Gfus/Ugdh/Uap1l1/Cyb5rl/Fcsk/Amdhd2/Gnpda1/Cyb5r4/Pmm1/Gne/Pmm2/Gnpnat1/Nagk/Pgm2/Nanp/Uxs1/Gnpda2/Gmppa/Cyb5r1/Npl/Gale/Nans |
| mmu03013 | RNA transport | 0,00132662 | Sap18b/Eif3j2/Rpp25/Nup107/Xpo1/Thoc5/Pom121/Eif2b3/Tpr/Sec13/Elac1/Semp2l2b/Tgs1/Pop5/Eef1a1/Eif1a/Eif2s1/Eif2b4/Eif3a/Eif4a1/Eif4a2/Eif4ebp1/Eif4ebp2/Eif4g2/Fxr1/Kpnb1/Eif3e/Nup155/Sumo2/Magoh/Nup50/Pabpc1/Pnn/Nup88/Eif4a3/Ran/Ranbp2/Rangap1/Rasl2-9/Pom121l2/Upf1/Rnps1/Cyfip1/Smn1/Sumo3/Rpp40/Eif4g1/Strap/Eif1/Eif2b1/Tacc3/Gemin5/Alyref/Eif2b2/Eif5/Nup153/Paip1/Ube2i/Sumo1/Eif2b5/Thoc1/Eif5b/Rpp38/Nup214/Pabpc4/Eif4g3/Nupl2/Nup133/Fxr2/Pabpc4l/Nup54/Eif4e2/Prmt5/Gemin4/Eif3b/Upf2/Nup85/Srrm1/Nxf1/Eif3g/Ddx20/Rpp30/Nup210/Eif3i/Eif3d/Alyref2/Acin1/Eif3c/Nxt1/Nup160/Snupn/Eif3f/Pop4/Magohb/Rae1/Upf3a/Rpp14/Eif2s2/Gemin6/Rpp21/Eif3h/Elac2/Eif1b/Nup35/Gemin7/Nup37/Nup43/Rpp25l/Trnt1/Nup205/Nup93/Nupl1/Seh1l/Xpo5/Ndc1/Xpot/Thoc3/Pop7/Eif4b/Senp2/Cyfip2/Nmd3 |
| mmu01040 | Biosynthesis of unsaturated fatty acids | 0,00132738 | Tecr/Acaa1a/Acox1/Elovl3/Hsd17b4/Elovl6/Acot2/Acot3/Scd1/Scd2/Scp2/Acaa1b/Acot1/Scd3/Hacd1/Elovl1/Hsd17b12/Fads2/Hacd3/Hacd4/Elovl5/Acot7/Hacd2/Elovl7/Fads1/Acox3/Elovl4 |
| mmu04215 | Apoptosis - multiple species | 0,00132738 | Apaf1/Birc3/Birc2/Birc5/Bak1/Bax/Bcl2/Bcl2l1/Bid/Bcl2l11/Birc6/Casp3/Casp7/Casp8/Casp9/Cycs/Fadd/Bbc3/Septin4/Tnfrsf1a/Mapk10/Mapk8/Mapk9/Bok/Becn1/Pmaip1/Diablo |
| mmu03040 | Spliceosome | 0,00137694 | Tra2a/Sf3b3/Snrpd2/Sf3b4/Srsf9/U2af1/Srsf1/Aqr/Dhx15/Ddx5/Srsf10/Hnrnpc/Hnrnpk/Hspa1l/Hspa1b/Hspa2/Magoh/Prpf8/Eif4a3/Slu7/Rbmxl1/Sf3a2/Sart1/Srsf2/Srsf3/Srsf5/Tra2b/Eftud2/Snrpc/Snrnp70/Snrpb/Snrpb2/Snrpd1/Snrpe/Snu13/Ddx46/Alyref/Dhx8/U2af2/Srsf7/Thoc1/Hnrnpa3/Pcbp1/Txnl4a/Lsm2/Cherp/Prpf19/Usp39/Sf3b2/Snrnp200/Lsm4/Hnrnpu/Plrg1/Rp9/Alyref2/Ppie/Tcerg1/Prpf40a/Acin1/Srsf4/Isy1/Sf3b6/Lsm7/Ppih/Sf3b5/Snw1/Lsm5/Magohb/Zmat2/Snrnp27/Ctnnbl1/Rbm22/Crnkl1/Prpf38b/Rbm25/Prpf18/Snrpd3/U2surp/Puf60/Srsf6/Snrpg/Bcas2/Syf2/Ppil1/Prpf6/Snrpa1/Prpf31/Dhx16/Snrpf/Prpf3/Cdc40/Ccdc12/Thoc3/Ddx23/Sf3a3/Smndc1/Lsm8/Hnrnpm/Rbm17/Lsm6/Sf3b1 |
| mmu00450 | Selenocompound metabolism | 0,00162326 | Cth/Sephs1/Sephs2/Sepsecs/Mars2/Pstk/Mars1/Inmt/Kyat3/Txnrd3/Mtr/Papss1/Papss2/Txnrd1/Scly/Kyat1 |
| mmu05219 | Bladder cancer | 0,00168473 | Braf/Raf1/Ccnd1/Cdh1/Cdk4/Cdkn1a/Cdkn2a/Dapk2/Dapk3/E2f1/E2f3/Egf/Egfr/Erbb2/Fgfr3/Hbegf/Hras/Mdm2/Mmp9/Myc/Nras/Rb1/Src/Thbs1/Vegfa/E2f2/Map2k1/Map2k2/Mapk1/Mapk3/Rassf1/Dapk1/Rps6ka5 |
| mmu04666 | Fc gamma R-mediated phagocytosis | 0,00168473 | Raf1/Akt1/Akt2/Arf6/Arpc1b/Cdc42/Cfl1/Cfl2/Crk/Crkl/Asap1/Dnm2/Fcgr1/Gab2/Hck/Inppl1/Limk1/Limk2/Lyn/Marcks/Myo10/Pak1/Pik3ca/Pik3cd/Pik3r1/Pik3r2/Pik3r3/Pip5k1c/Pip5k1b/Pip5k1a/Prkcb/Prkcg/Prkcd/Prkce/Pla2g4a/Plcg1/Pld1/Pld2/Plpp1/Rac1/Scin/Sphk1/Syk/Asap2/Vasp/Vav1/Vav2/Gsn/Asap3/Plcg2/Akt3/Wasf2/Map2k1/Mapk1/Mapk3/Bin1/Pla2g4e/Plpp2/Pla2g6/Arpc3/Vav3/Rps6kb2/Plpp3/Pik3cb/Arpc2 |
| mmu04916 | Melanogenesis | 0,00213652 | Adcy3/Camk2d/Raf1/Adcy6/Adcy8/Adcy9/Calm1/Calm2/Calm3/Ctnnb1/Creb1/Creb3/Crebbp/Dvl1/Dvl2/Dvl3/Edn1/Ednrb/Fzd1/Fzd3/Fzd4/Fzd5/Fzd6/Fzd7/Fzd8/Fzd9/Gnai1/Gnai2/Gnai3/Gnao1/Gnaq/Gnas/Hras/Kit/Lef1/Kitl/Mitf/Nras/Prkaca/Prkacb/Prkcb/Prkcg/Plcb1/Plcb2/Plcb3/Plcb4/Pomc/Creb3l2/Tcf7/Tcf7l1/Tcf7l2/Wnt9a/Wnt1/Wnt10b/Wnt11/Wnt9b/Wnt2/Wnt2b/Wnt4/Wnt5a/Wnt5b/Wnt6/Map2k1/Map2k2/Mapk1/Mapk3/Creb3l1/Ep300/Gsk3b/Fzd2/Creb3l4 |
| mmu00240 | Pyrimidine metabolism | 0,00232357 | Upb1/Nt5m/Nt5c3/Dut/Entpd1/Entpd6/Entpd5/Dck/Nme7/Nme1/Nme2/Enpp1/Pnp/Rrm1/Rrm2/Enpp3/Entpd3/Tk1/Dtymk/Cmpk2/Tyms/Uck1/Umps/Upp1/Nt5c1a/Nt5e/Dctd/Rrm2b/Nt5c/Ctps/Nme6/Nme4/Dhodh/Tk2/Dctpp1/Cmpk1/Pnp2/Nt5c3b/Cda/Cant1/Upp2/Nt5c2/Uck2/Dpyd |
| mmu05142 | Chagas disease (American trypanosomiasis) | 0,00252635 | Ticam1/Akt1/Akt2/Bdkrb2/C1qa/Calr/Casp8/Cd247/Cflar/Chuk/Fadd/Fas/Fos/Gna11/Gna14/Gna15/Gnai1/Gnai2/Gnai3/Gnal/Gnao1/Gnaq/Gnas/Ifngr1/Ifngr2/Ikbkb/Il10/Jun/Smad2/Nfkb1/Nfkbia/Nos2/Pik3ca/Pik3cd/Pik3r1/Pik3r2/Pik3r3/Plcb1/Plcb2/Plcb3/Plcb4/Ppp2ca/Ppp2cb/Mapk11/Rela/Ccl5/Tgfb1/Tgfb2/Tgfb3/Tgfbr1/Tgfbr2/Tlr4/Tnfrsf1a/Traf6/Akt3/Tlr2/Map2k4/Mapk1/Mapk10/Mapk13/Mapk14/Mapk3/Mapk8/Mapk9/Irak4/Mapk12/Kng2/Ppp2r1a/Ppp2r2d/Ppp2r2a/Ppp2r1b/Pik3cb |
| mmu04392 | Hippo signaling pathway - multiple species | 0,00256758 | Ajuba/Lats1/Nf2/Pak1/Wwc1/Rassf4/Rassf2/Tead1/Tead2/Tead3/Tead4/Yap1/Csnk1e/Limd1/Frmd6/Fat4/Lats2/Stk3/Rassf1/Sav1/Mob1b/Rassf6/Wwtr1 |
| mmu04380 | Osteoclast differentiation | 0,0028056 | Acp5/Akt1/Akt2/Camk4/Chuk/Socs3/Socs1/Creb1/Csf1/Cyba/Fcgr1/Fcgr3/Fhl2/Fos/Fosb/Fosl1/Fosl2/Fyn/Gab2/Grb2/Ifnar1/Ifnar2/Ifngr1/Ifngr2/Ikbkb/Il1r1/Irf9/Jak1/Jun/Junb/Jund/Lck/Mitf/Nfatc1/Nfatc2/Nfkb1/Nfkb2/Nfkbia/Tnfrsf11b/Sqstm1/Pik3ca/Pik3cd/Pik3r1/Pik3r2/Pik3r3/Pparg/Ppp3ca/Ppp3cb/Ppp3cc/Ppp3r1/Mapk11/Sirpa/Rac1/Rela/Relb/Spi1/Stat1/Stat2/Syk/Tec/Tgfb1/Tgfb2/Tgfbr1/Tgfbr2/Tnfrsf11a/Tnfrsf1a/Traf2/Traf6/Plcg2/Akt3/Map2k1/Map2k6/Map2k7/Map3k7/Mapk1/Mapk10/Mapk13/Mapk14/Mapk3/Mapk8/Mapk9/Mapk12/Map3k14/Tyk2/Tab1/Tab2/Cyld/Pik3cb |
| mmu04066 | HIF-1 signaling pathway | 0,002918 | Camk2d/Egln1/Egln2/Egln3/Angpt1/Angpt2/Akt1/Akt2/Aldoa/Arnt/Bcl2/Cdkn1a/Cdkn1b/Crebbp/Edn1/Egf/Egfr/Eif4ebp1/Eno1/Eno2/Epo/Erbb2/Gapdh/Hif1a/Hk1/Hk2/Hmox1/Ifngr1/Ifngr2/Igf1/Igf1r/Il6ra/Insr/Ldha/Ldhb/Ltbr/Pfkfb3/Mknk1/Mknk2/Nfkb1/Nos2/Pfkl/Pfkm/Pik3ca/Pik3cd/Pik3r1/Pik3r2/Pik3r3/Prkcb/Prkcg/Plcg1/Rela/Rps6/Slc2a1/Stat3/Hkdc1/Tlr4/Tfrc/Vegfa/Vhl/Eno4/Pdk1/Plcg2/Akt3/Map2k1/Map2k2/Mapk1/Mapk3/Eif4e2/Ep300/Pfkp/Rbx1/Mtor/Rps6kb2/Elob/Pdhb/Cul2/Pik3cb |
| mmu04922 | Glucagon signaling pathway | 0,002918 | Acacb/Phkb/Gck/Ppp4r3b/Prkaa1/Acaca/Camk2d/Prkaa2/Prkab2/Prkag2/Pygb/Pygl/Akt1/Akt2/Atf2/Atf4/Calm1/Calm2/Calm3/Cpt1a/Creb1/Creb3/Crebbp/Fbp2/Fbp1/G6pc/Gnaq/Gnas/Gys1/Itpr1/Itpr2/Itpr3/Ldha/Ldhb/Sik1/Pde3b/Pfkl/Pfkm/Pgam1/Pkm/Prkaca/Prkacb/Plcb1/Plcb2/Plcb3/Plcb4/Ppara/Ppargc1a/Ppp3ca/Ppp3cb/Ppp3cc/Ppp3r1/Prkab1/Prkag1/Slc2a1/Slc2a2/Creb3l2/Creb5/Sik2/Akt3/Creb3l1/Ep300/Ppp4c/Pfkp/Foxo1/Pdhb/G6pc3/Phkg2/Crtc2/Pck2/Cpt1c/Creb3l4/Sirt1 |
| mmu00640 | Propanoate metabolism | 0,00310409 | Acacb/Aldh6a1/Acaca/Acat1/Acat2/Pcca/Acads/Acox1/Bckdha/Bckdhb/Dbt/Dld/Ldha/Ldhb/Mmut/Sucla2/Suclg2/Hibch/Abat/Acss3/Echdc1/Suclg1/Mlycd/Pccb/Acss1/Ehhadh/Acox3 |
| mmu04130 | SNARE interactions in vesicular transport | 0,00310409 | Bet1/Stx2/Sec22b/Snap23/Stx1a/Stx3/Stx4a/Vamp1/Vamp2/Vamp3/Vamp8/Bnip1/Stx16/Vamp4/Stx7/Gosr1/Vti1a/Vti1b/Vamp5/Stx8/Ykt6/Stx6/Use1/Stx17/Stx19/Stx18/Stx11 |
| mmu04933 | AGE-RAGE signaling pathway in diabetic complications | 0,00310409 | Agtr1a/Akt1/Akt2/Bax/Bcl2/Casp3/Ccnd1/Cdc42/Cdk4/Cdkn1b/Col4a1/Col1a1/Diaph1/Edn1/Egr1/F3/Hras/Icam1/Jak2/Jun/Smad2/Smad3/Smad4/Nfatc1/Nfkb1/Nras/Pik3ca/Pik3cd/Pik3r1/Pik3r2/Pik3r3/Pim1/Prkcb/Prkcd/Prkce/Prkcz/Plcb1/Plcb2/Plcb3/Plcb4/Plcd1/Plcg1/Mapk11/Rac1/Rela/Stat1/Stat3/Stat5a/Stat5b/Tgfb1/Tgfb2/Tgfb3/Tgfbr1/Tgfbr2/Thbd/Vegfa/Vegfb/Vegfc/Plcg2/Akt3/Mapk1/Mapk10/Mapk13/Mapk14/Mapk3/Mapk8/Mapk9/Mapk12/Foxo1/Plce1/Pik3cb |
| mmu05164 | Influenza A | 0,00316647 | Dnajc3/Xpo1/Try5/Ticam1/Raf1/Fdps/Actb/Actg1/Akt1/Akt2/Slc25a4/Apaf1/Bak1/Bax/Bid/Ciita/Casp3/Casp8/Casp9/Ccnd3/Cdk4/Cdk6/Chuk/Socs3/Crebbp/Cycs/Eif2s1/Fadd/Fas/Tlr3/H2-Ab1/H2-Eb1/H2-DMa/Icam1/Cxcl10/Ifnar1/Ifnar2/Ifngr1/Ifngr2/Ikbkb/Il18/Irf9/Jak1/Jak2/Kpna1/Kpna2/Kpna6/Mx2/Nfkb1/Nfkbia/Nfkbib/Pik3ca/Pik3cd/Pik3r1/Pik3r2/Pik3r3/Prkcb/Pml/Eif2ak2/Rab11b/Rela/Ccl5/Stat1/Stat2/Tmprss4/Trim25/Tlr4/Tnfrsf10b/Tnfrsf1a/Traf3/Tnfsf10/Prss2/Try4/Vdac1/Mavs/Ddx58/Ifna13/Hnrnpul1/Akt3/Rnasel/Ifna15/Oas1a/Map2k1/Map2k2/Mapk1/Mapk3/Irak4/Nlrx1/Ep300/Kpna7/Try10/Tmprss2/Nxf1/Rab11a/Irf7/Irf3/Cpsf4/Pabpn1/Tyk2/Adar/Tbk1/Nxt1/Ikbke/Rsad2/Rae1/Pycard/2210010C04Rik/Ifih1/Tradd/1810009J06Rik/Pik3cb |
| mmu04371 | Apelin signaling pathway | 0,00367833 | Adcy3/Prkaa1/Mef2bl/Mylk/Prkaa2/Prkab2/Prkag2/Raf1/Slc8a3/Adcy6/Adcy8/Adcy9/Agtr1a/Akt1/Akt2/Calm1/Calm2/Calm3/Camk4/Ccnd1/Cdh1/Egr1/Gnai1/Gnai2/Gnai3/Gnaq/Gnb1/Gnb4/Gnb5/Gng10/Gng12/Gng2/Gng5/Gng7/Gng8/Gngt2/Hdac5/Hras/Itpr1/Itpr2/Itpr3/Jag1/Klf2/Lipe/Smad2/Smad3/Smad4/Mef2a/Mef2c/Mef2d/Mras/Myl4/Nos2/Notch3/Nras/Nrf1/Pde3b/Prkaca/Prkacb/Prkce/Plat/Plcb1/Plcb2/Plcb3/Plcb4/Ppargc1a/Prkab1/Prkag1/Rps6/Rras/Slc8a1/Slc9a1/Sphk1/Spp1/Hdac4/Tfam/Tgfbr1/Pik3c3/Akt3/Map2k1/Map2k2/Mapk1/Mapk3/Pik3cg/Pik3r5/Becn1/Gabarap/Mtor/Rps6kb2/Gng11/Rras2/Pik3r4/Gabarapl2 |
| mmu00564 | Glycerophospholipid metabolism | 0,00428164 | Gpat4/Dgkz/Dgkg/Dgkq/Cds2/Ache/Chka/Pcyt1a/Dgka/Lpin1/Gpd1/Gpd2/Gnpat/Gpam/Lpcat3/Pemt/Lypla1/Pla2g1b/Pla2g4a/Pld1/Pld2/Pld3/Plpp1/Ptdss1/Pla2g15/Lpcat1/Chpt1/Etnk2/Dgkb/Mboat1/Lclat1/Plaat3/Lpgat1/Dgkd/Gpat3/Phospho1/Lypla2/Lpcat2/Ptdss2/Selenoi/Agpat3/Dgki/Pisd/Pla2g4e/Gpd1l/Dgkh/Pnpla6/Plpp2/Agpat5/Pla2g6/Dgke/Lpin2/Lpin3/Pla2g12a/Adprm/Crls1/Mboat2/Agpat2/Plpp3/Agpat4/Pcyt2/Etnppl/Plpp5/Gpcpd1/Pgs1/Cds1/Etnk1/Lpcat4 |
| mmu00510 | N-Glycan biosynthesis | 0,00445727 | Alg9/Mgat4b/Rpn1/Mgat5/Ddost/Dpagt1/Dpm2/Man2a2/B4galt1/Stt3a/Man1a2/Man2a1/Mgat1/Rpn2/St6gal1/Alg1/Mgat2/Alg12/Man1b1/Dolk/Man1c1/Mgat5b/Mgat4a/Glt28d2/Alg6/Alg8/B4galt2/Fut8/Alg2/Dolpp1/Srd5a3/B4galt3/Mogs/Alg5/Alg14/Stt3b/Dpm3/Tusc3 |
| mmu00620 | Pyruvate metabolism | 0,00473446 | Acacb/Me2/Acaca/Me3/Glo1/Acat1/Acat2/Aldh7a1/Aldh2/Aldh3a2/Dld/Fh1/Hagh/Ldha/Ldhb/Mdh2/Mdh1/Pcx/Pkm/Pklr/Dlat/Ldhd/Aldh9a1/Acyp1/Pdhb/Acss1/Aldh1b1/Pck2/Acyp2/Grhpr |
| mmu03430 | Mismatch repair | 0,00483845 | Rfc4/Lig1/Mlh1/Msh2/Msh6/Pcna/Pms2/Pold1/Pold2/Rfc1/Rfc2/Rpa2/Mlh3/Exo1/Ssbp1/Pold3/Rfc3/Pold4/Rfc5 |
| mmu04625 | C-type lectin receptor signaling pathway | 0,00511288 | Raf1/Akt1/Akt2/Bcl10/Bcl3/Calm1/Calm2/Calm3/Casp8/Chuk/Plk3/Egr2/Egr3/Hras/Ikbkb/Il10/Irf1/Irf9/Itpr1/Itpr2/Itpr3/Jun/Ksr1/Mapkapk2/Mdm2/Mras/Nfatc1/Nfatc2/Nfatc3/Nfkb1/Nfkb2/Nfkbia/Nras/Pak1/Pik3ca/Pik3cd/Pik3r1/Pik3r2/Pik3r3/Prkcd/Ppp3ca/Ppp3cb/Ppp3cc/Ppp3r1/Mapk11/Ptgs2/Ptpn11/Rela/Relb/Rras/Src/Stat1/Stat2/Cblb/Syk/Plcg2/Akt3/Il17d/Malt1/Mapk1/Mapk10/Mapk13/Mapk14/Mapk3/Mapk8/Mapk9/Mapk12/Card9/Map3k14/Ikbke/Clec7a/Pycard/Rras2/Arhgef12/Nfatc4/Cyld/Pik3cb |
| mmu01230 | Biosynthesis of amino acids | 0,00567882 | Psph/Psat1/Cth/Shmt2/Mat2b/Gpt2/Acy1/Asl/Aco1/Aco2/Aldoa/Mat1a/Arg1/Arg2/Ass1/Bcat2/Cbs/Cs/Eno1/Eno2/Gapdh/Got1/Got2/Idh1/Idh3b/Pah/Pcx/Pfkl/Pfkm/Pgam1/Pkm/Pklr/Rpia/Shmt1/Pycr1/Taldo1/Nags/Tkt/Tpi1/Eno4/Mat2a/Phgdh/Mtr/Sdsl/Idh2/Asns/Prps1l3/Pfkp/Aldh18a1/Pycrl/Rpe/Idh3a/Pycr2/Tha1/Gpt |
| mmu00511 | Other glycan degradation | 0,00598698 | Manba/Aga/Gba/Hexa/Hexb/Man2b1/Man2b2/Neu1/Engase/Gba2/Hexdc/Neu2/Neu3/Fuca2/Fuca1/Man2c1 |
| mmu04912 | GnRH signaling pathway | 0,00625973 | Adcy3/Camk2d/Raf1/Adcy6/Adcy8/Adcy9/Atf4/Cacna1d/Calm1/Calm2/Calm3/Cdc42/Cga/Egfr/Egr1/Gna11/Gnaq/Gnas/Gnrhr/Grb2/Hbegf/Hras/Itpr1/Itpr2/Itpr3/Jun/Mmp14/Nras/Prkaca/Prkacb/Prkcb/Prkcd/Pla2g4a/Plcb1/Plcb2/Plcb3/Plcb4/Pld1/Pld2/Mapk11/Ptk2b/Sos1/Sos2/Src/Mapk7/Map2k1/Map2k2/Map2k3/Map2k4/Map2k6/Map2k7/Map3k1/Map3k2/Map3k3/Mapk1/Mapk10/Mapk13/Mapk14/Mapk3/Mapk8/Mapk9/Mapk12/Pla2g4e |
| mmu03410 | Base excision repair | 0,00654291 | Parp1/Parp2/Apex1/Fen1/Hmgb1/Lig1/Mbd4/Nthl1/Ogg1/Pcna/Polb/Pold1/Pold2/Pole/Pole2/Tdg/Ung/Xrcc1/Neil3/Parp3/Mpg/Neil2/Pole4/Pold3/Pold4/Smug1/Neil1 |
| mmu04930 | Type II diabetes mellitus | 0,00818308 | Gck/Cacna1a/Cacna1d/Cacna1e/Cacna1g/Socs3/Socs1/Hk1/Hk2/Ikbkb/Insr/Irs1/Irs3/Pdx1/Pik3ca/Pik3cd/Pik3r1/Pik3r2/Pik3r3/Pkm/Prkcd/Prkce/Prkcz/Pklr/Slc2a2/Slc2a4/Hkdc1/Socs2/Mapk1/Mapk10/Mapk3/Mapk8/Mapk9/Irs2/Mtor/Pik3cb |
| mmu04659 | Th17 cell differentiation | 0,00818446 | Ahr/Runx1/Cd247/Chuk/Fos/Gata3/H2-Ab1/H2-Eb1/H2-DMa/Hif1a/Hsp90ab1/Hsp90aa1/Ifngr1/Ifngr2/Ikbkb/Il12rb1/Il1r1/Il1rap/Il4ra/Il6ra/Il6st/Irf4/Jak1/Jak2/Jak3/Jun/Lck/Smad2/Smad3/Smad4/Nfatc1/Nfatc2/Nfatc3/Nfkb1/Nfkbia/Nfkbib/Nfkbie/Plcg1/Ppp3ca/Ppp3cb/Ppp3cc/Ppp3r1/Mapk11/Rara/Rela/Rora/Rorc/Rxra/Rxrg/Stat1/Stat3/Stat5a/Stat5b/Stat6/Tgfb1/Tgfbr1/Tgfbr2/Zap70/Il17d/Mapk1/Mapk10/Mapk13/Mapk14/Mapk3/Mapk8/Mapk9/Mapk12/Tyk2/Mtor/Tbx21 |
| mmu04923 | Regulation of lipolysis in adipocytes | 0,00921923 | Adcy3/Npy/Adcy6/Adcy8/Adcy9/Adora1/Adrb1/Adrb2/Akt1/Akt2/Cga/Gnai1/Gnai2/Gnai3/Gnas/Insr/Irs1/Irs3/Lipe/Npr1/Npy1r/Pde3b/Pik3ca/Pik3cd/Pik3r1/Pik3r2/Pik3r3/Prkaca/Prkacb/Prkg1/Ptger3/Ptgs1/Ptgs2/Tshr/Plaat3/Akt3/Mgll/Irs2/Pnpla2/Abhd5/Pik3cb |
| mmu04728 | Dopaminergic synapse | 0,01106418 | Camk2d/Slc18a1/Akt1/Akt2/Arntl/Atf2/Atf4/Cacna1a/Cacna1d/Calm1/Calm2/Calm3/Clock/Comt/Creb1/Creb3/Atf6b/Slc6a3/Ddc/Fos/Gnai1/Gnai2/Gnai3/Gnal/Gnao1/Gnaq/Gnas/Gnb1/Gnb4/Gnb5/Gng10/Gng12/Gng2/Gng5/Gng7/Gng8/Gngt2/Itpr1/Itpr2/Itpr3/Kcnj9/Kif5a/Kif5b/Kif5c/Prkaca/Prkacb/Prkcb/Prkcg/Plcb1/Plcb2/Plcb3/Plcb4/Ppp1cb/Ppp1cc/Ppp1r1b/Ppp2ca/Ppp2cb/Ppp3ca/Ppp3cb/Ppp3cc/Mapk11/Scn1a/Creb3l2/Slc18a2/Arrb2/Ppp2r5d/Ppp2r5b/Ppp2r5a/Creb5/Ppp2r3a/Akt3/Mapk10/Mapk13/Mapk14/Mapk8/Mapk9/Creb3l1/Ppp2r5c/Ppp2r5e/Mapk12/Ppp2r1a/Ppp2r2d/Gsk3b/Ppp2r3c/Gsk3a/Gng11/Ppp2r2a/Ppp2r1b/Creb3l4 |
| mmu00100 | Steroid biosynthesis | 0,01121745 | Cel/Cyp27b1/Cyp51/Dhcr7/Fdft1/Hsd17b7/Lipa/Lss/Soat1/Sqle/Soat2/Sc5d/Cyp2r1/Msmo1/Tm7sf2/Dhcr24/Lbr |
| mmu04660 | T cell receptor signaling pathway | 0,01155855 | Raf1/Akt1/Akt2/Bcl10/Cd247/Cdc42/Cdk4/Chuk/Dlg1/Fos/Fyn/Grb2/Ptpn6/Hras/Ikbkb/Il10/Jun/Lck/Nck1/Nck2/Nfatc1/Nfatc2/Nfatc3/Nfkb1/Nfkbia/Nfkbib/Nfkbie/Nras/Pak1/Pdpk1/Pik3ca/Pik3cd/Pik3r1/Pik3r2/Pik3r3/Plcg1/Ppp3ca/Ppp3cb/Ppp3cc/Ppp3r1/Mapk11/Rela/Sos1/Sos2/Cblb/Pak6/Tec/Vav1/Vav2/Pak2/Zap70/Akt3/Malt1/Map2k1/Map2k2/Map2k7/Map3k7/Map3k8/Mapk1/Mapk10/Mapk13/Mapk14/Mapk3/Mapk8/Mapk9/Mapk12/Map3k14/Gsk3b/Vav3/Pik3cb |
| mmu00051 | Fructose and mannose metabolism | 0,01296539 | Mpi/Aldoa/Akr1b3/Akr1b7/Fbp2/Fbp1/Hk1/Hk2/Khk/Pfkfb3/Pfkfb2/Pfkl/Pfkm/Sord/Hkdc1/Gmds/Tpi1/Gfus/Tkfc/Fcsk/Pfkfb4/Pmm1/Tigar/Pmm2/Pfkp/Akr1b10/Gmppa |
| mmu00020 | Citrate cycle (TCA cycle) | 0,01304259 | Acly/Aco1/Aco2/Cs/Dld/Fh1/Idh1/Idh3b/Mdh2/Mdh1/Ogdh/Pcx/Sucla2/Suclg2/Dlat/Ogdhl/Idh2/Suclg1/Sdhd/Sdha/Sdhb/Idh3a/Pdhb/Pck2/Dlst |
| mmu04136 | Autophagy - other | 0,01304259 | Atg5/Ppp2ca/Ppp2cb/Atg9b/Pik3c3/Atg4d/Atg4c/Atg9a/Ulk2/Atg2a/Wipi1/Becn1/Gabarap/Mlst8/Mtor/Atg4b/Atg10/Atg3/Atg101/Atg7/Rptor/Wipi2/Pik3r4/Atg16l1/Gabarapl2 |
| mmu04979 | Cholesterol metabolism | 0,01368103 | Ldlrap1/Pcsk9/Cyp27a1/Abca1/Apoa1/Apoe/Apoh/Tspo/Cd36/Lrp2/Ldlr/Lipa/Lipg/Lpl/Lrp1/Lrpap1/Npc1/Pltp/Soat1/Sort1/Scarb1/Mylip/Vdac1/Vdac2/Vdac3/Soat2/Abcg5/Abcb11/Vapa/Nceh1/Vapb/Angptl4/Stard3/Angptl8/Npc2/Osbpl5 |
| mmu05012 | Parkinson disease | 0,01597219 | Ppif/Cox6b1/Slc18a1/Adora2a/Slc25a4/Apaf1/Atp5a1/Atp5b/Atp5c1/Atp5pb/Atp5g1/Atp5j/Casp3/Casp9/Cox5a/Cox5b/Cox6a1/Cox6c/Cox7a1/Cox7a2/Cox7c/Cox8a/Cycs/Slc6a3/Ube2j2/Gnai1/Gnai2/Gnai3/Gnal/Ndufa2/Ndufa4/Ndufs4/Ndufv1/Prkaca/Prkacb/Septin5/Cox7a2l/Snca/Slc18a2/Ubb/Ube2l3/Ube2g2/Uqcrc1/Vdac1/Vdac2/Vdac3/Ndufs8/Ndufs2/Ndufs1/Atp5g3/Ndufb6/Atp5o/Ndufs6/Ndufa4l2/Prkn/Ube2l6/Park7/Ndufs5/Atp5d/Ndufb5/Ndufa3/Ndufa9/Uqcr10/Ndufb9/Ndufc1/Ndufa12/Ndufa7/Cyc1/Uqcr11/Uqcrfs1/Lrrk2/Ndufb7/Sdhd/Sdha/Uqcrc2/Atp5e/Ube2g1/Ndufa6/Ndufb8/Ndufa10/Sdhb/Sncaip/Atp5g2/Ndufb4/Ndufc2/Ndufb2/Ndufa5/Ndufa8/Pink1/Ndufab1/Ndufv2/Uba7/Ndufs7 |
| mmu05132 | Salmonella infection | 0,01758084 | Dync2h1/Actb/Actg1/Arpc1b/Cd14/Cdc42/Dync1h1/Dync1i1/Dync1i2/Fos/Cxcl1/Ifngr1/Ifngr2/Il18/Jun/Klc1/Klc2/Nfkb1/Nos2/Pfn1/Pfn2/Mapk11/Rab7/Rac1/Rela/Rock1/Rock2/Tjp1/Tlr4/Klc3/Dync1li2/Dync1li1/Wasf2/Mapk1/Mapk10/Mapk13/Mapk14/Mapk3/Mapk8/Mapk9/Rilp/Flnb/Mapk12/Pkn1/Tlr5/Rhog/Arpc3/Pycard/Flnc/Plekhm2/Wasl/Arpc2 |
| mmu00600 | Sphingolipid metabolism | 0,01820171 | Asah1/Degs1/Galc/Gba/Neu1/Plpp1/Sgpl1/Smpd1/Smpd2/Sphk1/Sptlc2/Sgms1/Ugcg/Gba2/Acer2/Neu2/Cers6/Sptlc1/Sgpp2/Plpp2/Neu3/Gal3st1/Asah2/B4galt6/Smpd3/Acer3/Cers4/Plpp3/Degs2/Kdsr/Cers5/Sgms2/Cers2/Smpd4/Sgpp1 |
| mmu04658 | Th1 and Th2 cell differentiation | 0,02216645 | Maml1/Cd247/Chuk/Dll1/Dll3/Fos/Gata3/H2-Ab1/H2-Eb1/H2-DMa/Ifngr1/Ifngr2/Ikbkb/Il12rb1/Il4ra/Jag1/Jak1/Jak2/Jak3/Jun/Lck/Maf/Nfatc1/Nfatc2/Nfatc3/Nfkb1/Nfkbia/Nfkbib/Nfkbie/Notch1/Notch2/Notch3/Plcg1/Ppp3ca/Ppp3cb/Ppp3cc/Ppp3r1/Mapk11/Rbpj/Rela/Stat1/Stat4/Stat5a/Stat5b/Stat6/Zap70/Mapk1/Mapk10/Mapk13/Mapk14/Mapk3/Mapk8/Mapk9/Maml2/Mapk12/Maml3/Dll4/Tyk2/Tbx21 |
| mmu04611 | Platelet activation | 0,02263269 | Adcy3/Mylk/Orai1/Rap1a/Actb/Actg1/Adcy6/Adcy8/Adcy9/Akt1/Akt2/Col1a1/F2/F2r/Fyn/Gnai1/Gnai2/Gnai3/Gnaq/Gnas/Gp1bb/Gp5/Itga2/Itga2b/Itgb1/Itpr1/Itpr2/Itpr3/Lyn/Ppp1r12a/P2rx1/P2ry1/Pik3ca/Pik3cd/Pik3r1/Pik3r2/Pik3r3/Prkaca/Prkacb/Prkci/Prkcz/Pla2g4a/Plcb1/Plcb2/Plcb3/Plcb4/Ppp1cb/Ppp1cc/Prkg1/Mapk11/Ptgs1/Rasgrp2/Rock1/Rock2/Snap23/Src/Stim1/Syk/Tbxas1/Rap1b/Vamp8/Vasp/Plcg2/Gucy1a2/Akt3/Gp6/Mapk1/Mapk13/Mapk14/Mapk3/Mapk12/Pik3cg/Pik3r5/Pla2g4e/Gucy1b1/Apbb1ip/Gucy1a1/Myl12a/Myl12b/Arhgef12/Tln2/Pik3cb |
| mmu05014 | Amyotrophic lateral sclerosis (ALS) | 0,02277225 | Apaf1/Bad/Bax/Bcl2/Bcl2l1/Bid/Cat/Casp12/Casp3/Casp9/Ccs/Cycs/Daxx/Gpx1/Gpx3/Grin2d/Ppp3ca/Ppp3cb/Ppp3cc/Ppp3r1/Mapk11/Rac1/Slc1a2/Sod1/Tnfrsf1a/Tnfrsf1b/Map2k3/Map2k6/Map3k5/Mapk13/Mapk14/Rab5a/Mapk12/Nefh/Tomm40/Tomm40l/Gpx7/Derl1/Gpx8/Als2/Gpx6 |
| mmu00500 | Starch and sucrose metabolism | 0,02441146 | Amy2a4/Amy2a3/Amy2a2/Gck/Amy2a5/Pygb/Pygl/Amy1/G6pc/Gaa/Gpi1/Gys1/Hk1/Hk2/Enpp1/Enpp3/Hkdc1/Ugp2/Gyg/Treh/Pgm2/G6pc3/Pgm2l1/Gbe1/Agl |
| mmu04929 | GnRH secretion | 0,02447374 | Arrb1/Raf1/Akt1/Akt2/Cacna1d/Cacna1g/Cga/Esr2/Kcnn2/Kcnn3/Gna11/Gnaq/Hcn1/Hcn3/Hras/Itpr1/Itpr2/Itpr3/Kcnj9/Nras/Pik3ca/Pik3cd/Pik3r1/Pik3r2/Pik3r3/Prkcb/Prkcg/Plcb1/Plcb2/Plcb3/Plcb4/Spp1/Arrb2/Trpc1/Akt3/Map2k1/Map2k2/Mapk1/Mapk3/Kiss1/Gabbr1/Pik3cb/Gper1/Kcnn1 |
| mmu04620 | Toll-like receptor signaling pathway | 0,02563263 | Ticam1/Akt1/Akt2/Tirap/Casp8/Cd14/Chuk/Fadd/Fos/Tlr3/Cxcl10/Ifnar1/Ifnar2/Ikbkb/Jun/Ly96/Nfkb1/Nfkbia/Pik3ca/Pik3cd/Pik3r1/Pik3r2/Pik3r3/Mapk11/Rac1/Rela/Ripk1/Ccl5/Spp1/Stat1/Tlr4/Cd40/Traf3/Traf6/Ticam2/Ifna13/Akt3/Tlr2/Ifna15/Map2k1/Map2k2/Map2k3/Map2k4/Map2k6/Map2k7/Map3k7/Map3k8/Mapk1/Mapk10/Mapk13/Mapk14/Mapk3/Mapk8/Mapk9/Irak4/Irf5/Mapk12/Tlr5/Irf7/Irf3/Tollip/Tbk1/Ikbke/Tab1/Tab2/Pik3cb |
| mmu04950 | Maturity onset diabetes of the young | 0,02583239 | Gck/Hes1/Hhex/Mnx1/Foxa2/Foxa3/Hnf4a/Onecut1/Bhlha15/Neurod1/Nkx2-2/Nkx6-1/Pax6/Pdx1/Pklr/Slc2a2/Hnf1a/Hnf1b/Nr5a2/Hnf4g/Rfx6 |
| mmu04915 | Estrogen signaling pathway | 0,02633725 | Adcy3/Raf1/Adcy6/Adcy8/Adcy9/Akt1/Akt2/Atf2/Atf4/Bcl2/Calm1/Calm2/Calm3/Creb1/Creb3/Atf6b/Ctsd/Egfr/Esr1/Esr2/Fkbp4/Fkbp5/Fos/Gnai1/Gnai2/Gnai3/Gnao1/Gnaq/Gnas/Grb2/Hbegf/Hras/Hspa1l/Hspa1b/Hspa2/Hsp90ab1/Hsp90aa1/Itpr1/Itpr2/Itpr3/Jun/Kcnj9/Krt10/Krt18/Krt19/Mmp9/Ncoa1/Ncoa2/Ncoa3/Nras/Pik3ca/Pik3cd/Pik3r1/Pik3r2/Pik3r3/Prkaca/Prkacb/Prkcd/Plcb1/Plcb2/Plcb3/Plcb4/Pomc/Rara/Shc1/Shc3/Sos1/Sos2/Sp1/Src/Creb3l2/Tgfa/Creb5/Akt3/Map2k1/Map2k2/Mapk1/Mapk3/Creb3l1/Shc4/Gabbr1/Ebag9/Krt25/Pik3cb/Gper1/Creb3l4/Krt23 |
| mmu04216 | Ferroptosis | 0,02703946 | Atg5/Cp/Acsl1/Fth1/Ftl1/Gclc/Gclm/Lpcat3/Gss/Hmox1/Slc3a2/Slc11a2/Pcbp2/Slc39a14/Acsl6/Tfrc/Vdac2/Vdac3/Pcbp1/Slc7a11/Ncoa4/Acsl5/Slc40a1/Gpx4/Map1lc3a/Map1lc3b/Slc39a8/Steap3/Sat2/Atg7 |
| mmu04973 | Carbohydrate digestion and absorption | 0,03084718 | Amy2a4/Amy2a3/Amy2a2/Amy2a5/Akt1/Akt2/Amy1/Atp1a1/Atp1b1/Atp1b2/Atp1b3/Cacna1d/G6pc/Slc37a4/Hk1/Hk2/Pik3ca/Pik3cd/Pik3r1/Pik3r2/Pik3r3/Prkcb/Plcb1/Plcb2/Plcb3/Plcb4/Slc2a2/Slc5a1/Hkdc1/Akt3/Slc2a5/G6pc3/Pik3cb |
| mmu00062 | Fatty acid elongation | 0,03463287 | Tecr/Elovl3/Hadh/Elovl6/Acot2/Acot3/Ppt1/Hadhb/Acot1/Mecr/Hacd1/Elovl1/Ppt2/Hsd17b12/Hacd3/Hacd4/Elovl5/Acot7/Hacd2/Elovl7/Them4/Elovl4 |
| mmu04662 | B cell receptor signaling pathway | 0,03573211 | Raf1/Akt1/Akt2/Bcl10/Cd22/Cd79a/Cd81/Chuk/Fos/Grb2/Ptpn6/Hras/Ikbkb/Inppl1/Jun/Rac3/Lyn/Nfatc1/Nfatc2/Nfatc3/Nfkb1/Nfkbia/Nfkbib/Nfkbie/Nras/Pik3ca/Pik3cd/Pik3r1/Pik3r2/Pik3r3/Prkcb/Ppp3ca/Ppp3cb/Ppp3cc/Ppp3r1/Rac1/Rela/Sos1/Sos2/Syk/Vav1/Vav2/Plcg2/Akt3/Rasgrp3/Malt1/Dapp1/Map2k1/Map2k2/Mapk1/Mapk3/Gsk3b/Vav3/Pik3cb/Pik3ap1 |
| mmu04024 | cAMP signaling pathway | 0,03618546 | Adcy3/Camk2d/Orai1/Npy/Braf/Rap1a/Raf1/Pde4c/Acox1/Adcy6/Adcy8/Adcy9/Adora1/Adora2a/Adrb1/Adrb2/Akt1/Akt2/Amh/Atp1a1/Atp1b1/Atp1b2/Atp1b3/Atp2a2/Bad/Cacna1d/Calm1/Calm2/Calm3/Camk4/Cftr/Cga/Chrm1/Creb1/Creb3/Crebbp/Edn1/Edn2/F2r/Fos/Gli1/Glp1r/Gnai1/Gnai2/Gnai3/Gnas/Grin2d/Hhip/Htr1b/Htr1d/Htr6/Jun/Lipe/Rac3/Afdn/Ppp1r12a/Nfatc1/Nfkb1/Nfkbia/Npr1/Npy1r/Oxt/Pak1/Pde3b/Pde4a/Pde4b/Pik3ca/Pik3cd/Pik3r1/Pik3r2/Pik3r3/Prkaca/Prkacb/Pld1/Pld2/Pomc/Ppara/Ppp1cb/Ppp1cc/Ppp1r1b/Ptch1/Ptger2/Ptger3/Rac1/Rela/Rock1/Rock2/Rras/Slc9a1/Sst/Sstr1/Sstr2/Sox9/Creb3l2/Rap1b/Tiam1/Tshr/Vav1/Vav2/Vipr2/Rapgef3/Creb5/Akt3/Pde4d/Abcc4/Pde10a/Grin3a/Hcar1/Chrm2/Map2k1/Map2k2/Mapk1/Mapk10/Mapk3/Mapk8/Mapk9/Creb3l1/Ep300/Hcn4/Atp2b4/Gipr/Atp2a3/Gabbr1/Fxyd1/Rapgef4/Vav3/Rras2/Atp2b1/Plce1/Pik3cb/Creb3l4/Hcar2/Acox3/Sucnr1 |
| mmu00280 | Valine, leucine and isoleucine degradation | 0,03618546 | Il4i1b/Aldh6a1/Acat1/Acat2/Aldh7a1/Pcca/Acadm/Acaa1a/Acads/Aldh2/Aldh3a2/Aox1/Auh/Bcat2/Bckdha/Bckdhb/Dbt/Dld/Hadh/Hmgcl/Hmgcs2/Mmut/Hmgcs1/Hmgcll1/Hibch/Hadhb/Acaa1b/Acsf3/Abat/Ivd/Aldh9a1/Hibadh/Acadsb/Pccb/Acad8/Aldh1b1/Ehhadh/Mccc2/Aacs |
| mmu00515 | Mannose type O-glycan biosynthesis | 0,0365124 | Fut4/B4galt1/Large1/St3gal3/Pomgnt2/Rxylt1/Large2/Fkrp/Fktn/Mgat5b/B3gat2/B4galt2/B4galt3/Pomk/Crppa/B3galnt2/Chst10/Pomt1 |
| mmu04725 | Cholinergic synapse | 0,03686279 | Adcy3/Chrnb4/Camk2d/Ache/Chrna7/Adcy6/Adcy8/Adcy9/Akt1/Akt2/Atf4/Bcl2/Cacna1a/Cacna1d/Camk4/Chrm1/Chrm3/Creb1/Creb3/Fos/Fyn/Gna11/Gnai1/Gnai2/Gnai3/Gnao1/Gnaq/Gnb1/Gnb4/Gnb5/Gng10/Gng12/Gng2/Gng5/Gng7/Gng8/Gngt2/Hras/Itpr1/Itpr2/Itpr3/Jak2/Kcnj2/Kcnj4/Kcnq1/Nras/Pik3ca/Pik3cd/Pik3r1/Pik3r2/Pik3r3/Prkaca/Prkacb/Prkcb/Prkcg/Plcb1/Plcb2/Plcb3/Plcb4/Creb3l2/Kcnq5/Creb5/Akt3/Chrm2/Map2k1/Mapk1/Mapk3/Creb3l1/Pik3cg/Pik3r5/Gng11/Pik3cb/Creb3l4 |
| mmu00330 | Arginine and proline metabolism | 0,04296654 | Amd2/Carns1/Aldh7a1/Aldh2/Aldh3a2/Arg1/Arg2/Ckb/Ckm/Ckmt1/Gamt/Got1/Got2/Nos2/Oat/Odc1/P4ha1/P4ha2/Prodh/Srm/Pycr1/Aldh4a1/Smox/Azin2/Aldh18a1/Aldh9a1/Cndp2/Pycrl/Lap3/Gatm/L3hypdh/Hoga1/Pycr2/Sat2/Aldh1b1 |
| mmu00630 | Glyoxylate and dicarboxylate metabolism | 0,04366493 | Gldc/Shmt2/Acat1/Acat2/Pcca/Aco1/Aco2/Cat/Cs/Dld/Mdh2/Mdh1/Mmut/Shmt1/Glyctk/Amt/Hao2/Pccb/Hoga1/Gcsh/Acss1/Afmid/Grhpr |
| mmu04622 | RIG-I-like receptor signaling pathway | 0,04485458 | Isg15/Atg5/Casp8/Chuk/Fadd/Cxcl10/Ikbkb/Nfkb1/Nfkbia/Nfkbib/Mapk11/Rela/Ripk1/Tank/Trim25/Traf2/Traf3/Traf6/Tkfc/Mavs/Ddx58/Ifna13/Pin1/Ifna15/Map3k1/Map3k7/Mapk10/Mapk13/Mapk14/Mapk8/Mapk9/Nlrx1/Azi2/Mapk12/Irf7/Irf3/Tbk1/Ikbke/Sike1/Rnf125/Ifih1/Tradd/Sting1/Tbkbp1/Cyld/Dhx58 |
| mmu04918 | Thyroid hormone synthesis | 0,04584601 | Adcy3/Slc5a5/Adcy6/Adcy8/Adcy9/Atf2/Atf4/Atp1a1/Atp1b1/Atp1b2/Atp1b3/Pdia4/Canx/Cga/Creb1/Creb3/Atf6b/Gnaq/Gnas/Lrp2/Gpx1/Gpx3/Gsr/Hspa5/Itpr1/Itpr2/Itpr3/Prkaca/Prkacb/Prkcb/Prkcg/Plcb1/Plcb2/Plcb3/Plcb4/Creb3l2/Duox2/Tshr/Ttf1/Creb5/Slc26a4/Creb3l1/Duoxa2/Gpx7/Gpx8/Iyd/Ttf2/Gpx6/Creb3l4 |
| mmu00270 | Cysteine and methionine metabolism | 0,04947275 | Amd2/Il4i1b/Adi1/Psat1/Cth/Mat2b/Mat1a/Bcat2/Cbs/Cdo1/Dnmt1/Dnmt3a/Dnmt3b/Gclc/Gclm/Got1/Got2/Gss/Ldha/Ldhb/Mdh2/Mdh1/Srm/Tst/Ahcyl1/Kyat3/Mat2a/Phgdh/Mtr/Sdsl/Ahcy/Apip/Mtap/Mri1/Kyat1/Ahcyl2 |
| mmu05168 | Herpes simplex virus 1 infection | 6,9795E-17 | Gm2026/Gm3055/Zfp984/Gm14308/Zfp850/Gm10778/Zfp729b/Zfp282/Zfp956/E430018J23Rik/AW146154/Zfp119a/Ticam1/Srsf9/Eif2b3/H2-Q6/H2-Q9/Srsf1/Zfp607b/Akt1/Akt2/Apaf1/Birc3/Birc2/B2m/Bad/Bak1/Bax/Bcl2/Bcl2l1/Bid/Calr/Casp3/Casp8/Casp9/Chuk/Socs3/Cycs/Daxx/Eif2s1/Eif2ak3/Eif2b4/Eif4ebp1/Fadd/Fas/Tlr3/Pdia3/H2-Ab1/H2-Bl/H2-D1/H2-Eb1/H2-K1/H2-M3/H2-DMa/H2-Q1/H2-Q10/H2-Q2/H2-Q4/H2-T22/H2-T23/H2-T24/Eif2ak1/Ifnar1/Ifnar2/Ifngr1/Ifngr2/Ikbkb/Irf9/Itga5/Jak1/Jak2/Zfp87/Zfp617/Nfkb1/Nfkbia/Pik3ca/Pik3cd/Pik3r1/Pik3r2/Pik3r3/Pml/Pou2f1/Pou2f2/Pou2f3/Ppp1cb/Ppp1cc/Eif2ak2/Ptpn11/Zfp286/Zfp184/Rela/Rheb/Ccl5/Srsf2/Srsf3/Sp100/Src/Srpk1/Zfp871/Stat1/Stat2/Eif2b1/Syk/Zfp658/Zfp719/Zfp180/Zfp677/Zfp947/Zfp748/Zfp273/Zfp583/Tap2/Tapbp/Zfp354a/Cgas/Zfp879/AU041133/Alyref/Eif2b2/Zfp455/Zfp595/Tnfrsf1a/Traf2/Traf3/Traf5/Traf6/Zfp7/Eif2b5/Zfp160/Zfp472/Zfp81/H2-M10.5/Zfp959/Srsf7/Zfp1/Zfp101/Zfp11/Zfp13/Zfp26/Zfp28/Zfp30/Zfp37/Zfp39/Zfp40/Zfp46/Zfp51/Zfp52/Zfp54/Zfp57/Zfp60/Zfp61/Zfp85/Zfp9/Zfp90/Zfp93/Zfp94/Zfp97/Mavs/Zfp334/Ddx58/Zfp189/Ifna13/Tnfrsf14/Zfp12/Zfp212/Zfp954/Zfp418/Zfp772/Zfp114/Zfp790/Zfp940/Zfp420/Zfp382/Zfp939/AI987944/Zfp768/Zfp764/Zfp958/Zfp930/Zfp868/Zfp961/Zfp612/Zfp426/Zfp809/Zfp599/Zfp810/Zfp65/Zfp825/Zfp709/Zfp938/Zfp454/Zfp867/Akt3/Zfp458/Zfp874a/Zfp58/Zfp647/Zfp641/Zfp760/Zfp994/Zfp799/Zfp870/Zfp952/Zfp563/Zfp119b/Rnasel/Tlr2/Zfp53/Zfp68/Zfp267/Ifna15/Zfp933/A430033K04Rik/Zfp128/Zfp324/Zfp568/Zfp14/Zfp473/Zfp791/Zfp317/Zfp937/Map3k7/Irak4/Zfp747/Eif2ak4/Zfp398/Zfp354b/Zfp354c/Zfp459/Zfp853/Zfp786/Zfp78/Zfp82/Zfp866/Card9/Zfp780b/Gm5141/Rsl1/Zfp948/Zfp229/Zfp69/Zfp667/Zfp456/Zfp874b/Zfp950/Zfp457/Zfp708/Zfp268/Zfp141/Zfp975/Zfp560/Zfp960/Nxf1/Irf7/Irf3/Zfp316/Zfp607a/Zfp108/Tyk2/Alyref2/Zfp386/Zfp113/Tbk1/Ikbke/Zfp235/Zfp111/Mtor/Zfp109/Srsf4/Zfp112/Nectin1/Zfp872/Zfp551/Zfp963/Zfp951/Gm14322/Gm14430/Zfp808/Zfp964/Tsc1/Gm14391/Tab1/Gm14326/Zfp991/H2-T-ps/Gm8909/Gm14434/Zfp869/Zfp442/Zfp605/Zfp788/Zfp169/Hcfc2/Srsf6/Tab2/Zfp707/Zfp746/Zfp688/Bst2/Zfp715/2810021J22Rik/Zfp619/Zfp597/Zfp689/Zfp626/Zfp935/Ifih1/Zfp251/Tradd/Zfp949/2610008E11Rik/Zfp157/Zfp661/Zfp558/Zfp777/Sting1/Zfp248/Zfp74/Zfp974/Zfp763/Zfp383/Zfp946/Zfp84/Pik3cb/Zfp773/Zfp266/Zfp712/Zfp623/9130019O22Rik |
| mmu05225 | Hepatocellular carcinoma | 5,3319E-13 | Gstt3/Ddb2/Braf/Raf1/Actb/Actg1/Akt1/Akt2/Apc/Axin1/Axin2/Bad/Bak1/Bax/Bcl2l1/Ctnnb1/Ccnd1/Cdk4/Cdk6/Cdkn1a/Cdkn2a/Gadd45a/Dvl1/Dvl2/Dvl3/E2f1/E2f3/Egfr/Frat1/Fzd1/Fzd3/Fzd4/Fzd5/Fzd6/Fzd7/Fzd8/Fzd9/Gab1/Grb2/Gsta3/Gsta4/Gstm1/Gstm2/Gstm3/Gstm4/Gstm5/Gstp1/Gstt1/Gstt2/Gsto1/Hmox1/Hras/Igf1r/Lef1/Lrp5/Lrp6/Smad2/Smad3/Smad4/Met/Myc/Gadd45b/Nfe2l2/Nqo1/Nras/Pik3ca/Pik3cd/Pik3r1/Pik3r2/Pik3r3/Prkcb/Prkcg/Plcg1/Pten/Rb1/Shc1/Shc3/Smarca4/Smarcb1/Smarcc1/Sos1/Sos2/Wnt8a/Mgst2/Frat2/Tcf7/Tcf7l1/Tcf7l2/Wnt9a/Terc/Tert/Tgfa/Tgfb1/Tgfb2/Tgfb3/Tgfbr1/Tgfbr2/Wnt1/Wnt10b/Wnt11/Wnt9b/Wnt2b/Wnt5a/Wnt5b/Wnt6/Gstp3/Txnrd3/Plcg2/Akt3/Apc2/Gadd45g/Arid1b/E2f2/Map2k1/Map2k2/Mapk1/Mapk3/Brd7/Polk/Shc4/Peg12/Dpf1/Txnrd1/Keap1/Actl6a/Mgst1/Gsk3b/Mtor/Fzd2/Smarce1/Rps6kb2/Mgst3/Smarcd3/Smarca2/Gsto2/Gstm7/Dpf3/Phf10/Pik3cb/Smarcd2/Smarcd1/Csnk1a1/Arid1a |
| mmu04141 | Protein processing in endoplasmic reticulum | 2,5885E-12 | Dnajc3/Rpn1/Txndc5/Ssr1/Edem2/Selenos/Sec13/Atxn3/Derl2/Atf4/Bag1/Bak1/Bax/Bcl2/Hyou1/Pdia4/Calr/Canx/Capn1/Capn2/Atf6b/Cryaa/Cryab/Dnajc5/Ddost/Dnajc1/Eif2s1/Eif2ak3/Ube2j2/Sec63/Ganab/Pdia3/Hspa5/Eif2ak1/Hspa1l/Dnaja1/Hsph1/Hspa1b/Hspa2/Hsp90ab1/Hsp90aa1/Stt3a/Man1a2/Ppp1r15a/Nfe2l2/Hspa4l/P4hb/Plaa/Prkcsh/Eif2ak2/Edem1/Rad23a/Rad23b/Rnf185/Rpn2/Sar1a/Sec23a/Sec61g/Sel1l/Bag2/Skp1/Ube2d1/Ckap4/Os9/Nploc4/Sec24c/Traf2/Ube2g2/Marchf6/Wfs1/Xbp1/Atf6/Man1b1/Man1c1/Fbxo2/Amfr/Sec31b/Map2k7/Map3k5/Mapk10/Mapk8/Mapk9/Vcp/Cul1/Sec23b/Eif2ak4/Uggt1/Nsfl1c/Ero1a/Fbxo6/Prkn/Preb/Sec61a1/Rnf5/Ubqln1/Stub1/Rbx1/Dnaja2/Mbtps1/Ube2d2a/Dnajb12/Dnajb2/Mogs/Sec61a2/Ngly1/Ube4b/Herpud1/Hspbp1/Ssr2/Sar1b/Uggt2/Ubxn6/Dnajc10/Lman2/Ube2g1/Erp29/Ssr3/Ero1b/Derl1/Dnajb11/Stt3b/Sec31a/Sec62/Sec24d/Lman1/Derl3/Pdia6/Tram1/Syvn1/Svip/Sec24a/Ern1/Tusc3/Sil1/Rrbp1/Sec24b |
| mmu04140 | Autophagy - animal | 5,4136E-12 | Rubcn/Atg14/Prkaa1/Prkaa2/Raf1/Akt1/Akt2/Atg5/Bad/Bcl2/Bcl2l1/Bnip3/Rb1cc1/Cflar/Ctsb/Ctsd/Ctsl/Dapk2/Dapk3/Eif2s1/Eif2ak3/Hif1a/Hmgb1/Hras/Igf1r/Irs1/Irs3/Itpr1/Lamp1/Mtmr4/Rab8a/Mras/Nras/Sqstm1/Pdpk1/Pik3ca/Pik3cd/Pik3r1/Pik3r2/Pik3r3/Prkaca/Prkacb/Prkcd/Ppp2ca/Ppp2cb/Pten/Rab1a/Rab33b/Rab7/Rheb/Rras/Camkk2/Stk11/Tank/Atg9b/Zfyve1/Wdr41/Traf6/Ulk1/Vamp8/Pik3c3/Ambra1/Atg4d/Smcr8/Akt3/Atg4c/Atg9a/Map2k1/Map2k2/Map3k7/Mapk1/Mapk10/Mapk3/Mapk8/Mapk9/Eif2ak4/Ulk2/Atg2a/Irs2/Rragd/Wipi1/Rragc/Sh3glb1/Becn1/Tbk1/Gabarap/Mlst8/Mtor/Supt20/Rps6kb2/Nrbf2/Tsc1/Atg4b/Atg10/Rras2/Akt1s1/Stx17/Atg3/Atg101/Rraga/Trp53inp2/Dapk1/C9orf72/Atg16l2/Atg7/Mtmr3/Rptor/Ddit4/Pik3cb/Wipi2/Pik3r4/Atg16l1/Uvrag/Ern1/Gabarapl2/Mtmr14/Deptor |
| mmu04110 | Cell cycle | 6,2289E-12 | E2f4/Espl1/Cdc20/Abl1/Atm/Bub1/Bub1b/Bub3/Ccna2/Ccnb2/Ccnd1/Ccnd2/Ccnd3/Ccne1/Ccne2/Cdc25a/Cdc25b/Cdc25c/Cdk1/Cdc45/Cdc7/Cdk2/Cdk4/Cdk6/Cdk7/Cdkn1a/Cdkn1b/Cdkn1c/Cdkn2a/Cdkn2b/Cdkn2c/Cdkn2d/Chek1/Crebbp/Smc3/Gadd45a/E2f1/E2f3/E2f5/Hdac2/Mad1l1/Smad2/Smad3/Smad4/Mcm3/Mcm2/Mcm4/Mcm5/Mcm6/Anapc1/Mdm2/Myc/Gadd45b/Orc1/Orc2/Pcna/Plk1/Rad21/Rb1/Rbl1/Rbl2/Stag1/Tfdp2/Skp1/Tfdp1/Tgfb1/Tgfb2/Tgfb3/Cdc14b/Ttk/Wee1/Ywhae/Ywhag/Ywhah/Ywhaq/Ywhaz/Zbtb17/Cdc14a/Cdc6/Gadd45g/E2f2/Atr/Orc5/Ccnb1/Pkmyt1/Cul1/Dbf4/Skp2/Pttg1/Ep300/Hdac1/Anapc4/Cdc23/Ywhab/Sfn/Mad2l1/Anapc7/Fzr1/Rbx1/Orc6/Gsk3b/Anapc5/Cdc26/Ccnh/Cdc16/Mad2l2 |
| mmu05220 | Chronic myeloid leukemia | 6,2289E-12 | Ddb2/Braf/Raf1/Bcr/Abl1/Akt1/Akt2/Bad/Bak1/Bax/Bcl2l1/Runx1/Cbl/Ccnd1/Cdk4/Cdk6/Cdkn1a/Cdkn1b/Cdkn2a/Chuk/Crk/Crkl/Ctbp1/Ctbp2/Gadd45a/E2f1/E2f3/Mecom/Gab2/Grb2/Hdac2/Hras/Ikbkb/Smad3/Smad4/Mdm2/Myc/Gadd45b/Nfkb1/Nfkbia/Nras/Pik3ca/Pik3cd/Pik3r1/Pik3r2/Pik3r3/Ptpn11/Rb1/Rela/Shc1/Shc3/Sos1/Sos2/Stat5a/Stat5b/Tgfb1/Tgfb2/Tgfb3/Tgfbr1/Tgfbr2/Akt3/Gadd45g/E2f2/Map2k1/Map2k2/Mapk1/Mapk3/Polk/Shc4/Hdac1/Pik3cb |
| mmu04144 | Endocytosis | 9,9131E-12 | Ldlrap1/Dnm3/Chmp7/Rab31/Vps37c/Gbf1/Wwp1/Arrb1/Grk2/H2-Q6/H2-Q9/Vps4a/Ap2a1/Ap2a2/Ap2m1/Arf1/Arf3/Arf5/Arf6/Arpc1b/Capza1/Capza2/Capzb/Cav1/Cav2/Cbl/Cdc42/Clta/Cxcr4/Dab2/Asap1/Dnm1/Dnm2/Egfr/Ehd1/Epn1/Epn2/Eps15/Eps15l1/Acap3/Fgfr2/Fgfr3/Fgfr4/Grk4/Grk5/H2-Bl/H2-D1/H2-K1/H2-M3/H2-Q1/H2-Q10/H2-Q2/H2-Q4/H2-T22/H2-T23/H2-T24/Hgs/Hras/Hspa1l/Hspa1b/Hspa2/Igf1r/Igf2r/Itch/Kif5a/Kif5b/Kif5c/Ldlr/Smad2/Smad3/Mdm2/Rab8a/Nedd4/Pdcd6ip/Pdgfra/Pip5k1c/Pip5k1b/Pip5k1a/Prkci/Prkcz/Pld1/Pld2/Pml/Cyth1/Cyth2/Cyth3/Rab10/Rab11b/Rab4a/Rab5b/Rab5c/Rab7/Vps37d/Sh3gl2/Sh3gl1/Vps4b/Src/Chmp6/Stam/Cblb/Arfgef1/Asap2/Arap2/Agap3/Wipf1/Rab11fip3/Psd4/Eea1/Rufy1/Arrb2/Git1/Tgfbr1/Tgfbr2/Zfyve16/Traf6/Tfrc/Tsg101/Ubb/Washc5/Vps45/H2-M10.5/Snx32/Sh3glb2/Pip5kl1/Arfgap1/Spg20/Zfyve9/Asap3/Iqsec1/Ap2s1/Psd3/Chmp1a/Grk6/Git2/Rab11fip4/Rab5a/Snf8/Spg21/Washc2/Vps25/Vps26a/Bin1/Washc4/Zfyve27/Grk3/Vps37b/Agap1/Rab11fip5/Rab11a/Rabep1/Snx3/Sh3glb1/Stam2/Arpc3/Snx1/Ehd3/Pard6b/Arfgap3/Smurf2/Chmp4c/Chmp3/H2-T-ps/Gm8909/Vps28/Chmp1b/Washc3/Cltc/Rnf41/Snx2/Wipf2/Washc1/Chmp2b/Chmp2a/Vps26b/Snx4/Snx5/Arap1/Smap2/Vps36/Stambp/Epn3/Ist1/Snx6/Mvb12b/Dnajc6/Wasl/Mvb12a/Psd2/Cltb/Rab11fip2/Chmp4b/Rab11fip1/Smurf1/Arpc2/Arfgap2/Rbsn/Acap2/Cblc/Nedd4l/Usp8/Pard6g/Pard3/Smap1/Ehd4/Arfgef2 |
| mmu05212 | Pancreatic cancer | 4,6133E-11 | Ddb2/Braf/Raf1/Akt1/Akt2/Bad/Bak1/Bax/Bcl2l1/Brca2/Casp9/Ccnd1/Cdc42/Cdk4/Cdk6/Cdkn1a/Cdkn2a/Chuk/Gadd45a/E2f1/E2f3/Egf/Egfr/Erbb2/Ikbkb/Jak1/Rac3/Smad2/Smad3/Smad4/Gadd45b/Nfkb1/Pik3ca/Pik3cd/Pik3r1/Pik3r2/Pik3r3/Pld1/Pld2/Rac1/Rac2/Rad51/Rb1/Rela/Ralgds/Ralbp1/Stat1/Stat3/Tgfa/Tgfb1/Tgfb2/Tgfb3/Tgfbr1/Tgfbr2/Vegfa/Akt3/Gadd45g/E2f2/Map2k1/Mapk1/Mapk10/Mapk3/Mapk8/Mapk9/Polk/Rala/Mtor/Rps6kb2/Ralb/Pik3cb |
| mmu04120 | Ubiquitin mediated proteolysis | 4,6133E-11 | Ube3c/Fbxw11/Nhlrc1/Fbxo4/Wwp1/Ddb2/Cdc20/Ube2q2/Aire/Ube3b/Birc3/Birc2/Brca1/Birc6/Btrc/Cbl/Socs3/Socs1/Ddb1/Ube2j2/Ubox5/Ube4a/Trip12/Herc2/Itch/Anapc1/Mgrn1/Mdm2/Pias2/Nedd4/Pml/Rnf7/Siah1a/Cblb/Skp1/Ube2d1/Cdc34/Ube2o/Ube2e2/Traf6/Ube2m/Ube2e3/Ube2e1/Ube2l3/Ube2i/Uba3/Ube2b/Ube2g2/Ube2h/Ube3a/Vhl/Pias3/Fbxo2/Uba6/Fbxw8/Herc1/Klhl9/Rhobtb2/Cop1/Map3k1/Cul3/Ube2z/Cul1/Skp2/Prpf19/Fbxw7/Keap1/Prkn/Uba2/Anapc4/Cdc23/Ube2k/Anapc7/Fzr1/Stub1/Rbx1/Sae1/Pias1/Ube2d2a/Ube2l6/Pias4/Anapc5/Ube4b/Ppil2/Smurf2/Cdc26/Cul7/Ube2w/Wwp2/Fancl/Ube2g1/Herc4/Ube2r2/Elob/Ube2f/Ube2c/Trim37/Rhobtb1/Trim32/Cdc16/Ube2q1/Ubr5/Cul2/Ercc8/Herc3/Syvn1/Uba7/Cul5/Smurf1/Det1/Ube2ql1/Ube2s/Cblc/Nedd4l/Ube2n |
| mmu04550 | Signaling pathways regulating pluripotency of stem cells | 1,6822E-10 | Raf1/Acvr1/Acvr1b/Acvr2a/Acvr2b/Akt1/Akt2/Apc/Zfhx3/Axin1/Axin2/Bmi1/Bmpr1a/Bmpr1b/Bmpr2/Ctnnb1/Dvl1/Dvl2/Dvl3/Lefty1/Smarcad1/Fgf2/Fgfr1/Fgfr2/Fgfr3/Fgfr4/Fzd1/Fzd3/Fzd4/Fzd5/Fzd6/Fzd7/Fzd8/Fzd9/Grb2/Hand1/Hesx1/Onecut1/Hras/Id1/Id2/Id3/Id4/Igf1/Igf1r/Il6st/Inhba/Inhbb/Inhbe/Isl1/Jak1/Jak2/Jak3/Jarid2/Klf4/Lhx5/Lif/Lifr/Smad1/Smad2/Smad3/Smad4/Smad5/Meis1/Myc/Nras/Otx1/Pax6/Pik3ca/Pik3cd/Pik3r1/Pik3r2/Pik3r3/Mapk11/Rest/Skil/Stat3/Wnt8a/Tbx3/Tcf7/Tcf3/Wnt9a/Wnt1/Wnt10b/Wnt11/Wnt9b/Wnt2b/Wnt5a/Wnt5b/Wnt6/Pcgf2/Akt3/Apc2/Kat6a/Esrrb/Map2k1/Map2k2/Mapk1/Mapk13/Mapk14/Mapk3/Acvr1c/Mapk12/Lefty2/Rif1/Smad9/Gsk3b/Fzd2/Pcgf3/Pcgf1/Pcgf6/Pik3cb/Pcgf5/Setdb1 |
| mmu05210 | Colorectal cancer | 2,5062E-10 | Ddb2/Braf/Raf1/Akt1/Akt2/Apc/Birc5/Areg/Axin1/Axin2/Bad/Bak1/Bax/Bcl2/Bcl2l11/Casp3/Casp9/Ctnnb1/Ccnd1/Cdkn1a/Cycs/Gadd45a/Egf/Egfr/Fos/Grb2/Hras/Jun/Lef1/Rac3/Bbc3/Smad2/Smad3/Smad4/Mlh1/Msh2/Msh6/Myc/Gadd45b/Nras/Pik3ca/Pik3cd/Pik3r1/Pik3r2/Pik3r3/Rac1/Rac2/Ralgds/Sos1/Sos2/Tcf7/Tcf7l1/Tcf7l2/Tgfa/Tgfb1/Tgfb2/Tgfb3/Tgfbr1/Tgfbr2/Akt3/Apc2/Gadd45g/Map2k1/Map2k2/Mapk1/Mapk10/Mapk3/Mapk8/Mapk9/Polk/Rala/Gsk3b/Mtor/Pmaip1/Rps6kb2/Ralb/Appl1/Pik3cb |
| mmu05161 | Hepatitis B | 2,7514E-10 | Ticam1/Ddb2/Braf/Raf1/Akt1/Akt2/Tirap/Apaf1/Birc5/Atf2/Atf4/Bad/Bax/Bcl2/Bid/Casp3/Casp8/Casp9/Ccna2/Ccne1/Ccne2/Cdk2/Cdkn1a/Chuk/Creb1/Creb3/Crebbp/Atf6b/Cycs/Ddb1/E2f1/E2f3/Egr2/Egr3/Fadd/Fas/Fos/Tlr3/Grb2/Hras/Hspg2/Ifnar1/Ikbkb/Jak1/Jak2/Jak3/Jun/Smad3/Smad4/Mmp9/Myc/Nfatc1/Nfatc2/Nfatc3/Nfkb1/Nfkbia/Nras/Pcna/Pik3ca/Pik3cd/Pik3r1/Pik3r2/Pik3r3/Prkcb/Prkcg/Mapk11/Ptk2b/Rb1/Rela/Sos1/Sos2/Src/Stat1/Stat2/Stat3/Stat4/Stat5a/Stat5b/Stat6/Creb3l2/Tgfb1/Tgfb2/Tgfb3/Tgfbr1/Tgfbr2/Tlr4/Traf3/Traf6/Vdac3/Ticam2/Ywhaq/Ywhaz/Mavs/Ddx58/Ifna13/Creb5/Akt3/Tlr2/Ifna15/E2f2/Map2k1/Map2k2/Map2k4/Map2k6/Map2k7/Map3k1/Map3k7/Mapk1/Mapk10/Mapk13/Mapk14/Mapk3/Mapk8/Mapk9/Creb3l1/Irak4/Mapk12/Ep300/Irf7/Irf3/Ywhab/Tyk2/Tbk1/Ikbke/Tab1/Tab2/Ifih1/Nfatc4/Pik3cb/Creb3l4 |
| mmu04068 | FoxO signaling pathway | 6,088E-10 | Prkaa1/Prkaa2/Prkab2/Prkag2/Braf/Raf1/Akt1/Akt2/Atm/Bcl6/Bcl2l11/Bnip3/Cat/Ccnb2/Ccnd1/Ccnd2/Ccng2/Cdk2/Cdkn1a/Cdkn1b/Cdkn2b/Cdkn2d/Chuk/Plk3/Crebbp/Gadd45a/S1pr1/Egf/Egfr/G6pc/Grb2/Foxg1/Hras/Igf1/Igf1r/Ikbkb/Insr/Irs1/Irs3/Klf2/Sgk3/Smad3/Smad4/Mdm2/Gadd45b/Nlk/Nras/Pdpk1/Pik3ca/Pik3cd/Pik3r1/Pik3r2/Pik3r3/Plk1/Prkab1/Prkag1/Mapk11/Pten/Rbl2/Sgk1/Slc2a4/Plk2/Sos1/Sos2/Stat3/Stk11/Plk4/Tgfb1/Tgfb2/Tgfb3/Tgfbr1/Tgfbr2/Tnfsf10/Akt3/Gadd45g/Usp7/Map2k1/Map2k2/Mapk1/Mapk10/Mapk13/Mapk14/Mapk3/Mapk8/Mapk9/Homer1/Homer2/Homer3/Ccnb1/Sgk2/Csnk1e/Skp2/Mapk12/Ep300/Foxo6/Irs2/Foxo1/Foxo3/Gabarap/Stk4/Fbxo25/Fbxo32/G6pc3/Setd7/Pck2/Pik3cb/Gabarapl2/Sirt1 |
| mmu04010 | MAPK signaling pathway | 6,1766E-10 | Mapkapk3/Arrb1/Braf/Rap1a/Raf1/Rasa2/Angpt1/Angpt2/Akt1/Akt2/Areg/Atf2/Atf4/Cacna1a/Cacna1c/Cacna1d/Cacna1e/Cacna1g/Cacna2d1/Cacnb1/Cacnb2/Cacnb3/Cacnb4/Casp3/Cd14/Cdc25b/Cdc42/Chuk/Crk/Crkl/Csf1/Daxx/Gadd45a/Dusp2/Efna1/Efna2/Efna3/Efna4/Efna5/Egf/Egfr/Elk4/Epha2/Erbb2/Erbb3/Erbb4/Mecom/Fas/Fgf1/Fgf15/Fgf18/Fgf2/Fgf5/Fgf8/Fgf9/Fgfr1/Fgfr2/Fgfr3/Fgfr4/Flt3l/Flt4/Fos/Gna12/Gng12/Grb2/Nr4a1/Hras/Hspa1l/Hspb1/Hspa1b/Hspa2/Igf1/Igf1r/Ikbkb/Il1r1/Il1rap/Insr/Jun/Jund/Kdr/Kit/Stmn1/Rac3/Mapkapk2/Mapkapk5/Max/Mef2c/Met/Kitl/Mknk1/Mknk2/Mras/Mapt/Myc/Gadd45b/Nf1/Nfatc1/Nfatc3/Nfkb1/Nfkb2/Nlk/Nras/Dusp8/Pak1/Pdgfa/Pdgfb/Pdgfra/Pgf/Prkaca/Prkacb/Prkcb/Prkcg/Pla2g4a/Ppm1a/Ppm1b/Ppp3ca/Ppp3cb/Ppp3cc/Ppp3r1/Ppp5c/Mapk11/Mapk8ip1/Dusp1/Ptprr/Rac1/Rac2/Rasgrp2/Rasgrf1/Rela/Relb/Rps6ka1/Rps6ka2/Rras/Sos1/Sos2/Rap1b/Arrb2/Tek/Taok1/Tgfa/Tgfb1/Tgfb2/Tgfb3/Tgfbr1/Tgfbr2/Rasa1/Tnfrsf1a/Traf2/Traf6/Vegfa/Vegfb/Vegfc/Pak2/Map4k3/Dusp7/Akt3/Gadd45g/Map2k5/Mapk7/Rasgrp3/Dusp5/Map2k1/Map2k2/Map2k4/Map2k6/Map2k7/Map3k1/Map3k12/Map3k2/Map3k3/Map3k5/Map3k7/Map3k8/Map4k1/Map4k2/Mapk1/Mapk10/Mapk13/Mapk14/Mapk3/Mapk8/Mapk9/Irak4/Map4k4/Ecsit/Flnb/Mapk12/Mapk8ip3/Dusp4/Taok3/Taok2/Map3k6/Map3k14/Pdgfc/Stk3/Rps6ka4/Lamtor3/Cacna2d2/Cacna1h/Stk4/Dusp10/Map3k20/Tab1/Rras2/Dusp6/Tab2/Flnc/Dusp16/Tradd/Map3k13/Pdgfd/Dusp3/Rps6ka5/Rapgef2/Ntf5/Cacng7 |
| mmu04152 | AMPK signaling pathway | 1,079E-09 | Acacb/Prkaa1/Acaca/Prkaa2/Prkab2/Prkag2/Akt1/Akt2/Cab39/Ccna2/Ccnd1/Cd36/Cftr/Cpt1a/Creb1/Creb3/Eef2/Eef2k/Eif4ebp1/Fasn/Fbp2/Fbp1/G6pc/Gys1/Hmgcr/Hnf4a/Elavl1/Igf1/Igf1r/Insr/Irs1/Irs3/Lipe/Pfkfb3/Rab8a/Pdpk1/Pfkfb2/Pfkl/Pfkm/Pik3ca/Pik3cd/Pik3r1/Pik3r2/Pik3r3/Pparg/Ppargc1a/Ppp2ca/Ppp2cb/Prkab1/Prkag1/Rab10/Rab11b/Rheb/Scd1/Scd2/Slc2a4/Camkk2/Srebf1/Creb3l2/Stk11/Ppp2r5d/Ulk1/Ppp2r5b/Ppp2r5a/Stradb/Creb5/Ppp2r3a/Akt3/Map3k7/Creb3l1/Ppp2r5c/Ppp2r5e/Pfkfb4/Scd3/Irs2/Ppp2r1a/Ppp2r2d/Pfkp/Foxo1/Foxo3/Mlycd/Mtor/Tbc1d1/Rps6kb2/Rab2a/Ppp2r3c/Tsc1/Akt1s1/Rab14/G6pc3/Adipor2/Cab39l/Ppp2r2a/Strada/Adipor1/Ppp2r2b/Ppp2r1b/Crtc2/Rptor/Pck2/Pik3cb/Cpt1c/Creb3l4/Sirt1 |
| mmu04390 | Hippo signaling pathway | 1,3612E-09 | Fbxw11/Csnk1d/Scrib/Actb/Actg1/Amh/Apc/Birc2/Birc5/Areg/Axin1/Axin2/Bmp2/Bmp7/Bmp8a/Bmpr1a/Bmpr1b/Bmpr2/Btrc/Ctnna1/Ctnnb1/Ccnd1/Ccnd2/Ccnd3/Cdh1/Patj/Dlg1/Dlg4/Dvl1/Dvl2/Dvl3/Fgf1/Fzd1/Fzd3/Fzd4/Fzd5/Fzd6/Fzd7/Fzd8/Fzd9/Gli2/Id1/Id2/Ajuba/Lats1/Lef1/Llgl1/Bbc3/Crb1/Smad1/Smad2/Smad3/Smad4/Smad7/Myc/Nf2/Prkci/Prkcz/Ppp1cb/Ppp1cc/Ppp2ca/Ppp2cb/Snai2/Wnt8a/Trp53bp2/Wwc1/Tcf7/Tcf7l1/Tcf7l2/Ctnna3/Tead1/Tead2/Tead3/Tead4/Wnt9a/Llgl2/Tgfb1/Tgfb2/Tgfb3/Tgfbr1/Tgfbr2/Trp73/Wnt1/Wnt10b/Wnt11/Wnt9b/Wnt2b/Wnt5a/Wnt5b/Wnt6/Yap1/Ywhae/Ywhag/Ywhah/Ywhaq/Ywhaz/Apc2/Dlg2/Crb2/Gdf6/Csnk1e/Limd1/Frmd6/Lats2/Ppp2r1a/Ppp2r2d/Ywhab/Pals1/Stk3/Rassf1/Gsk3b/Fzd2/Pard6b/Sav1/Mob1b/Ppp2r2a/Ppp2r2b/Rassf6/Ppp2r1b/Pard6g/Pard3/Nkd1/Wwtr1 |
| mmu01522 | Endocrine resistance | 2,7228E-09 | Adcy3/Braf/Raf1/Adcy6/Adcy9/Akt1/Akt2/Bad/Bax/Bcl2/Bik/Ccnd1/Cdk4/Cdkn1a/Cdkn1b/Cdkn2a/Cdkn2c/Dll1/Dll3/E2f1/E2f3/Egfr/Erbb2/Esr1/Esr2/Ptk2/Fos/Gnas/Grb2/Hbegf/Hras/Igf1/Igf1r/Jag1/Jun/Mdm2/Mmp9/Ncoa3/Notch1/Notch2/Notch3/Nras/Pik3ca/Pik3cd/Pik3r1/Pik3r2/Pik3r3/Prkaca/Prkacb/Med1/Mapk11/Rb1/Ncor1/Shc1/Shc3/Sos1/Sos2/Sp1/Src/Adcy2/Akt3/E2f2/Map2k1/Map2k2/Mapk1/Mapk10/Mapk13/Mapk14/Mapk3/Mapk8/Mapk9/Shc4/Abcb11/Mapk12/Dll4/Mtor/Rps6kb2/Carm1/Pik3cb/Gper1 |
| mmu05223 | Non-small cell lung cancer | 3,6347E-09 | Ddb2/Braf/Raf1/Akt1/Akt2/Bad/Bak1/Bax/Casp9/Ccnd1/Cdk4/Cdk6/Cdkn1a/Cdkn2a/Gadd45a/E2f1/E2f3/Egf/Egfr/Erbb2/Fhit/Grb2/Hras/Jak3/Gadd45b/Nras/Pdpk1/Pik3ca/Pik3cd/Pik3r1/Pik3r2/Pik3r3/Prkcb/Prkcg/Plcg1/Rb1/Rxra/Rxrg/Sos1/Sos2/Stat3/Stat5a/Stat5b/Tgfa/Rarb/Plcg2/Akt3/Gadd45g/E2f2/Map2k1/Map2k2/Mapk1/Mapk3/Polk/Rassf5/Rassf1/Foxo3/Stk4/Pik3cb/Eml4 |
| mmu04150 | mTOR signaling pathway | 3,7276E-09 | Prkaa1/Prkaa2/Atp6v1h/Prr5/Braf/Raf1/Sec13/Akt1/Akt2/Atp6v1a/Atp6v1e1/Cab39/Chuk/Dvl1/Dvl2/Dvl3/Eif4ebp1/Lpin1/Fzd1/Fzd3/Fzd4/Fzd5/Fzd6/Fzd7/Fzd8/Fzd9/Grb10/Grb2/Hras/Igf1/Igf1r/Ikbkb/Insr/Irs1/Lrp5/Lrp6/Nprl3/Slc3a2/Nras/Pdpk1/Pik3ca/Pik3cd/Pik3r1/Pik3r2/Pik3r3/Prkcb/Prkcg/Pten/Rheb/Rps6/Rps6ka1/Rps6ka2/Sgk1/Slc7a5/Sos1/Sos2/Stk11/Wnt8a/Fnip1/Wnt9a/Tnfrsf1a/Ulk1/Wnt1/Wnt10b/Wnt11/Wnt9b/Wnt2b/Wnt5a/Wnt5b/Wnt6/Stradb/Mapkap1/Sesn2/Akt3/Mios/Map2k1/Map2k2/Mapk1/Mapk3/Slc38a9/Wdr24/Eif4e2/Skp2/Depdc5/Ulk2/Wdr59/Fnip2/Rragd/Rragc/Nprl2/Clip1/Gsk3b/Lamtor3/Mlst8/Mtor/Fzd2/Rps6kb2/Tsc1/Lamtor4/Atp6v1f/Atp6v1g1/Atp6v1c1/Lamtor1/Tbc1d7/Akt1s1/Rraga/Lamtor5/Atp6v1c2/Cab39l/Telo2/Castor1/Seh1l/Strada/Rptor/Ddit4/Pik3cb/Atp6v1e2/Eif4b/Rictor/Castor2/Lamtor2/Deptor |
| mmu05224 | Breast cancer | 4,8314E-09 | Ddb2/Braf/Raf1/Akt1/Akt2/Apc/Axin1/Axin2/Bak1/Bax/Brca1/Brca2/Ctnnb1/Ccnd1/Cdk4/Cdk6/Cdkn1a/Gadd45a/Dll1/Dll3/Dvl1/Dvl2/Dvl3/E2f1/E2f3/Egf/Egfr/Erbb2/Esr1/Esr2/Fgf1/Fgf15/Fgf18/Fgf2/Fgf5/Fgf8/Fgf9/Fgfr1/Flt4/Fos/Frat1/Fzd1/Fzd3/Fzd4/Fzd5/Fzd6/Fzd7/Fzd8/Fzd9/Grb2/Hes1/Hes5/Hey1/Hey2/Hras/Igf1/Igf1r/Jag1/Jun/Kit/Lef1/Lrp5/Lrp6/Myc/Gadd45b/Ncoa1/Ncoa3/Nfkb2/Notch1/Notch2/Notch3/Nras/Pik3ca/Pik3cd/Pik3r1/Pik3r2/Pik3r3/Pten/Rb1/Shc1/Shc3/Sos1/Sos2/Sp1/Wnt8a/Frat2/Tcf7/Tcf7l1/Tcf7l2/Wnt9a/Wnt1/Wnt10b/Wnt11/Wnt9b/Wnt2b/Wnt5a/Wnt5b/Wnt6/Akt3/Apc2/Gadd45g/E2f2/Map2k1/Map2k2/Mapk1/Mapk3/Polk/Shc4/Peg12/Dll4/Heyl/Gsk3b/Mtor/Fzd2/Rps6kb2/Pik3cb/Csnk1a1 |
| mmu04360 | Axon guidance | 5,2337E-09 | Unc5a/Unc5b/Camk2d/Sema3d/Raf1/Abl1/Srgap1/Rhod/Bmp7/Bmpr1b/Bmpr2/Cdc42/Cdk5/Cfl1/Cfl2/Cxcr4/Dpysl2/Efna1/Efna2/Efna3/Efna4/Efna5/Efnb2/Enah/Epha1/Epha2/Epha3/Epha4/Epha5/Epha7/Ephb3/Ephb4/Ephb6/Plxnb2/Ptk2/Fes/Srgap2/Fyn/Fzd3/Gnai1/Gnai2/Gnai3/Hras/Itgb1/Limk1/Limk2/Rac3/Ntng2/Met/Nck1/Nck2/Neo1/Nfatc2/Nfatc3/Nras/Nrp1/Ntn1/Ntn3/Pak1/Pik3ca/Pik3cd/Pik3r1/Pik3r2/Pik3r3/Prkcz/Plcg1/Plxna1/Plxna2/Ppp3ca/Ppp3cb/Ppp3cc/Ppp3r1/Ptch1/Ptpn11/Rac1/Rac2/Robo3/Rock1/Rock2/Rras/Ryk/Cxcl12/Sema3a/Sema3c/Sema3f/Sema4a/Sema4b/Sema4c/Sema4d/Sema4f/Sema6a/Sema6b/Sema6c/Sema7a/Shh/Slit2/Slit3/Src/Unc5d/Pak6/Sema6d/Rasa1/Trpc1/Unc5c/Rnd1/Pak2/Wnt5a/Wnt5b/Ablim1/Pdk1/Ablim2/Ssh1/Plcg2/Plxnb1/Ssh2/Plxna4/Rgma/Ssh3/Srgap3/Mapk1/Mapk3/Sema4g/Robo2/Ephb1/Ablim3/Smo/Rgs3/Plxnc1/Gsk3b/Ntn4/Pard6b/Dpysl5/Myl12b/Arhgef12/Nfatc4/Pik3cb/Ntng1/Pard6g/Pard3 |
| mmu04218 | Cellular senescence | 6,8908E-09 | Traf3ip2/Fbxw11/E2f4/Raf1/H2-Q6/H2-Q9/Akt1/Akt2/Slc25a4/Atm/Zfp36l1/Zfp36l2/Btrc/Cacna1d/Calm1/Calm2/Calm3/Capn1/Capn2/Ccna2/Ccnb2/Ccnd1/Ccnd2/Ccnd3/Ccne1/Ccne2/Cdc25a/Cdk1/Cdk2/Cdk4/Cdk6/Cdkn1a/Cdkn2a/Cdkn2b/Chek1/Gadd45a/E2f1/E2f3/E2f5/Eif4ebp1/Foxm1/Gata4/H2-Bl/H2-D1/H2-K1/H2-M3/H2-Q1/H2-Q10/H2-Q2/H2-Q4/H2-T22/H2-T23/H2-T24/Hipk1/Hipk2/Hipk3/Hras/Hus1/Itpr1/Itpr2/Itpr3/Smad2/Smad3/Mapkapk2/Mdm2/Mras/Mre11a/Mybl2/Myc/Gadd45b/Nfatc1/Nfatc2/Nfatc3/Nfkb1/Nras/Sqstm1/Pik3ca/Pik3cd/Pik3r1/Pik3r2/Pik3r3/Ppp1cb/Ppp1cc/Ppp3ca/Ppp3cb/Ppp3cc/Ppp3r1/Mapk11/Pten/Rad1/Rad50/Rad9a/Rb1/Rbbp4/Rbl1/Rbl2/Rela/Rheb/Rras/Mcu/Tgfb1/Tgfb2/Tgfb3/Tgfbr1/Tgfbr2/Vdac1/Vdac2/Vdac3/H2-M10.5/Lin54/Rad9b/Akt3/Ets1/Gadd45g/E2f2/Atr/Map2k1/Map2k2/Map2k6/Mapk1/Mapk13/Mapk14/Mapk3/Ccnb1/Nbn/Mapk12/Rassf5/Foxo1/Foxo3/Mtor/Trpm7/Trpv4/Tsc1/H2-T-ps/Gm8909/Rras2/Ppid/Lin9/Nfatc4/Pik3cb/Lin37/Sirt1 |
| mmu04910 | Insulin signaling pathway | 8,3139E-09 | Acacb/Phkb/Gck/Rhoq/Ppp1r3e/Prkaa1/Trip10/Acaca/Rapgef1/Prkaa2/Prkab2/Prkag2/Braf/Pygb/Pygl/Raf1/Akt1/Akt2/Bad/Calm1/Calm2/Calm3/Cbl/Socs3/Socs1/Crk/Crkl/Eif4ebp1/Fasn/Fbp2/Fbp1/Flot1/Flot2/G6pc/Grb2/Gys1/Hk1/Hk2/Hras/Ikbkb/Inppl1/Insr/Irs1/Irs3/Lipe/Mknk1/Mknk2/Nras/Pde3b/Pdpk1/Pik3ca/Pik3cd/Pik3r1/Pik3r2/Pik3r3/Prkaca/Prkacb/Prkci/Prkcz/Pklr/Ppargc1a/Ppp1cb/Ppp1cc/Inpp5k/Prkab1/Prkag1/Prkar1a/Prkar1b/Prkar2a/Prkar2b/Ptpn1/Ptprf/Pygm/Rheb/Rps6/Sorbs1/Shc1/Shc3/Slc2a4/Sos1/Sos2/Srebf1/Cblb/Hkdc1/Socs2/Ppp1r3d/Akt3/Sh2b2/Ppp1r3b/Map2k1/Map2k2/Mapk1/Mapk10/Mapk3/Mapk8/Mapk9/Eif4e2/Shc4/Irs2/Ppp1r3c/Exoc7/Foxo1/Gsk3b/Mtor/Rps6kb2/Tsc1/G6pc3/Phkg2/Rptor/Pck2/Pik3cb |
| mmu04520 | Adherens junction | 1,1127E-08 | Baiap2/Actn1/Acp1/Actb/Actg1/Ctnna1/Ctnnb1/Ctnnd1/Cdc42/Cdh1/Crebbp/Csnk2a1/Csnk2a2/Csnk2b/Egfr/Erbb2/Fer/Fgfr1/Fyn/Igf1r/Insr/Lef1/Rac3/Smad3/Smad4/Met/Afdn/Nlk/Ptpn1/Ptprf/Ptprj/Ptprm/Nectin2/Rac1/Rac2/Sorbs1/Snai2/Snai1/Src/Tcf7/Tcf7l1/Tcf7l2/Ctnna3/Tgfbr1/Tgfbr2/Tjp1/Vcl/Yes1/Farp2/Wasf2/Map3k7/Mapk1/Mapk3/Iqgap1/Ep300/Lmo7/Nectin1/Nectin3/Actn4/Nectin4/Wasl/Pard3/Ssx2ip |
| mmu05226 | Gastric cancer | 1,2599E-08 | Ddb2/Braf/Raf1/Akt1/Akt2/Apc/Axin1/Axin2/Bak1/Bax/Bcl2/Ctnna1/Ctnnb1/Ccnd1/Ccne1/Ccne2/Cdh1/Cdk2/Cdkn1a/Cdkn1b/Cdkn2b/Cdx2/Gadd45a/Dvl1/Dvl2/Dvl3/E2f1/E2f3/Egf/Egfr/Erbb2/Fgf1/Fgf15/Fgf18/Fgf2/Fgf5/Fgf8/Fgf9/Fgfr2/Frat1/Fzd1/Fzd3/Fzd4/Fzd5/Fzd6/Fzd7/Fzd8/Fzd9/Gab1/Grb2/Hras/Jup/Lef1/Lrp5/Lrp6/Smad2/Smad3/Smad4/Met/Mlh1/Myc/Gadd45b/Nras/Abcb1b/Abcb1a/Pik3ca/Pik3cd/Pik3r1/Pik3r2/Pik3r3/Rb1/Rxra/Rxrg/Shc1/Shc3/Shh/Sos1/Sos2/Wnt8a/Frat2/Tcf7/Tcf7l1/Tcf7l2/Ctnna3/Wnt9a/Terc/Tert/Tgfb1/Tgfb2/Tgfb3/Tgfbr1/Tgfbr2/Rarb/Wnt1/Wnt10b/Wnt11/Wnt9b/Wnt2b/Wnt5a/Wnt5b/Wnt6/Akt3/Apc2/Gadd45g/E2f2/Map2k1/Map2k2/Mapk1/Mapk3/Polk/Shc4/Peg12/Gsk3b/Mtor/Fzd2/Rps6kb2/Pik3cb/Csnk1a1 |
| mmu05205 | Proteoglycans in cancer | 1,7441E-08 | Camk2d/Ank2/Braf/Raf1/Actb/Actg1/Akt1/Akt2/Ank1/Ank3/Casp3/Ctnnb1/Cav1/Cav2/Cbl/Ccnd1/Cd44/Cd63/Cdc42/Cdkn1a/Col1a1/Ctsl/Cttn/Ddx5/Egfr/Erbb2/Erbb3/Erbb4/Esr1/Drosha/Ptk2/Fas/Fgf2/Fgfr1/Fzd1/Fzd3/Fzd4/Fzd5/Fzd6/Fzd7/Fzd8/Fzd9/Gab1/Gpc1/Grb2/Hbegf/Hif1a/Hras/Sdc2/Hspg2/Igf1/Igf1r/Ihh/Itga2/Itga5/Itgav/Itgb1/Itgb5/Itpr1/Itpr2/Itpr3/Kdr/Arhgef1/Smad2/Mdm2/Met/Mmp9/Mras/Myc/Ppp1r12a/Nras/Pak1/Pdcd4/Pdpk1/Pik3ca/Pik3cd/Pik3r1/Pik3r2/Pik3r3/Prkaca/Prkacb/Prkcb/Prkcg/Plau/Plcg1/Ppp1cb/Ppp1cc/Mapk11/Ptch1/Ptpn11/Pxn/Rac1/Rdx/Rock1/Rock2/Rps6/Rras/Shh/Slc9a1/Sos1/Sos2/Src/Stat3/Wnt8a/Sdc1/Sdc4/Wnt9a/Tgfb1/Tgfb2/Thbs1/Tiam1/Timp3/Tlr4/Vav1/Vav2/Vegfa/Ezr/Vtn/Wnt1/Wnt10b/Wnt11/Wnt9b/Wnt2b/Wnt5a/Wnt5b/Wnt6/Ppp1r12c/Plcg2/Akt3/Tlr2/Map2k1/Map2k2/Mapk1/Mapk13/Mapk14/Mapk3/Flnb/Mapk12/Iqgap1/Smo/Frs2/Ppp1r12b/Mir21a/Mtor/Vav3/Fzd2/Rps6kb2/Rras2/Flnc/Arhgef12/Plce1/Pik3cb/Eif4b/Tfap4 |
| mmu01521 | EGFR tyrosine kinase inhibitor resistance | 4,0233E-08 | Nrg2/Braf/Raf1/Akt1/Akt2/Bad/Bax/Bcl2/Bcl2l1/Bcl2l11/Egf/Egfr/Eif4ebp1/Erbb2/Erbb3/Fgf2/Fgfr2/Fgfr3/Gab1/Gas6/Grb2/Hras/Igf1/Igf1r/Il6ra/Jak1/Jak2/Kdr/Met/Nf1/Nras/Pdgfa/Pdgfb/Pdgfra/Pik3ca/Pik3cd/Pik3r1/Pik3r2/Pik3r3/Prkcb/Prkcg/Plcg1/Pten/Rps6/Shc1/Shc3/Sos1/Sos2/Src/Stat3/Nrg1/Tgfa/Vegfa/Plcg2/Akt3/Map2k1/Map2k2/Mapk1/Mapk3/Eif4e2/Shc4/Pdgfc/Foxo3/Gsk3b/Mtor/Rps6kb2/Pdgfd/Pik3cb |
| mmu05215 | Prostate cancer | 4,7337E-08 | Etv5/Braf/Raf1/Akt1/Akt2/Atf4/Bad/Bcl2/Casp9/Ctnnb1/Ccnd1/Ccne1/Ccne2/Cdk2/Cdkn1a/Cdkn1b/Chuk/Creb1/Creb3/Crebbp/E2f1/E2f3/Egf/Egfr/Erbb2/Fgfr1/Fgfr2/Grb2/Gstp1/Hras/Hsp90ab1/Hsp90aa1/Igf1/Igf1r/Ikbkb/Lef1/Mdm2/Mmp9/Nfkb1/Nfkbia/Nras/Pdgfa/Pdgfb/Pdgfra/Pdpk1/Pik3ca/Pik3cd/Pik3r1/Pik3r2/Pik3r3/Plau/Pten/Rb1/Rela/Sos1/Sos2/Spint1/Creb3l2/Tcf7/Tcf7l1/Tcf7l2/Zeb1/Tgfa/Gstp3/Creb5/Akt3/Insrr/E2f2/Map2k1/Map2k2/Mapk1/Mapk3/Creb3l1/Ep300/Tmprss2/Pdgfc/Foxo1/Gsk3b/Mtor/Pdgfd/Pik3cb/Creb3l4 |
| mmu04071 | Sphingolipid signaling pathway | 5,782E-08 | Raf1/Adora1/Akt1/Akt2/Asah1/Bax/Bcl2/Bid/Ctsd/Degs1/S1pr1/S1pr3/Fcer1g/Fyn/Gab2/Gna12/Gnai1/Gnai2/Gnai3/Gnaq/S1pr2/Hras/Rac3/Abcc1/Nfkb1/Nras/Nsmaf/Pdpk1/Pik3ca/Pik3cd/Pik3r1/Pik3r2/Pik3r3/Prkcb/Prkcg/Prkce/Prkcz/Plcb1/Plcb2/Plcb3/Plcb4/Pld1/Pld2/Ppp2ca/Ppp2cb/Mapk11/Pten/Rac1/Rac2/Rela/Rock1/Rock2/Sgpl1/Smpd1/Smpd2/Sphk1/Sptlc2/Sgms1/Ppp2r5d/Tnfrsf1a/Traf2/Ppp2r5b/Ppp2r5a/Acer2/Ppp2r3a/Akt3/Cers6/Map2k1/Map2k2/Map3k5/Mapk1/Mapk10/Mapk13/Mapk14/Mapk3/Mapk8/Mapk9/Sptlc1/Ppp2r5c/Ppp2r5e/Mapk12/Kng2/Sgpp2/Ppp2r1a/Ppp2r2d/Asah2/Ppp2r3c/Cers4/Degs2/Tradd/Cers5/Ppp2r2a/Ppp2r2b/Ppp2r1b/Sgms2/Pik3cb/Cers2/Sgpp1/S1pr5 |
| mmu04210 | Apoptosis | 6,3069E-08 | Raf1/Actb/Actg1/Parp1/Parp2/Akt1/Akt2/Apaf1/Birc3/Birc2/Birc5/Atf4/Atm/Bad/Bak1/Bax/Bcl2/Bcl2l1/Bid/Hrk/Bcl2l11/Capn1/Capn2/Casp2/Casp3/Casp6/Casp7/Casp8/Casp9/Cflar/Chuk/Ctsb/Ctsc/Ctsd/Ctsh/Ctsl/Cycs/Daxx/Gadd45a/Dffa/Dffb/Eif2s1/Eif2ak3/Endog/Fadd/Fas/Fos/Hras/Ikbkb/Il3ra/Itpr1/Itpr2/Itpr3/Jun/Lmna/Lmnb1/Lmnb2/Bbc3/Mcl1/Gadd45b/Nfkb1/Nfkbia/Nras/Pdpk1/Pik3ca/Pik3cd/Pik3r1/Pik3r2/Pik3r3/Septin4/Ptpn13/Rela/Ripk1/Sptan1/Tnfrsf10b/Tnfrsf1a/Traf1/Traf2/Tnfsf10/Tuba1a/Tuba1b/Tuba4a/Tuba1c/Ctso/Parp3/Akt3/Tubal3/Gadd45g/Map2k1/Map2k2/Map3k5/Mapk1/Mapk10/Mapk3/Mapk8/Mapk9/Tuba8/Map3k14/Ctsf/Pidd1/Pmaip1/Ctsz/Diablo/Dab2ip/Tradd/Pik3cb/Ern1 |
| mmu04137 | Mitophagy - animal | 6,6706E-08 | Usp30/Atg5/Atf4/Bcl2l1/Bnip3/Bnip3l/Csnk2a1/Csnk2a2/Csnk2b/E2f1/Eif2ak3/Usp15/Hif1a/Hras/Jun/Mfn2/Mitf/Mras/Cited2/Nras/Sqstm1/Rab7/Rela/Rras/Sp1/Src/Atg9b/Tfeb/Rhot2/Ubb/Ulk1/Ambra1/Tbc1d17/Atg9a/Mapk10/Mapk8/Mapk9/Prkn/Tax1bp1/Becn1/Tbk1/Foxo3/Gabarap/Rhot1/Tomm7/Fis1/Tbc1d15/Rras2/Mfn1/Pink1/Optn/Pgam5/Calcoco2/Usp8/Gabarapl2/Bcl2l13 |
| mmu04935 | Growth hormone synthesis, secretion and action | 1,0306E-07 | Adcy3/Raf1/Adcy6/Adcy9/Akt1/Akt2/Atf2/Atf4/Cacna1c/Cacna1d/Socs3/Socs1/Creb1/Creb3/Crebbp/Atf6b/Bcar1/Crk/Crkl/Ptk2/Fos/Ghr/Gna11/Gnai1/Gnai2/Gnai3/Gnaq/Gnas/Grb2/Hras/Igf1/Igfals/Irs1/Irs3/Itpr1/Itpr2/Itpr3/Jak2/Junb/Nras/Pik3ca/Pik3cd/Pik3r1/Pik3r2/Pik3r3/Prkaca/Prkacb/Prkcb/Prkcg/Plcb1/Plcb2/Plcb3/Plcb4/Plcg1/Mapk11/Shc1/Shc3/Sst/Sstr1/Sstr2/Sos1/Sos2/Stat1/Stat3/Stat5a/Stat5b/Creb3l2/Adcy2/Socs2/Creb5/Plcg2/Akt3/Map2k1/Map2k2/Map2k4/Map2k6/Map3k1/Mapk1/Mapk10/Mapk13/Mapk14/Mapk3/Mapk8/Mapk9/Creb3l1/Shc4/Mapk12/Ep300/Irs2/Gsk3b/Mtor/Pik3cb/Creb3l4 |
| mmu05213 | Endometrial cancer | 1,1096E-07 | Ddb2/Braf/Raf1/Akt1/Akt2/Apc/Axin1/Axin2/Bad/Bak1/Bax/Casp9/Ctnna1/Ctnnb1/Ccnd1/Cdh1/Cdkn1a/Gadd45a/Egf/Egfr/Erbb2/Grb2/Hras/Lef1/Mlh1/Myc/Gadd45b/Nras/Pdpk1/Pik3ca/Pik3cd/Pik3r1/Pik3r2/Pik3r3/Pten/Sos1/Sos2/Tcf7/Tcf7l1/Tcf7l2/Ctnna3/Akt3/Apc2/Gadd45g/Map2k1/Map2k2/Mapk1/Mapk3/Polk/Foxo3/Gsk3b/Pik3cb |
| mmu05203 | Viral carcinogenesis | 1,477E-07 | H4c17/Scrib/Cdc20/Actn1/H2-Q6/H2-Q9/Rasa2/Atf2/Atf4/Atp6v0d1/Bad/Bak1/Bax/Casp3/Casp8/Ccna2/Ccnd1/Ccnd2/Ccnd3/Ccne1/Ccne2/Cdk1/Cdc42/Cdk2/Cdk4/Cdk6/Cdkn1a/Cdkn1b/Cdkn2a/Cdkn2b/Chek1/Creb1/Creb3/Crebbp/Atf6b/Ddb1/Dlg1/Egr2/Egr3/Kat2a/Grb2/Gtf2h1/Gtf2h4/H2-Bl/H2-D1/H2-K1/H2-M3/H2-Q1/H2-Q10/H2-Q2/H2-Q4/H2-T22/H2-T23/H2-T24/Hdac2/Hdac3/Hdac5/Hnrnpk/Hpn/Hras/Il6st/Irf9/Jak1/Jak3/Jun/Ltbr/Hdac10/Lyn/Mad1l1/Mapkapk2/Mdm2/Nfkb1/Nfkb2/Nfkbia/Nras/Kat2b/Pik3ca/Pik3cd/Pik3r1/Pik3r2/Pik3r3/Pkm/Prkaca/Prkacb/Polb/Eif2ak2/Psmc1/Pxn/Rac1/Rb1/Rbl1/Rbl2/Rbpj/Rel/Rela/Scin/Sp100/Src/Stat3/Stat5a/Stat5b/Creb3l2/Hdac4/Syk/Traf1/Traf2/Traf3/Traf5/Ube3a/Vdac3/H2-M10.5/Ywhae/Ywhag/Ywhah/Ywhaq/Ywhaz/Gsn/Gtf2b/Creb5/Hdac11/Vac14/Gtf2a2/Tbpl1/Gtf2h2/Usp7/Mapk1/Mapk3/Creb3l1/Skp2/H4c3/H4c4/H4c6/H4c9/H4c11/H4c12/H4c18/H2bc1/H2bc6/H2bc11/H2bc13/H2bc14/H2bc18/H4f16/H4c1/H4c2/Ep300/Hdac1/Irf7/Irf3/Ywhab/Hdac7/Snd1/Pmaip1/Actn4/Snw1/H2-T-ps/Gm8909/Gtf2e2/Ubr4/H2bl1/H4c8/Tradd/Pik3cb/Creb3l4/H2bu2/Gtf2a1/Dnaja3/H4c14 |
| mmu05222 | Small cell lung cancer | 1,7851E-07 | Ddb2/Akt1/Akt2/Apaf1/Birc3/Birc2/Bak1/Bax/Bcl2/Bcl2l1/Casp3/Casp9/Ccnd1/Ccne1/Ccne2/Cdk2/Cdk4/Cdk6/Cdkn1a/Cdkn1b/Cdkn2b/Chuk/Cycs/Gadd45a/E2f1/E2f3/Ptk2/Fhit/Ikbkb/Itga2/Itga2b/Itga3/Itga6/Itgav/Itgb1/Lama1/Lama3/Lama4/Lama5/Lamb1/Lamb2/Lamb3/Lamc2/Max/Myc/Gadd45b/Nfkb1/Nfkbia/Nos2/Pik3ca/Pik3cd/Pik3r1/Pik3r2/Pik3r3/Pten/Ptgs2/Rb1/Rela/Rxra/Rxrg/Rarb/Traf1/Traf2/Traf3/Traf4/Traf5/Traf6/Zbtb17/Lamc1/Akt3/Gadd45g/E2f2/Polk/Skp2/Cks2/Pik3cb |
| mmu05165 | Human papillomavirus infection | 2,0336E-07 | Tada3/Tubg1/Tubg2/Maml1/Itga9/Scrib/Ticam1/Atp6v1h/Raf1/H2-Q6/H2-Q9/Atp6v0b/Akt1/Akt2/Apc/Atm/Atp6v1a/Atp6v0d1/Atp6v1e1/Atp6v0e/Atp6v0a1/Atp6v0c/Axin1/Axin2/Bad/Bak1/Bax/Casp3/Casp8/Ctnnb1/Ccna2/Ccnd1/Ccnd2/Ccnd3/Ccne1/Ccne2/Cdc42/Cdk2/Cdk4/Cdk6/Cdkn1a/Cdkn1b/Chad/Chuk/Patj/Col1a1/Comp/Creb1/Creb3/Crebbp/Dlg1/Dvl1/Dvl2/Dvl3/E2f1/Egf/Egfr/Eif4ebp1/Fadd/Ptk2/Fas/Tlr3/Fzd1/Fzd3/Fzd4/Fzd5/Fzd6/Fzd7/Fzd8/Fzd9/Gnas/Grb2/Magi1/H2-Bl/H2-D1/H2-K1/H2-M3/H2-Q1/H2-Q10/H2-Q2/H2-Q4/H2-T22/H2-T23/H2-T24/Hdac2/Hes1/Hes2/Hes5/Hey1/Hey2/Hras/Ifnar1/Ifnar2/Ikbkb/Irf1/Irf9/Itga2/Itga2b/Itga3/Itga5/Itga6/Itga7/Itgav/Itgb1/Itgb5/Itgb6/Itgb7/Jag1/Jak1/Lama1/Lama3/Lama4/Lama5/Lamb1/Lamb2/Lamb3/Lamc2/Lfng/Llgl1/Mdm2/Mx2/Nfkb1/Notch1/Notch2/Notch3/Nras/Pik3ca/Pik3cd/Pik3r1/Pik3r2/Pik3r3/Pkm/Prkaca/Prkacb/Prkci/Prkcz/Ppp2ca/Ppp2cb/Eif2ak2/Psen1/Psmc1/Pten/Ptger4/Ptgs2/Itgb4/Pxn/Rb1/Rbl1/Rbl2/Rbpj/Rela/Rfng/Rheb/Sos1/Sos2/Spp1/Stat1/Stat2/Creb3l2/Wnt8a/Tcf7/Tcf7l1/Tcf7l2/Wnt9a/Llgl2/Tert/Ppp2r5d/Thbs1/Atp6v0a2/Tnfrsf1a/Traf3/Ube3a/Vegfa/Vtn/Wnt1/Wnt10b/Wnt11/Wnt9b/Wnt2b/Wnt5a/Wnt5b/Wnt6/H2-M10.5/Crb3/Ppp2r5b/Lamc1/Ppp2r5a/Ifna13/Oasl1/Creb5/Ppp2r3a/Tbpl1/Akt3/Apc2/Dlg2/Oasl2/Itga8/Ifna15/Atr/Map2k1/Map2k2/Mapk1/Mapk3/Creb3l1/Ppp2r5c/Ppp2r5e/Slc9a3r1/Maml2/Tcirg1/Itgb8/Ep300/Maml3/Hdac1/Ppp2r1a/Ppp2r2d/Irf3/Tyk2/Hes6/Heyl/Pals1/Foxo1/Tbk1/Ikbke/Gsk3b/Mtor/Fzd2/Pard6b/Rps6kb2/Ppp2r3c/Tsc1/Atp6v1f/Atp6v1g1/Atp6v1c1/H2-T-ps/Gm8909/Atp6v1c2/Ubr4/Tradd/Ppp2r2a/Ppp2r2b/Ppp2r1b/Nfx1/Pik3cb/Atp6v1e2/Creb3l4/Hes7/Csnk1a1/Pard6g/Pard3 |
| mmu04070 | Phosphatidylinositol signaling system | 2,0741E-07 | Inpp5f/Synj1/Dgkz/Pi4kb/Dgkg/Dgkq/Cds2/Impa2/Pip4k2c/Calm1/Calm2/Calm3/Dgka/Inpp1/Inpp5b/Inppl1/Itpr1/Itpr2/Itpr3/Mtmr4/Inpp5j/Pik3c2a/Pik3c2g/Pik3ca/Pik3cd/Pik3r1/Pik3r2/Pik3r3/Pip5k1c/Pip4k2a/Pip5k1b/Pip5k1a/Prkcb/Prkcg/Plcb1/Plcb2/Plcb3/Plcb4/Plcd1/Plcg1/Inpp5k/Pten/Synj2/Inpp5a/Dgkb/Itpk1/Mtmr6/Pi4ka/Pik3c3/Dgkd/Ppip5k2/Itpka/Itpkc/Inpp4b/Plcg2/Pik3c2b/Bpnt2/Inpp4a/Ip6k1/Dgki/Itpkb/Ppip5k1/Dgkh/Mtmr7/Impa1/Dgke/Inpp5e/Pi4k2b/Ipmk/Pip4p2/Plce1/Mtmr3/Cds1/Pik3cb/Ippk/Ip6k2/Mtmr2/Sacm1l/Pi4k2a/Mtmr14 |
| mmu04115 | p53 signaling pathway | 2,189E-07 | Ddb2/Apaf1/Atm/Bax/Bcl2/Bcl2l1/Bid/Casp3/Casp8/Casp9/Ccnb2/Ccnd1/Ccnd2/Ccnd3/Ccne1/Ccne2/Ccng1/Ccng2/Cd82/Cdk1/Cdk2/Cdk4/Cdk6/Cdkn1a/Cdkn2a/Chek1/Cycs/Gadd45a/Ei24/Sesn1/Fas/Igf1/Bbc3/Mdm2/Mdm4/Gadd45b/Pten/Rrm2/Siah1a/Thbs1/Trp73/Zmat3/Sesn2/Gadd45g/Atr/Cop1/Ccnb1/Gtse1/Siva1/Rrm2b/Ppm1d/Sfn/Pidd1/Pmaip1/Perp/Shisa5/Rprm/Steap3/Aifm2/Sesn3/Gorab |
| mmu04931 | Insulin resistance | 2,5673E-07 | Acacb/Ppp1r3e/Prkaa1/Prkaa2/Prkab2/Prkag2/Pygb/Pygl/Ptpa/Akt1/Akt2/Cd36/Socs3/Cpt1a/Creb1/Creb3/G6pc/Gfpt1/Gfpt2/Gys1/Ikbkb/Insr/Irs1/Ppargc1b/Nfkb1/Nfkbia/Pdpk1/Pik3ca/Pik3cd/Pik3r1/Pik3r2/Pik3r3/Prkcb/Prkcd/Prkce/Prkcz/Ppara/Ppargc1a/Ppp1cb/Ppp1cc/Prkab1/Prkag1/Pten/Ptpn1/Ptpn11/Ptprf/Pygm/Rela/Rps6ka1/Rps6ka2/Slc2a1/Slc2a2/Slc2a4/Srebf1/Mlxip/Stat3/Creb3l2/Tbc1d4/Mlx/Tnfrsf1a/Nr1h3/Nr1h2/Trib3/Ppp1r3d/Creb5/Akt3/Ppp1r3b/Mapk10/Mapk8/Mapk9/Creb3l1/Slc27a1/Slc27a2/Slc27a3/Slc27a4/Irs2/Ppp1r3c/Foxo1/Gsk3b/Mtor/Mlxipl/Rps6kb2/G6pc3/Crtc2/Pck2/Pik3cb/Oga/Creb3l4 |
| mmu04668 | TNF signaling pathway | 2,6124E-07 | Akt1/Akt2/Birc3/Birc2/Atf2/Atf4/Bcl3/Casp3/Casp7/Casp8/Cebpb/Cflar/Chuk/Socs3/Creb1/Creb3/Atf6b/Csf1/Edn1/Fadd/Fas/Fos/Cxcl1/Icam1/Cxcl10/Ifi47/Ikbkb/Il15/Irf1/Itch/Jag1/Jun/Junb/Lif/Mmp14/Mmp9/Nfkb1/Nfkbia/Pik3ca/Pik3cd/Pik3r1/Pik3r2/Pik3r3/Mapk11/Ptgs2/Rela/Ripk1/Ccl5/Cx3cl1/Creb3l2/Tnfaip3/Tnfrsf1a/Tnfrsf1b/Traf1/Traf2/Traf3/Traf5/Vegfc/Creb5/Akt3/Nod2/Map2k1/Map2k4/Map2k6/Map2k7/Map3k5/Map3k7/Map3k8/Mapk1/Mapk10/Mapk13/Mapk14/Mapk3/Mapk8/Mapk9/Creb3l1/Mapk12/Map3k14/Ripk3/Rps6ka4/Tab1/Tab2/Dab2ip/Tradd/Pgam5/Rps6ka5/Dnm1l/Mlkl/Pik3cb/Creb3l4 |
| mmu04012 | ErbB signaling pathway | 2,7339E-07 | Nrg2/Camk2d/Braf/Raf1/Abl1/Abl2/Akt1/Akt2/Areg/Bad/Btc/Cbl/Cdkn1a/Cdkn1b/Crk/Crkl/Egf/Egfr/Eif4ebp1/Erbb2/Erbb3/Erbb4/Ptk2/Gab1/Grb2/Hbegf/Hras/Jun/Myc/Nck1/Nck2/Nras/Pak1/Pik3ca/Pik3cd/Pik3r1/Pik3r2/Pik3r3/Prkcb/Prkcg/Plcg1/Shc1/Shc3/Sos1/Sos2/Src/Stat5a/Stat5b/Cblb/Nrg1/Pak6/Tgfa/Pak2/Plcg2/Akt3/Map2k1/Map2k2/Map2k4/Map2k7/Mapk1/Mapk10/Mapk3/Mapk8/Mapk9/Shc4/Gsk3b/Mtor/Rps6kb2/Pik3cb/Nrg4 |
| mmu04015 | Rap1 signaling pathway | 6,7614E-07 | Prkd2/Adcy3/Rapgef1/Braf/Rap1a/Raf1/Rap1gap/Actb/Actg1/Adcy6/Adcy9/Adora2a/Adora2b/Angpt1/Angpt2/Akt1/Akt2/Calm1/Calm2/Calm3/Ctnnb1/Ctnnd1/Cdc42/Cdh1/Cnr1/Bcar1/Crk/Crkl/Csf1/Efna1/Efna2/Efna3/Efna4/Efna5/Egf/Egfr/Enah/Epha2/F2r/Fgf1/Fgf15/Fgf18/Fgf2/Fgf5/Fgf8/Fgf9/Fgfr1/Fgfr2/Fgfr3/Fgfr4/Flt4/Gnai1/Gnai2/Gnai3/Gnao1/Gnaq/Gnas/Lpar1/Grin1/Magi1/Hras/Id1/Igf1/Igf1r/Insr/Itga2b/Itgam/Itgb1/Kdr/Kit/Rac3/Met/Kitl/Afdn/Mras/Nras/P2ry1/Pdgfa/Pdgfb/Pdgfra/Pfn1/Pfn2/Pgf/Pik3ca/Pik3cd/Pik3r1/Pik3r2/Pik3r3/Prkcb/Prkcg/Prkci/Prkd1/Prkcz/Plcb1/Plcb2/Plcb3/Plcb4/Plcg1/Mapk11/Rapgef6/Rac1/Rac2/Rasgrp2/Ralgds/Rras/Sipa1/Src/Adcy2/Rap1b/Tek/Sipa1l1/Rapgef5/Thbs1/Tiam1/Vasp/Vav1/Vav2/Vegfa/Vegfb/Vegfc/Rapgef3/Farp2/Akt3/Dock4/Fyb/Rasgrp3/Sipa1l2/Map2k1/Map2k2/Map2k6/Mapk1/Mapk13/Mapk14/Mapk3/Mapk12/Magi2/Lpar2/Rassf5/Pdgfc/Rala/Rapgef4/Vav3/Pard6b/Ralb/Lpar3/Tln2/Pdgfd/Plce1/Sipa1l3/Pik3cb/Prkd3/Rapgef2/Pard6g/Pard3/Magi3 |
| mmu04530 | Tight junction | 7,3214E-07 | Arhgef18/Synpo/Scrib/Prkaa1/Prkaa2/Prkab2/Prkag2/Actn1/Rap1a/Actb/Actg1/Cacna1d/Runx1/Ccnd1/Cd1d1/Cd1d2/Cdc42/Cdk4/Cftr/Patj/Cldn1/Cldn3/Cttn/Dlg1/Erbb2/Gata4/Magi1/Hspa4/Itgb1/F11r/Jun/Arhgef2/Llgl1/Rab8a/Afdn/Mpdz/Myh11/Myh9/Myl6/Nedd4/Nf2/Ocln/Pcna/Prkaca/Prkacb/Prkce/Prkci/Prkcz/Ppp2ca/Ppp2cb/Prkab1/Prkag1/Rapgef6/Rac1/Rdx/Rock1/Rock2/Src/Stk11/Myl6b/Llgl2/Tiam1/Marveld2/Tjp1/Tjp2/Tuba1a/Tuba1b/Tuba4a/Tuba1c/Vasp/Ezr/Crb3/Micall2/Rab8b/Tubal3/Dlg2/Actr3b/Map2k7/Map3k1/Map3k5/Mapk10/Mapk8/Mapk9/Slc9a3r1/Tjp3/Whamm/Ppp2r1a/Ppp2r2d/Cldn7/Tuba8/Epb41l4b/Cldn6/Cldn8/Myh13/Pals1/Amotl2/Ybx3/Cldn10/Pard6b/Actn4/Actr2/Myh7b/Jam2/Myl12b/Cgnl1/Rab13/Arhgap17/Cgn/Cldn23/Myh14/Ppp2r2a/Igsf5/Ppp2r2b/Marveld3/Ppp2r1b/Tjap1/Actr3/Amotl1/Rapgef2/Myh10/Nedd4l/Jam3/Pard6g/Pard3 |
| mmu05211 | Renal cell carcinoma | 7,3214E-07 | Rapgef1/Braf/Rap1a/Raf1/Egln1/Egln2/Egln3/Akt1/Akt2/Arnt/Bad/Cdc42/Cdkn1a/Crebbp/Crk/Crkl/Epas1/Fh1/Gab1/Grb2/Hif1a/Hras/Jun/Met/Nras/Pak1/Pdgfb/Pik3ca/Pik3cd/Pik3r1/Pik3r2/Pik3r3/Ptpn11/Rac1/Slc2a1/Sos1/Sos2/Pak6/Rap1b/Tgfa/Tgfb1/Tgfb2/Tgfb3/Vegfa/Vhl/Pak2/Akt3/Ets1/Map2k1/Map2k2/Mapk1/Mapk3/Ep300/Rbx1/Elob/Cul2/Pik3cb/Prcc |
| mmu01524 | Platinum drug resistance | 7,7551E-07 | Gstt3/Akt1/Akt2/Apaf1/Birc3/Birc2/Birc5/Atm/Atp7b/Bad/Bak1/Bax/Bcl2/Bcl2l1/Bid/Brca1/Casp3/Casp8/Casp9/Cdkn1a/Cdkn2a/Cycs/Erbb2/Ercc1/Fadd/Fas/Gsta3/Gsta4/Gstm1/Gstm2/Gstm3/Gstm4/Gstm5/Gstp1/Gstt1/Gstt2/Gsto1/Bbc3/Mdm2/Mlh1/Msh2/Msh6/Pdpk1/Pik3ca/Pik3cd/Pik3r1/Pik3r2/Pik3r3/Rev3l/Slc31a1/Mgst2/Top2a/Top2b/Gstp3/Xpa/Akt3/Map3k5/Mapk1/Mapk3/Mgst1/Pmaip1/Mgst3/Gsto2/Gstm7/Pik3cb |
| mmu04146 | Peroxisome | 9,6429E-07 | Pex12/Nudt19/Pecr/Acaa1a/Acox1/Cat/Ephx2/Acsl1/Gnpat/Hmgcl/Hsd17b4/Idh1/Phyh/Acot8/Amacr/Mpv17/Mvk/Nos2/Prdx1/Pex11a/Pex11b/Pex16/Pex7/Pipox/Pex19/Abcd3/Abcd4/Pex2/Pex5/Scp2/Slc25a17/Sod1/Hmgcll1/Paox/Acsl6/Xdh/Pex6/Agps/Mpv17l2/Acaa1b/Eci2/Decr2/Slc27a2/Abcd2/Idh2/Dhrs4/Acsl5/Ech1/Prdx5/Hao2/Pex14/Pex3/Mlycd/Pxmp4/Pex10/Far1/Nudt7/Nudt12/Pmvk/Eci3/Pex11g/Ddo/Pex13/Pex26/Crot/Ehhadh/Gstk1/Acox3/Mpv17l |
| mmu05418 | Fluid shear stress and atherosclerosis | 1,2583E-06 | Gstt3/Prkaa1/Prkaa2/Actb/Actg1/Acvr1/Acvr2a/Acvr2b/Akt1/Akt2/Ass1/Bcl2/Bmpr1a/Bmpr1b/Bmpr2/Calm1/Calm2/Calm3/Ctnnb1/Cav1/Cav2/Chuk/Ctsl/Cyba/Edn1/Ptk2/Fos/Gpc1/Gsta3/Gsta4/Gstm1/Gstm2/Gstm3/Gstm4/Gstm5/Gstp1/Gstt1/Gstt2/Gsto1/Hmox1/Hsp90ab1/Hsp90aa1/Sdc2/Icam1/Ikbkb/Il1r1/Itga2b/Itgav/Jun/Kdr/Klf2/Arhgef2/Rac3/Sumo2/Mef2a/Mef2c/Mmp9/Nfe2l2/Nfkb1/Nqo1/Nppc/Sqstm1/Pdgfa/Pdgfb/Pik3ca/Pik3cd/Pik3r1/Pik3r2/Pik3r3/Prkcz/Mapk11/Dusp1/Rac1/Rac2/Rela/Sumo3/Src/Sdc1/Sdc4/Mgst2/Thbd/Tnfrsf1a/Txn1/Sumo1/Vegfa/Gstp3/Akt3/Map2k5/Mapk7/Map2k4/Map2k6/Map2k7/Map3k5/Map3k7/Mapk10/Mapk13/Mapk14/Mapk8/Mapk9/Mapk12/Keap1/Txn2/Mgst1/Pias4/Trpv4/Mgst3/Gsto2/Gstm7/Pik3cb |
| mmu00562 | Inositol phosphate metabolism | 1,4795E-06 | Inpp5f/Fig4/Synj1/Aldh6a1/Pi4kb/Impa2/Pip4k2c/Inpp1/Inpp5b/Inppl1/Mtmr4/Inpp5j/Minpp1/Pik3c2a/Pik3c2g/Pik3ca/Pik3cd/Pip5k1c/Pip4k2a/Pip5k1b/Pip5k1a/Plcb1/Plcb2/Plcb3/Plcb4/Plcd1/Plcg1/Inpp5k/Pten/Synj2/Inpp5a/Itpk1/Mtmr6/Tpi1/Pi4ka/Pik3c3/Pip5kl1/Itpka/Itpkc/Inpp4b/Plcg2/Pik3c2b/Bpnt2/Inpp4a/Plch1/Pik3cg/Itpkb/Mtmr7/Impa1/Inpp5e/Pi4k2b/Ipmk/Isyna1/Plce1/Mtmr3/Pik3cb/Ippk/Mtmr2/Sacm1l/Pi4k2a/Mtmr14 |
| mmu04310 | Wnt signaling pathway | 2,3061E-06 | Fbxw11/Prickle1/Lgr4/Camk2d/Semp2l2b/Apc/Axin1/Axin2/Btrc/Cacybp/Ctnnb1/Ccnd1/Ccnd2/Ccnd3/Crebbp/Csnk2a1/Csnk2a2/Csnk2b/Ctbp1/Ctbp2/Dvl1/Dvl2/Dvl3/Lgr5/Fosl1/Frat1/Fzd1/Fzd3/Fzd4/Fzd5/Fzd6/Fzd7/Fzd8/Fzd9/Invs/Jun/Lef1/Lrp5/Lrp6/Rac3/Smad3/Smad4/Myc/Nfatc1/Nfatc2/Nfatc3/Nlk/Prkaca/Prkacb/Prkcb/Prkcg/Plcb1/Plcb2/Plcb3/Plcb4/Ppard/Ppp3ca/Ppp3cb/Ppp3cc/Ppp3r1/Psen1/Rac1/Rac2/Rock2/Ryk/Sfrp2/Sfrp1/Siah1a/Sox17/Rnf43/Daam1/Wnt8a/Frat2/Skp1/Tcf7/Tcf7l1/Tcf7l2/Wnt9a/Wnt1/Wnt10b/Wnt11/Wnt9b/Wnt2b/Wnt5a/Wnt5b/Wnt6/Vangl1/Apc2/Wif1/Prickle2/Map3k7/Mapk10/Mapk8/Mapk9/Ror1/Ror2/Cul1/Csnk1e/Peg12/Ep300/Znrf3/Sfrp5/Rbx1/Ruvbl1/Gsk3b/Fzd2/Ctnnbip1/Chd8/Bambi/Rspo3/Nfatc4/Cby1/Sost/Senp2/Daam2/Notum/Tbl1xr1/Csnk1a1/Vangl2/Nkd1 |
| mmu04710 | Circadian rhythm | 2,5985E-06 | Fbxw11/Csnk1d/Prkaa1/Prkaa2/Prkab2/Prkag2/Arntl/Btrc/Clock/Creb1/Cry1/Cry2/Npas2/Per1/Per2/Per3/Prkab1/Prkag1/Rora/Rorc/Bhlhe40/Skp1/Nr1d1/Rorb/Cul1/Csnk1e/Fbxl3/Rbx1/Bhlhe41 |
| mmu05135 | Yersinia infection | 2,7315E-06 | Ticam1/Baiap2/Pkn2/Arhgef28/Actb/Actg1/Akt1/Akt2/Arf6/Cdc42/Chuk/Bcar1/Crk/Crkl/Ptk2/Fos/Ikbkb/Il18/Itga5/Itgb1/Jun/Arhgef1/Limk1/Rac3/Nfatc1/Nfatc2/Nfatc3/Nfkb1/Nfkbia/Pik3ca/Pik3cd/Pik3r1/Pik3r2/Pik3r3/Pip5k1c/Pip5k1b/Pip5k1a/Plcg1/Mapk11/Ptk2b/Pxn/Rac1/Rac2/Rela/Rock1/Rock2/Rps6ka1/Rps6ka2/Src/Wipf1/Tlr4/Traf2/Traf6/Vav1/Vav2/Zap70/Akt3/Fyb/Wasf2/Actr3b/Map2k1/Map2k2/Map2k4/Map2k6/Map2k7/Map3k7/Mapk1/Mapk10/Mapk13/Mapk14/Mapk3/Mapk8/Mapk9/Git2/Irak4/Mapk12/Pkn1/Dock1/Arhgef7/Irf3/Skap2/Tbk1/Gsk3b/Vav3/Tab1/Actr2/Pycard/Wipf2/Tab2/Arhgef12/Wasl/Actr3/Pik3cb |
| mmu05166 | Human T-cell leukemia virus 1 infection | 2,7866E-06 | Trrap/Xpo1/Adcy3/Espl1/Cdc20/Fdps/H2-Q6/H2-Q9/Adcy6/Adcy9/Akt1/Akt2/Slc25a4/Atf2/Atf4/Atm/B2m/Bax/Bcl2l1/Bub1b/Bub3/Tspo/Calr/Canx/Ccna2/Ccnb2/Ccnd1/Ccnd2/Ccnd3/Ccne1/Ccne2/Cdk2/Cdk4/Cdkn1a/Cdkn2a/Cdkn2b/Cdkn2c/Chek1/Chuk/Creb1/Creb3/Crebbp/Atf6b/Dlg1/E2f1/E2f3/Egr1/Egr2/Elk4/Fos/Fosl1/Kat2a/H2-Ab1/H2-Bl/H2-D1/H2-Eb1/H2-K1/H2-M3/H2-DMa/H2-Q1/H2-Q10/H2-Q2/H2-Q4/H2-T22/H2-T23/H2-T24/Hras/Icam1/Ikbkb/Il15/Il15ra/Il1r1/Jak1/Jak3/Jun/Ltbr/Mad1l1/Smad2/Smad3/Smad4/Anapc1/Msx1/Msx3/Myc/Nfatc1/Nfatc2/Nfatc3/Nfkb1/Nfkb2/Nfkbia/Nfyb/Nras/Nrp1/Kat2b/Pik3ca/Pik3cd/Pik3r1/Pik3r2/Pik3r3/Prkaca/Prkacb/Polb/Ppp3ca/Ppp3cb/Ppp3cc/Ppp3r1/Pten/Ran/Rasl2-9/Rb1/Rela/Relb/Spi1/Slc2a1/Stat5a/Stat5b/Creb3l2/Adcy2/Tcf3/Tert/Tgfb1/Tgfb2/Tgfb3/Tgfbr1/Tgfbr2/Tnfrsf1a/Cd40/Vdac1/Vdac2/Vdac3/H2-M10.5/Zfp36/Creb5/Vac14/Tbpl1/Akt3/Ets1/Ets2/E2f2/Atr/Map2k1/Map2k2/Map2k4/Map3k1/Map3k3/Mapk1/Mapk10/Mapk3/Mapk8/Mapk9/Creb3l1/Pttg1/Ep300/Crtc1/Anapc4/Cdc23/Map3k14/Mad2l1/Gps2/Anapc7/Anapc5/Cdc26/H2-T-ps/Gm8909/Cdc16/Crtc3/Tln2/Ranbp3/Nfatc4/Crtc2/Pik3cb/Creb3l4/Kat5 |
| mmu05231 | Choline metabolism in cancer | 5,454E-06 | Slc44a1/Dgkz/Raf1/Dgkg/Dgkq/Akt1/Akt2/Chka/Pcyt1a/Dgka/Egf/Egfr/Eif4ebp1/Fos/Grb2/Hif1a/Hras/Jun/Rac3/Nras/Pdgfa/Pdgfb/Pdgfra/Pdpk1/Pik3ca/Pik3cd/Pik3r1/Pik3r2/Pik3r3/Pip5k1c/Pip5k1b/Pip5k1a/Prkcb/Prkcg/Lypla1/Pla2g4a/Plcg1/Pld1/Pld2/Plpp1/Rac1/Rac2/Ralgds/Rheb/Slc22a5/Sos1/Sos2/Sp1/Chpt1/Slc44a3/Dgkb/Dgkd/Akt3/Wasf2/Map2k1/Map2k2/Mapk1/Mapk10/Mapk3/Mapk8/Mapk9/Slc22a4/Dgki/Dgkh/Plpp2/Pdgfc/Dgke/Slc22a21/Mtor/Rps6kb2/Tsc1/Plpp3/Slc44a2/Slc44a4/Pdgfd/Gpcpd1/Pik3cb |
| mmu03018 | RNA degradation | 6,6168E-06 | Pan2/Cnot6/Skiv2l/Exosc4/Btg1/Btg2/Btg3/Ddx6/Eno1/Eno2/Hspd1/Hspa9/Pabpc1/Pfkl/Pfkm/Cnot7/Tent4a/Dis3l/Tent4b/Ttc37/Tob1/Patl1/Eno4/Exosc2/Pabpc4/Cnot6l/Cnot3/Cnot1/Edc4/Xrn1/Xrn2/Pabpc4l/Lsm2/Dcp1b/Zcchc7/Edc3/Pabpc1l/Lsm4/Exosc9/Exosc10/Cnot4/Pfkp/Tob2/C1d/Lsm7/Wdr61/Exosc3/Lsm5/Exosc7/Exosc1/Lsm1/Mphosph6/Cnot8/Dcps/Dcp2/Pnpt1/Cnot2/Dhx36/Exosc6/Pan3/Parn/Nudt16/Dcp1a/Lsm8/Lsm6/Cnot10 |
| mmu04142 | Lysosome | 6,6479E-06 | Gga1/Ap4e1/Atp6v1h/Gusb/Manba/Abca2/Laptm4b/Atp6v0b/Acp2/Acp5/Aga/Ap1b1/Ap1g1/Ap1g2/Ap1m1/Ap1m2/Ap1s1/Ap3b1/Ap3b2/Ap3d1/Ap3s1/Ap3s2/Ap4m1/Ap4s1/Arsb/Asah1/Atp6v0d1/Atp6v0a1/Atp6v0c/Scarb2/Cd63/Tpp1/Cln3/Clta/Ctsb/Ctsc/Ctsd/Ctse/Ctsh/Ctsl/Dnase2a/Gaa/Galc/Gba/Gm2a/Hexa/Hexb/Hyal1/Igf2r/Lamp1/Lipa/M6pr/Man2b1/Laptm4a/Naga/Neu1/Npc1/Slc11a1/Slc11a2/Ctsa/Ppt1/Lgmn/Psap/Pla2g15/Smpd1/Sort1/Cln5/Gnptg/Atp6v0a2/Ctso/Slc17a5/Ap1s3/Gga3/Tcirg1/Naglu/Nagpa/Gnptab/Galns/Hgsnat/Cd164/Ppt2/Abcb9/Ctsf/Litaf/Sumf1/Ctsz/Ap3m2/Cltc/Npc2/Fuca1/Gga2/Cltb/Gns/Mcoln1 |
| mmu05169 | Epstein-Barr virus infection | 6,6479E-06 | Ddb2/H2-Q6/H2-Q9/Akt1/Akt2/Apaf1/B2m/Bak1/Bax/Bcl2/Bid/Bcl2l11/Calr/Casp3/Casp8/Casp9/Ccna2/Ccnd1/Ccnd2/Ccnd3/Ccne1/Ccne2/Entpd1/Cd247/Cd44/Cdk2/Cdk4/Cdk6/Cdkn1a/Cdkn1b/Chuk/Cr2/Cycs/Gadd45a/E2f1/E2f3/Fadd/Fas/Pdia3/H2-Ab1/H2-Bl/H2-D1/H2-Eb1/H2-K1/H2-M3/H2-DMa/H2-Q1/H2-Q10/H2-Q2/H2-Q4/H2-T22/H2-T23/H2-T24/Hdac2/Hes1/Icam1/Cxcl10/Ifnar1/Ifnar2/Ikbkb/Irf9/Jak1/Jak3/Jun/Lyn/Mdm2/Psmd7/Myc/Gadd45b/Nedd4/Nfkb1/Nfkb2/Nfkbia/Nfkbib/Nfkbie/Pik3ca/Pik3cd/Pik3r1/Pik3r2/Pik3r3/Mapk11/Eif2ak2/Psmc1/Psmc2/Psmc3/Psmc5/Psmd4/Rac1/Rb1/Rbpj/Rela/Relb/Ripk1/Sem1/Ncor2/Stat1/Stat2/Stat3/Syk/Tap2/Tapbp/Entpd3/Psmd2/Tnfaip3/Cd40/Traf2/Traf3/Traf5/Traf6/Psmd3/Vim/H2-M10.5/Mavs/Ddx58/Ifna13/Plcg2/Akt3/Gadd45g/Psmc4/Psmd13/Tlr2/Ifna15/E2f2/Usp7/Map2k4/Map2k6/Map2k7/Map3k7/Mapk10/Mapk13/Mapk14/Mapk8/Mapk9/Irak4/Polk/Skp2/Mapk12/Hdac1/Map3k14/Irf7/Irf3/Tyk2/Adrm1/Tbk1/Ikbke/Psmd8/Psmd14/Sap30/Snw1/Psmd6/Tab1/H2-T-ps/Gm8909/Cir1/Psmd12/Psmc6/Tab2/Psmd11/Psmd1/Tradd/Pik3cb |
| mmu04722 | Neurotrophin signaling pathway | 6,6479E-06 | Rapgef1/Camk2d/Irak2/Braf/Rap1a/Raf1/Abl1/Akt1/Akt2/Atf4/Bad/Bax/Bcl2/Calm1/Calm2/Calm3/Camk4/Cdc42/Crk/Crkl/Gab1/Arhgdig/Grb2/Hras/Ikbkb/Irs1/Jun/Sh2b3/Zfp369/Mapkapk2/Nfkb1/Nfkbia/Nfkbib/Nfkbie/Nras/Pdpk1/Pik3ca/Pik3cd/Pik3r1/Pik3r2/Pik3r3/Prkcd/Plcg1/Mapk11/Psen1/Psen2/Ptpn11/Arhgdia/Rac1/Rela/Rps6ka1/Rps6ka2/Sh2b1/Shc1/Shc3/Sort1/Sos1/Sos2/Rap1b/Traf6/Trp73/Ywhae/Plcg2/Akt3/Sh2b2/Map2k5/Mapk7/Map2k1/Map2k2/Map2k7/Map3k1/Map3k3/Map3k5/Mapk1/Mapk10/Mapk13/Mapk14/Mapk3/Mapk8/Mapk9/Irak4/Shc4/Mapk12/Frs2/Foxo3/Gsk3b/Zfp110/Prdm4/Rps6ka5/Pik3cb/Kidins220/Ntf5 |
| mmu04919 | Thyroid hormone signaling pathway | 6,7153E-06 | Raf1/Actb/Actg1/Akt1/Akt2/Atp1a1/Atp1b1/Atp1b2/Atp1b3/Atp2a2/Bad/Casp9/Ctnnb1/Ccnd1/Crebbp/Dio2/Esr1/Myh7/Gata4/Kat2a/Hdac2/Hdac3/Hif1a/Hras/Itgav/Mdm2/Myc/Ncoa1/Ncoa2/Ncoa3/Notch1/Notch2/Notch3/Nras/Kat2b/Pdpk1/Pfkfb2/Pik3ca/Pik3cd/Pik3r1/Pik3r2/Pik3r3/Prkaca/Prkacb/Prkcb/Prkcg/Plcb1/Plcb2/Plcb3/Plcb4/Plcd1/Plcg1/Med1/Rheb/Rxra/Rxrg/Ncor1/Slc2a1/Slc9a1/Src/Stat1/Tbc1d4/Med16/Thra/Thrb/Plcg2/Med17/Akt3/Med24/Map2k1/Map2k2/Mapk1/Mapk3/Med13/Ep300/Med12l/Hdac1/Rcan2/Rcan1/Pfkp/Foxo1/Gsk3b/Mtor/Med4/Med27/Med30/Slc16a10/Plce1/Pik3cb/Med13l |
| mmu01200 | Carbon metabolism | 8,975E-06 | H6pd/Psph/Gck/Gldc/Aldh6a1/Me2/Psat1/Shmt2/Gpt2/Me3/Pgd/Acat1/Acat2/Pcca/Acads/Aco1/Aco2/Acox1/Adh5/Aldoa/Aldoc/Cat/Cs/Dld/Eno1/Eno2/Esd/Fbp2/Fbp1/Fh1/Gapdh/Glud1/Got1/Got2/Gpi1/Hk1/Hk2/Idh1/Idh3b/Me1/Mdh2/Mdh1/Mthfr/Mmut/Ogdh/Pcx/Pfkl/Pfkm/Pgam1/Pkm/Pklr/Rpia/Shmt1/Sucla2/Suclg2/Taldo1/Hkdc1/Tkt/Tpi1/Tkfc/Eno4/Hibch/Aldob/Dlat/Glyctk/Phgdh/Ogdhl/Sdsl/Idh2/Prps1l3/Amt/Hao2/Pfkp/Suclg1/Pgls/Rpe/Pccb/Sdhd/Sdha/Sdhb/Idh3a/Gcsh/Pdhb/Acss1/Adpgk/Ehhadh/Idnk/Gpt/Dlst/Acox3/Hadha |
| mmu05214 | Glioma | 1,1568E-05 | Ddb2/Camk2d/Braf/Raf1/Akt1/Akt2/Bak1/Bax/Calm1/Calm2/Calm3/Camk4/Ccnd1/Cdk4/Cdk6/Cdkn1a/Cdkn2a/Gadd45a/E2f1/E2f3/Egf/Egfr/Grb2/Hras/Igf1/Igf1r/Mdm2/Gadd45b/Nras/Pdgfa/Pdgfb/Pdgfra/Pik3ca/Pik3cd/Pik3r1/Pik3r2/Pik3r3/Prkcb/Prkcg/Plcg1/Pten/Rb1/Shc1/Shc3/Sos1/Sos2/Tgfa/Plcg2/Akt3/Gadd45g/E2f2/Map2k1/Map2k2/Mapk1/Mapk3/Polk/Shc4/Camk1/Mtor/Pik3cb |
| mmu04330 | Notch signaling pathway | 1,2548E-05 | Maml1/Adam17/Crebbp/Ctbp1/Ctbp2/Dll1/Dll3/Dvl1/Dvl2/Dvl3/Dtx1/Kat2a/Hdac2/Hes1/Hes5/Hey1/Hey2/Jag1/Lfng/Notch1/Notch2/Notch3/Numb/Numbl/Kat2b/Psen1/Psen2/Rbpj/Rfng/Atxn1/Ncor2/Dtx4/Aph1b/Dtx3l/Aph1a/Maml2/Ep300/Maml3/Hdac1/Atxn1l/Dll4/Heyl/Snw1/Cir1/Aph1c/Dtx2 |
| mmu04350 | TGF-beta signaling pathway | 1,3891E-05 | E2f4/Amhr2/Acvr1/Acvr1b/Acvr2a/Acvr2b/Amh/Bmp2/Bmp7/Bmp8a/Bmpr1a/Bmpr1b/Bmpr2/Cdkn2b/Chrd/Crebbp/E2f5/Lefty1/Fst/Id1/Id2/Id3/Id4/Inhba/Inhbb/Inhbe/Smad1/Smad2/Smad3/Smad4/Smad5/Smad6/Smad7/Myc/Nbl1/Neo1/Pitx2/Ppp2ca/Ppp2cb/Rbl1/Rock1/Sp1/Thsd4/Skp1/Tfdp1/Tgfb1/Tgfb2/Tgfb3/Tgfbr1/Tgfbr2/Tgif1/Thbs1/Zfyve16/Tgif2/Zfyve9/Gdf6/Rgma/Mapk1/Mapk3/Ltbp1/Acvr1c/Cul1/Lefty2/Ep300/Ppp2r1a/Smad9/Rbx1/Rps6kb2/Smurf2/Hamp2/Bambi/Rgmb/Ppp2r1b/Smurf1 |
| mmu03050 | Proteasome | 1,9011E-05 | Psme4/Psmb9/Psmb8/Psmd7/Psma2/Psma3/Psmb1/Psmb4/Psmb5/Psmb6/Psmb7/Psmc1/Psmc2/Psmc3/Psmc5/Psmd4/Psme1/Psme2/Psme3/Sem1/Psmd2/Psmd3/Psmf1/Psmc4/Psmd13/Psma1/Psma5/Psma6/Psma7/Psmb2/Psmb3/Adrm1/Psmd8/Psmd14/Psmd6/Pomp/Psmd12/Psmc6/Psmd11/Psmd1 |
| mmu04211 | Longevity regulating pathway | 2,7607E-05 | Adcy3/Prkaa1/Prkaa2/Prkab2/Prkag2/Ehmt2/Adcy6/Adcy9/Akt1/Akt2/Atg5/Atf2/Atf4/Bax/Camk4/Cat/Rb1cc1/Creb1/Creb3/Atf6b/Eif4ebp1/Sesn1/Hras/Igf1/Igf1r/Insr/Irs1/Irs3/Nfkb1/Nras/Pik3ca/Pik3cd/Pik3r1/Pik3r2/Pik3r3/Prkaca/Prkacb/Pparg/Ppargc1a/Prkab1/Prkag1/Rela/Rheb/Camkk2/Creb3l2/Stk11/Adcy2/Ulk1/Sesn2/Creb5/Akt3/Creb3l1/Eif4e2/Irs2/Foxo1/Foxo3/Mtor/Rps6kb2/Tsc1/Akt1s1/Atg101/Adipor2/Adipor1/Appl1/Rptor/Pik3cb/Sesn3/Ehmt1/Creb3l4/Sirt1 |
| mmu05170 | Human immunodeficiency virus 1 infection | 3,1814E-05 | Fbxw11/Raf1/H2-Q6/H2-Q9/Akt1/Akt2/Ap1b1/Ap1g1/Ap1g2/Ap1m1/Ap1m2/Ap1s1/Atm/B2m/Bad/Bak1/Bax/Bcl2/Bcl2l1/Bid/Btrc/Calm1/Calm2/Calm3/Calr/Casp3/Casp8/Casp9/Ccnb2/Cd247/Cdc25c/Cdk1/Cfl1/Cfl2/Chek1/Chuk/Cxcr4/Crk/Crkl/Cycs/Ddb1/Fadd/Ptk2/Fas/Fos/Gna11/Gnai1/Gnai2/Gnai3/Gnao1/Gnaq/Gnb1/Gng10/Gng12/Gng2/Gng5/Gng7/Gng8/Pdia3/H2-Bl/H2-D1/H2-K1/H2-M3/H2-Q1/H2-Q10/H2-Q2/H2-Q4/H2-T22/H2-T23/H2-T24/Hras/Ikbkb/Itpr1/Itpr2/Itpr3/Jun/Limk1/Limk2/Rac3/Nfatc1/Nfatc2/Nfatc3/Nfkb1/Nfkbia/Nras/Pak1/Pik3ca/Pik3cd/Pik3r1/Pik3r2/Pik3r3/Prkcb/Prkcg/Plcg1/Ppp3ca/Ppp3cb/Ppp3cc/Ppp3r1/Mapk11/Ptk2b/Pxn/Rac1/Rac2/Rela/Ripk1/Rnf7/Tap2/Tapbp/Skp1/Pak6/Cgas/Tlr4/Tnfrsf1a/Tnfrsf1b/Traf2/Traf5/Traf6/Wee1/Pak2/H2-M10.5/Ifna13/Plcg2/Akt3/Tlr2/Ifna15/Atr/Ap1s3/Map2k1/Map2k2/Map2k6/Map2k7/Map3k7/Mapk1/Mapk10/Mapk13/Mapk14/Mapk3/Mapk8/Mapk9/Irak4/Ccnb1/Cul1/Mapk12/Trim12c/Dcaf1/Irf3/Samhd1/Rbx1/Tbk1/Mtor/Rps6kb2/Gng11/Tab1/H2-T-ps/Gm8909/Elob/Tab2/Bst2/Tradd/Sting1/Nfatc4/Pik3cb/Cul5/Apobec3 |
| mmu04928 | Parathyroid hormone synthesis, secretion and action | 3,3551E-05 | Adcy3/Arrb1/Braf/Raf1/Pde4c/Adcy6/Adcy9/Atf2/Atf4/Bcl2/Bglap2/Runx2/Cdkn1a/Creb1/Creb3/Atf6b/Cyp27b1/Egfr/Egr1/Fgfr1/Slc34a3/Fos/Gata3/Gcm1/Gna11/Gna12/Gnai1/Gnai2/Gnai3/Gnaq/Gnas/Hbegf/Itpr1/Itpr2/Itpr3/Jund/Mafb/Arhgef1/Lrp5/Lrp6/Mef2a/Mef2c/Mef2d/Mmp14/Mmp15/Mmp16/Naca/Nr4a2/Pde4a/Pde4b/Prkaca/Prkacb/Prkcb/Prkcg/Plcb1/Plcb2/Plcb3/Plcb4/Pld1/Pld2/Pthlh/Rxra/Rxrg/Sp1/Creb3l2/Adcy2/Arhgef11/Arrb2/Vdr/Creb5/Pde4d/Mmp17/Mmp25/Map2k1/Mapk1/Mapk3/Creb3l1/Slc9a3r1/Sost/Akap13/Creb3l4 |
| mmu05160 | Hepatitis C | 3,5857E-05 | Ticam1/Braf/Raf1/Akt1/Akt2/Apaf1/Bad/Bak1/Bax/Bid/Casp3/Casp8/Casp9/Ctnnb1/Ccnd1/Cd81/Cdk2/Cdk4/Cdk6/Cdkn1a/Cflar/Chuk/Socs3/Cldn1/Cldn3/Cycs/E2f1/E2f3/Egf/Egfr/Eif2s1/Eif2ak3/Fadd/Fas/Tlr3/Grb2/Hras/Eif2ak1/Cxcl10/Ifnar1/Ifnar2/Ikbkb/Eif3e/Irf9/Jak1/Ldlr/Mx2/Myc/Nfkb1/Nfkbia/Nras/Ocln/Pik3ca/Pik3cd/Pik3r1/Pik3r2/Pik3r3/Ppara/Ppp2ca/Ppp2cb/Eif2ak2/Psme3/Rb1/Rela/Ripk1/Rxra/Sos1/Sos2/Scarb1/Stat1/Stat2/Stat3/Tnfrsf1a/Traf2/Traf3/Traf6/Nr1h3/Ywhae/Ywhag/Ywhah/Ywhaq/Ywhaz/Mavs/Ddx58/Ifna13/Akt3/Rnasel/Ifna15/E2f2/Map2k1/Map2k2/Mapk1/Mapk3/Eif2ak4/Ppp2r1a/Ppp2r2d/Cldn7/Irf7/Irf3/Ywhab/Cldn6/Cldn8/Tyk2/Pias1/Tbk1/Ikbke/Gsk3b/Rsad2/Cldn10/Tradd/Cldn23/Ppp2r2a/Ppp2r2b/Ppp2r1b/Pik3cb |
| mmu00310 | Lysine degradation | 4,0376E-05 | Nsd2/Ehmt2/Acat1/Acat2/Prdm2/Aldh7a1/Aldh2/Aldh3a2/Dld/Ezh1/Ezh2/Hadh/Nsd1/Plod1/Pipox/Ash1l/Setd1b/Dot1l/Dhtkd1/Prdm9/Kmt2a/Kmt5b/Smyd2/Kmt2c/Kmt5c/Setd1a/Nsd3/Colgalt1/Hykk/Setd2/Setdb2/Plod2/Plod3/Colgalt2/Gcdh/Aass/Kmt2d/Aldh9a1/Suv39h2/Kmt5a/Smyd3/Aldh1b1/Phykpl/Setd7/Ehhadh/Setmar/Ehmt1/Dlst/Setdb1/Hadha |
| mmu04934 | Cushing syndrome | 4,6056E-05 | Adcy3/Camk2d/Orai1/Braf/Rap1a/Adcy6/Adcy9/Ahr/Aip/Apc/Arnt/Atf2/Atf4/Axin1/Axin2/Cacna1c/Cacna1d/Cacna1g/Ctnnb1/Ccnd1/Ccne1/Ccne2/Cdk2/Cdk4/Cdk6/Cdkn1a/Cdkn1b/Cdkn2a/Cdkn2b/Cdkn2c/Creb1/Creb3/Atf6b/Crhr2/Cyp11a1/Dvl1/Dvl2/Dvl3/E2f1/E2f3/Egfr/Wdr5/Fh1/Fzd1/Fzd3/Fzd4/Fzd5/Fzd6/Fzd7/Fzd8/Fzd9/Gna11/Gnai1/Gnai2/Gnai3/Gnaq/Gnas/Nr4a1/Itpr1/Itpr2/Itpr3/Ldlr/Lef1/Men1/Pbx1/Pde8a/Prkaca/Prkacb/Plcb1/Plcb2/Plcb3/Plcb4/Pomc/Rasd1/Rb1/Sp1/Scarb1/Creb3l2/Wnt8a/Adcy2/Rbbp5/Tcf7/Tcf7l1/Tcf7l2/Kmt2a/Rap1b/Wnt9a/Pde8b/Wnt1/Wnt10b/Wnt11/Wnt9b/Wnt2b/Wnt5a/Wnt5b/Wnt6/Creb5/Apc2/Ash2l/E2f2/Map2k1/Map2k2/Mapk1/Mapk3/Nr5a1/Creb3l1/Nceh1/Kmt2d/Gsk3b/Fzd2/Cacna1h/Wdr5b/Creb3l4/Usp8 |
| mmu04340 | Hedgehog signaling pathway | 5,3675E-05 | Csnk1g2/Fbxw11/Csnk1d/Arrb1/Grk2/Bcl2/Btrc/Ccnd1/Ccnd2/Gas1/Gli1/Gli2/Lrp2/Hhip/Ihh/Kif3a/Kif7/Mgrn1/Prkaca/Prkacb/Ptch1/Shh/Spop/Csnk1g1/Arrb2/Mosmo/Sufu/Cul3/Cul1/Megf8/Csnk1e/Smo/Grk3/Gsk3b/Cdon/Smurf2/Csnk1g3/Smurf1/Spopl/Csnk1a1 |
| mmu04932 | Non-alcoholic fatty liver disease (NAFLD) | 5,6113E-05 | Prkaa1/Prkaa2/Prkab2/Prkag2/Cox6b1/Akt1/Akt2/Atf4/Bax/Bid/Bcl2l11/Casp3/Casp7/Casp8/Cdc42/Cebpa/Socs3/Cox5a/Cox5b/Cox6a1/Cox6c/Cox7a1/Cox7a2/Cox7c/Cox8a/Cycs/Eif2s1/Eif2ak3/Fas/Ikbkb/Il6ra/Insr/Irs1/Itch/Jun/Ndufa2/Ndufa4/Ndufs4/Ndufv1/Nfkb1/Pik3ca/Pik3cd/Pik3r1/Pik3r2/Pik3r3/Pklr/Ppara/Prkab1/Prkag1/Rac1/Rela/Rxra/Cox7a2l/Srebf1/Mlxip/Mlx/Tgfb1/Tnfrsf1a/Traf2/Nr1h3/Uqcrc1/Xbp1/Ndufs8/Ndufs2/Ndufs1/Ndufb6/Akt3/Map3k5/Mapk10/Mapk8/Mapk9/Irs2/Ndufs6/Ndufa4l2/Gsk3b/Mlxipl/Ndufs5/Gsk3a/Ndufb5/Ndufa3/Ndufa9/Uqcr10/Ndufb9/Ndufc1/Ndufa12/Cyc1/Uqcr11/Uqcrfs1/Ndufb7/Sdhd/Sdha/Uqcrc2/Ndufa6/Ndufb8/Ndufa10/Sdhb/Ndufb4/Ndufc2/Ndufb2/Ndufa5/Ndufa8/Adipor2/Ndufab1/Adipor1/Ndufv2/Pik3cb/Ndufs7/Ern1 |
| mmu04114 | Oocyte meiosis | 6,3742E-05 | Fbxw11/Adcy3/Espl1/Cdc20/Camk2d/Adcy6/Adcy9/Btrc/Bub1/Calm1/Calm2/Calm3/Ccnb2/Ccne1/Ccne2/Cdc25c/Cdk1/Cdk2/Cpeb1/Smc3/Igf1/Igf1r/Itpr1/Itpr2/Itpr3/Mad1l1/Anapc1/Mos/Prkaca/Prkacb/Plk1/Ppp1cb/Ppp1cc/Ppp2ca/Ppp2cb/Ppp3ca/Ppp3cb/Ppp3cc/Ppp3r1/Mapk11/Rps6ka1/Rps6ka2/Slk/Aurka/Cpeb3/Adcy2/Skp1/Ppp2r5d/Ppp2r5b/Ywhae/Ywhag/Ywhah/Ywhaq/Ywhaz/Ppp2r5a/Cpeb2/Map2k1/Mapk1/Mapk13/Mapk14/Mapk3/Ccnb1/Pkmyt1/Ppp2r5c/Ppp2r5e/Cul1/Mapk12/Pttg1/Spdye4b/Stag3/Ppp2r1a/Anapc4/Cdc23/Ywhab/Mad2l1/Anapc7/Rbx1/Rec8/Anapc5/Cdc26/Fbxo5/Cpeb4/Cdc16/Spdya/Mad2l2/Sgo1/Ppp2r1b/Fbxo43 |
| mmu04810 | Regulation of actin cytoskeleton | 6,6474E-05 | Brk1/Itga9/Mylk/Baiap2/Actn1/Braf/Raf1/Actb/Actg1/Pip4k2c/Apc/Arpc1b/Cdc42/Cfl1/Cfl2/Chrm1/Chrm3/Cxcr4/Bcar1/Crk/Crkl/Diaph1/Egf/Egfr/Enah/F2/F2r/Ptk2/Fgf1/Fgf15/Fgf18/Fgf2/Fgf5/Fgf8/Fgf9/Fgfr1/Fgfr2/Fgfr3/Fgfr4/Gna12/Gng12/Lpar1/Hras/Itga2/Itga2b/Itga3/Itga5/Itga6/Itga7/Itgam/Itgav/Itgb1/Itgb5/Itgb6/Itgb7/Arhgef1/Limk1/Limk2/Rac3/Mos/Mras/Myh9/Mylpf/Ppp1r12a/Nras/Pak1/Pdgfa/Pdgfb/Pdgfra/Pfn1/Pfn2/Pik3ca/Pik3cd/Pik3r1/Pik3r2/Pik3r3/Pip5k1c/Pip4k2a/Pip5k1b/Pip5k1a/Ppp1cb/Ppp1cc/Itgb4/Pxn/Rac1/Rac2/Rdx/Rock1/Rock2/Rras/Scin/Cxcl12/Cyfip1/Slc9a1/Sos1/Sos2/Src/Pak6/Git1/Tiam1/Spata13/Vav1/Vav2/Vcl/Ezr/Pak2/Arhgef4/Gsn/Ssh1/Ppp1r12c/Ssh2/Apc2/Insrr/Itga8/Wasf2/Chrm2/Ssh3/Map2k1/Map2k2/Mapk1/Mapk3/Iqgap1/Fgd3/Itgb8/Abi2/Ppp1r12b/Dock1/Itgad/Kng2/Iqgap3/Nckap1/Lpar2/Arhgef7/Iqgap2/Pdgfc/Arpc3/Diaph3/Vav3/Actn4/Rras2/Myl12b/Arhgef12/Pdgfd/Myh14/Wasl/Pik3cb/Arpc2/Cyfip2/Myh10 |
| mmu05163 | Human cytomegalovirus infection | 6,6474E-05 | Adcy3/Raf1/H2-Q6/H2-Q9/Adcy6/Adcy9/Akt1/Akt2/Atf2/Atf4/B2m/Bak1/Bax/Bid/Calm1/Calm2/Calm3/Calr/Casp3/Casp8/Casp9/Ctnnb1/Ccnd1/Cdk4/Cdk6/Cdkn1a/Cdkn2a/Chuk/Cxcr4/Creb1/Creb3/Atf6b/Bcar1/Crk/Crkl/Cycs/E2f1/E2f3/Egfr/Eif4ebp1/Fadd/Ptk2/Fas/Gna11/Gna12/Gnai1/Gnai2/Gnai3/Gnao1/Gnaq/Gnas/Gnb1/Gng10/Gng12/Gng2/Gng5/Gng7/Gng8/Grb2/Pdia3/H2-Bl/H2-D1/H2-K1/H2-M3/H2-Q1/H2-Q10/H2-Q2/H2-Q4/H2-T22/H2-T23/H2-T24/Hras/Ikbkb/Il10rb/Il1r1/Il6ra/Itgav/Itpr1/Itpr2/Itpr3/Jak1/Arhgef1/Rac3/Mdm2/Myc/Nfatc1/Nfatc2/Nfatc3/Nfkb1/Nfkbia/Nras/Pdgfra/Pik3ca/Pik3cd/Pik3r1/Pik3r2/Pik3r3/Prkaca/Prkacb/Prkcb/Prkcg/Plcb1/Plcb2/Plcb3/Plcb4/Ppp3ca/Ppp3cb/Ppp3cc/Ppp3r1/Mapk11/Ptger2/Ptger3/Ptger4/Ptgs2/Ptk2b/Pxn/Rac1/Rac2/Rb1/Rela/Rheb/Ripk1/Rock1/Rock2/Ccl5/Cx3cl1/Cxcl12/Sos1/Sos2/Sp1/Src/Stat3/Creb3l2/Adcy2/Arhgef11/Tap2/Tapbp/Cgas/Tnfrsf1a/Traf2/Traf5/Vegfa/H2-M10.5/Ifna13/Creb5/Akt3/Ifna15/E2f2/Map2k1/Map2k2/Map2k6/Mapk1/Mapk13/Mapk14/Mapk3/Creb3l1/Mapk12/Irf3/Tbk1/Gsk3b/Mtor/Rps6kb2/Tsc1/Gng11/H2-T-ps/Gm8909/Arhgef12/Tradd/Sting1/Nfatc4/Pik3cb/Akap13/Creb3l4 |
| mmu04914 | Progesterone-mediated oocyte maturation | 6,6474E-05 | Adcy3/Braf/Kif22/Raf1/Adcy6/Adcy9/Akt1/Akt2/Bub1/Ccna2/Ccnb2/Cdc25a/Cdc25b/Cdc25c/Cdk1/Cdk2/Cpeb1/Gnai1/Gnai2/Gnai3/Hsp90ab1/Hsp90aa1/Igf1/Igf1r/Mad1l1/Anapc1/Mos/Pde3b/Pik3ca/Pik3cd/Pik3r1/Pik3r2/Pik3r3/Prkaca/Prkacb/Plk1/Mapk11/Rps6ka1/Rps6ka2/Stk10/Aurka/Cpeb3/Adcy2/Cpeb2/Akt3/Map2k1/Mapk1/Mapk10/Mapk13/Mapk14/Mapk3/Mapk8/Mapk9/Ccnb1/Pkmyt1/Mapk12/Spdye4b/Anapc4/Cdc23/Mad2l1/Anapc7/Fzr1/Anapc5/Cdc26/Cpeb4/Cdc16/Spdya/Mad2l2/Pik3cb |
| mmu04917 | Prolactin signaling pathway | 6,6474E-05 | Gck/Raf1/Akt1/Akt2/Ccnd1/Ccnd2/Cga/Cish/Socs3/Socs1/Esr1/Esr2/Fos/Grb2/Hras/Irf1/Jak2/Nfkb1/Nras/Pik3ca/Pik3cd/Pik3r1/Pik3r2/Pik3r3/Mapk11/Prlr/Socs7/Rela/Shc1/Shc3/Slc2a2/Sos1/Sos2/Src/Stat1/Stat3/Stat5a/Stat5b/Socs2/Tnfrsf11a/Akt3/Map2k1/Map2k2/Mapk1/Mapk10/Mapk13/Mapk14/Mapk3/Mapk8/Mapk9/Shc4/Mapk12/Socs6/Socs5/Foxo3/Gsk3b/Pik3cb |
| mmu00561 | Glycerolipid metabolism | 0,0001254 | Gpat4/Dgkz/Dgkg/Dgkq/Aldh7a1/Aldh2/Aldh3a2/Akr1b3/Akr1b7/Cel/Dgka/Dgat1/Lpin1/Gpam/Lipg/Lpl/Pnliprp1/Pnliprp2/Plpp1/Dgkb/Mboat1/Lclat1/Tkfc/Dgkd/Gpat3/Glyctk/Mgll/Agpat3/Dgki/Dgkh/Plpp2/Agpat5/Dgke/Aldh9a1/Akr1a1/Lpin2/Lpin3/Pnpla2/Mboat2/Agpat2/Dgat2/Akr1b10/Plpp3/Agpat4/Mogat1/Pnlip/Agk/Plpp5/Aldh1b1 |
| mmu01212 | Fatty acid metabolism | 0,0001254 | Tecr/Acaca/Acat1/Acat2/Acadl/Acadm/Acadvl/Acaa1a/Acads/Acox1/Elovl3/Cpt1a/Cpt2/Acsl1/Fasn/Hadh/Hsd17b4/Elovl6/Ppt1/Scd1/Scd2/Scp2/Acsl6/Mcat/Hadhb/Cbr4/Acaa1b/Acsf3/Mecr/Scd3/Hacd1/Acsl5/Elovl1/Ppt2/Hsd17b12/Fads2/Hacd3/Hacd4/Acadsb/Elovl5/Hacd2/Oxsm/Ehhadh/Elovl7/Fads1/Cpt1c/Acox3/Elovl4/Hadha |
| mmu05145 | Toxoplasmosis | 0,00012773 | Ppif/Akt1/Akt2/Alox5/Birc3/Birc2/Bad/Bcl2/Bcl2l1/Casp3/Casp8/Casp9/Chuk/Socs1/Cycs/Gnai1/Gnai2/Gnai3/Gnao1/H2-Ab1/H2-Eb1/H2-DMa/Hspa1l/Hspa1b/Hspa2/Irgm1/Ifngr1/Ifngr2/Igtp/Ikbkb/Il10rb/Itga6/Itgb1/Jak1/Jak2/Lama1/Lama3/Lama4/Lama5/Lamb1/Lamb2/Lamb3/Lamc2/Ldlr/Ly96/Nfkb1/Nfkbia/Nfkbib/Nos2/Pdpk1/Mapk11/Rela/Stat1/Stat3/Tgfb1/Tgfb2/Tgfb3/Tlr4/Tnfrsf1a/Cd40/Traf6/Lamc1/Akt3/Tlr2/Map2k6/Map3k7/Mapk1/Mapk10/Mapk13/Mapk14/Mapk3/Mapk8/Mapk9/Irak4/Mapk12/Pik3cg/Pik3r5/Tyk2/Tab1/Tab2 |
| mmu03460 | Fanconi anemia pathway | 0,00014759 | Fancf/Faap24/Fancm/Rad51c/Blm/Brca1/Brca2/Ercc1/Fanca/Fancc/Hes1/Mlh1/Pms2/Rad51/Eme2/Rev3l/Rpa2/Fanci/Cenpx/Fancd2/Top3a/Top3b/Rmi2/Usp1/Palb2/Brip1/Atr/Poli/Eme1/Polk/Fan1/Ercc4/Slx4/Rev1/Fancg/Fancl/Ube2t/Wdr48/Cenps/Telo2/Faap100/Fance |
| mmu05016 | Huntington disease | 0,00015593 | Dnal1/Ppif/Dnah1/Cox6b1/Slc25a4/Ap2a1/Ap2a2/Ap2m1/Apaf1/Atp5a1/Atp5b/Atp5c1/Atp5pb/Atp5g1/Atp5j/Bax/Casp3/Casp8/Casp9/Clta/Cox5a/Cox5b/Cox6a1/Cox6c/Cox7a1/Cox7a2/Cox7c/Cox8a/Creb1/Creb3/Crebbp/Cycs/Dctn1/Dlg4/Dnah11/Gnaq/Gpx1/Gpx3/Grin1/Hap1/Hdac2/Htt/Itpr1/Bbc3/Ndufa2/Ndufa4/Ndufs4/Ndufv1/Nrf1/Plcb1/Plcb2/Plcb3/Plcb4/Pparg/Ppargc1a/Rest/Polr2a/Polr2c/Polr2j/Cox7a2l/Sod1/Sp1/Creb3l2/Hip1/Tfam/Rcor1/Tgm2/Uqcrc1/Vdac1/Vdac2/Vdac3/Ndufs8/Ndufs2/Dnah7b/Ndufs1/Atp5g3/Taf4/Ndufb6/Polr2b/Creb5/Ap2s1/Tbpl1/Creb3l1/Atp5o/Dnah2/Ep300/Dnah6/Ndufs6/Ndufa4l2/Dnai2/Hdac1/Dnal4/Ndufs5/Atp5d/Ndufb5/Ndufa3/Ndufa9/Uqcr10/Ndufb9/Ndufc1/Ndufa12/Polr2e/Cyc1/Uqcr11/Uqcrfs1/Ndufb7/Sdhd/Sdha/Uqcrc2/Atp5e/Ndufa6/Ndufb8/Ndufa10/Cltc/Gpx7/Dctn4/Sdhb/Polr2g/Atp5g2/Ndufb4/Ndufc2/Ndufb2/Ndufa5/Ndufa8/Dnai1/Polr2d/Gpx8/Dctn2/Ndufab1/Taf4b/Ndufv2/Ift57/Cltb/Ndufs7/Gpx6/Dnali1/Creb3l4 |
| mmu04714 | Thermogenesis | 0,00016114 | Adcy3/Kdm3a/Prkaa1/Prkaa2/Prkab2/Prkag2/Cox6b1/Actb/Actg1/Adcy6/Adcy9/Atf2/Atp5a1/Atp5b/Atp5c1/Atp5pb/Atp5g1/Atp5j/Atp5k/Bmp8a/Cnr1/Cox17/Cox5a/Cox5b/Cox6a1/Cox6c/Cox7a1/Cox7a2/Cox7c/Cox8a/Cpt1a/Cpt2/Creb1/Creb3/Acsl1/Fgfr1/Gnas/Grb2/Hras/Lipe/Ndufa2/Ndufa4/Ndufs4/Ndufv1/Npr1/Nras/Prkaca/Prkacb/Pparg/Ppargc1a/Prkab1/Prkag1/Prkg1/Mapk11/Rheb/Rps6/Rps6ka1/Rps6ka2/Cox7a2l/Smarca4/Smarcb1/Smarcc1/Sos1/Sos2/Creb3l2/Adcy2/Acsl6/Uqcrc1/Ndufs8/Cox15/Ndufs2/Ndufs1/Atp5g3/Ndufb6/Cox18/Creb5/Mgll/Arid1b/Map3k5/Mapk13/Mapk14/Creb3l1/Atp5l/Kdm3b/Atp5o/Mapk12/Dpf1/Frs2/Zfp516/Ndufs6/Ndufa4l2/Acsl5/Sirt6/Coa3/Actl6a/Mlst8/Mtor/Slc25a20/Smarce1/Atp5j2/Rps6kb2/Ndufs5/Tsc1/Atp5d/Ndufb5/Ndufa3/Ndufa9/Uqcr10/Ndufb9/Cox16/Cox20/Ndufc1/Ndufa12/Cyc1/Uqcr11/Uqcrfs1/Pnpla2/Ndufb7/Sdhd/Sdha/Smarcd3/Uqcrc2/Atp5e/Ndufa6/Smarca2/Ndufb8/Ndufa10/Akt1s1/Sdhb/Coa6/Atp5g2/Cox19/Coa4/Ndufb4/Ndufc2/Ndufb2/Ndufa5/Ndufa8/Ndufaf4/Ndufaf1/Coa7/Dpf3/Ndufab1/Prdm16/Ndufv2/Rptor/Ndufs7/Ndufaf2/Ndufaf6/Cpt1c/Creb3l4/Klb/Smarcd2/Smarcd1/Arid1a/Kdm1a |
| mmu05216 | Thyroid cancer | 0,00018171 | Ddb2/Tpr/Braf/Bak1/Bax/Ctnnb1/Ccnd1/Cdh1/Cdkn1a/Gadd45a/Hras/Lef1/Myc/Gadd45b/Nras/Pparg/Ret/Rxra/Rxrg/Tcf7/Tcf7l1/Tcf7l2/Tfg/Gadd45g/Map2k1/Map2k2/Mapk1/Mapk3/Polk/Ncoa4/Tpm3/Ccdc6 |
| mmu05218 | Melanoma | 0,00018307 | Ddb2/Braf/Raf1/Akt1/Akt2/Bad/Bak1/Bax/Ccnd1/Cdh1/Cdk4/Cdk6/Cdkn1a/Cdkn2a/Gadd45a/E2f1/E2f3/Egf/Egfr/Fgf1/Fgf15/Fgf18/Fgf2/Fgf5/Fgf8/Fgf9/Fgfr1/Hras/Igf1/Igf1r/Mdm2/Met/Mitf/Gadd45b/Nras/Pdgfa/Pdgfb/Pdgfra/Pik3ca/Pik3cd/Pik3r1/Pik3r2/Pik3r3/Pten/Rb1/Akt3/Gadd45g/E2f2/Map2k1/Map2k2/Mapk1/Mapk3/Polk/Pdgfc/Pdgfd/Pik3cb |
| mmu05167 | Kaposi sarcoma-associated herpesvirus infection | 0,00019782 | Atg14/Ticam1/Raf1/H2-Q6/H2-Q9/Angpt2/Akt1/Akt2/Bak1/Bax/Bid/Calm1/Calm2/Calm3/Casp3/Casp8/Casp9/Ctnnb1/Ccnd1/Cdk4/Cdk6/Cdkn1a/Chuk/Creb1/Crebbp/Cycs/E2f1/E2f3/Fadd/Fas/Fgf2/Fos/Tlr3/Gnb1/Gng10/Gng12/Gng2/Gng5/Gng7/Gng8/Cxcl1/H2-Bl/H2-D1/H2-K1/H2-M3/H2-Q1/H2-Q10/H2-Q2/H2-Q4/H2-T22/H2-T23/H2-T24/Hck/Hif1a/Hras/Icam1/Ifnar1/Ifnar2/Ifngr1/Ikbkb/Il6st/Irf9/Itpr1/Itpr2/Itpr3/Jak1/Jak2/Jun/Lyn/Mapkapk2/Myc/Nfatc1/Nfatc2/Nfatc3/Nfkb1/Nfkbia/Nras/Pdgfb/Pik3ca/Pik3cd/Pik3r1/Pik3r2/Pik3r3/Plcg1/Ppp3ca/Ppp3cb/Ppp3cc/Ppp3r1/Mapk11/Eif2ak2/Ptgs2/Rac1/Rb1/Rela/Src/Stat1/Stat2/Stat3/Syk/Tnfrsf1a/Traf2/Traf3/Ubb/Vegfa/H2-M10.5/Pik3c3/Zfp36/Ifna13/Plcg2/Akt3/Ifna15/E2f2/Map2k1/Map2k2/Map2k4/Map2k6/Map2k7/Mapk1/Mapk10/Mapk13/Mapk14/Mapk3/Mapk8/Mapk9/Prex1/Mapk12/Pik3cg/Pik3r5/Ep300/Irf7/Irf3/Rcan1/Tyk2/Becn1/Tbk1/Gabarap/Ikbke/Gsk3b/Mtor/Gng11/H2-T-ps/Gm8909/Atg3/Tradd/Nfatc4/Pik3cb/Gabarapl2 |
| mmu00230 | Purine metabolism | 0,00023474 | Ak6/Nt5m/Adcy3/Gmpr2/Nt5c3/Atic/Ampd2/Pde4c/Ada/Adcy6/Adcy9/Adk/Adsl/Adssl1/Adss/Ak1/Ak2/Ak4/Ampd3/Aprt/Entpd1/Entpd2/Entpd6/Entpd5/Dck/Fhit/Gart/Gda/Gucy2e/Guk1/Itpa/Nme7/Nme1/Nme2/Npr1/Pde1b/Pde1c/Pde3b/Pde4a/Pde4b/Pde6d/Pde7a/Pde8a/Pde9a/Enpp1/Pkm/Pklr/Pnp/Rrm1/Rrm2/Pde2a/Enpp3/Adcy2/Entpd3/Pde8b/Xdh/Enpp4/Gmps/Prune1/Ak5/Npr2/Nt5c1a/Ppat/Gucy1a2/Pfas/Pde4d/Impdh1/Impdh2/Nt5e/Papss1/Papss2/Pde10a/Pde5a/Dguok/Pde7b/Prps1l3/Rrm2b/Nt5c/Nudt5/Gucy1b1/Nme6/Ak3/Nme4/Gucy1a1/Gmpr/Adprm/Ntpcr/Pgm2/Pnp2/Nt5c3b/Hddc3/Ak8/Nudt16/Cant1/Nt5c2/Urah/Allc |
| mmu04962 | Vasopressin-regulated water reabsorption | 0,00025023 | Adcy3/Dync2h1/Adcy6/Adcy9/Aqp2/Aqp3/Creb1/Creb3/Dctn1/Dync1h1/Dync1i1/Dync1i2/Arhgdig/Gnas/Nsf/Prkaca/Prkacb/Arhgdia/Rab11b/Rab5b/Rab5c/Creb3l2/Stx4a/Dync2li1/Vamp2/Dctn6/Creb5/Dync1li2/Dync1li1/Creb3l1/Rab5a/Rab11a/Dctn4/Dynll2/Dctn2/Creb3l4 |
| mmu05010 | Alzheimer disease | 0,00025395 | Cox6b1/Adam10/Adam17/Apaf1/Apbb1/Apoe/App/Atp2a2/Atp5a1/Atp5b/Atp5c1/Atp5pb/Atp5g1/Atp5j/Bad/Bid/Cacna1c/Cacna1d/Calm1/Calm2/Calm3/Capn1/Capn2/Casp3/Casp7/Casp8/Casp9/Cdk5/Cdk5r1/Cox5a/Cox5b/Cox6a1/Cox6c/Cox7a1/Cox7a2/Cox7c/Cox8a/Cycs/Eif2ak3/Fadd/Fas/Gapdh/Gnaq/Grin1/Grin2d/Itpr1/Itpr2/Itpr3/Lpl/Lrp1/Mme/Mapt/Ndufa2/Ndufa4/Ndufs4/Ndufv1/Plcb1/Plcb2/Plcb3/Plcb4/Ppp3ca/Ppp3cb/Ppp3cc/Ppp3r1/Psen1/Psen2/Rtn3/Cox7a2l/Snca/Aph1b/Tnfrsf1a/Uqcrc1/Ndufs8/Aph1a/Atf6/Ndufs2/Ndufs1/Atp5g3/Ndufb6/Bace1/Mapk1/Mapk3/Atp5o/Ndufs6/Ndufa4l2/Bace2/Gsk3b/Ndufs5/Atp5d/Ndufb5/Ndufa3/Ndufa9/Uqcr10/Ndufb9/Ndufc1/Ndufa12/Cyc1/Uqcr11/Uqcrfs1/Ndufb7/Sdhd/Sdha/Uqcrc2/Atp5e/Ndufa6/Ndufb8/Ndufa10/Sdhb/Atp5g2/Ndufb4/Ndufc2/Ndufb2/Ndufa5/Aph1c/Ndufa8/Rtn4/Ndufab1/Ndufv2/Ndufs7/Ern1 |
| mmu05235 | PD-L1 expression and PD-1 checkpoint pathway in cancer | 0,00029003 | Ticam1/Raf1/Akt1/Akt2/Tirap/Cd247/Chuk/Csnk2a1/Csnk2a2/Csnk2b/Egf/Egfr/Fos/Hif1a/Hras/Ifngr1/Ifngr2/Ikbkb/Jak1/Jak2/Jun/Nfatc1/Nfatc2/Nfatc3/Nfkb1/Nfkbia/Nfkbib/Nfkbie/Nras/Pik3ca/Pik3cd/Pik3r1/Pik3r2/Pik3r3/Plcg1/Ppp3ca/Ppp3cb/Ppp3cc/Ppp3r1/Mapk11/Pten/Ptpn11/Rela/Stat1/Stat3/Tlr4/Traf6/Ticam2/Zap70/Akt3/Tlr2/Map2k1/Map2k2/Map2k6/Map3k3/Mapk1/Mapk13/Mapk14/Mapk3/Mapk12/Batf/Mtor/Rps6kb2/Cd274/Pik3cb/Eml4 |
| mmu04072 | Phospholipase D signaling pathway | 0,00032695 | Dnm3/Adcy3/Dgkz/Grm3/Grm6/Raf1/Dgkg/Dgkq/Adcy6/Adcy9/Akt1/Akt2/Arf1/Arf6/Dgka/Dnm1/Dnm2/Egf/Egfr/F2/F2r/Fcer1g/Fyn/Gab1/Gab2/Gna12/Gnas/Lpar1/Grb2/Grm8/Hras/Insr/Kit/Kitl/Mras/Nras/Pdgfa/Pdgfb/Pdgfra/Pik3ca/Pik3cd/Pik3r1/Pik3r2/Pik3r3/Pip5k1c/Pip5k1b/Pip5k1a/Pla2g4a/Plcb1/Plcb2/Plcb3/Plcb4/Plcg1/Pld1/Pld2/Plpp1/Cyth1/Cyth2/Cyth3/Ptk2b/Ptpn11/Ralgds/Rheb/Rras/Shc1/Shc3/Sos1/Sos2/Sphk1/Syk/Adcy2/Dgkb/Rapgef3/Dgkd/Plcg2/Akt3/Map2k1/Map2k2/Mapk1/Mapk3/Shc4/Agpat3/Pik3cg/Dgki/Pik3r5/Dgkh/Plpp2/Agpat5/Lpar2/Pdgfc/Rala/Dgke/Rapgef4/Mtor/Ralb/Tsc1/Lpar3/Rras2/Lpar6/Agpat2/Plpp3/Agpat4/Pdgfd/Pik3cb |
| mmu01040 | Biosynthesis of unsaturated fatty acids | 0,00032695 | Tecr/Acaa1a/Acox1/Elovl3/Hsd17b4/Elovl6/Acot2/Acot3/Scd1/Scd2/Scp2/Acot5/Acaa1b/Acot1/Scd3/Hacd1/Elovl1/Hsd17b12/Fads2/Hacd3/Hacd4/Elovl5/Acot7/Hacd2/Elovl7/Fads1/Acox3/Elovl4 |
| mmu05017 | Spinocerebellar ataxia | 0,00032969 | Atg14/Atxn3/Afg3l1/Akt1/Akt2/Atp2a2/Cacna1a/Rb1cc1/Dab1/Fgf14/Gnaq/Grin1/Grin2d/Itpr1/Itpr2/Itpr3/Kcnc3/Pik3ca/Pik3cd/Pik3r1/Pik3r2/Pik3r3/Prkcb/Prkcg/Plcb1/Plcb2/Plcb3/Plcb4/Rbpj/Rora/Atxn1/Atxn2/Slc1a6/Sp1/Sptbn2/Traf2/Ulk1/Vldlr/Xbp1/Pik3c3/Twnk/Ambra1/Gtf2b/Atxn2l/Tbpl1/Akt3/Map3k5/Mapk10/Mapk8/Mapk9/Ulk2/Atg2a/Atxn1l/Wipi1/Atxn10/Becn1/Kcnd3/Mtor/Nrbf2/Oma1/Atg101/Afg3l2/Cic/Opa1/Pik3cb/Wipi2/Pik3r4/Ern1/Pum1/Pum2/Kat5 |
| mmu05100 | Bacterial invasion of epithelial cells | 0,00032969 | Dnm3/Actb/Actg1/Arpc1b/Ctnna1/Ctnnb1/Cav1/Cav2/Cbl/Cd2ap/Cdc42/Cdh1/Clta/Bcar1/Crk/Crkl/Cttn/Dnm1/Dnm2/Elmo2/Elmo1/Ptk2/Gab1/Itga5/Itgb1/Met/Pik3ca/Pik3cd/Pik3r1/Pik3r2/Pik3r3/Pxn/Rac1/Septin8/Shc1/Shc3/Src/Ctnna3/Vcl/Elmo3/Wasf2/Shc4/Dock1/Septin11/Septin9/Septin1/Rhog/Arpc3/Arhgef26/Cltc/Mad2l2/Wasl/Cltb/Pik3cb/Arpc2/Arhgap10 |
| mmu04370 | VEGF signaling pathway | 0,00032969 | Mapkapk3/Raf1/Akt1/Akt2/Bad/Casp9/Cdc42/Ptk2/Hras/Hspb1/Kdr/Rac3/Mapkapk2/Nfatc2/Nras/Pik3ca/Pik3cd/Pik3r1/Pik3r2/Pik3r3/Prkcb/Prkcg/Pla2g4a/Plcg1/Ppp3ca/Ppp3cb/Ppp3cc/Ppp3r1/Mapk11/Ptgs2/Pxn/Rac1/Rac2/Sphk1/Src/Vegfa/Plcg2/Akt3/Map2k1/Map2k2/Mapk1/Mapk13/Mapk14/Mapk3/Mapk12/Pik3cb |
| mmu05221 | Acute myeloid leukemia | 0,00032969 | Braf/Raf1/Cebpe/Akt1/Akt2/Bad/Runx1/Ccna2/Ccnd1/Cd14/Cebpa/Chuk/Eif4ebp1/Fcgr1/Grb2/Hras/Ikbkb/Itgam/Jup/Kit/Lef1/Myc/Nfkb1/Nras/Per2/Pik3ca/Pik3cd/Pik3r1/Pik3r2/Pik3r3/Pim1/Pml/Ppard/Rara/Rela/Spi1/Sos1/Sos2/Stat3/Stat5a/Stat5b/Tcf7/Tcf7l1/Tcf7l2/Zbtb16/Akt3/Map2k1/Map2k2/Mapk1/Mapk3/Mtor/Rps6kb2/Dusp6/Pik3cb |
| mmu05202 | Transcriptional misregulation in cancer | 0,00038326 | Etv5/Nsd2/Ddb2/Jmjd1c/Supt3/Cebpe/Birc3/Atf1/Atm/Bak1/Bax/Bcl2l1/Bcl6/Bmi1/Runx2/Runx1/Ccna2/Ccnd2/Cd14/Cdkn1a/Cdkn1b/Cdkn2c/Cebpa/Cebpb/Gadd45a/Ddx5/Elk4/Etv1/Etv6/Ewsr1/Bmp2k/Ptk2/Fcgr1/Fli1/H3c14/H3f3a/H3f3b/Hdac2/Hhex/Hmga2/Hpgd/Id2/Igf1/Igf1r/Itgam/Itgb7/Jup/Klf3/Ldb1/Lmo2/Smad1/Maf/Max/Mdm2/Mef2c/Meis1/Men1/Met/Mitf/Mlf1/Aff1/Mmp9/Myc/Gadd45b/Nfkb1/Mycn/Nr4a3/Pax3/Pax5/Pbx1/Pbx3/Pdgfa/Etv4/Per2/Cdk14/Plau/Pml/Pparg/Prom1/Rara/Rel/Rela/Rxra/Rxrg/Ncor1/Spi1/Six1/Six4/Sp1/Spint1/Dot1l/Slc45a3/Kmt2a/Zeb1/Tcf3/Tgfbr2/Tlx1/Cd40/Traf1/Zbtb17/Fus/Zbtb16/Gadd45g/Fev/H3c7/Ss18/Polk/H3c3/H3c4/H3c2/H3c6/H3c10/H3c11/H3c13/Hdac1/Tmprss2/Fut8/Nupr1/Foxo1/Mllt1/Dusp6/Aspscr1/Mllt3/Taf15/Nfkbiz/Prcc |
| mmu04066 | HIF-1 signaling pathway | 0,00038326 | Camk2d/Egln1/Egln2/Egln3/Angpt1/Angpt2/Akt1/Akt2/Aldoa/Aldoc/Arnt/Bcl2/Cdkn1a/Cdkn1b/Crebbp/Edn1/Egf/Egfr/Eif4ebp1/Eno1/Eno2/Epo/Erbb2/Gapdh/Hif1a/Hk1/Hk2/Hmox1/Ifngr1/Ifngr2/Igf1/Igf1r/Il6ra/Insr/Ldha/Ldhb/Ltbr/Pfkfb3/Mknk1/Mknk2/Nfkb1/Nos2/Pfkl/Pfkm/Pik3ca/Pik3cd/Pik3r1/Pik3r2/Pik3r3/Prkcb/Prkcg/Plcg1/Rela/Rps6/Slc2a1/Stat3/Hkdc1/Tek/Tlr4/Tfrc/Vegfa/Vhl/Eno4/Pdk1/Aldob/Plcg2/Akt3/Map2k1/Map2k2/Mapk1/Mapk3/Eif4e2/Ep300/Pfkp/Rbx1/Mtor/Rps6kb2/Elob/Pdhb/Cul2/Pik3cb |
| mmu03030 | DNA replication | 0,00038514 | Rfc4/Fen1/Lig1/Mcm3/Mcm2/Mcm4/Mcm5/Mcm6/Pcna/Pola2/Pold1/Pold2/Pole/Pole2/Prim1/Prim2/Rfc1/Rfc2/Rnaseh1/Rpa2/Dna2/Ssbp1/Pole4/Rnaseh2b/Pold3/Rnaseh2c/Rfc3/Rnaseh2a/Pold4/Rfc5 |
| mmu03013 | RNA transport | 0,00041021 | Sap18b/Eif3j2/Rpp25/Nup107/Xpo1/Thoc5/Pom121/Eif2b3/Tpr/Sec13/Elac1/Semp2l2b/Tgs1/Pop5/Eef1a1/Eef1a2/Eif1a/Eif2s1/Eif2b4/Eif3a/Eif4a1/Eif4a2/Eif4ebp1/Eif4ebp2/Eif4g2/Fxr1/Kpnb1/Eif3e/Nup155/Sumo2/Magoh/Nup50/Pabpc1/Pnn/Nup88/Eif4a3/Ran/Ranbp2/Rangap1/Rasl2-9/Pom121l2/Upf1/Rnps1/Cyfip1/Smn1/Sumo3/Rpp40/Eif4g1/Strap/Eif1/Eif2b1/Tacc3/Gemin5/Alyref/Eif2b2/Eif5/Nup153/Paip1/Ube2i/Sumo1/Eif2b5/Thoc1/Eif5b/Rpp38/Nup214/Pabpc4/Eif4g3/Nupl2/Nup133/Fxr2/Pabpc4l/Nup54/Eif4e2/Prmt5/Gemin4/Eif3b/Upf2/Pabpc1l/Nup85/Srrm1/Nxf1/Eif3g/Ddx20/Rpp30/Nup210/Eif3i/Eif3d/Alyref2/Acin1/Eif3c/Nxt1/Nup160/Snupn/Eif3f/Pop4/Magohb/Rae1/Upf3a/Rpp14/Eif2s2/Gemin6/Rpp21/Eif3h/Elac2/Eif1b/Nup35/Gemin7/Nup37/Nup43/Rpp25l/Trnt1/Nup205/Nup93/Nupl1/Seh1l/Xpo5/Ndc1/Xpot/Thoc3/Pop7/Eif4b/Senp2/Cyfip2/Nmd3 |
| mmu04510 | Focal adhesion | 0,0004214 | Itga9/Mylk/Rapgef1/Actn1/Braf/Rap1a/Raf1/Actb/Actg1/Akt1/Akt2/Birc3/Birc2/Arhgap5/Bad/Bcl2/Capn2/Ctnnb1/Cav1/Cav2/Ccnd1/Ccnd2/Ccnd3/Cdc42/Chad/Col1a1/Comp/Bcar1/Crk/Crkl/Diaph1/Egf/Egfr/Erbb2/Ptk2/Flt4/Fyn/Grb2/Hras/Igf1/Igf1r/Itga2/Itga2b/Itga3/Itga5/Itga6/Itga7/Itgav/Itgb1/Itgb5/Itgb6/Itgb7/Jun/Kdr/Lama1/Lama3/Lama4/Lama5/Lamb1/Lamb2/Lamb3/Lamc2/Parvb/Rac3/Met/Mylpf/Ppp1r12a/Pak1/Pdgfa/Pdgfb/Pdgfra/Pdpk1/Pgf/Pik3ca/Pik3cd/Pik3r1/Pik3r2/Pik3r3/Pip5k1c/Prkcb/Prkcg/Ppp1cb/Ppp1cc/Pten/Itgb4/Pxn/Rac1/Rac2/Rasgrf1/Rock1/Rock2/Shc1/Shc3/Sos1/Sos2/Spp1/Src/Pak6/Rap1b/Thbs1/Vasp/Vav1/Vav2/Vcl/Vegfa/Vegfb/Vegfc/Vtn/Pak2/Lamc1/Zyx/Ppp1r12c/Akt3/Itga8/Map2k1/Mapk1/Mapk10/Mapk3/Mapk8/Mapk9/Shc4/Flnb/Itgb8/Ppp1r12b/Dock1/Pdgfc/Gsk3b/Vav3/Parva/Actn4/Myl12b/Flnc/Tln2/Pdgfd/Pik3cb |
| mmu05217 | Basal cell carcinoma | 0,00045225 | Ddb2/Apc/Axin1/Axin2/Bak1/Bax/Bmp2/Ctnnb1/Cdkn1a/Gadd45a/Dvl1/Dvl2/Dvl3/Fzd1/Fzd3/Fzd4/Fzd5/Fzd6/Fzd7/Fzd8/Fzd9/Gli1/Gli2/Hhip/Kif7/Lef1/Gadd45b/Ptch1/Shh/Wnt8a/Tcf7/Tcf7l1/Tcf7l2/Wnt9a/Wnt1/Wnt10b/Wnt11/Wnt9b/Wnt2b/Wnt5a/Wnt5b/Wnt6/Apc2/Gadd45g/Sufu/Polk/Smo/Gsk3b/Fzd2 |
| mmu04540 | Gap junction | 0,00048027 | Adcy3/Csnk1d/Raf1/Adcy6/Adcy9/Adrb1/Cdk1/Egf/Egfr/Gja1/Gna11/Gnai1/Gnai2/Gnai3/Gnaq/Gnas/Lpar1/Grb2/Hras/Itpr1/Itpr2/Itpr3/Nras/Pdgfa/Pdgfb/Pdgfra/Prkaca/Prkacb/Prkcb/Prkcg/Plcb1/Plcb2/Plcb3/Plcb4/Prkg1/Sos1/Sos2/Src/Adcy2/Tjp1/Tuba1a/Tuba1b/Tuba4a/Tuba1c/Tubb2a/Tubb3/Tubb5/Tubb4b/Gucy1a2/Tubal3/Map2k5/Mapk7/Map2k1/Map2k2/Map3k2/Mapk1/Mapk3/Tuba8/Gucy1b1/Pdgfc/Gucy1a1/Tubb6/Pdgfd/Tubb2b |
| mmu03440 | Homologous recombination | 0,00049677 | Babam2/Rad51c/Atm/Bard1/Blm/Brca1/Brca2/Mre11a/Pold1/Pold2/Rad50/Rad51/Rad51b/Rad51d/Rad52/Rad54l/Rpa2/Uimc1/Sem1/Top3a/Top3b/Rbbp8/Palb2/Topbp1/Brip1/Eme1/Nbn/Ssbp1/Xrcc2/Rad54b/Pold3/Babam1/Pold4/Abraxas1 |
| mmu01230 | Biosynthesis of amino acids | 0,00054372 | Psph/Psat1/Cth/Shmt2/Mat2b/Gpt2/Acy1/Asl/Aco1/Aco2/Aldoa/Aldoc/Mat1a/Arg1/Arg2/Ass1/Bcat1/Bcat2/Cbs/Cs/Eno1/Eno2/Gapdh/Got1/Got2/Idh1/Idh3b/Pah/Pcx/Pfkl/Pfkm/Pgam1/Pkm/Pklr/Rpia/Shmt1/Pycr1/Taldo1/Nags/Tkt/Tpi1/Eno4/Aldob/Mat2a/Phgdh/Mtr/Sdsl/Idh2/Asns/Prps1l3/Pfkp/Aldh18a1/Pycrl/Rpe/Idh3a/Pycr2/Tha1/Gpt |
| mmu04920 | Adipocytokine signaling pathway | 0,0005785 | Acacb/Prkaa1/Prkaa2/Prkab2/Prkag2/Npy/Agrp/Akt1/Akt2/Cd36/Chuk/Socs3/Cpt1a/Acsl1/G6pc/Ikbkb/Irs1/Irs3/Jak2/Nfkb1/Nfkbia/Nfkbib/Nfkbie/Pomc/Ppara/Ppargc1a/Prkab1/Prkag1/Ptpn11/Rela/Rxra/Rxrg/Slc2a1/Slc2a4/Camkk2/Stat3/Stk11/Acsl6/Tnfrsf1a/Tnfrsf1b/Traf2/Akt3/Mapk10/Mapk8/Mapk9/Irs2/Acsl5/Mtor/G6pc3/Adipor2/Tradd/Adipor1/Pck2/Cpt1c |
| mmu00670 | One carbon pool by folate | 0,0005785 | Mthfsl/Aldh1l1/Mthfs/Shmt2/Atic/Mthfd1/Dhfr/Gart/Mthfd2/Mthfr/Shmt1/Aldh1l2/Tyms/Mtr/Mthfd1l/Amt/Mthfd2l/Mtfmt |
| mmu00900 | Terpenoid backbone biosynthesis | 0,00059538 | Fdps/Acat1/Acat2/Fntb/Fnta/Ggps1/Hmgcr/Hmgcs2/Mvk/Mvd/Rce1/Hmgcs1/Zmpste24/Idi1/Nus1/Pdss1/Icmt/Pcyox1/Dhdds/Pmvk/Pdss2 |
| mmu00480 | Glutathione metabolism | 0,00059538 | Gstt3/Ggct/Pgd/Ggt1/Gclc/Gclm/Gpx1/Gpx3/Gsr/Gss/Gsta3/Gsta4/Gstm1/Gstm2/Gstm3/Gstm4/Gstm5/Gstp1/Gstt1/Gstt2/Gsto1/Idh1/Anpep/Odc1/Rrm1/Rrm2/Ggt7/Srm/Mgst2/Gstp3/Ggt5/Idh2/Rrm2b/Mgst1/Gpx4/Txndc12/Nat8f1/Mgst3/Lap3/Gpx7/Chac2/Gsto2/Gstm7/Nat8/Chac1/Gpx8/Ggt6/Oplah/Gpx6/Gstk1 |
| mmu04213 | Longevity regulating pathway - multiple species | 0,00059955 | Adcy3/Prkaa1/Prkaa2/Prkab2/Prkag2/Adcy6/Adcy9/Akt1/Akt2/Atg5/Cat/Cryab/Eif4ebp2/Hdac2/Foxa2/Hras/Hspa1l/Hspa1b/Hspa2/Igf1/Igf1r/Insr/Irs1/Irs3/Nras/Pik3ca/Pik3cd/Pik3r1/Pik3r2/Pik3r3/Prkaca/Prkacb/Prkab1/Prkag1/Clpb/Sod1/Adcy2/Akt3/Irs2/Hdac1/Foxo1/Foxo3/Mtor/Rps6kb2/Akt1s1/Rptor/Pik3cb/Sirt1 |
| mmu04151 | PI3K-Akt signaling pathway | 0,00060806 | Itga9/Prkaa1/Prkaa2/Pkn2/Raf1/Angpt1/Angpt2/Akt1/Akt2/Areg/Atf2/Atf4/Bad/Bcl2/Bcl2l1/Bcl2l11/Brca1/Casp9/Ccnd1/Ccnd2/Ccnd3/Ccne1/Ccne2/Cdc37/Cdk2/Cdk4/Cdk6/Cdkn1a/Cdkn1b/Chad/Chrm1/Chuk/Col1a1/Comp/Creb1/Creb3/Atf6b/Csf1/Efna1/Efna2/Efna3/Efna4/Efna5/Egf/Egfr/Eif4ebp1/Epha2/Epo/Epor/Erbb2/Erbb3/Erbb4/F2r/Ptk2/Fgf1/Fgf15/Fgf18/Fgf2/Fgf5/Fgf8/Fgf9/Fgfr1/Fgfr2/Fgfr3/Fgfr4/Flt3l/Flt4/G6pc/Ghr/Gnb1/Gng10/Gng12/Gng2/Gng5/Gng7/Gng8/Lpar1/Grb2/Magi1/Gys1/Nr4a1/Hras/Hsp90ab1/Hsp90aa1/Ifnar1/Ifnar2/Igf1/Igf1r/Ikbkb/Il3ra/Il4ra/Il6ra/Il7/Insr/Irs1/Itga2/Itga2b/Itga3/Itga5/Itga6/Itga7/Itgav/Itgb1/Itgb5/Itgb6/Itgb7/Jak1/Jak2/Jak3/Kdr/Kit/Lama1/Lama3/Lama4/Lama5/Lamb1/Lamb2/Lamb3/Lamc2/Sgk3/Mcl1/Mdm2/Met/Kitl/Myb/Myc/Nfkb1/Nras/Osmr/Pdgfa/Pdgfb/Pdgfra/Pdpk1/Pgf/Pik3ca/Pik3cd/Pik3r1/Pik3r2/Pik3r3/Ppp2ca/Ppp2cb/Prlr/Pten/Itgb4/Rac1/Rbl2/Rela/Rheb/Rps6/Rxra/Sgk1/Sos1/Sos2/Spp1/Creb3l2/Stk11/Syk/Tek/Ppp2r5d/Tgfa/Thbs1/Tlr4/Vegfa/Vegfb/Vegfc/Vtn/Ppp2r5b/Ywhae/Ywhag/Ywhah/Ywhaq/Ywhaz/Lamc1/Ppp2r5a/Ifna13/Creb5/Ppp2r3a/Akt3/Tlr2/Itga8/Ifna15/Chrm2/Phlpp2/Pkn3/Map2k1/Map2k2/Mapk1/Mapk3/Creb3l1/Ppp2r5c/Ppp2r5e/Eif4e2/Sgk2/Pik3cg/Pik3r5/Pkn1/Itgb8/Magi2/Ppp2r1a/Ppp2r2d/Lpar2/Ywhab/Pdgfc/Foxo3/Gsk3b/Mlst8/Mtor/Rps6kb2/Ppp2r3c/Tsc1/Lpar3/Gng11/Lpar6/G6pc3/Pdgfd/Ppp2r2a/Ppp2r2b/Ppp2r1b/Crtc2/Rptor/Pck2/Ddit4/Pik3cb/Eif4b/Them4/Creb3l4/Ntf5/Pik3ap1 |
| mmu04922 | Glucagon signaling pathway | 0,00068893 | Acacb/Phkb/Gck/Ppp4r3b/Prkaa1/Acaca/Camk2d/Prkaa2/Prkab2/Prkag2/Pygb/Pygl/Akt1/Akt2/Atf2/Atf4/Calm1/Calm2/Calm3/Cpt1a/Creb1/Creb3/Crebbp/Fbp2/Fbp1/G6pc/Gnaq/Gnas/Gys1/Itpr1/Itpr2/Itpr3/Ldha/Ldhb/Sik1/Pde3b/Pfkl/Pfkm/Pgam1/Pkm/Prkaca/Prkacb/Plcb1/Plcb2/Plcb3/Plcb4/Ppara/Ppargc1a/Ppp3ca/Ppp3cb/Ppp3cc/Ppp3r1/Prkab1/Prkag1/Pygm/Slc2a1/Slc2a2/Creb3l2/Adcy2/Creb5/Sik2/Akt3/Creb3l1/Ep300/Ppp4c/Pfkp/Foxo1/Pdhb/G6pc3/Phkg2/Crtc2/Pck2/Cpt1c/Creb3l4/Sirt1 |
| mmu04666 | Fc gamma R-mediated phagocytosis | 0,00072214 | Raf1/Akt1/Akt2/Arf6/Arpc1b/Cdc42/Cfl1/Cfl2/Crk/Crkl/Asap1/Dnm2/Fcgr1/Gab2/Hck/Inppl1/Limk1/Limk2/Lyn/Marcks/Marcksl1/Myo10/Pak1/Pik3ca/Pik3cd/Pik3r1/Pik3r2/Pik3r3/Pip5k1c/Pip5k1b/Pip5k1a/Prkcb/Prkcg/Prkcd/Prkce/Pla2g4a/Plcg1/Pld1/Pld2/Plpp1/Rac1/Rac2/Scin/Sphk1/Syk/Asap2/Vasp/Vav1/Vav2/Gsn/Asap3/Plcg2/Akt3/Wasf2/Map2k1/Mapk1/Mapk3/Bin1/Plpp2/Pla2g6/Arpc3/Vav3/Rps6kb2/Plpp3/Pik3cb/Arpc2 |
| mmu03420 | Nucleotide excision repair | 0,00074808 | Rfc4/Ddb2/Cdk7/Ddb1/Ercc1/Ercc2/Ercc3/Gtf2h1/Gtf2h4/Lig1/Mnat1/Pcna/Pold1/Pold2/Pole/Pole2/Rad23a/Rad23b/Rfc1/Rfc2/Rpa2/Xpa/Xpc/Ercc5/Gtf2h2/Ercc6/Ercc4/Rbx1/Ccnh/Pole4/Pold3/Rfc3/Pold4/Ercc8/Rfc5 |
| mmu05230 | Central carbon metabolism in cancer | 0,00077544 | Sco2/Gck/Raf1/Akt1/Akt2/Egfr/Erbb2/Fgfr1/Fgfr2/Fgfr3/Gls/Hif1a/Hk1/Hk2/Hras/Kit/Ldha/Ldhb/Met/Myc/Nras/Pdgfra/Pfkl/Pfkm/Pgam1/Pik3ca/Pik3cd/Pik3r1/Pik3r2/Pik3r3/Pkm/Pten/Ret/Slc1a5/Slc2a1/Slc2a2/Slc7a5/Hkdc1/Gls2/Pdk1/Akt3/Map2k1/Map2k2/Mapk1/Mapk3/Tigar/Sirt6/Pfkp/Mtor/Pdhb/Pik3cb |
| mmu00520 | Amino sugar and nucleotide sugar metabolism | 0,00080058 | Gck/Uap1/Cyb5r3/Pgm3/Mpi/Cmas/Gfpt1/Gfpt2/Galk1/Gpi1/Hexa/Hexb/Hk1/Hk2/Hkdc1/Ugp2/Gmds/Gfus/Ugdh/Uap1l1/Cyb5rl/Fcsk/Amdhd2/Gnpda1/Cyb5r4/Pmm1/Gne/Pmm2/Gnpnat1/Nagk/Pgm2/Nanp/Uxs1/Gnpda2/Gmppa/Cyb5r1/Npl/Gale/Nans |
| mmu00640 | Propanoate metabolism | 0,00080058 | Acacb/Aldh6a1/Acaca/Acat1/Acat2/Pcca/Acads/Acox1/Bckdha/Bckdhb/Dbt/Dld/Ldha/Ldhb/Mmut/Sucla2/Suclg2/Hibch/Abat/Acss3/Echdc1/Suclg1/Mlycd/Pccb/Acss1/Ehhadh/Acox3/Hadha |
| mmu04014 | Ras signaling pathway | 0,00108629 | Rap1a/Raf1/Abl1/Abl2/Rasa2/Angpt1/Angpt2/Akt1/Akt2/Arf6/Bad/Bcl2l1/Calm1/Calm2/Calm3/Cdc42/Chuk/Csf1/Efna1/Efna2/Efna3/Efna4/Efna5/Egf/Egfr/Epha2/Fgf1/Fgf15/Fgf18/Fgf2/Fgf5/Fgf8/Fgf9/Fgfr1/Fgfr2/Fgfr3/Fgfr4/Flt3l/Flt4/Gab1/Gab2/Gnb1/Gng10/Gng12/Gng2/Gng5/Gng7/Gng8/Grb2/Grin1/Hras/Igf1/Igf1r/Ikbkb/Insr/Kdr/Kit/Ksr1/Rac3/Met/Kitl/Afdn/Mras/Nf1/Nfkb1/Nras/Pak1/Pdgfa/Pdgfb/Pdgfra/Pgf/Pik3ca/Pik3cd/Pik3r1/Pik3r2/Pik3r3/Prkaca/Prkacb/Prkcb/Prkcg/Pla2g1b/Pla2g4a/Plcg1/Pld1/Pld2/Ptpn11/Rab5b/Rab5c/Rac1/Rac2/Rasgrp2/Rasa3/Rasal1/Rasgrf1/Rel/Rela/Ralgds/Rgl1/Rgl2/Ralbp1/Rras/Shc1/Shc3/Sos1/Sos2/Pak6/Rap1b/Tek/Rapgef5/Tgfa/Rasa1/Tiam1/Vegfa/Vegfb/Vegfc/Pak2/Plaat3/Zap70/Rasal2/Plcg2/Akt3/Ets1/Ets2/Syngap1/Rasgrp3/Map2k1/Map2k2/Mapk1/Mapk10/Mapk3/Mapk8/Mapk9/Rab5a/Shc4/Rasal3/Ksr2/Pla2g6/Rasa4/Rassf5/Pdgfc/Rala/Rassf1/Tbk1/Stk4/Ralb/Gng11/Pla2g12a/Exoc2/Rras2/Pdgfd/Plce1/Pik3cb/Ntf5 |
| mmu04215 | Apoptosis - multiple species | 0,00122588 | Apaf1/Birc3/Birc2/Birc5/Bak1/Bax/Bcl2/Bcl2l1/Bid/Bcl2l11/Birc6/Casp3/Casp7/Casp8/Casp9/Cycs/Fadd/Bbc3/Septin4/Tnfrsf1a/Mapk10/Mapk8/Mapk9/Bok/Becn1/Pmaip1/Diablo |
| mmu00051 | Fructose and mannose metabolism | 0,00137569 | Mpi/Aldoa/Aldoc/Akr1b3/Akr1b7/Fbp2/Fbp1/Hk1/Hk2/Khk/Pfkfb3/Pfkfb2/Pfkl/Pfkm/Sord/Hkdc1/Gmds/Tpi1/Gfus/Tkfc/Aldob/Fcsk/Pfkfb4/Pmm1/Tigar/Pmm2/Pfkp/Akr1b10/Gmppa |
| mmu00620 | Pyruvate metabolism | 0,00148469 | Acacb/Me2/Acaca/Me3/Glo1/Acat1/Acat2/Aldh7a1/Aldh2/Aldh3a2/Dld/Fh1/Hagh/Ldha/Ldhb/Me1/Mdh2/Mdh1/Pcx/Pkm/Pklr/Dlat/Ldhd/Aldh9a1/Acyp1/Pdhb/Acss1/Aldh1b1/Pck2/Acyp2/Grhpr |
| mmu00450 | Selenocompound metabolism | 0,00151192 | Cth/Sephs1/Sephs2/Sepsecs/Mars2/Pstk/Mars1/Inmt/Kyat3/Txnrd3/Mtr/Papss1/Papss2/Txnrd1/Scly/Kyat1 |
| mmu05219 | Bladder cancer | 0,00154348 | Braf/Raf1/Ccnd1/Cdh1/Cdk4/Cdkn1a/Cdkn2a/Dapk2/Dapk3/E2f1/E2f3/Egf/Egfr/Erbb2/Fgfr3/Hbegf/Hras/Mdm2/Mmp9/Myc/Nras/Rb1/Src/Thbs1/Vegfa/E2f2/Map2k1/Map2k2/Mapk1/Mapk3/Rassf1/Dapk1/Rps6ka5 |
| mmu03040 | Spliceosome | 0,00211467 | Tra2a/Sf3b3/Snrpd2/Sf3b4/Srsf9/U2af1/Srsf1/Aqr/Dhx15/Ddx5/Srsf10/Hnrnpc/Hnrnpk/Hspa1l/Hspa1b/Hspa2/Magoh/Prpf8/Eif4a3/Slu7/Rbmxl1/Sf3a2/Sart1/Srsf2/Srsf3/Tra2b/Eftud2/Snrpc/Snrnp70/Snrpb/Snrpb2/Snrpd1/Snrpe/Snu13/Ddx46/Alyref/Dhx8/U2af2/Srsf7/Thoc1/Hnrnpa3/Pcbp1/Txnl4a/Lsm2/Cherp/Prpf19/Usp39/Sf3b2/Snrnp200/Lsm4/Hnrnpu/Plrg1/Rp9/Alyref2/Ppie/Tcerg1/Prpf40a/Acin1/Srsf4/Isy1/Sf3b6/Lsm7/Ppih/Sf3b5/Snw1/Lsm5/Magohb/Zmat2/Snrnp27/Ctnnbl1/Rbm22/Crnkl1/Prpf38b/Rbm25/Prpf18/Snrpd3/U2surp/Puf60/Srsf6/Snrpg/Bcas2/Syf2/Ppil1/Prpf6/Snrpa1/Prpf31/Dhx16/Snrpf/Prpf3/Cdc40/Ccdc12/Thoc3/Ddx23/Sf3a3/Smndc1/Lsm8/Hnrnpm/Rbm17/Lsm6/Sf3b1 |
| mmu00513 | Various types of N-glycan biosynthesis | 0,00224917 | Alg9/Mgat4b/Rpn1/Ddost/Man2a2/Hexa/Hexb/Stt3a/Man1a2/Man2a1/Mgat1/Rpn2/St3gal3/Alg1/Mgat2/Alg12/Man1b1/Man1c1/Hexdc/Mgat4a/Glt28d2/B4galnt3/B4galnt4/B4galt2/Fut8/Alg2/B4galt3/Alg14/Stt3b/Chst8/Chst9/Tusc3 |
| mmu04979 | Cholesterol metabolism | 0,00227271 | Ldlrap1/Pcsk9/Cyp27a1/Abca1/Apoa1/Apoe/Apoh/Tspo/Cd36/Lrp2/Lcat/Ldlr/Lipa/Lipg/Lpl/Lrp1/Lrpap1/Npc1/Pltp/Soat1/Sort1/Scarb1/Mylip/Vdac1/Vdac2/Vdac3/Soat2/Abcg5/Abcb11/Angptl3/Vapa/Nceh1/Vapb/Angptl4/Stard3/Angptl8/Npc2/Osbpl5 |
| mmu04392 | Hippo signaling pathway - multiple species | 0,00239216 | Ajuba/Lats1/Nf2/Pak1/Wwc1/Rassf4/Rassf2/Tead1/Tead2/Tead3/Tead4/Yap1/Csnk1e/Limd1/Frmd6/Fat4/Lats2/Stk3/Rassf1/Sav1/Mob1b/Rassf6/Wwtr1 |
| mmu04130 | SNARE interactions in vesicular transport | 0,00297397 | Bet1/Stx2/Sec22b/Snap23/Stx1a/Stx3/Stx4a/Vamp1/Vamp2/Vamp3/Vamp8/Bnip1/Stx16/Vamp4/Stx7/Gosr1/Vti1a/Vti1b/Vamp5/Stx8/Ykt6/Stx6/Use1/Stx17/Stx19/Stx18/Stx11 |
| mmu04930 | Type II diabetes mellitus | 0,00317181 | Gck/Cacna1a/Cacna1c/Cacna1d/Cacna1e/Cacna1g/Socs3/Socs1/Hk1/Hk2/Ikbkb/Insr/Irs1/Irs3/Pdx1/Pik3ca/Pik3cd/Pik3r1/Pik3r2/Pik3r3/Pkm/Prkcd/Prkce/Prkcz/Pklr/Slc2a2/Slc2a4/Hkdc1/Socs2/Mapk1/Mapk10/Mapk3/Mapk8/Mapk9/Irs2/Mtor/Pik3cb |
| mmu00564 | Glycerophospholipid metabolism | 0,00401587 | Gpat4/Dgkz/Dgkg/Dgkq/Cds2/Ache/Chka/Pcyt1a/Dgka/Lpin1/Gpd1/Gpd2/Gnpat/Gpam/Lpcat3/Lcat/Pemt/Lypla1/Pla2g1b/Pla2g4a/Pld1/Pld2/Pld3/Plpp1/Ptdss1/Pla2g15/Lpcat1/Chpt1/Etnk2/Dgkb/Mboat1/Lclat1/Plaat3/Lpgat1/Dgkd/Gpat3/Phospho1/Lypla2/Lpcat2/Ptdss2/Selenoi/Agpat3/Dgki/Pisd/Gpd1l/Dgkh/Pnpla6/Plpp2/Agpat5/Pla2g6/Dgke/Lpin2/Lpin3/Pla2g12a/Adprm/Crls1/Mboat2/Agpat2/Plpp3/Agpat4/Pcyt2/Etnppl/Plpp5/Gpcpd1/Pgs1/Cds1/Etnk1/Lpcat4 |
| mmu00062 | Fatty acid elongation | 0,00407445 | Tecr/Elovl3/Hadh/Elovl6/Acot2/Acot3/Ppt1/Acot5/Hadhb/Acot1/Mecr/Hacd1/Elovl1/Ppt2/Hsd17b12/Hacd3/Hacd4/Elovl5/Acot7/Hacd2/Elovl7/Them4/Elovl4/Hadha |
| mmu05142 | Chagas disease (American trypanosomiasis) | 0,00427237 | Ticam1/Akt1/Akt2/C1qa/Calr/Casp8/Cd247/Cflar/Chuk/Fadd/Fas/Fos/Gna11/Gna14/Gna15/Gnai1/Gnai2/Gnai3/Gnal/Gnao1/Gnaq/Gnas/Ifngr1/Ifngr2/Ikbkb/Jun/Smad2/Nfkb1/Nfkbia/Nos2/Pik3ca/Pik3cd/Pik3r1/Pik3r2/Pik3r3/Plcb1/Plcb2/Plcb3/Plcb4/Ppp2ca/Ppp2cb/Mapk11/Rela/Ccl5/Tgfb1/Tgfb2/Tgfb3/Tgfbr1/Tgfbr2/Tlr4/Tnfrsf1a/Traf6/Akt3/Tlr2/Map2k4/Mapk1/Mapk10/Mapk13/Mapk14/Mapk3/Mapk8/Mapk9/Irak4/Mapk12/Kng2/Ppp2r1a/Ppp2r2d/Ppp2r2a/Ppp2r2b/Ppp2r1b/Pik3cb |
| mmu04380 | Osteoclast differentiation | 0,00436454 | Acp5/Akt1/Akt2/Camk4/Chuk/Socs3/Socs1/Creb1/Csf1/Cyba/Fcgr1/Fcgr3/Fhl2/Fos/Fosb/Fosl1/Fosl2/Fyn/Gab2/Grb2/Ifnar1/Ifnar2/Ifngr1/Ifngr2/Ikbkb/Il1r1/Irf9/Jak1/Jun/Junb/Jund/Mitf/Nfatc1/Nfatc2/Nfkb1/Nfkb2/Nfkbia/Tnfrsf11b/Sqstm1/Pik3ca/Pik3cd/Pik3r1/Pik3r2/Pik3r3/Pparg/Ppp3ca/Ppp3cb/Ppp3cc/Ppp3r1/Mapk11/Sirpa/Rac1/Rela/Relb/Spi1/Stat1/Stat2/Syk/Tec/Tgfb1/Tgfb2/Tgfbr1/Tgfbr2/Tnfrsf11a/Tnfrsf1a/Traf2/Traf6/Plcg2/Akt3/Map2k1/Map2k6/Map2k7/Map3k7/Mapk1/Mapk10/Mapk13/Mapk14/Mapk3/Mapk8/Mapk9/Mapk12/Map3k14/Tyk2/Tab1/Tab2/Cyld/Pik3cb |
| mmu05164 | Influenza A | 0,0046586 | Dnajc3/Xpo1/Try5/Ticam1/Raf1/Fdps/Actb/Actg1/Akt1/Akt2/Slc25a4/Apaf1/Bak1/Bax/Bid/Casp3/Casp8/Casp9/Ccnd3/Cdk4/Cdk6/Chuk/Socs3/Crebbp/Cycs/Eif2s1/Fadd/Fas/Tlr3/H2-Ab1/H2-Eb1/H2-DMa/Icam1/Cxcl10/Ifnar1/Ifnar2/Ifngr1/Ifngr2/Ikbkb/Il18/Irf9/Jak1/Jak2/Kpna1/Kpna2/Kpna6/Mx2/Nfkb1/Nfkbia/Nfkbib/Pik3ca/Pik3cd/Pik3r1/Pik3r2/Pik3r3/Prkcb/Pml/Eif2ak2/Rab11b/Rela/Ccl5/Stat1/Stat2/Tmprss4/Trim25/Tlr4/Tnfrsf10b/Tnfrsf1a/Traf3/Tnfsf10/Prss2/Try4/Vdac1/Mavs/Ddx58/Ifna13/Hnrnpul1/Akt3/Rnasel/Ifna15/Map2k1/Map2k2/Mapk1/Mapk3/Irak4/Nlrx1/Ep300/Kpna7/Try10/Tmprss2/Nxf1/Rab11a/Irf7/Irf3/Cpsf4/Pabpn1/Tyk2/Adar/Tbk1/Nxt1/Ikbke/Rsad2/Rae1/Pycard/2210010C04Rik/Ifih1/Tradd/1810009J06Rik/Pik3cb/Calcoco2 |
| mmu03430 | Mismatch repair | 0,00468568 | Rfc4/Lig1/Mlh1/Msh2/Msh6/Pcna/Pms2/Pold1/Pold2/Rfc1/Rfc2/Rpa2/Mlh3/Exo1/Ssbp1/Pold3/Rfc3/Pold4/Rfc5 |
| mmu04625 | C-type lectin receptor signaling pathway | 0,00469762 | Raf1/Akt1/Akt2/Bcl10/Bcl3/Calm1/Calm2/Calm3/Casp8/Chuk/Plk3/Egr2/Egr3/Fcer1g/Hras/Ikbkb/Irf1/Irf9/Itpr1/Itpr2/Itpr3/Jun/Ksr1/Mapkapk2/Mdm2/Mras/Nfatc1/Nfatc2/Nfatc3/Nfkb1/Nfkb2/Nfkbia/Nras/Pak1/Pik3ca/Pik3cd/Pik3r1/Pik3r2/Pik3r3/Prkcd/Ppp3ca/Ppp3cb/Ppp3cc/Ppp3r1/Mapk11/Ptgs2/Ptpn11/Rela/Relb/Rras/Src/Stat1/Stat2/Cblb/Syk/Plcg2/Akt3/Il17d/Malt1/Mapk1/Mapk10/Mapk13/Mapk14/Mapk3/Mapk8/Mapk9/Mapk12/Card9/Map3k14/Ikbke/Clec7a/Pycard/Rras2/Arhgef12/Nfatc4/Cyld/Pik3cb |
| mmu04929 | GnRH secretion | 0,00582453 | Arrb1/Raf1/Akt1/Akt2/Cacna1c/Cacna1d/Cacna1g/Cga/Esr2/Kcnn2/Kcnn3/Gna11/Gnaq/Hcn1/Hcn3/Hras/Itpr1/Itpr2/Itpr3/Kcnj9/Nras/Pik3ca/Pik3cd/Pik3r1/Pik3r2/Pik3r3/Prkcb/Prkcg/Plcb1/Plcb2/Plcb3/Plcb4/Spp1/Arrb2/Trpc1/Akt3/Map2k1/Map2k2/Mapk1/Mapk3/Kiss1/Gabbr1/Cacna1h/Pik3cb/Gper1/Kcnn1 |
| mmu00511 | Other glycan degradation | 0,00582453 | Manba/Aga/Gba/Hexa/Hexb/Man2b1/Man2b2/Neu1/Engase/Gba2/Hexdc/Neu2/Neu3/Fuca2/Fuca1/Man2c1 |
| mmu03410 | Base excision repair | 0,00632446 | Parp1/Parp2/Apex1/Fen1/Hmgb1/Lig1/Mbd4/Nthl1/Ogg1/Pcna/Polb/Pold1/Pold2/Pole/Pole2/Tdg/Ung/Xrcc1/Neil3/Parp3/Mpg/Neil2/Pole4/Pold3/Pold4/Smug1/Neil1 |
| mmu04916 | Melanogenesis | 0,00662785 | Adcy3/Camk2d/Raf1/Adcy6/Adcy9/Calm1/Calm2/Calm3/Ctnnb1/Creb1/Creb3/Crebbp/Dvl1/Dvl2/Dvl3/Edn1/Fzd1/Fzd3/Fzd4/Fzd5/Fzd6/Fzd7/Fzd8/Fzd9/Gnai1/Gnai2/Gnai3/Gnao1/Gnaq/Gnas/Hras/Kit/Lef1/Kitl/Mitf/Nras/Prkaca/Prkacb/Prkcb/Prkcg/Plcb1/Plcb2/Plcb3/Plcb4/Pomc/Creb3l2/Wnt8a/Adcy2/Tcf7/Tcf7l1/Tcf7l2/Wnt9a/Wnt1/Wnt10b/Wnt11/Wnt9b/Wnt2b/Wnt5a/Wnt5b/Wnt6/Map2k1/Map2k2/Mapk1/Mapk3/Creb3l1/Ep300/Gsk3b/Fzd2/Creb3l4 |
| mmu00280 | Valine, leucine and isoleucine degradation | 0,00880081 | Il4i1b/Aldh6a1/Acat1/Acat2/Aldh7a1/Pcca/Acadm/Acaa1a/Acads/Aldh2/Aldh3a2/Aox1/Auh/Bcat1/Bcat2/Bckdha/Bckdhb/Dbt/Dld/Hadh/Hmgcl/Hmgcs2/Mmut/Hmgcs1/Hmgcll1/Hibch/Hadhb/Acaa1b/Acsf3/Abat/Ivd/Aldh9a1/Hibadh/Acadsb/Pccb/Acad8/Aldh1b1/Ehhadh/Mccc2/Aacs/Hadha |
| mmu04923 | Regulation of lipolysis in adipocytes | 0,00880081 | Adcy3/Npy/Adcy6/Adcy9/Adora1/Adrb1/Adrb2/Akt1/Akt2/Cga/Gnai1/Gnai2/Gnai3/Gnas/Insr/Irs1/Irs3/Lipe/Npr1/Npy1r/Pde3b/Pik3ca/Pik3cd/Pik3r1/Pik3r2/Pik3r3/Prkaca/Prkacb/Prkg1/Ptger3/Ptgs1/Ptgs2/Adcy2/Tshr/Plaat3/Akt3/Mgll/Irs2/Pnpla2/Abhd5/Pik3cb |
| mmu04933 | AGE-RAGE signaling pathway in diabetic complications | 0,00955206 | Akt1/Akt2/Bax/Bcl2/Casp3/Ccnd1/Cdc42/Cdk4/Cdkn1b/Col1a1/Diaph1/Edn1/Egr1/F3/Hras/Icam1/Jak2/Jun/Smad2/Smad3/Smad4/Nfatc1/Nfkb1/Nras/Pik3ca/Pik3cd/Pik3r1/Pik3r2/Pik3r3/Pim1/Prkcb/Prkcd/Prkce/Prkcz/Plcb1/Plcb2/Plcb3/Plcb4/Plcd1/Plcg1/Mapk11/Rac1/Rela/Stat1/Stat3/Stat5a/Stat5b/Tgfb1/Tgfb2/Tgfb3/Tgfbr1/Tgfbr2/Thbd/Vegfa/Vegfb/Vegfc/Plcg2/Akt3/Mapk1/Mapk10/Mapk13/Mapk14/Mapk3/Mapk8/Mapk9/Mapk12/Foxo1/Plce1/Pik3cb |
| mmu00510 | N-Glycan biosynthesis | 0,00992245 | Alg9/Mgat4b/Rpn1/Mgat5/Ddost/Dpagt1/Dpm2/Man2a2/Ganab/Stt3a/Man1a2/Man2a1/Mgat1/Rpn2/St6gal1/Alg1/Mgat2/Alg12/Man1b1/Dolk/Man1c1/Mgat4a/Glt28d2/Alg6/Alg8/B4galt2/Fut8/Alg2/Dolpp1/Srd5a3/B4galt3/Mogs/Alg5/Alg14/Stt3b/Dpm3/Tusc3 |
| mmu04926 | Relaxin signaling pathway | 0,00992245 | Adcy3/Arrb1/Raf1/Adcy6/Adcy9/Akt1/Akt2/Atf2/Atf4/Col1a1/Creb1/Creb3/Atf6b/Edn1/Egfr/Fos/Gna15/Gnai1/Gnai2/Gnai3/Gnao1/Gnas/Gnb1/Gng10/Gng12/Gng2/Gng5/Gng7/Gng8/Grb2/Hras/Jun/Smad2/Mmp9/Nfkb1/Nfkbia/Nos2/Nras/Pik3ca/Pik3cd/Pik3r1/Pik3r2/Pik3r3/Prkaca/Prkacb/Prkcz/Plcb1/Plcb2/Plcb3/Plcb4/Mapk11/Rela/Rln1/Shc1/Shc3/Sos1/Sos2/Src/Creb3l2/Adcy2/Arrb2/Tgfb1/Tgfbr1/Tgfbr2/Vegfa/Vegfb/Vegfc/Creb5/Akt3/Map2k1/Map2k2/Map2k4/Map2k7/Mapk1/Mapk10/Mapk13/Mapk14/Mapk3/Mapk8/Mapk9/Creb3l1/Shc4/Mapk12/Gng11/Pik3cb/Creb3l4 |
| mmu00240 | Pyrimidine metabolism | 0,01073276 | Nt5m/Nt5c3/Dut/Entpd1/Entpd6/Entpd5/Dck/Nme7/Nme1/Nme2/Enpp1/Pnp/Rrm1/Rrm2/Enpp3/Entpd3/Tk1/Dtymk/Cmpk2/Tyms/Uck1/Umps/Upp1/Nt5c1a/Nt5e/Dctd/Rrm2b/Nt5c/Ctps/Nme6/Nme4/Dhodh/Tk2/Dctpp1/Cmpk1/Pnp2/Nt5c3b/Cda/Cant1/Nt5c2/Uck2/Dpyd |
| mmu00100 | Steroid biosynthesis | 0,01076163 | Cel/Cyp27b1/Cyp51/Dhcr7/Fdft1/Hsd17b7/Lipa/Lss/Soat1/Sqle/Soat2/Sc5d/Cyp2r1/Msmo1/Tm7sf2/Dhcr24/Lbr |
| mmu00020 | Citrate cycle (TCA cycle) | 0,01257183 | Acly/Aco1/Aco2/Cs/Dld/Fh1/Idh1/Idh3b/Mdh2/Mdh1/Ogdh/Pcx/Sucla2/Suclg2/Dlat/Ogdhl/Idh2/Suclg1/Sdhd/Sdha/Sdhb/Idh3a/Pdhb/Pck2/Dlst |
| mmu04136 | Autophagy - other | 0,01257183 | Atg5/Ppp2ca/Ppp2cb/Atg9b/Pik3c3/Atg4d/Atg4c/Atg9a/Ulk2/Atg2a/Wipi1/Becn1/Gabarap/Mlst8/Mtor/Atg4b/Atg10/Atg3/Atg101/Atg7/Rptor/Wipi2/Pik3r4/Atg16l1/Gabarapl2 |
| mmu04659 | Th17 cell differentiation | 0,01313588 | Ahr/Runx1/Cd247/Chuk/Fos/Gata3/H2-Ab1/H2-Eb1/H2-DMa/Hif1a/Hsp90ab1/Hsp90aa1/Ifngr1/Ifngr2/Ikbkb/Il12rb1/Il1r1/Il1rap/Il4ra/Il6ra/Il6st/Irf4/Jak1/Jak2/Jak3/Jun/Smad2/Smad3/Smad4/Nfatc1/Nfatc2/Nfatc3/Nfkb1/Nfkbia/Nfkbib/Nfkbie/Plcg1/Ppp3ca/Ppp3cb/Ppp3cc/Ppp3r1/Mapk11/Rara/Rela/Rora/Rorc/Rxra/Rxrg/Stat1/Stat3/Stat5a/Stat5b/Stat6/Tgfb1/Tgfbr1/Tgfbr2/Zap70/Il17d/Mapk1/Mapk10/Mapk13/Mapk14/Mapk3/Mapk8/Mapk9/Mapk12/Tyk2/Mtor/Tbx21 |
| mmu04728 | Dopaminergic synapse | 0,01600587 | Camk2d/Slc18a1/Akt1/Akt2/Arntl/Atf2/Atf4/Cacna1a/Cacna1c/Cacna1d/Calm1/Calm2/Calm3/Clock/Comt/Creb1/Creb3/Atf6b/Slc6a3/Ddc/Fos/Gnai1/Gnai2/Gnai3/Gnal/Gnao1/Gnaq/Gnas/Gnb1/Gng10/Gng12/Gng2/Gng5/Gng7/Gng8/Itpr1/Itpr2/Itpr3/Kcnj9/Kif5a/Kif5b/Kif5c/Prkaca/Prkacb/Prkcb/Prkcg/Plcb1/Plcb2/Plcb3/Plcb4/Ppp1cb/Ppp1cc/Ppp1r1b/Ppp2ca/Ppp2cb/Ppp3ca/Ppp3cb/Ppp3cc/Mapk11/Scn1a/Creb3l2/Slc18a2/Arrb2/Ppp2r5d/Ppp2r5b/Ppp2r5a/Creb5/Ppp2r3a/Akt3/Mapk10/Mapk13/Mapk14/Mapk8/Mapk9/Creb3l1/Ppp2r5c/Ppp2r5e/Mapk12/Ppp2r1a/Ppp2r2d/Gsk3b/Ppp2r3c/Gsk3a/Gng11/Ppp2r2a/Ppp2r2b/Ppp2r1b/Creb3l4 |
| mmu04915 | Estrogen signaling pathway | 0,01600587 | Adcy3/Raf1/Adcy6/Adcy9/Akt1/Akt2/Atf2/Atf4/Bcl2/Calm1/Calm2/Calm3/Creb1/Creb3/Atf6b/Ctsd/Egfr/Esr1/Esr2/Fkbp4/Fkbp5/Fos/Gnai1/Gnai2/Gnai3/Gnao1/Gnaq/Gnas/Grb2/Hbegf/Hras/Hspa1l/Hspa1b/Hspa2/Hsp90ab1/Hsp90aa1/Itpr1/Itpr2/Itpr3/Jun/Kcnj9/Krt10/Krt18/Krt19/Mmp9/Ncoa1/Ncoa2/Ncoa3/Nras/Pik3ca/Pik3cd/Pik3r1/Pik3r2/Pik3r3/Prkaca/Prkacb/Prkcd/Plcb1/Plcb2/Plcb3/Plcb4/Pomc/Rara/Shc1/Shc3/Sos1/Sos2/Sp1/Src/Creb3l2/Adcy2/Tgfa/Creb5/Krt39/Akt3/Map2k1/Map2k2/Mapk1/Mapk3/Creb3l1/Shc4/Gabbr1/Ebag9/Krt25/Pik3cb/Gper1/Creb3l4/Krt23 |
| mmu05132 | Salmonella infection | 0,01647183 | Dync2h1/Actb/Actg1/Arpc1b/Cd14/Cdc42/Dync1h1/Dync1i1/Dync1i2/Fos/Cxcl1/Ifngr1/Ifngr2/Il18/Jun/Klc1/Klc2/Nfkb1/Nos2/Pfn1/Pfn2/Mapk11/Rab7/Rac1/Rela/Rock1/Rock2/Tjp1/Tlr4/Klc3/Dync1li2/Dync1li1/Wasf2/Mapk1/Mapk10/Mapk13/Mapk14/Mapk3/Mapk8/Mapk9/Rilp/Flnb/Mapk12/Pkn1/Tlr5/Rhog/Arpc3/Pycard/Flnc/Plekhm2/Wasl/Arpc2 |
| mmu00600 | Sphingolipid metabolism | 0,01728748 | Asah1/Degs1/Galc/Gba/Neu1/Plpp1/Sgpl1/Smpd1/Smpd2/Sphk1/Sptlc2/Sgms1/Ugcg/Gba2/Acer2/Neu2/Cers6/Sptlc1/Sgpp2/Plpp2/Neu3/Gal3st1/Asah2/B4galt6/Smpd3/Acer3/Cers4/Plpp3/Degs2/Kdsr/Cers5/Sgms2/Cers2/Smpd4/Sgpp1 |
| mmu04912 | GnRH signaling pathway | 0,01877054 | Adcy3/Camk2d/Raf1/Adcy6/Adcy9/Atf4/Cacna1c/Cacna1d/Calm1/Calm2/Calm3/Cdc42/Cga/Egfr/Egr1/Gna11/Gnaq/Gnas/Grb2/Hbegf/Hras/Itpr1/Itpr2/Itpr3/Jun/Mmp14/Nras/Prkaca/Prkacb/Prkcb/Prkcd/Pla2g4a/Plcb1/Plcb2/Plcb3/Plcb4/Pld1/Pld2/Mapk11/Ptk2b/Sos1/Sos2/Src/Adcy2/Mapk7/Map2k1/Map2k2/Map2k4/Map2k6/Map2k7/Map3k1/Map3k2/Map3k3/Mapk1/Mapk10/Mapk13/Mapk14/Mapk3/Mapk8/Mapk9/Mapk12 |
| mmu05014 | Amyotrophic lateral sclerosis (ALS) | 0,02168951 | Apaf1/Bad/Bax/Bcl2/Bcl2l1/Bid/Cat/Casp3/Casp9/Ccs/Cycs/Daxx/Gpx1/Gpx3/Grin1/Grin2d/Nefm/Ppp3ca/Ppp3cb/Ppp3cc/Ppp3r1/Mapk11/Rac1/Slc1a2/Sod1/Tnfrsf1a/Tnfrsf1b/Map2k6/Map3k5/Mapk13/Mapk14/Rab5a/Mapk12/Nefh/Tomm40/Tomm40l/Gpx7/Derl1/Gpx8/Als2/Gpx6 |
| mmu05012 | Parkinson disease | 0,02260019 | Ppif/Cox6b1/Slc18a1/Adora2a/Slc25a4/Apaf1/Atp5a1/Atp5b/Atp5c1/Atp5pb/Atp5g1/Atp5j/Casp3/Casp9/Cox5a/Cox5b/Cox6a1/Cox6c/Cox7a1/Cox7a2/Cox7c/Cox8a/Cycs/Slc6a3/Ube2j2/Gnai1/Gnai2/Gnai3/Gnal/Ndufa2/Ndufa4/Ndufs4/Ndufv1/Prkaca/Prkacb/Septin5/Cox7a2l/Snca/Slc18a2/Ubb/Ube2l3/Ube2g2/Uqcrc1/Vdac1/Vdac2/Vdac3/Ndufs8/Ndufs2/Ndufs1/Atp5g3/Ndufb6/Atp5o/Ndufs6/Ndufa4l2/Prkn/Ube2l6/Park7/Ndufs5/Atp5d/Ndufb5/Ndufa3/Ndufa9/Uqcr10/Ndufb9/Ndufc1/Ndufa12/Cyc1/Uqcr11/Uqcrfs1/Lrrk2/Ndufb7/Sdhd/Sdha/Uqcrc2/Atp5e/Ube2g1/Ndufa6/Ndufb8/Ndufa10/Sdhb/Sncaip/Atp5g2/Ndufb4/Ndufc2/Ndufb2/Ndufa5/Ndufa8/Pink1/Ndufab1/Ndufv2/Uba7/Ndufs7 |
| mmu00500 | Starch and sucrose metabolism | 0,02343411 | Amy2a4/Amy2a3/Amy2a2/Gck/Amy2a5/Pygb/Pygl/Amy1/G6pc/Gaa/Gpi1/Gys1/Hk1/Hk2/Enpp1/Pygm/Enpp3/Hkdc1/Ugp2/Gyg/Pgm2/G6pc3/Pgm2l1/Gbe1/Agl |
| mmu04024 | cAMP signaling pathway | 0,02415217 | Adcy3/Camk2d/Orai1/Npy/Braf/Rap1a/Raf1/Pde4c/Acox1/Adcy6/Adcy9/Adora1/Adora2a/Adrb1/Adrb2/Akt1/Akt2/Amh/Atp1a1/Atp1b1/Atp1b2/Atp1b3/Atp2a2/Bad/Cacna1c/Cacna1d/Calm1/Calm2/Calm3/Camk4/Cftr/Cga/Chrm1/Creb1/Creb3/Crebbp/Edn1/Edn2/F2r/Fos/Gli1/Glp1r/Gnai1/Gnai2/Gnai3/Gnas/Grin1/Grin2d/Hhip/Htr1b/Htr1d/Htr6/Jun/Lipe/Rac3/Afdn/Ppp1r12a/Nfatc1/Nfkb1/Nfkbia/Npr1/Npy1r/Oxt/Pak1/Pde3b/Pde4a/Pde4b/Pik3ca/Pik3cd/Pik3r1/Pik3r2/Pik3r3/Prkaca/Prkacb/Pld1/Pld2/Pomc/Ppara/Ppp1cb/Ppp1cc/Ppp1r1b/Ptch1/Ptger2/Ptger3/Rac1/Rac2/Rela/Rock1/Rock2/Rras/Slc9a1/Sst/Sstr1/Sstr2/Sox9/Creb3l2/Adcy2/Rap1b/Tiam1/Tshr/Vav1/Vav2/Vipr2/Rapgef3/Creb5/Ffar2/Akt3/Pde4d/Abcc4/Pde10a/Chrm2/Map2k1/Map2k2/Mapk1/Mapk10/Mapk3/Mapk8/Mapk9/Creb3l1/Ep300/Hcn4/Atp2b4/Gipr/Gabbr1/Fxyd1/Rapgef4/Vav3/Rras2/Atp2b1/Plce1/Pik3cb/Creb3l4/Hcar2/Acox3/Sucnr1 |
| mmu04950 | Maturity onset diabetes of the young | 0,02506347 | Gck/Hes1/Hhex/Mnx1/Foxa2/Foxa3/Hnf4a/Onecut1/Bhlha15/Neurod1/Nkx2-2/Nkx6-1/Pax6/Pdx1/Pklr/Slc2a2/Hnf1a/Hnf1b/Nr5a2/Hnf4g/Rfx6 |
| mmu00270 | Cysteine and methionine metabolism | 0,02514785 | Amd2/Il4i1b/Adi1/Psat1/Cth/Mat2b/Mat1a/Bcat1/Bcat2/Cbs/Cdo1/Dnmt1/Dnmt3a/Dnmt3b/Gclc/Gclm/Got1/Got2/Gss/Ldha/Ldhb/Mdh2/Mdh1/Srm/Tst/Ahcyl1/Kyat3/Mat2a/Phgdh/Mtr/Sdsl/Ahcy/Apip/Mtap/Mri1/Kyat1/Ahcyl2 |
| mmu04216 | Ferroptosis | 0,0260124 | Atg5/Cp/Acsl1/Fth1/Ftl1/Gclc/Gclm/Lpcat3/Gss/Hmox1/Slc3a2/Slc11a2/Pcbp2/Slc39a14/Acsl6/Tfrc/Vdac2/Vdac3/Pcbp1/Slc7a11/Ncoa4/Acsl5/Slc40a1/Gpx4/Map1lc3a/Map1lc3b/Slc39a8/Steap3/Sat2/Atg7 |
| mmu04973 | Carbohydrate digestion and absorption | 0,02961914 | Amy2a4/Amy2a3/Amy2a2/Amy2a5/Akt1/Akt2/Amy1/Atp1a1/Atp1b1/Atp1b2/Atp1b3/Cacna1d/G6pc/Slc37a4/Hk1/Hk2/Pik3ca/Pik3cd/Pik3r1/Pik3r2/Pik3r3/Prkcb/Plcb1/Plcb2/Plcb3/Plcb4/Slc2a2/Slc5a1/Hkdc1/Akt3/Slc2a5/G6pc3/Pik3cb |
| mmu04611 | Platelet activation | 0,03195803 | Adcy3/Mylk/Orai1/Rap1a/Actb/Actg1/Adcy6/Adcy9/Akt1/Akt2/Col1a1/F2/F2r/Fcer1g/Fyn/Gnai1/Gnai2/Gnai3/Gnaq/Gnas/Gp1bb/Gp5/Itga2/Itga2b/Itgb1/Itpr1/Itpr2/Itpr3/Arhgef1/Lyn/Ppp1r12a/P2rx1/P2ry1/Pik3ca/Pik3cd/Pik3r1/Pik3r2/Pik3r3/Prkaca/Prkacb/Prkci/Prkcz/Pla2g4a/Plcb1/Plcb2/Plcb3/Plcb4/Ppp1cb/Ppp1cc/Prkg1/Mapk11/Ptgs1/Rasgrp2/Rock1/Rock2/Snap23/Src/Stim1/Syk/Adcy2/Tbxas1/Rap1b/Vamp8/Vasp/Plcg2/Gucy1a2/Akt3/Gp6/Mapk1/Mapk13/Mapk14/Mapk3/Mapk12/Pik3cg/Pik3r5/Gucy1b1/Gucy1a1/Myl12b/Arhgef12/Tln2/Pik3cb |
| mmu04662 | B cell receptor signaling pathway | 0,03372 | Raf1/Akt1/Akt2/Bcl10/Cd22/Cd81/Chuk/Cr2/Fos/Grb2/Hras/Ikbkb/Inppl1/Jun/Rac3/Lyn/Nfatc1/Nfatc2/Nfatc3/Nfkb1/Nfkbia/Nfkbib/Nfkbie/Nras/Pik3ca/Pik3cd/Pik3r1/Pik3r2/Pik3r3/Prkcb/Ppp3ca/Ppp3cb/Ppp3cc/Ppp3r1/Rac1/Rac2/Rela/Sos1/Sos2/Syk/Vav1/Vav2/Plcg2/Akt3/Rasgrp3/Malt1/Dapp1/Map2k1/Map2k2/Mapk1/Mapk3/Gsk3b/Vav3/Pik3cb/Pik3ap1 |
| mmu04371 | Apelin signaling pathway | 0,03372 | Adcy3/Prkaa1/Mef2bl/Mylk/Prkaa2/Prkab2/Prkag2/Raf1/Slc8a3/Adcy6/Adcy9/Akt1/Akt2/Calm1/Calm2/Calm3/Camk4/Ccnd1/Cdh1/Egr1/Gnai1/Gnai2/Gnai3/Gnaq/Gnb1/Gng10/Gng12/Gng2/Gng5/Gng7/Gng8/Hdac5/Hras/Itpr1/Itpr2/Itpr3/Jag1/Klf2/Lipe/Smad2/Smad3/Smad4/Mef2a/Mef2c/Mef2d/Mras/Myl4/Nos2/Notch3/Nras/Nrf1/Pde3b/Prkaca/Prkacb/Prkce/Plcb1/Plcb2/Plcb3/Plcb4/Ppargc1a/Prkab1/Prkag1/Rps6/Rras/Slc8a1/Slc9a1/Sphk1/Spp1/Hdac4/Adcy2/Tfam/Tgfbr1/Pik3c3/Akt3/Map2k1/Map2k2/Mapk1/Mapk3/Pik3cg/Pik3r5/Becn1/Gabarap/Mtor/Rps6kb2/Gng11/Rras2/Pik3r4/Gabarapl2 |
| mmu04658 | Th1 and Th2 cell differentiation | 0,03403331 | Maml1/Cd247/Chuk/Dll1/Dll3/Fos/Gata3/H2-Ab1/H2-Eb1/H2-DMa/Ifngr1/Ifngr2/Ikbkb/Il12rb1/Il4ra/Jag1/Jak1/Jak2/Jak3/Jun/Maf/Nfatc1/Nfatc2/Nfatc3/Nfkb1/Nfkbia/Nfkbib/Nfkbie/Notch1/Notch2/Notch3/Plcg1/Ppp3ca/Ppp3cb/Ppp3cc/Ppp3r1/Mapk11/Rbpj/Rela/Stat1/Stat4/Stat5a/Stat5b/Stat6/Zap70/Mapk1/Mapk10/Mapk13/Mapk14/Mapk3/Mapk8/Mapk9/Maml2/Mapk12/Maml3/Dll4/Tyk2/Tbx21 |
| mmu04620 | Toll-like receptor signaling pathway | 0,03836158 | Ticam1/Akt1/Akt2/Tirap/Casp8/Cd14/Chuk/Fadd/Fos/Tlr3/Cxcl10/Ifnar1/Ifnar2/Ikbkb/Jun/Ly96/Nfkb1/Nfkbia/Pik3ca/Pik3cd/Pik3r1/Pik3r2/Pik3r3/Mapk11/Rac1/Rela/Ripk1/Ccl5/Spp1/Stat1/Tlr4/Cd40/Traf3/Traf6/Ticam2/Ifna13/Akt3/Tlr2/Ifna15/Map2k1/Map2k2/Map2k4/Map2k6/Map2k7/Map3k7/Map3k8/Mapk1/Mapk10/Mapk13/Mapk14/Mapk3/Mapk8/Mapk9/Irak4/Irf5/Mapk12/Tlr5/Irf7/Irf3/Tollip/Tbk1/Ikbke/Tab1/Tab2/Pik3cb |
| mmu04921 | Oxytocin signaling pathway | 0,04036449 | Adcy3/Prkaa1/Mylk/Camk2d/Prkaa2/Prkab2/Prkag2/Raf1/Actb/Actg1/Adcy6/Adcy9/Cacna1c/Cacna1d/Cacna2d1/Cacnb1/Cacnb2/Cacnb3/Cacnb4/Calm1/Calm2/Calm3/Camk4/Ccnd1/Cd38/Cdkn1a/Eef2/Eef2k/Egfr/Fos/Gnai1/Gnai2/Gnai3/Gnao1/Gnaq/Gnas/Hras/Itpr1/Itpr2/Itpr3/Jun/Kcnj2/Kcnj4/Kcnj9/Mef2c/Myl6/Ppp1r12a/Nfatc1/Nfatc2/Nfatc3/Npr1/Nras/Oxt/Prkaca/Prkacb/Prkcb/Prkcg/Pla2g4a/Plcb1/Plcb2/Plcb3/Plcb4/Ppp1cb/Ppp1cc/Ppp3ca/Ppp3cb/Ppp3cc/Ppp3r1/Prkab1/Prkag1/Ptgs2/Rgs2/Rock1/Rock2/Camkk2/Src/Adcy2/Myl6b/Npr2/Ppp1r12c/Gucy1a2/Map2k5/Mapk7/Map2k1/Map2k2/Mapk1/Mapk3/Pik3cg/Pik3r5/Ppp1r12b/Camk1/Gucy1b1/Rcan1/Cacna2d2/Gucy1a1/Nfatc4/Cacng7 |
| mmu00330 | Arginine and proline metabolism | 0,04126691 | Amd2/Carns1/Aldh7a1/Aldh2/Aldh3a2/Arg1/Arg2/Ckb/Ckm/Ckmt1/Gamt/Got1/Got2/Nos2/Oat/Odc1/P4ha1/P4ha2/Prodh/Srm/Pycr1/Aldh4a1/Smox/Azin2/Aldh18a1/Aldh9a1/Cndp2/Pycrl/Lap3/Gatm/L3hypdh/Hoga1/Pycr2/Sat2/Aldh1b1 |
| mmu00630 | Glyoxylate and dicarboxylate metabolism | 0,04236444 | Gldc/Shmt2/Acat1/Acat2/Pcca/Aco1/Aco2/Cat/Cs/Dld/Mdh2/Mdh1/Mmut/Shmt1/Glyctk/Amt/Hao2/Pccb/Hoga1/Gcsh/Acss1/Afmid/Grhpr |
| mmu04622 | RIG-I-like receptor signaling pathway | 0,04275873 | Atg5/Casp8/Chuk/Fadd/Cxcl10/Ikbkb/Nfkb1/Nfkbia/Nfkbib/Mapk11/Rela/Ripk1/Tank/Trim25/Traf2/Traf3/Traf6/Tkfc/Mavs/Ddx58/Ifna13/Ifne/Pin1/Ifna15/Map3k1/Map3k7/Mapk10/Mapk13/Mapk14/Mapk8/Mapk9/Nlrx1/Azi2/Mapk12/Irf7/Irf3/Tbk1/Ikbke/Sike1/Rnf125/Ifih1/Tradd/Sting1/Tbkbp1/Cyld/Dhx58 |
| mmu04918 | Thyroid hormone synthesis | 0,04362624 | Adcy3/Slc5a5/Adcy6/Adcy9/Atf2/Atf4/Atp1a1/Atp1b1/Atp1b2/Atp1b3/Pdia4/Canx/Cga/Creb1/Creb3/Atf6b/Gnaq/Gnas/Lrp2/Gpx1/Gpx3/Gsr/Hspa5/Itpr1/Itpr2/Itpr3/Prkaca/Prkacb/Prkcb/Prkcg/Plcb1/Plcb2/Plcb3/Plcb4/Creb3l2/Adcy2/Duox2/Tshr/Ttf1/Creb5/Slc26a4/Creb3l1/Duoxa2/Gpx7/Gpx8/Iyd/Ttf2/Gpx6/Creb3l4 |
| mmu04660 | T cell receptor signaling pathway | 0,04500185 | Raf1/Akt1/Akt2/Bcl10/Cd247/Cdc42/Cdk4/Chuk/Dlg1/Fos/Fyn/Grb2/Hras/Ikbkb/Jun/Nck1/Nck2/Nfatc1/Nfatc2/Nfatc3/Nfkb1/Nfkbia/Nfkbib/Nfkbie/Nras/Pak1/Pdpk1/Pik3ca/Pik3cd/Pik3r1/Pik3r2/Pik3r3/Plcg1/Ppp3ca/Ppp3cb/Ppp3cc/Ppp3r1/Mapk11/Rela/Sos1/Sos2/Cblb/Pak6/Tec/Vav1/Vav2/Pak2/Zap70/Akt3/Malt1/Map2k1/Map2k2/Map2k7/Map3k7/Map3k8/Mapk1/Mapk10/Mapk13/Mapk14/Mapk3/Mapk8/Mapk9/Mapk12/Map3k14/Gsk3b/Vav3/Pik3cb |
| mmu03022 | Basal transcription factors | 0,04863225 | Taf5l/Cdk7/Ercc2/Ercc3/Gtf2h1/Gtf2h4/Gtf2i/Mnat1/Taf3/Taf6/Taf6l/Taf5/Taf4/Gtf2b/Gtf2a2/Tbpl1/Gtf2h2/Taf7/Taf10/Taf2/Gtf2ird1/Taf8/Taf12/Ccnh/Gtf2e2/Gtf2f2/Taf15/Taf4b/Gtf2a1/Gtf2f1/Taf13 |
| mmu05168 | Herpes simplex virus 1 infection | 8,6732E-17 | Gm2026/Gm3055/Zfp984/Gm14308/Zfp850/Gm10778/Zfp729b/Zfp282/Zfp956/E430018J23Rik/AW146154/Zfp119a/Ticam1/Srsf9/Eif2b3/H2-Q6/H2-Q9/Srsf1/Zfp607b/Akt1/Akt2/Apaf1/Birc3/Birc2/B2m/Bad/Bak1/Bax/Bcl2/Bcl2l1/Bid/Calr/Casp3/Casp8/Casp9/Chuk/Socs3/Cycs/Daxx/Eif2s1/Eif2ak3/Eif2b4/Eif4ebp1/Fadd/Fas/Tlr3/Pdia3/H2-Ab1/H2-Bl/H2-D1/H2-Eb1/H2-K1/H2-M3/H2-DMa/H2-Q1/H2-Q2/H2-Q4/H2-T22/H2-T23/H2-T24/Eif2ak1/Ifnar1/Ifnar2/Ifngr1/Ifngr2/Ikbkb/Il12b/Irf9/Itga5/Jak1/Jak2/Zfp87/Zfp617/Nfkb1/Nfkbia/Pik3ca/Pik3cd/Pik3r1/Pik3r2/Pik3r3/Pml/Pou2f1/Pou2f2/Pou2f3/Ppp1ca/Ppp1cb/Ppp1cc/Eif2ak2/Ptpn11/Zfp286/Zfp184/Rela/Rheb/Ccl5/Srsf2/Srsf3/Src/Srpk1/Zfp871/Stat1/Stat2/Eif2b1/Syk/Zfp658/Zfp719/Zfp180/Zfp677/Zfp947/Zfp748/Zfp273/Zfp583/Tap2/Tapbp/Zfp354a/Cgas/Zfp879/AU041133/Alyref/Eif2b2/Zfp455/Zfp595/Tnfrsf1a/Traf2/Traf3/Traf5/Traf6/Zfp7/Eif2b5/Zfp160/Zfp472/Zfp81/Zfp959/Srsf7/Zfp1/Zfp101/Zfp11/Zfp13/Zfp26/Zfp28/Zfp30/Zfp37/Zfp39/Zfp40/Zfp46/Zfp51/Zfp52/Zfp54/Zfp57/Zfp60/Zfp61/Zfp85/Zfp9/Zfp90/Zfp93/Zfp94/Zfp97/Zik1/Zim1/Mavs/Zfp334/Ddx58/Zfp189/Ifna13/Tnfrsf14/Zfp12/Zfp212/Zfp954/Zfp418/Zfp772/Zfp114/Zfp790/Zfp940/Zfp420/Zfp382/Zfp939/AI987944/Zfp768/Zfp764/Zfp958/Zfp930/Zfp868/Zfp961/Zfp612/Zfp426/Zfp809/Zfp599/Zfp810/Zfp65/Zfp825/Zfp709/Zfp938/Zfp454/Zfp867/Akt3/Zfp458/Zfp874a/Zfp58/Zfp647/Zfp641/Zfp760/Zfp994/Zfp799/Zfp870/Zfp952/Zfp563/Zfp119b/Rnasel/Tlr2/Zfp53/Zfp68/Zfp267/Ifna15/Zfp933/A430033K04Rik/Zfp128/Zfp324/Zfp568/Zfp14/Zfp473/Zfp791/Zfp317/Zfp937/Map3k7/Irak4/Zfp747/Eif2ak4/Zfp398/Zfp354b/Zfp354c/Zfp459/Zfp853/Zfp786/Zfp78/Zfp82/Zfp866/Card9/Zfp780b/Gm5141/Rsl1/Zfp948/Zfp229/Zfp69/Zfp667/Zfp456/Zfp874b/Zfp950/Zfp457/Zfp708/Zfp268/Zfp141/Zfp975/Zfp560/Zfp960/Nxf1/Irf7/Irf3/Zfp316/Zfp607a/Zfp108/Tyk2/Alyref2/Zfp386/Zfp113/Tbk1/Ikbke/Zfp235/Zfp111/Mtor/Zfp109/Srsf4/Zfp112/Nectin1/Zfp872/Zfp551/Zfp963/Zfp951/Gm14322/Gm14430/Zfp808/Zfp964/Tsc1/Gm14391/Tab1/Gm14326/Zfp991/H2-T-ps/Gm8909/Gm14434/Zfp869/Zfp442/Zfp605/Zfp788/Zfp169/Hcfc2/Srsf6/Tab2/Zfp707/Zfp746/Zfp688/Bst2/Zfp715/2810021J22Rik/Zfp619/Zfp597/Zfp689/Zfp626/Zfp935/Ifih1/Zfp251/Tradd/Zfp949/2610008E11Rik/Zfp157/Zfp661/Zfp558/Zfp777/Sting1/Zfp248/Zfp74/Zfp974/Zfp763/Zfp946/Zfp84/Pik3cb/Zfp773/Zfp266/Zfp712/Zfp623/9130019O22Rik |
| **H3K27me3** | | | |
| mmu00830 | Retinol metabolism | 4,7053E-06 | Cyp3a41b/Cyp2c50/Cyp26a1/Cyp2a12/Cyp2b13/Cyp2b19/Cyp2b9/Cyp2c29/Cyp2c37/Cyp2c38/Cyp2c39/Cyp2c40/Cyp3a11/Cyp3a16/Cyp4a12b/Cyp4a14/Aldh1a2/Aox2/Cyp2b23/Cyp3a44/Cyp2c54/Cyp4a30b/Cyp3a41a/Cyp3a25/Cyp4a31/Cyp2c66/Aox3/Aox4/Cyp2c65/Rdh10 |
| mmu04080 | Neuroactive ligand-receptor interaction | 1,2235E-05 | Grm3/Grm5/Npy/Grik4/Chrna3/Chrna5/Gabra5/Adcyap1/Adra1b/Adra1a/Calcb/Calca/Calcr/Drd2/Drd3/Drd5/Ednrb/Fpr2/Fpr3/Fshb/Gabra1/Gabra2/Gabrb1/Gabrg1/Gabrg2/Glra1/Glrb/Gria1/Gria2/Grid1/Grid2/Grik2/Grin1/Grin2b/Grm1/Grm8/Gzma/Htr1a/Htr1b/Htr1f/Htr5a/Htr7/Iapp/Lep/Lepr/Mc3r/Nmbr/Npy2r/Npy5r/Npy6r/Oprm1/P2ry1/Pdyn/Prlr/Ptgdr/Ptgfr/Sst/Ghsr/Tac1/Tacr1/Vip/Grp/Prlhr/Rxfp3/Grin3a/Chrm2/Avpr1b/Rxfp1/Hcrtr2/Avpr1a/Nts/Glp2r |
| mmu00140 | Steroid hormone biosynthesis | 0,00019622 | Cyp3a41b/Cyp2c50/Cyp1b1/Cyp21a1/Cyp2b13/Cyp2b19/Cyp2b9/Cyp2c29/Cyp2c37/Cyp2c38/Cyp2c39/Cyp2c40/Cyp2d9/Cyp2e1/Cyp3a11/Cyp3a16/Cyp2b23/Cyp3a44/Cyp2c54/Cyp3a41a/Hsd17b12/Cyp3a25/Cyp2c66/Cyp2d40/Cyp2c65/Srd5a2 |
| mmu00591 | Linoleic acid metabolism | 0,00095136 | Cyp3a41b/Cyp2c50/Cyp2c29/Cyp2c37/Cyp2c38/Cyp2c39/Cyp2c40/Cyp2e1/Cyp3a11/Cyp3a16/Cyp2j12/Cyp3a44/Cyp2c54/Cyp3a41a/Cyp3a25/Cyp2c66/Cyp2c65 |
| mmu00590 | Arachidonic acid metabolism | 0,00128998 | Cyp2c50/Cbr3/Cyp2b13/Cyp2b19/Cyp2b9/Cyp2c29/Cyp2c37/Cyp2c38/Cyp2c39/Cyp2c40/Cyp2e1/Cyp4a12b/Cyp4a14/Gpx5/Ptgis/Cyp2j12/Cyp2b23/Cyp2c54/Cyp4a30b/Hpgds/Cyp4a31/Cyp2c66/Cyp2c65/Gpx6 |
| mmu04742 | Taste transduction | 0,00792905 | Scn2a/Gabra5/Adcy8/Gabra1/Gabra2/Grm1/Htr1a/Htr1b/Htr1f/P2ry1/Pde1a/Pde1c/Prkacb/Scn9a/Scnn1g/Tas2r139/Tas2r109/Tas2r118/Tas2r144/Tas2r140/Htr3b/Tas2r103 |
| mmu05204 | Chemical carcinogenesis | 0,00817418 | Cyp3a41b/Cyp2c50/Cyp1b1/Cyp2b13/Cyp2b19/Cyp2b9/Cyp2c29/Cyp2c37/Cyp2c38/Cyp2c39/Cyp2c40/Cyp2e1/Cyp3a11/Cyp3a16/Cyp2b23/Cyp3a44/Cyp2c54/Cyp3a41a/Hpgds/Cyp3a25/Mgst3/Cyp2c66/Cyp2c65 |
| mmu05032 | Morphine addiction | 0,02642491 | Gabra5/Adcy8/Gabra1/Gabra2/Gabrb1/Gabrg1/Gabrg2/Gnao1/Kcnj5/Kcnj6/Oprm1/Pde1a/Pde1c/Pde4b/Prkacb/Adcy2/Pde4d/Pde10a/Pde7b/Pde3a/Gng11 |
| mmu05033 | Nicotine addiction | 0,02855664 | Gabra5/Slc17a6/Gabra1/Gabra2/Gabrb1/Gabrg1/Gabrg2/Gria1/Gria2/Grin1/Grin2b/Grin3a |
| mmu04080 | Neuroactive ligand-receptor interaction | 1,2415E-12 | Grm5/Npy/Glra3/Chrna3/Chrna5/Gabra5/Chrna6/Chrna7/Adcyap1/Adra1b/Adra1a/Adra1d/Adrb3/Agtr1a/Calcb/Calca/Calcr/Cck/Crhr1/Crhr2/Ctsg/Drd2/Drd3/Drd4/Drd5/Ednra/Ednrb/Rxfp2/Fpr2/Fpr1/Fpr3/Fshb/Gabra1/Gabra2/Gabra6/Gabrb1/Gabrd/Gabrg1/Gabrg2/Gabrg3/Gcg/Gpr83/Glra1/Glrb/Gnrhr/Gria1/Gria2/Grid1/Grid2/Grik1/Grik2/Grik3/Grin1/Grin2a/Grin2b/Grm1/Grm8/Gzma/Hc/Hrh1/Htr1a/Htr1b/Htr1f/Htr2a/Htr4/Htr5a/Htr5b/Htr7/Iapp/Lep/Lepr/Lhcgr/Mc3r/Nmbr/Npy2r/Npy5r/Npy6r/Ntsr1/Oprm1/Pdyn/Prlr/Ptgdr/Sst/Sstr4/Ghsr/Tac1/Tacr1/Nmur2/Tshb/Tshr/Ucn/Vip/Grp/Prlhr/Npbwr1/Rxfp3/Gabbr2/Grin3a/Chrm2/Rxfp1/Hcrtr2/Avpr1a/Lpar3/Nts/Cysltr2/Ucn3/Glp2r |
| mmu05033 | Nicotine addiction | 0,00010051 | Gabra5/Chrna6/Chrna7/Cacna1b/Slc17a6/Gabra1/Gabra2/Gabra6/Gabrb1/Gabrd/Gabrg1/Gabrg2/Gabrg3/Gria1/Gria2/Grin1/Grin2a/Grin2b/Grin3a |
| mmu05032 | Morphine addiction | 0,00068557 | Gabra5/Adcy8/Cacna1b/Gabra1/Gabra2/Gabra6/Gabrb1/Gabrd/Gabrg1/Gabrg2/Gabrg3/Gnao1/Gng2/Kcnj5/Kcnj6/Oprm1/Pde1a/Pde1b/Pde1c/Pde4b/Prkacb/Prkcg/Adcy2/Adcy5/Pde4d/Pde10a/Pde11a/Gabbr2/Pde3a/Gng11 |
| mmu04726 | Serotonergic synapse | 0,00093887 | Cyp2c50/Alox12e/Alox15/Alox5/Cacna1b/Cyp2c29/Cyp2c37/Cyp2c38/Cyp2c39/Cyp2c40/Cyp2d9/Cyp2j5/Kcnn2/Gabrb1/Gnao1/Gng2/Htr1a/Htr1b/Htr1f/Htr2a/Htr4/Htr5a/Htr5b/Htr7/Kcnd2/Kcnj5/Kcnj6/Prkacb/Prkcg/Adcy5/Cyp2j13/Cyp2j12/Cyp2d12/Htr3b/Gng11/Cyp2c66/Cyp2d40/Cyp2c65 |
| mmu04514 | Cell adhesion molecules (CAMs) | 0,00093887 | H2-Q6/H2-Q9/Cd34/Cd86/Cdh2/Cldn1/Cntn1/Glycam1/H2-Ab1/H2-Bl/H2-M10.1/H2-M9/H2-Q1/H2-Q10/H2-Q2/H2-T3/Sdc2/Itga4/Ntng2/Ncam1/Nrxn1/Nrxn2/Nrxn3/Nlgn1/Ptprc/Sele/Selp/Cntn2/Cd40/Vcam1/H2-M10.4/H2-M11/Cd226/H2-M5/Itga8/Lrrc4c/Lrrc4b/Nrcam/Itgb8/Cadm1/Cldn14/Nectin3/Cntnap2/Jam2/Ntng1/Jam3 |
| mmu04742 | Taste transduction | 0,00990934 | Scn2a/Gabra5/Asic2/Adcy8/Gabra1/Gabra2/Gabra6/Grm1/Htr1a/Htr1b/Htr1f/Pde1a/Pde1b/Pde1c/Prkacb/Scn9a/Scnn1g/Gabbr2/Tas2r139/Tas2r109/Tas2r118/Tas2r138/Tas2r144/Tas2r140/Htr3b/Tas2r103 |
| mmu04940 | Type I diabetes mellitus | 0,00990934 | H2-Q6/H2-Q9/Cd86/Gad1/Gad2/Gzmb/H2-Ab1/H2-Bl/H2-M10.1/H2-M9/H2-Q1/H2-Q10/H2-Q2/H2-T3/Il12b/Il1b/Ins1/Ins2/Ptprn/H2-M10.4/H2-M11/H2-M5 |
| mmu04727 | GABAergic synapse | 0,00990934 | Gabra5/Adcy8/Cacna1b/Gabra1/Gabra2/Gabra6/Gabrb1/Gabrd/Gabrg1/Gabrg2/Gabrg3/Slc6a13/Gad1/Gad2/Gnao1/Gng2/Kcnj6/Prkacb/Prkcg/Adcy2/Adcy5/Slc6a1/Gabbr2/Slc6a11/Slc12a5/Gng11 |
| mmu05320 | Autoimmune thyroid disease | 0,01644517 | H2-Q6/H2-Q9/Cd86/Gzmb/H2-Ab1/H2-Bl/H2-M10.1/H2-M9/H2-Q1/H2-Q10/H2-Q2/H2-T3/Ifna1/Tg/Cd40/Tshb/Tshr/H2-M10.4/H2-M11/H2-M5/Ifna15/Ifna12/Ifna14 |
| mmu04512 | ECM-receptor interaction | 0,01644517 | Col2a1/Col4a3/Col9a1/Col1a2/Ibsp/Itga4/Itgb3/Lama1/Lama3/Lamc2/Reln/Thbs4/Tnc/Tnr/Itga8/Col6a6/Itgb8/Tnn/Frem1/Sv2b/Col6a5/Dspp/Col6a4/Sv2c/Tnxb |
| mmu00590 | Arachidonic acid metabolism | 0,01797491 | Cyp2c50/Cbr3/Alox12e/Alox15/Alox5/Cbr1/Cyp2b19/Cyp2c29/Cyp2c37/Cyp2c38/Cyp2c39/Cyp2c40/Cyp2e1/Cyp2j5/Cyp4a12b/Cyp4a14/Ptgis/Cyp2j13/Cyp2j12/Cyp4a12a/Cyp4a30b/Hpgds/Cyp4a31/Cyp2c66/Cyp2c65 |
| mmu04950 | Maturity onset diabetes of the young | 0,02029426 | Neurog3/Hhex/Mnx1/Iapp/Ins1/Ins2/Neurod1/Nkx6-1/Pax4/Rfx6/Mafa |
| mmu04750 | Inflammatory mediator regulation of TRP channels | 0,02351786 | Cyp2c50/Asic2/Adcy8/Alox12e/Cyp2c29/Cyp2c37/Cyp2c38/Cyp2c39/Cyp2c40/Cyp2j5/Cyp4a12b/Cyp4a14/Hrh1/Htr2a/Il1b/Prkacb/Prkcg/Prkch/Prkcq/Adcy2/Adcy5/Cyp2j13/Cyp2j12/Mapk10/Mapk8/Trpa1/Cyp4a12a/Cyp4a30b/Asic5/Cyp4a31/Cyp2c66/Cyp2c65 |
| mmu04020 | Calcium signaling pathway | 0,0246514 | Mylk/Grm5/Slc8a3/Chrna7/Adcy8/Adra1b/Adra1a/Adra1d/Adrb3/Agtr1a/Cacna1b/Cacna1e/Camk4/Drd5/Ednra/Ednrb/Erbb4/Gnal/Grin1/Grin2a/Grm1/Hrh1/Htr2a/Htr4/Htr5a/Htr5b/Htr7/Lhcgr/Nos1/Ntsr1/Pde1a/Pde1b/Pde1c/Pdgfra/Prkacb/Prkcg/Ppp3r1/Ryr2/Ryr3/Adcy2/Tacr1/Chrm2/Avpr1a/Cysltr2 |
| mmu00591 | Linoleic acid metabolism | 0,02706817 | Cyp3a41b/Cyp2c50/Alox15/Cyp2c29/Cyp2c37/Cyp2c38/Cyp2c39/Cyp2c40/Cyp2e1/Cyp2j5/Cyp3a16/Cyp2j13/Cyp2j12/Cyp3a25/Cyp2c66/Cyp2c65 |
| mmu04060 | Cytokine-cytokine receptor interaction | 0,04666969 | Il1rl2/Bmp10/Bmp4/Bmp5/Bmp6/Ccr6/Ccr1/Ccr3/Ccr2/Csf2ra/Ifna1/Ifnb1/Il12b/Il12rb2/Il18rap/Il1b/Il18r1/Il2ra/Il2rb/Il3/Il5ra/Il7r/Il9/Inhba/Lep/Lepr/Il1rl1/Ngfr/Tnfrsf11b/Prlr/Ccl1/Ccl11/Ccl12/Ccl2/Ccl3/Cxcl2/Cxcl5/Il23r/Il36g/Cd40/Tnfrsf8/Tnfsf11/Cd70/Tnfsf8/Il20ra/Gdf7/Tnfsf18/Gdf6/Ifna15/Ifna12/Tnfrsf19/Ifna14/Il22/Tslp/Il36a/Cxcl14/Crlf2/Il20/Il21/Il36b/Il33 |
