## Supplementary Table S1 for "EZH2 deletion does not impact acinar cell regeneration but restricts progression to pancreatic cancer in mice"

**Supplemental Table S1. Significant Gene Ontology pathways enriched in *Mist1^creERT/+^KRAS^G12D^* *vs* WT according to RNA-sequencing**

| **ID** | **Description** | **P adjust** | **Gene ID** |
| --- | --- | --- | --- |
| GO:0150146 | cell junction disassembly | 0.001275977 | C1qa/Tgfb3/Ngef/Dusp3/Iqsec1/Abcc8 |
| GO:0009914 | hormone transport | 0.004476534 | G6pc2/Slc16a10/Ttr/Gal/Hnf1b/Rfx6/Scg5/Snap25/Ptprn/Baiap3/Cltrn/Ucn3/Nr1d1/Selenom/Chga/Gipr/Neurod1/Anxa5/Abcc8/Ica1 |
| GO:0046879 | hormone secretion | 0.008077706 | G6pc2/Slc16a10/Gal/Hnf1b/Rfx6/Scg5/Snap25/Ptprn/Baiap3/Cltrn/Ucn3/Nr1d1/Selenom/Chga/Gipr/Neurod1/Anxa5/Abcc8/Ica1 |
| GO:0035592 | establishment of protein localization to extracellular region | 0.015677302 | G6pc2/Afm/Hnf1b/Rfx6/Ins2/Tgfb3/Ptprn/Baiap3/Cltrn/Ucn3/Nr1d1/Cd34/Ankrd1/Chga/Gipr/Rhbdf1/Neurod1/Anxa5/Ang/Abcc8/Ica1 |
| GO:0071692 | protein localization to extracellular region | 0.015738106 | G6pc2/Afm/Hnf1b/Rfx6/Ins2/Tgfb3/Ptprn/Baiap3/Cltrn/Ucn3/Nr1d1/Cd34/Ankrd1/Chga/Gipr/Rhbdf1/Neurod1/Anxa5/Ang/Abcc8/Ica1 |
| GO:0034764 | positive regulation of transmembrane transport | 0.017498712 | Slc1a2/Gal/Ins2/Dpp6/Iapp/Wnk4/Cltrn/Lrrc55/Rapgef3/F2/Gstm7/Rhoq/Hap1/Abcc8 |
| GO:0010721 | negative regulation of cell development | 0.017498712 | Spock1/Bmp7/Nexmif/Sema4g/Tnr/Ngef/Dynlt1b/Ccl11/Kremen1/F2/Fuom/Nr1d1/Inpp5j/Hes1/B2m/Trak2/Sema3d/Abcc8 |
| GO:0031018 | endocrine pancreas development | 0.017498712 | Hnf1b/Rfx6/Ins1/Iapp/Gipr/Hes1/Neurod1 |
| GO:1901381 | positive regulation of potassium ion transmembrane transport | 0.017498712 | Gal/Dpp6/Wnk4/Lrrc55/Rapgef3/Abcc8 |
| GO:0030072 | peptide hormone secretion | 0.017498712 | G6pc2/Slc16a10/Hnf1b/Rfx6/Ptprn/Baiap3/Cltrn/Ucn3/Nr1d1/Chga/Gipr/Neurod1/Anxa5/Abcc8/Ica1 |
| GO:0010959 | regulation of metal ion transport | 0.017498712 | Cntn1/Fxyd3/Gal/Dpp6/Vdr/Plcb4/Iapp/Atp2c2/Wnk4/Lrrc55/Adcyap1r1/Rapgef3/F2/Atp2b2/Gstm7/Hes1/B2m/Hap1/Abcc8 |
| GO:0009306 | protein secretion | 0.017700567 | G6pc2/Hnf1b/Rfx6/Ins2/Tgfb3/Ptprn/Baiap3/Cltrn/Ucn3/Nr1d1/Cd34/Ankrd1/Chga/Gipr/Rhbdf1/Neurod1/Anxa5/Ang/Abcc8/Ica1 |
| GO:0042445 | hormone metabolic process | 0.017700567 | Slc16a10/Ttr/Gal/Scg5/Pcsk6/Duox2/Dio1/Iyd/Pcsk1n/Selenom/Pcsk2/Egr1 |
| GO:0030073 | insulin secretion | 0.018034921 | G6pc2/Hnf1b/Rfx6/Ptprn/Baiap3/Cltrn/Ucn3/Nr1d1/Chga/Gipr/Neurod1/Abcc8/Ica1 |
| GO:0050708 | regulation of protein secretion | 0.018034921 | G6pc2/Rfx6/Ins2/Tgfb3/Baiap3/Cltrn/Ucn3/Nr1d1/Cd34/Ankrd1/Chga/Gipr/Rhbdf1/Anxa5/Ang/Abcc8/Ica1 |
| GO:1904064 | positive regulation of cation transmembrane transport | 0.018716907 | Gal/Dpp6/Iapp/Wnk4/Cltrn/Lrrc55/Rapgef3/F2/Gstm7/Hap1/Abcc8 |
| GO:0002791 | regulation of peptide secretion | 0.018863686 | G6pc2/Rfx6/Ins2/Tgfb3/Baiap3/Cltrn/Ucn3/Nr1d1/Cd34/Cd74/Ankrd1/Chga/Gipr/Rhbdf1/Anxa5/Ang/Abcc8/Ica1 |
| GO:0032102 | negative regulation of response to external stimulus | 0.02651814 | Mfhas1/Lgals9/Dusp1/Ins2/Sema4g/Tnr/Pyy/Kremen1/F2/Nr1d1/Cd34/Dusp3/Tspan6/Apod/Anxa5/Sema3d/Abcc8 |
| GO:0045745 | positive regulation of G protein-coupled receptor signaling pathway | 0.02651814 | Pde5a/Acpp/Slc39a14/F2/Chga |
| GO:0043268 | positive regulation of potassium ion transport | 0.026684306 | Gal/Dpp6/Wnk4/Lrrc55/Rapgef3/Abcc8 |
| GO:0016486 | peptide hormone processing | 0.029284342 | Scg5/Pcsk6/Pcsk1n/Pcsk2 |
| GO:0008277 | regulation of G protein-coupled receptor signaling pathway | 0.029284342 | Ramp1/Pde5a/Acpp/Dynlt1b/Slc39a14/F2/Grk3/Chga/Gipr |
| GO:0034767 | positive regulation of ion transmembrane transport | 0.029984192 | Gal/Dpp6/Iapp/Wnk4/Cltrn/Lrrc55/Rapgef3/F2/Gstm7/Hap1/Abcc8 |
| GO:0140448 | signaling receptor ligand precursor processing | 0.032894494 | Scg5/Pcsk6/Pcsk1n/Pcsk2 |
| GO:0001706 | endoderm formation | 0.041272697 | Dusp1/Hnf1b/Dusp5/Dusp4/Vtn |
| GO:0043270 | positive regulation of ion transport | 0.041366008 | Cntn1/Gal/Dpp6/Iapp/Atp2c2/Wnk4/Cltrn/Lrrc55/Adcyap1r1/Rapgef3/F2/Atp2b2/Gstm7/Hap1/Abcc8 |
| GO:0046883 | regulation of hormone secretion | 0.047170348 | G6pc2/Gal/Rfx6/Scg5/Snap25/Baiap3/Cltrn/Ucn3/Nr1d1/Chga/Gipr/Anxa5/Abcc8/Ica1 |
